## Supplemental Figures 1-3 for "The incidence of candidate binding sites for β-arrestin in Drosophila neuropeptide GPCRs"

**S1 Figure.** ***Drosophila*** **Rhodopsin-like GPCRs with BBS-like sequences in ICL2 following the conserved P (capitalized)**

No other Rhodopsin-like GPCRs had a BBS-like sequence in either *D. melanogaster* or *D. virilis*

**AstA R1**

**M(180)** drflavvh**P**v***tsmslrt***ernat

**V(200)** drflavvh**P**v***tsmslrt***ernat

**CCKL R 17D1**

**M(272)** eryyaich**P**lr***srtwqt***inh

**V(136)** eryyaich**P**lr***srtwqt***inh

**CCKL R 17D3**

**M(208)** eryyaich**P**lr***srswqt***ish

**V(212)** eryyaich**P**lr***srswqt***is

**PK2 R1**

**M(205)** eryiaich**P**frqh***tmskls***rai

**V(208)** eryiaich**P**frqh***tmskls***rai

**PK2 R2**

**M(164)** eryiaich**P**frqh***tmskls***rav

**V(171)** eryiaich**P**frqh***tmskls***rai

***S2. Figure.***

***Drosophila* Rhodopsin-like GPCRs do not contain a conserved P**

**6 AAs past the DRY sequence of ICL2**

Asterisk marks the +6 position

SP R has one (bold and capitalized) but it is 7 AAs past the DRY sequence

**LGR1**

```
                   *
M(593)      erwlaitqamylnhriklrpaa
V(575)      erwfaithamylnkritlrqaa
                   *
```

**Moody PC**

```
                   *
M(133)      nryvmithhglyariykrh
V(142)      nryvmithhgcyariykrh
                   *
```

**Rickets**

```
                   *
M(848)      ernyaithaihlnkrlslkqag
V(889)      ernyaithaihlnkrlslrqag
                   *
```

**SP R**

```
                   *
M(192)      qryiyvchaPmartwctmprvrr
V(256)      qryiyvchaPmartwctmprvr
                   *
```

*S3 Figure*

**The BBS-like sequence in ICL3 of the TrissinR display sequence refinement due to alternative slicing among its PB, PC and PD isoforms**

The BBS-like sequence is shown in **RED** and bold
Alternative splicing alters the final two AAs of the BBS-like sequence in PC (asterisks)
Alternative splicing also alters the peptide sequence immediately following the BBS (underlined)

```
Dm PB(469):  RKQSSKYEKRGVSITESQLDNCKVSLEADRPIVSACRKTSFYHHG
Dv PB(501):  RKHSSKYEKRGVSITESQLDNCKVSLEADRPIVSACRKTSFYHHS
                                 **
Dm PC(469):  RKQSSKYEKRGVSITESQVSLEADRPIVSACRKTSFYHHG
Dv PC(501):  RKHSSKYEKRGVSITESQVSLEADRPIVSACRKTSFYHHS
                                 **
Dm PD(469):  RKQSSKYEKRGVSITESQLDNCKVSLEADRPIVSACRKTSFYHHG
Dv PD(501):  RKHSSKYEKRGVSITESQLDNCKVSLEADRPIVSACRKTSFYHHS
```
