## Supplementary material for "The incidence of candidate binding sites for β-arrestin in Drosophila neuropeptide GPCRs": S1 Table

### S1 Table.
### Alternative D. melanogaster neuropeptide GPCR CT isoforms

* This entry lists the # of BBS in the *melanogaster* GPCR/the number of BBS which are conserved in *virilis*

** alt splicing produces a longer CT but lacking any additional BBS's

**** alt splicing produces an isoform with an unspliced intron that includes a BBS; not present in *virilis*

***** alt splicing produces different CTs but neither contains a BBS

| | Protein Isoform (alternative CTs) | Genbank Ref. # | Exp. Variance | # BBS * (total/conserved) | # BBS-like(* total/conserved) | Figure |
|---|---|---|---|---|---|---|
| **AKH R** | | | | | | 3 |
| | PA | AAF52426 | | 1/1 | 1/1 | |
| | PB | AAN10595 | | | | |
| | PC | AAS64647 | alt. splicing ** | 1/? | 1/? | |
| | PD | AGB92685 | | | | |
| **AstA-R1** | | | | | | 3 |
| | PB | AAF45884.3 | | 1/1 | 1/1 | |
| | PD | AAG22404 | | | | |
| **AstA-R2** | | | | | | 3, 23 |
| | PA | AAF56809 | | 0/0 | 0/0 | |
| | PB | AFH06669 | alt. splicing | 1/1 | 0/0 | |
| | PC | AGB96422 | | | | |
| **AstC-R1** | | | | | | 3 |
| | PA | AAF49259.2 | | 2/1 | 0/0 | |
| **AstC-R2** | | | | | | 3 |
| | PB | AAN11677.2 | | 1/1 | 0/0 | |
| | PD | AGB94708.1 | stop suppr. | 1/1 | 0/0 | |
| | PE | AAZ66058 | | | | |
| | PF | ALI30485 | stop suppr. | 1/1 | 0/0 | |

| | | | | | | |
|---|---|---|---|---|---|---|
| **CAPA R** | | | | | | 3, 14 |
| | PB | AAS65092.1 | | 2/0 | 0/0 | |
| | PC | AGB94868 | alt. splicing | 3/? | | |
| **CCAP-R** | | | | | | 3 |
| | PB | AAS65092.1 | | 1/1 | 0/0 | |
| | PC | AGB94868 | alt. splicing | | | |
| **CCHa 1 R** | | | | | | 5 |
| | PA | AAF57819 | | 1/0 | 0/0 | |
| **CCHa2 R** | | | | | | 5, 16 |
| | PA | AAF57285.4 | | 0/0 | 0/0 | |
| | PB | ACZ94340 | alt. splicing | | | |
| | PC | QCD26194 | stop suppr. | | | |
| **CCKLR-17D1 R** | | | | | | 3 |
| | PA | ABW09450 | | 1/1 | 1/1 | |
| **CCKLR-17D3 R** | | | | | | 3 |
| | PB | AAF48879 | | 2/1 | 3/3 | |
| **CG4313** | | | | | | 9 |
| | PC | AHN59270 | | | | |
| | PD | AAF45710 | | 0/0 | 0/0 | |
| | PE | AHN59271 | | | | |
| **CG12290** | | | | | | 9 |
| | PA | AAF56578 | | 2/1 | 1/1 | |
| | PB | AHN57539 | | | | |
| **CG13229** | | | | | | 9 |
| | PA | AAF58717 | | 1/1 | 0/0 | |
| | PB | AGB93413 | | | | |
| | PC | AGB93414 | | | | |
| **CG13575** | | | | | | 9 |
| | PA | AAF47188 | | 1/1 | 0/0 | |
| **CG13995** | | | | | | 9 |
| | PA | AAF52333 | | 0/0 | 2/2 | |
| **CG30340** | | | | | | 9, 21 |

| | | | | | | |
|---|---|---|---|---|---|---|
| | PA | AAM71077 | | 0/0 | 0/0 | |
| **CG32547** | | | | | | 8 |
| | PC | AAX52506 | | 4/4 | 1/1 | |
| | PD | AGB95524 | | | | |
| **CG33639** | | | | | | 9, 20 |
| | PA | AAF48813 | | 0/? | 0/? | |
| | PB | ADV37751 | | | | |
| | PC | ADV37752 | | | | |
| | PD | AGB95518 | lt. splicing **** | 0/0 | 0/0 | |
| **CNMa R** | | | | | | 5 |
| | PA | AAF50229.3 | | 1/1 | 1/0 | 5 |
| **Crz R** | | | | | | 6 |
| | PA | AAF49928 | | 3/3 | 1/1 | |
| | PB | AGB94448 | stop suppr. | 3/? | 1/? | |
| **ETH R** | | | | | | 6 |
| | PA | AAF55872 | | 1/1 | 0/0 | |
| | PB | AAS65191 | alt. splicing | 3/3 | 0/0 | |
| | PC | AHN57438 | | | | |
| **FMRFa R** | | | | | | 4 |
| | PA | AAF47700 | | 2/2 | 0/0 | |
| | PB | AGB94042 | | | | |
| | PC | AHN57950 | | | | |
| **Lg R1** | | | | | | 7, 17 |
| | PB | AAN13752 | | | | |
| | PA | AAF55460 | | 0/0 | 0/0 | |
| **Lg R3** | | | | | | 7 |
| | PA | AAF56490 | | 1/1 | 1/1 | 7 |
| **Lg R4** | | | | | | 7 |
| | PB | ABW09404 | | 1/1 | 2/2 | 7 |
| | PC | AHN59662 | | | | |
| **Lk R** | | | | | | 6 |
| | PA | AAF50775.2 | | 2/2 | 0/0 | 6 |

| | | | | | | |
|---|---|---|---|---|---|---|
| **moody** | | | | | | 8 |
| | PA | NP_569970.2 | | 4/2 | 0/0 | 8 |
| | PC | NP_001188535.1 | alt. splicing | 1/0 | 0/0 | |
| **MS R1** | | | | | | 7 |
| | PA | AAF47635.2 | | 1/0 | 1/0 | 7 |
| | PB | AGB94019 | stop suppr. | | | |
| **Ms R2** | | | | | | 7 |
| | PA | AAF47633 | | 1/1 | 0/0 | 7 |
| | PB | AAN12219 | | | | |
| | PC | AGB94018 | stop suppr. | | | |
| **NPF R** | | | | | | 7, 18 |
| | PA | AAF51909 | | 1/0 | 0/0 | |
| | PB | AFH06264 | alt. splicing | | | |
| | PC | AFH06265 | | | | |
| | PD | AFH06266 | | | | |
| **PK1 R** | | | | | | 4 |
| | PD | AAX52950 | | 2/1 | 3/1 | |
| | PE | AFH06407 | | | | |
| **PK2 R1** | | | | | | 4 |
| | PA | AAF54930 | | 1/1 | 1/1 | |
| **PK2 R2** | | | | | | 4 |
| | PA | AAF54929.2 | | 3/2 | 1/1 | |
| | PB | AAN13555.1 | | | | |
| **Proc R** | | | | | | 7, 12 |
| | PA | AAF45980.2 | | | | |
| | PB | AAN09130.1 | | | | |
| | PC | AAX52477.1 | | 1/0 | 1/0 | |
| | PD | AAX52478.1 | | | | |
| | PE | AHN59339 | | | | |
| **rk** | | | | | | 8 |
| | PA | AAF53367 | | 5/3 | 0/0 | |
| **Rya R** | | | | | | 5 |

| | | | | | | |
|---|---|---|---|---|---|---|
| | PA | AAF56655.3 | | 1/1 | 1/1 | |
| | PB | AHN57551 | alt. splicing | 2/2 | 2/2 | |
| | PC | AHN57552 | alt. splicing | 0/? | 0/? | |
| **SIFa R** | | | | | | 6 |
| | PA | AAN13859.2 | | 7/7 | 0/0 | |
| | PB | ACZ94970 | | | | |
| **sNPF R** | | | | | | 4 |
| | PA | AAF49074 | | 1/1 | 1/1 | |
| | PB | AGB94779 | | | | |
| **SP R** | | | | | | 5 |
| | PA | AAF46037 | | 1/1 | 1/1 | |
| | PB | AHN59364 | | | | |
| | PC | AHN59363 | | | | |
| | PD | QJC18359 | stop suppr. | | | |
| **Tk R 86C** | | | | | | 5, 13 |
| | PA | AAF54544.1 | | 2/0 | 1/1 | |
| | PB | ABW08638 | alt. splicing**** | 3/? | 1/? | |
| **Tk R 99D** | | | | | | 5 |
| | PA | AAF56979.2 | | 2/1 | 0/0 | |
| | PB | ACZ95066 | | | | |
| | PC | AGB96471 | stop suppr. | | | |
| **Tre1** | | | | | | 9 |
| | PA | NP_524792.1 | | 0/0 | 2/2 | |
| **Trissin R** | | | | | | 6 |
| | PB | AAF52294 | | 2/2 | 2/2 | |
| | PC | AAF52292 | | | | |
| | PD | AGB92644 | alt. splicing | 3/? | 3/? | |
| | PE | AHN54156 | | | | |
| **Dh31 R** | | | | | | 10 |
| | PA | AAN16138 | | 1/1 | 0/0 | 10 |
| | PB | AGB93482 | | | | |
| | PC | AGB93483 | stop suppr. | 3/3 | 0/0 | |

| | | | | | | |
|---|---|---|---|---|---|---|
| **Hector/CG4395** | | | | | | 10, 19 |
| | PA | AAF48216 | | 0/0 | 0/0 | |
| | PB | AHN59644 | | | | |
| **PDF R** | | | | | | 10, 15, 22 |
| | PA | AAF45788 | | 2/0 | 1/1 | |
| | PB | AFH07215 | | | | |
| | PC | AHN59297 | same as PD | | | |
| | PD | AHN59298 | alt. splicing | 4/2 | 1/1 | |
| **Dh44 R1** | | | | | | 11 |
| | PA | AAF58250 | | 2/2 | 1/1 | |
| **Dh44 R2** | | | | | | 11 |
| | PA | AAF58501 | | 1/1 | 1/1 | |
| | PB | AAM68690 | alt. splicing | 1/1 | 0/0 | |
