## Supplementary material for "The incidence of candidate binding sites for β-arrestin in Drosophila neuropeptide GPCRs": S 1 Text

***S1. Text***

**Collated and Annotated D. melanogaster and D. virilis GPCR sequences**

**M** = ***D. melanogaster*** **V** = ***D. virilis***

**BBS candidates in bold**

Predicted TM domains

*D. melanogaster*

*D. virilis*

**M AKH R – PA (Genbank ref # AAF52426)**

1 makvaeendh rdlsnwsnvn dtngtihltk dmvfndghrl sitvysilfv istignstvl
61 ylltkrrlrg plridimlmh laiadlmvtl llmpmeivwa wtvqwlstdl mcrlmsffrv
121 fglylssyvm vcisldryfa ilkplkrsyn rgrimlacaw lgsvvcsipq aflfhleehp
181 avtgyfqcvi fn***sfrsdfd***e klyqaasmcs myafplimfi ycygaiylei yrksqrvlkd
241 viaerfrrsn ddvlsrakkr tlkmtitivi vfiicwtpyy tismwywldk hsagkinpll
301 rkalfifast nscmnplvyg lynirgrmnn nnpsvnnrht slsnrldssn qlmqkqltnn
361 sllngrgqvm aaavsattkl anvvslkgta ngngsaaaag tvpitppltv tiaplatdde
421 andd***sclsav t***ircqdqspi rqk

(***sfrsdfe*** sequence in ECL2)

**M AKH R-PC (Genbank ref #** AAS64647)

1 makvaeendh rdlsnwsnvn dtngtihltk dmvfndghrl sitvysilfv istignstvl
61 ylltkrrlrg plridimlmh laiadlmvtl llmpmeivwa wtvqwlstdl mcrlmsffrv
121 fglylssyvm vcisldryfa ilkplkrsyn rgrimlacaw lgsvvcsipq aflfhleehp
181 avtgyfqcvi fn***sfrsdfd***e klyqaasmcs myafplimfi ycygaiylei yrksqrvlkd
241 viaerfrrsn ddvlsrakkr tlkmtitivi vfiicwtpyy tismwywldk hsagkinpll
301 rkalfifast nscmnplvyg lynirgrmnn nnpsvnnrht slsnrldssn qlmqkqltnn
361 sllngrgqvm aaavsattkl anvvslkgta ngngsaaaag tvpitppltv tiaplatdde
421 andd***sclsav t***ircqdqspi rqkcgdsiel tsvvk

**V AKH R PC (Genbank ref #**XP_002051398.2)

1 maqsgevnev vydhrilrdw snvnntngtm hlskdmifnd ghrlsitvys ilfvistign
61 stvlylltkr rlrgplridi mlmhlaiadl mvtlllmple iawawtvqwr stdlmcrlms
121 ffrvfglyls sfvmvcisld ryyailkplq rsynrgriml acawlgsvic sipqaflfhl
181 eehpivkgyf qcv***tfhsfvs efd***nwlyqia tmcamyafpl iafiycygai yleiyrknqr
241 vhkdviaerf rrsnddvlsr akkrtlkmti sivivfiicw tpyyficmwy sldktsvdkv
301 nslvrkalfi fastnscmnp lvyglynirg rmnnnnnvsv nnrhtslsnr ldssnqllqk
361 pintlpnnng nvmaaavaat tklahvvrlk tngaagdsqs pvaaaplaap ldindv***svs***
421 ***vvt***tkcqeqt pkqpktpiic lncgdsieva svekt

**M AstA R1 PB (Genbank Ref #** AAF45884.3)

1 maghqslall latlisswpk aswgatgngs iisvsnssgn nyaftsehtd hsdhnandsm
61 eydaesvale rivsti
77 vpvffgiigf agllgnglvi lvvvanqqmr sttnlliinl avsdilfvif cvpftatdyv
147 lpewpfgnvw ckfvqymivv tchcsvytlv lmsfdrflav vhpv***tsmslr t***ernatlaim
207 cawitivtta ipvalshsvr iyqyhgnagt acvfsteeei wslvgfqvsf flssyvaplt
267 licflymgml arlwksapgc kpsaesrkgk rrvtrmvvvv vlafaicwlp ihvilvlkal
327 nlyggshlsv iiqiishvva ytnscinpil yaflsdnfrk afrkvvwcgs ppplmtnqqv
387 ***tkttrt***atgn gtsnieml

**V AstA R1 PB (Genbank Ref #** XP_002055362.1)

1 mglsshrlam alaltvlisf wptgshgaaa vdcadwlsps ecsnstgnnn nnnnnnnkig
61 nnssnyaags taldhlnssm rpdnllemdl eqdgdnwple rivsii
107 vpvffgiigf agllgnalvi lvvvvnqqmr sttnlliinl avsdilfvif cvpftatdyv
167 lpewpfgnlw ckfvqymivv tchcsvytlv lmsfdrflav vhpv***tsmslr t***ernatlaim
227 cawitivtta ipvalahsvr iyqyhgragt acvfsteeev wslvgfqvsf flssyvaplt
287 licflymgml arlwksapgc kpsaesrkgk rrvtrmvvvv vlafaicwlp ihvilvlkal
347 nmyggthltv iiqiishvla ytnscinpil yaflsdnfrk afrkvvwcgs pppivtnqqm
380 ***tkttrt***atgn gtsnieml

**M AstA R2 PA** **(Genbank Ref #** AAF56809)

1 menttmlani slnatrneen itsfftdeew laingtlpwi vgfffgviai tgffgnllvi

61 lvvvfnnnmr sttnlmivnl aaadlmfvil cipftatdym vyywpygrfw crsvqylivv

121 tafasiytlv lmsidrflav vhpirsrmmr tenitliaiv tlwivvlvvs vpvafthdvv

181 vdydakknit ygmctfttnd flgprtyqvt ffissyllpl miisglymrm imrlwrqgtg

241 vrmskesqrg rkrvtrlvvv vviafaslwl pvqlillks ldvietntlt klviqvtaqt

301 layssscinp llyaflsenf rkafyk/avnc ssryqnytsd lppprktsca rtsttgl

**V AstA R2 (PA-like) (Genbank Ref #** XP_002053983.1)

1 mnlsnitltl psnsswlesq lelttassnl dnstlssfya eaeairatvr wvvpfffgii

61 aisgffgnll vilvvllnkn mhsttnlliv nlaaadllfv ifcvpftaid yvtqhwpfgk

121 mwcrsvqyli vvtayasiyt lvlmsidrfl avvhpirsrm lrtehitkia iftlwtvvlt

181 vsmpvtfahd vvvdydnqtn vtyamcryid ndvldlstfq vsffissyll plmvisglyv

241 rmimrlwhqg tgvrmskesq rgrkrvtrlv vvvviafasl wlpvqlilll kaldmyeins

301 mfnvilqiva htmaytssci npllyaflsd nfrkafyk/ai ncsnryhnyt sdlppprkts

361 cgrtsttgl

**M Ast-A R2-PB** **(Genbank Ref #** AAF56809)

1 menttmlani slnatrneen itsfftdeew laingtlpwi vgfffgviai tgffgnllvi

61 lvvvfnnnmr sttnlmivnl aaadlmfvil cipftatdym vyywpygrfw crsvqylivv

121 tafasiytlv lmsidrflav vhpirsrmmr tenitliaiv tlwivvlvvs vpvafthdvv

181 vdydakknit ygmctfttnd flgprtyqvt ffissyllpl miisglymrm imrlwrqgtg

241 vrmskesqrg rkrvtrlvvv vviafaslwl pvqlillks ldvietntlt klviqvtaqt

301 layssscinp llyaflsenf rkafyk/glqs nrlgmw***ttth qd***vssektty

**V AstA R2 (PB-like) (not annotated; obtained by direct inspection of genomic DNA – see SXX document [AstA R2] isoform annotations)**

1 mnlsnitltl psnsswlesq lelttassnl dnstlssfya eaeairatvr wvvpfffgii

61 aisgffgnll vilvvllnkn mhsttnlliv nlaaadllfv ifcvpftaid yvtqhwpfgk

121 mwcrsvqyli vvtayasiyt lvlmsidrfl avvhpirsrm lrtehitkia iftlwtvvlt

181 vsmpvtfahd vvvdydnqtn vtyamcryid ndvldlstfq vsffissyll plmvisglyv

241 rmimrlwhqg tgvrmskesq rgrkrvtrlv vvvviafasl wlpvqlilll kaldmyeins

301 mfnvilqiva htmaytssci npllyaflsd nfrkafyk/ek ikikpn***swhs vgt***rgrrsap

361 lcftyf

**M AstC R1 – PA ((Genbank Ref #** AAF56809)

```
  1 mftwlmmdvl qfvkgemtad seanatnwyn tneslyttel nhrwisgsst iqpeeslygt
 61 dlptyqhcia trnsfadlft vvlygfvcii glfgntlviy vvlrfskmqt vtniyilnla
121 vadecfligi pfllytmric swrfgefmck aymvstsits ftssifllim sadryiavch
181 pisspryrtl hiakvvsaia wstsavlmlp vilyastveq edginyscni mwpdaykkhs
241 gttfilytff lgfatplcfi lsfyylvirk lrsvgpkpgt kskekrrahr kvtrlvltvi
301 svyilcwlph wisqvalihs npaqrdlsrl eiliflllga lvysnsavnp ilyaflsenf
361 rksffkaftc mnkqdinaql qlepsvftkq gskkrggskr lltsnpqipp llplnagnnn
421 sstttssttt aektgttgtq kscnsngkvt appenliicl seqqeafctt arrgsgavqq
481 tdl
```

**V AstC R1 – PA (Genbank Ref #** XP_002048373.1)

```
  1 mwllivglhl vggsptesqp ewllmgntst apynypndti ysthsseylp ttgssihstd
 61 lptyqhciat rnsfadlftv vlyglvcivg lfgntlviyv vlrfskmqtv tniyilnlai
121 adecfligip fllytmrics wrfgelmcka ymvstsitsf tssifllims adrymavchp
181 isspryrtlh nakvvsalaw stsavlmlpv mlyastveqe dginyscnim wpdaykkhsg
241 ttfilytffl gfatplcfil sfyylvirkl rsvgpkhttk skekrrahrk vtrlvltvit
301 vyiscwlphw msqlalinsn paqrdlsrle iliflllgal vysnsavnpi lyaflsenfr
361 ksffkaftcm tkqdinaqlq lepsvftkqg srrrggsrrl lttnpqqqqe qqppllalhv
421 gnnnsststs stttaektgs tnapkscssn gkltvpgsst apeaenliic lseqheafct
481 ttrrgsslvq qtdl
```

**M AstC R2 – PB (Genbank Ref #** AAN11677.2)

1 meggwwrggg gggrlggkai meghstpnga aashrnnstr tniatngcah sgillfvlta

61 mtltslitpt eqlavapngt tlhqlesves esypsingtq netmvtsvrp hldhrnrptq

121 qngshyleyd ddgpdcsysy nfilklitmi lyalvciigl fgntlviyvv mrfskmqtvt

181 niyilnlaia decfligipf llytmqvgnw pfgnymckay mvstsitsft ssiflimsa

241 dryiavchpi sspryrtpfv sklvsafawm tsvllmlpvi lfastvqssn gnvscniewp

301 dtqnshtdst filyslvlgf atpltfilvf yclvirklht vgpkhkskek krshrkvtkl

361 vltvisayif cwlphwisqv alissapqrc asrlelavfl acgclsysns amnpilyafl

421 sdnfkksfmk actcaarkdv naqlqlensf fpkfgkgrqs erllggngkg gaqrgaltkk

481 kclatrnnna pma***ttttttt tttgt***davtc lqppvhqvpa eiqvgnpatv lvvnaetnnc

541 kppvlhtdl

**V AstC R2 PB (Genbank Ref #** XP_002048370.2)

1 mkgypkpnga anrschmdhv hggcgvllfl ltaltltsli tpteqltats tsvatlangt

61 atvqpmptyt kssdadaaad sdidvdvnse gfspalelsd yykhmnanlv nggsgsgfif

121 psgfngtlpi gfggdppngs gpynpmqgcr rdgsfgleli tmilyalvci vglfgntlvi

181 yvvlrfskmq tvtniyilnl aiadecflig ipfllytmqv gnwpfgnymc kaylvstsvt

241 qftssiflli msadryiavc hpisspryrt pfvskvvsav awttsvllml pvilfastfe

301 sgpghvscsi nwpeafniqs dsafilyslv lgfvtplifi mifyclvirk lhtvgpkhks

361 kekkrshrkv tklvltvitv yimcwlphwi sqvalinstp gcasrlelav flacgclsys

421 nsamnpilya flsdnfkksf mkactcaark dvnaqlqlen sffpkfgkgr qserligpna

481 aannskakkr alanarnnna qm***tttttttt agt***dvvtsiq pvaithtape vtptagaall

541 vvnaetnckp pvlhtdl

**M AstC R2 – PD (Genbank Ref #** AGB94708.1)

1 meggwwrggg gggrlggkai meghstpnga aashrnnstr tniatngcah sgillfvlta

61 mtltslitpt eqlavapngt tlhqlesves esypsingtq netmvtsvrp hldhrnrptq

121 qngshyleyd ddgpdcsysy nfilklitmi lyalvciigl fgntlviyvv mrfskmqtvt

181 niyilnlaia decfligipf llytmqvgnw pfgnymckay mvstsitsft ssiflimsa

241 dryiavchpi sspryrtpfv sklvsafawm tsvllmlpvi lfastvqssn gnvscniewp

301 dtqnshtdst filyslvlgf atpltfilvf yclvirklht vgpkhkskek krshrkvtkl

361 vltvisayif cwlphwisqv alissapqrc asrlelavfl acgclsysns amnpilyafl

421 sdnfkksfmk actcaarkdv naqlqlensf fpkfgkgrqs erllggngkg gaqrgaltkk

481 kclatrnnna pma***ttttttt tttgtdavt***c lqppvhqvpa eiqvgnpatv lvvnaetnnc

541 kppvlhtdl**x** drapsmplet vvfiarr

**X** = stop suppression

**M AstC R2 – PF (Genbank Ref #** AGB94708.1)

1 meggwwrggg gggrlggkai meghstpnga aashrnnstr tniatngcah sgillfvlta

61 mtltslitpt eqlavapngt tlhqlesves esypsingtq netmvtsvrp hldhrnrptq

121 qngshyleyd ddgpdcsysy nfilklitmi lyalvciigl fgntlviyvv mrfskmqtvt

181 niyilnlaia decfligipf llytmqvgnw pfgnymckay mvstsitsft ssiflllimsa

241 dryiavchpi sspryrtpfv sklvsafawm tsvllmlpvi lfastvqssn gnvscniewp

301 dtqnshtdst filyslvlgf atpltfilvf yclvirklht vgpkhkskek krshrkvtkl

361 vltvisayif cwlphwisqv alissapqrc asrlelavfl acgclsysns amnpilyafl

421 sdnfkksfmk actcaarkdv naqlqlensf fpkfgkgrqs erllggngkg gaqrgaltkk

481 kclatrnnna pma***ttttttt*** ***tttgtdavt***c lqppvhqvpa eiqvgnpatv lvvnaetnnc

541 kppvlhtdl**x** drapsmplet vvfiarr**x**dh qveldldtai dcqaiarqpe cll

**X** = stop suppression

**M CAPA R PB (Genbank Ref #** AGB94708.1)

```
  1 mnsstdptfs elnasftntp dtlfatsvss dpshgfgeed yacgtfncsp kefvafvlgp
 61 qtlplykavl itiifggifi tgvvgnllvc iviirhsamh tatnyylfsl avsdllyllf
121 glptevflyw hqypdlfgmp fckirafise actyvsvfti vafsmerfla ichplhlyam
181 vgfkrairii talwivsfis aipfgllsdi qylnypldhs rieesafcsm spkivneipv
241 fevsfciffv ipmiliilly grmgakirsr tnqklgvqqg tnnretrnsq mrkktvirml
301 aavvitffvc wfpfhlqrli flyaknmdny ldinealfsi agfayyvsc//t vnpivysvms
361 rryrvafrel lcgkavgayy nsgfardhss fressaydrv hs//vhvrasqh pnkfetdsss
421 anrvlikkty slplpk/nads tvlsttdivi vlenshtvce epkvendiwi eneetci
(/ = intron)
```

**V CAPA R X2 PB (Genbank Ref #** XP_002047623.2)

```
  1 mnmnmstnms mdtnlstylg tsdatalpyp gmddygcphm nctamefvqf vlgpqtlplh
 61 kallisiifs gifitgvlgn vlvcmviirh aamhtatnyy lfslavsdll ylllglpaev
121 flywhqypyl fglpfcklra fvseactyvs vftivafsme rflaichplh vcamsgfqra
181 lriitalwiv sflsaipfgv kteiqylnfp n/dgsrilesa fcsielefpe efplfevsfc
241 iffiipmili illygrmgaq irsratdklg/vqqgsrnres rssqkkkravirml/aavvit
301 ffvcwfpfhl qrlwflyakn ianyqdvnew lfsiagfayy vs//ctinpivy nvmsqryrva
361 fkeilcgkka gayynsgfar dqssfirdes sfrrgssatp nlrgstrysr vsang/madcs
421 llntttkivi vlgnnspqrd vdrnipeete akkqen
```

**M CCAP R PC (Genbank Ref #** AGB94708.1)

1 mlhlrlfdss lyytlasase ssglasstst ersfngtqga ggvaaggesl tptdvaavnl
61 tyftpaishv mlapttiatt tasatmvqiq ttaapshdle tggnstssdp gefdnlnsfy
121 fyeteqfavl wilftvivlg nsavlfvmfi nknrksrmny fikqlala/dl cvgllnvltd
181 iiwritiswr agnlackair fsqvcvtyss tyvlvamsid rydaithpmn fsks/wkrarh
241 lvagawlisa lfslpilvly eekliqghpq cwielgspia wqvymslvsa tlfaipalii
301 sacyaiivkt iwakgsifvp t/eragfgaap arrassrgii prakvktvkm tltivfvfii
361 cwspyiifdl lqvfgqiphs qtniaiatfi qslaplnsaa npliyclfss qvfrtlsrfp
421 pfkwftccck syrnnsqqnr chtvgrrlhn ***scdsmrtltt sltvsrrs***tn ktnarvvice
481 rptkvvtvpa msev

**V CCAP R PC (Genbank Ref #** XP_002053465.1)

1 mlhlrlfdss lyytlasvss gmlpppqasn gsqltltgaa agiapaestv nltyftpais
61 hvmlaptptt asaaqtteaa ttttatptta stttpgsdfe aaaataasgd nltspyagel
121 dnlnsfyfye teqfavlwil ftiivlgnsa vlfvmfinkn rksrmnyfir qlaladlcvg
181 llnvltdiiw ritiswragn vackvirfsq vcvtysstyv lvamsidryd aithpmnfsk
241 swkrarhlva gawllsalfs lpilvlyeek liqghpqcwi elgspmawqi ymclvsaalf
301 avpaliisac yaiivktiwa kgsifvpter vgfggaatrr assrgiipra kvktvkmtlt
361 ivfvfilcws pyiifdllqv fgqiphsqtn iaiatfiqsl aplnsaanpl iyclfssqvf
421 rtlsrfppfk wltcccksyr nnsqqnrcht vgrrlhn***scd smrtlttslt vsrrs***tnktn
481 arvvicerpn kvitvpamse v

**M CCAP R PD (Genbank Ref #** AAS65092.1)

1 mlhlrlfdss lyytlasase ssglasstst ersfngtqga ggvaaggesl tptdvaavnl
61 tyftpaishv mlapttiatt tasatmvqiq ttaapshdle tggnstssdp gefdnlnsfy
121 fyeteqfavl wilftvivlg nsavlfvmfi nknrksrmny fikqlala/dl cvgllnvltd
181 iiwritiswr agnlackair fsqvcvtyss tyvlvamsid rydaithpmn fsks/wkrarh
241 lvagawlisa lfslpilvly eekliqghpq cwielgspia wqvymslvsa tlfaipalii
301 sacyaiivkt iwakgsifvp t/eragfgaap arrassrgii prakvktvkm tltivfvfii
361 cwspyiifdl lqvfgqiphs qtniaiatfi qslaplnsaa npliyclfss qvfrtlsrfp
421 pfkwftccck syrnnsqqnr chtvgrrlhn ***scdsmrtltt sltvsrrs***tn ktnarvvice
481 rptkvvtvpa msev**laht srkrsafsih gpt***sfsdae***f l

** Stop Suppression

**V CCAP R (PD-like) (not annotated – obtained by direct inspection of genomic DNA; see SXX Document de novo [CCAP R] isoform annotations**)

1 mlhlrlfdss lyytlasvss gmlpppqasn gsqltltgaa agiapaestv nltyftpais
61 hvmlaptptt asaaqtteaa ttttatptta stttpgsdfe aaaataasgd nltspyagel
121 dnlnsfyfye teqfavlwil ftiivlgnsa vlfvmfinkn rksrmnyfir qlaladlcvg
181 llnvltdiiw ritiswragn vackvirfsq vcvtysstyv lvamsidryd aithpmnfsk

241 swkrarhlva gawllsalfs lpilvlyeek liqghpqcwi elgspmawqi ymclvsaalf

301 avpaliisac yaiivktiwa kgsifvpter vgfggaatrr assrgiipra kvktvkmtlt

361 ivfvfilcws pyiifdllqv fgqiphsqtn iaiatfiqsl aplnsaanpl iyclfssqvf

421 rtlsrfppfk wltccccksyr nnsqqnrcht vgrrlhn***scd smrtlttslt vsrrs***tnktn

481 arvvicerpn kvitvpamse v**lahstrk rstfavhgpt ***sfsdae***fl

**M CCHa1 R-PA (Genbank Ref #** AAF57819)

```
  1 mianlvsmet dlamnigldt sgeaptalpp mpnvtetlwd lamvvsqstq wplldtgsse
 61 nfselvttet pyvpygrrpe tyivpilfal ifvvgvlgng tlivvflsvr qmrnvpntyi
121 lslaladllv iittvplast vytveywpyg sflcslsefm kdvsigvsvf tltalsgdry
181 faivdplrkf hahgggrrat rmtlatavsi wllailcglp aligsnlkhl gineksivic
241 ypypeewgin yaksmvllhf lvyyaiplvv iavfyvlial hlmysasvpg eiqgavrqvr
301 arrkvavtvl afvvifgicf lpyhvfflwf yfwptaqddy nafwhvlriv aycmsfansc
361 anpvalyfvs gafrkhfnry lfcrgasgrr kkrgqhdtfc mhrdtsltst askrfqsrhs
421 cyqstirscr lqettittlp nggnqngani savelalpvl qapghneaha ppsygflpln
481 eivqqtrssp akfqeslln
```

**V CCHa1 R (Genbank Ref #** XP_002049859.1)

```
  1 mmtsvlsdis tiamdmelgs gsesplspln atralwelam tttaetplfd engslvpvte
 61 ipyvpygrrl etyivpilfa iifvvgvlgn gtlivvflsv rqmrnvpnty ilslaladll
121 vilttvplvs tvyaveywpw gsflcsvsef mkdvsigvsv ftltalssdr yfaivdplrk
181 fhahgggrra trmtlaiavs iwllaiicgl paligsnlkp vginqeksiv icypypeawg
241 dnyaklmvml hflvyyaipl viiaafyvmi alhlmysasv pgemqgavrq vrarrkvavt
301 vlafvvifgi cflpyhvffl wfyywptaqq dynmfwhvlr ivgfcmsfan scanpvalyf
361 vsgafrkhfn rylfcrgisg rrkkrnqhnd tfcmhrntsl tstaskrfqs rhscyqstvr
421 scrlqettit tlpngglnga pstavthnea pgyeftplsd fgplkasqla qrlqespln
```

**M CCHa2 R-PA (Genbank Ref #** AAF57285.4)

```
  1 myaslmdvgq tlaarladsd gngandsgll atgqgleqeq eglaldmghn asadggivpy
 61 vpvldrpety ivtvlytlif ivgvlgngtl viiffrhrsm rnipntyils laladllvil
121 vcvpvativy tqeswpfern mcriseffkd isigvsvftl talsgeryca ivnplrklqt
181 kpltvftavm iwilaillgm psvlfsdiks ypvftatgnm tievcspfrd peyakfmvag
241 kalvyyllpl siigalyimm akrlhmsarn mpgeqqsmqs rtqararlhv armvvafvvv
301 fficffpyhv felwyhfypt aeedfdefwn vlrivgfcts flnscvnpva lycvsgvfrq
361 hfnrylccic vkrqphlrqh statgmmdnt svmsmrrsty vggtagnlra slhrnsnhgv
421 ggagggvggg vgsgrvgsfh rqdsmplqhg nahgggaggg ssglgaggrt aavsek/sfin
481 ryesgvmry
```

**V CCHa-2 R PA (Genbank Ref #**XP_002049178.2)

```
  1 mpknmlaalm dmsqtlaasl ayaplesnaa ataaaaaaaa vvlnvsqlgn ssqldgslat
 61 aaatttttav ttststhnas geeypqyvkv ldrpetyivt vlytlifivg vlgngtlvii
121 ffrhrsmrni pntyilslal adllvilvcv pvativytqe swpfernmcr iteffkdisi
181 gvsvftltal sgerycaivn plrklqtkpl tvftaviiwv faimlgmpsf vvsdiqgytl
241 ptpngnitie vcspfrskiy akymvvakas iyylvplsii gvlyimmakr lhisardmpg
301 eqlsiqsrsq ararrhvarm vvafvvvffi cffpyhvfel wyhfyptaee dfddfwhvvr
361 ivgfctsfln scvnpvalyc vsgvfrqhfn rylccicvkr qphlrqhsta tgvmdtsvts
421 mrrstyvggg gggavggsla ahraslhmnn nhgvavgggg ggggrggsfh rqdsmplqha
481 gsgnghahnv ggpgagigra siinek/slik rydertry
```

**M CCHa2 R PB (Genbank Ref #** AAF57285.4)

```
  1 myaslmdvgq tlaarladsd gngandsgll atgqgleqeq eglaldmghn asadggivpy
 61 vpvldrpety ivtvlytlif ivgvlgngtl viiffrhrsm rnipntyils laladllvil
121 vcvpvativy tqeswpfern mcriseffkd isigvsvftl talsgeryca ivnplrklqt
181 kpltvftavm iwilaillgm psvlfsdiks ypvftatgnm tievcspfrd peyakfmvag
241 kalvyyllpl siigalyimm akrlhmsarn mpgeqqsmqs rtqararlhv armvvafvvv
301 fficffpyhv felwyhfypt aeedfdefwn vlrivgfcts flnscvnpva lycvsgvfrq
361 hfnrylccic vkrqphlrqh statgmmdnt svmsmrrsty vggtagnlra slhrnsnhgv
421 ggagggvggg vgsgrvgsfh rqdsmplqhg nahgggaggg ssglgaggrt aavsekr
```

**M CCHa2 R PC** stop suppression **(Genbank Ref #** QCD26194)

```
  1  myaslmdvgq tlaarladsd gngandsgll atgqgleqeq eglaldmghn asadggivpy
 61  vpvldrpety ivtvlytlif ivgvlgngtl viiffrhrsm rnipntyils laladllvil
121  vcvpvativy tqeswpfern mcriseffkd isigvsvftl talsgeryca ivnplrklqt
181  kpltvftavm iwilaillgm psvlfsdiks ypvftatgnm tievcspfrd peyakfmvag
241  kalvyyllpl siigalyimm akrlhmsarn mpgeqqsmqs rtqararlhv armvvafvvv
301  fficffpyhv felwyhfypt aeedfdefwn vlrivgfcts flnscvnpva lycvsgvfrq
361  hfnrylccic vkrqphlrqh statgmmdnt svmsmrrsty vggtagnlra slhrnsnhgv
```

421  ggagggvggg vgsgrvgsfh rqdsmplqhg nahgggaggg ssglgaggrt aavsekr**x**gt
481

**M CCK-R 17D1 PA (Genbank Ref #** ABW09450)

```
  1 mlprlcadac rqcfakiarr dthrgtrtpy gcadtqsrpk pnfllrevde vcctaasasp
 61 rllvlfrdhk rasffgltid afyhylrqal plakeaaihl nasneisavg dgvtitgtpg
121 dllnysglel dlgldldlnl dmdlattpss stlapavtvr tpgnrsvvrv sadvpiwvvp
181 cysaillcav vgnllvvltl vqnrrmrtit nvfllnlais dillgvfcmp vtlvgtllrh
241 fifgellckl iqfaqaasva vsswtlvais ceryyaichp lrsrtwqtin hankiiaiiw
301 lgslvcmtpi aafsqlmpts rpglrkcreq wpadslnyer aynlfldlal lvlpllalsf
361 tylfitrtly vsmrneramn fgssgpevtt sssaavaeag sqrrangshc qsldtivphq
421 hnphqqhhhh sqyyydyghc gskrrlisgg gpcegrrhly cmrsasvksl rhqqinggggg
481 tlsgtgagng eccsrvhrmr qqmqlqqqgy vsdnesrrks lsqpslrite aglrrsnetk
541 sleskkrvvk mlfvlvleff icwtplyvin tmtmllgptv yeyvgytsis flqllaysss
601 ccnpitycfm nasfrrafvd tfkgmrvcer lcapccfwrr rsknetnlsv agnsialans
661 vmsshtiles prl
```

**V CCK-R 17D1-like (Genbank Ref #** XP_032295977.1)

```
  1 msgslaseat mtsatasatv mptplvpsnr smsrviadvp iwvipcysii llcavvgnll
 61 vvltlvqnrr mrtitnvfll nlaisdillg vlcmpvtlvg tllrnfifge slckliqfaq
121 aasvavsswt lvaisceryy aichplrsrt wqtinhanki iafiwlgslv cmtpiaifsq
181 lmptsrqglr kcreqwpans lgyeraynif lnlallvlpl malsfaylfi trtlyvsmrn
241 eramnfgssg pdvglttnss ssnnnssctg irrtygnsny lmrmqmrqea laidgsakld
301 mllqqqkasg ppqyyyaegy tqggskrrlf gscdgrrhly cmrsasvksl rqqqqqqql
361 ggsgdccarm qrmrqqqlnm tagndgerrk slstpslrit eatlrrsnes ksleskkrvv
421 kmlfvlvlef ficwtplyvi ntmtmligpv vyeyvdytai sflqllayss sccnpitycf
481 mnasfrrafv dtfkgmrlcd ggrfgfcrrr sknetnlsva gnsialansa msshtilesp
541 rl
```

**M CCK R 17D3 PB (Genbank Ref #** ABW09450**)**

1 mfnyeegdad qaamaaaaay ralldyyana psaaghivsl nvapyngtgn ggtvslagna
61 tssygddrd gymdtepsdl vtelafslgt ssspspsstp asssststgm pvwlipsysm
121 illfavlgnl lvistlvqnr rmrtitnvfl lnlaisdmll gvlcmpvtlv gtllrnfifg
181 eflcklfqfs qaasvavssw tlvaiscery yaichplr***sr*** ***swqt***ishayk iigfiwlggi
241 lcmtpiavfs qliptsrpgy ckcrefwpdq gyelfynill dflllvlpll vlcvayilit
301 rtlyvgmakd sgrilqqslp vsattaggsa pnpgtssssn cilvltatav ynensnnnng
361 nsegsagggs tnma***tttltt*** rp***taptvitt*** ***ttttt***vtlak tsspsirvhd aalrrsneak
421 tleskkrvvk mlfvlvleff icwtplyvin tmvmligpvv yeyvdytais flqllaysss
481 ccnpitycfm nasfrrafvd tfkglpwrrg agasggvgga aggglsasqa gagpgaya***sa***
541 ***ntnis***lnpgl amgmgtwr***sr*** ***srhe***flnavv ttnsaaaavn spql

**M CCK-R 17D3-like (Genbank Ref #** XP_032295927.1**)**

1 mysasaedaa saaatykall dyyanarsaa shivsltlap lneslslgla eagngnasgn
61 anssasyedd alsaenifli tesvatessr gaaaaangva vgsgrsssss aptempawli
121 psyslillca vvgnllvist lmqnrrmrti tnlfllnlai sdmllgvlcm pvtlvgtllr
181 nfifgeflck liqfaqassv avsswtlvai sceryyaich plr***srswqt***i shaykiigfi
241 wlggilcmtp iavfsrlipt srpgfckcre hwpdqgyerf ynimldlill vlpllvlcaa
301 yilitrtlyv gmnvgkdarm pa***ssgsaqt***v ataaiaatpg ssscvlvlna aseynessnn
361 nnaattstta aa***tattttta*** ***tatttatatt*** ***tlttittttl*** ***ttvt***pakssn aspslrihda
421 alrrsnetrt leskkrvvkm lfvlvleffi cwtplyvint lsmfigqtly eyidyttisf
481 lqllaysssc cnpitycfmn asfrrafvdt fkglpwkrng taagglsasq ggvpnlsnpn
541 pglamgmdtw r***srsrnd***qll nsvvytnsaa aaadspql

**M CNMa R (Genbank Ref #** ABW09450)

```
  1 mdmeyitsss gnitattead fssslgesnv teynttemda nesagedeem lriaffighf
 61 vhqyyipvlc ctgsignils vfvffrtklr klsssfylaa lavsdtcfla glfaqwlnfl
121 nvdiynqnyf cqfftffsyl asfcsvwfvv aftverfiav iyplkrqtmc tvrrakivlf
181 cltlvgclhc lpyiviakpv fmpklnttic dlnseykeql alfnywdtiv vyavpfttia
241 vlntctgctv wkfatvrrtl tmhkmkpqtn smpsnssnss ggassavasy rlsaslkrqk
301 stgthpsgqh nvanrqtddq eqqqsqqhq inncqhhcei tqkparrkvq nssqlkvtkm
361 llivstvfvc lnlpscllri eaywetesar nqnstialqy ifhaffitnf ginfvlycvs
421 gqnfrkavls ifrrvssaqr eagntqvtvs eycrntgtst rrrmmtqhcw nemhelhplk
```

**V CNMa R (Genbank Ref #** XP_002047272.3)

```
  1 manpyepttm tamsntsqdi sapttttttt tattatvttt lsyttdssrn isidleymsv
 61 edeemlhiaf lisdfvnryy vpiicctgsi gnilsvfvff mtklrklsss fylaalaisd
121 tcflcglfmq wlnflnvniy nqnyfcqfft fisylasfcs vwfvvaftve rfiavmyplk
181 rqimctvrra kivllgltla gcvhcvpyil iakpvyspkl ndticdlnts ykeqlalfny
241 wdsivvyavp fttitvlntc tgctvwkfat vrrtltmhkm kpqitnvpan attgggvaaa
301 tyrisaslrr qkstgthpsg qhsvsrqteq qqqqqqqrh darsqhhcei tqktgrrkvq
361 nssqlkvtkm llivstvfvc lnlpscllri etywetqtsk tqnttivlqy ifnaffitnf
421 ginfvlycvs gqnfrkavls ifrrvssaqr egitqvtvse ycrntgtstr rrmmtqhcwn
481 emhelhplr
```

**M 4313 R PD (Genbank Ref #** AAF45710)

1 mfppksgptp hpisialpls dpyatdhmad qdavlpvpls dsldldvdvd lnlnmnlnln
61 vddrrqvlfe gysdelltia wvacivfiiv gvpgnlltiv alsrgrqtrn staifiinls
121 csdllfgcfn lplaastfke rawthsdllc rlfpmlrygl lavsllsvsl itinryiiia
181 hprqypriyq rrylalmvag twittfsimi ptwrgvwgif gldvsigscs imhdrygrsp
241 keflfiaafm vpcicivicy arifllvrka airagtagkt nvsdvtpssa pqhqiqamat
301 pkkpekvtts sgeanepiag rpfvveenla yiddnastds lpisysirrr dqdqqpppvd
361 anvvlkerek erdrdqekvs lgr***sqtqle***m gkthgknpi***t tslrttsftr fs***prkshyas
421 mgntsnassi ypgrmsakdr rllkmilvif vwfvicylpi tvakiwksat evhwfniagy
481 lliylttcin pliyvlmsse yrraywnllr chgspdtqkq rnqanakrkh lesnrqvkat
541 t

**V 4313 R PD-like (Genbank Ref #** XP_002057364.1)

1 mihqldsqaq gldiqllgvp gavaqplala rkaggtnrsn smdqenelfe gysdelltfa
61 wiacivfiiv gvpgnlltiv alsrgkqtrn staifiinls csdllfgcfn lplaastfke
121 rawthsdllc rlfpllrygl lavsllsvsl itinryiiia hprqypriyq rrylalmvag
181 twlvtfsimi ptwrgvwgrf gldtsigscs ilhdkydrsp keflfmaafm lpcvcivicy
241 arifllvrqa amragaksse laitppiqtp tqtqkppvad kpkdkskvks kdrqqeldkd
301 adfgtsrpfv vaeslayidd nassesfpis ysikqeapid anvvlqdnnl annkhsvsgs
361 apatgpatad atvnk***sqsql e***qgrnsknpi ***ttslrtsftr fs***prkshyvs mgntsnassi
421 ypgrmsvkdr rllkmilvif vwfvicylpi tvakiwksan dvhvfnimgy lliylttcin
481 pliyvlmsse yrraywnllr chqteqqqqq qralkkhles nralkt

**M 12290 R PA (Genbank Ref #** AAF45710) **(starts just before TM2)**

1 maidllilal llvsflinll alcafwitpg lrttanrfti nllainligc cilaptlflg
61 lpgksaeast snaetleffs kpgnhqvilr rngqlveqdg vvvrrnisen gdtvetffkc
121 natycrelti dergdggfvi tetetheenl safeslptea pilppvqlrc wsidmtaalg
181 alavllvvgd twcavtdplr yhsrisgvkt wifialtwvv gilfgalsaf rvldfeadal
241 fsrqrrlavt yfnisstnsi fgvvyasvyf iviillpfgf vcgmywrifs eargnglrmr
301 qngsspllqs alnltagqqa aqanqfsnsl cvhrh***sissa*** ***ss***hggnsslg lgglqmqidq
361 rqqprsspsc lrrdsaakvl lptisddggs daesgagvql mpvqehslsd rnqnimltlq
421 tasgeikrny sarqlpllgt ssqdlretnr lqgirqvhss pnlhkytelr qdslseecgs
481 phllghaqrq qqqqlhlqhq qqqhhhhqqh hphfssprhq qhghalqipa ihaspkalsy
541 msslrhrlsn asslfkyree sraarisilv vvmfvvsylp fgllvllqsr lsaanfggss
601 qlaifmilla nlsspfifay rnkrvrrgvk rlfgldsssg lqrnc***sssvk*** ***tngt***agpaas
661 gaqlqrns***sk*** ***lsqyssns***ck yltpqsslvs qvpvhttltl rpnsscstii nfggargsad
721 sdeqppatpp ptvavapppt rpqrpklqrg itivehiait ptmpqkfqnr rarlfdmffr
781 sskklqagcq sqslptev

**V 12290 R (Genbank Ref #** XP_002053408.1)

1 maidflilav llisfiinll alcafwitpg lrttanrfti nlliinligc cilaptlfls
61 gfkslsgqee qlsgvnaagd siefyskpgn hqltirhhgq lvekdgivlr knitsgnsns
121 ndssgdtiem iykcnatycr eltidergdg gliitetetr ednlsvsvhs ntslpavqlr
181 cwsidmtaal galavllvvg dtwcavtdpl ryhsrisgvk awifitltwv vgivfgalsa
241 frvldfetdt llsrqrrlaa tyfnisstss ifgviyacvy fiviillpfg fvcgmywrif
301 seargnglrm rqngsspllq salnltnhaq aapatpyans lcvhrh***siss*** ***ass***hggggvl
361 sgglqmqidq raprnspscl rrdsaakvll ptisddgsdv dvesshggqq lmsvpeqtva
421 drnqnilltl qtasgeikrn ysarqlpllg tssqdlrelh rlqgirqvhs spnlhkytel
481 rqd***sltlse***e cssphllqah rqqqqhsrql hqqqqqlqql paqqqlhthf stgpqqvaag
541 halqipaiha spkalsymss lrhrlsnass lfkyreesra arisilvvvm fvvsylpfgl
601 lvllqsrlsa anftgstqla ifmillanls spfifayrnk rvrrgvkrlf gldaasalkr
661 qhssslknhg hsaaassapq lqrns***srlsq*** ***yssns***crylt pqsslvsqcq tttpvhttlt
721 lrpnsscsti inygghhkgs adsdevpsle atpppqrqtr pklqrgitiv ehihiagspp
781 kfqsrrglld mffrgskkmq sscseampte v

**M 13229 R PA (Genbank Ref #** AAF58717)

```
  1 mmqetgnqmg qthmhqrvpf ndtvlkdyhl tstdiekfvk lwqeyqmknm tpqvdecqgy
 61 cqgeiynwlr aynsihgyvs lmicifgtia nilnimvltr kemaktpinn ilkwlavadm
121 fvmleyipyt syqyiymgpg ekdlsytwav cllvhmhftq ilhtisiglt vtlavwryva
181 irhpnggcan fllahsreai llpfilspil clptyfvfqv retydvdkvn seamyhvyfd
241 kdsvlyrfnf wihsvlikll pcgilivisa vlmhvlceas rrrlklrdyn npakyaiqln
301 lnetkskkpp rcdrrndrtt lllvavlvlf litefpqgll gllsgvmekc ffahcyppfg
361 elmdllalin aavgfvlygl mskqfrttfr slfmkrhfgs temtrltrvt ttcv
```

**V 13229 R PA (Genbank Ref #** XP_015024303.1)

```
  1 mqetnmaqrq rephvalnet llkdleitss diqdfvkllf dfqkknhsqq decqgycqge
 61 iynwlrayng ihgyvsllic ifgtianiln imvltrrema kapinnilkw lavadmfvml
121 eyipyttyqy iymkpgekdl syawavyllv hmhftqilht isigltvtla vwryvairhp
181 ngscanflla hsreailfpf iispivclpt yfvfkvretl evdtrehevm yhvyfdvdsm
241 lfrfnfwihs viikllpcci ltvislvlmh vlceasrrrl klkdydnptk yaiqlnlnet
301 ksrrpprcdr rndrttlllv avlilflvte fpqgllglls gvlekcffah cyppfgelmd
361 llalinaavg fvlyglmskq frttfrslff krhfgssemt rltrvtttcv
```

**M 13575 R PA (Genbank Ref #** AAF58717**)**

1 mallhytfdq lelylewafa qhgeatpips iqpypgvfvg dlsqlnrfkr hafsavvgtl
61 fvlafcgnls tlyvnsrrkl rpffraclis lacsdlvssi fctvsymaqf qaqylqlwti
121 ggfmckfvpf itttsvlsgs ltlvaialdr ylavmrpvlg fwspdkrfst lsmlliwacs
181 igssgpllgi ydyrkiylld vedsseesee vvtavpeelv vtelemvhmc lagdhdvgly
241 yvilftlifl pcivsflwln aviarqlwlr rhyhqeqqeq hqepkegqfk tmanggdllm
301 pstlvsamgv avpfaldntp lppkstvnap gkkttaaala rearhrkmvv vvllmmavfi
361 clrlpawvfl imrlygsyse pidwllyfsf gilnlfscal npifytfltq tirtltlvkh
421 kiqgflgcpp gkvpdgmptd qmdksgcccg lrpptftwrc hpsrdraaat virdvdqpdp
481 psdqvqpdps slrrflsykq evftiykqcg ds***ssasies***s a

**V 13575 R PA (Genbank Ref #** EDW60637.1**)**

1 malrsliidt ddlypivias dlcqihdtpr yifmalvtvl flctfvgnvs alyvntrrkl
61 rpffraclis lacsdliycv nfttsntamf naeyleywil gpfmcnfvpf vntttvlcss
121 fmlvaialdr ymairraaig iwnpgfvfcg vciagiwlac maaavplffi ytpiqvyiqn
181 tdelliseld qatmcvgrrt qigiynsvsl slvfvpciva fvflnatiar qlwqlrhqqr
241 nlqqqqqqqr eqdqprfvhl lnkpettyam mtafsvaasf dmstaqltgl ppplplplpe
301 klspaaaarv arhrrmvrvv llmmgafmcl rlpawtfllm rvygsfsspv swlfyfsfgl
361 lnltscalnp lfytflpqti rvlsklkral srllccrras klesdatmpq etaerarrcl
421 ccglqvtwrc hlksppaaag svvtqtvavi eapsaasslp pvghckddyk dlaiynen***sl***
481 ***qttasmks***sr

**M 13995 R PA (Genbank Ref #** AAF52333)

```
  1 mnrdnlqqww ensyrrqhpe ptddlgldsa elhlalqepn qlpadydygn fslgnpydvd
 61 sehsispltl lllavsyglv vfggvvgnst lvltlcsass vrlrnpllla vciadllvtg
121 isapvtllnl amnrrtrslp lvlckvihyv qvmpvsasti sffmlsldry atvkhprlaq
181 lrqrrylhvs lallswlasa aistpflfay kiiaksmvvk gggaanttpn pvsisctsdl
241 ganamfmsfi ifhtiavfvl pgigvllnhy gvrrklcals ltaraahgel plpipilrrq
301 thmvivtgcp naqqaacggg ttaddtsngn gtgtgggpma vspgdiqlht lqprqpgsag
361 salepgsyrs snpispramr eirahsqrqr inragrgpat pgiplpqtst lrsrrhlanm
421 liasavifia cwaphvfcif yknfgnnqqc sqtsvyfsll lgyfysaisp viywalnhns
481 lrqspcapii rlrsmqnflr srfrthtapp ppsstneaal gafnpklikl tpkqyraqas
541 shyly
```

**V 13995 R PA (Genbank Ref #** XP_015028190.1)

```
  1 mnsnnlqqww ensyrrqhqq qptvdsnidn dndsdateql hyalleqnnf ggnsggidgn
 61 fggqlsssfd ydsygnftlt npydidtgad tehaipslti lllaisyglv vfggvvgnst
121 lvltlcsass vrlrnpllla vciadllvtg isapvtllnl amnrgtrslp lllcklihyv
181 qvmpvaasti sffmlsldry atvkhprlaq lrqrrylhvs lailswiasa aistpflfay
241 kiiaksviik gagtgttpnp vsisctselg anamfmsfii fhtiavfvlp gvgvllnhyg
301 vrrklcalsl taraahgelp lpmpilrrqt hmvivtgcan aqqagcggat taddtsngng
361 ngggqiainp gdiqlhnlqp crpgssaale pgsyrssnpi spramreira hsqrqrifra
421 grgpatpgip lpqtstlrsr rhlanmlias alifivcwap hvfcifyknf gykqycskts
481 vyfslllgyf ysaispviyw alnhntlrqs pcapiirlrs mqnflrsrfr shtvppaass
541 tneaalgafn pklikltpkq yraqasshyl y
```

**M 30340 R PA (Genbank Ref #** AAF52333)

```
  1 masvsssddf dfgkwdfpae riwlhkpnge itwkictflp liafglygnf smvyviatnr
 61 slrsptnlii anmavadllt laicpamfmv ndfyqnyqlg cvgcklegfl vvvflitavl
121 nlsvvsydrl taivlpmetr ltirgvqivv vctwvsgill asplafyrsy rvrvwknfte
181 ryckentsvl pkywyvliti lvwlplgiml icyiaifykl dryekrvlsr enpltvsykr
241 svaktlfivv vvfaalrlpf tilvvlreky fgedvsvssg mqlfwyisqy lmflnaavnp
301 liygfnnenf rrayyqiswv rrwrdatqmk kfsrspdhcc ycafmkngkr tseaaqkagn
361 lekdiskdms saqqsakstk ivenefvsei eadgfi
```

**V 30340 R PA (Genbank Ref #** XP_032293011.1)

```
  1 mtaynysiqq fdfsqwdfpa eriwlhkane eiawkiisfl pliifglygn yiliyliatn
 61 ralrsptnli ianmamadfl tllicpamfl indfyqnyql gcvgcklegf lvvvflitav
121 lnlsvvsydr ltaivlpqet rltlcgariv iagtwlagll lalplaiyrq yrvriwrnft
181 eryckenmtv lpkywyvlit vlvwlplgim licytaifvk ldryekrvls renplsvryk
241 rsvaktlfiv vivfvllrlp ftifvvlrek yystessvdc gmkyfsyfsq ylifvnaavn
301 piiygfnnen frrayaqiac mqkrraanan rihhclycdf iqnnksgqan aeqrsdkeis
361 qsaaretkkl gatsnidetl mpqlkgegfi
```

**M 32547 R PC** **(Genbank Ref #** AAX52506)

1 mspaeqlrlv gvsdayqlaa ssggaggggg gggggggggg lggyggggsg gdaggsgekm
61 kdvpdkyvta lshfldwhsn gtvdmerlsg pilkssiksv ywlfliqyaa lallgvvlnv
121 iivvyimyhr lykdvthafi inlalchfvq calvlpvslm vmliqnwifg qflcfflpml
181 qdiplhvami shiliawdrm rwlndplkgr lpgfvcccat wltgmvialp ypiytiyvel
241 gdympqlsgi glcvvnlmdd mqeytrglfl lmycgpaill sylyirtsqe lrppdgpfav
301 mmyehradlr mrqrns***stss ve***prhlsggg vaglsngggn sarsydlysa eldvhrekrk
361 qrnfgsmaat qvvcmcplmi lrfarlslee tyenakhfdf tylmfvwvaf lptvifpciy
421 asqilprdeq erlrgyfrls skrkqksqrr sdagggsgvg gssredsiek deasnttsvh
481 haaephkhss klrherevrl pghgssaggd rytgapyrqg kdrererererd rdrerdrdre
541 rererqragg rgagdagvvn nlgkdvrgkk hdvkiniisd gvghgatshs haahpahrkq
601 tggnnnnnnn ssrsrtnpgq rhggskltad cl***snvtast***f cngsgsvtag atgngngpsg
661 vlipddssts nygdgeessv vssmmvppqr wpgsgggvlr hkdv***sfsecs stfssstle***r
721 dleimdqler ersmdiqeml qrererekvr rqlpdiekly aqrspkgkrg eaagagagtg
781 aglalvmpgs dplsslsq***sr sisteyslcs*** **tlet**sgagvs vghdevlphh leeveeevvp
841 pdfydsytpg gtqnlppnag gavpvataav payqyqysrn rlsrkssgss sghhhhghhh
901 gqhggggvgg vaggrgskrd sfnslngtdl iagafceldp tqplnkmavi egrmrrsspr
961 na***sfssgsgv sgrs***smksss rdydyghgss shsahpeldf renifael

**V 32547 R PC** **(Genbank Ref #** XP_002059063.1)

1 mppaehhvat pqaagnddfq leastsaekw kdvpdkyllv iahllaghsn etedmerfng
61 pilkasiksv ywlfliqyaa lallgvilni aiivyimyhr lykdvthafi inlafchfvq
121 calvlpislm vmllqnwifg qflcfflpml qdiplhvami shiliawdrm rwlndplkgr
181 ipgfvcccat wltgmvialp ypiytiyvel gdylpqlsgi glcvvnlmdd mqeytrglfl
241 imycgpavll sylyirtsqe lrppdgpfav mmyehrvdlr mrqrn***sstss e***prthsgggv
301 aglsnghgts trsydlysae ldvcrekrkq rnfgsmaatq vvclcplmil rfarlsleet
361 yenqkhfdft ylmfvwvafl ptvifpciya sqilprdeqe rlrgyfrlsa krkpksqrrs
421 nag***tgslqd***d siekdehtnt tsvavnvagq tqppaesqkh ppkgnavslr herspghgls
481 stdrysgaph rqgqsavker drerdrdrer erdrdrerer erererg rdr ererrerggh
541 vlsgrgagda gvvnnlgkhm rgkkhdvkin iisdghgstg sgrkqpp***sss snns***sgrskp
601 aagqarpkpt ieym***svsnns*** vsnltassyc ngnnsallddd sgtsnygdge essvvssvmh
661 ppqrwphvag lrhk**di*****sfse csstfssst***l erdleiidll erersmdiqe mmqreqqgek
721 vrlsvggvgr qlpdieklya qrspkpkrds gygydkgala lvmpgsdpls msla***sssqpt***
781 ***eyslcs***nlqe hqlhqqaqqr lghlqqqqlq qqqqqlhqhq qldeeevpqd fydsytpggs
841 nnlppavqav qvaapapppp tyqyqytrrl srksspgpsh hvpgggtarg skrdsfnsln
901 gtdliagafc eldptqplnk maglegrmrr ssprnasfn***s nsars***smksr dyghgsthsa
961 hpeldfreni fael

**M 33639 R PA (Genbank Ref #** AAF48813) **(PA-specific underlined)**

1 mitrlyntee dpaycsfiwg snltssvdvl aanatsvfss dlrddfyrdv edprteslre
61 ycyglvlpii camgiignvl nlvvltrrnm rgtayiymra ystaallaiv faipfgirml
121 vhkdrgqwee fgpafytahl elylgngclg vgvmmllvlt ieryvsvchp gfarpvmgpp
181 gvvvfltcla tvivylpsif rgelikcilg ssdvyvylrr dntiyqqtif yrvykimlev
241 ifklvptlvi gglnmrimmv yrrtcerrrk mvlsrphaqg hghghghghg hghghahghg
301 ylkdddprkf aeerrlflll gstsilflvc vspmailhmt iasevypsfp fqvfrasanl
361 lelinysltf yiyclfsedf rntlvrtikw pwlkgkfchq aehevsaspp atagtvavag
421 tgnghvsifh paipaltltp aepderprca ngvlh

**M 33639 R PD (Genbank Ref #**AGB95518.1)

1 mitrlyntee dpaycsfiwg snltssvdvl aanatsvfss dlrddfyrdv edprteslre
61 ycyglvlpii camgiignvl nlvvltrrnm rgtayiymra ystaallaiv faipfgirml
121 vhkdrgqwee fgpafytahl elylgngclg vgvmmllvlt ieryvsvchp gfarpvmgpp
181 gvvvfltcla tvivylpsif rgelikcilg ssdvyvylrr dntiyqqtif yrvykimlev
241 ifklvptlvi gglnmrimmv yrrtcerrrk mvlsrphaqg hghghghghg hghghahghg
301 ylkdddprkf aeerrlflll gstsilflvc vspmailhmt iasevypsfp fqvfrasanl
361 lelinysltf yiyclfsedf rntlvrtikw pwlkgkfchq aehenptngp gvpmacftkv
421 drghqkhhit ttsglgrsss i

**V 33639 R PD-like (Genbank Ref #** AGB95518)

1 mitrlyntee dpaycsfiwg anltssndlv lgttansthi yandlrddly advedprtes
61 lreycyglml pvicalgiig nvlnlivltr rnmrgtayiy mraystaall aivfaipfgi
121 rmlvhkdrgq weefgpafyt ahlelflgng clgvgvmmll vltieryvsv chpgftrpvm
181 gppgvvvflt cfatfiiylp sifrgelikc mltsnnvyvy lrrdnniyqr tifysvykim
241 levifklipt vliaglnlri mlvyrrtcer rrqmvltran yvkdddprkf aeerrlflll
301 gstsilfllc vspmailhmt iasevlpsfp fqvfralanl lelinysitf yiyclfsedf
361 rntlmrtikw pwlksklchq vdetqtikgv pmvrfdkvtl rnhhitttsg igrsssi

the *D melanogaster* PB and PC isoforms matches PA; PD differs by an alternative splice but does not introduce a BBS. The only *D virilis* isoform received by Blastp search was orthologous to PD.

**M CNMa R PA (Genbank Ref** AAF50229.3)

1 mdmeyitsss gnitattead fssslgesnv teynttemda nesagedeem lriaffighf

61 vhqyyipvlc ctgsignils vfvffrtklr klsssfylaa lavsdtcfla glfaqwlnfl

121 nvdiynqnyf cqfftffsyl asfcsvwfvv aftverfiav iyplkrqtmc tvrrakivlf

181 cltlvgclhc lpyiviakpv fmpklnttic dlnseykeql alfnywdtiv vyavpfttia

241 vlntctgctv wkfatvrrtl tmhkmkpqtn ***smpsnssnss*** ggassavasy rlsaslkrqk

301 stgthpsgqh nvanrqtddq eqqqqsqqhq inncqhhcei tqkparrkvq nssqlkvtkm

361 llivstvfvc lnlpscllri eaywetesar nqnstialqy ifhaffitnf ginfvlycvs

421 gqnfrkavls ifrrvssaqr eagn***tqvtvs*** ***e***ycrntgtst rrrmmtqhcw nemhelhplk

**V CNMa R PA (Genbank Ref #** XP_002047272.3)

1 manpyepttm tamsntsqdi sapttttttt tattatvttt lsyttdssrn isidleymsv

61 edeemlhiaf lisdfvnryy vpiicctgsi gnilsvfvff mtklrklsss fylaalaisd

121 tcflcglfmq wlnflnvniy nqnyfcqfft fisylasfcs vwfvvaftve rfiavmyplk

181 rqimctvrra kivllgltla gcvhcvpyil iakpvyspkl ndticdlnts ykeqlalfny

241 wdsivvyavp fttitvlntc tgctvwkfat vrrtltmhkm kpqitnvpan attgggvaaa

301 tyrisaslrr qkstgthpsg qh***svsrqt***eq qqqqqqqqrh darsqhhcei tqktgrrkvq

361 nssqlkvtkm llivstvfvc lnlpscllri etywetqtsk tqnttivlqy ifnaffitnf

421 ginfvlycvs gqnfrkavls ifrrvssaqr egi***tqvtvse*** ycrntgtstr rrmmtqhcwn

481 emhelhplr

**M CRZ R PA (Genbank Ref #** AAF49928)

1 medewgsfdr lpsvpsasmd letenevvsn wstlanftrl vagaapeivn ytlnmidvgv

61 gmatdisnls vsttplpaya isnssslaht nsrheappma eqvpehvmdh apqlsrsgll

121 kvyvlavmal fsllgnllti wniyktrisr rnsrhtwsai yslmfhlsia dvlvtwfcii

181 geaawcytvq wlaneltckl vklfqmfsly lstyvlvlig vdrwiavkyp mkslnmakrc

241 hrllggtyil slvlslpqff ifhvargpfv eefyqcvthg fytadwqeqm yatftlvftf

301 llplcilfgt yms***tfrtiss*** ***se***kmfqgskl anystaklpt qtnrqrlihk akmkslrisv

361 viiiaflicw tpyyvmmimf mflnpdkrlg ddlqdaifff gmsnslvnpl iygafhlcpg

421 kggkssgggg nnnayslnrg dsqr***tpsmlt*** ***avtqvd***gtgg ssrqmrafrq qsyyr***sssng***

481 ***t***agpgaapfk eqvgllhvgp gngtpgg***svs*** ***sgat***pqlirk gsallarqps clreqehqqr

541 lllhekpstl vlsydsqrgg vgvgvasgll dnnervssv

**V CRZ R P (Genbank Ref #** XP_032290616.1)

1 megledihaha rlpvgnttvp ihsdlldnay netdsnwstv anftrliiaa asevapnian

61 ytlnmldvgi alateatnae satstampgs nrgteaayai mtnsrlnsst lghinslhel

121 spvaeqvpeh vmdhapqlsr sgllkvyvlt vmalfsllgn lltiwniykt ritrrnsrht

181 wsaiyslmfh lsiadvlvtg fcligeaawc ytvqwlanel tcklvklfqm fslylstyvl

241 vligvdrwia vkypmkslnm akrchrllgg tyilslvlsl pqffifhvar gpfveefyqc

301 vthgfytavw qeqmyatftl vftfllplci lfgtyms***tfr*** ***tissse***kmfq gsklanystt

361 kqlptqtnrq rlihkakmks lrisvviiia flicwtpyyv mmiifmfwnp dkrlgddlqd

421 aifffgmsns lvnpliygaf hlcpgksnks gtgggnnnay slnrgdsqr***t*** ***psmltavtqv***

481 ***d***aaggsarqm rtfrqqsyyr ***sssngt***aggp fkeqvgllqv ggaggsgatp hmmrk***sssvr***

541 ***s***phpnhssai atrhsgclre qeelllqtkp stlvlnydsq rggvgvgvan gnklmtgasl

601 rvdnkecvss v

**M ETH R PA** **(Genbank Ref #** AAF47700)

```
  1 mlpqipsyir ttamffcivi mllgvvgnvm vpivivktkd mrnstniflt nlsiadllvl
 61 lvctptvlve vntrpetwvl ghemckavpf veltvahasv ltilaisfer yyaiceplka
121 gyvctkgrai licvlawgia alftspilwv aeyklaeyid gssvavcltq aisdwtlaff
181 lmtisvffvv pfvtlvvlyg iiarnlvsnr aamlrarptk pelslkarkq vvlmlgavvl
241 sffvcllpfr vltlwiilst dqtlhdlglv ryysllyfcr imlylnsamn pilynlmstk
301 frrgfkrlcq dagrlllelv tlgrrkedss rgrrgtlslg mgtntntntn ssnatgatss
361 silsrssnrr csedisrtrl kiemqmpcgs dleamamlqh stlgkgiarr vsdsrlmplr
421 nhqprrhkpq isfdeeslee nkrseakipt kcreklpgia reivnltent l
```

**V ETH R PA-like** **(Genbank Ref #** XP_002058475.1)

```
  1 mlpqipsyir ttamffciii mllgvignvm vpivivktkd mrnstniflt nlsiadllvl
 61 lvctptvlve vntrpetwvl ghemckavpf veltvahasv ltilaisfer yyaiceplka
121 gyvctkarai licvlawgia alftspilwv aeyklveyid gssvpvcltq aigvstvgff
181 lmtisvffvl pflmlvvlyg iiarnlvsnq gamlrarptk pelslkarkq vvlmlgavvl
241 sffvcllpfr vltlwiilst eqtlhdmglv ryysllyfcr imlylnsamn pilynlmstk
301 frkgfarlch dvarfllkll tlgrrranpa dsrgrsgtls tgtgtqtnss natsatnssi
361 lsrssnrrgs edisrtrlki elqlpcgsdl eamamlqnst lrskrrqsds rllgekkprr
421 pkpqisfdea avgrapketv ipa
```

**M ETH-R PB** **(Genbank Ref #** AAF47700)

```
  1 mlpqipsyir ttamffcivi mllgvvgnvm vpivivktkd mrnstniflt nlsiadllvl
 61 lvctptvlve vntrpetwvl ghemckavpf veltvahasv ltilaisfer yyaiceplka
121 gyvctkgrai licvlawgia alftspiiai stysvepygd gtdapvctta adgfwsifyf
181 vgcitvfffl pfgilvllya aiaykllrpn nafhrptspq pqqpsggats gssqvpstkg
241 nshqqsngmr khrkqvifml vavvssffvc llpfraftlw vilasaedve glgiagyynl
301 lyfsrfmlyl nsamnpilyn lmsskfrsgf wrllltclgq rphhhhrhhy hqrqhptagg
361 sgrnastrqe qdaeegaala gttsarhprr tlrreatfli nsistssgtd rrttsssawrs
421 nslsisglse rergilgaai igttaatvtt aclqerrask i
```

**V ETH R PB-like** **(Genbank Ref #** XP_015024858.1)

```
  1 mlpqipsyir ttamffciii mllgvignvm vpivivktkd mrnstniflt nlsiadllvl
 61 lvctptvlve vntrpetwvl ghemckavpf veltvahasv ltilaisfer yyaiceplka
121 gyvctkarai licvlawgia alftspiiai ssysvepygd gteapvctta adgfwpilyf
181 vgtitlfffl pfgillllya aiaykllrpn safhrptspq gptptaataa snkgnshqqs
241 ngmrrhrkqv ifmlvavvas ffvcllpfra ftlwvitass edverlgiag yynllyfsrf
301 mlylnsamnp ilynlmsskf rsgfwrllls clgrrphhhr hhhhyptiae rslrqtqrrq
361 dhepteavpg tdnaqrprrt lrreatflin sistssgter htsstwrsns lstslglger
421 erdrgalgaa iigttaatvt taclqerras kt
```

**M FMRFa R PA (Genbank Ref #** AAF47700)

1 msgtavarll lrlelpspgv mppppptdydy ggpisddefl asamategpt vrydlfpqnn
61 sqptlqivln htevqtdlqy phyedlgldp dpnwtriced vynpllennr iefwvcgvli
121 nivgvlgilg niismiilsr pqmrssinyl ltglarcdtv liitsillfg ipsiypytgh
181 ffgyynyvyp fispavfpig miaqtasiym tftvtleryv avchplkara lctygrakiy
241 fivcvcfsla ynmprfwevl tvtypepgkd vilhcvrpsr lrrsetyini yihwcylivn
301 yiipfltlai lncliyrqvk ranrerqrls rsekreigla tmllcvvivf fmlnflplvl
361 niseafysti dhkitkisnl litinssvnf liyiifgekf krifllliffk rrlsrdqpdl
421 ihyessisnn gdgtlnhrss grfsrhg***tqr*** ***stttt***ylvat ggpggggcgg gggnnslnnv
481 rltqvsgspg lvkikrnrap spgpvvyfpa remqr***sastt*** ***ns***ttnnntsi gydwtlpdsk
541 klghvssgf

**V FMRFa R PA (Genbank Ref #** XP_002047744.1)

1 msaaaarlll rleqpvpgvm pppapaeydw spisvdqfla satgrsqeda vryeladplg
61 stlapliydv gqrllfngss gtqngndtyt dlfpqmedmg depnwtrice evynpqlenn
121 riefwvcgvi lnivgilgil gnvismiils rpqmrssiny lltglarcdt mliissmllf
181 gipsiypytg qffgyynyvy pfispamfpi gmiaqtasiy mtftvtlery vavchplrar
241 alctygraki yfivcvcfal aynmprfwev ltvsyqlpns tdvlhcvrps plrrnptyin
301 iyihwcyliv nyiipfltla ilncliyrqv kranrerqrl srsekreigl atmllcvviv
361 ffmlnflplv lniseafysv idhkvtkvsn llitinssvn fliyiifgek fkrifllliff
421 krrlsrdqpd lihyessisn ngdgtinhrs sgrfsrhg***tq*** ***rsttt***tylva tgggggnssl
481 nsvrltqvsg spgllkikrn rapspgpvvy ypgpremhr***s*** ***vstsns***ttnn ntalgydwtv
541 dgkklghvss gf

**M LGR1 PA (Genbank Ref #** AAF55460)

```
  1 mekhpslsqr mgttyrprkg lkclsfefqc rlllhhlllt slsgrhfvya tsavggalsa
 61 nnchdihhgf dvypnltavs laqstdtplt atmprsawkc ccwnasnqae evecrcegdg
121 lnrvpqtltl piqrltiasa glprlrhtgl kvygstlldv aftdclqlel iqdgafanlt
181 llrtiyitna pkltflskdv flgisdtvdi iriinsgltr vpdlghlpph nilqmidldn
241 nqitridsks ikvktaqlil tnneisyvdd saffgskiak lslkenkklq mmhpnafdgi
301 iditeldlss tslvglpsag lqniealyiq nthtlktips iynfrnlqra ylthsfhcca
361 fqfpsrhdpq rhaqrmleie kwrkqcksds gtrkerstld npfnmpedfg sfggtddsat
421 ditpitfasf dymaddtmnk gtfhekiiln pgddssaelc gnftfrkpni ecypmpndln
481 pcedvmgyqw lrisvwivva lavvgnvavl tvilsirpes tpvprflmch lafadlclgl
541 ylllvacida hsmgeyfnfa ydwqyglgck vagfltvfas hlsvftltvi tierwlaitq
601 amylnhrikl rpaalimlgg wiysmlmssl plfgisnyss tsiclpmenr dvydtiylia
661 ilgsngvafs iiavcyaqiy lslgretrqa hqnspgelsv akkmallvft nfacwspiaf
721 fgltalagyp linvtkskil lvffyplnsc adpylyailt sqyrqdlftl lsklglcqqs
781 alkykdslsg qattrftihg siqrhssltc kmqtvmgaet qkmlknsedy v
```

**V LGR1 PA-like (Genbank Ref #** XP_032289845.1)

```
  1 mkcmviisql dfltfillsl ahgtrqantn chdnnngfnt vfnnlsidng netdmsvimt
 61 qaptmstnpt sdasvwkccc weatnqnefe crcegealtr vpqtlklpll rltiasaglp
121 rlrsmglkvy attlldvafi dclqleaiqn gafsnltflr tiyisnapkl tylpknvfeg
181 isdtieiiri insgltsvpd fgylppnnil qmidldnnqi sridsksiqv ktaqfvlann
241 dihfiddsaf lgskiaklsl kdnrrltdvh pnafygiidi teldlsstsl vslpsaglqt
301 vevlyiinth tlktipsiyn fqnlqrahlt hsfhccafqf psrhdpkrha erlqelqkwr
361 eqcnverdly rdvdkklkkn tsvkdtagvh ytnqsgtatd nlltdaasns ydymadstmn
421 nigifheqit inpddnqlae ycgnftfrnp diqcfpmpna lnpcedvmgy qwlrisvwiv
481 valaivgnla vltvtlsiks espsvprfli chlafadlcl glylllliasi dahsmgeyfn
541 yafdwqyglg ckiagfltvf ashlsiftlt iitierwfai thamylnkri tlrqaagiml
601 tgwiysiims slplfgisny sstsiclpme irdiydsiyl ililgcnfva ftiiaicysq
661 iylslgqetr rarrnnpgem svakkmlllv finftcgapi affgltalag cplinvtksk
721 illvffypln scadpylyai ltsqyqqdll tflsklgicr qnalkykhsd slhgtshyti
781 rgsieqqssl cqkpqqegaa etqtmlknne dyv
```

**M LGR3 PA (Genbank Ref #** AAF56490)

1 mvygrsiavg fclmtvvlll aavifylslg pcpaasfacd ngtlcvprrq mcdsrndcad
61 ssdenpvecg llygskeiad kivrnaiekk qqrlisavsn asgadsttsm vprnqsltln
121 mtcdivtypk acqcgqgtil ycgryaklrr fprlssevtn liiirnnltl rdnifanftr
181 lqkltlkynn isrvplgsfs glfhlerlel shnnvshlph gvflglhslq wlflvnnhlh
241 hlpveqlrff rrlewlvlsr nrltlrnvql pkiptlyevy ldfnrieyig eetfsqldnl
301 hlldlqhnli thihgrafan ltnmrdirlv gnpikelsge tflhntrlea lslalmpihi
361 ssslmeplni sflnltgiry dhidfeains mrnltyiiyd rffycsmtpr vrmckpstdg
421 vssfqdllsk pvlrysawvm atltiagnvl vlwgrfiyrd envavtmvir nlaladmlmg
481 fylvtigvqd yryrneyykv vldwitswqc tligtlavss sevsmlilaf mslerfllia
541 dpfrghrsig nrvmwlalic iwitgvglav apvllwrtst lpyygsysgt cfplhiheaf
601 pmgwlysafv flgvnllllv miamlytall isiwrtrsa***t pltlld***cefa vrfffivltd
661 flcwvpiivm kiwvffnyni sddiyawlvv fvlplnsavn pllytfttpk yrnqiflrgw
721 kkitsrkrae agngnvattt tg***tatgss***qh pddftifaka amrch

**V LGR3 PA (Genbank Ref #** XP_002056481.1)

1 mvygrsiavg icfmtvvvvl sgliyylslg pcptgsfacn ndiqcvpqrq mcdkhldced
61 gsdenpvecg nlygskelad kivrnaiekk kqqqqqqen qrlmivtsns sawdspslag
121 srnqsmpipc aittyprqcq cgggtmlycg rfaklrrwpr iseevtnlfi irnnvtlren
181 lfanltrlqk ltlkhnnisr lppgcfsglp hlerlelghn nfsqlphfil kdlpalqwlf
241 lvnnqlrsfp mdhlaamhkl ewlvlshnrl tlrneqlsks pklveifldn nrieyigekt
301 faqldnlell dlqynlithi hvkafanlta isdirlignp ikelsgetfl ynthlealsl
361 aymplpidrn lmkslnvsfl nltgieferi dfaalneirn lkyivfdrfy ycsmtphvlm
421 ckpksdgvss lkdllskpsl rhstwfmatl siagnlmvlw grfiyrdenv avtmvirnla
481 ladllmgfyl liigvqdfrf rdeyhkvard wisswqckai gtlavsssev smlilafmsl
541 erflliadpf rghraisvri iyfsllciwv tgvglaitpv viw***sssttd***f ygthsgtcfp
601 lfiheayplg wqysafvflg vnlllllmia llytallisi wrtrsa***tpls lld***cefavrf
661 ffivltdvlc wapiiamkiw vffkynisdd iyawlvvfil plnsavnpll ytfttpkyrn
721 qillrgwkki tsrkrtetgh gnvantttg***t atgss***qnpde yttatgiksm plaltlsh

**M LGR4 PB (Genbank Ref #** ABW09404)

```
  1 mciahlpitf tlaillaias negaqgvesa trtaieairt gigtkpetei adateaeapv
 61 revisllgii dgaesdilvp daddkcpggy fhcnttaqcv pqrancdgsv dcddasdevn
121 cvnevdakyw dhlyrkqpfg rhdnlrigec lwpnenfscp crgdeilcrf qqltdiperl
181 pqhdlatldl tgnnfetihe tffselpdvd slvlkfcsir eiashafdrl adnplrtlym
241 ddnklphlpe hffpegnqls ililarnhlh hlkrsdflnl qklqeldlrg nrignfeaev
301 farlpnlevl ylnenhlkrl dpdrfprtll nlhtlslayn qiediaantf pfprlrylfl
361 agnrlshird etfcnlsnlq glhlnenrie gfdleafacl knlssllltg nrfqtldsrv
421 lknltsldyi yfswfhlcsa amnvrvcdph gdgissklhl ldnqilrgsv wvmasiavvg
481 nllvllgryf yksrsnvehs lylrhlaasd flmgiyltli acadisfrge yikyeetwrh
541 sgvcafagfl stfscqsstl lltlvtwdrl msvtrplkpr dtekvrivlr llllwgisfg
601 laaapllpnp yfgshfygnn gvclslhihd pyakgweysa llfilvntls lifilfsyir
661 mlqairdsgg gmrsthsgre nvvatrfaii vttdcacwlp iivvklaals gceispdlya
721 wlavlvlpvn salnpvlytl ttaafkqqlr rychtlpscs lvnnetrsqt qtayesglsv
781 slahlgggvg ggsgrkrmsh rqmsyl
```

**V LGR4 PB (Genbank Ref #** XP_015025567.1)

```
  1 mtimmmplls lgalvallsl tsmtlasity sveqprgtir ssvgqgntmf mlgssvvdae
 61 ademdeldsg illpdekdnc pagyfhcntt aqcvpqranc dgsvecddrs dewncvnevd
121 akywdhlfrk ssfgrqgmdd kpigvcywpn aknfscpcrg neilcrfqql talppqlptn
181 nlstldltgn nfgvidetff gglpaveslv lklcaireia shafdrlaai plktlymddn
241 elsrlpehff apgnqlrili larnqlssls sgdfrylqhl edldlrgnli shfeaqvfaq
301 lhslevlyln nnrlqqlrpg mfpsnlvhlh tlslahnqis siaantftfp qlrhlflagn
361 qlsyirdgtf cnlsylqglh lnenqiekfd lhafdclenl sslltgnrf ktlepavlqn
421 lssldfiyfs wfhlcraalh vrvcdprgdg isstfhlldn qllrgsvwim asiavvgnll
481 vlagryfyks rsnvehslyl rhlavsdflm giyltliaca disfrgqyir heeswrhsgf
541 cafagflstf scqsstlllt lvtwdrlmsv trplkprdte kirivlrlll vwgisfglaa
601 apllpndyfg ahfygnngvc lslhihdpya kgweysallf icintislif ilfsyvrmlq
661 airdsgggmr sthsgresvv atrfaiivtt dcacwlpiiv vkvaalsgca ispdlyawla
721 vlvlpvnsal npvlytltta afkqqlrryc htlptcslgn netrsqtqtg yesglsvsla
781 hlgggggagg giggsgrkrm shrhmsyl
```

**M LK R PA (Genbank Ref #** ABW09404)

1 mamdlieqes rleflpgaee eaeferlyaa paeivallsi fyggisivav igntlviwvv

61 attrqmrtvt nmyianlafa dviiglfcip fqfqaallqs wnlpwfmcsf cpfvqalsvn

121 vsvftltaia idrhraiinp lrarptkfvs kfiiggiwml allfavpfai afrveelter

181 frennetynv trpfcmnknl sddqlqsfry tlvfvqylvp fcvisfvyiq mavrlwgtra

241 pgnaqdsrdi tllknkkkvi kmliivviif glcwlplqly nilyvtipei ndyhfisivw

301 fccdwlamsn scynpfiygi ynekfkrefn krfaacfckf ktsmdaher***t*** ***fsmhtras***si

361 rstyanssmr irsnlfgpar ggvnngkpgl hmprvhgsga nsgiyngssg qnnnvngqhh

421 qhqsvvtfaa tpgvsapgvg vamppwrrnn fkplhpnvie ceddvalmel psttppseel

481 asgagvqlal lsressscic eqefg***sqtec*** ***d***gtcilsevs rvhlpgsqak dkdagkslwq

541 pl

**V LR R PA (Genbank Ref #** XP_002047380.1)

1 meadiiynqh iheellpgae eeaeferlya apaeivalls ifyggisiva vigntlviwv

61 vattrqmrtv tnmyianlaf adviiglfsi pfqfqaallq rwnlpyfmcg fcpfvqalsv

121 nvsvftltai aidrhraiin plrarptkfi skfiiagiwl lalvfaapfp iafrveeitd

181 rfrendvyfn ltrpfcmnkn lsdeqlkayr yalvfvqylv pfcvisfvyi qmavrlwgth

241 apgnaqdsrd itllknkkkv ikmliivvvi fglcwlplql ynilyvtipe indyhfisiv

301 wfccdwlams nscynpfiyg iynekfkref nrrfaacfck fktslepher ***tfsmhtras***s

361 irstyanssm rirnnffavr nagngktgln igrsayqang ngtgtgianv netssynnng

421 hqtvvtfags gstmppwrrn qfkplhpnva eceddlalme lpstneeapp smaeagvpla

481 lfkaesress sciceqefg***s*** ***rtecd***gtcil sevsrvqqpa tkgpdgvggi g***sgsese***ilw

541 qpl

**M moody PA (Genbank Ref #** NP_569970.2**)**

1 msdettisle dgyppleallt tmvppadatg fsqslltfaa vmtflimivg icgnlltvva
61 llkcpkvrnv aaafiislci adllfcalvl pfqglrfvqg twrhgqvlcr lipfiqygni
121 gvsllciami tinryvmith hglyariykr hwiavmiaac wlfsygmqlp tllgewgrfg
181 ydsrlqtcsi mtddhghssk ttlfitafvi pclviiacya kifwvvhkse qrlkrhatkq
241 nsipnnlrpl astgsgalps gaecqpsnrv ssdssssfsi dvpetapsgk qqptrvkdqr
301 evrakrnewr itkmvlaifl sfvvcylpit ivkvadknve hpslhicsyi llylsacinp
361 iiyvimnkqy rkayktvvfc qparlllpfg ktngassaae kwkdtglsnn h***srtivsqms***
421 ***ggt***gaasgag tatgtaavav mqtppevqqa qalemvsrgp dlisksnlpq pnvtppppsv
481 ltatpng***sns nslt***lrlplk knnhcy***tnsg fns***stpspss glgigissss iyrpgvgslg
541 sgsasirrit mvgddiilee eelpptppat sapttpappp pssplhpl***st dsstttis***gg
601 avvagssapk patptphiym nvdspkrnqy ymdrntnava pesdsgpant satvsisgsk
661 ltakmkfpkd

**V moody PA-like (not annotated; obtained by direct inspection of genomic DNA – see SXX Document – "de novo moody isoform annotations"))**

1 mnddttasps pglghgvddm ltaaptgtas gldavdesgf sqslltfaai mtflimivgi
61 cgnfltvval lkcpkvrnva aafiislcia dllfcavvlp fqglrfvqgt wrhgpvlcrl
121 ipfiqygnig vsllciamit inryvmithh gcyariykrh wiaimiaacw lvsygmqlpt
181 llgawgrfgy darlqtcsim sdahghsskt tlfitafvip clviiacyak ifwvvhkseq
241 rlkrhankqn sipnnlrpva aaspagqdgs vqptaaasrv ssdsssystd vpegkqpppq
301 srvkdqrevr akrnewritk mvlaiflsfv icylpitivk vadkdvehps lhifsyimly
361 lsacinpiiy vimnkqyrka yktvvlcqpa rlllpfgktn gassaaekwk dtglsnnh***sr***
421 ***tlvsqms***aga aaaaaaaaaa avaaasastp ppspasdcqs lpatkaspqs lelmsrcpdl
481 inrtnqpplv tpppp***svtla t***aqp***sssnss nssnsggs***rl plkksnhsyi nngfns***sghs***
541 ***qns***avyrpaa gsppsaaaap lrritmvgdd iileeeelpa vpsp***tatals vskvt***kspiy
601 mnvnspkrnq sysdkasvkd alqqasvpsd qgkdqqdask vpikfpnpkd

**M moody PC** **(Genbank Ref #** NP_001188535.1)

1 msdettisle dgypplealt tmvppadatg fsqslltfaa vmtflimivg icgnlltvva

61 llkcpkvrnv aaafiislci adllfcalvl pfqglrfvqg twrhgqvlcr lipfiqygni

121 gvsllciami tinryvmith hglyariykr hwiavmiaac wlfsygmqlp tllgewgrfg

181 ydsrlqtcsi mtddhghssk ttlfitafvi pclviiacya kifwvvhkse qrlkrhatkq

241 nsipnnlrpl astgsgalps gaecqpsnrv ssdssssfsi dvpetapsgk qqptrvkdqr

301 evrakrnewr itkmvlaifl sfvvcylpit ivkvadknve hpslhicsyi llylsacinp

361 iiyvimnkqy rkayktvvfc qparlllpfg ktngassaae meryrveqqp qphnrlpdvg

421 gngsrigsrd gngngsgggd adptggpt***sp*** ***slgd***gvagtg pnkqieppaa eryasatlgp

481 hgdtqrkqqq qphpataaqe eqsllhqqrl qqqhsqpqqr lgdrdqqqlh lparswltge

541 rlrldpshnn gggrhntggr gaaahasgdf cadnagtspt liptasavhg linnnnqrrs

601 sgsgeqctka gdanaphlhe crqpeekpil hgs

**V moody PC-like** **(Genbank Ref #** XP_002057362.1)

1 mnddttasps pglghgvddm ltaaptgtas gldavdesgf sqslltfaai mtflimivgi

61 cgnfltvval lkcpkvrnva aafiislcia dllfcavvlp fqglrfvqgt wrhgpvlcrl

121 ipfiqygnig vsllciamit inryvmithh gcyariykrh wiaimiaacw lvsygmqlpt

181 llgawgrfgy darlqtcsim sdahghsskt tlfitafvip clviiacyak ifwvvhkseq

241 rlkrhankqn sipnnlrpva aaspagqdgs vqptaaasrv ssdsssystd vpegkqpppq

301 srvkdqrevr akrnewritk mvlaiflsfv icylpitivk vadkdvehps lhifsyimly

361 lsacinpiiy vimnkqyrka yktvvlcqpa rlllpfgktn gassaaekwk emeryrieqq

421 pqphpclpdv rrrsssssgg gggggcsgig vdaaavarir lpvlagdqsv a***svtgad***vpm

481 ***srsnksnes***t aigyapaaig nagngsteqq qqqqqqqqrr qsaaaqeeqs llhqqrlqqq

541 raqpeqrrls arswlpavgs ggaiashhhg ggryhtgggg ***tarsaqsdgd*** gavsqqsnqi

601 ahlherqqse ekpil

**M Ms R1 PA (Genbank Ref # AAF47635.2)**

```
  1 masgnnetep lycgsgmdnf htsyknmhgy vslvvcilgt iantlniivl trremrsptn
 61 ailtglavad lavmleyipy tihdyiltds lpreeklsys wacfikfhsi faqvlhtisi
121 wltvtlavwr yiavgypqkn rvwcgmrtti itittayvvc vlvvspslyl itaiteyvdq
181 ldmngkvins ipmtqyvidy rnellsarta alnatptsap lnetvwlnas tlltstttaa
241 pptpspvvrn vtvyrlyhsd lalhnaslqn atfliysvvi klipcialti lsvrlilall
301 eakrrrkklt skpatpgasn gtkspangka adrprknskt lekekqtdrt trmllavlll
361 flitefpqgi mgllnavlgd vfylqcylrl sdlmdilali nssinfilyc smskqfrttf
421 tllfrpkfld kwlpvaqdem aaaraersav apvlekgrqq pqvvmasttt nitqvtnl
```

**V MS R1 PA-like (Genbank Ref # XP_002047415.1)**

```
  1 maggsndttp lycgtgtdnf htsyknmhgy vslvvcilgt iantlniivl trremrsptn
 61 ailtglavad lavmleyipy tvhdyilkds lpreqklsyg wacfikfhsi faqvlhtisi
121 wltvtlavwr yiavghpqkn rvwcgmrtti itittayvvc vlvvspslyl itaiaefmdq
181 mdvegnliss ipmsqyvidy rnemqakmsv avnatptspt anvtqwlnms vlatspmppt
241 pvppaslvvr nvtvyrlyhs dlalhntslq tatfliysvl iklipcialt ilsvrlimal
301 leakrrrkkl tskpaasngt ktlvngksae rprknsktle kekqtdrttr mllavlllfl
361 itefpqgilg llnsvlgnef lmqcylrlsd lmdvlalins sinfilycsm skqfrstftl
421 lfrpkfldkw lpvaqdelaa sradglqrsv vaavpdkyqs thkqpqvvtl aattttnnat
481 evtnl
```

**M MsR1 PB (Genbank Ref # AAF47635.2)** (stop suppression)

```
  1 masgnnetep lycgsgmdnf htsyknmhgy vslvvcilgt iantlniivl trremrsptn
 61 ailtglavad lavmleyipy tihdyiltds lpreeklsys wacfikfhsi faqvlhtisi
121 wltvtlavwr yiavgypqkn rvwcgmrtti itittayvvc vlvvspslyl itaiteyvdq
181 ldmngkvins ipmtqyvidy rnellsarta alnatptsap lnetvwlnas tlltstttaa
241 pptpspvvrn vtvyrlyhsd lalhnaslqn atfliysvvi klipcialti lsvrlilall
301 eakrrrkklt skpatpgasn gtkspangka adrprknskt lekekqtdrt trmllavlll
361 flitefpqgi mgllnavlgd vfylqcylrl sdlmdilali nssinfilyc smskqfrttf
421 tllfrpkfld kwlpvaqdem aaaraersav apvlekgrqq pqvvmasttt nitqvtnlxh
481 rrsrgrrtll srllsvlkrg rrrssgeggg vggggaplag ndavepafqa ivvvvdkvsg
541 atenqlytae qarivt
```

**M Ms R2 PA (Genbank Ref #** AAF47633)

1 mvtnmsqphy cgtgiddfht nykyfhgyfs livcilgtia ntlniivltr remrsptnai
61 ltglavadla vmleyipytv hdyilsvrlp reeqlsyswa cfikfhsvfp qvlhtisiwl
121 tvtlavwryi avsypqrnri wcgmrttlit iatayvvcvl vvspwlylvt aiakfletld
181 angktiasvp lsqyildynr qdevtmqvms sttpdvswai psdsangtav sllslttvip
241 lttlstgvtt ssslgernvt vyklyhsala lrdrqfrnat fliysvlikl ipcfaltils
301 vrligallea krrrkilach aandmqpivn gkvviptqpk sckllekekq tdrttrmlla
361 vlllflvtef pqgimgllnv llgdafflqc ylklsdlmdi lalinssinf ilycsmsrqf
421 rstfallfrp rwldkwlpls qhdgegrvgg sgglggyggy grqrllhtda v***sksmaid***lg
481 lttqvtnv

**V MS R2 PA (Genbank Ref #** XP_015031310.1)

1 mvtnmsqphy cgasvddfht nykyihgyfs livcilgtla ntlniivltr remrsptnai
61 ltglavadla vmleyipytv hdyilsarlp reqqlsyswa cfikfhsifa qvlhtisiwl
121 tvtlavwryi avsypqrnri wcgmrtttit iatayvvcvl vvspwlylvs siakfletld
181 adgktirsvp lsqyildynn eehitmqvls sttpdmawsl apgssnttav nllgpftdlp
241 tttmttttvp ydvgernvtv yklyhselal hdrplrnatf liysvlikli pcfaltvlsv
301 rligalleak krrkilacha andmqpivng kvvtpsqpks ckllekekqt drttrmllav
361 lllflitefp qgimgllnal lgdaflmqcy lklsdlmdil alinssinfi lycsmsrqfr
421 stftllfrpr wldkwlplsq hdgdardgag grldgygrqr lvhtdav***sks*** ***maid***lglttq
481 vtnv

**M MsR2 PB (GenbankRef #** AAN12219) **(stop suppression)**

1 mvtnmsqphy cgtgiddfht nykyfhgyfs livcilgtia ntlniivltr remrsptnai
61 ltglavadla vmleyipytv hdyilsvrlp reeqlsyswa cfikfhsvfp qvlhtisiwl
121 tvtlavwryi avsypqrnri wcgmrttlit iatayvvcvl vvspwlylvt aiakfletld
181 angktiasvp lsqyildynr qdevtmqvms sttpdvswai psdsangtav sllslttvip
241 lttlstgvtt ssslgernvt vyklyhsala lrdrqfrnat fliysvlikl ipcfaltils
301 vrligallea krrrkilach aandmqpivn gkvviptqpk sckllekekq tdrttrmlla
361 vlllflvtef pqgimgllnv llgdafflqc ylklsdlmdi lalinssinf ilycsmsrqf
421 rstfallfrp rwldkwlpls qhdgegrvgg sgglggyggy grqrllhtda vsksmaidlg
481 lttqvtnvxq essgraamsa aaggaaasva lalaatdvdg cppatdaavs tndislvekl
541 hlqpsprgta issgqhrrrr sgsgtkciwp ttdwlrklrn qkareteqss eqdielgkss
601 inrrssvllm vllsssdevk akavlvseqp pspadedved aidalwl

**M NPF R PA** **(Genbank Ref #** AAF51909)

```
  1 miismnqtep aqladgehls gyasssnsvr ylddrhpldy ldlgtvhaln ttaintsdln
 61 etgsrpldpv lidrflsnra vdspwyhmli smygvlivfg algntlvvia virkpimrta
121 rnlfilnlai sdlllclvtm pltlmeilsk ywpygscsil cktiamlqal cifvstisit
181 aiafdryqvi vyptrdslqf vgavtilagi walalllasp lfvykelint dtpallqqig
241 lqdtipycie dwpsrngrfy ysifslcvqy lvpilivsva yfgiynklks ritvvavqas
301 saqrkvergr rmkrtnclli siaiifgvsw lplnffnlya dmerspvtqs mlvryaichm
361 igmssacsnp llygwlndnf rkefqellcr csdtnvalng httgcnvqaa arrrrklgae
421 lskgelkllg pggaqsgtag gegglaatdf mtghhegglr saitesvalt dhnpvpsevt
481 klmpr
```

Dm NPFR isoforms do not differ by BBS, only by alternative CT sequences and a small exon

**V NPF R** **(Genbank Ref #** XP_002058565.3)

  1 miiamnrtef gtlpqfessa eifkaiarns nfdligerrh lanfpslhtg dvsssssssi
 61 snsisnninn ssnsnsssnn ishsnmtill lnstnnesnf mpadmdpvlm dqylhnraie
121 spwyhlliam ysvlivfgam gnimvviavv rkpimrtarn lfilnlaisd lllclvtmpl
181 tlmeilskfw pygscavlck tiatlqalsi fvstisitai afdryqvivy ptrdslqfvg
241 availagiwi laltlasplf iykqlismdm ppvlqrlgvp hrisyciedw plsdgrfyys
301 ifslcvqylv pilivsvayf giynklk***sri tvvtvqs***ssq rkvergrrmk rtnrllisia
361 iifgvswlpl nffnlyadlq hpsavtqrml vayaichmig mssacsnpll ygwlndnfrk
421 efqellcrst estnvalhgh ttgsnmqaaa arrrrkhelq llgnsvqrga sdgdclggms
481 iaatefntrh tvng***trsavt*** esvaltespm psemttlvpr

**M Pk1 R PD (Genbank Ref #** AAX52950)

1 msagnmshdl gpprdplaiv ipvtvvysli fitgvvgnis tcivikknrs mhtatnyylf

61 slaisdflll lsgvpqevsy iwskypyvfg eyicigrgll aetsanatvl titaftvery

121 iaichpflgq amsklsrair iivlvwimai vtaipqaaqf giehysgveq cgivrvivkh

181 sfqlstfiff lapmsiilvl ylligvhlyr stlvegpasv arrqqlksvp sdtilyrygg

241 ***sgtams***fngg gsgagtaglm ggsgaqlssv rgrlnhygtr rvlrmlvavv vcfflcwapf

301 haqrliaiya pargaklrdq hefvytvmty vsgvlyylst cinpllynim shkfreafka

361 vlfgkkv***skg slns***rnnies rrlrral***tns sqt***qrfsies aeqpkpsimq nptnkppvaa

421 qyamigvqvn

**V Pk1 R (Genbank Ref #** XP_002054764.2)

1 mspastppai nqsaatitql lgaqrdplai vipvtvvycl ifltgvvgni stcivikknr

61 smhtatnyyl fslaisdfml llsgvpqemy fiwskypyvf geyfcigrgl laetsanatv

121 ltitaftver ymaichpflg qamsklsrai riivliwlla vltaipqaaq fginsyagvd

181 kcvvvrvivq hsfqlstfif ffapmsiilv lylciglhly rssvigggsp tapvtessar

241 rqqplkavas dtilyryags ssvlannagp qlssvrgrlt hygtrrvlrm lvavvicffl

301 cwapfhaqrl iaiyaparga qlhdqhelly tvmtyvsgvl yylstcinpl lynlmsnkfr

361 eafkavllgk kyskgmqnsr hqlesrrlrr ***tttlnsstqr dsieskht***lm qtalnerllq

421 tksqlvl

**M Pk2 R1 PA (Genbank Ref #** AAF54930)

```
  1 mlqqgvaitia ndsnddginq sfmahvspsp nqspsigvgi giasstmanp sespemlllk
 61 ndkflthvah llnittenls nllgstngtn astmaadspv desltlrtal tvcyalifva
121 gvlgnlitci visrnnfmht atnfylfnla vsdlillvsg ipqelynlwy pdmypftdam
181 cimgsvlsem aanatvltit aftveryiai chpfrqhtms klsraikfif aiwlaaflla
241 lpqamqfsvv yqnegysctm endfyahvfa vsgfiffggp mtaicvlyvl igvklkrsrl
301 lqslprrtfd anrglnaqgr virmlvavav afflcwapfh aqrlmavygl nlinigisrd
361 afndyfrild ytsgvlyfls tcinpllyni mshkfreafk itltrqfgla rnhhhqqsqh
421 hqhnysallr qngsmrlqpa scsvnnnale pygsyrvvqf rcrdanhqls lqdsirtttt
481 tttinsnsma agngvgggag gggggrrlrk qelygpgpgt avphrmlqaq vsqlsslgda
541 nslleaevvd rhyasgrakr allatksgal lvtppqsgdp sevsqpatrl kltrvisrrd
601 evantstppf cgshslpdpe tcqsasvagr ssrkfpwrkr rqktedpsse gltygspksq
```

**V PK2 R1 PA-like (Genbank Ref #** XP_002056087.1)

```
  1 mmqgvdfala sdnddglnqs fmahmlpnar pslrpstsrs psatsspell llhndkflth
 61 vaqmlnmtte nltnllaans tngssatati tattsttnma nsteesptll iilticyali
121 fvagvlgnli tcivisrnnf mhtatnfylf nlavsdlill vsgipqelyn lwcpnsypft
181 dgicivesvl semaanatvl titaftvery iaichpfrqh tmsklsraik fifaiwlaaf
241 llalpqamqf svvnqdngys ctmennfyah vfavsgfiff cgpmtaicvl yvligmklkr
301 srllqslprr aydanrglna qsrvirmlva vavafflcwa pfhaqrlmav ygvslinacr
361 crdafndyfh ildytsgvly flstcinpll ynimshkfre afkitltrqf glarnhqqqs
421 qyrqhnysal mrlqgsmrlq pvscsnnnna lepygsyrvv qfrcrdanhq lslqdsirtn
481 tttttinsss lagagaasvp gsgggcggsg vgggnggasg rrlrkqefya tapgsavphr
541 llqsqvsrls dgnshslldt nvadvgrhca agrakrtlla tnngalllat aqpddaqppt
601 rlklsrvisr rdepaysgss slpeqetnns tnttdpngge lrkfpwrkkr qkqnvannne
661 tgggrngyat pksl
```

**M Pk2 R2 PA (Genbank Ref #** AAF54929.2)

1 mavkmlptns sgvlatdlql fhnekfllnl tqvlnisadn ltsllqglep eellptvtpm
61 tplsllatls vgyalifiag vlgnlitciv isrnnfmhta tnfylfnlai sdmillcsgm
121 pqdlynlwhp dnypfsdsic ilesvlseta anatvltita ftveryiaic hpfrqh***tmsk***
181 ***ls***ravkfifa iwiaalllal pqaiqfsvvm qgmgtsctmk ndffahvfav sgflffggpm
241 taicvlyvli gvklkrsrll qalprrcydv nrgisaqtrv irmlvavava fficwapfha
301 qrlmavygst sgiesqwfnd vfsildytsg vlyflstcin pllynimshk freafkvtla
361 rhfglggknq grglphtysa lrrnqtgslr lh***ttdsvrtt mtsmatttt***g lngsangsgn
421 g***tttgqs***vrl nrvsldsvqm qgqnrsrqdl fdnprrmlq***t qisqlss***vgd ahslleedlq
481 fpgeplqrqp tmcsideltd dlaisrsrlk ltritrppgg vtggvaggst tgaagsggvs
541 gdessgkvrk akvkvlksss pfkglrtkfn wrarrkgshk phekgatvng gdteeraaf

**V PK2 R2 PA-like (Genbank Ref #** XP_032289636.1)

1 msammlatnv salgegtppl selqmfhsek fllnltqvln isadnltsll qelephdlle
61 qsprpmathm gllatlsvgy alifvagvlg nlitcivisr nnfmhtatnf ylfnlaisdl
121 illcsgmpqd lynlwhpnny pfsdgicile svlsetaana tvltitsftv eryiaichpf
181 rqh***tmskls***r aikfifaiwi aalllalpqa iqfsvvtqga gssctmnndf fahvfavsgf
241 lffggpmtai cvlyvligik lkrsrllqal prrsydvnrg isaqtrvirm lvavavaffi
301 cwapfhaqrl mavygstfgi esqwfndvfn ilnytsgvly flstcinpll ynimshkfre
361 afkvtlarqf rlsgkhqgqg lphnysalrr nqtgslrlh***t dsvrttttlt t***lngssngaa
421 vggcrssqrl qrgsl***sssha t***llstq***srsr qd***lfaarvva adaaattaag aaaaatagaa
481 ahsqphr***tlq tqisqlss***vg dahslleael qlpeehyela arhkcphaam gaidelscqs
541 svvggaaaag lsrarlkltr itrhpqmqts gaalasepaa ttataaataa sasasavaks
601 tgsgkvrkak akrsntlkgl raklnwrgrh kesandsadg qqmphssvas tpsgvy

**M Proc R PC (Genbank Ref #** AAX52477.1)

```
  1 mtmsststat atstatatld eanatvgemf sdadmaevrh vvq rilvpcv fvigllgnsv
 61 siyvltr krm rcttniylta laitdiaylt cqlilslqhy dypkyhfkly wqly gyfvwl
121 cdsfgyisiy iavcfti erf iairyplkrq tfcteslakk viaavaifcl lstlst afeh
181 titigtrqid dayqpcnqtv anispmpppp vavtpplatp plptpatiwq spdsamestt
241 sgssnqlvdw gsgsgdgepe niprhrrhwq ssgfvtlptl rktleeqdqk vadaaqrsgv
301 tesllqlwrr krsaenhnin ntdafafnvt eycqnvtyyn hglselgyde ly sylwnlft
361 llvfvvfpll llatfn sili llvhrsknlr gdltnassir rtkrksnsgl kgsvsqenrv
421 titliavvlm fivcqlpwai ylivnqymei qigtq vvagn vcnllaslha asnfflycvl
481 sdkyrktvre litgyryrrr harnntslyv phttttltqi ngdhygsnyg gagsrrnrnt
541 grlia
```

(residues 513–518 "ttttlt" in bold italic)

**V Proc R PC-like (Genbank Ref #** XP_015026487.1)

```
  1 mttlaatsts tampvdgigt akmfsdadva evrhvvqril vpcvfvigll gnsvsiyvlt
 61 rkrmrcttni ylsalaitdi ayltfvlils lkhyeyikyh celywrlygf imwlcdacay
121 isiyiavcft ierfiairyp lkrqtfctes lakkviaava lfcllttlst afehtydinw
181 klidgayrpc nltlanvspt paqptlataw hvdehannaa tprylepfdl ssgsgsgage
241 sdhipsrqhr lpastsasvg vtaaatepat aaatasllql srarrsedny dseassnnny
301 nynnnsafaf niteycqnmt vytnglsslg qnalyfniws vytlivfvvl plfvlatfnc
361 flillvhrsk slrgdltnas sirrtkrksn sgitgsvsqe nrvtitliav vllfivcqlp
421 waiylilvqy veiemniqri agnvcnllva inaaanffly cvlsdkyrkt vrelitgyry
481 rhrharnnis lyaphttttl ngdgagggga sgygssysga ssrrcraksa varrlia
```

(residues 389–394 "sn sgit" and 515–522 "ssysga ss" in bold italic)

**M rkts PA (Genbank Ref #** AAF53367**)**

```
   1 maarcrwswr lalcplllql llqllllpps amghdetken papdmqnsqe qepyvhlqhl
  61 qqqqqqnpqt vqqlsqitvn rtsksasvtp tgirenvmlp sadpekeaqi lyekslqeyh
 121 gsqlstasta tdviagkrtl hsicerwlqk hchctgslev lrlscrgigi lavpvnlpne
 181 vvvldlgnnn ltkleansff mapnleeltl sdnsiinmdp nafyglaklk rlslqncglk
 241 slppqsfqgl aqltslqlng nalvsldgdc lghlqklrtl rlegnlfyri ptnalaglrt
 301 lealnlgsnl ltiindedfp rmpnlivlll krnqimkisa galknltalk vlelddnlis
 361 slpeglskls qlqelsitsn rlrwindtel prsmqmldmr anplstispg afrgmsklrk
 421 lilsdvrtlr sfpeleacha leilkldrag iqevpanlcr qtprlkslel ktnslkripn
 481 lsscrdlrll dlssnqieki qgkpfnglkq lndlllsynr ikalpqdafq gipklqlldl
 541 egneisyihk eafsgftale dlnlgnnifp elpesglral lhlktfnnpk lrefpppdtf
 601 priqtlilsy ayhccaflpl vamssqkkts qvqeavlfps daefdmtlwn nsmmniwpqm
 661 hnlskqlgas mhdpwetain fneeqlqtqt ggqiatsyme eyfeehdvsg patgygfgtg
 721 lfsgmstedf qpgsvqclpm pgpflpcadl fdwwtlrcgv wvvfllsllg ngtvvfvllc
 781 srskmdvprf lvcnlaaadf fmgiylgila ivdaatlgef rmfaipwqms vlcqlsgfla
 841 vlsselsvyt lavitlerny aithaihlnk rlslkqagyi msvgwvfali malmplvgvs
 901 dyrkfavclp fetttgpasl tyvislmfin gcafltlmgc ylkmywairg sqawntndsr
 961 iakrmallvf tdflcwspia ffsitaifgl qlisleqaki ftvfvlplns ccnpflyaim
1021 tkqfkkdcvt lckhfeesrv vggggpggrg avartkrgdl ppllpaaav ahppgcrclr
1081 mlpsempnwh kmeqtpsmwq rlrtfccgen rrrrkqrrqp qqrrqrayta aaanpyqyqf
1141 aelrqqrqnr assissenfc ssrssswrhg ppssapvppg ncsmplkmle phahphghgr
1201 rrhsawlitr ktsqdsnlss srndssasat taststfrls rssagsstpl psiiahngka
1261 qldavkprlv rqeavqeeed sspprlgvrf lptipsaads svvmedgdsa ntgvasflgm
1321 plpgassgfl iapttaatts pppvvlqpak pppdpndapl
```

**V Rk PA-like (Genbank Ref #** XP_002057499.2**)**

```
   1 maahsrrrrk qkrsdagrav trvaaatsit afglqlsmpp lllltlllmr tttalltpts
  61 mhlnetkenr gtakqlsgis vnnmsstpta tptttptatp vgirenvmlp ssdpereaqi
 121 lyekslqeyh gkaaaaaaaa aasassnvds ngvsgsgsgs atkttarslh svcelwllkh
 181 chctgslenl klscrsigil avpvnlpsev lfldlgnnnl trleansffm vphleeltls
 241 dnsiinmdpy afyglaklkr lslqncglka laphsfqgls qlvslqlngn alvsldgncl
 301 gnlqqlrtlr legnlfyrip tnalaglktl ealnlgsnll tiindedfpr mpnlivlllk
 361 rnqimkisag alknltalkv lelddnliss lpeglgklpq lqelsmtsnr lrwindtelp
 421 rsmqildlra nplstittga frgmsklrkl ilsdvrtlrn fpeleachal eilkldragi
 481 qevpsnlcrq tprlkslelk tnslksipnl sscrdlrlld lssnqietlq grpfhglkql
 541 hdlllsynri ktlpqdafqg ipklqlldle gneivhihkd afaaftaled lnlgnnifph
 601 lpeaglrall hlktfnnpkl refpppdtfp riqtlilsya yhccaflplv amsaqrktsq
 661 vqeavlfpsd aefdmtlwnn smmniwpqmh nlskqlgaam hdlwdsplnn ynlnedlqqs
 721 ptgsqsassy meeyfdehdv sgpgtgygfg tglfsgitad dlqpgsvqcl pmpgpflpca
 781 dlfdwwtlrc gvwvvfllal lgngtvvfvl vcsrskmdvp rflvcnlaaa dffmgiylgi
```

841 laivdaatlg efrmfaipwq msllcqlagf lavlsselsv ytlavitler nyaithaihl

901 nkrlslrqag yimsvgwifa lcmallpllg vsdyrkfavc lpfetttgva sltyvislmf

961 ingcafltlm gcylkmywai rgsqawntnd sriakrmall vftdflcwsp iaffsitaif

1021 glqlisleqa kiftvfvlpl nsccnpflya imtkqfkkdc vtlckhfeet rvvgggataa

1081 araargkrgg elppplpa***t ataat***aagva hppgcrclrm lpsempnwhk meqttptlwq

1141 rlkrlccgqr qrrkqrrqpq qrrqraytaa aanpyqyqfa elrqqrknra ssissenfcs

1201 srssswrhga nsstpapqsn csmplklmep gqshthgrrr hsawlitrkt sqdsnlsssr

1261 nd***ssasatta stst***frl***srs sags***stplps iiahngkqqh etltkprlvr qeavqeeeds

1321 spprlgvrfl p***tipsaad***ss vimddgdsan aaagsagclg mpqpgvssgf lvtppaqlqp

1381 ekpppapnda pg

**M RYa R PA (Genbank Ref #** AAF56655.3)

```
  1 mehhnshllp ggsekmyyia hqqpmlrned dnyqegyfir pdpasliynt talpaddegs
 61 nygygstttl sglqfetyni tvmmnfscdd ydllsedmws sayfkiivym lyipififal
121 igngtvcyiv ystprmrtvt nyfiaslaig dilmsffcvp ssfislfiln ywpfglalch
181 fvnysqavsv lvsaytlvai sidryiaimw plkpritkry atfiiagvwf ialatalpip
241 ivsgldipms pwhtkcekyi cremwpsrtq eyyytlslfa lqfvvplgvl iftyaritir
301 vwakrppgea etnrdqrmar skrkmvkmml tvvivftccw lpfnilqlll ndeefahwdp
361 lpyvwfafhw lamshccynp iiycymnarf rsgfvqlmhr mpglrrwccl rsvgdrmnat
421 sgtgpalpln rmntsttyis arrkpratsl ranplscget splr
```

(In line 421, "tsttyis" is printed in bold italics.)

**V RYa R PA (Genbank Ref #** XP_002053912.2)

```
  1 mefnsfvwqr rrldlqrtvl nderlyyiap qqpllrnedd yqdtagalmy nssstelslg
 61 aedadyisst patehfnvtv llnfsctdvs shsddlwssd yfksvvylly ipififallg
121 ngivcyivqs tprmrtvtny fianlalgdi lmslfcvpss fisqyilnyw pfgivlchfv
181 nysqvvsvlv saytlvaisi dryiaimwpl rpritkryak fiiagvwfia latafpipvv
241 srlmvpssiw hekcekyicr evwpsteqdy yytlalftlq fivpllvlif tytriaiavw
301 gkrppgeaen srdqrmarsk rkmikmmltv vivftscwlp fnilqlmlnd eefanwkplp
361 yvwfafhwla mshscynpii ycymnarfrg gflqimyrvp glrrccclhr ylrsrgersy
421 eatg//tedafh lhrvntctty istrrklrtn smqm//sqfsca ettvlr
```

(In line 421, "tctty is", "sqfsca" and "et" are printed in bold italics.)

**M RYa R PB (Genbank Ref #** AAF56655.3) (difference from PA underlined)

```
  1 mehhnshllp ggsekmyyia hqqpmlrned dnyqegyfir pdpasliynt talpaddegs
 61 nygygstttl sglqfetyni tvmmnfscdd ydllsedmws sayfkiivym lyipififal
121 igngtvcyiv ystprmrtvt nyfiaslaig dilmsffcvp ssfislfiln ywpfglalch
181 fvnysqavsv lvsaytlvai sidryiaimw plkpritkry atfiiagvwf ialatalpip
241 ivsgldipms pwhtkcekyi cremwpsrtq eyyytlslfa lqfvvplgvl iftyaritir
301 vwakrppgea etnrdqrmar skrkmvkmml tvvivftccw lpfnilqlll ndeefahwdp
361 lpyvwfafhw lamshccynp iiycymnarf rsgfvqlmhr mpglrrwccl rsvgdrmnat
421 sgemttkyhr hvgdalfrkp kicir/ngsst ssqsnehihh lhqrsskats difasepiiv
481 rrdvttavav isknktdspv rrsgssggte anirstef
```

(Underlined as printed: in line 421, everything from "emttkyhr" to the end of the line; line 481 in full. Printed in bold italics: "sst ssqs" in line 421 and "sgssggt" in line 481.)

**V Rya R PB-like (**not annotated; predicted based on inspection of genomic DNA documented in SXX Document – "de novo RYaR annotations")

```
  1 mefnsfvwqr rrldlqrtvl nderlyyiap qqpllrnedd yqdtagalmy nssstelslg
 61 aedadyisst patehfnvtv llnfsctdvs shsddlwssd yfksvvylly ipififallg
121 ngivcyivqs tprmrtvtny fianlalgdi lmslfcvpss fisqyilnyw pfgivlchfv
181 nysqvvsvlv saytlvaisi dryiaimwpl rpritkryak fiiagvwfia latafpipvv
241 srlmvpssiw hekcekyicr evwpsteqdy yytlalftlq fivpllvlif tytriaiavw
301 gkrppgeaen srdqrmarsk rkmikmmltv vivftscwlp fnilqlmlnd eefanwkplp
361 yvwfafhwla mshscynpii ycymnarfrg gflqimyrvp glrrccclhr ylrsrgersy
```

421 eatg//emtkynrrngdglvrkpkirir/nrrcipptscehlhhlhqhktkaahefyane/pifm
crdhsvavapgtaqepkprlqprqptgsfhln***sasyls***stqf

**M RYa R PC (Genbank Ref #** AHN57552)

```
  1 mehhnshllp ggsekmyyia hqqpmlrned dnyqegyfir pdpasliynt talpaddegs
 61 nygygstttl sglqfetyni tvmmnfscdd ydllsedmws sayfkiivym lyipififal
121 igngtvcyiv ystprmrtvt nyfiaslaig dilmsffcvp ssfislfiln ywpfglalch
181 fvnysqavsv lvsaytlvai sidryiaimw plkpritkry atfiiagvwf ialatalpip
241 ivsgldipms pwhtkcekyi cremwpsrtq eyyytlslfa lqfvvplgvl iftyaritir
301 vwakrppgea etnrdqrmar skrkmvkmml tvvivftccw lpfnilqlll ndeefahwdp
361 lpyvwfafhw lamshccynp iiycymnarf rsgfvqlmhr mpglrrwccl rsvgdrmnat
421 sgemttkyhr hvgdalfrkp kicircktlh lvsvsvflfv llrfwfi
```

**M SIF R PA (Genbank Ref #** AAN13859.2)

1 mmaasgrirk rkhkshtsgd vpstttsvpm piptmapgkm vaetmeeaaa lagdynnfth
61 nfvdlqnlls fnelngtsgs ggtavsslgs ssaiklnnsa itdtllgtvl ttatatvapa
121 assllatlaa tttasargsl agkslaiada tsstyysnll nlspattsli saaaatksyn
181 dsalrweqld gsvdfgfdpl yrhslamsmv ycvayivvfl vglignsfvi avvlraprmr
241 tvtnyfivnl aiadilvivf clpatligni fvpwmlgwlm ckfvpyiqgv svaasvysli
301 avsldrfiai wwplkqmtkr rarimiigiw vialvttipw llffdlvpae evfsdalvsa
361 ysqpqflcqe vwppgtdgnl yfllanlvac yllpmslitl cyvliwikvs trsipgeskd
421 aqmdrmqqks kvkvikmlva vvilfvlswl plyvifarik fgsdisqeef eilkkvmpva
481 qwlgssnsci npilysvnkk yrrgfaaiik srsccgrlry ydnvaia***sst tstrks***shyh
541 qnssrkspss kgnavsyiye hnslrrhnmm lkqdsnlsqq mllkqdshgs rqflikqe***ss***
601 ***csdas***girrp lcqqd***sngsk vslskqdsiv syme***arrsag hglndtlvdr dsvsmdvgrr
661 qga***tpsslld*** krqkfvkqd***s visfvd***qrpe qrrhqlvkqd ***svisfad***qrr gllhkqdslm
721 anrtgdapth hvsilkk***tds qlsygssts***p rrnadlye

**V SIFa R PA-like (Genbank Ref #** XP_002058467.3)

1 mavggrtrkr khrshaagdt ptttaaaaaa agtaatataa itnssssssgn tpatgglgrg
61 wpmleqrdll edvtqgagea nnfthnfvdl qnllnfndaa ssnnsnnssn nvvgvsfsps
121 aiklsngait dtllgtiltt atatvapaas slissltaat atattttaat atsssssqla
181 aigmpgaviv adatsssyya sllnmspatt slitaaaatk syndsllrwd qldgnvdfgf
241 dplyrhslam sivycvayiv vflvgligns fviavvlrap rmrtvtnyfi vnlaiadilv
301 ivfclpatli gnlfvpwmlg wlmckfvpyi qgvsvaasvy sliavsldrf iaiwwplkqm
361 tkrrarimii giwvialvtt ipwllffdlv paeevfsdal vssytqpqyl cqevwppgtd
421 gnlyfllanl vacyllpmsl itlcyvliwi kvstrsipge lskdaqmdrm qqkskvkvik
481 mlvavvilfv lswlplyvif arikfgsdis qeefeilkkv mplaqwlgss nscinpilys
541 vnkkyrrgfa aiiksrsccg rlryydnvai a***ssttstrks*** shyhpngsrk spsspglrkt
601 navsyiyehn slrrhnlmmk qdsnlsqqml lkqdshgsrq flikqe***sscs das***gtrrllc
661 qqd***sngskvs lskqdsivsy me***srrvaala aqer***svdsal t***qqd***tisies*** rrggaqa***tpa***
721 ***slld***krqkfv kqd***svisfvd*** qrpeprrhql vkqd***svisfa d***qrrgllhkq d***slmtnrs***gd
781 apthhvsilk ktdsqlsyg***t sssssas***pr rnvelye

**M sNPF R PA (Genbank Ref #** AAF49074)

1 manlswlsti tttsssists qlplvsttnw sltspgttsa iladvaasde drsggiihnq
61 fvqiffyvly atvfvlgvfg nvlvcyvvlr nramqtvtni fitnlalsdi llcvlavpft
121 plytfmgrwa fgrslchlvs faqgcsiyis tltltsiaid ryfviiypfh prmklstcig
181 iivsiwvial latvpygmym kmtnelvngt qtgnetlvea tlmlngsfva qgsgfieapd
241 stsatqaymq vmtagstgpe mpyvrvycee nwpseqyrkv fgaitttlqf vlpffiisic
301 yvwisvklnq rarakpg***sks srre***eadrdr kkrtnrmlia mvavfglswl pinvvnifdd
361 fddksnewrf yilfffvahs iamsstcynp flyawlnenf r//kefkhvlpc fnpsnnniin
421 itrgynrsdr ntcgprlhhg kgdggmgggs ldaddqdeng itqetclpke klliiprept
481 ygngtgavsp ilsgrginaa lvhggdhqmh qlqpshhqqv eltrrirrrt detdgdylds
541 gdeqtvevrf setpfvstdn t***tgisile***ts tshcqdsdvm velgeaigag ggaelgrrin

**V sNPF R PA-like (Genbank Ref #**XP_002048039.1)

1 manvsnetvn awlvsvstql phvlgidvgv nwssssttst ttmaipsssm gttttastvs
61 slssdantda aadadksgii hnqfvqiffy vlyttvfvlg vfgnvlvcyv vlrnramqtv
121 tnifitnlal sdillcvlav pftplytfmg rwafgrtlch lvsfaqgcsi yistltltsi
181 aidryfviiy pfhprmklst cigiivsiwv iallatvpyg mymkmtnevm dnttqlvgan
241 qtsssrygny glatpdatsa aqaymqvmtd gvtvceenwp sehyrkvfga itttlqfvlp
301 ffiisicyvw isvklnqrar akpg***skssrr e***eadrdrkkr tnrmliamva vfglswlpin
361 lvnifddfdd ksnewrlyml fffvahsiam sstcynpfly awlnenfrke fkhvlpcfnp
421 snnniinitr gynrsdrntc gprlhhgnge ggaggsldad dqddngitqe tclpkeklli
481 ipreptygng ngavspilsg rginaallha sqpqqqqqqv qqqqveltrr irrrtdddtd
541 fidsgdeqtv evrfsetpfv ssdnt***tgism le***tsesqfqd sgelpelsav vvgdasrrcn

**M SP R PA (Genbank Ref #** AAF46037**)**

1 mdnytdvlyq yrlapsaspe memeladprq mvrgfhlptn esqleipdyg nesldypnyq

61 qmvggpcrme dnnisywnlt cdspleyamp lygycmpfll iitiisnsli vlvlskksma

121 tptnfvlmgm aicdmltvif papglwymyt fgnhykplhp vsmclaysif neimpamcht

181 isvwltlala vqryiyvcha pmartwctmp rvrrctayia llaflhqlpr ffdrtymplv

241 iewngsptev chletsmwvh dyigvdlyyt

271 syylfrvlfv hllpciilvt lnillfaamr qaqerrkllf renrkkeckk lretncttlm

331 livvvsvfll aeipiavvta mhivssliie fldyglanic imltnfflvf sypinfgiyc

391 gmsrqfretf keiflgrlma kkd***sstkys***i vngar***tctnt net***vl

**V SP R PA-01ike (Genbank Ref #**XP_002055695.1)

1 marsvdqsll ieleevstlt aaatavakss smdnntvysy disitdvlyq wsaaasgara

61 iaqsaasgpl ptasvaavae rlpelvgsml etrasqasln esqfllvdya nsmndsldya

121 syqqqlgsse crqmdgnmsy wnltcdspld yalplygycm pfllfmtiis nslivlvlsk

181 ksmatptnfv lmgmaicdll tvvfpapglw ymytfgnhyk pmhpvsmcla ysifneimpa

241 mchtisvwlt lalavqryiy vchapmartw ctmprvrrct fyiallaflh qlprffdrty

301 mpmeiewngn etevchlets lwvheyvgvd lyyt

335 syylfrvlfv nllpciilvt lnillfaalr qaqerrkllf renrkkeckr lrdsncttlm

395 livvvsvfli aeipiavvta mhivssliie fldygianif imltnfflvl sypinfgiyc

391 gmsrqfretf reifmgrvag kke***ssskys***i vngprtc***tnt net***il

**M SPR PD (Genbank Ref #** AAF46037**) (stop suppression)**

1 mdnytdvlyq yrlapsaspe memeladprq mvrgfhlptn esqleipdyg nesldypnyq

61 qmvggpcrme dnnisywnlt cdspleyamp lygycmpfll iitiisnsli vlvlskksma

121 tptnfvlmgm aicdmltvif papglwymyt fgnhykplhp vsmclaysif neimpamcht

181 isvwltlala vqryiyvcha pmartwctmp rvrrctayia llaflhqlpr ffdrtymplv

241 iewngsptev chletsmwvh dyigvdlyyt

271 syylfrvlfv hllpciilvt lnillfaamr qaqerrkllf renrkkeckk lretncttlm

331 livvvsvfll aeipiavvta mhivssliie fldyglanic imltnfflvf sypinfgiyc

391 gmsrqfretf keiflgrlma kkd***sstkys***i vngar***tctnt net***vl**xx**lvm lvprrgssdh

451 rrssststtt tttktiggsm iiggeasaqh qhlvthhlqt hsqpsqqrrv stmdiiteer

511 il

**M Tk R 86C PA (Genbank Ref #** AAF46037**)**

```
  1 mseivdtell vnctilavrr felnsivntt llgslnrtev vsllssiidn rdnlesinea
 61 kdflteclfp sptrpyelpw eqktiwaiif glmmfvaiag ngivlwivtg hrsmrtvtny
121 fllnlsiadl lmsslncvfn fifmlnsdwp fgsiyctinn fvanvtvsts vftlvaisfd
181 ryiaivhplk rrtsrrkvri ilvliwalsc vlsapcllys simtkhyyng ksrtvcfmmw
241 pdgryptsma dyaynliilv ltygipmivm licyslmgrv lwgsrsigen tdrqmesmks
301 krkvvrmfia ivsifaicwl pyhlffiyay hnnqvastky vqhmylgfyw lamsnamvnp
361 liyywmnkrf rmyfqriicc ccvgltrhrf dspksrltnk nssnrhtrae tksqwkrstm
421 etqiqqapvt sscreqrsaq qqqppgsgtn raavecimer padgsssplc lsinnsiger
481 qrvkikyisc dednnpvels pkqm
```

**V Tk 86C R PA- (Genbank Ref #**XP_002053610.1**)**

(virilis lost the intron/exon encoding the additional CT sequence

```
  1 mseivdtell vnctilavrr felntivntt llntlnrtev vgllsgiien rdnldsinea
 61 kdflteclfp sptrpyelpw eqktiwaivf glmmfvaiag ngivlwivtg hrsmrtvtny
121 fllnlsiadl lmstlncvfn fifmvnsdwp fgsiyctinn fvanvtvsts vftlvaisfd
181 ryiaivhplk rrtsrrkvrf ilvliwalsc vlsapcllys simtkhyyng ksrtvcfmmw
241 pdgryptsmt dyvynvtilv ltygipmivm licyslmgrv lwgsrsigen tdrqmesmks
301 krkvvrmfia ivsifaicwl pyhlffiyay hnnqvastky vqhmylgfyw lamsnamvnp
361 iiyywmnkr/f rmyfqriifc cclglmryrf espksrmank nssnrhtrae tksqwkrstm
421 etqiqqmpkt ssrdkdagvq glnhtaveci ierpiddnss piclsiknsa gerqrvkiky
481 iscdednnpi eegsennssh dsnhshghgh scgrshgqna kaiqql
```

(155) virilis adds 25 bp at end (new exon or read through?) which includes a BBS

**M Tk R 86C PB (Genbank Ref # ABW08638)**

**(underline is the difference with D melanogaster PA, due to an unspliced intron)**

1 mseivdtell vnctilavrr felnsivntt llgslnrtev vsllssiidn rdnlesinea

61 kdflteclfp sptrpyelpw eqktiwaiif glmmfvaiag ngivlwivtg hrsmrtvtny

121 fllnlsiadl lmsslncvfn fifmlnsdwp fgsiyctinn fvanvtvsts vftlvaisfd

181 ryiaivhplk rrtsrrkvri ilvliwalsc vlsapcllys simtkhyyng ksrtvcfmmw

241 pdgryptsma dyaynliilv ltygipmivm licyslmgrv lwg***srsige***n tdrqmesmks

301 krkvvrmfia ivsifaicwl pyhlffiyay hnnqvastky vqhmylgfyw lamsnamvnp

361 liyywmnkrf rmyfqriicc ccvgltrhrf d***spksrlt***nk nssnrhtrgg ytvahslpns

421 ***spptqt***lla vlaqtltqpk pqtqlllshh sphptqpsaa etksqwkrst metqiqqapv

481 ***tsscre***qrsa qqqqppgsgt nraavecime rpadgssspl clsinnsige rqrvkikyis

541 cdednnpvel spkqm

**V Tk R 86C PB**

1 mseivdtell vnctilavrr felntivntt llntlnrtev vgllsgiien rdnldsinea

61 kdflteclfp sptrpyelpw eqktiwaivf glmmfvaiag ngivlwivtg hrsmrtvtny

121 fllnlsiadl lmstlncvfn fifmvnsdwp fgsiyctinn fvanvtvsts vftlvaisfd

181 ryiaivhplk rrtsrrkvrf ilvliwalsc vlsapcllys simtkhyyng ksrtvcfmmw

241 pdgryptsmt dyvynvtilv ltygipmivm licyslmgrv lwg***srsige***n tdrqmesmks

301 krkvvrmfia ivsifaicwl pyhlffiyay hnnqvastky vqhmylgfyw lamsnamvnp

361 iiyywmnk/at ecrrfntlv vmghgnsdii i***sstess***fmvg ysgs/icclg lmryrfespk

421 srmanknssnr htraetksq wkrstmetqi qqmpktssrd kdagvqglnh taveciierp

481 iddnsspicls iknsagerq rvkikyiscd ednnpieegs enns***shdsnh*** ***s***hghghscgr

541 shgqna kaiqql

ATECRRFNTLVVMGHGNSDIII

***SSTESS***FMVGYSGSI

**M Tk R 99D PA (Genbank Ref #** ABW08638)

```
  1 menrsdfead dygdiswsnw snwstpagvl fsamssvlsa snhtplpdfg qelalstssf
 61 nhsqtlstdl pavgdvedaa edaaasmetg sfafvvpwwr qvlwsilfgg mvivatggnl
121 ivvwivmttk rmrtvtnyfi vnlsiadamv sslnvtfnyy ymldsdwpfg efycklsqfi
181 amlsicasvf tlmaisidry vaiirplqpr mskrcnlaia aviwlastli scpmmiiyrt
241 eevpvrglsn rtvcypewpd gptnhstmes lyniliiilt yflpivsmtv tysrvgielw
301 gsktigectp rqvenvrskr rvvkmmivvv lifaicwlpf hsyfiitscy paiteapfiq
361 elylaiywla msnsmynpii ycwmnsrfry gfkmvfrwcl fvrvgtepfs rrenltsrys
421 csgspdhnri krndtqksil ytcpsspksh rishsgtgrs atlrnslpae slssggsggg
481 ghrkrlsyqq emqqrwsgpn satavtnsss tanttqlls
```

**V Tk R 99D PA-like (Genbank Ref # XP_032294790)**

```
  1 mldsittttt gagqvmenne elldntwsnw styapvliyn amnsvlanqt plsseygpsl
 61 aynlsrvlpt gitglpaadv dedaatpvas fafvvpwwrq vlwsilfggm vivatggnli
121 vvwivlttkr mrtvtnyfiv nlsiadamvs slnvtfnyyy mldsdwifge fyckvsqfia
181 mlsicasvft lmaisidryv aimkplqprm skrrnlaiaa liwlsstlis cpmllffrte
241 evpvtmenkt rivcfpewpd gqtnhskqeh iyniliilt yflpiismtv tysrvgielw
301 gsktigeytp rqtenvrskr rvvkmmivvv lifgfcwlpf htyfivtscy paiteapfiq
361 elylviywla msnsmynpii ycwmnsrfry gfkmvfrwcp fvnvgaesln rrenltsrys
421 csgspdhnri krndtqksil yacpsspksq rvshcgkdds lsgkdlslsh afsysysvag
481 vpprslpvqq lsssrrgqrs syqqemqerw sgdkssnsln sttecrttql ls
```

***tnhstme*** – present in ECL

**M Tk R 99D PC (Genbank ref #ABW08638)**

```
  1 menrsdfead dygdiswsnw snwstpagvl fsamssvlsa snhtplpdfg qelalstssf
 61 nhsqtlstdl pavgdvedaa edaaasmetg sfafvvpwwr qvlwsilfgg mvivatggnl
121 ivvwivmttk rmrtvtnyfi vnlsiadamv sslnvtfnyy ymldsdwpfg efycklsqfi
181 amlsicasvf tlmaisidry vaiirplqpr mskrcnlaia aviwlastli scpmmiiyrt
241 eevpvrglsn rtvcypewpd gptnhstmes lyniliiilt yflpivsmtv tysrvgielw
301 gsktigectp rqvenvrskr rvvkmmivvv lifaicwlpf hsyfiitscy paiteapfiq
361 elylaiywla msnsmynpii ycwmnsrfry gfkmvfrwcl fvrvgtepfs rrenltsrys
421 csgspdhnri krndtqksil ytcpsspksh rishsgtgrs atlrnslpae slssggsggg
481 ghrkrlsyqq emqqrwsgpn satavtnsss tanttqllsx qpaiqvvppe mqtqivcssp
541 ynnnyrranp glsdrdssdd ktwl
```

**M Tre1 PA (Genbank Ref #** NP_524792.1 )

```
        1 mdqdmgmatg yfqdadmqmd epaaatqsiy phsatlfaai sacvfvtigv lgnlitllal
       61 lksptireha ttafvislsi sdllfcsfsl pltavrffqe swtfgttlck ifpvifygnv
      121 avsllsmvgi tlnryiliac hsrysqiykp kfitlqllfv wavsfllllp pilgiwgemg
      181 ldeatfscti lkkegrsikk tlfvigfllp clviivsysc iyitvlhqkk kirnhdnfqi
      241 aaakgssssg ggsymtttct rkarednrlt vmmvtiflcf lvcflplmla nvvddernts
      301 ypwlhiiasv mawassvinp iiyaasnrny rvayykifal lkfwgeplsp mpsrnyhqsk
      361 nskelsgvir stplfhavqk nsinqmcqty sv
```

**V Tre1 PA-like (Genbank Ref #** XP_002056799.1)

```
        1 mdaptqsiyp hsatlfaaic acvfvtigvf gnlitllall ksptirehat tafvislsis
       61 dlffcsfslp ltavrffqes wtfgstlcki fpvifygnva vsllsmvgit lnryiliach
      121 srysqiykpk litlqlvfvw avsfllllpp ilgiwgemgl deatfsctil kkegksikkt
      181 lfligfllpc lviiisysci yitvlhqkkk irshdnfqig aaagaktgsa sgsyvtttst
      241 rkarednrlt vmmvtiflcf licflplmla nvvdderktn ypwlhiiasv mawassvinp
      301 iiyaasnrny rvayykifal lkfwgeplsp mpsrnyhqsk nskelsgvir stplfhavqk
      361 nsinqmcqty sv
```

both present in ICL1 and 3

adjusted TM3 prediction to end just before SRYSQIYKP

**M Trissin R PB (Genbank Ref #** AAF52294)

```
  1 mimtmmqtvr awqqesdveh rkqhkqrwrp dgahisaayd lnsdnddghh rvvhnqnngs
 61 pnsspnqsts afrqrqphhp ptgqqpprlp ctvthfsahw ktllillltll sastltasan
121 vtstisppin gsstdyilly gesttslvpa lttglsgdgs gaviedeeda ekaseyifdr
181 tdvriifitl ytlvfcccff gnllvilvvt lsrrlrsitn fflanlafad fcvglfcvmq
241 nlsiyliesw vfgeflcrmy qfvhslsyta sifilvvicm eryfaivhpi tckqiltaar
301 lrmvivtvwi tsavystpkf vfsktiknih tqdgqeeeic vldremfnsk lldminfvll
361 yvmpllvmtv lyskiaialw rssrgltphv vqhqhqqpqq pscqdigmgm hnsmyhhhph
421 hhhhhhqhhq lqsaassagv vgvglggggg ggpgpslasg gssttslsrk qsskyekrgv
481 sitesqldnc kvsleadrpi vsacrktsfy hhghahhqra gnasvgggsg gagagathms
541 hsssnvlrar rgvvrmliif vltfalcnlp yharkmwqyw srsyrgdsnf nalltpltfl
601 vtyfnsgvnp llyaflsrnf rkgmkelllc swkkgkgkss snssmhhkrk alqthslptd
661 tthigneql
```

**V Trissin R PB* (not annotated; predicted based on inspection of genomic DNA documented in SXX Document – "de novo TrissinR annotations")**

```
  1 msvhslqgms tstaalptsc wqrqrqrlrr rrqqeteaas hrrahkqmrr tgsanisaay
 61 ednddnnnnn nnnynnnnnn nnnskasksp ttttnysisg lpdcqqqkte qsthhkltts
121 ihpralpwlf lspllllvll vdrslstagn ysslalnesg iestlpinat tailatаapt
181 ggsgaattal alsepttteq adeedaenss eyvfdrtdvr iifitlytiv fcccffgnll
241 vilvvtlsrr lrsitnffla nlafadfcvg lfcvmqnlsi ylidswvfge flcrmyqfvh
301 slsytasifi lvvicmeryf aivhpitckq iltaarlrmv ivtvwitsav ystpkfvfsk
361 tiknihtedg qeeeicvldr emfnsklldm infvllyvmp llvmtvlysk iaialwrssr
421 glsphvtqhq qqqqqhqqq qqqaiqensm slhnsmyhhh hhhpqhphhh qlhqqhqlpa
481 saaaaaassl asgssstsls rkhsskyekr gvsitesqld nckvsleadr pivsacrkts
541 fyhhshshnq rqgnggaggg gathmshsss nvlrarrgvv rmliifvltf alcnlpyhar
601 kmwqywsrsy rgdsnfnall tpltflvtyf nsgvnpllya flsrnfrkgm kelllcswkk
661 gkgksssnss mhhkrkalqt hslptdtthi gneql
```

**M Trissin PC (Genbank Ref #** AAF52294)

```
  1 mimtmmqtvr awqqesdveh rkqhkqrwrp dgahisaayd lnsdnddghh rvvhnqnngs
 61 pnsspnqsts afrqrqphhp ptgqqpprlp ctvthfsahw ktllillltll sastltasan
121 vtstisppin gsstdyilly gesttslvpa lttglsgdgs gaviedeeda ekaseyifdr
181 tdvriifitl ytlvfcccff gnllvilvvt lsrrlrsitn fflanlafad fcvglfcvmq
241 nlsiyliesw vfgeflcrmy qfvhslsyta sifilvvicm eryfaivhpi tckqiltaar
301 lrmvivtvwi tsavystpkf vfsktiknih tqdgqeeeic vldremfnsk lldminfvll
361 yvmpllvmtv lyskiaialw rssrgltphv vqhqhqqpqq pscqdigmgm hnsmyhhhph
421 hhhhhhqhhq lqsaassagv vgvglggggg ggpgpslasg gssttslsrk qsskyekrgv
481 sitesqvsle adrpivsacr ktsfyhhgha hhqragnasv gggsggagag athmshsssn
```

541 vlrarrgvvr mliifvltfa lcnlpyhark mwqywsrsyr gdsnfnallt pltflvtyfn
601 sgvnpllyaf lsrnfrkgmk elllcswkkg kgk***sssnss***m hhkrkalqth ***slptdtt***hig
661 neql

**PC form does not include the micro exon(at 486) present in PB which inserts ~6 AA; in both there is a precise BBS, but sequence varies (TESQLD vs. TESQVS)**

**V Trissin R PC-like (Genbank Ref #** XP_002051478.2)

1 msvhslqgms tstaalptsc wqrqrqrlrr rrqqeteaas hrrahkqmrr tgsanisaay
61 ednddnnnnn nnnynnnnnn nnnskasksp ttttnysisg lpdcqqqkte qsthhkltts
121 ihpralpwlf lsplllllvll vdrslstagn ysslalnesg iestlpinat tailataapt
181 ggsgaattal alseptteq adeedaenss eyvfdrtdvr iifitlytiv fcccffgnll
241 vilvvtlsrr lrsitnffla nlafadfcvg lfcvmqnlsi ylidswvfge flcrmyqfvh
301 slsytasifi lvvicmeryf aivhpitckq iltaarlrmv ivtvwitsav ystpkfvfsk
361 tiknihtedg qeeeicvldr emfnsklldm infvllyvmp llvmtvlysk iaialwrssr
421 glsphvtqhq qqqqqhqqq qqqaiqensm slhnsmyhhh hhhpqhphhh qlhqqhqlpa
481 saaaaaas***sl asgssstsls*** rkhsskyekr gvsi***tesqvs*** leadrpivsa crktsfyhhs
541 hshnqrqgng gaggggа***thm shss***snvlra rrgvvrmlii fvltfalcnl pyharkmwqy
601 wsrsyrgdsn fnalltpltf lvtyfnsgvn pllyaflsrn frkgmkelll cswkkgkgk***s***
661 ***ssnss***mhhkr kalqth***slpt dtt***higneql

**M Trissin R PD (Genbank Ref #** AGB92644**)**

1 mimtmmqtvr awqqesdveh rkqhkqrwrp dgahisaayd lnsdnddghh rvvhnqnngs
61 pnsspnqsts afrqrqphhp ptgqqpprlp ctvthfsahw ktllillltll sastltasan
121 vtstisppin gsstdyilly gesttslvpa lttglsgdgs gaviedeeda ekaseyifdr
181 tdvriifitl ytlvfcccff gnllvilvvt lsrrlrsitn fflanlafad fcvglfcvmq
241 nlsiyliesw vfgeflcrmy qfvhslsyta sifilvvicm eryfaivhpi tckqiltaar
301 lrmvivtvwi tsavystpkf vfsktiknih tqdgqeeeic vldremfnsk lldminfvll
361 yvmpllvmtv lyskiaialw rssrgltphv vqhqhqqpqq pscqdigmgm hnsmyhhhph
421 hhhhhhqhhq lqsaassagv vgvglggggg ggpgp***slasg gssttsls***rk qsskyekrgv
481 si***tesqld***nc kvsleadrpi vsacrktsfy hhghahhqra gnasvgggsg gagaga***thms***
541 ***hss***snvlrar rgvvrmliif vltfalcnlp yharkmwqyw srsyrgdsnf nalltpltfl
601 vtyfnsgvnp llyaflsrnf rkgmkelllc swkkgkgk***ss snss***mhhkrk alqsafftp

**V Trissin PD* (not annotated; predicted based on inspection of genomic DNA documented in SXX Document – "de novo TrissinR annotations"**)

1 msvhslqgms tstaalptsc wqrqrqrlrr rrqqeteaas hrrahkqmrr tgsanisaay
61 ednddnnnnn nnnynnnnnn nnnskasksp ttttnysisg lpdcqqqkte qsthhkltts
121 ihpralpwlf lsplllllvll vdrslstagn ysslalnesg iestlpinat tailataapt

181 ggsgaattal alsepttteq adeedaenss eyvfdrtdvr iifitlytiv fcccffgnll

241 vilvvtlsrr lrsitnffla nlafadfcvg lfcvmqnlsi ylidswvfge flcrmyqfvh

301 slsytasifi lvvicmeryf aivhpitckq iltaarlrmv ivtvwitsav ystpkfvfsk

361 tiknihtedg qeeeicvldr emfnsklldm infvllyvmp llvmtvlysk iaialwrssr

421 glsphvtqhq qqqqqqhqqq qqqaiqensm slhnsmyhhh hhhpqhphhh qlhqqhqlpa

481 saaaaaaas***sl asgssstsls*** rkhsskyekr gvsi***tesqld*** nckvsleadr pivsacrkts

541 fyhhshshnq rqgnggaggg ga***thmshss***s nvlrarrgvv rmliifvltf alcnlpyhar

601 tskmwqywsr syrgdsnfna lltpltflvt yfnsgvnpll yaflsrnfrk gmkelllcsw

661 kkgkgk***sssn ss***mhhkrkal qvsantkyyr vtlsiwh

**M Dh31 R PA (Genbank Ref #**XP_032292141.1)

1 msdqignpna tfsgsgsgsg tnvasiaesv aesgpdfdal raacetrlna sgqlagsggp
61 gaeagthcag tfdgwlcwpd tavgtsayel cpdfitgfdp aryahkecgl dgewfkhplt
121 nktwsnyttc vnledlnwrh tvnlisevgy gtsllaills lailgyfksl kcaritlhmn
181 lfasfaanns lwlvwyllvm pnsellhqsp mrcvalhitl hyfllsnysw mlcegfylht
241 vlvaafisek rlvkwliafg wgspaivifv ysmarglggt pednrhcwmn qtnyqnilmv
301 pvcismflnl lflcnivrvv llklnapasi qgscgpsrtv lqafratlll vpllglqyil
361 tpfrpapkhp wentyeiisa ftasfqglcv ailfcfcnge viaqmkrkwr mmcfsnrprt
421 nsy***tatqvs***f vrcgpplpge ekv

**V Dh31 R PA-like (Genbank Ref #** AAN16138.1 )

1 maeqtsnsss sssgsvssss sssgsshiaa esgpdfdalr aacnarlnss hqltgkrrcd
61 mphaachstv qfayklvslg sycagtfdgw lcwpdtaags sayelcpdfi tgfdparyah
121 kecgedgewf khpltnktws nyttcvnldd larnhnvnli yevgysisll aillslails
181 yfkslkcari tlhmnlfssf aansslwliw ylvvvpntel vqlspgycva lhiilhyfll
241 tnyswmlceg fylhtvlvaa fisekklvkw liafgwcspa iviciyglar gftgsweqnl
301 hcwmtdtdfn yilivpvcis iflnllflcn ivrvvllkln apasiqgscg psrtvlqafr
361 atlllvpllg lqymltpfrp gqehpleyty qvisaftasf qglcvatlfc ffngeviaqv
421 krkwrtvcfs nrprtnsy***ta tqvs***fvrcgp pvpgeekv

**M Dh31 R PC (Genbank Ref #** AGB93483)

1 msdqignpna tfsgsgsgsg tnvasiaesv aesgpdfdal raacetrlna sgqlagsggp

61 gaeagthcag tfdgwlcwpd tavgtsayel cpdfitgfdp aryahkecgl dgewfkhplt

121 nktwsnyttc vnledlnwrh tvnlisevgy gtsllaills lailgyfksl kcaritlhmn

181 lfasfaanns lwlvwyllvm pnsellhqsp mrcvalhitl hyfllsnysw mlcegfylht

241 vlvaafisek rlvkwliafg wgspaivifv ysmarglggt pednrhcwmn qtnyqnilmv

301 pvcismflnl lflcnivrvv llklnapasi qgscgpsrtv lqafratlll vpllglqyil

361 tpfrpapkhp wentyeiisa ftasfqglcv ailfcfcnge viaqmkrkwr mmcfsnrprt

421 nsy***tatqvs***f vrcgpplpge ekvxlkdmsa krrasagpqh hqphhqshql stdqqrar***sq***

481 ***slassss***fle gwrdrmpflk rrqtidhsrq sqplmeegge tvgqaksapg avdrptlmtt

541 iaedvae***tgt aat***shaasts daaaaaavga ggatgaeegh pngmgtvivr meragqqhma

601 deal

**V DH31 R PC-like (Genbank Ref #** XP_032292141.1) **(stop suppression)**

1 maeqtsnsss sssgsvssss sssgsshiaa esgpdfdalr aacnarlnss hqltgkrrcd

61 mphaachstv qfayklvslg sycagtfdgw lcwpdtaags sayelcpdfi tgfdparyah

121 kecgedgewf khpltnktws nyttcvnldd larnhnvnli yevgysisll aillslails

181 yfkslkcari tlhmnlfssf aansslwliw ylvvvpntel vqlspgycva lhiilhyfll

241 tnyswmlceg fylhtvlvaa fisekklvkw liafgwcspa iviciyglar gftgsweqnl

301 hcwmtdtdfn yilivpvcis iflnllflcn ivrvvllkln apasiqgscg psrtvlqafr

361 atlllvpllg lqymltpfrp gqehpleyty qvisaftasf qglcvatlfc ffngeviaqv

421 krkwrtvcfs nrprtnsy***ta tqvs***fvrcgp pvpgeekvxl kdlsasakrr ssaphrphqq

481 qqqqqqqqqq qqlqqqqqhl laqqqrarsq slgm***tgatss s***srlidgwrh klrllrrpsv

541 dnahqrqplm eetagnsqpq maantqaead rptlmttiae dvaeaattta ***tatgittttt***

601 ***t***nnateqrng tgmglglvsr sglglapgig vvivsvdand qqhiadeal

**M Hector R PA (Genbank Ref #** AGB93483)

```
  1 mattssdses qnvdvasqaq tqdnlriflk hlyaecvfry qnvtydtddp sfslgpatdy
 61 dsdlpenfsp vprylenaam negvidmrnv deelaekeel matvvsatma tnqkenrlfc
121 plnfdgylcw prtpagtvls qycpdfvegf nrkflahktc lengswyrhp vsnqtwsnyt
181 ncvdyedlef rqfinelyvk gyalsllall isiiiflgfk slrctririh vhlfaslact
241 cvawilwyrl vversetiae nplwciglhl vvhyfmlvny fwmfceglhl hlvlvvvfvk
301 dtivmrwfiv iswfspipia ivyglarhfs spdnkhcwit dslylwifsv pitlsllasf
361 iflinvlrvi vrklhpqsaq paplairkav ratiilvplf glqhfllpyr pdagtqldhf
421 yqmlsvvlvs lqgfvvsflf cfanhdvtfa irtllnkllp slvtpppags ntgqmatttp
481 srelgv
```

**V Hector R PA-like (Genbank Ref # XP_032296576)**

```
  1 mallraaltm ataayfemtv sgamppqpqd nlrtflkhly aecvyryqnd tataqlatep
 61 ddggllllety tmipryleqa vlnegtidmq dvdeeaasen elyatvlsat matnhhshnt
121 semetlycpv nfdgylcwpr tpagtvlsqy cpdfvegfns kflahktcle tgswfrhpvs
181 nqtwsnytnc vdyedfqfrq fvnelyvkgy alsllalfis iviflgfksl rctririhvh
241 lfaslactci awilwyrlvv ehteqiaenp pwcialhlvv hyfmlvnyfw mfceglhlhl
301 vlvvvfvkdt ivmrwfklls wllpllfvlp ygvarhfsan dnahcwmnds fylwifsvpi
361 tlsllasfif linvlrvivr klhpqsaqpa plairkavra tiilvplfgl qhfllpyrpd
421 agtqldrfyq llsvvlvslq gfvvsflfcf anhdvtfamr tllnklmptl vapppagsnt
481 gqlatttpsr elgv
```

**M PDF R PA (Genbank Ref #** AAF45788)

```
  1 mtllsnildc ggcisaqrft rllrqsgssg pspsaptagt fesksmlept sshslatgrv
 61 pllhdfdast tespgtyvld gvarvaqlal eptvmdalpd sdteqvlgnl nssapwnltl
121 asaaatnfen csalfvnytl pqtglycnwt wdtllcwppt pagvlarmnc pggfhgvdtr
181 kfairkceld grwgsrpnat evnppgwtdy gpcykpeiir lmqqmgskdf dayidiarrt
241 rtleivglcl slfalivsll ifctfrslrn nrtkihknlf vamvlqviir ltlyldqfrr
301 gnkeaatnts lsvientpyl ceasyvlley artamfmwmf iegl ylhnmv tvavfqgsfp
361 lkffsrlgwc vpilmttvwa rctvmymdts lgeclwnynl tpyywilegp rlavillnfc
421 flvniirvlv mklrqsqasd ieqtrkavra aivllpllgi tnllhqlapl ktatnfavws
481 ygthfltsfq gffialiycf lngevravll kslatqlsvr ghpewapkra smysgaynta
541 pdtdavqpag dpsatgkris ppnkrlngrk pssasivmih epqqrqrlmp rlqnkarekg
601 kdrvektdae aepdptishi hskeagsars rtrgskwimg icfrgqkvlr vpsassvppe
661 svvfelseq
```

**V PDF R PA-like (Genbank Ref #** XP_032288784.1)

```
  1 mptaapssnr tvsthhsttm tststtygtt sttqaaltsf epmavtaagm dvtmagtepl
 61 sstsiptfwn ssintasans ydncsalfan ytqpttviyc nwtwdsllcw pptpagatah
121 mhcpagyhgv dtrkfanrkc eldghwagrp nsteqkptgw tdygpcykpe virlmqeikd
181 vnlymdiaqr trtleiiglc lslfaliisl mifcafrslr nnrtkihknl fvamvlqviv
241 rltlyldqfr rgksdsannt slsvientpy lceasyvlle yartamfmwm fieglylhnm
301 itvavfqgnf plvffsllgw gmpvlmtfvw vqctaifmdt slgdclwnyn ltpyywileg
361 prltvimlnf fflvniirvl vmklrqsqas eieqtrkavr aaivllpllg itnllhlvpa
421 lktawkfaiw syvthfltsf qgffialiyc flngevravm lksiavwlsv rghpewapkr
481 psmysgaynt apdtdpqlkq gdpqqsgkrl sqstkrsnsr kassvtivis tepqihryvp
541 rrnnnnrast gsarvrgilk ateepasgsa vgqrirstdd gasttgrnsn wmfglcfrgq
601 kvlrvppass vppesvvfel seq
```

***M PDF R PD*** **(Genbank Ref #** AHN59298**)**

1 mtllsnildc ggcisaqrft rllrqsgssg pspsaptagt fesksmlept sshslatgrv

61 pllhdfdast tespgtyvld gvarvaqlal eptvmdalpd sdteqvlgnl nssapwnltl

121 asaaatnfen csalfvnytl pqtglycnwt wdtllcwppt pagvlarmnc pggfhgvdtr

181 kfairkceld grwgsrpnat evnppgwtdy gpcykpeiir lmqqmgskdf dayidiarrt

241 rtleivglcl slfalivsll ifctfrslrn nrtkihknlf vamvlqviir ltlyldqfrr

301 gnkeaatnts lsvientpyl ceasyvlley artamfmwmf ieglylhnmv tvavfqgsfp

361 lkffsrlgwc vpilmttvwa rctvmymdts lgeclwnynl tpyywilegp rlavillnfc

421 flvniirvlv mklrq***sqasd ie***qtrkavra aivllpllgi tnllhqlapl ktatnfavws

481 ygthfltsfq gffialiycf lngevravll k***slatqls***vr ghpewapkra smysgaynta

541 pdtdavqpag dpsatgkris ppnkrlngrk pssasivmih epqqrqrlmp rlqnkarekg

601 kdrvektdae aepdptishi hskeagsar***s rtrgs***kwimg icfrgqkdkc vmpgsqktqq

661 ifmtsqmppt stlaava***tti tttsttt***aa k***ttiasiati at***mtkskaka kaiskshqiq

721 mpka

**V PDF R PD-like (Genbank Ref #** XP_032288761.1**)**

1 mptaapssnr tvsthhsttm tststtygtt sttqaaltsf epmavtaagm dvtmagtepl

61 sstsiptfwn ssintasans ydncsalfan ytqpttviyc nwtwdsllcw pptpagatah

121 mhcpagyhgv dtrkfanrkc eldghwagrp nsteqkptgw tdygpcykpe virlmqeikd

181 vnlymdiaqr trtleiiglc lslfaliisl mifcafrslr nnrtkihknl fvamvlqviv

241 rltlyldqfr rgksdsannt slsvientpy lceasyvlle yartamfmwm fieglylhnm

301 itvavfqgnf plvffsllgw gmpvlmtfvw vqctaifmdt slgdclwnyn ltpyywileg

361 prltvimlnf fflvniirvl vmklrq***sqas eie***qtrkavr aaivllpllg itnllhlvpa

421 lktawkfaiw syvthfltsf qgffialiyc flngevravm lksiavwlsv rghpewapkr

481 psmysgaynt apdtdpqlkq gdpqqsgkrl sqstkrsnsr kassvtivis tepqihryvp

541 rrnnnnrast gsarvrgilk ateepasgsa vgqrirstdd gasttgrnsn wmfglcfrgq

601 knkcvipnaq vsqqifmtsq lp***tataattt tttt***vaaaaa a***ttttaat***av aaa***tttsris***

661 saaaaaaaaa aailqkqqtp ka

**M Dh44 R1 PA (Genbank Ref #** AAF58250)

```
  1 msdhnhidsv nasgsdplld lhnldgiges velqclvqeh ieastygnds ghcltqfdsi
 61 lcwprtargt lavlqcmdel qgihydsskn atrfchangt wekytnydac ahlpapesvp
121 efevivelpt iiyyigytls lvslslaliv fayfkelrcl rntihanlff tyimsalfwi
181 lllsvqisir sgvgsciali tlfhfftltn ffwmlvegly lymlvvktfs gdnlrfniya
241 sigwggpalf vvtwavaksl tvtystpeky eincpwmqet hvdwiyqgpv cavliinltf
301 llrimwvlit klrsantvet rqyrkaakal lvliplfgit ylvvlagpse sglmghmfav
361 lravllstqg fsvslfycfl nsevrnalrh histwrdtrt iqlnqnrryt tksfskgggs
421 praesmrplt syygrgkres cvssatttl vgqhaplslh rgsnnalhtm ptlaanamss
481 gstlsvmpra isplmrqgle ensv
```

**V Dh44 R1 PA-like (Genbank Ref #** XP_032292296.1)

```
  1 mskdnnnnnn nnngnnnpia sealnssvnd alwsldnldg inqsvelhcl lqqqieatty
 61 gnasdhcltq fdtilcwprt argtlavlqc mdelqgihyd ssknatrfch sngtwaqytd
121 ydacahlpae tqtvpefeti velptiiyyi gyalslvslt lalivfayyk elrclrntih
181 anlfftyims alfwilllsv qisirsglss cialvtlfhf ftltnffwml veglylymlv
241 vktfsgdnir fniyasigwg gpalfvvtwa vaksltvtyn nnmekydinc pwmeetnvdw
301 ifqgpvcavl iinltfllri mwvlitklrs antvetrqyr kaakallvli plfgitylvv
361 lagpsesglm ghmfavlrav llstqgfwvs lfycflnsev rnalrhhast wrdtrniqrn
421 qnrryttksf skgggsprae smrpltsyyg rgkrescvss atttlvgqh aplslhrgsn
481 nalhmlptni agtgstlsvm praisplmkg leensv
```

**M Dh44 R2 PA (Genbank Ref #** AAF58501**)**

1 madddlralv dslddasqed lakvianfsv dmlqrasali gaqqgssggq lqnrtlqcqq
61 qqqreeeqas lealasggkr ilqcpssfds vlcwprtnag slavlpcfee fkgvhydttd
121 natrfcfpng twdhysdydr chqnsgsipv vpdfspnvel paiiyaggyf lsfatlvval
181 iiflsfkdlr clrntihanl fltyitsall wiltlflqvi ttessqagci tlvimfqyfy
241 ltnffwmfve glylytlvvq tfssdnisfi iyaligwgcp avcilvwsia kafaphlene
301 hfngleidca wmreshidwi fkvpaslall vnlvflirim wvlitklr***sa htlet***rqyyk
361 askallvlip lfgityllvl tgpeqgisrn lfeairafli stqgffvalf ycflnsevrq
421 tlrhgftrwr esrnihrnss iknr/rhrask dy***slrsrte***s lrltstspip tghye

**V DH44 R2 PA-like (Genbank Ref #** XP_032292527.1**)**

1 maedelqalv erlddasaen ianaianfsl emlqrasali gtqqphsgdi linrtledqc
61 kqqaeqqdal yssphlshtg idqyssskas adkptlycpt sfdsvlcwpr tsastwailp
121 cfeefkgvhy dttenatrfc hangtwnhys nysschqqlg svpvvpdfsa svdlpaiiya
181 ggyfisfatl vvaliiflsf kdlrclrnti hanlfltyit sallwiltlf lqvittessq
241 agcitlvimf qyfyltnfsw mfveglylyt lvvqtfssen isfviyalig wgcpalcilf
301 wsiakafash lenehfngle iectwmresh idwifkgpas lailinlvfl irimwvlitk
361 lr***sahtlet***r qyykaskall vliplfgity llvltgpeqg isrnlfeamr afllstqgff
421 valfycflns evrqtlrhrf irwresrnih rssslknr/rh rtskdy***sqrs rte***slrltss
481 spvpaghfe

**M DH44 R2 PB (Genbank Ref #** AAM68690**)**

1 madddlralv dslddasqed lakvianfsv dmlqrasali gaqqgssggq lqnrtlqcqq
61 qqqreeeqas lealasggkr ilqcpssfds vlcwprtnag slavlpcfee fkgvhydttd
121 natrfcfpng twdhysdydr chqnsgsipv vpdfspnvel paiiyaggyf lsfatlvval
181 iiflsfkdlr clrntihanl fltyitsall wiltlflqvi ttessqagci tlvimfqyfy
241 ltnffwmfve glylytlvvq tfssdnisfi iyaligwgcp avcilvwsia kafaphlene
301 hfngleidca wmreshidwi fkvpaslall vnlvflirim wvlitklr***sa htlet***rqyyk
361 askallvlip lfgityllvl tgpeqgisrn lfeairafli stqgffvalf ycflnsevrq
421 tlrhgftrwr esrnihrnss iknr/steecv iclrpsphtr lgslqryhsi ditdfv

**V DH44 R2 PB-like (Genbank Ref #** XP_032292526.1**)**

1 maedelqalv erlddasaen ianaianfsl emlqrasali gtqqphsgdi linrtledqc
61 kqqaeqqdal yssphlshtg idqyssskas adkptlycpt sfdsvlcwpr tsastwailp
121 cfeefkgvhy dttenatrfc hangtwnhys nysschqqlg svpvvpdfsa svdlpaiiya
181 ggyfisfatl vvaliiflsf kdlrclrnti hanlfltyit sallwiltlf lqvittessq
241 agcitlvimf qyfyltnfsw mfveglylyt lvvqtfssen isfviyalig wgcpalcilf
301 wsiakafash lenehfngle iectwmresh idwifkgpas lailinlvfl irimwvlitk
361 lr***sahtlet***r qyykaskall vliplfgity llvltgpeqg isrnlfeamr afllstqgff
421 valfycflns evrqtlrhrf irwresrnih rssslknr/st eecviclrps lhtrvgslkh
481 chsidltdfv
