## Supplementary material for "The incidence of candidate binding sites for β-arrestin in Drosophila neuropeptide GPCRs": S2 Text

AstA R2

*D melanogaster* has two isoforms that differ in CT: one with and one without BBS

PA does not have a BBS
PB has a BBS

Alternative splicing: PB specific exon comes before PA-specific exon by about 800 bp
http://flybase.org/jbrowse/?data=data%2Fjson%2Fdmel&loc=3R%3A28735576..28746319&tracks=Gene_span%2CGene%2Cprote_in&highlight=

Blastp search reveals only a single isoform in *D virilis*, without a BBS and its exon structure resembles the PA of melanogaster.

Search for an exon in region that would be the PB-specific exon area (in between the penultimate and final exons

Here is genome viewer for *D virilis* gene:
https://www.ncbi.nlm.nih.gov/genome/gdv/browser/protein/?id=XP_002053983.1

translate and look for PB-like CT sequence
melanogaster:
LLYAFLSENF RKAFYK (splice)GLQSNRLGMW**TTTHQD**VSSEKTTY

bp#1 = bp# 8481011 of the shotgun sequence

here is one candidate splicing in at bp 5299:
(splice) EKIKIKPN***SWHSVGT***RGRRSAPLCFTYF *
At the approximate position corresponding to that in *D melanogaster* (see below), 4 AAs longer, but splices correctly and contains a BBS

```
   A  E  T  K  K  A  N  V  M  S  A  R  A  I  K  N  Q  *  K  T        F1
     R  K  P  K  K  Q  M  *  *  A  L  G  Q  L  K  I  N  R  K  L      F2
      G  N  Q  K  S  K  C  N  E  R  S  G  N  *  K  S  I  E  N  C     F3
  1 GCGGAAACCAAAAAAGCAAATGTAATGAGCGCTCGGGCAATTAAAAATCAATAGAAAACT 60
    ----:----|----:----|----:----|----:----|----:----|----:----|

    V  D  R  K  R  H  S  M  N  L  S  N  I  T  L  T  L  P  S  N       F1
     *  T  E  S  A  T  A  *  I  *  A  T  *  L  *  P  C  P  A  T      F2
      R  Q  K  A  P  Q  H  E  S  K  Q  H  N  S  D  P  A  Q  Q  Q     F3
 61 GTAGACAGAAAGCGCCACAGCATGAATCTAAGCAACATAACTCTGACCCTGCCCAGCAAC 120
    ----:----|----:----|----:----|----:----|----:----|----:----|

    S  S  W  L  E  S  Q  L  E  L  T  T  A  S  S  N  L  D  N  S       F1
     A  A  G  W  S  R  S  W  S  *  P  Q  R  A  A  T  W  T  I  Q      F2
      Q  L  A  G  V  A  A  G  A  D  H  S  E  Q  Q  P  G  Q  F  N     F3
121 AGCAGCTGGCTGGAGTCGCAGCTGGAGCTGACCACAGCGAGCAGCAACCTGGACAATTCA 180
    ----:----|----:----|----:----|----:----|----:----|----:----|

    T  L  S  S  F  Y  A  E  A  E  A  I  R  A  T  V  R  W  V  V       F1
     L  Y  P  H  S  M  P  K  R  R  P  Y  G  P  L  C  A  G  W  C      F2
```

```
      F  I  L  I  L  C  R  S  G  G  H  T  G  H  C  A  L  G  G  A    F3
181 ACTTTATCCTCATTCTATGCCGAAGCGGAGGCCATACGGGCCACTGTGCGCTGGGTGGTG 240
    ----:----|----:----|----:----|----:----|----:----|----:----|

    P  F  F  F  G  I  I  A  I  S  G  F  F  G  N  L  L  V  I  L      F1
     H  S  F  L  A  L  S  P  S  P  V  S  L  A  T  C  W  S  Y  W     F2
      I  L  F  W  H  Y  R  H  L  R  F  L  W  Q  P  V  G  H  I  G    F3
241 CCATTCTTTTTTGGCATTATCGCCATCTCCGGTTTCTTTGGCAACCTGTTGGTCATATTG 300
    ----:----|----:----|----:----|----:----|----:----|----:----|

    V  V  L  L  N  K  N  M  H  S  T  T  N  L  L  I  V  N  L  A      F1
     W  C  C  *  T  R  T  C  T  P  R  Q  I  C  *  L  S  T  W  P     F2
      G  V  A  E  Q  E  H  A  L  H  D  K  S  A  D  C  Q  L  G  R    F3
301 GTGGTGTTGCTGAACAAGAACATGCACTCCACGACAAATCTGCTGATTGTCAACTTGGCC 360
    ----:----|----:----|----:----|----:----|----:----|----:----|

    A  A  D  L  L  F  V  I  F  C  V  P  F  T  A  I  D  Y  V  T      F1
     P  P  I  C  C  S  S  S  S  A  Y  H  S  R  Q  L  T  M  *  H     F2
      R  R  S  A  V  R  H  L  L  R  T  I  H  G  N  *  L  C  D  T    F3
361 GCCGCCGATCTGCTGTTCGTCATCTTCTGCGTACCATTCACGGCAATTGACTATGTGACA 420
    ----:----|----:----|----:----|----:----|----:----|----:----|

    Q  H  W  P  F  G  K  M  W  C  R  S  V  Q  Y  L  I  V  V  T      F1
     S  I  G  P  L  A  R  C  G  A  A  V  S  S  I  *  *  W  S  P     F2
      A  L  A  L  W  Q  D  V  V  P  Q  C  P  V  F  D  S  G  H  R    F3
421 CAGCATTGGCCCTTTGGCAAGATGTGGTGCCGCAGTGTCCAGTATTTGATAGTGGTCACC 480
    ----:----|----:----|----:----|----:----|----:----|----:----|

    A  Y  A  S  I  Y  T  L  V  L  M  S  I  D  R  F  L  A  V  V      F1
     P  M  L  P  S  I  R  S  S  *  C  P  L  I  A  S  W  P  S  F     F2
      L  C  F  H  L  Y  A  R  P  D  V  H  *  S  L  L  G  R  R  S    F3
481 GCCTATGCTTCCATCTATACGCTCGTCCTGATGTCCATTGATCGCTTCTTGGCCGTCGTT 540
    ----:----|----:----|----:----|----:----|----:----|----:----|

    H  P  I  R  S  R  M  L  R  T  E  H  I  T  K  I  A  I  F  T      F1
     I  P  Y  A  L  A  C  C  A  P  S  I  S  P  R  *  P  Y  L  R     F2
      S  H  T  L  S  H  A  A  H  R  A  Y  H  Q  D  S  H  I  Y  A    F3
541 CATCCCATACGCTCTCGCATGCTGCGCACCGAGCATATCACCAAGATAGCCATATTTACG 600
    ----:----|----:----|----:----|----:----|----:----|----:----|

    L  W  T  V  V  L  T  V  S  M  P  V  T  F  A  H  D  V  V  V      F1
     S  G  R  W  C  *  L  S  P  C  R  S  L  L  R  M  M  W  W  *     F2
      L  D  G  G  A  D  C  L  H  A  G  H  F  C  A  *  C  G  G  K    F3
601 CTCTGGACGGTGGTGCTGACTGTCTCCATGCCGGTCACTTTTGCGCATGATGTGGTGGTA 660
    ----:----|----:----|----:----|----:----|----:----|----:----|

    S  V  C  V  W  H  T  A  S  G  P  V  A  C  C  A  R  M  C  C      F1
     V  F  V  C  G  I  Q  P  Q  G  L  L  P  A  V  P  E  C  A  A     F2
      C  L  C  V  A  Y  S  L  R  A  C  C  L  L  C  P  N  V  L  R    F3
661 AGTGTTTGTGTGTGGCATACAGCCTCAGGGCCTGTTGCCTGCTGTGCCCGAATGTGCTGC 720
    ----:----|----:----|----:----|----:----|----:----|----:----|

    E  *  Q  R  P  A  K  I  Y  C  C  A  S  S  A  P  R  *  M  S      F1
     N  N  R  G  Q  L  K  F  I  A  A  H  H  R  H  L  V  R  C  P     F2
      I  T  E  A  S  *  N  L  L  L  R  I  I  G  T  S  L  D  V  P    F3
721 GAATAACAGAGGCCAGCTAAAATTTATTGCTGCGCATCATCGGCACCTCGTTAGATGTCC 780
    ----:----|----:----|----:----|----:----|----:----|----:----|

    R  A  A  A  P  R  Q  W  A  K  T  L  L  L  W  I  W  L  W  P      F1
     E  P  L  L  H  V  N  G  Q  K  H  F  C  C  G  S  G  F  G  L     F2
      S  R  C  S  T  S  M  G  K  N  T  F  V  V  D  L  A  L  A  F    F3
781 CGAGCCGCTGCTCCACGTCAATGGGCAAAAACACTTTTGTTGTGGATCTGGCTTTGGCCT 840
    ----:----|----:----|----:----|----:----|----:----|----:----|
```

```
     L  P  S  C  P  R  F  Y  F  V  I  V  V  I  V  F  F  S  S  *    F1
      Y  L  A  A  H  V  F  I  L  L  L  L  L  L  F  F  F  L  L  D   F2
       T  *  L  P  T  F  L  F  C  Y  C  C  Y  C  F  F  F  F  L  I  F3
 841 TTACCTAGCTGCCCACGTTTTTATTTTGTTATTGTTGTTATTGTTTTTTTTTCTTCTTGA 900
     ----:----|----:----|----:----|----:----|----:----|----:----|

     F  Y  L  S  L  F  F  L  F  T  F  T  C  L  Y  S  E  F  G  R    F1
      F  I  *  V  Y  S  F  F  S  H  L  R  V  Y  I  Q  N  S  V  D   F2
       L  F  K  F  I  L  S  F  H  I  Y  V  F  I  F  R  I  R  *  M  F3
 901 TTTTATTTAAGTTTATTCTTTCTTTTCACATTTACGTGTTTATATTCAGAATTCGGTAGA 960
     ----:----|----:----|----:----|----:----|----:----|----:----|

     C  G  R  L  F  C  F  C  Y  *  F  A  I  *  N  A  *  I  L  I    F1
      A  G  A  C  S  V  F  V  T  N  L  L  F  E  M  H  K  F  *  L   F2
       R  A  P  V  L  F  L  L  L  I  C  Y  L  K  C  I  N  F  N  Y  F3
 961 TGCGGGCGCCTGTTCTGTTTTTGTTACTAATTTGCTATTTGAAATGCATAAATTTTAATT 1020
     ----:----|----:----|----:----|----:----|----:----|----:----|

     I  E  L  I  Q  N  T  H  E  R  S  S  C  W  S  Y  G  Q  Y  H    F1
      *  S  *  Y  R  T  H  T  S  E  A  V  A  G  H  M  A  N  T  I   F2
       R  A  D  T  E  H  T  R  A  K  Q  L  L  V  I  W  P  I  P  S  F3
1021 ATAGAGCTGATACAGAACACACACGAGCGAAGCAGTTGCTGGTCATATGGCCAATACCAT 1080
     ----:----|----:----|----:----|----:----|----:----|----:----|

     L  Q  I  I  C  A  T  A  I  Y  N  L  Q  L  D  F  L  L  C  L    F1
      S  K  S  Y  V  P  Q  L  F  T  I  C  S  W  I  S  Y  S  A  Y   F2
       P  N  H  M  C  H  S  Y  L  Q  F  A  V  G  F  L  T  L  L  I  F3
1081 CTCCAAATCATATGTGCCACAGCTATTTACAATTTGCAGTTGGATTTCTTACTCTGCTTA 1140
     ----:----|----:----|----:----|----:----|----:----|----:----|

     F  S  C  P  K  N  *  W  I  I  D  H  R  H  T  D  T  Q  T  H    F1
      S  V  A  P  K  T  N  G  S  L  I  T  D  T  Q  T  H  R  H  T   F2
       Q  L  P  Q  K  L  M  D  H  *  S  Q  T  H  R  H  T  D  T  Q  F3
1141 TTCAGTTGCCCCAAAAACTAATGGATCATTGATCACAGACACACAGACACACAGACACAC 1200
     ----:----|----:----|----:----|----:----|----:----|----:----|

     R  Q  A  T  G  Y  R  P  G  I  R  C  V  *  *  V  S  G  D  M    F1
      D  R  Q  Q  A  T  G  R  A  Y  A  A  Y  D  K  S  L  A  I  W   F2
       T  G  N  R  L  Q  A  G  H  T  L  R  M  I  S  L  W  R  Y  G  F3
1201 AGACAGGCAACAGGCTACAGGCCGGGCATACGCTGCGTATGATAAGTCTCTGGCGATATG 1260
     ----:----|----:----|----:----|----:----|----:----|----:----|

     A  L  M  L  V  S  G  *  S  A  H  L  T  L  A  L  P  S  F  I    F1
      P  *  C  W  F  L  G  R  V  R  I  *  R  S  P  C  H  H  L  S   F2
       L  D  A  G  F  W  V  E  C  A  F  N  A  R  L  A  I  I  Y  P  F3
1261 GCCTTGATGCTGGTTTCTGGGTAGAGTGCGCATTTAACGCTCGCCTTGCCATCATTTATC 1320
     ----:----|----:----|----:----|----:----|----:----|----:----|

     H  L  P  *  R  R  *  C  F  Y  D  Q  C  P  E  C  E  Q  Q  V    F1
      I  C  H  K  G  V  N  V  F  M  I  N  A  L  S  V  N  N  K  L   F2
       F  A  I  K  A  L  M  F  L  *  S  M  P  *  V  *  T  T  S  *  F3
1321 CATTTGCCATAAAGGCGTTAATGTTTTTATGATCAATGCCCTGAGTGTGAACAACAAGTT 1380
     ----:----|----:----|----:----|----:----|----:----|----:----|

     N  C  L  I  S  A  A  K  Q  T  S  W  R  A  N  L  L  L  L  P    F1
      I  V  *  *  A  P  P  N  K  H  L  G  V  Q  I  C  C  C  C  P   F2
       L  F  N  K  R  R  Q  T  N  I  L  A  C  K  F  V  A  A  A  H  F3
1381 AATTGTTTAATAAGCGCCGCCAAACAAACATCTTGGCGTGCAAATTTGTTGCTGCTGCCC 1440
     ----:----|----:----|----:----|----:----|----:----|----:----|

     I  S  L  F  S  Q  W  T  E  *  N  F  Y  V  L  A  I  K  L  I    F1
      F  L  C  L  A  N  G  R  S  K  T  F  M  C  *  Q  S  N  *  S   F2
```

```
      F  F  V  *  P  M  D  G  V  K  L  L  C  A  S  N  Q  I  N  R   F3
1441 ATTTCTTTGTTTAGCCAATGGACGGAGTAAAACTTTTATGTGCTAGCAATCAAATTAATC 1500
     ----:----|----:----|----:----|----:----|----:----|----:----|

     V  F  K  L  I  S  N  I  F  K  Y  F  F  S  I  N  S  Q  F  P    F1
      Y  L  N  L  F  Q  I  S  S  N  I  S  F  Q  L  I  R  N  F  L   F2
       I  *  T  Y  F  K  Y  L  Q  I  F  L  F  N  *  F  A  I  S  C  F3
1501 GTATTTAAACTTATTTCAAATATCTTCAAATATTTCTTTTCAATTAATTCGCAATTTCCT 1560
     ----:----|----:----|----:----|----:----|----:----|----:----|

     A  N  C  T  *  I  *  *  L  F  W  K  K  L  S  E  I  R  L  K    F1
      P  T  A  L  K  Y  N  N  Y  F  G  K  N  *  A  K  *  D  *  R   F2
       Q  L  H  L  N  I  I  I  I  L  E  K  I  K  R  N  K  I  K  E  F3
1561 GCCAACTGCACTTAAATATAATAATTATTTTGGAAAAAATTAAGCGAAATAAGATTAAAG 1620
     ----:----|----:----|----:----|----:----|----:----|----:----|

     K  M  L  M  V  I  *  S  R  C  G  T  H  W  E  Q  I  Y  I  L    F1
      K  C  L  W  L  F  K  A  D  A  V  H  I  G  N  R  Y  I  Y  *   F2
       N  A  Y  G  Y  L  K  Q  M  R  Y  T  L  G  T  D  I  Y  I  K  F3
1621 AAAATGCTTATGGTTATTTAAAGCAGATGCGGTACACATTGGGAACAGATATATATATTA 1680
     ----:----|----:----|----:----|----:----|----:----|----:----|

     K  S  P  *  S  F  I  F  L  K  C  F  Y  I  N  L  N  Y  R  L    F1
      K  A  L  D  H  S  F  F  *  N  V  F  I  S  I  *  T  I  A  S   F2
       K  P  L  I  I  H  F  S  K  M  F  L  Y  Q  F  E  L  S  P  Q  F3
1681 AAAAGCCCTTGATCATTCATTTTTCTAAAATGTTTTTATATCAATTTGAACTATCGCCTC 1740
     ----:----|----:----|----:----|----:----|----:----|----:----|

     S  F  W  Y  G  S  T  F  V  G  A  R  S  S  S  R  K  S  S  H    F1
      A  F  G  M  V  P  H  L  W  V  L  E  V  R  A  V  K  A  H  I   F2
       L  L  V  W  F  H  I  C  G  C  S  K  F  E  P  *  K  L  T  F  F3
1741 AGCTTTTGGTATGGTTCCACATTTGTGGGTGCTCGAAGTTCGAGCCGTAAAAGCTCACAT 1800
     ----:----|----:----|----:----|----:----|----:----|----:----|

     F  F  E  F  K  I  F  I  I  I  F  S  R  S  *  K  *  C  I  I    F1
      S  L  N  L  K  Y  L  L  L  F  F  Q  E  A  E  N  N  V  L  *   F2
       L  *  I  *  N  I  Y  Y  Y  F  F  K  K  L  K  I  M  Y  Y  K  F3
1801 TTCTTTGAATTTAAAATATTTATTATTATTTTTCAAGAAGCTGAAAATAATGTATTATA 1860
     ----:----|----:----|----:----|----:----|----:----|----:----|

     R  *  R  I  L  I  S  N  T  V  I  T  K  F  I  L  K  K  F  K    F1
      G  N  A  F  *  Y  P  I  R  L  L  R  N  L  Y  *  R  N  L  R   F2
       V  T  H  F  N  I  Q  Y  G  Y  Y  E  I  Y  I  K  E  I  *  D  F3
1861 AGGTAACGCATTTTAATATCCAATACGGTTATTACGAAATTTATATTAAAGAAATTTAAG 1920
     ----:----|----:----|----:----|----:----|----:----|----:----|

     I  N  L  Y  I  *  L  I  K  L  K  P  A  D  S  S  I  Y  S  P    F1
      L  I  Y  I  F  N  *  L  N  *  N  Q  P  I  L  Q  S  T  L  H   F2
       *  F  I  Y  L  T  N  *  T  K  T  S  R  F  F  N  L  L  S  I  F3
1921 ATTAATTTATATATTTAACTAATTAAACTAAAACCAGCCGATTCTTCAATCTACTCTCCA 1980
     ----:----|----:----|----:----|----:----|----:----|----:----|

     Y  Y  F  R  W  N  P  *  K  S  R  D  S  T  Q  L  L  R  P  H    F1
      I  I  F  D  G  I  H  K  N  L  E  T  Q  L  S  C  L  G  H  I   F2
       L  F  S  M  E  S  I  K  I  S  R  L  N  S  A  A  *  A  T  Y  F3
1981 TATTATTTTCGATGGAATCCATAAAAATCTCGAGACTCAACTCAGCTGCTTAGGCCACAT 2040
     ----:----|----:----|----:----|----:----|----:----|----:----|

     T  R  T  A  A  F  L  F  S  H  F  F  F  F  G  Y  I  Y  L  I    F1
      L  E  R  Q  L  F  Y  S  A  I  F  F  F  L  D  I  F  I  *  F   F2
       *  N  G  S  F  F  I  Q  P  F  F  F  F  W  I  Y  L  F  N  L  F3
2041 ACTAGAACGGCAGCTTTTTTATTCAGCCATTTTTTTTTTTTTGGATATATTTATTTAATT 2100
     ----:----|----:----|----:----|----:----|----:----|----:----|
```

```
      Y  A  P  K  S  F  *  H  *  C  K  R  N  G  N  K  V  K  C  L    F1
       M  H  P  K  V  S  S  I  N  V  S  E  M  E  T  K  L  N  V  Y   F2
        C  T  Q  K  F  L  A  L  M  *  A  K  W  K  Q  S  *  M  F  I  F3
2101 TATGCACCCAAAAGTTTCTAGCATTAATGTAAGCGAAATGGAAACAAAGTTAAATGTTTA 2160
     ----:----|----:----|----:----|----:----|----:----|----:----|

      *  N  S  N  G  S  K  L  Q  A  E  S  K  A  A  L  S  S  R  N    F1
       K  T  L  M  G  Q  N  C  K  Q  N  Q  K  L  L  F  H  R  V  T   F2
        K  L  *  W  V  K  I  A  S  R  I  K  S  C  S  F  I  A  *  L  F3
2161 TAAAACTCTAATGGGTCAAAATTGCAAGCAGAATCAAAAGCTGCTCTTTCATCGCGTAAC 2220
     ----:----|----:----|----:----|----:----|----:----|----:----|

      Y  S  L  L  L  F  K  Y  F  R  S  K  R  F  S  K  R  *  F  E    F1
       I  R  F  C  Y  L  S  T  S  D  Q  S  D  L  A  S  V  N  L  N   F2
        F  A  F  V  I  *  V  L  Q  I  K  A  I  *  Q  A  L  I  *  I  F3
2221 TATTCGCTTTTGTTATTTAAGTACTTCAGATCAAAGCGATTTAGCAAGCGTTAATTTGAA 2280
     ----:----|----:----|----:----|----:----|----:----|----:----|

      *  C  S  E  N  A  N  Q  L  I  G  W  K  R  L  G  Q  L  P  H    F1
       N  V  R  K  T  L  I  S  L  L  A  G  N  V  W  D  S  C  H  I   F2
        M  F  G  K  R  *  S  A  Y  W  L  E  T  S  G  T  A  A  T  S  F3
2281 TAATGTTCGGAAAACGCTAATCAGCTTATTGGCTGGAAACGTCTGGGACAGCTGCCACAT 2340
     ----:----|----:----|----:----|----:----|----:----|----:----|

      L  N  S  F  P  V  M  L  Q  R  S  S  L  A  K  I  Q  R  N  S    F1
       *  I  A  F  L  *  C  F  N  E  V  H  *  Q  K  Y  N  A  I  P   F2
        K  *  L  S  C  N  A  S  T  K  F  I  S  K  N  T  T  Q  F  P  F3
2341 CTAAATAGCTTTCCTGTAATGCTTCAACGAAGTTCATTAGCAAAAATACAACGCAATTCC 2400
     ----:----|----:----|----:----|----:----|----:----|----:----|

      P  T  Y  G  Y  I  F  Q  E  L  C  V  C  V  C  E  V  G  I  F    F1
       L  L  M  D  T  F  S  R  N  C  V  C  V  C  V  R  W  G  Y  F   F2
        Y  L  W  I  H  F  P  G  T  V  C  V  C  V  *  G  G  D  I  F  F3
2401 CCTACTTATGGATACATTTTCCAGGAACTGTGTGTGTGTGTGTGTGAGGTGGGGATATTT 2460
     ----:----|----:----|----:----|----:----|----:----|----:----|

      L  *  V  W  L  R  P  I  H  N  Q  I  V  A  C  N  *  T  Y  L    F1
       C  E  C  G  *  G  L  F  I  I  K  *  S  P  V  I  K  R  I  L   F2
        V  S  V  V  E  A  Y  S  *  S  N  S  R  L  *  L  N  V  S  Y  F3
2461 TTGTGAGTGTGGTTGAGGCCTATTCATAATCAAATAGTCGCCTGTAATTAAACGTATCTT 2520
     ----:----|----:----|----:----|----:----|----:----|----:----|

      I  A  V  G  V  F  I  N  E  M  D  H  L  L  Y  I  H  C  F  C    F1
       S  P  L  A  S  L  S  M  K  W  T  I  C  C  T  F  I  V  F  V   F2
        R  R  W  R  L  Y  Q  *  N  G  P  F  V  V  H  S  L  F  L  L  F3
2521 ATCGCCGTTGGCGTCTTTATCAATGAAATGGACCATTTGTTGTACATTCATTGTTTTTGT 2580
     ----:----|----:----|----:----|----:----|----:----|----:----|

      C  C  L  C  V  V  I  V  T  *  Y  F  I  K  W  Y  F  T  L  L    F1
       V  V  C  V  L  L  L  *  L  D  I  L  L  N  G  I  S  R  S  F   F2
        L  F  V  C  C  Y  C  N  L  I  F  Y  *  M  V  F  H  A  P  F  F3
2581 TGTTGTTTGTGTGTTGTTATTGTAACTTGATATTTATTAAATGGTATTTCACGCTCCTT 2640
     ----:----|----:----|----:----|----:----|----:----|----:----|

      F  F  T  P  G  L  L  A  A  S  L  L  L  F  V  L  F  L  F  V    F1
       F  S  R  P  D  C  W  L  R  R  F  C  C  L  F  C  F  C  L  F   F2
        F  H  A  R  T  V  G  C  V  A  F  V  V  C  F  V  F  V  C  F  F3
2641 TTTTTCACGCCCGGACTGTTGGCTGCGTCGCTTTTGTTGTTTGTTTTGTTTTTGTTTGTT 2700
     ----:----|----:----|----:----|----:----|----:----|----:----|

      L  L  G  P  R  I  K  L  A  K  *  Q  P  K  L  N  I  Y  V  C    F1
       C  L  A  R  G  L  S  W  P  N  S  S  P  N  *  I  F  M  C  V   F2
```

```
      A  W  P  E  D  *  A  G  Q  I  A  A  Q  I  E  Y  L  C  V  C   F3
2701 TTGCTTGGCCCGAGGATTAAGCTGGCCAAATAGCAGCCCAAATTGAATATTTATGTGTGT 2760
     ----:----|----:----|----:----|----:----|----:----|----:----|

     V  C  R  *  Q  T  H  T  H  T  H  T  P  A  L  S  H  V  H  A    F1
      C  V  D  N  K  H  T  H  T  H  T  H  P  H  S  H  M  C  M  Q   F2
       V  S  I  T  N  T  H  T  H  T  H  T  R  T  L  T  C  A  C  K  F3
2761 GTGTGTCGATAACAAACACACACACACACACACACCCGCACTCTCACATGTGCATGCA 2820
     ----:----|----:----|----:----|----:----|----:----|----:----|

     N  K  S  K  T  C  D  K  T  A  *  L  T  H  V  *  S  T  C  Q    F1
      T  K  A  K  R  A  I  K  R  H  N  *  L  M  F  D  Q  H  V  N   F2
       Q  K  Q  N  V  R  *  N  G  I  I  N  S  C  L  I  N  M  S  I  F3
2821 AACAAAAGCAAAACGTGCGATAAAACGGCATAATTAACTCATGTTTGATCAACATGTCAA 2880
     ----:----|----:----|----:----|----:----|----:----|----:----|

     L  T  V  S  Y  *  L  Y  S  Y  Y  I  Q  L  S  I  V  Y  S  K    F1
      *  Q  L  A  I  N  Y  I  H  I  T  Y  S  Y  L  L  S  T  A  K   F2
       N  S  *  L  L  I  I  F  I  L  H  T  V  I  Y  C  L  Q  Q  K  F3
2881 TTAACAGTTAGCTATTAATTATATTCATATTACATACAGTTATCTATTGTCTACAGCAAA 2940
     ----:----|----:----|----:----|----:----|----:----|----:----|

     K  N  R  K  N  N  R  I  *  *  *  *  Q  Y  L  F  A  A  Y  T    F1
      K  T  E  K  T  I  E  Y  N  N  N  N  N  I  Y  L  L  P  I  P   F2
       K  Q  K  K  Q  *  N  I  I  I  I  I  T  I  S  I  C  C  L  Y  P  F3
2941 AAAAACAGAAAAAAACAATAGAATATAATAATAATAACAATATCTATTTGCTGCCTATACC 3000
     ----:----|----:----|----:----|----:----|----:----|----:----|

     Q  Y  N  I  V  A  T  N  C  G  A  Y  A  R  R  N  M  N  I  K    F1
      N  T  I  *  *  P  R  I  A  A  H  M  P  V  V  I  *  T  L  N   F2
       I  Q  Y  S  S  H  E  L  R  R  I  C  P  S  *  Y  E  H  *  I  F3
3001 CAATACAATATAGTAGCCACGAATTGCGGCGCATATGCCCGTCGTAATATGAACATTAAA 3060
     ----:----|----:----|----:----|----:----|----:----|----:----|

     Y  R  *  S  I  A  A  L  I  D  S  C  L  C  L  I  A  E  P  L    F1
      I  D  D  L  L  R  H  *  L  T  A  V  Y  A  L  S  Q  S  H  L   F2
       *  M  I  Y  C  G  I  D  *  Q  L  F  M  P  Y  R  R  A  T  *  F3
3061 TATAGATGATCTATTGCGGCATTGATTGACAGCTGTTTATGCCTTATCGCAGAGCCACTT 3120
     ----:----|----:----|----:----|----:----|----:----|----:----|

     E  L  G  Q  F  S  F  C  L  W  M  K  R  F  L  T  H  S  T  T    F1
      S  W  V  S  S  L  F  V  Y  G  *  K  D  F  *  H  T  Q  Q  Q   F2
       V  G  S  V  L  F  L  F  M  D  E  K  I  F  D  T  L  N  N  K  F3
3121 GAGTTGGGTCAGTTCTCTTTTTGTTTATGGATGAAAAGATTTTTGACACACTCAACAACA 3180
     ----:----|----:----|----:----|----:----|----:----|----:----|

     K  S  H  *  K  S  Q  Q  S  Q  L  V  A  V  V  V  V  V  E  L    F1
      N  R  I  K  N  H  N  R  A  N  L  L  L  L  L  L  L  L  S  *   F2
       I  A  L  K  I  T  T  E  P  T  C  C  C  C  C  C  C  *  V  K  F3
3181 AAATCGCATTAAAAATCACAACAGAGCCAACTTGTTGCTGTTGTTGTTGTTGTTGAGTTA 3240
     ----:----|----:----|----:----|----:----|----:----|----:----|

     R  T  *  K  S  H  P  N  T  T  N  *  I  F  N  *  R  T  P  T    F1
      E  H  E  N  P  I  Q  I  Q  L  I  K  F  S  I  S  A  R  P  Q   F2
       N  M  K  I  P  S  K  Y  N  *  L  N  F  Q  L  A  H  A  H  K  F3
3241 AGAACATGAAAATCCCATCCAAATACAACTAATTAAATTTTCAATTAGCGCACGCCCACA 3300
     ----:----|----:----|----:----|----:----|----:----|----:----|

     N  F  N  C  L  A  L  A  *  G  Q  I  N  W  H  K  S  P  W  P    F1
      I  S  T  A  L  L  W  P  E  G  K  S  I  G  T  N  L  L  G  Q   F2
       F  Q  L  P  C  F  G  L  R  A  N  Q  L  A  Q  I  S  L  A  N  F3
3301 AATTTCAACTGCCTTGCTTTGGCCTGAGGGCAAATCAATTGGCACAAATCTCCTTGGCCA 3360
     ----:----|----:----|----:----|----:----|----:----|----:----|
```

```
     T  F  W  G  I  V  *  L  L  V  T  T  T  R  E  A  I  N  N  R     F1
      H  F  G  A  L  F  S  C  L  *  Q  Q  Q  E  R  P  *  I  I  V    F2
       I  L  G  H  C  L  A  A  C  N  N  N  K  R  G  H  K  *  S  F   F3
3361 ACATTTTGGGGCATTGTTTAGCTGCTTGTAACAACAACAAGAGAGGCCATAAATAATCGT 3420
     ----:----|----:----|----:----|----:----|----:----|----:----|

     S  W  Y  A  F  L  *  S  F  *  *  Y  F  I  L  T  M  R  C  D     F1
      L  G  M  H  F  C  R  V  F  N  S  I  L  Y  *  Q  C  A  A  T    F2
       L  V  C  I  S  V  E  F  L  I  V  F  Y  I  N  N  A  L  R  L   F3
3421 TCTTGGTATGCATTTCTGTAGAGTTTTTAATAGTATTTTATATTAACAATGCGCTGCGAC 3480
     ----:----|----:----|----:----|----:----|----:----|----:----|

     C  A  V  N  K  I  V  N  K  K  R  A  *  G  I  F  F  F  Y  D     F1
      V  L  *  T  K  L  *  T  K  S  G  P  E  E  Y  F  F  F  T  T    F2
       C  C  K  Q  N  C  K  Q  K  A  G  L  R  N  I  F  F  L  R  Q   F3
3481 TGTGCTGTAAACAAAATTGTAAACAAAAAGCGGGCCTGAGGAATATTTTTTTTTTACGAC 3540
     ----:----|----:----|----:----|----:----|----:----|----:----|

     N  S  L  *  P  N  *  A  D  Q  G  Q  V  G  *  A  V  G  G  G     F1
      I  V  Y  D  P  T  E  Q  T  R  A  R  W  G  E  R  W  A  V  G    F2
       *  F  M  T  Q  L  S  R  P  G  P  G  G  V  S  G  G  R  W  A   F3
3541 AATAGTTTATGACCCAACTGAGCAGACCAGGGCCAGGTGGGGTGAGCGGTGGGCGGTGGG 3600
     ----:----|----:----|----:----|----:----|----:----|----:----|

     R  V  P  K  *  *  V  R  H  V  K  R  Q  I  C  Y  P  P  V  T     F1
      G  C  P  N  S  E  C  V  M  *  R  G  K  Y  V  I  R  L  S  Q    F2
       G  A  Q  I  V  S  A  S  C  E  E  A  N  M  L  S  A  C  H  R   F3
3601 CGGGTGCCCAAATAGTGAGTGCGTCATGTGAAGAGGCAAATATGTTATCCGCCTGTCACA 3660
     ----:----|----:----|----:----|----:----|----:----|----:----|

     D  S  V  Q  G  S  Q  I  V  R  F  S  Y  A  S  Y  L  H  V  F     F1
      T  V  F  R  A  R  R  *  C  V  L  V  M  P  A  I  C  T  C  F    F2
       Q  C  S  G  L  A  D  S  A  F  *  L  C  Q  L  F  A  R  V  L   F3
3661 GACAGTGTTCAGGGCTCGCAGATAGTGCGTTTTAGTTATGCCAGCTATTTGCACGTGTTT 3720
     ----:----|----:----|----:----|----:----|----:----|----:----|

     C  F  S  Y  T  L  L  L  L  F  G  L  F  M  N  V  K  A  M  K     F1
      A  F  H  T  L  S  S  C  Y  L  V  Y  L  *  T  S  K  P  *  K    F2
       L  F  I  H  S  P  P  V  I  W  F  I  Y  E  R  Q  S  H  E  N   F3
3721 TGCTTTTCATACACTCTCCTCCTGTTATTTGGTTTATTTATGAACGTCAAAGCCATGAAA 3780
     ----:----|----:----|----:----|----:----|----:----|----:----|

     M  S  D  H  M  T  Y  A  *  Y  C  S  Y  I  Y  I  F  A  *  T     F1
      C  Q  T  T  *  R  M  R  N  T  V  V  I  Y  I  Y  L  H  R  H    F2
       V  R  P  H  D  V  C  V  I  L  *  L  Y  I  Y  I  C  I  D  I   F3
3781 ATGTCAGACCACATGACGTATGCGTAATACTGTAGTTATATATATATATTTGCATAGACA 3840
     ----:----|----:----|----:----|----:----|----:----|----:----|

     Y  V  C  L  F  W  T  G  R  L  *  *  P  D  E  R  H  L  R  H     F1
      M  Y  V  C  F  G  Q  V  D  Y  D  N  Q  T  N  V  T  Y  A  M    F2
       C  M  F  V  L  D  R  S  T  M  I  T  R  R  T  S  P  T  P  C   F3
3841 TATGTATGTTTGTTTTGGACAGGTCGACTATGATAACCAGACGAACGTCACCTACGCCAT 3900
     ----:----|----:----|----:----|----:----|----:----|----:----|

     V  P  L  H  R  Q  R  C  P  *  S  E  H  L  S  G  E  L  L  H     F1
      C  R  Y  I  D  N  D  V  L  D  L  S  T  F  Q  V  S  F  F  I    F2
       A  A  T  *  T  T  M  S  L  I  *  A  P  F  R  *  A  S  S  *   F3
3901 GTGCCGCTACATAGACAACGATGTCCTTGATCTGAGCACCTTTCAGGTGAGCTTCTTCAT 3960
     ----:----|----:----|----:----|----:----|----:----|----:----|

     K  L  V  S  A  A  P  D  G  H  Q  W  I  V  C  A  H  D  N  A     F1
      S  S  Y  L  L  P  L  M  V  I  S  G  L  Y  V  R  M  I  M  R    F2
```

```
       A  R  I  C  C  P  *  W  S  S  V  D  C  M  C  A  *  *  C  G    F3
3961 AAGCTCGTATCTGCTGCCCCTGATGGTCATCAGTGGATTGTATGTGCGCATGATAATGCG 4020
     ----:----|----:----|----:----|----:----|----:----|----:----|

     A  L  A  S  G  H  R  R  A  H  V  Q  G  V  A  A  W  S  E  A     F1
      L  W  H  Q  G  T  G  V  R  M  S  K  E  S  Q  R  G  R  K  R    F2
       S  G  I  R  A  Q  A  C  A  C  P  R  S  R  S  V  V  G  S  G   F3
4021 GCTCTGGCATCAGGGCACAGGCGTGCGCATGTCCAAGGAGTCGCAGCGTGGTCGGAAGCG 4080
     ----:----|----:----|----:----|----:----|----:----|----:----|

     G  H  P  S  G  G  C  C  C  H  C  L  R  L  A  L  A  A  R  A     F1
      V  T  R  L  V  V  V  V  V  I  A  F  A  S  L  W  L  P  V  Q    F2
       S  P  V  W  W  L  L  L  S  L  P  S  P  R  S  G  C  P  C  R   F3
4081 GGTCACCCGTCTGGTGGTTGTTGTTGTCATTGCCTTCGCCTCGCTCTGGCTGCCCGTGCA 4140
     ----:----|----:----|----:----|----:----|----:----|----:----|

     G  E  S  *  H  F  S  L  M  K  F  A  Q  S  F  I  P  F  V  F     F1
      V  S  L  N  T  L  A  *  *  S  L  P  N  R  S  F  L  L  Y  S    F2
       *  V  L  T  L  *  L  N  E  V  C  P  I  V  H  S  F  C  I  R   F3
4141 GGTGAGTCTTAACACTTTAGCTTAATGAAGTTTGCCCAATCGTTCATTCCTTTTGTATTC 4200
     ----:----|----:----|----:----|----:----|----:----|----:----|

     A  A  N  I  T  A  E  G  A  G  H  V  *  N  K  Q  H  V  *  C     F1
      Q  L  I  L  L  L  K  A  L  D  M  Y  E  I  N  S  M  F  N  V    F2
       S  *  Y  Y  C  *  R  R  W  T  C  M  K  *  T  A  C  L  M  S   F3
4201 GCAGCTAATATTACTGCTGAAGGCGCTGGACATGTATGAAATAAACAGCATGTTTAATGT 4260
     ----:----|----:----|----:----|----:----|----:----|----:----|

     H  T  A  D  S  G  P  Y  H  G  V  H  Q  L  L  H  Q  S  A  A     F1
      I  L  Q  I  V  A  H  T  M  A  Y  T  S  S  C  I  N  P  L  L    F2
       Y  C  R  *  W  P  I  P  W  R  T  P  A  P  A  S  I  R  C  S   F3
4261 CATACTGCAGATAGTGGCCCATACCATGGCGTACACCAGCTCCTGCATCAATCCGCTGCT 4320
     ----:----|----:----|----:----|----:----|----:----|----:----|

     L  R  L  S  L  R  Q  F  S  Q  G  L  L  Q  G  E  C  K  D  Q     F1
      Y  A  F  L  S  D  N  F  R  K  A  F  Y  K  V  S  A  K  T  R    F2
       T  P  F  S  Q  T  I  F  A  R  P  S  T  R  *  V  Q  R  P  G   F3
4321 CTACGCCTTTCTCTCAGACAATTTTCGCAAGGCCTTCTACAAGGTGAGTGCAAAGACCAG 4380
     ----:----|----:----|----:----|----:----|----:----|----:----|

     A  Y  Q  G  C  Q  A  I  V  K  L  F  P  S  Y  C  A  V  S  A     F1
      L  T  K  A  V  K  L  L  L  S  S  S  R  R  I  V  P  *  V  L    F2
       L  P  R  L  S  S  Y  C  *  A  L  P  V  V  L  C  R  E  C  W   F3
4381 GCTTACCAAGGCTGTCAAGCTATTGTTAAGCTCTTCCCGTCGTATTGTGCCGTGAGTGCT 4440
     ----:----|----:----|----:----|----:----|----:----|----:----|

     G  S  I  P  C  L  L  P  I  E  L  F  G  S  S  L  L  F  F  L     F1
      G  Q  Y  L  A  C  C  Q  L  S  C  L  A  L  L  F  Y  F  F  C    F2
       V  N  T  L  L  V  A  N  *  V  V  W  L  F  S  F  I  F  F  V   F3
4441 GGGTCAATACCTTGCTTGTTGCCAATTGAGTTGTTTGGCTCTTCTCTTTTATTTTTTTTG 4500
     ----:----|----:----|----:----|----:----|----:----|----:----|

     F  S  F  S  L  S  C  I  F  H  M  A  L  L  S  G  Y  P  R  P     F1
      F  R  S  L  C  H  A  F  F  I  W  L  Y  *  V  A  T  L  D  P    F2
       F  V  L  F  V  M  H  F  S  Y  G  S  I  K  W  L  P  S  T  Q   F3
4501 TTTTCGTTCTCTTTGTCATGCATTTTTCATATGGCTCTATTAAGTGGCTACCCTCGACCC 4560
     ----:----|----:----|----:----|----:----|----:----|----:----|

     N  L  W  I  Y  T  D  F  K  S  I  F  A  A  L  F  S  M  Q  L     F1
      I  C  G  F  I  Q  I  L  K  V  F  S  Q  R  Y  F  P  C  S  C    F2
       F  V  D  L  Y  R  F  *  K  Y  F  R  S  V  I  F  H  A  A  V   F3
4561 AATTTGTGGATTTATACAGATTTTAAAAGTATTTTCGCAGCGTTATTTTCCATGCAGCTG 4620
     ----:----|----:----|----:----|----:----|----:----|----:----|
```

```
     * K R Q V F D L I S N V L L L I L S N V S    F1
      K N V R Y L I * S Q M C F * F F Q M F P     F2
       K T S G I * F D L K C A F N S F K C F H    F3
4621 TAAAAACGTCAGGTATTTGATTTGATCTCAAATGTGCTTTTAATTCTTTCAAATGTTTCC 4680
     ----:----|----:----|----:----|----:----|----:----|----:----|

     I D N S I K L F T L P T V R H G K K K S      F1
      S I I Q L N Y L R C Q Q S D M E K K N H     F2
       R * F N * I I Y A A N S Q T W K K K I T    F3
4681 ATCGATAATTCAATTAAATTATTTACGCTGCCAACAGTCAGACATGGAAAAAAAAAATCA 4740
     ----:----|----:----|----:----|----:----|----:----|----:----|

     H K I I L E K R A Q C P N I C Q Q * A A      F1
      I K * Y * K N A R N A Q T F A S S R R L     F2
       * N N I R K T R A M P K H L P A V G G C    F3
4741 CATAAAATAATATTAGAAAAACGCGCGCAATGCCCAAACATTTGCCAGCAGTAGGCGGCT 4800
     ----:----|----:----|----:----|----:----|----:----|----:----|

     A L M S * V R K T S I R R V G R R C A Q      F1
      P * * V K C E K R A Y A A W D G D A P N     F2
       L N E L S A K N E H T P R G T E M R P T    F3
4801 GCCTTAATGAGTTAAGTGCGAAAAACGAGCATACGCCGCGTGGGACGGAGATGCGCCCAA 4860
     ----:----|----:----|----:----|----:----|----:----|----:----|

     Q L R Y K Y Q H F T A K L * N E P A S P      F1
      N C V I N T N I L Q Q N C E T S Q P A Q     F2
       I A L * I P T F Y S K I V K R A S Q P R    F3
4861 CAATTGCGTTATAAATACCAACATTTTACAGCAAAATTGTGAAACGAGCCAGCCAGCCCA 4920
     ----:----|----:----|----:----|----:----|----:----|----:----|

     G N Q I Q S N R N F Y F W Y L C E M N A      F1
      A I K F N Q I G I S I F G T C A R * M L     F2
       Q S N S I K S E F L F L V L V R D E C L    F3
4921 GGCAATCAAATTCAATCAAATCGGAATTTCTATTTTTGGTACTTGTGCGAGATGAATGCT 4980
     ----:----|----:----|----:----|----:----|----:----|----:----|

     W H Y L T Q R K C * N M E F * T S S K H      F1
      G I I L H S E N V E I W N F K Q V V S T     F2
       A L S Y T A K M L K Y G I L N K * * A H    F3
4981 TGGCATTATCTTACACAGCGAAAATGTTGAAATATGGAATTTTAAACAAGTAGTAAGCAC 5040
     ----:----|----:----|----:----|----:----|----:----|----:----|

     I A S N W R W L G T S S T S C S I * L L      F1
      L H Q I G A G L E Q A A Q A V A F S Y *     F2
       C I K L A L A W N K Q H K L * H L V I K    F3
5041 ATTGCATCAAATTGGCGCTGGCTTGGAACAAGCAGCACAAGCTGTAGCATTTAGTTATTA 5100
     ----:----|----:----|----:----|----:----|----:----|----:----|

     K H F F L C K S Q K V R * H Q Q Q A H V      F1
      S I F F C A N R K R F V D T N N K P T W     F2
       A F F F V Q I A K G S L T P T T S P R G    F3
5101 AAGCATTTTTTTTTGTGCAAATCGCAAAAGGTTCGTTGACACCAACAACAAGCCCACGTG 5160
     ----:----|----:----|----:----|----:----|----:----|----:----|

     A T L C R T L P D P G I H I M C R A P Q      F1
      P R S A G P C R T R E Y T L C A G H R S     F2
       H A L Q D P A G P G N T H Y V P G T A A    F3
5161 GCCACGCTCTGCAGGACCCTGCCGGACCCGGGAATACACATTATGTGCCGGGCACCGCAG 5220
     ----:----|----:----|----:----|----:----|----:----|----:----|

     L I P I Q L T C C A H * F F C S A A I Y      F1
      S F Q F N S L A A H I N S F A R R P F T     F2
```

```
       H  S  N  S  T  H  L  L  R  T  L  I  L  L  L  G  G  H  L  H    F3
5221 CTCATTCCAATTCAACTCACTTGCTGCGCACATTAATTCTTTTGCTCGGCGGCCATTTAC 5280
     ----:----|----:----|----:----|----:----|----:----|----:----|

     I  S  T  K  R  K  E  K  I  K  I  K  P  N  S  W  H  S  V  G      F1
      *  V  Q  N  V  R  K  K  S  K  *  N  Q  I  R  G  I  P  W  A     F2
       K  Y  K  T  *  G  K  N  Q  N  K  T  K  F  V  A  F  R  G  H    F3
5281 ATAAGTACAAAACGTAAGGAAAAAATCAAAATAAAACCAAATTCGTGGCATTCCGTGGGC 5340
     ----:----|----:----|----:----|----:----|----:----|----:----|

     T  R  G  R  R  S  A  P  L  C  F  T  Y  F  *  L  L  *  G  Q      F1
      L  E  G  G  D  R  L  H  Y  A  L  L  T  F  N  Y  F  R  D  R     F2
       S  R  A  E  I  G  S  T  M  L  Y  L  L  L  I  T  L  G  T  D    F3
5341 ACTCGAGGGCGGAGATCGGCTCCACTATGCTTTACTTACTTTTAATTACTTTAGGGACAG 5400
     ----:----|----:----|----:----|----:----|----:----|----:----|

     T  M  G  A  A  D  P  L  T  R  V  G  V  W  S  R  V  G  A  I      F1
      Q  W  V  R  Q  T  H  *  L  E  W  V  S  G  V  E  L  V  P  S     F2
       N  G  C  G  R  P  I  D  *  S  G  C  L  E  S  S  W  C  H  R    F3
5401 ACAATGGGTGCGGCAGACCCATTGACTAGAGTGGGTGTCTGGAGTCGAGTTGGTGCCATC 5460
     ----:----|----:----|----:----|----:----|----:----|----:----|

     D  A  K  A  V  C  N  I  S  F  Q  S  F  C  G  L  I  L  I  F      F1
      M  P  K  P  S  V  T  F  P  F  N  H  F  V  A  *  F  *  Y  F     F2
       C  Q  S  R  L  *  H  F  L  S  I  I  L  W  P  N  F  N  I  L    F3
5461 GATGCCAAAGCCGTCTGTAACATTTCCTTTCAATCATTTTGTGGCCTAATTTTAATATTT 5520
     ----:----|----:----|----:----|----:----|----:----|----:----|

     *  F  V  L  L  *  M  F  L  V  L  L  L  L  L  L  L  L  A  Q      F1
      N  L  F  C  C  E  C  F  W  C  C  C  C  C  C  C  C  W  R  S     F2
       I  C  F  V  V  N  V  F  G  A  V  V  A  V  A  A  V  G  A  V    F3
5521 TAATTTGTTTTGTTGTGAATGTTTTTGGTGCTGTTGTTGCTGTTGCTGCTGTTGGCGCAG 5580
     ----:----|----:----|----:----|----:----|----:----|----:----|

     F  F  P  L  C  A  K  S  C  F  G  T  L  H  I  N  *  A  W  P      F1
      F  F  H  F  A  Q  S  P  A  S  A  L  C  I  *  I  E  P  G  H     F2
       F  S  T  L  R  K  V  L  L  R  H  F  A  Y  K  L  S  L  A  I    F3
5581 TTTTTTCCACTTTGCGCAAAGTCCTGCTTCGGCACTTTGCATATAAATTGAGCCTGGCCA 5640
     ----:----|----:----|----:----|----:----|----:----|----:----|

     F  S  M  R  P  A  P  C  A  P  R  A  A  L  I  D  V  F  V  K      F1
      S  P  C  G  L  R  P  A  P  L  A  R  R  *  L  M  C  L  S  S     F2
       L  H  A  A  C  A  L  R  P  S  R  G  A  N  *  C  V  C  Q  V    F3
5641 TTCTCCATGCGGCCTGCGCCCTGCGCCCCTCGCGCGGCGCTAATTGATGTGTTTGTCAAG 5700
     ----:----|----:----|----:----|----:----|----:----|----:----|

     S  L  N  *  I  N  Y  I  N  *  I  A  A  L  A  L  A  I  V  L      F1
      H  *  I  R  L  I  T  L  I  K  L  L  R  W  R  W  L  L  C  W     F2
       I  K  L  D  *  L  H  *  L  N  C  C  V  G  A  G  Y  C  V  G    F3
5701 TCATTAAATTAGATTAATTACATTAATTAAATTGCTGCGTTGGCGCTGGCTATTGTGTTG 5760
     ----:----|----:----|----:----|----:----|----:----|----:----|

     V  Y  C  A  R  R  M  W  F  Y  F  C  F  P  F  F  F  L  L  S      F1
      F  T  V  R  G  A  C  G  F  I  F  V  F  H  F  F  F  C  Y  H     F2
       L  L  C  E  A  H  V  V  L  F  L  F  S  I  F  F  F  V  I  T    F3
5761 GTTTACTGTGCGAGGCGCATGTGGTTTTATTTTTGTTTTCCATTTTTTTTTTTGTTATCA 5820
     ----:----|----:----|----:----|----:----|----:----|----:----|

     L  F  V  A  L  N  P  C  P  D  R  Y  *  T  H  L  L  A  Y  Q      F1
      F  L  *  L  *  I  L  V  Q  T  G  I  E  R  T  Y  *  P  I  K     F2
       F  C  S  F  K  S  L  S  R  Q  V  L  N  A  P  I  S  L  S  N    F3
5821 CTTTTTGTAGCTTTAAATCCTTGTCCAGACAGGTATTGAACGCACCTATTAGCCTATCAA 5880
     ----:----|----:----|----:----|----:----|----:----|----:----|
```

M G * L L G * Q L V H Y A I S Q K R V N F1
W A N C S A D S S S I M Q L V K S V S M F2
G L I A R L T A R P L C N * S K A C Q W F3
5881 ATGGGCTAATTGCTCGGCTGACAGCTCGTCCATTATGCAATTAGTCAAAAGCGTGTCAAT 5940
----:----|----:----|----:----|----:----|----:----|----:----|

G F F F F F F F Y C F R I R C V E S A S F1
V F F F F F F F F I V L G F D V W S Q P P F2
F F F F F F F F L L F * D S M C G V S L P F3
5941 GGTTTTTTTTTTTTTTTTTTTTTTTATTGTTTTAGGATTCGATGTGTGGAGTCAGCCTCC 6000
----:----|----:----|----:----|----:----|----:----|----:----|

H R F W P K T D L I Q L S Q I K Y E L K F1
I D F G Q K P I * F N C P R L N M S S N F2
S I L A K N R F N S I V P D * I * A Q I F3
6001 CATCGATTTTGGCCAAAAACCGATTTAATTCAATTGTCCCAGATTAAATATGAGCTCAAA 6060
----:----|----:----|----:----|----:----|----:----|----:----|

F I C V H * M * C P W L Y L L Y S F N * F1
L F V Y I E C D A R G C I Y F I L L I N F2
Y L C T L N V M P V A V F T L F F * L M F3
6061 TTTATTTGTGTACATTGAATGTGATGCCCGTGGCTGTATTTACTTTATTCTTTTAATTAA 6120
----:----|----:----|----:----|----:----|----:----|----:----|

C * L Y * Y L L L L V I T Y * Y * I M * F1
V N F I D I Y F C W * * H I N I K * C N F2
L T L L I F T F A G N N I L I L N N V M F3
6121 TGTTAACTTTATTGATATTTACTTTTGCTGGTAATAACATATTAATATTAAATAATGTAA 6180
----:----|----:----|----:----|----:----|----:----|----:----|

* K I G M N Y L Q F I L N A F H Q K F C F1
E K * A * T I Y N L F * M L F I R N F V F2
K N R H E L F T I Y F K C F S S E I L F F3
6181 TGAAAAATAGGCATGAACTATTTACAATTTATTTTAAATGCTTTTCATCAGAAATTTTGT 6240
----:----|----:----|----:----|----:----|----:----|----:----|

L H F S H I I L I N S M K C Q F S H N Y F1
Y I F P T L S * * I Q * N V N F L T I T F2
T F F P H Y L D K F N E M S I F S Q L H F3
6241 TTACATTTTTCCCACATTATCTTGATAAATTCAATGAAATGTCAATTTTCTCACAATTAC 6300
----:----|----:----|----:----|----:----|----:----|----:----|

M N S D L I Y I S E L I P D L C K A * T F1
* I P I * S I * A N * F P I F A R P K H F2
E F R F N L Y K R T D S R S L Q G L N I F3
6301 ATGAATTCCGATTTAATCTATATAAGCGAACTGATTCCCGATCTTTGCAAGGCCTAAACA 6360
----:----|----:----|----:----|----:----|----:----|----:----|

S F A E M L I R N R E Y D F F F * M P I F1
L L L R C * S G I E N M I F F S E C Q S F2
F C * D A N P E * R I * F F F L N A N Q F3
6361 TCTTTTGCTGAGATGCTAATCCGGAATAGAGAATATGATTTTTTTTTCTGAATGCCAATC 6420
----:----|----:----|----:----|----:----|----:----|----:----|

N S Y D L Y E A I E T F D * K I E F Y L F1
I V M I C T R P * R H L I R K * S F I Y F2
* L * F V R G H R D I * L E N R V L F I F3
6421 AATAGTTATGATTTGTACGAGGCCATAGAGACATTTGATTAGAAAATAGAGTTTTATTTA 6480
----:----|----:----|----:----|----:----|----:----|----:----|

F E * * F K * K F C * S M F S S Q N K R F1
L N N S L N R S S V K V C F P A K I N D F2

```
      * I I V * I E V L L K Y V F Q P K * T T    F3
6481 TTTGAATAATAGTTTAAATAGAAGTTCTGTTAAAGTATGTTTTCCAGCCAAAATAAACGA 6540
     ----:----|----:----|----:----|----:----|----:----|----:----|

     R L L T D S Y D V A K S Y L C S V L C L     F1
      G F * P T P T T W P K V T C A L C C A C    F2
       A S N R L L R R G Q K L P V L C A V P V   F3
6541 CGGCTTCTAACCGACTCCTACGACGTGGCCAAAAGTTACCTGTGCTCTGTGCTGTGCCTG 6600
     ----:----|----:----|----:----|----:----|----:----|----:----|

     C V C D L C I M P R L Q N M C F L K Q L     F1
      V C A I Y V L C L G Y R I C V F * S S C    F2
       C V R S M Y Y A * A T E Y V F F K A V V   F3
6601 TGTGTGTGCGATCTATGTATTATGCCTAGGCTACAGAATATGTGTTTTTTAAAGCAGTTG 6660
     ----:----|----:----|----:----|----:----|----:----|----:----|

     Y N * Y * C K E I C D K A F G S V I I R     F1
      I I N I N A K R Y V I K H L A A * L F D    F2
       * L I L M Q R D M * * S I W Q R N Y S I   F3
6661 TATAATTAATATTAATGCAAAGAGATATGTGATAAAGCATTTGGCAGCGTAATTATTCGA 6720
     ----:----|----:----|----:----|----:----|----:----|----:----|

     * F C S K C C A C I P L H S M R H N Q I     F1
      N F A V N A V H A F R Y I V C A I I K S    F2
       I L Q * M L C M H S V T * Y A P * S N H   F3
6721 TAATTTTGCAGTAAATGCTGTGCATGCATTCCGTTACATAGTATGCGCCATAATCAAATC 6780
     ----:----|----:----|----:----|----:----|----:----|----:----|

     T T K M Q I Q V D A H S H T H T H T H T     F1
      Q P K C K Y R * M H T R T H T H T H T H    F2
       N Q N A N T G R C T L A H T H T H T H I   F3
6781 ACAACCAAAATGCAAATACAGGTAGATGCACACTCGCACACACACACACACACACACACA 6840
     ----:----|----:----|----:----|----:----|----:----|----:----|

     S I D G A Q K P * L I S L E R L H G N A     F1
      R L M G P K N R N L F P L S D C M E M P    F2
       D * W G P K T V T Y F P * A I A W K C Q   F3
6841 TCGATTGATGGGGCCCAAAAACCGTAACTTATTTCCCTTGAGCGATTGCATGGAAATGCC 6900
     ----:----|----:----|----:----|----:----|----:----|----:----|

     R V C A I A F I V V V V D V V V L * R R     F1
      E S V R L L L L L L L L M L L F Y D G A    F2
       S L C D C F Y C C C C * C C C S M T A H   F3
6901 AGAGTCTGTGCGATTGCTTTTATTGTTGTTGTTGTTGATGTTGTTGTTCTATGACGGCGC 6960
     ----:----|----:----|----:----|----:----|----:----|----:----|

     T N Q Q F W T V Q S E R K F I N H L Y A     F1
      L I N N F G R F N Q S A N S * I I Y M Q    F2
       * S T I L D G S I R A Q I H K S F I C K   F3
6961 ACTAATCAACAATTTTGGACGGTTCAATCAGAGCGCAAATTCATAAATCATTTATATGCA 7020
     ----:----|----:----|----:----|----:----|----:----|----:----|

     N A Q R N R R D C A D R G V G E W T H R     F1
      T P N G I E G T A P I E E W G N G P T D    F2
       R P T E * K G L R R * R S G G M D P Q I   F3
7021 AACGCCCAACGGAATAGAAGGGACTGCGCCGATAGAGGAGTGGGGGAATGGACCCACAGA 7080
     ----:----|----:----|----:----|----:----|----:----|----:----|

     * T V G I S T P Q T S * I I E Y * I K S     F1
      K L W A F Q H L K P V K * L N I E S N L    F2
       N C G H F N T S N Q L N N * I L N Q I *   F3
7081 TAAACTGTGGGCATTTCAACACCTCAAACCAGTTAAATAATTGAATATTGAATCAAATCT 7140
     ----:----|----:----|----:----|----:----|----:----|----:----|
```

```
      N  H  S  N  A  K  P  Q  S  I  V  W  A  S  R  L  D  R  G  I     F1
       I  I  Q  T  Q  S  H  S  P  L  C  G  Q  A  G  L  I  G  E  L    F2
        S  F  K  R  K  A  T  V  H  C  V  G  K  Q  A  *  *  G  N  C   F3
7141 AATCATTCAAACGCAAAGCCACAGTCCATTGTGTGGGCAAGCAGGCTTGATAGGGGAATT 7200
     ----:----|----:----|----:----|----:----|----:----|----:----|

      A  K  A  A  T  G  S  I  G  I  A  V  H  P  E  R  I  I  N  Q     F1
       L  R  Q  R  L  V  P  L  A  L  P  C  I  Q  N  V  S  L  I  K    F2
        *  G  S  D  W  F  H  W  H  C  R  A  S  R  T  Y  H  *  S  N   F3
7201 GCTAAGGCAGCGACTGGTTCCATTGGCATTGCCGTGCATCCAGAACGTATCATTAATCAA 7260
     ----:----|----:----|----:----|----:----|----:----|----:----|

      T  F  W  G  P  R  T  W  L  N  L  L  V  V  I  L  C  I  F  S     F1
       H  F  G  A  P  G  P  G  S  I  Y  S  S  L  F  Y  V  F  L  A    F2
        I  L  G  P  P  D  L  A  Q  F  T  R  R  Y  F  M  Y  F  *  H   F3
7261 ACATTTTGGGGCCCCCGGACCTGGCTCAATTTACTCGTCGTTATTTTATGTATTTTTAGC 7320
     ----:----|----:----|----:----|----:----|----:----|----:----|

      I  N  L  C  P  T  F  L  A  F  H  L  S  F  L  R  R  P  L  T     F1
       L  T  Y  A  R  H  F  S  H  F  I  F  R  F  S  V  G  H  *  L    F2
        *  L  M  P  D  I  S  R  I  S  S  F  V  S  P  *  A  I  N  C   F3
7321 ATTAACTTATGCCCGACATTTCTCGCATTTCATCTTTCGTTTCTCCGTAGGCCATTAACT 7380
     ----:----|----:----|----:----|----:----|----:----|----:----|

      A  R  I  A  T  T  I  T  H  L  I  C  R  R  R  A  R  R  P  A     F1
       L  E  S  L  P  Q  L  H  I  *  F  A  A  A  A  Q  D  V  L  R    F2
        S  N  R  Y  H  N  Y  T  S  D  L  P  P  P  R  K  T  S  C  G   F3
7381 GCTCGAATCGCTACCACAATTACACATCTGATTTGCCGCCGCGCAAGACGTCCTGCG 7440
     ----:----|----:----|----:----|----:----|----:----|----:----|

      A  G  P  P  R  R  V  C  K  R  Q  C  D  X                       F1
       Q  D  L  H  D  G  S  V  S  A  N  A  T                         F2
        R  T  S  T  T  G  L  *  A  P  M  R  R                        F3
7441 GCAGGACCTCCACGACGGGTCTGTAAGCGCCAATGCGACG 7480
     ----:----|----:----|----:----|----:----|

    Q  R  P  G  L  P  R  L  S  S  Y  C  *  A  L  P  V  V  L  C     F1
       K  D  Q  A  Y  Q  G  C  Q  A  I  V  K  L  F  P  S  Y  C  A    F2
        K  T  R  L  T  K  A  V  K  L  L  L  S  S  S  R  R  I  V  P   F3
   1 CAAAGACCAGGCTTACCAAGGCTGTCAAGCTATTGTTAAGCTCTTCCCGTCGTATTGTGC 60
     ----:----|----:----|----:----|----:----|----:----|----:----|

      R  E  C  W  V  N  T  L  L  V  A  N  *  V  V  W  L  F  S  F     F1
       V  S  A  G  S  I  P  C  L  L  P  I  E  L  F  G  S  S  L  L    F2
        *  V  L  G  Q  Y  L  A  C  C  Q  L  S  C  L  A  L  L  F  Y   F3
  61 CGTGAGTGCTGGGTCAATACCTTGCTTGTTGCCAATTGAGTTGTTTGGCTCTTCTCTTTT 120
     ----:----|----:----|----:----|----:----|----:----|----:----|

      I  F  F  V  F  V  L  F  V  M  H  F  S  Y  G  S  I  K  W  L     F1
       F  F  L  F  S  F  S  L  S  C  I  F  H  M  A  L  L  S  G  Y    F2
        F  F  C  F  R  S  L  C  H  A  F  F  I  W  L  Y  *  V  A  T   F3
 121 ATTTTTTTGTTTTCGTTCTCTTTGTCATGCATTTTTCATATGGCTCTATTAAGTGGCTA 180
     ----:----|----:----|----:----|----:----|----:----|----:----|

      P  S  T  Q  F  V  D  L  Y  R  F  *  K  Y  F  R  S  V  I  F     F1
       P  R  P  N  L  W  I  Y  T  D  F  K  S  I  F  A  A  L  F  S    F2
        L  D  P  I  C  G  F  I  Q  I  L  K  V  F  S  Q  R  Y  F  P   F3
 181 CCCTCGACCCAATTTGTGGATTTATACAGATTTTAAAAGTATTTTCGCAGCGTTATTTTC 240
     ----:----|----:----|----:----|----:----|----:----|----:----|

      H  A  A  V  K  T  S  G  I  *  F  D  L  K  C  A  F  N  S  F     F1
       M  Q  L  *  K  R  Q  V  F  D  L  I  S  N  V  L  L  I  L  S    F2
```

```
     C  S  C  K  N  V  R  Y  L  I  *  S  Q  M  C  F  *  F  F  Q   F3
 241 CATGCAGCTGTAAAAACGTCAGGTATTTGATTTGATCTCAAATGTGCTTTTAATTCTTTC 300
     ----:----|----:----|----:----|----:----|----:----|----:----|

     K  C  F  H  R  *  F  N  *  I  I  Y  A  A  N  S  Q  T  W  K    F1
      N  V  S  I  D  N  S  I  K  L  F  T  L  P  T  V  R  H  G  K   F2
       M  F  P  S  I  I  Q  L  N  Y  L  R  C  Q  Q  S  D  M  E  K  F3
nrlgmwttth qdvssektty

301 AAATGTTTCCATCGATAATTCAATTAAATTATTTACGCTGCCAACAGTCAGACATGGAAA 360
     ----:----|----:----|----:----|----:----|----:----|----:----|

     K  K  I  T  *  N  N  I  R  K  T  R  A  M  P  K  H  L  P  A    F1
      K  K  S  H  K  I  I  L  E  K  R  A  Q  C  P  N  I  C  Q  Q   F2
       K  N  H  I  K  *  Y  *  K  N  A  R  N  A  Q  T  F  A  S  S  F3
 361 AAAAAAATCACATAAAATAATATTAGAAAAACGCGCGCAATGCCCAAACATTTGCCAGCA 420
     ----:----|----:----|----:----|----:----|----:----|----:----|

     V  G  G  C  L  N  E  L  S  A  K  N  E  H  T  P  R  G  T  E    F1
      *  A  A  A  L  M  S  *  V  R  K  T  S  I  R  R  V  G  R  R   F2
       R  R  L  P  *  *  V  K  C  E  K  R  A  Y  A  A  W  D  G  D  F3
 421 GTAGGCGGCTGCCTTAATGAGTTAAGTGCGAAAAACGAGCATACGCCGCGTGGGACGGAG 480
     ----:----|----:----|----:----|----:----|----:----|----:----|

     M  R  P  T  I  A  L  *  I  P  T  F  Y  S  K  I  V  K  R  A    F1
      C  A  Q  Q  L  R  Y  K  Y  Q  H  F  T  A  K  L  *  N  E  P   F2
       A  P  N  N  C  V  I  N  T  N  I  L  Q  Q  N  C  E  T  S  Q  F3
 481 ATGCGCCCAACAATTGCGTTATAAATACCAACATTTTACAGCAAAATTGTGAAACGAGCC 540
     ----:----|----:----|----:----|----:----|----:----|----:----|

     S  Q  P  R  Q  S  N  S  I  K  S  E  F  L  F  L  V  L  V  R    F1
      A  S  P  G  N  Q  I  Q  S  N  R  N  F  Y  F  W  Y  L  C  E   F2
       P  A  Q  A  I  K  F  N  Q  I  G  I  S  I  F  G  T  C  A  R  F3
 541 AGCCAGCCCAGGCAATCAAATTCAATCAAATCGGAATTTCTATTTTTGGTACTTGTGCGA 600
     ----:----|----:----|----:----|----:----|----:----|----:----|

     D  E  C  L  A  L  S  Y  T  A  K  M  L  K  Y  G  I  L  N  K    F1
      M  N  A  W  H  Y  L  T  Q  R  K  C  *  N  M  E  F  *  T  S   F2
       *  M  L  G  I  I  L  H  S  E  N  V  E  I  W  N  F  K  Q  V  F3
 601 GATGAATGCTTGGCATTATCTTACACAGCGAAAATGTTGAAATATGGAATTTTAAACAAG 660
     ----:----|----:----|----:----|----:----|----:----|----:----|

     *  *  A  H  C  I  K  L  A  L  A  W  N  K  Q  H  K  L  *  H    F1
      S  K  H  I  A  S  N  W  R  W  L  G  T  S  S  T  S  C  S  I   F2
       V  S  T  L  H  Q  I  G  A  G  L  E  Q  A  A  Q  A  V  A  F  F3
 661 TAGTAAGCACATTGCATCAAATTGGCGCTGGCTTGGAACAAGCAGCACAAGCTGTAGCAT 720
     ----:----|----:----|----:----|----:----|----:----|----:----|

     L  V  I  K  A  F  F  F  V  Q  I  A  K  G  S  L  T  P  T  T    F1
      *  L  L  K  H  F  F  L  C  K  S  Q  K  V  R  *  H  Q  Q  Q   F2
       S  Y  *  S  I  F  F  C  A  N  R  K  R  F  V  D  T  N  N  K  F3
 721 TTAGTTATTAAAGCATTTTTTTTTGTGCAAATCGCAAAAGGTTCGTTGACACCAACAACA 780
     ----:----|----:----|----:----|----:----|----:----|----:----|

     S  P  R  G  H  A  L  Q  D  P  A  G  P  G  N  T  H  Y  V  P    F1
      A  H  V  A  T  L  C  R  T  L  P  D  P  G  I  H  I  M  C  R   F2
       P  T  W  P  R  S  A  G  P  C  R  T  R  E  Y  T  L  C  A  G  F3
 781 AGCCCACGTGGCCACGCTCTGCAGGACCCTGCCGGACCCGGGAATACACATTATGTGCCG 840
     ----:----|----:----|----:----|----:----|----:----|----:----|

     G  T  A  A  H  S  N  S  T  H  L  L  R  T  L  I  L  L  L  G    F1
      A  P  Q  L  I  P  I  Q  L  T  C  C  A  H  *  F  F  C  S  A   F2
       H  R  S  S  F  Q  F  N  S  L  A  A  H  I  N  S  F  A  R  R  F3
```

```
 841 GGCACCGCAGCTCATTCCAATTCAACTCACTTGCTGCGCACATTAATTCTTTTGCTCGGC 900
     ----:----|----:----|----:----|----:----|----:----|----:----|

     G  H  L  H  K  Y  K  T  *  G  K  N  Q  N  K  T  K  F  V  A    F1
      A  I  Y  I  S  T  K  R  K  E  K  I  K  I  K  P  N  S  W  H   F2
       P  F  T  *  V  Q  N  V  R  K  K  S  K  *  N  Q  I  R  G  I  F3
 901 GGCCATTTACATAAGTACAAAACGTAAGGAAAAAATCAAAATAAAACCAAATTCGTGGCA 960

     F  R  G  H  S  R  A  E  I  G  S  T  M  L  Y  L  L  L  I  T    F1
      S  V  G  T  R  G  R  R  S  A  P  L  C  F  T  Y  F  *  L  L   F2
       P  W  A  L  E  G  G  D  R  L  H  Y  A  L  L  T  F  N  Y  F  F3
 961 TTCCGTGGGCACTCGAGGGCGGAGATCGGCTCCACTATGCTTTACTTACTTTTAATTACT 1020
     ----:----|----:----|----:----|----:----|----:----|----:----|

     L  G  T  D  N  G  C  G  R  P  I  D  *  S  G  C  L  E  S  S    F1
      *  G  Q  T  M  G  A  A  D  P  L  T  R  V  G  V  W  S  R  V   F2
       R  D  R  Q  W  V  R  Q  T  H  *  L  E  W  V  S  G  V  E  L  F3
1021 TTAGGGACAGACAATGGGTGCGGCAGACCCATTGACTAGAGTGGGTGTCTGGAGTCGAGT 1080
     ----:----|----:----|----:----|----:----|----:----|----:----|

     W  C  H  R  C  Q  S  R  L  *  H  F  L  S  I  I  L  W  P  N    F1
      G  A  I  D  A  K  A  V  C  N  I  S  F  Q  S  F  C  G  L  I   F2
       V  P  S  M  P  K  P  S  V  T  F  P  F  N  H  F  V  A  *  F  F3
1081 TGGTGCCATCGATGCCAAAGCCGTCTGTAACATTTCCTTTCAATCATTTTGTGGCCTAAT 1140
     ----:----|----:----|----:----|----:----|----:----|----:----|

     F  N  I  L  I  C  F  V  V  N  V  F  G  A  V  V  A  V  A  A    F1
      L  I  F  *  F  V  L  L  *  M  F  L  V  L  L  L  L  L  L  L   F2
       *  Y  F  N  L  F  C  C  E  C  F  W  C  C  C  C  C  C  C  C  F3
1141 TTTAATATTTTAATTTGTTTTGTTGTGAATGTTTTTGGTGCTGTTGTTGCTGTTGCTGCT 1200
     ----:----|----:----|----:----|----:----|----:----|----:----|

     V  G  A  V  F  S  T  L  R  K  V  L  L  R  H  F  A  Y  K  L    F1
      L  A  Q  F  F  P  L  C  A  K  S  C  F  G  T  L  H  I  N  *   F2
       W  R  S  F  F  H  F  A  Q  S  P  A  S  A  L  C  I  *  I  E  F3
1201 GTTGGCGCAGTTTTTTCCACTTTGCGCAAAGTCCTGCTTCGGCACTTTGCATATAAATTG 1260
     ----:----|----:----|----:----|----:----|----:----|----:----|

     S  L  A  I  L  H  A  A  C  A  L  R  P  S  R  G  A  N  *  C    F1
      A  W  P  F  S  M  R  P  A  P  C  A  P  R  A  A  L  I  D  V   F2
       P  G  H  S  P  C  G  L  R  P  A  P  L  A  R  R  *  L  M  C  F3
1261 AGCCTGGCCATTCTCCATGCGGCCTGCGCCCTGCGCCCCTCGCGCGGCGCTAATTGATGT 1320
     ----:----|----:----|----:----|----:----|----:----|----:----|

     V  C  Q  V  I  K  L  D  *  L  H  *  L  N  C  C  V  G  A  G    F1
      F  V  K  S  L  N  *  I  N  Y  I  N  *  I  A  A  L  A  L  A   F2
       L  S  S  H  *  I  R  L  I  T  L  I  K  L  L  R  W  R  W  L  F3
1321 GTTTGTCAAGTCATTAAATTAGATTAATTACATTAATTAAATTGCTGCGTTGGCGCTGGC 1380
     ----:----|----:----|----:----|----:----|----:----|----:----|

     Y  C  V  G  L  L  C  E  A  H  V  V  L  F  L  F  S  I  F  F    F1
      I  V  L  V  Y  C  A  R  R  M  W  F  Y  F  C  F  P  F  F  F   F2
       L  C  W  F  T  V  R  G  A  C  G  F  I  F  V  F  H  F  F  F  F3
1381 TATTGTGTTGGTTTACTGTGCGAGGCGCATGTGGTTTTATTTTTGTTTTCCATTTTTTTT 1440
     ----:----|----:----|----:----|----:----|----:----|----:----|

     F  V  I  T  F  C  S  F  K  S  L  S  R  Q  V  L  N  A  P  I    F1
      L  L  S  L  F  V  A  L  N  P  C  P  D  R  Y  *  T  H  L  L   F2
       C  Y  H  F  L  *  L  *  I  L  V  Q  T  G  I  E  R  T  Y  *  F3
1441 TTTGTTATCACTTTTTGTAGCTTTAAATCCTTGTCCAGACAGGTATTGAACGCACCTATT 1500
     ----:----|----:----|----:----|----:----|----:----|----:----|

     S  L  S  N  G  L  I  A  R  L  T  A  R  P  L  C  N  *  S  K    F1
```

```
      A  Y  Q  M  G  *  L  L  G  *  Q  L  V  H  Y  A  I  S  Q  K     F2
       P  I  K  W  A  N  C  S  A  D  S  S  S  I  M  Q  L  V  K  S    F3
1501 AGCCTATCAAATGGGCTAATTGCTCGGCTGACAGCTCGTCCATTATGCAATTAGTCAAAA 1560
     ----:----|----:----|----:----|----:----|----:----|----:----|

     A  C  Q  W  F  F  F  F  F  F  F  L  L  F  *  D  S  M  C  G      F1
      R  V  N  G  F  F  F  F  F  F  F  Y  C  F  R  I  R  C  V  E     F2
       V  S  M  V  F  F  F  F  F  F  F  I  V  L  G  F  D  V  W  S    F3
1561 GCGTGTCAATGGTTTTTTTTTTTTTTTTTTTTTTTATTGTTTTAGGATTCGATGTGTGGA 1620
     ----:----|----:----|----:----|----:----|----:----|----:----|

     V  S  L  P  S  I  L  A  K  N  R  F  N  S  I  V  P  D  *  I      F1
      S  A  S  H  R  F  W  P  K  T  D  L  I  Q  L  S  Q  I  K  Y     F2
       Q  P  P  I  D  F  G  Q  K  P  I  *  F  N  C  P  R  L  N  M    F3
1621 GTCAGCCTCCCATCGATTTTGGCCAAAAACCGATTTAATTCAATTGTCCCAGATTAAATA 1680
     ----:----|----:----|----:----|----:----|----:----|----:----|

     *  A  Q  I  Y  L  C  T  L  N  V  M  P  V  A  V  F  T  L  F      F1
      E  L  K  F  I  C  V  H  *  M  *  C  P  W  L  Y  L  L  Y  S     F2
       S  S  N  L  F  V  Y  I  E  C  D  A  R  G  C  I  Y  F  I  L    F3
1681 TGAGCTCAAATTTATTTGTGTACATTGAATGTGATGCCCGTGGCTGTATTTACTTTATTC 1740
     ----:----|----:----|----:----|----:----|----:----|----:----|

     F  *  L  M  L  T  L  L  I  F  T  F  A  G  N  N  I  L  I  L      F1
      F  N  *  C  *  L  Y  *  Y  L  L  L  L  V  I  T  Y  *  Y  *     F2
       L  I  N  V  N  F  I  D  I  Y  F  C  W  *  *  H  I  N  I  K    F3
1741 TTTTAATTAATGTTAACTTTATTGATATTTACTTTTGCTGGTAATAACATATTAATATTA 1800
     ----:----|----:----|----:----|----:----|----:----|----:----|

     N  N  V  M  K  N  R  H  E  L  F  T  I  Y  F  K  C  F  S  S      F1
      I  M  *  *  K  I  G  M  N  Y  L  Q  F  I  L  N  A  F  H  Q     F2
       *  C  N  E  K  *  A  *  T  I  Y  N  L  F  *  M  L  F  I  R    F3
1801 AATAATGTAATGAAAAATAGGCATGAACTATTTACAATTTATTTTAAATGCTTTTCATCA 1860
     ----:----|----:----|----:----|----:----|----:----|----:----|

     E  I  L  F  T  F  F  P  H  Y  L  D  K  F  N  E  M  S  I  F      F1
      K  F  C  L  H  F  S  H  I  I  L  I  N  S  M  K  C  Q  F  S     F2
       N  F  V  Y  I  F  P  T  L  S  *  *  I  Q  *  N  V  N  F  L    F3
1861 GAAATTTTGTTTACATTTTTCCCACATTATCTTGATAAATTCAATGAAATGTCAATTTTC 1920
     ----:----|----:----|----:----|----:----|----:----|----:----|

     S  Q  L  H  E  F  R  F  N  L  Y  K  R  T  D  S  R  S  L  Q      F1
      H  N  Y  M  N  S  D  L  I  Y  I  S  E  L  I  P  D  L  C  K     F2
       T  I  T  *  I  P  I  *  S  I  *  A  N  *  F  P  I  F  A  R    F3
1921 TCACAATTACATGAATTCCGATTTAATCTATATAAGCGAACTGATTCCCGATCTTTGCAA 1980
     ----:----|----:----|----:----|----:----|----:----|----:----|

     G  L  N  I  F  C  *  D  A  N  P  E  *  R  I  *  F  F  F  L      F1
      A  *  T  S  F  A  E  M  L  I  R  N  R  E  Y  D  F  F  F  *     F2
       P  K  H  L  L  L  R  C  *  S  G  I  E  N  M  I  F  F  S  E    F3
1981 GGCCTAAACATCTTTTGCTGAGATGCTAATCCGGAATAGAGAATATGATTTTTTTTTCTG 2040
     ----:----|----:----|----:----|----:----|----:----|----:----|

     N  A  N  Q  *  L  *  F  V  R  G  H  R  D  I  *  L  E  N  R      F1
      M  P  I  N  S  Y  D  L  Y  E  A  I  E  T  F  D  *  K  I  E     F2
       C  Q  S  I  V  M  I  C  T  R  P  *  R  H  L  I  R  K  *  S    F3
2041 AATGCCAATCAATAGTTATGATTTGTACGAGGCCATAGAGACATTTGATTAGAAAATAGA 2100
     ----:----|----:----|----:----|----:----|----:----|----:----|

     V  L  F  I  *  I  I  V  *  I  E  V  L  L  K  Y  V  F  Q  P      F1
      F  Y  L  F  E  *  *  F  K  *  K  F  C  *  S  M  F  S  S  Q     F2
       F  I  Y  L  N  N  S  L  N  R  S  S  V  K  V  C  F  P  A  K    F3
2101 GTTTTATTTATTTGAATAATAGTTTAAATAGAAGTTCTGTTAAAGTATGTTTTCCAGCCA 2160
```

```
     ----:----|----:----|----:----|----:----|----:----|----:----|

     K  *  T  T  A  S  N  R  L  L  R  R  G  Q  K  L  P  V  L  C     F1
      N  K  R  R  L  L  T  D  S  Y  D  V  A  K  S  Y  L  C  S  V   F2
       I  N  D  G  F  *  P  T  P  T  T  W  P  K  V  T  C  A  L  C  F3
2161 AAATAAACGACGGCTTCTAACCGACTCCTACGACGTGGCCAAAAGTTACCTGTGCTCTGT 2220
     ----:----|----:----|----:----|----:----|----:----|----:----|

     A  V  P  V  C  V  R  S  M  Y  Y  A  *  A  T  E  Y  V  F  F     F1
      L  C  L  C  V  C  D  L  C  I  M  P  R  L  Q  N  M  C  F  L   F2
       C  A  C  V  C  A  I  Y  V  L  C  L  G  Y  R  I  C  V  F  *  F3
2221 GCTGTGCCTGTGTGTGTGCGATCTATGTATTATGCCTAGGCTACAGAATATGTGTTTTTT 2280
     ----:----|----:----|----:----|----:----|----:----|----:----|

     K  A  V  V  *  L  I  L  M  Q  R  D  M  *  *  S  I  W  Q  R     F1
      K  Q  L  Y  N  *  Y  *  C  K  E  I  C  D  K  A  F  G  S  V   F2
       S  S  C  I  I  N  I  N  A  K  R  Y  V  I  K  H  L  A  A  *  F3
2281 AAAGCAGTTGTATAATTAATATTAATGCAAAGAGATATGTGATAAAGCATTTGGCAGCGT 2340
     ----:----|----:----|----:----|----:----|----:----|----:----|

     N  Y  S  I  I  L  Q  *  M  L  C  M  H  S  V  T  *  Y  A  P     F1
      I  I  R  *  F  C  S  K  C  C  A  C  I  P  L  H  S  M  R  H   F2
       L  F  D  N  F  A  V  N  A  V  H  A  F  R  Y  I  V  C  A  I  F3
2341 AATTATTCGATAATTTTGCAGTAAATGCTGTGCATGCATTCCGTTACATAGTATGCGCCA 2400
     ----:----|----:----|----:----|----:----|----:----|----:----|

     *  S  N  H  N  Q  N  A  N  T  G  R  C  T  L  A  H  T  H  T     F1
      N  Q  I  T  T  K  M  Q  I  Q  V  D  A  H  S  H  T  H  T  H   F2
       I  K  S  Q  P  K  C  K  Y  R  *  M  H  T  R  T  H  T  H  T  F3
2401 TAATCAAATCACAACCAAAATGCAAATACAGGTAGATGCACACTCGCACACACACACACA 2460
     ----:----|----:----|----:----|----:----|----:----|----:----|

     H  T  H  I  D  *  W  G  P  K  T  V  T  Y  F  P  *  A  I  A     F1
      T  H  T  S  I  D  G  A  Q  K  P  *  L  I  S  L  E  R  L  H   F2
       H  T  H  R  L  M  G  P  K  N  R  N  L  F  P  L  S  D  C  M  F3
2461 CACACACACATCGATTGATGGGGCCCAAAAACCGTAACTTATTTCCCTTGAGCGATTGCA 2520
     ----:----|----:----|----:----|----:----|----:----|----:----|

     W  K  C  Q  S  L  C  D  C  F  Y  C  C  C  C  *  C  C  C  S     F1
      G  N  A  R  V  C  A  I  A  F  I  V  V  V  V  D  V  V  V  L   F2
       E  M  P  E  S  V  R  L  L  L  L  L  L  L  L  M  L  L  F  Y  F3
2521 TGGAAATGCCAGAGTCTGTGCGATTGCTTTTATTGTTGTTGTTGTTGATGTTGTTGTTCT 2580
     ----:----|----:----|----:----|----:----|----:----|----:----|

     M  T  A  H  *  S  T  I  L  D  G  S  I  R  A  Q  I  H  K  S     F1
      *  R  R  T  N  Q  Q  F  W  T  V  Q  S  E  R  K  F  I  N  H   F2
       D  G  A  L  I  N  N  F  G  R  F  N  Q  S  A  N  S  *  I  I  F3
2581 ATGACGGCGCACTAATCAACAATTTTGGACGGTTCAATCAGAGCGCAAATTCATAAATCA 2640
     ----:----|----:----|----:----|----:----|----:----|----:----|

     F  I  C  K  R  P  T  E  *  K  G  L  R  R  *  R  S  G  G  M     F1
      L  Y  A  N  A  Q  R  N  R  R  D  C  A  D  R  G  V  G  E  W   F2
       Y  M  Q  T  P  N  G  I  E  G  T  A  P  I  E  E  W  G  N  G  F3
2641 TTTATATGCAAACGCCCAACGGAATAGAAGGGACTGCGCCGATAGAGGAGTGGGGGAATG 2700
     ----:----|----:----|----:----|----:----|----:----|----:----|

     D  P  Q  I  N  C  G  H  F  N  T  S  N  Q  L  N  N  *  I  L     F1
      T  H  R  *  T  V  G  I  S  T  P  Q  T  S  *  I  I  E  Y  *   F2
       P  T  D  K  L  W  A  F  Q  H  L  K  P  V  K  *  L  N  I  E  F3
2701 GACCCACAGATAAACTGTGGGCATTTCAACACCTCAAACCAGTTAAATAATTGAATATTG 2760
     ----:----|----:----|----:----|----:----|----:----|----:----|

     N  Q  I  *  S  F  K  R  K  A  T  V  H  C  V  G  K  Q  A  *     F1
```

```
      I  K  S  N  H  S  N  A  K  P  Q  S  I  V  W  A  S  R  L  D    F2
       S  N  L  I  I  Q  T  Q  S  H  S  P  L  C  G  Q  A  G  L  I   F3
2761 AATCAAATCTAATCATTCAAACGCAAAGCCACAGTCCATTGTGTGGGCAAGCAGGCTTGA 2820
     ----:----|----:----|----:----|----:----|----:----|----:----|

     *  G  N  C  *  G  S  D  W  F  H  W  H  C  R  A  S  R  T  Y     F1
      R  G  I  A  K  A  A  T  G  S  I  G  I  A  V  H  P  E  R  I    F2
       G  E  L  L  R  Q  R  L  V  P  L  A  L  P  C  I  Q  N  V  S   F3
2821 TAGGGGAATTGCTAAGGCAGCGACTGGTTCCATTGGCATTGCCGTGCATCCAGAACGTAT 2880
     ----:----|----:----|----:----|----:----|----:----|----:----|

     H  *  S  N  I  L  G  P  P  D  L  A  Q  F  T  R  R  Y  F  M     F1
      I  N  Q  T  F  W  G  P  R  T  W  L  N  L  L  V  V  I  L  C    F2
       L  I  K  H  F  G  A  P  G  P  G  S  I  Y  S  S  L  F  Y  V   F3
2881 CATTAATCAAACATTTTGGGGCCCCCGGACCTGGCTCAATTTACTCGTCGTTATTTTATG 2940
     ----:----|----:----|----:----|----:----|----:----|----:----|

     Y  F  *  H  *  L  M  P  D  I  S  R  I  S  S  F  V  S  P  *     F1
      I  F  S  I  N  L  C  P  T  F  L  A  F  H  L  S  F  L  R  R    F2
       F  L  A  L  T  Y  A  R  H  F  S  H  F  I  F  R  F  S  V  G   F3
2941 TATTTTTAGCATTAACTTATGCCCGACATTTCTCGCATTTCATCTTTCGTTTCTCCGTAG 3000
     ----:----|----:----|----:----|----:----|----:----|----:----|

     A  I  N  C  S  N  R  Y  H  N  Y  T  S  D  L  P  P  P  R  K     F1
      P  L  T  A  R  I  A  T  T  I  T  H  L  I  C  R  R  R  A  R    F2
       H  *  L  L  E  S  L  P  Q  L  H  I  *  F  A  A  A  A  Q  D   F3
3001 GCCATTAACTGCTCGAATCGCTACCACAATTACACATCTGATTTGCCGCCGCCGCGCAAG 3060
     ----:----|----:----|----:----|----:----|----:----|----:----|

     T  S  C  G  R  T  S                                            F1
      R  P  A  A  G  P  X                                           F2
       V  L  R  Q  D  L                                             F3
3061 ACGTCCTGCGGCAGGACCTC 3080
```
