## Supplementary material for "The incidence of candidate binding sites for β-arrestin in Drosophila neuropeptide GPCRs": S3 Text

*D virilis CCAP R* genomic DNA:
[https://www.ncbi.nlm.nih.gov/nuccore/1805215796?report=graph&v=3415416:3456792](https://www.ncbi.nlm.nih.gov/nuccore/1805215796?report=graph&v=3415416:3456792)

In *D melanogaster*, PC and PD isoforms are predicted; the latter by stop suppression, and it contains an additional BBS, ***SFSADE***

Blastp searches of *D virilis* reveal only a predicted PC isoform.  Here looking for a STOP Suppression-generated PD isoform orthologue

The following sequence (starting at 3416271 bp of (NW_022587377.1), and including termination of the predicted PC orthologue, shows that *D virilis* also contains the potential for expression of a similar stop-suppressed PD isoform which features an additional BBS: *SFSDAE*

```
  Q  R  Y  D  S  W  R  I  Q  P  A  S  V  Q  R  L  R  S  T  G      F1
     R  G  M  I  A  G  A  F  N  P  Q  A  F  N  V  C  G  P  R  A     F2
      E  V  *  *  L  A  H  S  T  R  K  R  S  T  F  A  V  H  G  P    F3
  1 CAGAGGTATGATAGCTGGCGCATTCAACCCGCAAGCGTTCAACGTTTGCGGTCCACGGGC 60
    ----:----|----:----|----:----|----:----|----:----|----:----|

    R  R  P  S  A  M  R  S  F  Y  K  *  I  A  S  A  H  C  R  L      F1
     D  V  L  Q  R  C  G  V  S  I  S  K  L  P  A  P  T  V  G  Y     F2
      T  S  F  S  D  A  E  F  L  *  V  N  C  Q  R  P  L  *  A  I    F3
 61 CGACGTCCTTCAGCGATGCGGAGTTTCTATAAGTAAATTGCCAGCGCCCACTGTAGGCTA 120
    ----:----|----:----|----:----|----:----|----:----|----:----|

    *  E  A  *  T  I  S  A  I  V  V  C  A  *  R  M  P  K  S  N      F1
     R  R  L  R  Q  *  V  R  *  L  C  V  H  S  E  C  P  N  R  I     F2
      G  G  L  D  N  K  C  D  S  C  V  C  I  A  N  A  Q  I  E  Y    F3
121 TAGGAGGCTTAGACAATAAGTGCGATAGTTGTGTGTGCATAGCGAATGCCCAAATCGAAT 180
    ----:----|----:----|----:----|----:----|----:----|----:----|

    I  Y  T  Y  T  Y  M  H  N  V  R  V  L  R  C  A  T  N  F  N      F1
     Y  I  H  T  H  I  C  I  M  C  V  Y  L  G  V  L  R  I  S  M     F2
      I  Y  I  H  I  Y  A  *  C  A  C  T  *  V  C  Y  E  F  Q  C    F3
181 ATATATACATACACATATATGCATAATGTGCGTGTACTTAGGTGTGCTACGAATTTCAAT 240
    ----:----|----:----|----:----|----:----|----:----|----:----|

    A  C  I  S  H  C  N  L  F  *  V  F  M  Q  L  A  C  N  Y  Y      F1
     H  V  F  P  I  A  I  Y  S  K  Y  L  C  N  *  L  V  T  T  I     F2
      M  Y  F  P  L  Q  F  I  L  S  I  Y  A  I  S  L  *  L  L  *    F3
241 GCATGTATTTCCCATTGCAATTTATTCTAAGTATTTATGCAATTAGCTTGTAACTACTAT 300
    ----:----|----:----|----:----|----:----|----:----|----:----|

    K  T  K  L  I  Q  L  N  E  N  L  T  C  I  G  H  R  F  X         F1
     K  Q  N  L  F  N  *  M  R  T  *  H  A  L  G  I  D  L           F2
      N  K  T  Y  S  T  K  *  E  L  N  M  H  W  A  S  I  X          F3
301 AAAACAAAACTTATTCAACTAAATGAGAACTTAACATGCATTGGGCATCGATTTA 355
    ----:----|----:----|----:----|----:----|----:----|----:
```
