## Supplementary material for "The incidence of candidate binding sites for β-arrestin in Drosophila neuropeptide GPCRs": S4 Text

*D. virilis RYaR* Genomic DNA from analysis of:
https://www.ncbi.nlm.nih.gov/nuccore/1805215796?report=graph&v=7837313:7930656

Annotations of the *D. melanogaster RYaR* present three isoforms, PA, PB, and PC. Blastp search of *D virilis* recovers only a PA ortholog. Below is the inspection of the genomic DNA to search for possible PB and PC orthologs.

Here I present evidence for possible expression of RYaR PB in *D virilis*, but do not find a PC candidate.

bp #1 of the record below = bp #7839511 of the above link from the shotgun sequence found in the above link

Following D melanogaster gene structure, last two exons in *D virilis* should use alternative frames to produce PA and PB sequences:

The predicted open reading frames, and predicted splice donor and acceptor sites, are marked in **RED** for sequences common to PA and PB. After the begin to use alternative reading frames as described below, The PA sequence is in **GREEN** and the PB in **RED**

If the splice donor at bp 522 below is neglected, the run-on sequence produces a good match for the predicted PB isoform and includes candidate BBS within 10 bp of the protein end, as expected.

PC should be an ~18 AA extension of PB (KIKIR) to a stop – but that is not evident.

```
    I  V  R  G  R  G  H  I  C  L  A  *  I  N  I  L  L  H  F  A     F1
     *  C  G  D  G  G  T  S  A  W  L  K  *  T  F  Y  Y  I  S  P    F2
      S  A  G  T  G  A  H  L  L  G  L  N  K  H  F  I  T  F  R  L   F3
  1 ATAGTGCGGGGACGGGGGCACATCTGCTTGGCTTAAATAAACATTTTATTACATTTCGCC 60
    ----:----|----:----|----:----|----:----|----:----|----:----|

    L  I  L  A  L  F  S  A  L  T  D  D  *  N  D  A  D  C  C  N     F1
     *  F  L  L  C  S  L  L  L  Q  M  I  K  M  M  L  T  V  V  I    F2
      N  S  C  S  V  L  C  S  Y  R  *  L  K  *  C  *  L  L  *  L   F3
 61 TTAATTCTTGCTCTGTTCTCTGCTCTTACAGATGATTAAAATGATGCTGACTGTTGTAAT 120
    ----:----|----:----|----:----|----:----|----:----|----:----|

    C  I  H  Q  L  L  A  T  I  Q  H  T  T  G  G  W  C  G  G  S     F1
     V  F  T  S  C  W  L  P  F  N  I  L  Q  V  G  G  A  V  V  R    F2
      Y  S  P  A  A  G  Y  H  S  T  Y  Y  R  W  V  V  R  W  F  G   F3
121 TGTATTCACCAGCTGCTGGCTACCATTCAACATACTACAGGTGGGTGGTGCGGTGGTTCG 180
    ----:----|----:----|----:----|----:----|----:----|----:----|

    V  V  A  L  A  A  N  *  P  Q  G  A  Q  I  L  P  L  I  E  F     F1
     W  L  P  W  Q  Q  I  S  H  K  A  H  K  S  C  H  *  L  S  F    F2
      G  C  P  G  S  K  L  A  T  R  R  T  N  L  A  I  D  *  V  S   F3
181 GTGGTTGCCCTGGCAGCAAATTAGCCACAAGGCGCACAAATCTTGCCATTGATTGAGTTT 240
```

```
    ----:----|----:----|----:----|----:----|----:----|----:----|

    L  S  K  P  S  P  P  P  H  L  F  L  S  L  A  A  D  A  *  R     F1
     S  L  N  L  H  P  L  P  I  S  F  S  P  L  Q  L  M  L  N  D    F2
      L  *  T  F  T  P  S  P  S  L  S  L  P  C  S  *  C  L  T  M   F3
241 CTCTCTAAACCTTCACCCCCTCCCCATCTCTTTCTCTCCCTTGCAGCTGATGCTTAACGA 300
    ----:----|----:----|----:----|----:----|----:----|----:----|

    *  G  V  R  Q  L  E  T  V  A  L  C  V  V  R  L  S  L  A  G     F1
     E  E  F  A  N  W  K  P  L  P  Y  V  W  F  A  F  H  W  L  A    F2
      R  S  S  P  T  G  N  R  C  P  M  C  G  S  P  F  T  G  W  P   F3
301 TGAGGAGTTCGCCAACTGGAAACCGTTGCCCTATGTGTGGTTCGCCTTTCACTGGCTGGC 360
    ----:----|----:----|----:----|----:----|----:----|----:----|

    H  V  A  Q  L  L  Q  S  D  H  L  L  L  Y  E  C  P  F  P  W     F1
     M  S  H  S  C  Y  N  P  I  I  Y  C  Y  M  N  A  R  F  R  G    F2
      C  R  T  A  A  T  I  R  S  S  I  A  I  *  M  P  V  S  V  A   F3
361 CATGTCGCACAGCTGCTACAATCCGATCATCTATTGCTATATGAATGCCCGTTTCCGTGG 420
    ----:----|----:----|----:----|----:----|----:----|----:----|

    R  I  P  T  D  N  V  P  C  A  R  P  A  A  L  L  L  F  A  S     F1
     G  F  L  Q  I  M  Y  R  V  P  G  L  R  R  C  C  C  L  H  R    F2
      D  S  Y  R  *  C  T  V  C  P  A  C  G  A  A  A  V  C  I  D   F3
421 CGGATTCCTACAGATAATGTACCGTGTGCCCGGCCTGCGGCGCTGCTGCTGTTTGCATCG 480
    ----:----|----:----|----:----|----:----|----:----|----:----|

    I  F  A  Q  S  R  R  A  Q  L  *  S  N  R  *  D  D  L  Q  I     F1
     Y  L  R  S  R  G  E  R  S  Y  E  A  T  G  E  M  T  S  K  Y    F2
      I  C  A  V  A  A  S  A  A  M  K  Q  P  V  R  *  P  P  N  T   F3
481 ATATTTGCGCAGTCGCGGCGAGCGCAGCTATGAAGCAACCGGTGAGATGACCTCCAAATA 540
    ----:----|----:----|----:----|----:----|----:----|----:----|

    Q  S  P  K  W  *  W  S  S  Q  E  T  *  N  T  H  *  V  P  P     F1
     N  R  R  N  G  D  G  L  V  R  K  P  K  I  R  I  R  C  L  L    F2
      I  A  E  M  V  M  V  *  S  G  N  L  K  Y  A  L  G  A  S  S   F3
541 CAATCGCCGAAATGGTGATGGTCTAGTCAGGAAACCTAAAATACGCATTAGGTGCCTCCT 600
    ----:----|----:----|----:----|----:----|----:----|----:----|

    P  L  S  L  *  Y  I  I  M  Y  F  K  S  I  C  L  D  V  C  Q     F1
     P  Y  L  F  S  T  L  L  C  I  S  N  L  F  V  L  T  F  A  K    F2
      L  I  S  L  V  H  Y  Y  V  F  Q  I  Y  L  S  *  R  L  P  K   F3
601 CCCTTATCTCTTTAGTACATTATTATGTATTTCAAATCTATTTGTCTTGACGTTTGCCAA 660
    ----:----|----:----|----:----|----:----|----:----|----:----|

    K  K  K  R  K  E  N  K  *  I  S  K  *  Y  K  F  C  R  R  A     F1
     K  K  K  E  K  K  I  S  E  Y  Q  N  N  T  S  F  A  G  G  L    F2
      K  K  K  K  R  K  *  V  N  I  K  I  I  Q  V  L  Q  A  G  L   F3
661 AAAAAAAAAAGAAAAGAAAATAAGTGAATATCAAAATAATACAAGTTTTGCAGGCGGGCT 720
    ----:----|----:----|----:----|----:----|----:----|----:----|

    *  P  T  A  K  Y  S  F  T  V  Q  *  L  L  E  I  G  *  I  A     F1
     N  R  L  R  N  T  R  S  Q  S  S  N  C  L  K  *  A  K  *  P    F2
      T  D  C  E  I  L  V  H  S  P  V  I  A  *  N  R  L  N  S  Q   F3
721 TAACCGACTGCGAAATACTCGTTCACAGTCCAGTAATTGCTTGAAATAGGCTAAATAGCC 780
    ----:----|----:----|----:----|----:----|----:----|----:----|

    N  I  I  F  T  I  Y  L  L  F  R  F  T  Y  I  F  I  Y  F  F     F1
     I  L  Y  L  R  Y  I  Y  F  S  D  L  H  I  Y  L  F  I  F  S    F2
      Y  Y  I  Y  D  I  F  T  F  P  I  Y  I  Y  I  Y  L  F  F  L   F3
781 AATATTATATTTACGATATATTTACTTTTCCGATTTACATATATATTTATTTATTTTTTC 840
    ----:----|----:----|----:----|----:----|----:----|----:----|

    F  E  V  F  Q  Y  Y  Q  W  F  Q  L  V  N  C  G  S  L  Y  I     F1
```

```
      L  K  Y  F  N  I  I  S  G  S  S  L  *  T  V  G  A  Y  I  S    F2
       *  S  I  S  I  L  S  V  V  P  A  C  K  L  W  E  L  I  Y  Q   F3
 841 TTTGAAGTATTTCAATATTATCAGTGGTTCCAGCTTGTAAACTGTGGGAGCTTATATATC 900
     ----:----|----:----|----:----|----:----|----:----|----:----|

     N  I  I  L  D  F  Y  L  N  P  L  K  P  C  E  R  V  F  K  T     F1
      I  *  Y  *  T  F  I  *  T  P  *  N  L  V  N  E  Y  L  K  Q    F2
       Y  N  I  R  L  L  S  K  P  P  E  T  L  *  T  S  I  *  N  K   F3
 901 AATATAATATTAGACTTTTATCTAAACCCCCTGAAACCTTGTGAACGAGTATTTAAAACA 960
     ----:----|----:----|----:----|----:----|----:----|----:----|

     K  S  L  T  I  L  I  S  *  F  P  *  V  V  S  S  S  S  F  P     F1
      K  A  *  Q  F  *  L  A  S  S  P  R  *  L  V  P  R  V  F  H    F2
       K  P  N  N  S  D  *  L  V  P  L  G  S  *  F  L  E  F  S  I   F3
 961 AAAAGCCTAACAATTCTGATTAGCTAGTTCCCCTAGGTAGTTAGTTCCTCGAGTTTTCCA 1020
     ----:----|----:----|----:----|----:----|----:----|----:----|

     Y  A  H  A  C  C  T  H  S  T  K  T  L  S  L  H  A  *  A  Q     F1
      M  P  M  H  V  V  H  I  A  Q  K  P  Y  H  C  M  R  E  H  R    F2
       C  P  C  M  L  Y  T  *  H  K  N  P  I  T  A  C  V  S  T  E   F3
1021 TATGCCCATGCATGTTGTACACATAGCACAAAAACCCTATCACTGCATGCGTGAGCACAG 1080
     ----:----|----:----|----:----|----:----|----:----|----:----|

     N  F  T  Y  S  I  Y  Y  E  S  F  C  H  S  C  N  F  *  L  L     F1
      I  L  H  I  Q  F  I  M  K  A  F  V  T  L  V  I  F  N  C  *    F2
       F  Y  I  F  N  L  L  *  K  L  L  S  L  L  *  F  L  I  V  N   F3
1081 AATTTTACATATTCAATTTATTATGAAAGCTTTTGTCACTCTTGTAATTTTTAATTGTTA 1140
     ----:----|----:----|----:----|----:----|----:----|----:----|

     I  A  V  V  V  V  F  L  Y  S  T  P  P  E  K  E  P  K  M  H     F1
      L  L  L  L  L  F  F  C  I  P  R  R  L  K  R  N  R  R  C  I    F2
       C  C  C  C  C  F  S  V  F  H  A  A  *  K  G  T  E  D  A  F   F3
1141 ATTGCTGTTGTTGTTGTTTTTCTGTATTCCACGCCGCCTGAAAAGGAACCGAAGATGCAT 1200
     ----:----|----:----|----:----|----:----|----:----|----:----|

     S  T  Y  I  V  *  T  P  A  P  P  T  S  A  Q  D  E  S  C  A     F1
      P  P  T  S  C  E  H  L  H  H  L  H  Q  H  K  T  K  A  A  H    F2
       H  L  H  R  V  N  T  C  T  T  Y  I  S  T  R  R  K  L  R  T   F3
1201 TCCACCTACATCGTGTGAACACCTGCACCACCTACATCAGCACAAGACGAAAGCTGCGCA 1260
     ----:----|----:----|----:----|----:----|----:----|----:----|

     R  I  L  C  K  W  *  V  E  L  F  P  T  L  Y  S  H  I  T  I     F1
      E  F  Y  A  N  G  E  *  S  S  F  Q  H  Y  T  A  I  *  L  Y    F2
       N  S  M  Q  M  V  S  R  A  L  S  N  I  I  Q  P  Y  N  Y  I   F3
1261 CGAATTCTATGCAAATGGTGAGTAGAGCTCTTTCCAACATTATACAGCCATATAACTATA 1320
     ----:----|----:----|----:----|----:----|----:----|----:----|

     F  L  T  E  P  I  F  M  C  R  D  H  S  V  A  V  A  P  G  T     F1
      F  L  Q  S  Q  F  S  C  A  E  T  T  V  L  R  *  L  P  A  P    F2
       S  Y  R  A  N  F  H  V  P  R  P  Q  C  C  G  S  S  R  H  R   F3
1321 TTTCTTACAGAGCCAATTTTCATGTGCCGAGACCACAGTGTTGCGGTAGCTCCCGGCACC 1380
     ----:----|----:----|----:----|----:----|----:----|----:----|

     A  Q  E  P  K  P  R  L  Q  P  R  Q  P  T  G  S  F  H  L  N     F1
      R  R  S  Q  N  Q  G  S  S  H  A  N  Q  L  V  H  F  T  *  T    F2
       A  G  A  K  T  K  A  P  A  T  P  T  N  W  F  I  S  P  E  Q   F3
1381 GCGCAGGAGCCAAAACCAAGGCTCCAGCCACGCCAACCAACTGGTTCATTTCACCTGAAC 1440
     ----:----|----:----|----:----|----:----|----:----|----:----|

     S  A  S  Y  L  S  S  T  Q  F  *  N  T  E  A  *  N  K  *  V     F1
      A  P  A  T  *  A  P  H  S  F  E  I  Q  K  L  K  I  N  E  F    F2
       R  Q  L  P  E  L  H  T  V  L  K  Y  R  S  L  K  *  M  S  L   F3
1441 AGCGCCAGCTACCTGAGCTCCACACAGTTTTGAAATACAGAAGCTTAAAATAAATGAGTT 1500
```

```
     ----:----|----:----|----:----|----:----|----:----|----:----|

     *  T  K  D  V  L  N  L  H  *  Y  T  I  S  R  V  H  A  C  K    F1
      E  P  K  M  C  *  I  S  I  D  I  L  Y  L  E  Y  M  H  A  K   F2
       N  Q  R  C  A  E  S  P  L  I  Y  Y  I  *  S  T  C  M  Q  S  F3
1501 TGAACCAAAGATGTGCTGAATCTCCATTGATATACTATATCTAGAGTACATGCATGCAAA 1560
     ----:----|----:----|----:----|----:----|----:----|----:----|

     A  S  Y  V  S  T  Y  L  *  C  *  V  Y  Y  C  K  C  *  L  N    F1
      L  V  M  C  L  H  T  Y  S  A  K  C  I  I  V  N  V  D  S  M   F2
       *  L  C  V  Y  I  L  I  V  L  S  V  L  L  *  M  L  T  Q  C  F3
1561 GCTAGTTATGTGTCTACATACTTATAGTGCTAAGTGTATTATTGTAAATGTTGACTCAAT 1620
     ----:----|----:----|----:----|----:----|----:----|----:----|

     V  N  E  *  N  M  Y  Y  K  Y  P  K  I  N  V  H  R  N  T  I    F1
      *  M  N  R  I  C  I  T  N  I  Q  K  *  M  C  I  E  I  Q  L   F2
       K  *  I  E  Y  V  L  Q  I  S  K  N  K  C  A  *  K  Y  N  *  F3
1621 GTAAATGAATAGAATATGTATTACAAATATCCAAAAATAAATGTGCATAGAAATACAATT 1680
     ----:----|----:----|----:----|----:----|----:----|----:----|

     D  V  R  N  K  K  F  *  S  N  F  I  I  *  *  I  Y  K  R  N    F1
      T  F  E  I  K  N  F  N  L  I  L  L  Y  N  K  Y  I  K  E  I   F2
       R  S  K  *  K  I  L  I  *  F  Y  Y  I  I  N  I  *  K  K  Y  F3
1681 GACGTTCGAAATAAAAAATTTTAATCTAATTTTATTATATAATAAATATATAAAAGAAAT 1740
     ----:----|----:----|----:----|----:----|----:----|----:----|

     M  *  K  Y  K  T  N  I  Y  S  M  A  G  K  I  K  S  A  *  W    F1
      C  K  N  I  K  Q  I  Y  T  A  W  Q  A  K  *  K  V  H  N  G   F2
       V  K  I  *  N  K  Y  I  Q  H  G  R  Q  N  K  K  C  I  M  E  F3
1741 ATGTAAAAATATAAAACAAATATATACAGCATGGCAGGCAAAATAAAAAGTGCATAATGG 1800
     ----:----|----:----|----:----|----:----|----:----|----:----|

     K  L  T  E  T  L  F  L  F  R  S  I  P  V  A  K  C  I  V  I    F1
      N  *  Q  K  L  Y  F  Y  S  A  P  F  Q  S  Q  N  A  S  *  S   F2
       I  N  R  N  F  I  F  I  P  L  H  S  S  R  K  M  H  R  N  Q  F3
1801 AAATTAACAGAAACTTTATTTTTATTCCGCTCCATTCCAGTCGCAAAATGCATCGTAATC 1860
     ----:----|----:----|----:----|----:----|----:----|----:----|

     R  P  K  M  P  R  C  V  V  E  F  Y  L  I  N  A  Q  K  Y  A    F1
      D  P  K  C  L  G  A  W  W  N  F  I  *  L  M  H  K  S  M  Q   F2
       T  Q  N  A  *  V  R  G  G  I  L  F  N  *  C  T  K  V  C  S  F3
1861 AGACCCAAAATGCCTAGGTGCGTGGTGGAATTTTATTTAATTAATGCACAAAAGTATGCA 1920
     ----:----|----:----|----:----|----:----|----:----|----:----|

     A  L  Y  A  N  G  A  P  T  T  R  K  M  C  C  I  L  Q  R  A    F1
      H  F  M  P  T  E  H  Q  L  R  V  K  C  V  A  Y  F  K  E  Q   F2
       T  L  C  Q  R  S  T  N  Y  A  *  N  V  L  H  T  S  K  S  N  F3
1921 GCACTTTATGCCAACGGAGCACCAACTACGCGTAAAATGTGTTGCATACTTCAAAGAGCA 1980
     ----:----|----:----|----:----|----:----|----:----|----:----|

     T  C  K  L  R  I  R                                           F1
      R  V  N  C  V  Y  X                                          F2
       V  *  I  A  Y  T                                            F3
1981 ACGTGTAAATTGCGTATACG 2000
     ----:----|----:----|
```
