## Supplementary material for "The incidence of candidate binding sites for β-arrestin in Drosophila neuropeptide GPCRs": S5 Text

**Alternative isoforms of Tk R 86C**

The *D melanogaster* the RB isoform contains an unspliced intron, introducing additional sequence in the middle of the CT and it that contains a BBS.
Below I inspect the gDNA from *D virilis* and find that the corresponding intronic region does contain a potential exon that encodes a BBS. This may be the virilis PB isoform; but the BBS sequence is not at all similar and no other feature of the intervening peptide sequence corresponds. This example is not compelling and so I cannot conclude with much certainty that it is accurate and do not include it in the main Fugures.

*D virilis* genomic and transcript information:
https://www.ncbi.nlm.nih.gov/nuccore/1805215796?report=graph&v=5251816:5271610

Below bp#1 = 5266391 from the shotgun sequence in the link

only predicted RNA is a RA-like (XM_002053574), which in *D. melanogaster,* splices out the small 52 AA section containing an additional BBS in PB – so have to look at genomic data
The 7$^{th}$ TM is encoded starting at bp 3331
The penultimate intron starts at bp 3452 and ends at bp 4028:
The final intron starts at bp 4147 and ends at 4207
Search in those introns for a cryptic exon encoding a protein section (that matches that in *D. melanogaster* PB) of approximately 51 AA containing a BBS
Bp 3637 starts a potential exon that does include the following BBS: ***SSTESS***
However, the peptide sequence introduced by this potential exon is not close to 51 AAs and the BBS sequence does not match that of the D melanogaster GPCR

**conclude: virilis lost the exon encoding that additional CT sequence and extra BBS**

```
  S  F  S  P  Q  G  L  S  Y  *  M  L  V  S  F  T  H  A  S  L     F1
    L  S  P  H  R  D  F  L  T  E  C  L  F  P  S  P  T  R  P  Y    F2
     F  L  P  T  G  T  F  L  L  N  A  C  F  L  H  P  R  V  L  M   F3
 1 TCTTTCTCCCCACAGGGACTTTCTTACTGAATGCTTGTTTCCTTCACCCACGCGTCCTTA 60
   ----:----|----:----|----:----|----:----|----:----|----:----|

   *  V  A  V  G  T  E  N  D  L  G  H  C  I  W  A  D  D  V  R      F1
    E  L  P  W  E  Q  K  T  I  W  A  I  V  F  G  L  M  M  F  V    F2
     S  C  R  G  N  R  K  R  F  G  P  L  Y  L  G  *  *  C  S  W   F3
61 TGAGTTGCCGTGGGAACAGAAAACGATTTGGGCCATTGTATTTGGGCTGATGATGTTCGT 120
   ----:----|----:----|----:----|----:----|----:----|----:----|

   G  H  C  G  Q  W  N  C  S  L  D  C  Y  R  *  A  L  D  S  *      F1
```

```
     A  I  A  G  N  G  I  V  L  W  I  V  T  G  K  H  L  I  V  K    F2
      P  L  R  A  M  E  L  F  S  G  L  L  Q  V  S  T  *  *  L  N   F3
121 GGCCATTGCGGGCAATGGAATTGTTCTCTGGATTGTTACAGGTAAGCACTTGATAGTTAA 180
    ----:----|----:----|----:----|----:----|----:----|----:----|

    T  G  L  R  L  F  Y  S  *  K  W  K  K  N  L  S  A  T  L  K     F1
     Q  A  *  D  C  F  I  V  E  N  G  R  R  T  *  A  P  H  *  R    F2
      R  L  K  I  V  L  *  L  K  M  E  E  E  L  K  R  H  I  K  E   F3
181 ACAGGCTTAAGATTGTTTTATAGTTGAAAATGGAAGAAGAACTTAAGCGCCACATTAAAG 240
    ----:----|----:----|----:----|----:----|----:----|----:----|

    S  I  F  D  L  N  H  V  Y  A  T  R  S  Q  F  C  L  V  A  W     F1
     V  F  L  T  *  I  T  Y  T  P  R  G  R  N  F  V  W  S  L  G    F2
      Y  F  *  L  K  S  R  I  R  H  A  V  A  I  L  F  G  R  L  A   F3
241 AGTATTTTTGACTTAAATCACGTATACGCCACGCGGTCGCAATTTTGTTTGGTCGCTTGG 300
    ----:----|----:----|----:----|----:----|----:----|----:----|

    L  H  R  L  L  A  L  F  S  R  I  S  C  C  *  Y  P  V  D  F     F1
     C  I  D  C  *  R  C  F  Q  E  *  V  A  A  N  I  R  W  I  S    F2
      A  S  I  A  S  V  V  F  K  N  K  L  L  L  I  S  G  G  F  L   F3
301 CTGCATCGATTGCTAGCGTTGTTTTCAAGAATAAGTTGCTGCTAATATCCGGTGGATTTC 360
    ----:----|----:----|----:----|----:----|----:----|----:----|

    S  L  L  S  R  L  T  F  W  P  L  F  V  L  F  S  G  H  R  S     F1
     L  C  C  H  V  S  H  F  G  H  F  S  F  C  F  Q  D  I  A  A    F2
      F  V  V  T  S  H  I  L  A  T  F  R  F  V  F  R  T  S  Q  H   F3
361 TCTTTGTTGTCACGTCTCACATTTTGGCCACTTTTCGTTTTGTTTTCAGGACATCGCAGC 420
    ----:----|----:----|----:----|----:----|----:----|----:----|

    M  R  T  V  T  N  Y  F  L  L  N  L  S  I  A  D  L  L  M  S     F1
     C  A  R  S  Q  I  I  S  C  *  T  *  V  S  L  T  C  *  C  P    F2
      A  H  G  H  K  L  F  P  A  E  P  E  Y  R  *  P  A  N  V  H   F3
421 ATGCGCACGGTCACAAATTATTTCCTGCTGAACCTGAGTATCGCTGACCTGCTAATGTCC 480
    ----:----|----:----|----:----|----:----|----:----|----:----|

    T  L  N  C  V  F  N  F  I  F  M  V  N  S  D  W  P  F  G  S     F1
     P  *  I  V  S  L  T  L  Y  S  W  *  I  R  I  G  H  L  A  R    F2
      L  E  L  C  L  *  L  Y  I  H  G  E  F  G  L  A  I  W  L  D   F3
481 ACCTTGAATTGTGTCTTTAACTTTATATTCATGGTGAATTCGGATTGGCCATTTGGCTCG 540
    ----:----|----:----|----:----|----:----|----:----|----:----|

    I  Y  C  T  I  N  N  F  V  A  N  V  T  V  S  T  S  V  F  T     F1
     F  I  A  P  S  T  I  L  W  Q  T  *  R  F  Q  H  R  S  S  P    F2
      L  L  H  H  Q  Q  F  C  G  K  R  N  G  F  N  I  G  L  H  P   F3
541 ATTTATTGCACCATCAACAATTTTGTGGCAAACGTAACGGTTTCAACATCGGTCTTCACC 600
    ----:----|----:----|----:----|----:----|----:----|----:----|

    L  V  A  I  S  F  D  R  *  V  L  S  V  C  Y  N  L  H  C  I     F1
     S  L  P  S  P  S  T  G  E  C  *  V  F  A  I  I  F  I  A  S    F2
      R  C  H  L  L  R  Q  V  S  A  E  C  L  L  *  S  S  L  H  L   F3
601 CTCGTTGCCATCTCCTTCGACAGGTGAGTGCTGAGTGTTTGCTATAATCTTCATTGCATC 660
    ----:----|----:----|----:----|----:----|----:----|----:----|

    Y  T  R  I  R  I  G  F  R  L  C  I  R  V  L  S  L  Q  I  Y     F1
     I  P  A  S  A  S  A  S  V  C  A  F  V  F  Y  L  C  R  Y  I    F2
      Y  P  H  P  H  R  L  P  F  V  H  S  C  S  I  F  A  D  I  L   F3
661 TATACCCGCATCCGCATCGGCTTCCGTTTGTGCATTCGTGTTCTATCTTTGCAGATATAT 720
    ----:----|----:----|----:----|----:----|----:----|----:----|

    C  H  S  T  S  S  K  A  A  H  V  A  S  Q  S  A  I  H  S  G     F1
     A  I  V  H  P  L  K  R  R  T  S  R  R  K  V  R  F  I  L  V    F2
      P  *  Y  I  L  *  S  G  A  R  R  V  A  K  C  D  S  F  W  Y   F3
721 TGCCATAGTACATCCTCTAAAGCGGCGCACGTCGCGTCGCAAAGTGCGATTCATTCTGGT 780
```

```
     ----:----|----:----|----:----|----:----|----:----|----:----|

     T  D  L  G  T  E  L  C  P  L  R  A  L  P  A  L  L  Q  H  H    F1
      L  I  W  A  L  S  C  V  L  S  A  P  C  L  L  Y  S  S  I  M   F2
       *  S  G  H  *  A  V  S  S  P  R  P  A  C  S  T  P  A  S  *  F3
 781 ACTGATCTGGGCACTGAGCTGTGTCCTCTCCGCGCCCTGCCTGCTCTACTCCAGCATCAT 840
     ----:----|----:----|----:----|----:----|----:----|----:----|

     D  K  A  V  S  E  V  F  P  Q  Y  I  Y  D  L  A  G  P  T  K    F1
      T  K  Q  *  V  K  C  F  H  N  I  Y  M  T  L  P  A  P  Q  S   F2
       Q  S  S  K  *  S  V  S  T  I  Y  I  *  P  C  R  P  H  K  V  F3
 841 GACAAAGCAGTAAGTGAAGTGTTTCCACAATATATATATGACCTTGCCGGCCCCACAAAG 900
     ----:----|----:----|----:----|----:----|----:----|----:----|

     S  A  A  A  S  G  S  S  K  G  N  N  R  R  I  F  C  L  F  S    F1
      Q  Q  R  Q  V  A  A  A  R  A  T  T  A  A  Y  F  V  Y  F  P   F2
       S  S  G  K  W  Q  Q  Q  G  Q  Q  P  P  H  I  L  F  I  F  R  F3
 901 TCAGCAGCGGCAAGTGGCAGCAGCAAGGGCAACAACCGCCGCATATTTTGTTTATTTTCC 960
     ----:----|----:----|----:----|----:----|----:----|----:----|

     A  F  P  T  Y  Q  P  S  N  K  H  K  E  L  C  K  T  K  A  Q    F1
      L  F  L  P  T  N  Q  V  I  N  I  K  N  F  V  R  R  K  L  K   F2
       F  S  Y  L  P  T  K  *  *  T  *  R  T  L  *  D  E  S  S  R  F3
 961 GCTTTTCCTACCTACCAACCAAGTAATAAACATAAAGAACTTTGTAAGACGAAAGCTCAA 1020
     ----:----|----:----|----:----|----:----|----:----|----:----|

     D  T  F  H  V  D  T  F  S  H  E  R  T  T  C  S  A  V  A  R    F1
      I  L  F  T  *  T  L  S  H  T  S  A  P  P  A  A  Q  S  P  A   F2
       Y  F  S  R  R  H  F  L  T  R  A  H  H  L  Q  R  S  R  P  L  F3
1021 GATACTTTTCACGTAGACACTTTCTCACACGAGCGCACCACCTGCAGCGCAGTCGCCCGC 1080
     ----:----|----:----|----:----|----:----|----:----|----:----|

     C  V  P  C  P  T  C  T  S  I  L  I  D  S  L  M  P  L  S  A    F1
      V  Y  P  A  P  H  V  R  V  F  S  S  I  R  *  C  P  Y  P  Q   F2
       C  T  L  P  H  M  Y  E  Y  S  H  R  F  A  D  A  P  I  R  N  F3
1081 TGTGTACCCTGCCCCACATGTACGAGTATTCTCATCGATTCGCTGATGCCCCTATCCGCA 1140
     ----:----|----:----|----:----|----:----|----:----|----:----|

     T  C  C  I  P  H  Q  T  Y  I  L  G  N  I  M  Q  R  R  R  S    F1
      H  V  V  Y  P  I  R  P  T  Y  *  A  T  L  C  K  G  V  D  L   F2
       M  L  Y  T  P  S  D  L  H  I  R  Q  H  Y  A  K  A  S  I  *  F3
1141 ACATGTTGTATACCCCATCAGACCTACATATTAGGCAACATTATGCAAAGGCGTCGATCT 1200
     ----:----|----:----|----:----|----:----|----:----|----:----|

     R  H  L  K  L  N  K  P  S  N  L  T  F  I  *  I  L  R  K  F    F1
      G  T  *  S  *  T  S  Q  V  T  *  L  S  F  E  F  Y  A  N  L   F2
       A  P  E  A  K  Q  A  K  *  L  N  F  H  L  N  F  T  Q  I  C  F3
1201 AGGCACCTGAAGCTAAACAAGCCAAGTAACTTAACTTTCATTTGAATTTTACGCAAATTT 1260
     ----:----|----:----|----:----|----:----|----:----|----:----|

     A  H  F  N  E  P  N  G  T  F  T  *  *  N  *  S  I  F  Y  Y    F1
      L  T  S  M  N  Q  M  A  H  S  P  N  R  I  E  A  F  F  I  I   F2
       S  L  Q  *  T  K  W  H  I  H  L  I  E  L  K  H  F  L  L  F  F3
1261 GCTCACTTCAATGAACCAAATGGCACATTCACCTAATAGAATTGAAGCATTTTTTATTAT 1320
     ----:----|----:----|----:----|----:----|----:----|----:----|

     L  A  C  L  T  A  *  L  T  G  *  Q  P  G  W  L  A  Q  S  A    F1
      *  P  A  *  L  P  D  *  L  A  D  S  L  V  G  S  H  R  A  L   F2
       S  L  L  D  C  L  T  D  W  L  T  A  W  L  A  R  T  E  R  S  F3
1321 TTAGCCTGCTTGACTGCCTGACTGACTGGCTGACAGCCTGGTTGGCTCGCACAGAGCGCT 1380
     ----:----|----:----|----:----|----:----|----:----|----:----|

     L  D  I  K  R  C  L  G  *  *  T  I  A  L  A  N  A  S  L  Y    F1
```

```
      L  T  *  N  V  A  *  G  N  R  L  *  L  *  Q  M  Q  V  Y  I    F2
       *  H  K  T  L  P  R  V  I  D  Y  S  S  S  K  C  K  F  I  Y   F3
1381 CTTGACATAAAACGTTGCCTAGGGTAATAGACTATAGCTCTAGCAAATGCAAGTTTATAT 1440
     ----:----|----:----|----:----|----:----|----:----|----:----|

     I  E  I  F  I  T  L  W  E  S  N  L  Y  L  S  G  D  T  S  I     F1
      S  K  Y  L  *  H  Y  G  S  L  I  C  T  *  V  E  I  Q  A  S    F2
       R  N  I  Y  N  I  M  G  V  *  S  V  P  K  W  R  Y  K  H  H   F3
1441 ATCGAAATATTTATAACATTATGGGAGTCTAATCTGTACCTAAGTGGAGATACAAGCATC 1500
     ----:----|----:----|----:----|----:----|----:----|----:----|

     I  D  K  V  K  *  G  K  M  C  K  Y  W  A  *  N  K  A  L  S     F1
      *  T  K  L  N  K  V  K  C  V  N  I  G  P  E  I  K  H  F  P    F2
       R  Q  S  *  I  R  *  N  V  *  I  L  G  L  K  *  S  T  F  L   F3
1501 ATAGACAAAGTTAAATAAGGTAAAATGTGTAAATATTGGGCCTGAAATAAAGCACTTTCC 1560
     ----:----|----:----|----:----|----:----|----:----|----:----|

     L  R  F  A  L  *  E  K  I  *  T  V  L  *  Y  D  Q  A  Y  V     F1
      *  D  L  R  Y  K  R  K  F  E  L  C  F  S  M  T  R  H  M  Y    F2
       K  I  C  V  I  R  E  N  L  N  C  A  L  V  *  P  G  I  C  M   F3
1561 TTAAGATTTGCGTTATAAGAGAAAATTTGAACTGTGCTTTAGTATGACCAGGCATATGTA 1620
     ----:----|----:----|----:----|----:----|----:----|----:----|

     *  V  K  L  N  E  M  C  T  Y  I  Y  N  I  T  T  F  C  N  S     F1
      E  *  S  *  M  K  C  A  H  T  Y  I  I  *  Q  L  F  V  I  L    F2
       S  E  V  K  *  N  V  H  I  H  I  *  Y  N  N  F  L  *  F  L   F3
1621 TGAGTGAAGTTAAATGAAATGTGCACATACATATATAATATAACAACTTTTTGTAATTCT 1680
     ----:----|----:----|----:----|----:----|----:----|----:----|

     *  F  L  D  L  K  N  I  L  K  Q  V  R  G  S  S  R  E  L  P     F1
      N  S  *  T  *  K  I  Y  L  N  K  *  E  V  L  V  G  S  S  R    F2
       I  L  R  P  E  K  Y  T  *  T  S  K  R  F  *  S  G  A  P  D   F3
1681 TAATTCTTAGACCTGAAAAATATACTTAAACAAGTAAGAGGTTCTAGTCGGGAGCTCCCG 1740
     ----:----|----:----|----:----|----:----|----:----|----:----|

     T  R  G  Y  P  E  P  S  S  S  N  I  K  C  I  Y  I  Y  I  F     F1
      L  G  D  T  L  N  P  L  L  P  T  S  N  A  Y  I  Y  I  Y  S    F2
       *  G  I  P  *  T  L  F  F  Q  H  Q  M  H  I  Y  I  Y  I  L   F3
1741 ACTAGGGGATACCCTGAACCCTCTTCTTCCAACATCAAATGCATATATATATATATATTC 1800
     ----:----|----:----|----:----|----:----|----:----|----:----|

     Y  F  R  S  N  M  S  C  F  V  N  L  A  L  I  I  Y  Q  K  T     F1
      I  L  E  A  T  C  H  V  S  *  I  *  L  L  L  F  T  K  K  Q    F2
       F  *  K  Q  H  V  M  F  R  E  S  S  S  Y  Y  L  P  K  N  R   F3
1801 TATTTTAGAAGCAACATGTCATGTTTCGTGAATCTAGCTCTTATTATTTACCAAAAAACA 1860
     ----:----|----:----|----:----|----:----|----:----|----:----|

     G  Y  R  Y  R  L  P  G  N  G  V  R  Y  R  L  S  E  T  N  L     F1
      D  I  D  I  D  C  L  E  T  E  *  D  I  D  Y  R  K  Q  T  *    F2
       I  *  I  *  I  A  W  K  R  S  K  I  S  I  I  G  N  K  L  D   F3
1861 GGATATAGATATAGATTGCCTGGAAACGGAGTAAGATATCGATTATCGGAAACAAACTTG 1920
     ----:----|----:----|----:----|----:----|----:----|----:----|

     I  C  A  G  T  R  S  T  Y  I  Q  N  F  K  S  L  A  H  I  G     F1
      S  A  Q  A  L  G  A  P  T  S  K  I  S  S  L  *  L  I  *  V    F2
       L  R  R  H  *  E  H  L  H  P  K  F  Q  V  F  S  S  Y  R  F   F3
1921 ATCTGCGCAGGCACTAGGAGCACCTACATCCAAAATTTCAAGTCTTTAGCTCATATAGGT 1980
     ----:----|----:----|----:----|----:----|----:----|----:----|

     S  D  S  L  R  S  Y  I  R  T  D  G  Q  T  D  G  H  G  *  I     F1
      L  I  P  C  V  H  T  Y  G  R  T  D  R  Q  T  D  M  A  R  S    F2
       *  F  L  A  F  I  H  T  D  G  R  T  D  R  R  T  W  L  D  R   F3
1981 TCTGATTCCTTGCGTTCATACATACGGACGGACGGACAGACAGACGGACATGGCTAGATC 2040
```

```
     ----:----|----:----|----:----|----:----|----:----|----:----|

     D  S  A  I  D  A  D  *  E  Y  I  Y  F  M  G  P  E  M  L  R    F1
      T  R  L  L  M  L  I  K  S  I  Y  T  L  W  G  R  R  C  F  V   F2
       L  G  Y  *  C  *  L  R  V  Y  I  L  Y  G  A  G  D  A  S  F  F3
2041 GACTCGGCTATTGATGCTGATTAAGAGTATATATACTTTATGGGGCCGGAGATGCTTCGT 2100
     ----:----|----:----|----:----|----:----|----:----|----:----|

     S  A  C  Y  K  H  L  D  F  A  Q  I  Q  Y  T  L  I  P  I  F    F1
      L  P  V  T  S  I  W  I  L  H  K  Y  N  I  P  L  Y  P  F  L   F2
       C  L  L  Q  A  F  G  F  C  T  N  T  I  Y  P  Y  T  H  F  Y  F3
2101 TCTGCCTGTTACAAGCATTTGGATTTTGCACAAATACAATATACCCTTATACCCATTTTT 2160
     ----:----|----:----|----:----|----:----|----:----|----:----|

     I  G  F  R  V  *  N  *  L  S  N  T  S  H  E  *  *  *  L  I    F1
      L  G  S  G  Y  K  I  S  *  A  T  Q  V  T  N  D  D  S  *  L   F2
       W  V  Q  G  I  K  L  V  E  Q  H  K  S  R  M  M  I  A  N  Y  F3
2161 ATTGGGTTCAGGGTATAAAATTAGTTGAGCAACACAAGTCACGAATGATGATAGCTAATT 2220
     ----:----|----:----|----:----|----:----|----:----|----:----|

     I  E  K  Y  I  I  N  I  Y  I  H  I  H  I  Y  I  Y  I  V  V    F1
      L  K  S  I  *  *  I  Y  T  Y  I  Y  T  Y  I  Y  I  L  L  W   F2
       *  K  V  Y  N  K  Y  I  H  T  Y  T  H  I  Y  I  Y  C  C  G  F3
2221 ATTGAAAAGTATATAATAAATATATACATACATATACACATATATATATATATTGTTGTG 2280
     ----:----|----:----|----:----|----:----|----:----|----:----|

     E  K  Y  N  *  Q  I  *  V  S  H  I  *  H  *  W  R  *  T  N    F1
      R  N  I  T  S  R  Y  K  L  V  I  Y  N  T  D  G  D  K  Q  I   F2
       E  I  *  L  A  D  I  S  *  S  Y  I  T  L  M  E  I  N  K  S  F3
2281 GAGAAATATAACTAGCAGATATAAGTTAGTCATATATAACACTGATGGAGATAAACAAAT 2340
     ----:----|----:----|----:----|----:----|----:----|----:----|

     H  I  S  A  K  S  S  N  C  Y  C  *  S  R  V  Q  L  N  R  F    F1
      I  F  Q  R  N  H  L  I  A  I  V  E  V  A  F  N  *  I  D  L   F2
       Y  F  S  E  I  I  *  L  L  L  L  K  S  R  S  T  E  *  I  *  F3
2341 CATATTTCAGCGAAATCATCTAATTGCTATTGTTGAAGTCGCGTTCAACTGAATAGATTT 2400
     ----:----|----:----|----:----|----:----|----:----|----:----|

     K  Y  N  E  L  N  T  A  E  N  S  I  D  L  P  S  A  H  A  G    F1
      S  T  T  S  *  I  P  Q  R  I  Q  *  I  C  H  L  L  M  Q  V   F2
       V  Q  R  V  E  Y  R  R  E  F  N  R  S  A  I  C  S  C  R  *  F3
2401 AAGTACAACGAGTTGAATACCGCAGAGAATTCAATAGATCTGCCATCTGCTCATGCAGGT 2460
     ----:----|----:----|----:----|----:----|----:----|----:----|

     D  R  *  L  E  Q  F  K  C  S  S  W  P  T  V  R  P  S  N  Y    F1
      T  G  N  W  S  N  L  N  A  R  V  G  R  Q  F  A  Q  V  I  T   F2
       Q  V  T  G  A  I  *  M  L  E  L  A  D  S  S  P  K  *  L  P  F3
2461 GACAGGTAACTGGAGCAATTTAAATGCTCGAGTTGGCCGACAGTTCGCCCAAGTAATTAC 2520
     ----:----|----:----|----:----|----:----|----:----|----:----|

     L  T  A  S  C  I  V  N  G  S  R  I  G  C  *  Q  I  V  T  D    F1
      *  L  P  V  A  L  L  M  G  P  E  L  V  V  N  K  L  S  P  M   F2
       D  C  Q  L  H  C  *  W  V  P  N  W  L  L  T  N  C  H  R  W  F3
2521 CTGACTGCCAGTTGCATTGTTAATGGGTCCCGAATTGGTTGTTAACAAATTGTCACCGAT 2580
     ----:----|----:----|----:----|----:----|----:----|----:----|

     G  D  N  S  F  K  C  A  R  H  D  A  *  K  L  R  *  F  S  K    F1
      G  T  T  L  L  S  A  H  A  T  T  H  K  S  C  D  N  F  P  N   F2
       G  Q  L  F  *  V  R  T  P  R  R  I  K  A  A  I  I  F  Q  I  F3
2581 GGGGACAACTCTTTTAAGTGCGCACGCCACGACGCATAAAAGCTGCGATAATTTTCCAAA 2640
     ----:----|----:----|----:----|----:----|----:----|----:----|

     L  G  F  H  F  A  I  K  W  A  *  I  N  S  M  L  K  C  C  K    F1
```

```
      *  V  F  I  L  P  S  N  G  H  E  *  I  L  C  *  N  A  V  K    F2
       R  F  S  F  C  H  Q  M  G  M  N  K  F  Y  A  K  M  L  *  K   F3
2641 TTAGGTTTTCATTTTGCCATCAAATGGGCATGAATAAATTCTATGCTAAAATGCTGTAAA 2700
     ----:----|----:----|----:----|----:----|----:----|----:----|

     N  S  F  Q  H  V  A  F  R  L  D  A  S  R  L  T  L  R  R  H     F1
      T  P  F  N  M  L  L  S  V  W  T  L  L  V  *  L  *  G  V  I    F2
       L  L  S  T  C  C  F  P  S  G  R  F  S  F  N  F  E  A  S  *   F3
2701 AACTCCTTTCAACATGTTGCTTTCCGTCTGGACGCTTCTCGTTTAACTTTGAGGCGTCAT 2760
     ----:----|----:----|----:----|----:----|----:----|----:----|

     N  I  *  G  A  L  W  F  R  F  A  F  R  N  *  I  M  L  K  L     F1
      I  F  K  G  H  F  G  L  G  S  L  S  V  I  K  *  C  *  N  C    F2
       Y  L  R  G  T  L  V  *  V  R  F  P  *  L  N  N  V  E  T  V   F3
2761 AATATTTAAGGGGCACTTTGGTTTAGGTTCGCTTTCCGTAATTAAATAATGTTGAAACTG 2820
     ----:----|----:----|----:----|----:----|----:----|----:----|

     L  G  A  T  K  T  A  E  A  P  L  C  I  I  *  N  S  L  R  R     F1
      W  G  P  Q  K  R  Q  K  L  L  Y  A  L  Y  K  I  H  L  G  A    F2
       G  G  H  K  N  G  R  S  S  F  M  H  Y  I  K  F  T  *  A  R   F3
2821 TTGGGGGCCACAAAAACGGCAGAAGCTCCTTTATGCATTATATAAAATTCACTTAGGCGC 2880
     ----:----|----:----|----:----|----:----|----:----|----:----|

     G  E  S  F  V  Y  L  K  I  *  N  N  L  P  Q  S  R  *  R  *     F1
      A  K  V  L  F  T  L  K  F  K  I  I  Y  P  S  R  D  N  G  R    F2
       R  K  F  C  L  P  *  N  L  K  *  F  T  P  V  E  I  T  V  D   F3
2881 GGCGAAAGTTTTGTTTACCTTAAAATTTAAAATAATTTACCCCAGTCGAGATAACGGTAG 2940
     ----:----|----:----|----:----|----:----|----:----|----:----|

     M  I  Y  N  Y  L  T  V  I  T  T  E  N  P  V  Q  F  A  S  *     F1
      *  F  I  I  T  *  Q  L  L  Q  R  K  I  P  Y  S  L  L  H  D    F2
       D  L  *  L  L  D  S  Y  Y  N  G  K  S  R  T  V  C  F  M  M   F3
2941 ATGATTTATAATTACTTGACAGTTATTACAACGGAAAATCCCGTACAGTTTGCTTCATGA 3000
     ----:----|----:----|----:----|----:----|----:----|----:----|

     C  G  P  M  D  V  I  Q  H  R  *  Q  T  M  C  K  Y  S  N  D     F1
      V  A  R  W  T  L  S  N  I  D  D  R  L  C  V  S  I  R  M  M    F2
       W  P  D  G  R  Y  P  T  S  M  T  D  Y  V  *  V  F  E  *  C   F3
3001 TGTGGCCCGATGGACGTTATCCAACATCGATGACAGACTATGTGTAAGTATTCGAATGAT 3060
     ----:----|----:----|----:----|----:----|----:----|----:----|

     V  Y  H  Y  *  I  I  *  K  C  L  P  T  T  S  Y  N  V  T  I     F1
      F  T  I  I  K  L  S  E  N  V  Y  L  L  P  A  I  M  *  P  Y    F2
       L  P  L  L  N  Y  L  K  M  F  T  Y  Y  Q  L  *  C  D  H  I   F3
3061 GTTTACCATTATTAAATTATCTGAAAATGTTTACCTACTACCAGCTATAATGTGACCATA 3120
     ----:----|----:----|----:----|----:----|----:----|----:----|

     L  V  L  T  Y  G  I  P  M  I  V  M  L  I  C  Y  S  L  M  G     F1
      W  Y  *  H  M  A  Y  P  *  L  L  C  L  Y  A  T  R  L  W  V    F2
       G  I  N  I  W  H  T  H  D  C  Y  A  Y  M  L  L  A  Y  G  S   F3
3121 TTGGTATTAACATATGGCATACCCATGATTGTTATGCTTATATGCTACTCGCTTATGGGT 3180
     ----:----|----:----|----:----|----:----|----:----|----:----|

     R  V  L  W  G  S  R  S  I  G  E  N  T  D  R  Q  M  E  S  M     F1
      E  Y  S  G  A  V  A  L  L  G  R  T  R  I  V  R  W  N  R  *    F2
       S  T  L  G  Q  S  L  Y  W  G  E  H  G  S  S  D  G  I  D  E   F3
3181 CGAGTACTCTGGGGCAGTCGCTCTATTGGGGAGAACACGGATCGTCAGATGGAATCGATG 3240
     ----:----|----:----|----:----|----:----|----:----|----:----|

     K  S  K  R  K  V  V  R  M  F  I  A  I  V  S  I  F  A  I  C     F1
      N  Q  S  E  R  W  F  A  C  L  *  R  L  S  P  S  S  P  F  A    F2
       I  K  A  K  G  G  S  H  V  Y  S  D  C  L  H  L  R  H  L  L   F3
3241 AAATCAAAGCGAAAGGTGGTTCGCATGTTTATAGCGATTGTCTCCATCTTCGCCATTTGC 3300
```

```
     ----:----|----:----|----:----|----:----|----:----|----:----|

     W  L  P  Y  H  L  F  F  I  Y  A  Y  H  N  N  Q  V  A  S  T     F1
      G  C  P  I  I  C  S  S  S  M  P  I  T  T  I  R  W  H  P  Q    F2
       A  A  L  S  S  V  L  H  L  C  L  S  Q  Q  S  G  G  I  H  K   F3
3301 TGGCTGCCCTATCATCTGTTCTTCATCTATGCCTATCACAACAATCAGGTGGCATCCACA 3360
     ----:----|----:----|----:----|----:----|----:----|----:----|

     K  Y  V  Q  H  M  Y  L  G  F  Y  W  L  A  M  S  N  A  M  V     F1
      S  T  C  S  T  C  I  W  A  S  T  G  W  P  C  P  M  Q  W  *    F2
       V  R  A  A  H  V  S  G  L  L  L  A  G  H  V  Q  C  N  G  E   F3
3361 AAGTACGTGCAGCACATGTATCTGGGCTTCTACTGGCTGGCCATGTCCAATGCAATGGTG 3420
     ----:----|----:----|----:----|----:----|----:----|----:----|

     N  P  I  I  Y  Y  W  M  N  K  R  *  D  I  I  *  *  Y  I  *     F1
      I  P  *  F  I  T  G  *  T  R  G  E  I  *  Y  N  N  I  Y  R    F2
       S  H  N  L  L  L  D  E  Q  E  V  R  Y  N  I  I  I  Y  I  D   F3
3421 AATCCCATAATTTATTACTGGATGAACAAGAGGTGAGATATAATATAATAATATATATAG 3480
     ----:----|----:----|----:----|----:----|----:----|----:----|

     I  Y  N  I  Q  M  T  K  T  S  S  N  I  F  M  R  N  *  A  Q     F1
      Y  I  I  Y  K  *  L  K  H  L  Q  T  Y  L  *  E  I  K  H  K    F2
       I  *  Y  T  N  D  *  N  I  F  K  H  I  Y  E  K  L  S  T  K   F3
3481 ATATATAATATACAAATGACTAAAACATCTTCAAACATATTTATGAGAAATTAAGCACAA 3540
     ----:----|----:----|----:----|----:----|----:----|----:----|

     N  N  T  K  N  *  K  L  *  K  A  N  Q  I  *  D  *  S  K  N     F1
      T  T  Q  K  I  K  S  C  E  K  Q  I  K  Y  K  I  E  A  K  T    F2
       Q  H  K  K  L  K  V  V  K  S  K  S  N  I  R  L  K  Q  K  Q   F3
3541 AACAACACAAAAAAATTAAAAGTTGTGAAAAGCAAATCAAATATAAGATTGAAGCAAAAAC 3600
     ----:----|----:----|----:----|----:----|----:----|----:----|

     S  N  L  N  V  M  N  Q  *  K  Q  K  R  T  E  C  R  R  F  N     F1
      A  I  *  M  *  *  I  S  K  N  R  K  E  R  N  A  D  D  L  I    F2
       Q  F  K  C  N  E  S  V  K  T  E  K  N  G  M  P  T  I  *  Y   F3
3601 AGCAATTTAAATGTAATGAATCAGTAAAAACAGAAAAGAACGGAATGCCGACGATTTAAT 3660
     ----:----|----:----|----:----|----:----|----:----|----:----|

     T  L  V  V  M  G  H  G  N  S  D  I  I  I  S  S  T  E  S  S     F1
      P  L  W  S  W  V  M  E  I  V  I  L  L  *  V  L  P  N  L  P    F2
       P  C  G  H  G  S  W  K  *  *  Y  Y  Y  K  F  Y  R  I  F  L   F3
3661 ACCCTTGTGGTCATGGGTCATGGAAATAGTGATATTATTATAAGTTCTACCGAATCTTCC 3720
     ----:----|----:----|----:----|----:----|----:----|----:----|

     F  M  V  G  Y  S  G  S  M  L  M  Y  S  K  S  G  *  N  I  *     F1
      L  W  L  D  I  V  V  R  C  *  C  I  V  K  V  V  K  I  F  K    F2
       Y  G  W  I  *  W  F  D  V  N  V  *  *  K  W  L  K  Y  L  K   F3
3721 TTTATGGTTGGATATAGTGGTTCGATGTTAATGTATAGTAAAAGTGGTTAAAATATTTAA 3780
     ----:----|----:----|----:----|----:----|----:----|----:----|

     K  S  K  K  F  A  Y  A  R  T  *  F  L  N  N  M  I  Y  L  I     F1
      K  A  K  N  L  L  M  Q  E  L  N  F  *  I  I  *  Y  I  L  *    F2
       K  Q  K  I  C  L  C  K  N  L  I  F  E  *  Y  D  I  S  Y  R   F3
3781 AAAAGCAAAAAAATTTGCTTATGCAAGAACTTAATTTTTGAATAATATGATATATCTTATA 3840
     ----:----|----:----|----:----|----:----|----:----|----:----|

     E  V  K  T  L  R  F  L  *  S  Y  K  T  T  W  I  I  S  Q  N     F1
      K  S  K  L  S  D  S  Y  S  P  T  R  L  H  G  *  F  L  R  I    F2
       S  Q  N  S  P  I  P  I  V  L  Q  D  Y  M  D  N  F  S  E  Y   F3
3841 GAAGTCAAAACTCTCCGATTCCTATAGTCCTACAAGACTACATGGATAATTTCTCAGAAT 3900
     ----:----|----:----|----:----|----:----|----:----|----:----|

     I  I  C  F  I  G  S  I  R  V  *  *  T  I  K  K  *  N  T  N     F1
```

```
      S  Y  A  L  *  D  L  *  G  Y  N  K  Q  L  K  N  K  T  R  I     F2
       H  M  L  Y  R  I  Y  K  G  I  I  N  N  *  K  I  K  H  E  L    F3
3901 ATCATATGCTTTATAGGATCTATAAGGGTATAATAAACAATTAAAAAATAAAACACGAAT 3960
     ----:----|----:----|----:----|----:----|----:----|----:----|

     *  *  *  G  L  Q  K  Y  N  I  I  I  I  F  N  L  L  V  F  S     F1
      N  S  E  A  C  K  N  T  I  *  *  L  F  S  T  C  L  F  F  L    F2
       I  V  R  L  A  K  I  Q  Y  N  N  Y  F  Q  P  A  C  F  F  F   F3
3961 TAATAGTGAGGCTTGCAAAAATACAATATAATAATTATTTTCAACCTGCTTGTTTTTTCT 4020
     ----:----|----:----|----:----|----:----|----:----|----:----|

     F  I  D  F  A  C  T  F  N  G  *  S  S  A  V  A  W  G  L  C     F1
      L  *  I  S  H  V  L  S  T  D  N  L  L  L  L  L  G  A  Y  A    F2
       Y  R  F  R  M  Y  F  Q  R  I  I  F  C  C  C  L  G  L  M  R   F3
4021 TTTATAGATTTCGCATGTACTTTCAACGGATAATCTTCTGCTGTTGCTTGGGGCTTATGC 4080
     ----:----|----:----|----:----|----:----|----:----|----:----|

     A  I  D  L  S  R  R  R  A  G  W  P  T  R  T  A  R  I  A  I     F1
      L  *  I  *  V  A  E  E  Q  D  G  Q  Q  E  Q  L  E  S  P  Y    F2
       Y  R  F  E  S  P  K  S  R  M  A  N  K  N  S  S  N  R  H  T   F3
4081 GCTATAGATTTGAGTCGCCGAAGAGCAGGATGGCCAACAAGAACAGCTCGAATCGCCATA 4140
     ----:----|----:----|----:----|----:----|----:----|----:----|

     Q  E  V  R  T  K  G  H  C  K  Y  L  Y  H  V  I  F  P  I  G     F1
      K  R  Y  V  R  R  D  T  V  S  I  S  I  M  *  F  S  Q  S  E    F2
       R  G  T  Y  E  G  T  L  *  V  S  L  S  C  D  F  P  N  R  N   F3
4141 CAAGAGGTACGTACGAAGGGACACTGTAAGTATCTCTATCATGTGATTTTCCCAATCGGA 4200
     ----:----|----:----|----:----|----:----|----:----|----:----|

     M  C  S  R  N  Q  K  P  V  E  A  *  H  N  G  N  T  N  P  A     F1
      C  A  A  E  T  K  S  Q  W  K  R  S  T  M  E  T  Q  I  Q  Q    F2
       V  Q  P  K  P  K  A  S  G  S  V  A  Q  W  K  H  K  S  S  R   F3
4201 ATGTGCAGCCGAAACCAAAAGCCAGTGGAAGCGTAGCACAATGGAAACACAAATCCAGCA 4260
     ----:----|----:----|----:----|----:----|----:----|----:----|

     D  A  Q  N  F  V  P  R  *  G  C  G  C  P  G  S  *  P  H  G     F1
      M  P  K  T  S  S  R  D  K  D  A  G  V  Q  G  L  N  H  T  A    F2
       C  P  K  L  R  P  A  I  R  M  R  V  S  R  V  L  T  T  R  P   F3
4261 GATGCCCAAAACTTCGTCCCGCGATAAGGATGCGGGTGTCCAGGGTCTTAACCACACGGC 4320
     ----:----|----:----|----:----|----:----|----:----|----:----|

     R  R  V  H  N  R  A  T  H  R  *  Q  *  L  P  H  L  L  V  H     F1
      V  E  C  I  I  E  R  P  I  D  D  N  S  S  P  I  C  L  S  I    F2
       S  S  A  *  *  S  D  P  *  M  T  I  A  P  P  F  A  C  P  L   F3
4321 CGTCGAGTGCATAATAGAGCGACCCATAGATGACAATAGCTCCCCCATTTGCTTGTCCAT 4380
     ----:----|----:----|----:----|----:----|----:----|----:----|

     *  K  Q  C  R  R  A  A  A  R  K  N  *  I  H  F  L  R  *  G     F1
      K  N  S  A  G  E  R  Q  R  V  K  I  K  Y  I  S  C  D  E  D    F2
       K  T  V  P  A  S  G  S  A  *  K  L  N  T  F  P  A  M  R  I   F3
4381 TAAAAACAGTGCCGGCGAGCGGCAGCGCGTAAAAATTAAATACATTTCCTGCGATGAGGA 4440
     ----:----|----:----|----:----|----:----|----:----|----:----|

     *  Q  S  H  R  G  G  I  R  K  Q  *  Q  S  R  F  Q  S  Q  S     F1
      N  N  P  I  E  E  G  S  E  N  N  S  S  H  D  S  N  H  S  H    F2
       T  I  P  S  R  R  D  P  K  T  I  A  V  T  I  P  I  T  V  T   F3
4441 TAACAATCCCATCGAGGAGGGATCCGAAAACAATAGCAGTCACGATTCCAATCACAGTCA 4500
     ----:----|----:----|----:----|----:----|----:----|----:----|

     R  P  R  S  Q  L  R  P  Q  P  W  P  K  C  E  S  N  S  T  T     F1
      G  H  G  H  S  C  G  R  S  H  G  Q  N  A  K  A  I  Q  Q  L    F2
       A  T  V  T  V  A  A  A  A  M  A  K  M  R  K  Q  F  N  N  C   F3
4501 CGGCCACGGTCACAGTTGCGGCCGCAGCCATGGCCAAAATGCGAAAGCAATTCAACAACT 4560
```

```
     ----:----|----:----|----:----|----:----|----:----|----:----|

     V  M  G  K  F  L  *  G  F  A  L  K  R  X                        F1
      *  W  E  S  S  Y  R  V  S  L  *  K  E                          F2
       D  G  K  V  P  I  G  F  R  F  K  K  X                         F3
4561 GTGATGGGAAAGTTCCTATAGGGTTTCGCTTTAAAAAGAG 4600
     ----:----|----:----|----:----|----:----|
```
