## Supplementary material for "The incidence of candidate binding sites for β-arrestin in Drosophila neuropeptide GPCRs": S6 Text

***D virilis* gDNA report for CAPA R**

[https://www.ncbi.nlm.nih.gov/nuccore/1805215609?report=graph&v=8515583:8519076](https://www.ncbi.nlm.nih.gov/nuccore/1805215609?report=graph&v=8515583:8519076)

In *D melanogaster*, two isoforms are predicted. PB and PC are predicted: PC is a truncation due to an alternative splice to a location with potential for a single AA (V) prior to a termination. This splice acceptor immediately precedes the one for the full length PB isoform. Here I search for evidence of a PC isoform in *D virilis*.

The splice acceptor site around bp 1201 does not present an obvious candidate for a PC isoform.

gDNA fasta
[https://www.ncbi.nlm.nih.gov/nuccore/NW_022587564.1?report=fasta&from=8516083&to=8518578](https://www.ncbi.nlm.nih.gov/nuccore/NW_022587564.1?report=fasta&from=8516083&to=8518578)

bp#1 = 8518051 of the above link

```
translation:
*        L  V  R  A  E  K  S  W  K  A  T  N  M  N  M  N  M  S  T     F1
     D  W  L  G  Q  K  N  L  G  R  Q  R  I  *  I  *  I  *  V  R    F2
      I  G  *  G  R  K  I  L  E  G  N  E  Y  E  Y  E  Y  E  Y  E   F3
  1 TGATTGGTTAGGGCAGAAAAATCTTGGAAGGCAACGAATATGAATATGAATATGAGTACG 60
    ----:----|----:----|----:----|----:----|----:----|----:----|

    N  M  S  M  D  T  N  L  S  T  Y  L  G  T  S  D  A  T  A  L     F1
     I  *  V  W  I  R  T  *  V  H  I  W  A  L  R  M  R  L  H  C    F2
      Y  E  Y  G  Y  E  L  E  Y  I  F  G  H  F  G  C  D  C  I  A   F3
 61 AATATGAGTATGGATACGAACTTGAGTACATATTTGGGCACTTCGGATGCGACTGCATTG 120
    ----:----|----:----|----:----|----:----|----:----|----:----|

    P  Y  P  G  M  D  D  Y  G  C  P  H  M  N  C  T  A  M  E  F     F1
     P  I  L  E  W  T  T  M  D  A  L  T  *  I  V  R  Q  W  S  L    F2
      L  S  W  N  G  R  L  W  M  P  S  H  E  L  Y  G  N  G  V  C   F3
121 CCCTATCCTGGAATGGACGACTATGGATGCCCTCACATGAATTGTACGGCAATGGAGTTT 180
    ----:----|----:----|----:----|----:----|----:----|----:----|

    V  Q  F  V  L  G  P  Q  T  L  P  L  H  K  A  L  L  V  S  L     F1
     F  S  S  Y  S  A  H  K  P  C  R  C  I  R  P  Y  W  *  V  *    F2
      S  V  R  T  R  P  T  N  P  A  A  A  *  G  L  I  G  E  F  R   F3
181 GTTCAGTTCGTACTCGGCCCACAAACCCTGCCGCTGCATAAGGCCTTATTGGTGAGTTTA 240
    ----:----|----:----|----:----|----:----|----:----|----:----|

    D  K  C  S  R  *  R  H  I  N  L  *  T  I  Y  R  *  A  *  F     F1
     I  N  A  V  D  K  D  I  *  T  Y  K  L  F  I  D  K  H  N  F    F2
      *  M  Q  *  I  K  T  Y  K  L  I  N  Y  L  *  I  S  I  I  F   F3
241 GATAAATGCAGTAGATAAAGACATATAAACTTATAAACTATTTATAGATAAGCATAATTT 300
    ----:----|----:----|----:----|----:----|----:----|----:----|

    L  V  A  Y  S  *  L  A  Y  W  E  T  S  W  S  A  W  *  *  F     F1
     *  W  H  I  H  N  W  R  T  G  K  R  P  G  L  H  G  N  N  S    F2
      S  G  I  F  I  T  G  V  L  G  N  V  L  V  C  M  V  I  I  R   F3
301 TTAGTGGCATATTCATAACTGGCGTACTGGGAAACGTCCTGGTCTGCATGGTAATAATTC 360
    ----:----|----:----|----:----|----:----|----:----|----:----|

    G  M  R  Q  C  T  R  Q  R  I  I  T  C  S  V  W  P  C  P  I     F1
```

```
      A  C  G  N  A  H  G  N  E  L  L  P  V  Q  F  G  R  V  R  F     F2
       H  A  A  M  H  T  A  T  N  Y  Y  L  F  S  L  A  V  S  D  L    F3
 361 GGCATGCGGCAATGCACACGGCAACGAATTATTACCTGTTCAGTTTGGCCGTGTCCGATT 420
     ----:----|----:----|----:----|----:----|----:----|----:----|

     C  C  T  Y  F  *  V  G  K  L  I  N  Y  L  T  A  F  N  I  D      F1
      A  V  L  T  F  R  L  V  S  S  *  I  T  *  L  H  L  T  L  I     F2
       L  Y  L  L  L  G  W  *  A  H  K  L  P  N  C  I  *  H  *  F    F3
 421 TGCTGTACTTACTTTTAGGTTGGTAAGCTCATAAATTACCTAACTGCATTTAACATTGAT 480
     ----:----|----:----|----:----|----:----|----:----|----:----|

     F  N  D  D  T  R  I  A  G  G  G  I  P  I  L  A  S  V  S  I      F1
      L  M  M  T  L  G  L  P  A  E  V  F  L  Y  W  H  Q  Y  P  Y     F2
       *  *  *  H  *  D  C  R  R  R  Y  S  Y  T  G  I  S  I  H  I    F3
 481 TTTAATGATGACACTAGGATTGCCGGCGGAGGTATTCCTATACTGGCATCAGTATCCATA 540
     ----:----|----:----|----:----|----:----|----:----|----:----|

     F  I  W  S  A  V  L  Q  V  A  S  I  C  V  G  S  V  S  M  L      F1
      L  F  G  L  P  F  C  K  L  R  A  F  V  S  E  A  *  V  C  C     F2
       Y  L  V  C  R  F  A  S  C  E  H  L  C  R  K  R  K  Y  V  A    F3
 541 TTTATTTGGTCTGCCGTTTTGCAAGTTGCGAGCATTTGTGTCGGAAGCGTAAGTATGTTG 600
     ----:----|----:----|----:----|----:----|----:----|----:----|

     L  W  A  Q  V  K  F  *  L  E  H  N  T  T  G  A  H  M  Y  R      F1
      S  G  R  R  S  N  F  N  *  S  T  I  Q  Q  V  H  I  C  I  G     F2
       L  G  A  G  Q  I  L  I  R  A  Q  Y  N  R  C  T  Y  V  S  V    F3
 601 CTCTGGGCGCAGGTCAAATTTTAATTAGAGCACAATACAACAGGTGCACATATGTATCGG 660
     ----:----|----:----|----:----|----:----|----:----|----:----|

     C  S  Q  L  L  P  S  R  W  N  D  F  W  P  Y  A  I  P  C  T      F1
      V  H  N  C  C  L  L  D  G  T  I  S  G  H  M  P  S  P  A  R     F2
       F  T  I  V  A  F  S  M  E  R  F  L  A  I  C  H  P  L  H  V    F3
 661 TGTTCACAATTGTTGCCTTCTCGATGGAACGATTTCTGGCCATATGCCATCCCCTGCACG 720
     ----:----|----:----|----:----|----:----|----:----|----:----|

     C  V  Q  C  P  A  F  S  G  H  *  G  S  *  R  P  F  G  L  *      F1
      V  C  N  V  R  L  S  A  G  T  E  D  H  N  G  P  L  D  C  K     F2
       C  A  M  S  G  F  Q  R  A  L  R  I  I  T  A  L  W  I  V  S    F3
 721 TGTGTGCAATGTCCGGCTTTCAGCGGGCACTGAGGATCATAACGGCCCTTTGGATTGTAA 780
     ----:----|----:----|----:----|----:----|----:----|----:----|

     V  S  *  V  P  Y  H  S  V  S  R  L  K  F  N  I  *  I  F  Q      F1
      F  P  K  C  H  T  I  R  C  Q  D  *  N  S  I  F  E  F  S  K     F2
       F  L  S  A  I  P  F  G  V  K  T  E  I  Q  Y  L  N  F  P  N    F3
 781 GTTTCCTAAGTGCCATACCATTCGGTGTCAAGACTGAAATTCAATATTTGAATTTTCCAA 840
     ----:----|----:----|----:----|----:----|----:----|----:----|

     T  V  W  F  K  Y  I  V  Y  W  N  P  *  R  L  L  N  T  Y  R      F1
      R  Y  G  L  S  T  L  Y  I  G  I  H  N  D  C  *  T  L  T  D     F2
       G  M  V  *  V  H  C  I  L  E  S  I  T  I  A  E  H  L  Q  M    F3
 841 ACGGTATGGTTTAAGTACATTGTATATTGGAATCCATAACGATTGCTGAACACTTACAGA 900
     ----:----|----:----|----:----|----:----|----:----|----:----|

     W  I  S  Y  T  G  I  R  I  L  F  N  R  I  G  V  S  R  R  I      F1
      G  S  R  I  L  E  S  A  F  C  S  I  E  L  E  F  P  E  E  F     F2
       D  L  V  Y  W  N  P  H  S  V  Q  S  N  W  S  F  P  K  N  S    F3
 901 TGGATCTCGTATACTGGAATCCGCATTCTGTTCAATCGAATTGGAGTTTCCCGAAGAATT 960
     ----:----|----:----|----:----|----:----|----:----|----:----|

     P  L  V  R  S  L  L  L  H  I  F  H  Y  T  H  D  T  H  Y  I      F1
      P  L  F  E  V  S  F  C  I  F  F  I  I  P  M  I  L  I  I  L     F2
       P  C  S  K  S  P  F  A  Y  F  S  L  Y  P  *  Y  S  L  Y  C    F3
 961 CCCCTTGTTCGAAGTCTCCTTTTGCATATTTTTCATTATACCCATGATACTCATTATATT 1020
```

```
     ----:----|----:----|----:----|----:----|----:----|----:----|

     V  I  W  S  H  G  R  S  D  S  F  K  S  H  R  Q  I  G  *  A     F1
      L  Y  G  R  M  G  A  Q  I  R  S  R  A  T  D  K  L  G  K  L    F2
       Y  M  V  A  W  A  L  R  F  V  Q  E  P  P  T  N  W  V  S  L   F3
1021 GTTATATGGTCGCATGGGCGCTCAGATTCGTTCAAGAGCCACCGACAAATTGGGTAAGCT 1080
     ----:----|----:----|----:----|----:----|----:----|----:----|

     *  Q  N  F  K  P  E  R  I  R  E  *  I  Q  F  S  C  R  S  S     F1
      D  K  I  S  S  R  N  A  F  G  N  K  Y  N  S  L  A  G  V  Q    F2
       T  K  F  Q  A  G  T  H  S  G  I  N  T  I  L  L  Q  E  F  S   F3
1081 TGACAAAATTTCAAGCCGGAACGCATTCGGGAATAAATACAATTCTCTTGCAGGAGTTCA 1140
     ----:----|----:----|----:----|----:----|----:----|----:----|

     A  G  F  Q  E  S  R  I  A  *  L  A  E  E  K  A  S  C  Y  T     F1
      Q  G  S  R  N  R  E  S  R  S  S  Q  K  K  K  R  A  V  I  R    F2
       R  V  P  G  I  A  N  R  V  A  R  R  R  K  S  E  L  L  Y  E   F3
1141 GCAGGGTTCCAGGAATCGCGAATCGCGTAGCTCGCAGAAGAAAAAGCGAGCTGTTATACG 1200
     ----:----|----:----|----:----|----:----|----:----|----:----|

     N  V  G  *  G  M  H  I  P  I  S  Q  F  I  L  L  K  S  Y  T     F1
      M  L  G  K  E  C  T  F  L  F  L  N  L  Y  Y  L  K  A  I  P    F2
       C  W  V  R  N  A  H  S  Y  F  S  I  Y  I  T  *  K  L  Y  L   F3
1201 AATGTTGGGTAAGGAATGCACATTCCTATTTCTCAATTTATATTACTTAAAAGCTATACC 1260
     ----:----|----:----|----:----|----:----|----:----|----:----|

     F  K  *  P  P  W  *  S  H  F  S  S  A  G  F  H  F  I  C  K     F1
      L  N  S  R  R  G  N  H  I  F  R  L  L  V  S  I  S  S  A  K    F2
       *  I  A  A  V  V  I  T  F  F  V  C  W  F  P  F  H  L  Q  R   F3
1261 TTTAAATAGCCGCCGTGGTAATCACATTTTTCGTCTGCTGGTTTCCATTTCATCTGCAAA 1320
     ----:----|----:----|----:----|----:----|----:----|----:----|

     D  C  G  F  C  T  P  R  T  L  P  T  I  R  M  *  M  N  G  C     F1
      I  V  V  S  V  R  Q  E  H  C  Q  L  S  G  C  E  *  M  A  V    F2
       L  W  F  L  Y  A  K  N  I  A  N  Y  Q  D  V  N  E  W  L  F   F3
1321 GATTGTGGTTTCTGTACGCCAAGAACATTGCCAACTATCAGGATGTGAATGAATGGCTGT 1380
     ----:----|----:----|----:----|----:----|----:----|----:----|

     S  P  *  R  D  S  R  T  M  S  L  G  E  H  L  Q  L  F  *  T     F1
      L  H  S  G  I  R  V  L  C  L  L  V  S  I  C  S  Y  F  K  L    F2
       S  I  A  G  F  A  Y  Y  V  S  W  *  A  F  A  A  I  L  N  *   F3
1381 TCTCCATAGCGGGATTCGCGTACTATGTCTCTTGGTGAGCATTTGCAGCTATTTTAAACT 1440
     ----:----|----:----|----:----|----:----|----:----|----:----|

     N  I  N  L  S  L  I  I  D  S  L  F  L  T  S  T  I  N  P  I     F1
      I  L  I  *  V  *  *  L  I  H  C  F  S  P  A  Q  L  I  R  L    F2
       Y  *  S  K  F  N  N  *  F  I  V  S  H  Q  H  N  *  S  D  C   F3
1441 AATATTAATCTAAGTTTAATAATTGATTCATTGTTTCTCACCAGCACAATTAATCCGATT 1500
     ----:----|----:----|----:----|----:----|----:----|----:----|

     V  Y  N  V  M  S  Q  R  Y  R  V  A  F  K  E  I  L  C  G  K     F1
      C  T  M  S  C  H  S  A  I  G  L  P  S  R  R  Y  S  V  A  K    F2
       V  Q  C  H  V  T  A  L  S  G  C  L  Q  G  D  T  L  W  Q  K   F3
1501 GTGTACAATGTCATGTCACAGCGCTATCGGGTTGCCTTCAAGGAGATACTCTGTGGCAAA 1560
     ----:----|----:----|----:----|----:----|----:----|----:----|

     K  A  G  A  Y  Y  N  S  G  F  A  R  D  Q  S  S  F  I  R  D     F1
      R  L  G  P  I  T  T  A  D  L  P  E  I  N  R  V  S  L  G  M    F2
       G  W  G  L  L  Q  Q  R  I  C  Q  R  S  I  E  F  H  *  G  *   F3
1561 AAGGCTGGGGCCTATTACAACAGCGGATTTGCCAGAGATCAATCGAGTTTCATTAGGGAT 1620
     ----:----|----:----|----:----|----:----|----:----|----:----|

     E  S  S  F  R  R  G  S  S  A  T  P  N  L  R  G  S  T  R  Y     F1
```

```
      N  P  V  F  E  G  A  A  Q  L  L  P  I  C  V  E  A  L  A  I     F2
       I  Q  F  S  K  G  Q  L  S  Y  S  Q  S  A  W  K  H  S  L  *    F3
1621 GAATCCAGTTTTCGAAGGGGCAGCTCAGCTACTCCCAATCTGCGTGGAAGCACTCGCTAT 1680
     ----:----|----:----|----:----|----:----|----:----|----:----|

     S  R  V  S  A  V  S  T  E  Q  *  S  I  R  P  K  H  I  I  S      F1
      A  A  Y  L  L  *  V  Q  S  N  N  P  F  D  L  S  I  L  F  P     F2
       P  R  I  C  C  E  Y  R  A  I  I  H  S  T  *  A  Y  Y  F  L    F3
1681 AGCCGCGTATCTGCTGTGAGTACAGAGCAATAATCCATTCGACCTAAGCATATTATTTCC 1740
     ----:----|----:----|----:----|----:----|----:----|----:----|

     S  F  F  F  *  N  G  M  A  D  C  S  L  L  N  T  T  T  K  I      F1
      H  F  S  F  R  T  E  W  R  T  A  L  Y  *  T  Q  Q  Q  K  L     F2
       I  F  L  L  E  R  N  G  G  L  L  F  T  K  H  N  N  K  N  C    F3
1741 TCATTTTTCTTTTAGAACGGAATGGCGGACTGCTCTTTACTAAACACAACAACAAAAATT 1800
     ----:----|----:----|----:----|----:----|----:----|----:----|

     V  I  V  L  G  N  N  S  P  Q  R  D  V  D  R  N  I  P  E  E      F1
      L  L  F  L  A  T  T  R  P  K  E  M  W  T  E  T  Y  Q  K  K     F2
       Y  C  S  W  Q  Q  L  A  P  K  R  C  G  P  K  H  T  R  R  N    F3
1801 GTTATTGTTCTTGGCAACAACTCGCCCCAAAGAGATGTGGACCGAAACATACCAGAAGAA 1860
     ----:----|----:----|----:----|----:----|----:----|----:----|

     T  E  A  K  K  Q  E  N  *  M  Y  F  C  F  F  F  Y  N  N  N      F1
      L  R  P  K  N  R  K  T  E  C  I  S  V  F  F  F  I  I  I  I     F2
       *  G  Q  K  T  G  K  L  N  V  F  L  F  F  F  L  *  *  *  L    F3
1861 ACTGAGGCCAAAAAACAGGAAAACTGAATGTATTTCTGTTTTTTTTTTTATAATAATAAT 1920
     ----:----|----:----|----:----|----:----|----:----|----:----|

     Y  *  L  L  K  I  K  P  S  *                                    F1
      T  N  Y  S  K  *  N  R  A  X                                   F2
       L  I  T  Q  N  K  T  E  L  X                                  F3
1921 TACTAATTACTCAAAATAAAACCGAGCTAG 1950
     ----:----|----:----|----:----|
```
