## Supplementary material for "The incidence of candidate binding sites for β-arrestin in Drosophila neuropeptide GPCRs": S7 Text

***D. virilis TrissinR* Genomic DNA from analysis of:**
https://www.ncbi.nlm.nih.gov/nuccore/1805215602?report=graph&v=8059641:8102051

Annotations of the *D. melanogaster TrissinR* present three isoforms, PB, PC and PD. Blastp search recovers only a PB ortholog. Below is the inspection of the genomic DNA to search for possible PC and PD orthologs.

bp #1 of the record below = bp 88067341 from the shotgun sequence found in the above link

The predicted open reading frames, and predicted splice donor and acceptor sites, are marked in **RED**.

In *D melanogaster*, the PC isoform is created by use of the splice event (at comparable position to bp 265 below), which subtly alters the sequence within and immediately following the BBS-like sequence in ICL2. That potential exists in *D. virilis*, therefore I predict a PC isoform.

In *D melanogaster*, the PD isoform is created by neglect of the final splice event, leading to a Stop without including the second CT BBS. In the *D virilis* g DNA, if the orthologous splice site (at bp 6636 below) is neglected a PD orthologue would form with the sequence indicated, lacking the second BBS.

```
      I  N  S  S  S  N  N  N  T  N  S  N  N  N  R  L  S  K  R  I     F1
       S  T  A  A  A  T  T  T  P  T  A  T  T  T  G  Y  P  R  E  *    F2
        Q  Q  Q  Q  Q  Q  Q  H  Q  Q  Q  Q  Q  Q  A  I  Q  E  N  S   F3
   1 ATCAACAGCAGCAGCAACAACAACACCAACAGCAACAACAACAGGCTATCCAAGAGAATA 60
     ----:----|----:----|----:----|----:----|----:----|----:----|

      A  *  V  C  T  T  A  C  T  I  I  I  I  I  I  H  N  I  H  T     F1
       H  E  S  A  Q  Q  H  V  P  S  S  S  S  S  S  T  T  S  T  P    F2
        M  S  L  H  N  S  M  Y  H  H  H  H  H  H  P  Q  H  P  H  H   F3
  61 GCATGAGTCTGCACAACAGCATGTACCATCATCATCATCATCATCCACAACATCCACACC 120
     ----:----|----:----|----:----|----:----|----:----|----:----|

      T  I  N  S  T  N  S  I  S  C  Q  R  R  R  Q  Q  Q  Q  R  R     F1
       P  S  T  P  P  T  A  S  V  A  S  V  G  G  S  S  S  S  V  V    F2
        H  Q  L  H  Q  Q  H  Q  L  P  A  S  A  A  A  A  A  A  S  S   F3
 121 ACCATCAACTCCACCAACAGCATCAGTTGCCAGCGTCGGCGGCAGCAGCAGCAGCGTCGT 180
     ----:----|----:----|----:----|----:----|----:----|----:----|

      R  *  L  P  A  A  A  P  R  P  C  H  V  N  I  A  V  N  M  K     F1
       A  S  F  R  Q  Q  L  H  V  L  V  T  *  T  *  Q  *  I  *  K    F2
        L  A  S  G  S  S  S  T  S  L  S  R  K  H  S  S  K  Y  E  K   F3
 181 CGCTAGCTTCCGGCAGCAGCTCCACGTCCTTGTCACGTAAACATAGCAGTAAATATGAAA 240
     ----:----|----:----|----:----|----:----|----:----|----:----|

      S  V  E  S  A  S  P  R  V  R  *  V  L  Q  Q  Q  Q  L  I  T     F1
       A  W  S  Q  H  H  R  E  S  G  E  S  C  N  N  N  N  *  L  Q    F2
        R  G  V  S  I  T  E  S  Q  V  S  P  A  T  T  T  T  D  Y  R   F3
 241 AGCGTGGAGTCAGCATCACCGAGAGTCAGGTGAGTCCTGCAACAACAACAACTGATTACA 300
     ----:----|----:----|----:----|----:----|----:----|----:----|

      G  V  P  H  T  K  T  *  R  M  P  Q  S  I  Y  I  Y  M  Q  K     F1
       V  C  H  I  P  K  H  S  E  C  P  R  A  Y  I  S  I  C  K  R    F2
```

```
     C  A  T  Y  Q  N  I  A  N  A  P  E  H  I  Y  L  Y  A  K  D   F3
301 GGTGTGCCACATACCAAAACATAGCGAATGCCCCAGAGCATATATATCTATATGCAAAAG 360
    ----:----|----:----|----:----|----:----|----:----|----:----|

    I  E  *  L  I  L  M  S  S  Y  I  F  K  M  R  K  N  A  L  C    F1
     *  N  N  *  S  *  C  P  R  I  Y  L  R  *  E  K  M  L  C  A   F2
      R  I  T  N  P  N  V  L  V  Y  I  *  D  E  K  K  C  F  V  H  F3
361 ATAGAATAACTAATCCTAATGTCCTCGTATATATTTAAGATGAGAAAAAATGCTTTGTGC 420
    ----:----|----:----|----:----|----:----|----:----|----:----|

    I  V  L  N  P  E  Q  *  N  C  I  T  T  *  T  L  P  K  T  *    F1
     L  C  S  T  L  S  N  K  T  A  *  Q  L  E  H  Y  R  R  P  K   F2
      C  A  Q  P  *  A  I  K  L  H  N  N  L  N  T  T  E  D  L  K  F3
421 ATTGTGCTCAACCCTGAGCAATAAAACTGCATAACAACTTGAACACTACCGAAGACCTAA 480
    ----:----|----:----|----:----|----:----|----:----|----:----|

    S  L  P  *  I  L  I  R  L  G  *  G  R  F  C  K  H  L  G  T    F1
     V  C  H  K  F  *  S  D  *  D  R  E  G  S  A  S  I  L  E  L   F2
      F  A  I  N  F  N  Q  T  R  I  G  K  V  L  Q  A  S  W  N  Y  F3
481 AGTTTGCCATAAATTTTAATCAGACTAGGATAGGGAAGGTTCTGCAAGCATCTTGGAACT 540
    ----:----|----:----|----:----|----:----|----:----|----:----|

    I  F  K  T  R  N  L  F  K  Y  L  G  E  M  K  I  H  K  S  I    F1
     F  S  R  L  A  I  F  L  N  I  W  A  K  *  K  F  I  N  L  L   F2
      F  Q  D  S  Q  S  F  *  I  F  G  R  N  E  N  S  *  I  Y  Y  F3
541 ATTTTCAAGACTCGCAATCTTTTTAAATATTTGGGCGAAATGAAAATTCATAAATCTATT 600
    ----:----|----:----|----:----|----:----|----:----|----:----|

    I  R  P  A  Y  M  F  Y  F  V  A  V  P  E  I  A  F  C  *  K    F1
     Y  D  Q  H  I  C  F  I  L  *  L  F  R  K  *  R  F  V  K  N   F2
      T  T  S  I  Y  V  L  F  C  S  C  S  G  N  S  V  L  L  K  I  F3
601 ATACGACCAGCATATATGTTTTATTTTGTAGCTGTTCCGGAAATAGCGTTTTGTTAAAAA 660
    ----:----|----:----|----:----|----:----|----:----|----:----|

    L  L  E  F  *  I  F  V  R  N  A  I  K  *  V  Q  K  L  W  L    F1
     Y  *  N  S  K  F  S  S  E  M  R  L  N  E  F  K  S  C  G  C   F2
      I  R  I  L  N  F  R  P  K  C  D  *  M  S  S  K  V  V  V  V  F3
661 TTATTAGAATTCTAAATTTTCGTCCGAAATGCGATTAAATGAGTTCAAAAGTTGTGGTTG 720
    ----:----|----:----|----:----|----:----|----:----|----:----|

    S  I  T  S  R  I  N  W  M  Y  L  N  *  Y  E  E  E  I  G  Y    F1
     P  L  H  L  A  *  I  G  C  I  *  I  N  T  K  K  K  *  D  I   F2
      H  Y  I  S  H  K  L  D  V  F  K  L  I  R  R  R  N  R  I  F  F3
721 TCCATTACATCTCGCATAAATTGGATGTATTTAAATTAATACGAAGAAGAAATAGGATAT 780
    ----:----|----:----|----:----|----:----|----:----|----:----|

    L  Y  F  *  I  N  *  M  R  S  S  K  T  *  K  L  F  I  E  Q    F1
     C  T  F  E  S  I  E  C  V  Q  A  K  H  K  S  F  L  S  S  K   F2
      V  L  L  N  Q  L  N  A  F  K  Q  N  I  K  A  F  Y  R  A  K  F3
781 TTGTACTTTTGAATCAATTGAATGCGTTCAAGCAAAACATAAAAGCTTTTTATCGAGCAA 840
    ----:----|----:----|----:----|----:----|----:----|----:----|

    S  A  S  K  F  F  G  *  N  T  *  N  T  W  R  T  N  F  S  *    F1
     V  L  Q  S  S  S  V  K  I  H  K  I  L  G  E  Q  I  L  V  S   F2
      C  F  K  V  L  R  L  K  Y  I  K  Y  L  E  N  K  F  *  L  V  F3
841 AGTGCTTCAAAGTTCTTCGGTTAAAATACATAAAATACTTGGAGAACAAATTTTAGTTAG 900
    ----:----|----:----|----:----|----:----|----:----|----:----|

    S  F  K  T  H  V  A  W  L  S  V  *  V  A  L  C  L  V  K  L    F1
     R  L  K  L  T  L  L  G  F  Q  F  E  L  H  F  V  W  S  S  W   F2
      V  *  N  S  R  C  L  A  F  S  L  S  C  T  L  S  G  Q  V  G  F3
901 TCGTTTAAAACTCACGTTGCTTGGCTTTCAGTTTGAGTTGCACTTTGTCTGGTCAAGTTG 960
    ----:----|----:----|----:----|----:----|----:----|----:----|

    V  C  F  S  C  W  Q  L  E  *  V  S  K  *  L  A  V  A  Y  N    F1
```

```
      C  V  S  A  V  G  S  *  S  K  *  V  S  S  S  L  L  L  I  I    F2
       V  F  Q  L  L  A  A  R  V  S  K  *  V  A  R  C  C  L  *  *   F3
 961 GTGTGTTTCAGCTGTTGGCAGCTAGAGTAAGTAAGTAAGTAGCTCGCTGTTGCTTATAAT 1020
     ----:----|----:----|----:----|----:----|----:----|----:----|

     S  *  Y  L  L  K  L  S  E  R  L  S  Q  V  C  A  E  I  A  I     F1
      A  N  I  C  *  S  *  A  K  G  Y  R  K  F  A  L  K  L  Q  F    F2
       L  I  F  A  E  V  E  R  K  A  I  A  S  L  R  *  N  C  N  L   F3
1021 AGCTAATATTTGCTGAAGTTGAGCGAAAGGCTATCGCAAGTTTGCGCTGAAATTGCAATT 1080
     ----:----|----:----|----:----|----:----|----:----|----:----|

     W  P  T  W  T  L  P  P  P  G  C  R  A  A  K  L  P  V  C  R     F1
      G  Q  L  G  R  C  L  H  L  A  A  G  L  P  S  C  R  C  A  E    F2
       A  N  L  D  V  A  S  T  W  L  P  G  C  Q  A  A  G  V  P  S   F3
1081 TGGCCAACTTGGACGTTGCCTCCACCTGGCTGCCGGGCTGCCAAGCTGCCGGTGTGCCGA 1140
     ----:----|----:----|----:----|----:----|----:----|----:----|

     A  V  S  F  T  A  A  A  V  K  A  Y  *  K  V  S  V  S  M  K     F1
      Q  S  V  S  R  Q  Q  L  S  K  R  I  R  K  Y  Q  *  A  *  S    F2
       S  Q  F  H  G  S  S  C  Q  S  V  L  E  S  I  S  E  H  E  V   F3
1141 GCAGTCAGTTTCACGGCAGCAGCTGTCAAAGCGTATTAGAAAGTATCAGTGAGCATGAAG 1200
     ----:----|----:----|----:----|----:----|----:----|----:----|

     F  K  *  N  I  *  F  L  A  T  A  L  S  W  Q  Q  A  G  G  S     F1
      S  S  E  I  S  S  F  *  Q  L  H  *  V  G  S  K  L  A  A  H    F2
       Q  V  K  Y  L  V  F  S  N  C  I  E  L  A  A  S  W  R  L  M   F3
1201 TTCAAGTGAAATATCTAGTTTTTAGCAACTGCATTGAGTTGGCAGCAAGCTGGCGGCTCA 1260
     ----:----|----:----|----:----|----:----|----:----|----:----|

     C  G  V  C  A  I  Y  Q  L  A  T  Y  L  T  P  I  D  S  L  N     F1
      A  A  Y  A  Q  Y  I  N  L  P  L  T  *  P  Q  L  T  A  *  T    F2
       R  R  M  R  N  I  S  T  C  H  L  L  N  P  N  *  Q  P  E  Q   F3
1261 TGCGGCGTATGCGCAATATATCAACTTGCCACTTACTTAACCCCAATTGACAGCCTGAAC 1320
     ----:----|----:----|----:----|----:----|----:----|----:----|

     R  Y  L  A  L  S  *  A  E  V  M  I  I  R  L  S  T  T  T  Y     F1
      G  I  W  H  *  V  E  L  K  *  *  L  F  V  Y  P  Q  Q  L  T    F2
       V  S  G  I  E  L  S  *  S  N  D  Y  S  F  I  H  N  N  L  Q   F3
1321 AGGTATCTGGCATTGAGTTGAGCTGAAGTAATGATTATTCGTTTATCCACAACAACTTAC 1380
     ----:----|----:----|----:----|----:----|----:----|----:----|

     N  M  D  T  S  I  S  Y  I  T  C  P  L  K  W  L  *  S  S  E     F1
      I  W  T  H  P  S  A  T  S  P  A  L  *  S  G  S  N  Q  V  K    F2
       Y  G  H  I  H  Q  L  H  H  L  P  F  E  V  A  L  I  K  *  K   F3
1381 AATATGGACACATCCATCAGCTACATCACCTGCCCTTTGAAGTGGCTCTAATCAAGTGAA 1440
     ----:----|----:----|----:----|----:----|----:----|----:----|

     K  C  A  L  R  K  A  V  E  S  I  R  Y  V  F  Y  V  C  T  L     F1
      N  V  R  *  E  K  Q  L  R  V  Y  Y  V  M  C  S  M  C  V  L  *    F2
       M  C  A  K  K  S  S  *  E  Y  T  L  C  V  L  C  V  Y  S  N   F3
1441 AAATGTGCGCTAAGAAAAGCAGTTGAGAGTATACGTTATGTGTTCTATGTGTGTACTCTA 1500
     ----:----|----:----|----:----|----:----|----:----|----:----|

     M  I  Y  E  H  K  F  V  T  E  Q  R  F  G  S  W  G  *  I  T     F1
      *  F  M  N  I  N  L  L  P  S  N  G  L  V  A  G  G  K  S  L    F2
       D  L  *  T  *  I  C  Y  R  A  T  V  W  *  L  G  V  N  H  L   F3
1501 ATGATTTATGAACATAAATTTGTTACCGAGCAACGGTTTGGTAGCTGGGGGTAAATCACT 1560
     ----:----|----:----|----:----|----:----|----:----|----:----|

     *  V  S  C  T  K  H  N  L  N  K  K  G  S  Y  F  E  C  C  E     F1
      K  C  R  A  Q  S  I  T  S  I  K  R  E  A  I  L  S  A  A  K    F2
       S  V  V  H  K  A  *  P  Q  *  K  G  K  L  F  *  V  L  R  N   F3
1561 TAAGTGTCGTGCACAAAGCATAACCTCAATAAAAAGGGAAGCTATTTTGAGTGCTGCGAA 1620
     ----:----|----:----|----:----|----:----|----:----|----:----|
```

```
      T  N  V  I  Y  K  E  F  M  *  S  F  Y  M  H  T  L  V  G  *     F1
       P  T  *  F  I  K  S  L  C  K  V  F  T  C  I  L  *  *  D  R    F2
        Q  R  N  L  *  R  V  Y  V  K  F  L  H  A  Y  S  S  R  I  G   F3
1621 ACCAACGTAATTTATAAAGAGTTTATGTAAAGTTTTTACATGCATACTCTAGTAGGATAG 1680
     ----:----|----:----|----:----|----:----|----:----|----:----|

      A  I  F  Y  *  Y  K  C  *  F  E  Y  *  L  C  *  F  T  I  I     F1
       R  Y  F  I  N  I  N  V  N  L  N  I  N  S  A  N  L  Q  *  *    F2
        D  I  L  L  I  *  M  L  I  *  I  L  T  L  L  I  Y  N  N  N   F3
1681 GCGATATTTTATTAATATAAATGTTAATTTGAATATTAACTCTGCTAATTTACAATAATA 1740
     ----:----|----:----|----:----|----:----|----:----|----:----|

      T  F  L  P  Q  Y  F  C  I  I  N  L  L  Y  A  F  L  *  P  *     F1
       H  F  F  P  N  I  F  V  *  L  I  Y  S  T  P  F  Y  D  L  N    F2
        I  S  S  P  I  F  L  Y  N  *  F  T  L  R  L  F  M  T  L  I   F3
1741 ACATTTCTTCCCCAATATTTTTGTATAATTAATTTACTCTACGCCTTTTTATGACCTTAA 1800
     ----:----|----:----|----:----|----:----|----:----|----:----|

      S  *  A  S  C  L  T  L  F  V  F  Q  C  S  F  S  L  S  F  S     F1
       P  K  L  A  V  *  R  S  L  C  F  N  V  L  S  L  S  L  F  L    F2
        L  S  *  L  F  N  A  L  C  V  S  M  F  F  L  S  L  F  F  Y   F3
1801 TCCTAAGCTAGCTGTTTAACGCTCTTTGTGTTTCAATGTTCTTTCTCTCTCTCTTTTTCT 1860
     ----:----|----:----|----:----|----:----|----:----|----:----|

      I  S  K  S  H  A  V  Y  M  C  L  S  L  F  F  F  S  V  F  W     F1
       S  L  N  L  T  L  S  I  C  V  S  L  S  F  S  F  L  F  F  G    F2
        L  *  I  S  R  C  L  Y  V  S  L  S  L  F  L  F  C  F  L  A   F3
1861 ATCTCTAAATCTCACGCTGTCTATATGTGTCTCTCTCTCTTTTTCTTTTCTGTTTTTTGG 1920
     ----:----|----:----|----:----|----:----|----:----|----:----|

      L  F  I  L  P  L  T  T  T  I  F  F  G  S  P  L  T  I  S  Q     F1
       F  L  F  Y  H  *  Q  Q  Q  F  F  L  A  P  H  *  Q  Y  H  N    F2
        F  Y  F  T  T  N  N  N  N  F  F  W  L  P  T  N  N  I  T  T   F3
1921 CTTTTTATTTTACCACTAACAACAACAATTTTTTTGGCTCCCCACTAACAATATCACAA 1980
     ----:----|----:----|----:----|----:----|----:----|----:----|

      R  Q  Q  Q  R  N  T  Q  H  T  Q  K  L  A  *  *  L  Q  G  G     F1
       D  N  N  N  A  I  H  N  T  H  K  N  *  L  D  N  C  K  V  G    F2
        T  T  T  T  Q  Y  T  T  H  T  K  I  S  L  I  I  A  R  W  V   F3
1981 CGACAACAACAACGCAATACACAACACACACAAAAATTAGCTTGATAATTGCAAGGTGGG 2040
     ----:----|----:----|----:----|----:----|----:----|----:----|

      Y  V  *  I  V  L  K  P  M  P  K  R  F  K  L  C  T  A  Q  T     F1
       M  S  E  *  S  L  S  Q  C  Q  N  A  L  S  S  V  L  H  R  L    F2
        C  L  N  S  P  *  A  N  A  K  T  L  *  A  L  Y  C  T  D  *   F3
2041 TATGTCTGAATAGTCCTTAAGCCAATGCCAAAACGCTTTAAGCTCTGTACTGCACAGACT 2100
     ----:----|----:----|----:----|----:----|----:----|----:----|

      N  C  *  S  *  C  L  M  L  *  H  L  L  L  M  Q  V  S  L  E     F1
       T  V  N  R  N  A  *  C  Y  N  I  Y  Y  S  C  R  S  R  W  R    F2
        L  L  I  V  M  P  N  A  I  T  S  I  T  H  A  G  L  A  G  G   F3
2101 AACTGTTAATCGTAATGCCTAATGCTATAACATCTATTACTCATGCAGGTCTCGCTGGAG 2160
     ----:----|----:----|----:----|----:----|----:----|----:----|

      A  D  R  P  I  V  S  A  C  R  K  T  S  F  Y  H  H  S  H  S     F1
       R  I  G  P  L  *  V  L  V  A  K  R  A  S  I  I  T  P  T  R    F2
        G  S  A  H  C  E  C  L  S  Q  N  E  L  L  S  S  L  P  L  A   F3
2161 GCGGATCGGCCCATTGTGAGTGCTTGTCGCAAAACGAGCTTCTATCATCACTCCCACTCG 2220
     ----:----|----:----|----:----|----:----|----:----|----:----|

      H  N  Q  R  Q  G  N  G  G  A  G  G  G  G  A  T  H  M  S  H     F1
       T  I  N  D  R  A  M  V  V  P  A  E  V  A  P  H  T  C  H  I    F2
        Q  S  T  T  G  Q  W  W  C  R  R  R  W  R  H  T  H  V  T  F   F3
2221 CACAATCAACGACAGGGCAATGGTGGTGCCGGCGGAGGTGGCGCCACACACATGTCACAT 2280
     ----:----|----:----|----:----|----:----|----:----|----:----|
```

```
     S  S  S  N  V  L  R  A  R  R  G  V  V  R  M  L  I  I  F  V    F1
      H  R  A  M  C  Y  A  L  A  G  A  W  S  A  C  *  *  Y  L  C   F2
       I  E  Q  C  A  T  R  S  Q  G  R  G  P  H  A  D  N  I  C  A  F3
2281 TCATCGAGCAATGTGCTACGCGCTCGCAGGGGCGTGGTCCGCATGCTGATAATATTTGTG 2340
     ----:----|----:----|----:----|----:----|----:----|----:----|

     L  T  F  A  L  C  N  L  P  Y  H  A  R  K  M  W  Q  Y  W  *    F1
      *  H  S  R  S  A  I  C  R  I  M  R  A  K  C  G  N  T  G  E   F2
       D  I  R  A  L  Q  S  A  V  S  C  A  Q  N  V  A  I  L  V  S  F3
2341 CTGACATTCGCGCTCTGCAATCTGCCGTATCATGCGCGCAAAATGTGGCAATACTGGTGA 2400
     ----:----|----:----|----:----|----:----|----:----|----:----|

     V  H  S  N  K  Q  L  K  N  I  C  I  *  K  H  F  D  I  S  M    F1
      Y  T  V  T  N  N  *  R  I  Y  V  Y  E  S  I  S  T  F  L  *   F2
       T  Q  *  Q  T  I  K  E  Y  M  Y  M  K  A  F  R  H  F  Y  E  F3
2401 GTACACAGTAACAAACAATTAAAGAATATATGTATATGAAAGCATTTCGACATTTCTATG 2460
     ----:----|----:----|----:----|----:----|----:----|----:----|

     K  N  F  K  N  V  R  Q  F  Q  I  Y  L  S  *  R  V  I  K  I    F1
      K  T  L  K  M  F  V  N  F  K  Y  T  *  V  R  E  L  *  K  S   F2
       K  L  *  K  C  S  S  I  S  N  I  L  E  L  E  S  Y  K  N  Q  F3
2461 AAAAACTTTAAAAATGTTCGTCAATTTCAAATATACTTGAGTTAGAGAGTTATAAAAATC 2520
     ----:----|----:----|----:----|----:----|----:----|----:----|

     R  A  Q  P  D  *  *  Q  L  *  V  G  R  L  A  A  P  F  Y  I    F1
      E  L  S  L  I  N  D  N  C  E  L  G  D  *  L  L  L  L  F  I  F   F2
       S  S  A  *  L  M  T  T  V  S  W  A  I  S  C  S  F  L  Y  L  F3
2521 AGAGCTCAGCCTGATTAATGACAACTGTGAGTTGGGCGATTAGCTGCTCCTTTTTATATT 2580
     ----:----|----:----|----:----|----:----|----:----|----:----|

     *  N  I  N  K  N  S  *  M  K  L  K  F  K  N  Y  Y  S  *  K    F1
      E  I  L  T  K  I  V  K  *  N  *  S  L  K  I  I  I  V  K  K   F2
       K  Y  *  Q  K  *  L  N  E  I  E  V  *  K  L  L  *  L  K  K  F3
2581 TGAAATATTAACAAAAATAGTTAAATGAAATTGAAGTTTAAAAATTATTATAGTTAAAAA 2640
     ----:----|----:----|----:----|----:----|----:----|----:----|

     K  N  I  R  A  R  F  F  A  V  Y  *  G  R  Q  S  A  P  L  S    F1
      R  T  S  G  H  V  F  L  L  Y  T  R  V  A  S  Q  R  P  C  L   F2
       E  H  Q  G  T  F  F  C  C  I  L  G  S  P  V  S  A  L  V  F  F3
2641 AAGAACATCAGGGCACGTTTTTTGCTGTATACTAGGGTCGCCAGTCAGCGCCCTTGTCT 2700
     ----:----|----:----|----:----|----:----|----:----|----:----|

     F  P  L  S  S  S  S  S  S  S  S  T  S  S  S  T  Y  R  P  S    F1
      F  H  Y  H  H  H  H  H  H  R  Q  H  L  R  R  H  T  A  H  Q   F2
       S  I  I  I  I  I  I  I  I  V  N  I  F  V  D  I  P  P  I  S  F3
2701 TTTCCATTATCATCATCATCATCATCATCGTCAACATCTTCGTCGACATACCGCCCATCA 2760
     ----:----|----:----|----:----|----:----|----:----|----:----|

     A  P  F  Y  V  M  Q  T  V  E  S  H  L  H  I  A  C  P  S  A    F1
      R  H  F  M  L  C  K  Q  L  R  V  I  C  I  *  R  A  H  L  L   F2
       A  I  L  C  Y  A  N  S  *  E  S  F  A  Y  S  V  P  I  C  C  F3
2761 GCGCCATTTTATGTTATGCAAACAGTTGAGAGTCATTTGCATATAGCGTGCCCATCTGCT 2820
     ----:----|----:----|----:----|----:----|----:----|----:----|

     V  P  *  I  I  L  L  L  G  H  A  N  N  *  T  V  N  *  S  G    F1
      C  R  K  *  Y  S  Y  L  A  M  Q  T  I  K  L  S  I  E  V  G   F2
       A  V  N  N  T  P  T  W  P  C  K  Q  L  N  C  Q  L  K  W  D  F3
2821 GTGCCGTAAATAATACTCCTACTTGGCCATGCAAACAATTAAACTGTCAATTGAAGTGGG 2880
     ----:----|----:----|----:----|----:----|----:----|----:----|

     T  E  A  P  D  *  L  L  K  C  F  C  S  P  S  A  T  I  L  L    F1
      L  K  R  Q  T  D  C  *  S  A  S  A  P  P  Q  P  Q  F  C  S   F2
       *  S  A  R  L  I  A  K  V  L  L  L  P  L  S  H  N  F  A  L  F3
2881 ACTGAAGCGCCAGACTGATTGCTAAAGTGCTTCTGCTCCCCCTCAGCCACAATTTTGCTC 2940
```

```
     ----:----|----:----|----:----|----:----|----:----|----:----|

     * L L C R A R L N V F H T V R C S N R I    F1
      N C Y A E L V L M F S I Q S V A P I E Y   F2
       I V M Q S S S * C F P Y S P L L Q S N I  F3
2941 TAATTGTTATGCAGAGCTCGTCTTAATGTTTTCCATACAGTCCGTTGCTCCAATCGAATA 3000
     ----:----|----:----|----:----|----:----|----:----|----:----|

     Y T S T L S * E I T D Q N S E L Q H K L    F1
      I L Q H S L K K L L T R T L N F S I N *   F2
       Y F N T L L R N Y * P E L * T S A * I R  F3
3001 TATACTTCAACACTCTCTTAAGAAATTACTGACCAGAACTCTGAACTTCAGCATAAATTA 3060
     ----:----|----:----|----:----|----:----|----:----|----:----|

     G C L P I A * P K V S F S N S Y L F R C    F1
      A A Y Q * P N R K * A L A I L I F S V A   F2
       L L T N S L T E S K L * Q F L S F P L H  F3
3061 GGCTGCTTACCAATAGCCTAACCGAAAGTAAGCTTTAGCAATTCTTATCTTTTCCGTTGC 3120
     ----:----|----:----|----:----|----:----|----:----|----:----|

     I S L Y * I L A K * Q N * F L L Q A S K    F1
      L V C I E Y W P N N R I N F C C K L R N   F2
       * F V L N I G Q I T E L I S V A S F E T  F3
3121 ATTAGTTTGTATTGAATATTGGCCAAATAACAGAATTAATTTCTGTTGCAAGCTTCGAAA 3180
     ----:----|----:----|----:----|----:----|----:----|----:----|

     L I T I Y S K I F K S * R * N * M L L N    F1
      * * Q Y I Q K Y L N H E D E I K C Y * T   F2
       N N N I F K N I * I M K M K L N V I K H  F3
3181 CTAATAACAATATATTCAAAAATATTTAAATCATGAAGATGAAATTAAATGTTATTAAAC 3240
     ----:----|----:----|----:----|----:----|----:----|----:----|

     I F * L Y Y L S Y Y L L F R Y V I * L I    F1
      F F N Y I I Y L I I Y Y L D M S Y N * Y   F2
       F L I I L F I L L F I I * I C H I I N I  F3
3241 ATTTTTAATTATATTATTTATCTTATTATTTATTATTTAGATATGTCATATAATTAATA 3300
     ----:----|----:----|----:----|----:----|----:----|----:----|

     L Y Y F P N S L K W C P K S * Y T I Y K    F1
      C I I F Q T L * S G V Q K A N I L S T N   F2
       V L F S K L F K V V S K K L I Y Y L Q I  F3
3301 TTGTATTATTTTCCAAACTCTTTAAAGTGGTGTCCAAAAAGCTAATATACTATCTACAAA 3360
     ----:----|----:----|----:----|----:----|----:----|----:----|

     Y T F N T C R I * I I N C K V S Q V L S    F1
      T P L T P V V F K L * I V K C H K S C L   F2
       H L * H L S Y L N Y K L * S V T S P V S  F3
3361 TACACCTTTAACACCTGTCGTATTTAAATTATAAATTGTAAAGTGTCACAAGTCCTGTCT 3420
     ----:----|----:----|----:----|----:----|----:----|----:----|

     R L P R M L S N T E P I S I W C Q L W Y    F1
      V C P E C C Q T Q N Q S P F G V S S G I   F2
       S A Q N A V K H R T N L H L V S A L V S  F3
3421 CGTCTGCCCAGAATGCTGTCAAACACAGAACCAATCTCCATTTGGTGTCAGCTCTGGTAT 3480
     ----:----|----:----|----:----|----:----|----:----|----:----|

     Q S I G H S I * I I A L N L L L R E F L    F1
      N Q L G I Q Y E L L L * I C F * G N F C   F2
       I N W A F N M N Y C F K F A F K G I F V  F3
3481 CAATCAATTGGGCATTCAATATGAATTATTGCTTTAAATTTGCTTTTAAGGGAATTTTTG 3540
     ----:----|----:----|----:----|----:----|----:----|----:----|

     * S P * S L L P I Y S I S N N Y W A S H    F1
      D H L E A C C Q F I P L A T I I G H H M   F2
       I T L K L A A N L F H * Q Q L L G I T W  F3
```

```
3541 TGATCACCTTGAAGCTTGCTGCCAATTTATTCCATTAGCAACAATTATTGGGCATCACAT 3600
     ----:----|----:----|----:----|----:----|----:----|----:----|

     G  K  *  S  N  K  K  K  N  G  K  G  L  L  L  L  L  F  Q  P    F1
      A  N  S  L  I  K  K  K  K  M  V  K  V  C  S  C  Y  C  F  N  L F2
       Q  I  V  *  *  K  K  K  W  *  R  S  A  P  V  I  V  S  T  S  F3
3601 GGCAAATAGTCTAATAAAAAAAAAAATGGTAAAGGTCTGCTCCTGTTATTGTTTCAACCT 3660
     ----:----|----:----|----:----|----:----|----:----|----:----|

     P  L  D  P  S  C  F  C  L  G  F  P  F  T  C  F  Y  F  F  A    F1
      R  S  T  P  A  V  F  V  W  V  F  L  L  L  A  F  I  F  L  L   F2
       A  R  P  Q  L  F  L  F  G  F  S  F  Y  L  L  L  F  F  C  Y  F3
3661 CCGCTCGACCCCAGCTGTTTTTGTTTGGGTTTTCCTTTTACTTGCTTTTATTTTTTTGCT 3720
     ----:----|----:----|----:----|----:----|----:----|----:----|

     I  G  L  W  F  F  V  G  L  V  *  F  S  H  Y  T  T  A  Y  G    F1
      L  A  F  G  F  L  W  V  S  F  D  F  P  I  I  R  Q  L  M  V   F2
       W  P  L  V  F  C  G  S  R  L  I  F  P  L  Y  D  S  L  W  S  F3
3721 ATTGGCCTTTGGTTTTTTGTGGGTCTCGTTTGATTTTCCCATTATACGACAGCTTATGGT 3780
     ----:----|----:----|----:----|----:----|----:----|----:----|

     R  L  S  E  G  F  S  G  Q  W  P  V  V  R  *  L  A  A  W  I    F1
      V  *  V  R  D  S  V  A  S  G  Q  W  S  D  D  W  L  H  G  *   F2
       F  E  *  G  I  Q  W  P  V  A  S  G  P  M  I  G  C  M  D  S  F3
3781 CGTTTGAGTGAGGGATTCAGTGGCCAGTGGCCAGTGGTCCGATGATTGGCTGCATGGATA 3840
     ----:----|----:----|----:----|----:----|----:----|----:----|

     A  M  S  W  Q  L  P  R  *  F  D  I  N  N  L  T  P  T  D  L    F1
      Q  C  H  G  S  Y  P  D  D  S  I  *  T  I  *  R  P  Q  T  W   F2
       N  V  M  A  A  T  Q  M  I  R  Y  K  Q  F  N  A  H  R  L  G  F3
3841 GCAATGTCATGGCAGCTACCCAGATGATTCGATATAAACAATTTAACGCCCACAGACTTG 3900
     ----:----|----:----|----:----|----:----|----:----|----:----|

     V  A  E  S  *  K  S  H  R  N  K  *  K  S  E  M  E  A  K  L    F1
      W  Q  S  L  K  S  R  T  E  T  N  K  K  A  K  W  K  Q  N  *   F2
       G  R  V  L  K  V  A  Q  K  Q  I  K  K  R  N  G  S  K  I  E  F3
3901 GTGGCAGAGTCTTAAAAGTCGCACAGAAACAAATAAAAAAGCGAAATGGAAGCAAAATTG 3960
     ----:----|----:----|----:----|----:----|----:----|----:----|

     K  M  *  N  A  K  K  *  T  F  I  K  A  *  F  R  Q  A  K  S    F1
      K  C  E  M  P  R  S  K  H  L  S  K  H  D  L  D  R  P  S  P   F2
       N  V  K  C  Q  E  V  N  I  Y  Q  S  M  I  *  T  G  Q  V  Q  F3
3961 AAAATGTGAAATGCCAAGAAGTAAACATTTATCAAAGCATGATTTAGACAGGCCAAGTCC 4020
     ----:----|----:----|----:----|----:----|----:----|----:----|

     R  Y  I  S  I  I  F  A  D  T  T  R  C  E  Y  S  G  *  N  S    F1
      D  I  S  Q  L  F  L  L  T  R  H  V  V  S  T  L  A  E  I  R   F2
       I  Y  L  N  Y  F  C  *  H  D  T  L  *  V  L  W  L  K  F  V  F3
4021 AGATATATCTCAATTATTTTTGCTGACACGACACGTTGTGAGTACTCTGGCTGAAATTCG 4080
     ----:----|----:----|----:----|----:----|----:----|----:----|

     C  I  V  F  M  W  Q  T  K  I  N  K  F  Y  L  M  P  F  Q  L    F1
      A  L  F  L  C  G  K  R  K  *  I  N  F  I  *  C  L  S  S  Y   F2
       H  C  F  Y  V  A  N  E  N  K  *  I  L  F  N  A  F  P  A  T  F3
4081 TGCATTGTTTTTATGTGGCAAACGAAAATAAATAAATTTTATTTAATGCCTTTCCAGCTA 4140
     ----:----|----:----|----:----|----:----|----:----|----:----|

     H  I  S  H  I  T  K  N  A  R  K  K  A  P  S  E  N  A  F  Q    F1
      I  F  H  I  L  L  K  M  R  E  K  K  P  R  A  K  M  H  F  S   F2
       Y  F  T  Y  Y  *  K  C  A  K  K  S  P  E  R  K  C  I  S  A  F3
4141 CATATTTCACATATTACTAAAAATGCGCGAAAAAAAGCCCCGAGCGAAAATGCATTTCAG 4200
     ----:----|----:----|----:----|----:----|----:----|----:----|

     L  F  C  A  C  C  T  *  N  I  L  L  Y  E  L  N  *  Y  F  W    F1
      Y  F  V  P  V  A  L  K  I  Y  Y  Y  M  N  *  T  N  I  F  G   F2
```

```
      I  L  C  L  L  H  L  K  Y  I  I  I  *  I  K  L  I  F  L  G   F3
4201 CTATTTTGTGCCTGTTGCACTTAAAATATATTATTATATGAATTAAACTAATATTTTTGG 4260
     ----:----|----:----|----:----|----:----|----:----|----:----|

     G  A  S  R  A  K  R  P  D  C  K  Q  I  L  *  S  I  *  P  K    F1
      E  Q  A  E  Q  N  D  Q  I  V  N  K  F  Y  E  V  F  S  Q  N   F2
       S  K  P  S  K  T  T  R  L  *  T  N  F  M  K  Y  L  A  K  I  F3
4261 GGAGCAAGCCGAGCAAAACGACCAGATTGTAAACAAATTTTATGAAGTATTTAGCCAAAA 4320
     ----:----|----:----|----:----|----:----|----:----|----:----|

     Y  I  M  C  E  *  E  I  A  C  F  A  K  R  S  S  L  I  N  V    F1
      I  *  C  V  N  K  K  L  R  V  S  Q  N  A  P  V  *  *  T  L   F2
       Y  N  V  *  I  R  N  C  V  F  R  K  T  L  Q  F  N  K  R  *  F3
4321 TATATAATGTGTGAATAAGAAATTGCGTGTTTCGCAAAACGCTCCAGTTTAATAAACGTT 4380
     ----:----|----:----|----:----|----:----|----:----|----:----|

     N  K  *  L  M  A  N  I  N  K  *  Y  L  Y  K  Y  K  F  Q  A    F1
      I  S  S  *  W  P  T  *  I  N  N  T  Y  I  N  I  N  S  K  P   F2
       *  V  V  N  G  Q  H  K  *  I  I  L  I  *  I  *  I  P  S  H  F3
4381 AATAAGTAGTTAATGGCCAACATAAATAAATAATACTTATATAAATATAAATTCCAAGCC 4440
     ----:----|----:----|----:----|----:----|----:----|----:----|

     T  N  K  K  N  T  N  M  T  A  C  *  P  D  F  L  S  S  L  F    F1
      P  I  K  K  K  T  Q  I  *  Q  L  V  N  Q  T  F  *  A  V  Y  L   F2
       Q  *  K  K  H  K  Y  D  S  L  L  T  R  L  F  E  Q  F  I  *  F3
4441 ACCAATAAAAAAAACACAAATATGACAGCTTGTTAACCAGACTTTTTGAGCAGTTTATTT 4500
     ----:----|----:----|----:----|----:----|----:----|----:----|

     D  T  L  S  F  K  L  K  F  L  G  I  *  I  I  K  I  C  Y  P    F1
      T  L  *  A  L  S  *  N  S  *  A  Y  E  L  S  K  Y  V  T  L   F2
       H  F  K  L  *  A  K  I  L  R  H  M  N  Y  Q  N  M  L  P  Y  F3
4501 GACACTTTAAGCTTTAAGCTAAAATTCTTAGGCATATGAATTATCAAAATATGTTACCCT 4560
     ----:----|----:----|----:----|----:----|----:----|----:----|

     M  R  S  S  R  D  Y  I  A  T  A  R  N  I  K  I  K  S  R  S    F1
      C  A  V  Q  E  I  I  L  Q  L  H  G  I  *  K  L  N  R  D  P   F2
       A  Q  F  K  R  L  Y  C  N  C  T  E  Y  K  N  *  I  E  I  L  F3
4561 ATGCGCAGTTCAAGAGATTATATTGCAACTGCACGGAATATAAAAATTAAATCGAGATCC 4620
     ----:----|----:----|----:----|----:----|----:----|----:----|

     L  *  T  F  P  P  F  C  R  Y  I  R  I  V  Q  L  A  N  A  E  F    F1
      Y  R  H  F  P  P  F  A  D  I  *  E  *  C  S  S  Q  M  L  N  L   F2
       I  D  I  S  L  L  Q  I  Y  K  N  S  A  A  R  K  C  *  I  *  F3
4621 TTATAGACATTTCCCTTTTGCAGATATATAAGAATAGTGCAGCTCGCAAATGCTGAATTT 4680
     ----:----|----:----|----:----|----:----|----:----|----:----|

     D  E  N  I  L  E  S  R  I  F  V  H  I  R  T  K  F  S  I  D    F1
      M  K  I  Y  W  K  A  E  Y  L  S  I  *  E  P  S  F  R  *  I   F2
       *  K  Y  I  G  K  Q  N  I  C  P  Y  K  N  Q  V  F  D  R  S  F3
4681 GATGAAAATATATTGGAAAGCAGAATATTTGTCCATATAAGAACCAAGTTTTCGATAGAT 4740
     ----:----|----:----|----:----|----:----|----:----|----:----|

     L  S  Y  S  N  Y  M  I  *  W  S  V  K  L  I  H  S  V  I  *    F1
      F  L  I  A  T  I  *  F  S  G  P  *  N  L  Y  I  R  S  F  N   F2
       F  L  *  Q  L  Y  D  L  V  V  R  E  T  Y  T  F  G  H  L  T  F3
4741 CTTTCTTATAGCAACTATATGATTTAGTGGTCCGTGAAACTTATACATTCGGTCATTTAA 4800
     ----:----|----:----|----:----|----:----|----:----|----:----|

     H  I  F  N  I  V  N  K  Y  K  M  D  V  T  *  I  S  K  L  N    F1
      I  F  S  I  L  *  T  N  I  K  W  M  *  L  K  F  Q  S  L  I   F2
       Y  F  Q  Y  C  K  Q  I  *  N  G  C  N  L  N  F  K  A  *  Y  F3
4801 CATATTTCAATATTGTAAACAATATAAAATGGATGTAACTTAAATTTCAAAGCTTAAT 4860
     ----:----|----:----|----:----|----:----|----:----|----:----|

     I  *  A  S  A  Y  V  Q  V  E  L  V  P  R  D  L  L  T  G  Y    F1
```

```
      F  K  L  L  P  M  S  K  L  S  S  Y  L  E  I  Y  *  Q  G  I    F2
       L  S  F  C  L  C  P  S  *  A  R  T  *  R  F  T  N  R  V  F   F3
4861 ATTTAAGCTTCTGCCTATGTCCAAGTTGAGCTCGTACCTAGAGATTTACTAACAGGGTAT 4920
     ----:----|----:----|----:----|----:----|----:----|----:----|

     L  L  P  K  N  N  T  I  N  V  I  C  C  H  I  S  L  S  S  G    F1
      Y  C  P  K  T  T  Q  L  M  L  S  A  A  I  L  V  *  V  L  V   F2
       I  A  Q  K  Q  H  N  *  C  Y  L  L  P  Y  *  S  E  F  W  S  F3
4921 TTATTGCCCAAAAACAACACAATTAATGTTATCTGCTGCCATATTAGTCTGAGTTCTGGT 4980
     ----:----|----:----|----:----|----:----|----:----|----:----|

     H  L  Y  G  V  A  L  M  N  *  S  L  C  N  C  V  I  W  F  I    F1
      I  Y  M  E  *  L  *  *  T  E  A  Y  V  I  V  S  F  G  S  F   F2
       F  I  W  S  S  F  N  E  L  K  L  M  *  L  C  H  L  V  H  L  F3
4981 CATTTATATGGAGTAGCTTTAATGAACTGAAGCTTATGTAATTGTGTCATTTGGTTCATT 5040
     ----:----|----:----|----:----|----:----|----:----|----:----|

     *  C  C  H  A  L  S  I  A  A  T  H  T  Q  F  C  C  T  L  Y    F1
      N  A  V  M  R  C  Q  L  L  R  L  I  P  S  F  V  A  R  Y  I   F2
       M  L  S  C  A  V  N  C  C  D  S  Y  P  V  L  L  H  V  I  F  F3
5041 TAATGCTGTCATGCGCTGTCAATTGCTGCGACTCATACCCAGTTTTGTTGCACGTTATAT 5100
     ----:----|----:----|----:----|----:----|----:----|----:----|

     L  Y  *  C  V  T  R  Q  Q  Q  Q  D  S  A  C  D  Q  A  R  D    F1
      Y  I  N  A  L  P  D  S  N  N  K  T  Q  R  A  T  R  H  V  I   F2
       I  L  M  R  Y  Q  T  A  T  T  R  L  S  V  R  P  G  T  *  S  F3
5101 TTATATTAATGCGTTACCAGACAGCAACAACAAGACTCAGCGTGCGACCAGGCACGTGAT 5160
     ----:----|----:----|----:----|----:----|----:----|----:----|

     H  Y  *  P  R  K  H  S  T  R  K  V  S  W  L  L  S  M  G  *    F1
      I  I  S  P  E  N  T  L  R  G  R  S  A  G  C  C  L  W  A  N   F2
       L  L  A  Q  K  T  L  Y  E  E  G  Q  L  A  A  V  Y  G  L  M  F3
5161 CATTATTAGCCCAGAAAACACTCTACGAGGAAGGTCAGCTGGCTGCTGTCTATGGGCTAA 5220
     ----:----|----:----|----:----|----:----|----:----|----:----|

     C  W  T  C  Q  P  L  C  H  R  L  M  S  A  Y  Y  K  D  C  T    F1
      A  G  P  A  S  H  Y  A  I  A  L  C  R  P  I  I  R  T  A  L   F2
       L  D  L  P  A  I  M  P  S  P  Y  V  G  L  L  *  G  L  H  W  F3
5221 TGCTGGACCTGCCAGCCATTATGCCATCGCCTTATGTCGGCCTATTATAAGGACTGCACT 5280
     ----:----|----:----|----:----|----:----|----:----|----:----|

     G  S  G  E  C  S  P  H  A  A  T  A  G  H  H  G  H  C  F  M    F1
      G  Q  A  N  V  R  P  M  R  R  Q  L  A  T  M  A  I  A  S  W   F2
       V  R  R  M  F  A  P  C  G  D  S  W  P  P  W  P  L  L  H  G  F3
5281 GGGTCAGGCGAATGTTCGCCCCATGCGGCGACAGCTGGCCACCATGGCCATTGCTTCATG 5340
     ----:----|----:----|----:----|----:----|----:----|----:----|

     G  C  Q  F  I  F  R  L  Y  F  E  F  A  S  Q  L  *  K  P  L    F1
      V  A  N  L  Y  L  D  F  I  L  N  L  Q  V  N  C  K  S  H  L   F2
       L  P  I  Y  I  *  T  L  F  *  I  C  K  S  I  V  K  A  T  *  F3
5341 GGTTGCCAATTTATATTTAGACTTTATTTTGAATTTGCAAGTCAATTGTAAAAGCCACTT 5400
     ----:----|----:----|----:----|----:----|----:----|----:----|

     D  N  S  S  C  E  Q  S  S  S  S  S  K  A  K  R  T  F  K  G  A    F1
      T  T  Q  V  A  S  R  V  A  A  A  K  Q  K  G  P  L  R  V  P   F2
       Q  L  K  L  R  A  E  *  Q  Q  Q  S  K  K  D  L  *  G  C  Q  F3
5401 GACAACTCAAGTTGCGAGCAGAGTAGCAGCAGCAAAGCAAAAAGGACCTTTAAGGGTGCC 5460
     ----:----|----:----|----:----|----:----|----:----|----:----|

     N  Q  L  A  T  P  P  S  P  V  T  R  R  P  F  L  S  V  A  S    F1
      I  S  *  P  H  P  L  R  Q  L  H  A  A  H  F  C  Q  L  Q  V   F2
       S  V  S  H  T  P  F  A  S  Y  T  P  P  I  F  V  S  C  K  Y  F3
5461 AATCAGTTAGCCACACCCCCTTCGCCAGTTACACGCCGCCCATTTTTGTCAGTTGCAAGT 5520
     ----:----|----:----|----:----|----:----|----:----|----:----|
```

```
     T  A  Q  I  V  N  L  Q  T  T  K  K  K  T  F  *  Q  L  T  Q    F1
      Q  R  K  L  S  T  C  K  Q  R  K  K  K  H  F  D  S  *  L  K   F2
       S  A  N  C  Q  L  A  N  N  E  K  K  N  I  L  T  V  D  S  K  F3
5521 ACAGCGCAAATTGTCAACTTGCAAACAACGAAAAAAAAAACATTTTGACAGTTGACTCAA 5580
     ----:----|----:----|----:----|----:----|----:----|----:----|

     S  L  S  F  V  H  L  E  M  K  I  L  C  M  R  L  D  N  I  L    F1
      V  *  A  L  S  T  W  K  *  K  Y  C  V  C  D  *  T  I  S  C   F2
       F  E  L  C  P  L  G  N  E  N  I  V  Y  A  T  R  Q  Y  P  A  F3
5581 AGTTTGAGCTTTGTCCACTTGGAAATGAAAATATTGTGTATGCGACTAGACAATATCCTG 5640
     ----:----|----:----|----:----|----:----|----:----|----:----|

     Q  T  K  R  P  N  A  I  Q  I  L  C  N  I  H  L  F  I  N  S    F1
      K  P  S  A  Q  M  P  Y  K  F  Y  V  I  Y  I  Y  L  L  I  Q   F2
       N  Q  A  P  K  C  H  T  N  F  M  *  Y  T  F  I  Y  *  F  K  F3
5641 CAAACCAAGCGCCCAAATGCCATACAAATTTTATGTAATATACATTTATTTATTAATTCA 5700
     ----:----|----:----|----:----|----:----|----:----|----:----|

     K  L  S  F  *  S  *  I  N  Q  R  C  A  T  K  T  P  S  I  R    F1
      N  S  V  F  N  H  K  S  T  R  D  V  Q  L  R  H  P  A  L  E   F2
       T  Q  F  L  I  I  N  Q  P  E  M  C  N  *  D  T  Q  H  *  N  F3
5701 AAACTCAGTTTTTAATCATAAATCAACCAGAGATGTGCAACTAAGACACCCAGCATTAGA 5760
     ----:----|----:----|----:----|----:----|----:----|----:----|

     I  *  I  S  I  D  W  E  I  N  P  I  F  N  I  C  I  I  N  F    F1
      Y  K  L  V  *  I  G  K  *  I  P  Y  L  I  F  V  *  L  I  L   F2
       I  N  *  Y  R  L  G  N  K  S  H  I  *  Y  L  Y  N  *  F  *  F3
5761 ATATAAATTAGTATAGATTGGGAAATAAATCCCATATTTAATATTTGTATAATTAATTTT 5820
     ----:----|----:----|----:----|----:----|----:----|----:----|

     N  W  K  Y  I  *  L  P  P  K  V  *  N  T  *  T  C  Y  F  K    F1
      I  G  N  I  F  N  Y  P  Q  K  Y  K  I  L  R  H  V  I  L  K   F2
       L  E  I  Y  L  T  T  P  K  S  I  K  Y  L  D  M  L  F  *  N  F3
5821 AATTGGAAATATATTTAACTACCCCCAAAAGTATAAAATACTTAGACATGTTATTTTAAA 5880
     ----:----|----:----|----:----|----:----|----:----|----:----|

     T  E  A  K  K  Y  T  Q  P  L  K  G  N  T  Q  V  G  Q  Q  L    F1
      Q  K  L  K  S  T  H  S  L  *  K  G  I  H  K  S  G  S  S  *   F2
       R  S  *  K  V  H  T  A  F  E  R  E  Y  T  S  R  A  A  V  K  F3
5881 ACAGAAGCTAAAAAGTACACACAGCCTTTGAAAGGGAATACACAAGTCGGCAGCAGTTA 5940
     ----:----|----:----|----:----|----:----|----:----|----:----|

     S  K  W  A  D  T  L  S  V  R  S  S  Y  A  F  Y  L  K  T  P    F1
      V  S  G  Q  I  P  *  V  F  V  P  A  M  H  S  T  *  R  R  R   F2
       *  V  G  R  Y  L  K  C  S  F  Q  L  C  I  L  L  K  D  A  E  F3
5941 AGTAAGTGGGCAGATACCTTAAGTGTTCGTTCCAGCTATGCATTCTACTTAAAGACGCCG 6000
     ----:----|----:----|----:----|----:----|----:----|----:----|

     K  N  F  A  L  L  E  K  L  R  E  A  W  T  I  P  K  K  Q  L    F1
      K  T  L  L  S  W  K  S  Y  A  K  R  G  Q  F  Q  R  S  S  Y   F2
       K  L  C  S  P  G  K  V  T  R  S  V  D  N  S  K  E  A  A  T  F3
6001 AAAAACTTTGCTCTCCTGGAAAAGTTACGCGAAGCGTGGACAATTCCAAAGAAGCAGCTA 6060
     ----:----|----:----|----:----|----:----|----:----|----:----|

     Q  I  S  S  N  T  C  Q  S  S  S  N  S  W  K  T  K  T  E  K    F1
      K  S  V  Q  I  L  V  R  V  R  Q  I  A  G  R  R  K  Q  R  K   F2
       N  Q  F  K  Y  L  S  E  F  V  K  *  L  E  D  E  N  R  E  R  F3
6061 CAAATCAGTTCAAATACTTGTCAGAGTTCGTCAAATAGCTGGAAGACGAAAACAGAGAAA 6120
     ----:----|----:----|----:----|----:----|----:----|----:----|

     E  T  S  G  C  K  *  C  E  K  T  L  S  A  N  V  P  K  W  K    F1
      R  R  V  V  A  S  S  A  K  K  L  F  Q  Q  M  F  L  N  G  R   F2
       D  E  W  L  Q  V  V  R  K  N  S  F  S  K  C  S  *  M  E  D  F3
6121 GAGACGAGTGGTTGCAAGTAGTGCGAAAAAACTCTTTCAGCAAATGTTCCTAAATGGAAG 6180
     ----:----|----:----|----:----|----:----|----:----|----:----|
```

```
     T  N  P  I  T  R  F  I  *  N  *  L  I  Y  M  I  R  *  Y  V     F1
      L  T  R  *  P  D  L  F  K  I  N  L  Y  I  *  *  D  N  M  F    F2
       *  P  D  N  P  I  Y  L  K  L  T  Y  I  Y  D  K  I  I  C  L   F3
6181 ACTAACCCGATAACCCGATTTATTTAAAATTAACTTATATATATGATAAGATAATATGTT 6240
     ----:----|----:----|----:----|----:----|----:----|----:----|

     *  A  E  R  R  V  G  N  L  Y  E  N  Q  L  E  N  V  V  S  R     F1
      K  Q  S  V  E  W  A  I  F  M  K  I  N  L  K  M  L  F  H  E    F2
       S  R  A  *  S  G  Q  S  L  *  K  S  T  *  K  C  C  F  T  K   F3
6241 TAAGCAGAGCGTAGAGTGGGCAATCTTTATGAAAATCAACTTGAAAATGTTGTTTCACGA 6300
     ----:----|----:----|----:----|----:----|----:----|----:----|

     R  E  D  Q  E  C  F  L  A  H  V  Y  L  V  A  G  T  *  M  L     F1
      E  K  T  R  S  A  F  W  H  M  F  I  *  L  L  V  H  K  C  *    F2
       R  R  P  G  V  L  S  G  T  C  L  F  S  C  W  Y  I  N  A  S   F3
6301 AGAGAAGACCAGGAGTGCTTTCTGGCACATGTTTATTTAGTTGCTGGTACATAAATGCTA 6360
     ----:----|----:----|----:----|----:----|----:----|----:----|

     A  I  F  T  Y  T  K  T  *  Y  I  F  F  F  Y  A  *  F  Q  V     F1
      L  Y  L  H  I  L  R  L  N  I  Y  F  F  S  M  H  N  S  R  S    F2
       Y  I  Y  I  Y  *  D  L  I  Y  I  F  F  L  C  I  I  P  G  H   F3
6361 GCTATATTTACATATACTAAGACTTAATATATATTTTTTTCTATGCATAATTCCAGGTC 6420
     ----:----|----:----|----:----|----:----|----:----|----:----|

     T  F  I  S  R  *  F  Q  F  *  C  A  A  N  A  A  H  F  S  G     F1
      R  S  Y  R  G  D  S  N  F  N  A  L  L  T  P  L  T  F  L  V    F2
       V  H  I  A  V  I  P  I  L  M  R  C  *  R  R  S  L  F  W  S   F3
6421 ACGTTCATATCGCGGTGATTCCAATTTTAATGCGCTGCTAACGCCGCTCACTTTTCTGGT 6480
     ----:----|----:----|----:----|----:----|----:----|----:----|

     H  V  L  Q  F  G  C  *  S  P  A  L  C  F  S  Q  S  Q  F  S     F1
      T  Y  F  N  S  G  V  N  P  L  L  Y  A  F  L  S  R  N  F  R    F2
       R  T  S  I  R  V  L  I  P  C  F  M  L  F  S  V  A  I  F  A   F3
6481 CACGTACTTCAATTCGGGTGTTAATCCCCTGCTTTATGCTTTTCTCAGTCGCAATTTTCG 6540
     ----:----|----:----|----:----|----:----|----:----|----:----|

     Q  G  H  E  G  A  V  A  L  L  L  E  E  G  Q  G  Q  V  V  L     F1
      K  G  M  K  E  L  L  L  C  S  W  K  K  G  K  G  K  S  S  S    F2
       R  A  *  R  S  C  C  S  A  P  G  R  R  A  R  A  S  R  P  P   F3
6541 CAAGGGCATGAAGGAGCTGTTGCTCTGCTCCTGGAAGAAGGGCAAGGGCAAGTCGTCCTC 6600
     ----:----|----:----|----:----|----:----|----:----|----:----|

     Q  F  V  N  A  S  Q  T  Q  G  A  A  G  K  C  *  H  Q  I  L     F1
      N  S  S  M  H  H  K  R  K  A  L  Q  V  S  A  N  T  K  Y  Y    F2
       I  R  Q  C  I  T  N  A  R  R  C  R  *  V  L  T  P  N  I  I   F3
6601 CAATTCGTCAATGCATCACAAACGCAAGGCGCTGCAGGTAAGTGCTAACACCAAATATTA 6660
     ----:----|----:----|----:----|----:----|----:----|----:----|

     S  S  H  F  K  Y  L  A  L  S  N  T  W  S  V  F  R  L  S  A     F1
      R  V  T  L  S  I  W  H  *  A  T  L  G  L  C  F  D  S  L  Q    F2
       E  S  L  *  V  F  G  I  E  Q  H  L  V  C  V  S  T  L  C  R   F3
6661 TCGAGTCACTTTAAGTATTTGGCATTGAGCAACACTTGGTCTGTGTTTCGACTCTCTGCA 6720
     ----:----|----:----|----:----|----:----|----:----|----:----|

     D  P  L  A  A  H  G  Y  H  A  H  W  Q  R  A  A  M  M  P  E     F1
      T  H  S  L  P  T  D  T  T  H  I  G  N  E  Q  L  *  C  P  S    F2
       P  T  R  C  P  R  I  P  R  T  L  A  T  S  S  Y  D  A  R  A   F3
6721 GACCCACTCGCTGCCCACGGATACCACGCACATTGGCAACGAGCAGCTATGATGCCCGAG 6780
     ----:----|----:----|----:----|----:----|----:----|----:----|

     L  L  E  D  A  L  Q  A  L  R  S  S  L  V  E  P  I  G  G  R     F1
      C  W  K  T  H  F  K  L  F  G  R  R  S  *  N  P  S  V  A  A    F2
       A  G  R  R  T  S  S  S  S  V  V  A  R  R  T  H  R  W  P  L   F3
6781 CTGCTGGAAGACGCACTTCAAGCTCTTCGGTCGTCGCTCGTAGAACCCATCGGTGGCCGC 6840
```

```
     ----:----|----:----|----:----|----:----|----:----|----:----|

     S  C  R  S  H  L  A  A  I  L  L  *  F  S  K  K  P              F1
      R  A  G  R  I  *  R  Q  Y  Y  C  D  S  L  R  S  X             F2
       V  P  V  A  S  S  G  N  I  I  V  I  L  *  E  A               F3
6841 TCGTGCCGGTCGCATCTAGCGGCAATATTATTGTGATTCTCTAAGAAGCC 6890
```
