## Supplementary material for "The incidence of candidate binding sites for β-arrestin in Drosophila neuropeptide GPCRs": S8 Text

Moody R

*D melanogaster* moody has 2 predicted protein isoforms annotated (PA and PC), but *D virilis* moody has only one

These two isoforms result from alternative splicing and use of different reading frames from a common exon domain.

From blastp searches, I only retrieve the PC moody isoform from *D virilis*.

In *D virilis* genomic DNA, here I find a potential PA isoform, as in *D melanogaster*, as an alternative splice site which catches a different reading frame.

*D virilis moody* gDNA:
https://www.ncbi.nlm.nih.gov/genome/gdv/browser/protein/?id=XP_002057362.1
bp #1 = bp 16633201of shotgun sequence found in the link

The PA isoform is displayed in **GREEN** and the PC isoform in **RED**. The corresponding predicted splice acceptors sites in similar colors.

### Sequences alignment:

```
DNA:  ggtatgtaccatgggctatggccagttcagttgcatgcgaaccacatccactaagtcacg
+1fr: ·G··M··Y··H··G··L··W··P··V··Q··L··H··A··N··H··I··H··*··V··T·
+2fr:  ·V··C··T··M··G··Y··G··Q··F··S··C··M··R··T··T··S··T··K··S··R·
+3fr:   ·Y··V··P··W··A··M··A··S··S··V··A··C··E··P··H··P··L··S··H··A·
```

```
DNA:  ctatccgttcgttttagagaaatggaaagataccggattgagcaacaaccacagccgcac
+1fr: ·L··S··V··R··F··R··E··M··E··R··Y··R··I··E··Q··Q··P··Q··P··H·
+2fr:  ·Y··P··F··V··L··E··K··W··K··D··T··G··L··S··N··N··H··S··R··T·
+3fr:   ·I··R··S··F··*··R··N··G··K··I··P··D··*··A··T··T··T··A··A··P·
```

```
DNA:  ccttgtctcccagatgtccgccggcgcagcagcagcagcagcggcggcggcggcggc
+1fr: ·P··C··L··P··D··V··R··R··R··S··S··S··S··S··G··G··G··G··G··G·
+2fr:  ·L··V··S··Q··M··S··A··G··A··A··A··A··A··A··A··A··A··A··A··A
+3fr:   ·L··S··P··R··C··P··P··A··Q··Q··Q··Q··Q··R··R··R··R··R··R··R·
```

```
DNA:  ggttgcagcggcatcggcgtcgacgccgccgccgtcgcccgcatccgattgccagtcctt
+1fr: ·G··C··S··G··I··G··V··D··A··A··A··V··A··R··I··R··L··P··V··L·
+2fr:  ·V··A··A··A··S··A··S··T··P··P··P··S··P··A··S··D··C··Q··S··L·
+3fr:   ·L··Q··R··H··R··R··R··R··R··R··R··R··P··H··P··I··A··S··P··C·
```

```
DNA:  gccggcgaccaaagcgtcgcctcagtcactggagctgatgtcccgatgtccagatctaat
+1fr: ·A··G··D··Q··S··V··A··S··V··T··G··A··D··V··P··M··S··R··S··N·
+2fr:  ·P··A··T··K··A··S··P··Q··S··L··E··L··M··S··R··C··P··D··L··I·
+3fr:   ·R··R··P··K··R··R··L··S··H··W··S··*··C··P··D··V··Q··I··*··*·

DNA:  aaatcgaacgaatcaaccgccattggttacgcccccgccgccatcggtaacgctggcaac
+1fr: ·K··S··N··E··S··T··A··I··G··Y··A··P··A··A··I··G··N··A··G··N·
+2fr:  ·N··R··T··N··Q··P··P··L··V··T··P··P··P··P··S··V··T··L··A··T·
+3fr:   ·I··E··R··I··N··R··H··W··L··R··P··R··R··H··R··*··R··W··Q··R·

DNA:  ggctcaaccgagcagcagcaacagcagcaacagcagcaacagcggcggcagtcggctgcc
+1fr: ·G··S··T··E··Q··Q··Q··Q··Q··Q··Q··Q··Q··Q··R··R··Q··S··A··A·
+2fr:  ·A··Q··P··S··S··S··N··S··S··N··S··S··N··S··G··G··S··R··L··P·
+3fr:   ·L··N··R··A··A··A··T··A··A··T··A··A··T··A··A··A··V··G··C··R·

DNA:  gctcaagaagagcaatcactcctacatcaacaacggcttcaacagcagcgggcacagcca
+1fr: ·A··Q··E··E··Q··S··L··L··H··Q··Q··R··L··Q··Q··Q··R··A··Q··P·
+2fr:  ·L··K··K··S··N··H··S··Y··I··N··N··G··F··N··S··S··G··H··S··Q·
+3fr:   ·S··R··R··A··I··T··P··T··S··T··T··A··S··T··A··A··G··T··A··R·

DNA:  gaacagcgccgtttatcggcccgcagctggctccccgccgtcggcagcggcggcgccatt
+1fr: ·E··Q··R··R··L··S··A··R··S··W··L··P··A··V··G··S··G··G··A··I·
+2fr:  ·N··S··A··V··Y··R··P··A··A··G··S··P··P··S··A··A··A··A··P··L·
+3fr:   ·T··A··P··F··I··G··P··Q··L··A··P··R··R··R··Q··R··R··R··H··C·

DNA:  gcgtcgcatcaccatggtggggacgatatcatactggaggaggaggaactgcccgcagt
+1fr: ·A··S··H··H··H··G··G··G··R··Y··H··T··G··G··G··G··T··A··R··S·
+2fr:  ·R··R··I··T··M··V··G··D··D··I··I··L··E··E··E··E··L··P··A··V·
+3fr:   ·V··A··S··P··W··W··G··T··I··S··Y··W··R··R··R··N··C··P··Q··C·

DNA:  gcccagtccgacggcgacggcgctgtcagtcagcaaagtaaccaaatcgcccatctacat
+1fr: ·A··Q··S··D··G··D··G··A··V··S··Q··Q··S··N··Q··I··A··H··L··H·
+2fr:  ·P··S··P··T··A··T··A··L··S··V··S··K··V··T··K··S··P··I··Y··M
+3fr:   ·P··V··R··R··R··R··R··C··Q··S··A··K··*··P··N··R··P··S··T··*·

DNA:  gaacgtcaacagtccgaagagaaaccaatcttatagcgacaaggcttcggtcaaggatgc
+1fr: ·E··R··Q··Q··S··E··E··K··P··I··L··*··R··Q··G··F··G··Q··G··C·
+2fr:  ·N··V··N··S··P··K··R··N··Q··S··Y··S··D··K··A··S··V··K··D··A·
+3fr:   ·T··S··T··V··R··R··E··T··N··L··I··A··T··R··L··R··S··R··M··H·

DNA:  actgcagcaggcatctgtcccatccgaccagggcaaggatcagcaggatgctagcaaagt
+1fr: ·T··A··A··G··I··C··P··I··R··P··G··Q··G··S··A··G··C··*··Q··S·
+2fr:  ·L··Q··Q··A··S··V··P··S··D··Q··G··K··D··Q··Q··D··A··S··K··V·
+3fr:   ·C··S··R··H··L··S··H··P··T··R··A··R··I··S··R··M··L··A··K··C·

DNA:  gcccataaagtttccaaatccaaaagactaacgaaatgctttaattccaaatatcgtaac
+1fr: ·A··H··K··V··S··K··S··K··R··L··T··K··C··F··N··S··K··Y··R··N·
+2fr:   P··I··K··F··P··N··P··K··D··*··R··N··A··L··I··P··N··I··V··T·
```

```
+3fr:    ·P··*··S··F··Q··I··Q··K··T··N··E··M··L··*··F··Q··I··S··*··L·


DNA:  tgttacattccccccacgccccaccccaccaccttcctcc
+1fr: ·C··Y··I··P··P··T··P··H··P··T··T··F··L·
+2fr:  ·V··T··F··P··P··R··P··T··P··P··P··S··S·
+3fr:   ·L··H··S··P··H··A··P··P··H··H··L··P·
```
