## Supplementary material for "The incidence of candidate binding sites for β-arrestin in Drosophila neuropeptide GPCRs": S9 Text

***S9. Text* Multi-species analysis of Proc R**

**Supporting Figure 11**

CLUSTAL Line-ups; Genbank Reference IDs below
5th, 6th and 7th Predicted TM domains in YELLOW
BBS sequences in RED

```
Grimshawi          -----MTT-----------------------LTATSTAMPGDGNAAAKMFSDADVAEVRH        32
Mojavensis         --MSTLAA-----------------------ATSTSTAMPVDAIAAAKMFSDADVAEVRH        35
Virilism           ---MTTLA-----------------------ATSTSTAMPVDGIGTAKMFSDADVAEVRH        34
Bipectinate        MTMFSLLTATLTATATATATATVTATATSTESEAEAATLNETEAAGTEMFSDADVVEVRR        60
Anannassae         MTMFSLLTATLTATATATATATATVTPAE--------AETEAEAAGSEMFSDADVAEVRR        52
Serrata            MTMSS-----------------------------TSSQSSDGNETVGEMFSEADTAEVRH        31
Kikkawei           --MSS-----------------------------TSSQSSDGNETVGEMFSEAETAEVRH        29
Fichsuphila        --MSSTSTS------------------TTTATLTATEAATVGNATVGEMFSDAEMAEVRH        40
Eugracilis         MTMSTTSTLTA--------------------TATATATETLGNATVGEMFSDADMAEVRH        40
Rhopaloa           ----------------------------------MEVAATLGNATVGEMFSDADMAEVRH        26
Elegans            --MS----------------------STSTSTEAAPEAATLGNATVGEMFSDADMAEVRH        36
Takahashi          MTMSTTSTATL--------------TEAA--AAAAAEAATLGNATVGEMFSDADMAEVRH        44
Suzuki             MTMSSTSTATL--------------TEAAAEAAAAAEAATLGNATVGEMFSDADMAEVRH        46
Biarmipes          MTMSSTVTATL--------------TEAA--AAAEAEAATLGNATAGEMFSDADMAEVRH        44
Erecta             MTMSSTSTATA--------------TATST----ATATLAEANATVGEMFSDADMAEVRH        42
Melanogaster       MTMSS--TSTA--------------TATST----ATATLDEANATVGEMFSDADMAEVRH        40
Simulans           MTM----SSTS--------------TATST----ATATLAEANATVGEMFSDADMAEVRH        38
Mauritania         MTM----SSTS--------------TATST----ATATLAEANATVGEMFSDADMAEVRH        38
Sechellia          MTM----SSTS--------------TATST----ATATLAEANATVGEMFSDADMAEVRH        38
                                                                 :***:*:  .***:

Grimshawi          VVQRILVPCVFVIGLLGNSVSIYVLTRKRMRCTTNIYLSALAITDIAYLTFVLILSLKHY        92
Mojavensis         VVQRILVPCVFVIGLLGNSVSIYVLTRKRMRCTTNIYLTALAITDIAYLTFVLILSLKHY        95
Virilism           VVQRILVPCVFVIGLLGNSVSIYVLTRKRMRCTTNIYLSALAITDIAYLTFVLILSLKHY        94
Bipectinate        VVQRILVPCVFVIGLLGNSVSIYVLTRKRMRCTTNIYLTALAITDIAYLTFQLILSFQHY        120
Anannassae         VVQRILVPCVFVIGLLGNSVSIYVLTRKRMRCTTNIYLSALAITDIAYLTFQLILSLQHY        112
Serrata            VVQRILVPCVFVIGLLGNSVSIYVLTRKRMRCTTNIYLTALAITDIAYLTFQLILSLQHY        91
Kikkawei           VVQRILVPCVFVIGLLGNSVSIYVLTRKRMRCTTNIYLTALAITDIAYLTFQLILSLQHY        89
Fichsuphila        VVQRILVPCVFVIGLLGNSVSIYVLTRKRMRCTTNIYLTALAITDIAYLTCQLILSIQHY        100
Eugracilis         VVQRILVPCVFVIGLLGNSVSIYVLTRKRMRCTTNIYLTALAITDIAYLTCQLILSLQHY        100
Rhopaloa           VVQRILVPCVFVIGLLGNSVSIYVLTRKRMRCTTNIYLTALAITDIAYLTCQLILSLQHY        86
Elegans            VVQRILVPCVFVIGLLGNSVSIYVLTRKRMRCTTNIYLTALAITDIAYLTCQLILSLQHY        96
Takahashi          VVQRILVPCVFVIGLLGNSVSIYVLTRKRMRCTTNIYLTALAITDIAYLTCQLILSLQHY        104
Suzuki             VVQRILVPCVFVIGLLGNSVSIYVLTRKRMRCTTNIYLTALAITDIAYLTCQLILSLQHY        106
Biarmipes          VVQRILVPCVFVIGLLGNSVSIYVLTRKRMRCTTNIYLTALAITDIAYLTCQLILSLQHY        104
Erecta             VVQRILVPCVFVIGLLGNSVSIYVLTRKRMRCTTNIYLTALAITDIAYLTCQLILSLQHY        102
Melanogaster       VVQRILVPCVFVIGLLGNSVSIYVLTRKRMRCTTNIYLTALAITDIAYLTCQLILSLQHY        100
Simulans           VVQRILVPCVFVIGLLGNSVSIYVLTRKRMRCTTNIYLTALAITDIAYLTCQLILSLQHY        98
Mauritania         VVQRILVPCVFVIGLLGNSVSIYVLTRKRMRCTTNIYLTALAITDIAYLTCQLILSLQHY        98
Sechellia          VVQRILVPCVFVIGLLGNSVSIYVLTRKRMRCTTNIYLTALAITDIAYLTCQLILSLQHY        98
                   **************************************:***********  ****::**

Grimshawi          EYIKYHCELYWRLYGFVMWLCDACAYISIYIAVCFTIERFIAIRYPLKRQTFCTESLAKK        152
Mojavensis         EYIKYHCELYWLLYGFVVWLCDACAYISIYIAVCFTIERFIAIRYPLKRQTFCTESLAKK        155
Virilism           EYIKYHCELYWRLYGFIMWLCDACAYISIYIAVCFTIERFIAIRYPLKRQTFCTESLAKK        154
Bipectinate        DYVKYHCEIYWQFYGYFVWLCDSCAYISIYIAVCFTIERFIAIRYPLKRQTFCTESLAKK        180
Anannassae         DYFKYHSEIYWQLYGYVVWLCDSSAYISIYIAVCFTIERFIAIRYPLKRQTFCTESLAKK        172
Serrata            DYVKYHCEIYWQLYGIFVWLCDSSGYISIYIAVCFTIERFIAIRYPLKRQTFCTESLAKK        151
Kikkawei           DYVKYHCEIYWQLYGIFVWLCDSSGYISIYIAVCFTIERFIAIRYPLKRQTFCTESLAKK        149
Fichsuphila        DYAKYHLQIFWELYGYFVWLCDSFGYISIYIAVCFTIERFIAIRYPLKRQTFCTESLAKK        160
Eugracilis         DYTKFHVEIYWQLYGYFVWLCDSFGYISIYIAVCFTIERFIAIRYPLKRQTFCTESLAKK        160
Rhopaloa           DYPKYHFEIYYQLFGYFVWLCDSFGYISIYIAVCFTIERFIAIRYPLKRQTFCTESLAKK        146
Elegans            DYPKYHLEIYYQLYGYFVWLCDSFGYISIYIAVCFTIERFIAIRYPLKRQTFCTESLAKK        156
Takahashi          DYPKYNFKIYWQLYGYFVWLCDSFGYISIYIAVCFTIERFIAIRYPLKRQTFCTESLAKK        164
Suzuki             DYPKYHLKIYWQLYGYFVWLCDSFGYISIYIAVCFTIERFIAIRYPLKRQTFCTESLAKK        166
Biarmipes          DYPKYHLKIYWQLYGYFVWLCDSFGYISIYIAVCFTIERFIAIRYPLKRQTFCTESLAKK        164
Erecta             DYTKYHLKLYWQLYGYFVWLCDSFGYISIYIAVCFTIERFIAIRYPLKRQTFCTESLAKK        162
```

```
Melanogaster     DYPKYHFKLYWQLYGYFVWLCDSFGYISIYIAVCFTIERFIAIRYPLKRQTFCTESLAKK     160
Simulans         DYPKYHFKLYWQLYGYFVWLCDSFGYISIYIAVCFTIERFIAIRYPLKRQTFCTESLAKK     158
Mauritania       DYPKYHFKLYWQLYGYFVWLCDSFGYISIYIAVCFTIERFIAIRYPLKRQTFCTESLAKK     158
Sechellia        DYPKYHFKLYWQLYGYFVWLCDSFGYISIYIAVCFTIERFIAIRYPLKRQTFCTESLAKK     158
                 :* *:: :::: ::* .:****: .***********************************

Grimshawi        VIAAVALFCLLTTLSTAFEHTYDIDWKLIDGAYRPCNQTLANVSPTPAH-----------     201
Mojavensis       VIAAVALFCLLTTLSTAFEHTYDINWKLIDGAYRPCNLTLANVSPMPTPTPTSTPMPTPA     215
Virilism         VIAAVALFCLLTTLSTAFEHTYDINWKLIDGAYRPCNLTLANVSPTPAQ-----------     203
Bipectinate      VIAAVALFCLLSTLSTAFEHSFARKFRLIDDDYRPCNQTLANVSPQPLALTPSLSMQAPS     240
Anannassae       VIAAVALFCLLSTLSTAFEHSYARGYRLIDDAYRPCNQTLANVSPQPQPLIPSLHMQAPS     232
Serrata          VIAAVSLFCLLSTLSTAFEHTIKVDWKLIDGAYQPCNQTLANVSPTPPSTS---------     202
Kikkawei         VIAAVSLFCLLSTLSTAFEHTIKVDWKLIDGAYQNCNQTLANVSPMPPSTS---------     200
Fichsuphila      VIAAVAVFCLLSTLSTAFEHTITVGWKLIDGAYQPCNQTEANFSPMPPSFALG------T     214
Eugracilis       VIAAVAFFCLLSTLSTAFEHTITVGSKLIDDAYQPCNQTLANYSPMPTS--S--------     210
Rhopaloa         VIAAVALFCLLSTLSTAFEHTITVGYKLIDDAYQPCNLTEANISPTPPAPPF--------     198
Elegans          VIAAVAVFCLLSTLSTAFEHTITVGWKLIDDAYQPCNQTVANFSPTPPVPSF--------     208
Takahashi        VIAAVAIFCLLSTLSTAFEHTITVGWKQIDDAYQPCNQTVANISPTPPP-S---------     214
Suzuki           VIAAVAVFCLLSTLSTAFEHTITVGSKLIDDAYQPCNQTVANISPTPPP-S---------     216
Biarmipes        VIAAVAVFCLLSTLSTAFEHTITVGSRLIDEAYQPCNQSVANISPTPPA-F---------     214
Erecta           VIAAVAIFCLLSTLSTAFEHTITMGSRQIDDAYQPCNQTVANISPTPPPPS---------     213
Melanogaster     VIAAVAIFCLLSTLSTAFEHTITIGTRQIDDAYQPCNQTVANISPMPP-PP---------     210
Simulans         VIAAVAIFCLLSTLSTAFEHTITIGTRQIDDAYQPCNQTLANISPTPPPPP---------     209
Mauritania       VIAAVAIFCLLSTLSTAFEHTITIGTRQIDDAYQPCNQTLANISPTPPPPP---------     209
Sechellia        VIAAVAIFCLLSTLSTAFEHTITIGTRQIDDAYQPCNQTLANISPTPPPPL---------     209
                 *****:.****:********:     : **  *: ** : ** ** *

Grimshawi        -------------QTPGT-AWHVNENASMSN-------TTPS------NVEPPLDLGSGA     234
Mojavensis       -------------QTMTT-AWHIEEPAGNPA-------TSRYLEPFDLSSDSGSGSGSGA     254
Virilism         -------------PTLAT-AWHVDEHANNAA-------TPRYLEPFDLSS----GSGSGA     238
Bipectinate      PLDSTTPLPTAMLPTPVTPHMQDLT-----------RKPEKPLGV--LVSDVFSGGGSGD     287
Anannassae       PSASTTPLPTVMLPTPVTPHWQDSRDSQD----LAGRSTEKPLGGLFLAADVFSGGGSGD     288
Serrata          ---LTPALSTPPLVTPAT-IWQEQDQERESLQDFFTT-ESSSVKSSRLLLDFGSGSG--D     255
Kikkawei         ---VTPALSTPPMVTPPT-IWKEKEL----EQNFFTQ-ESSTAKSSRLLFDFGSGSG--D     249
Fichsuphila      PPLATPPLATPPLATPPT-VWQDFSTES-P---------TAIAGSSSQLIDWGSGSG--D     261
Eugracilis       -VAATPPLATPPLPTPAT-IWQSTELNTES---------TTTAGKSNVLIDWGSGSG--D     257
Rhopaloa         -AVATPPLATPPRPTPAT-I--SQDFTTES---------T-TAGSSNLLVDWGSGSG--D     242
Elegans          -AVATPPLATPPLPTPAT-I--SQHFSTES---------T-TAGSSNLLFDWGSGSG--D     252
Takahashi        -AAATPPLVTPPLPTPAT-IWQSQDLTTES---------T-TAGSSNLLADWGSGSG--D     260
Suzuki           -VAATPPLVTPPLSTPAT-IWQSQDFTTES---------T-TAGSSNLLVDWGSGSG--D     262
Biarmipes        -VAATPPLVTPPLSTPAT-IWKSQDSTTES---------T-TAGSSNLLVDWGSGSG--D     260
Erecta           -AAATPPLATPPLPTPAT-IWQSQDYSTES---------T-TAGSTSLLVDWGSGSGDGD     261
Melanogaster     -VAVTPPLATPPLPTPAT-IWQSPDSAMES---------T-TSGSSNQLVDWGSGSG--D     256
Simulans         -VAATPPLATPPLPTPAT-VWQSPDLAMES---------T-TAGSSNLLVDWGSGSG--D     255
Mauritania       -VAATPPLATPPLPTPAT-AWQSPDFAMES---------T-TAGSSNLLVDWGSGSG--D     255
Sechellia        -GEATPPLATPPLPTPAT-VWQSPDFAMES---------T-TAGSSNLLVDWGSGSG--D     255
                               *  *                                    . *

Grimshawi        -GESDHIPSRQRRLPASTSAFVG----------------VTAPATEPTAAAATVSFLHLS     277
Mojavensis       -GEPDHIPSRQRRLPAFTSASVG----------------VTAAATEPTTASATASLLQLS     297
Virilism         -GESDHIPSRQHRLPASTSASVG----------------VTAAATEPATAAATASLLQLS     281
Bipectinate      GGVPEYKAEKRWPLHSPGFRTTPPMSKVLQQQDIDQDEDQ-----DQNPDHVTESLQFVS     342
Anannassae       GGVPE----KRWHLPSSGFRTTPSMSKVIQEQDIDQD--Q-----DQSPDVVTESLQFHS     337
Serrata          -GEPVNIPRLRRHWQSSGFVTPPTLRRTLQDQDRVLENRGKIGQKQEMVQVTTESLLQLL     314
Kikkawei         -GEPVNIPRLRRHWQSSGFVTPPTLRKTLQDQERELENGGKIGQKQELVQVTTESLLQLL     308
Fichsuphila      -GEPENIPRHRRHWQSTGFVTLPALRRTLEEQEQQVRQPAEEEEEQDEGSRVTESLLQLL     320
Eugracilis       -GEAENIPRHRRQWQSRGFVTLPTLRKTLEEQETDQQQ------DEEQGSSATESLLQLL     310
Rhopaloa         -GEPDNIPRHRRHWQSSGFVTLPNLRKTLEEQDQDQEV-----EEGEQGSRVTESLLQLL     296
Elegans          -GEPENIPRHRRHWQSSGFVTLPNLRKTLEEQDQDEAV-----E--EQGSRVTESLLQLL     304
Takahashi        -GEPDNIPRQRRHWQSSGFVTLPTLRKTLEEQEQDQDVQ---EGEGELGSRVTESLLQLL     316
Suzuki           -GEPENIPRHRRHRQSSGFVTLPTLRKTLEEQDQEG-------DEEEPGSRVTESLLQLL     314
Biarmipes        -GEPENIPRHRRHWHSSGFVTLPTLRKTLEEQDQEG-------EAGEPGSRVTESLLQLL     312
Erecta           -GEPENIPRRRRHWQSSGLVTLPALRKTLEEQDQDQDQE---GGEEEEGSGVTESLLQLL     317
Melanogaster     -GEPENIPRHRRHWQSSGFVTLPTLRKTLEEQDQ----K---VADAAQRSGVTESLLQLW     308
Simulans         -GEPENIPRHRRHWQSSGFVTLPALRKTLEEQDP----E---GADAGQGSGVTESLLQLW     307
Mauritania       -GEPENIPRHRRHWQSSGFVTLPTLRKTLEEQDP----E---GADAGQGSGVTQSLLQLW     307
Sechellia        -GEPENITRHRRHRQSSGFVTLPTLRKTLEEQDP----E---GADAGQGSGVTESLLQLW     307
                  *        :    :                                   .* *:

Grimshawi        RARRNEENYDNDANFKNNYSNNNNNNNSNSNNNNNNNNNSNKNNSNSSAFEFNVTEYCQNM    337
Mojavensis       RARRSEDNYNYNNDANNNNNN--------NNNDSGNDNNNNNNNNNSAFAFNITEYCQNM    349
```

```
Virilism          RARRSEDNYDSEASSN----------------------NNYNYNNNSAFAFNITEYCQNM        319
Bipectinate       RRLKSL----------------------------------SEVNTTEAFAFNITEYCQNM        368
Anannassae        RRLRST----------------------------------SDVNTTEAFAFNVTEYCQNV        363
Serrata           RRKRSTD-----------------------------------HNITDAFAFNVTEYCQNM        339
Kikkawei          RSKRSTDH----------------------------------HNITDAFAFNVTEYCQNM        334
Fichsuphila       RSKRSADNNNDY--------------------------NNDNSNNTDAFAFNVTEYCQNM        354
Eugracilis        LRSKRSAENNNN-------------------------NNNNNINNTDAFAFNVTEYCQNV        345
Rhopaloa          RRKRNNN-----------------------------------INNTDAFAFNVTEYCQNV        321
Elegans           RRKRNNNDNNNNNNNNNNN--NNNNNNNNNNNNNNNNNNNNNNNNTDAFAFNVTEYCQNV        362
Takahashi         RRKRSAQN--------------------------------SNSNNTDAFAFNVTEYCQNV        344
Suzuki            RRKRSAGNN-------------------------------SN-NNTDAFAFNVTEYCQNV        342
Biarmipes         RRKRSTGNN-------------------------------NNINNTDAFAFNVTEYCQNV        341
Erecta            RRKRRAEN--------------------------------HNINNTDAFAFNVTEYCQNV        345
Melanogaster      RRKRSAEN--------------------------------HNINNTDAFAFNVTEYCQNV        336
Simulans          RRKRSAEI--------------------------------HNINNTDAFAFNVTEYCQNV        335
Mauritania        RRKRSAEI--------------------------------HNINNTDAFAFNVTEYCQNV        335
Sechellia         RRKRSAEI--------------------------------HNINKTDAFAFNVTEYCQNV        335
                     :                                       . ..** **:******:

Grimshawi         TVYANDLSSLGQNALYINIWSVYSLIVFVLLPLLVLATFNCFLILLVHRSKSLRGDLTNA        397
Mojavensis        TFYTNGLSSLGQNALYFNIWSVYTLIVFVLLPLFVLATFNCFLILLVHRSKSLRGDLTNA        409
Virilism          TVYTNGLSSLGQNALYFNIWSVYTLIVFVVLPLFVLATFNCFLILLVHRSKSLRGDLTNA        379
Bipectinate       TFFLHTSSELGMNDLYASVWNLFTLLVFVLLPLLLLATFNSFLILLVHRSKNLRGDLTNA        428
Anannassae        TYFFHASSELGMNYLYASVWNMFTLLVFVLSPLLLLATFNSFLILLVHRSKNLRGDLTNA        423
Serrata           TLYNHGLSELGMNELYGNIWNVFTLLVFVVLPLLLLVTFNSFLILLVHRSKSLRGDLTNA        399
Kikkawei          TLYNHGPSELGMNELYANIWNVFTLLVFVVLPLLLLVTFNSFLILLVHRSKSLRGDLTNA        394
Fichsuphila       TVFGHGYSELGMDELYSNLWSMVTLLIFVVLPLLLLATFNSFLILLVHRSKNLRGDLTNA        414
Eugracilis        TIYNHGLSELGQDELYSNLWNMFTLLVFVVLPLLLLATFNSFLILLVHRSKNLRGDLTNA        405
Rhopaloa          TIYNHGTSALANDELYSNLWNLFTLLVFVVFPLLLLATFNSFLILLVHRSKSLRGDLTNA        381
Elegans           TIYNHGTSELGKDELYSNLWNMFTLLVFVVFPLLLLATFNSFLILLVHRSKSLRGDLTNA        422
Takahashi         TFYNHGLSELGKDELYSNLWNMFTLLVFVVFPLLLLATFNTFLILLVHRSKNLRGDLTNA        404
Suzuki            TFYNHGPSELGMDELYSNLWNMFTLLVFVVFPLLLLATFNTFLILLVHRSKNLRGDLTNA        402
Biarmipes         TFYNHGPSELGMDELYSNLWNLFTLLVFVVLPLLLLATFNTFLILLVHRSKNLRGDLTNA        401
Erecta            TFYNHGLSELGYDELYSYLWNLFTLLVFVVFPLLLLATFNSFLILLVHRSKNLRGDLTNA        405
Melanogaster      TYYNHGLSELGYDELYSYLWNLFTLLVFVVFPLLLLATFNSILILLVHRSKNLRGDLTNA        396
Simulans          TYYNHGLSELGYDELYSYLWNLFTLLVFVVFPLLLLATFNSILILLVHRSKNLRGDLTNA        395
Mauritania        TYYNHGLSELGYDELYSYLWNLFTLLVFVVFPLLLLATFNSILILLVHRSKNLRGDLTNA        395
Sechellia         TYYNHGLSELGYDELYSYLWNLFTLLVFVVFPLLLLATFNSILILLVHRSKNLRGDLTNA        395
                  * : :  * *. : **  :*.: :*::**: **::*.*** :*********.********

Grimshawi         SSMRRTKRKSTSGLTGSVSQENRVTITLIAVVLLFIVCQLPWAIYLILVQYMDIELNIQR        457
Mojavensis        SSIRRTKRKSNSGLTGSVSQENRVTITLIAVVLLFIVCQLPWAIYLILVQYVDIEMNIQR        469
Virilism          SSIRRTKRKSNSGITGSVSQENRVTITLIAVVLLFIVCQLPWAIYLILVQYVEIEMNIQR        439
Bipectinate       NSIRRTKRKSNSGITGTVSQENRVTITLIAVVLLFIVCQLPWAIYLILGQYMEIAPSTQV        488
Anannassae        NSIRRTKRKSNTGITGTVSQENRVTITLIAVVLLFIVCQLPWAIYLILEQYMEIHGSTQV        483
Serrata           SSIRRTKRKSSSGIKGSVSQENRVTITLIAVVLMFIVCQLPWAIYLILNTYMDIQVGTQL        459
Kikkawei          SSIRRTKRKSSSGIKGSVSQENRVTITLIAVVLMFIVCQLPWAIYLILSTYMEIQVGTQL        454
Fichsuphila       SSIRRTKRKSNSGIKSSVSQEHRVTITLIAVVLMFIVCQLPWAIYLVVEQYMTIQVSTQV        474
Eugracilis        SSIRRTKRKSNTGIKGSVSQENRVTITLIAVVLMFIVCQLPWAIYLVVNQYMDIQVGTQV        465
Rhopaloa          SSIRRTKRKSNSGLKGSVSQENRVTITLIAVVLMFIVCQLPWAIYLVLSQYMNFQLGTQV        441
Elegans           SSIRRTKRKSNSGIKGSVSQENRVTITLIAVVLMFIFCQLPWAIYLILSQYMDFQIGTQV        482
Takahashi         SSIRRTKRKSNSGIKGSVSQENRVTITLIAVVLMFIVCQLPWAIYLVVNQYMDIQVGTQV        464
Suzuki            SSIRRTKRKSSSGIKGSVSQENRVTITLIAVVLMFIVCQLPWAIYLVVNQYMDIQVGTQV        462
Biarmipes         SSIRRTKRKSSSGIKGSVSQENRVTITLIAVVLMFIVCQLPWAIYLVVNQYMDIQFGTQV        461
Erecta            SSIRRTKRKSNSGLKGSVSQENRVTITLIAVVLMFIVCQLPWAIYLIVNQYKEIQVGTQV        465
Melanogaster      SSIRRTKRKSNSGLKGSVSQENRVTITLIAVVLMFIVCQLPWAIYLIVNQYMEIQIGTQV        456
Simulans          SSIRRTKRKSNSGLKGSVSQENRVTITLIAVVLMFIVCQLPWAIYLIVNQYMEIQVGTQV        455
Mauritania        SSIRRTKRKSNSGLKGSVSQENRVTITLIAVVLMFIVCQLPWAIYLIVNQYMEIQVGTQV        455
Sechellia         SSIRRTKRKSNSGLKGSVSQENRVTITLIAVVLMFIVCQLPWAIYLIVNQYMEIQVGTQV        455
                  .*:*******.:*:..:****:***********:**.*********::  *  :  . *

Grimshawi         IAGNVCNLLVAINAAANFFLYCVLSDKYRKTVRELITGYRYHHRHARNNFSLYMPHTTTT        517
Mojavensis        IAGNVCNLLVAINAAANFFLYCVLSDKYRKTVRELVTGYRYRHRHARNNISLYVPHTSTT        529
Virilism          IAGNVCNLLVAINAAANFFLYCVLSDKYRKTVRELITGYRYRHRHARNNISLYAPHTTTT        499
Bipectinate       VAGNVCNLLAAFNAASNFFLYCVLSDKYRKTVRELITGYRYRRRHMRNNTSLYVPHTTTT        548
Anannassae        VAGNICNLLAAFNAASNFFLYCVLSDKYRKTVRELITGYRYRRRHMRNNTSLYVPHTTTT        543
Serrata           VAGNVFNLLAALNAASNFFLYCVLSDKYRKTVRELITGYRYRRRHARNNTSLYVPHTTTT        519
Kikkawei          VAGNVFNLLAALNAASNFFLYCVLSDKYRKTVRELITGYRYRRRHARNNTSLYVPHTTTT        514
Fichsuphila       VAGNVCNLLASLHAASNFFLYCVLSDKYRKTVRELITGYRYRHRHNRNNTSLYVPHTTTT        534
Eugracilis        VAGNVCNLLASLHAASNFFLYCVLSDKYRKTVRELITGYRYRRRHARNNTSLYVPHTTTT        525
Rhopaloa          VAGNVCNLLASLHAASNFFLYCVLSDKYRKTVRELITGYRYRRRHIRNNTSLYVPQTTTT        501
```

Elegans VAGNVCNLLASLHAASNFFLYCVLSDKYRKTVRELITGYRYRRRHARNNTSVYVPH**TTTT** 542
Takahashi VAGNVCNLLASLHAASNFFLYCVLSDKYRKTVRELITGYRYRRRHARNNTSLYVPH**TTTT** 524
Suzuki VAGNVCNLLASLHAASNFFLYCVLSDKYRKTVRELITGYRYKRRHARNNTSLYVPQ**TTTT** 522
Biarmipes VAGNVCNLLASLHAASNFFLYCVLSDKYRKTVRELITGYRYKRRHARNNTSLYVPQ**TTTT** 521
Erecta VAGNVCNLLASLHAASNFFLYCVLSDKYRKTVRELITGYRYRRRHARNNTSLYVPH**TTTT** 525
Melanogaster VAGNVCNLLASLHAASNFFLYCVLSDKYRKTVRELITGYRYRRRHARNNTSLYVPH**TTTT** 516
Simulans VAGNVCNLLASLHAASNFFLYCVLSDKYRKTVRELITGYRYRRRHARNNTSLYVPH**TTTT** 515
Mauritania VAGNVCNLLASLHAASNFFLYCVLSDKYRKTVRELITGYRYRRRHARNNTSLYVPH**TTTT** 515
Sechellia VAGNVCNLLASLHAASNFFLYCVLSDKYRKTVRELITGYRYRRRHARNNTSLYVPH**TTTT** 515
:***: ***.:::**:*******************:*****::** *** *:* *:*:**

Grimshawi TNGDSASASGGASGYGSYRNANSRRCR------PTGRLIA 551
Mojavensis LNGDGASSVGSGGGYNAYGGASSRRCRGKAAV--ARRLIA 567
Virilism LNGDGAGGGGASGYG**SSYSGAS**SRRCRAKSAV--ARRLIA 537
Bipectinate **LT**HINGDR---GGGGSYYGGAGNRRSRNKSA--ALGRLIA 583
Anannassae **LT**HINGDR---GGGGSYYGGAGNRRSQNKSA--AMGRLIA 578
Serrata **LS**HINGHD--------NYGGTGSRRSRNNNKSMATGRLIA 551
Kikkawei **LS**HINGHD--H--GGSHYGGAGSRRSRNNNKSMATGRLIA 550
Fichsuphila **LT**QINGDH—YGGGGGHYGGAGSRRTRN------TTGRLIA 567
Eugracilis **LT**QINGDH--YG---SNYGGAGSRRNR------NINRLIA 554
Rhopaloa **LT**QINGDH--YG---SNYGGAGSRRSR------NTSRLIT 530
Elegans **LT**QINGDH--YG---SNYGGAGSRRSR------NTSRLIT 571
Takahashi **LT**QINGDH--YG---SNYGGAGSRRNR------NTNRLIA 553
Suzuki **LT**QINGDH--YG---SNYGGAGSRRNR------NTNRLIA 551
Biarmipes **LT**QINGDH--YG---SNYGGAGSRRNR------NTNRLIA 550
Erecta **LT**QINGDH--YG---SNYGGAGSRRNR------NTARLIA 554
Melanogaster **LT**QINGDH--YG---SNYGGAGSRRNR------NTGRLIA 545
Simulans **LT**QINGDH--YG---SNYGGNGSRRNR------NTGRLIA 544
Mauritania **LT**QINGDH--YG---SNYGGNGSRRNR------NTGRLIA 544
Sechellia **LT**QINGDH--YG---SNYGGNGSRRNR------NTGRLIA 544

Melanogaster **NP_001014723.1**

1 mtmsststat atstatatld eanatvgemf sdadmaevrh vvqrilvpcv fvigllgnsv
61 siyvltrkrm rcttniylta laitdiaylt cqlilslqhy dypkyhfkly wqlygyfvwl
121 cdsfgyisiy iavcftierf iairyplkrq tfcteslakk viaavaifcl lstlstafeh
181 titigtrqid dayqpcnqtv anispmpppp vavtpplatp plptpatiwq spdsamestt
241 sgssnqlvdw gsgsgdgepe niprhrrhwq ssgfvtlptl rktleeqdqk vadaaqrsgv
301 tesllqlwrr krsaenhnin ntdafafnvt eycqnvtyyn hglselgyde lysylwnlft
361 llvfvvfpll llatfnsili llvhrsknlr gdltnassir rtkrksnsgl kgsvsqenrv
421 titliavvlm fivcqlpwai ylivnqymei qigtqvvagn vcnllaslha asnfflycvl
481 sdkyrktvre litgyryrrr harnntslyv phttttltqi ngdhygsnyg gagsrrnrnt
541 grlia

Simulans **XP_016038098.1**

1 mtmsststat statatlaea natvgemfsd admaevrhvv qrilvpcvfv igllgnsvsi
61 yvltrkrmrc ttniyltala itdiayltcq lilslqhydy pkyhfklywq lygyfvwlcd
121 sfgyisiyia vcftierfia iryplkrqtf cteslakkvi aavaifclls tlstafehti
181 tigtrqidda yqpcnqtlan isptpppppv aatpplatpp lptpatvwqs pdlamestta
241 gssnllvdwg sgsgdgepen iprhrrhwqs sgfvtlpalr ktleeqdpeg adagqgsgvt
301 esllqlwrrk rsaeihninn tdafafnvte ycqnvtyynh glselgydel ysylwnlftl
361 lvfvvfplll latfnsilil lvhrsknlrg dltnassirr tkrksnsglk gsvsqenrvt
421 itliavvlmf ivcqlpwaiy livnqymeiq vgtqvvagnv cnllaslhaa snfflycvls
481 dkyrktvrel itgyryrrrh arnntslyvp httttltqin gdhygsnygg ngsrrnrntg
541 rlia

Suzuki **XP_016939418.1**

1 mtmsststat lteaaaeaaa aaeaatlgna tvgemfsdad maevrhvvqr ilvpcvfvig
61 llgnsvsiyv ltrkrmrctt niyltalait diayltcqli lslqhydypk yhlkiywqly
121 gyfvwlcdsf gyisiyiavc ftierfiair yplkrqtfct eslakkviaa vavfcllstl
181 stafehtitv gskliddayq pcnqtvanis ptpppsvaat pplvtpplst patiwqsqdf
241 ttesttagss nllvdwgsgs gdgepenipr hrrhrqssgf vtlptlrktl eeqdqegdee
301 epgsrvtesl lqllrrkrsa gnnsnnntda fafnvteycq nvtfynhgps elgmdelysn
361 lwnmftllvf vvfpllllat fntflillvh rsknlrgdlt nassirrtkr ksssgikgsv
421 sqenrvtitl iavvlmfivc qlpwaiylvv nqymdiqvgt qvvagnvcnl laslhaasnf
481 flycvlsdky rktvrelitg yrykrrharn ntslyvpqtt ttltqingdh ygsnyggags

```
541 rrnrntnrli a
```

Mauritania **XP_033170840.1**

```
  1 mtmsststat statatlaea natvgemfsd admaevrhvv qrilvpcvfv igllgnsvsi
 61 yvltrkrmrc ttniyltala itdiayltcq lilslqhydy pkyhfklywq lygyfvwlcd
121 sfgyisiyia vcftierfia iryplkrqtf cteslakkvi aavaifclls tlstafehti
181 tigtrqidda yqpcnqtlan isptpppppv aatpplatpp lptpatawqs pdfamestta
241 gssnllvdwg sgsgdgepen iprhrrhwqs sgfvtlptlr ktleeqdpeg adagqgsgvt
301 qsllqlwrrk rsaeihninn tdafafnvte ycqnvtyynh glselgydel ysylwnlftl
361 lvfvvfplll latfnsilil lvhrsknlrg dltnassirr tkrksnsglk gsvsqenrvt
421 itliavvlmf ivcqlpwaiy livnqymeiq vgtqvvagnv cnllaslhaa snfflycvls
481 dkyrktvrel itgyryrrrh arnntslyvp htttltqin gdhygsnygg ngsrrnrntg
541 rlia
```

Sechellia **XP_002036980.1**

```
  1 mtmsststat statatlaea natvgemfsd admaevrhvv qrilvpcvfv igllgnsvsi
 61 yvltrkrmrc ttniyltala itdiayltcq lilslqhydy pkyhfklywq lygyfvwlcd
121 sfgyisiyia vcftierfia iryplkrqtf cteslakkvi aavaifclls tlstafehti
181 tigtrqidda yqpcnqtlan isptppppplg eatpplatpp lptpatvwqs pdfamestta
241 gssnllvdwg sgsgdgepen itrhrrhrqs sgfvtlptlr ktleeqdpeg adagqgsgvt
301 esllqlwrrk rsaeihnink tdafafnvte ycqnvtyynh glselgydel ysylwnlftl
361 lvfvvfplll latfnsilil lvhrsknlrg dltnassirr tkrksnsglk gsvsqenrvt
421 itliavvlmf ivcqlpwaiy livnqymeiq vgtqvvagnv cnllaslhaa snfflycvls
481 dkyrktvrel itgyryrrrh arnntslyvp htttltqin gdhygsnygg ngsrrnrntg
541 rlia
```

Serrata **KAH8385187.1**

```
  1 mtmsststssqs sdgnetvgem fseadtaevr hvvqrilvpc vfvigllgns vsiyvltrkr
 61 mrcttniylt alaitdiayl tfqlilslqh ydyvkyhcei ywqlygifvw lcdssgyisi
121 yiavcftier fiairyplkr qtfcteslak kviaavslfc llstlstafe htikvdwkli
181 dgayqpcnqt lanvsptpps tsltpalstp plvtpatiwq eqdqereslq dfftesssv
241 kssrllldfg sgsgdgepvn iprlrrhwqs sgfvtpptlr rtlqdqdrvl enrgkigqkq
301 emvqvttesl lqllrrkrst dhnitdafaf nvteycqnmt lynhglselg mnelygniwn
361 vftllvfvvl pllllvtfns flillvhrsk slrgdltnas sirrtkrkss sgikgsvsqe
421 nrvtitliav vlmfivcqlp waiylilnty mdiqvgtqlv agnvfnllaa lnaasnffly
481 cvlsdkyrkt vrelitgyry rrrharnnts lyvphttttl shinghdnyg gtgsrrsrnn
541 nksmatgrli a
```

Erecta **XP_001976911.1**

```
  1 mtmsststat atatstatat laeanatvge mfsdadmaev rhvvqrilvp cvfvigllgn
 61 svsiyvltrk rmrcttniyl talaitdiay ltcqlilslq hydytkyhlk lywqlygyfv
121 wlcdsfgyis iyiavcftie rfiairyplk rqtfctesla kkviaavaif cllstlstaf
181 ehtitmgsrq iddayqpcnq tvanisptpp ppsaaatppl atpplptpat iwqsqdyste
241 sttagstsll vdwgsgsgdg dgepeniprr rrhwqssglv tlpalrktle eqdqdqdqeg
301 geeeegsgvt esllqllrrk rraenhninn tdafafnvte ycqnvtfynh glselgydel
361 ysylwnlftl lvfvvfplll latfnsflil lvhrsknlrg dltnassirr tkrksnsglk
421 gsvsqenrvt itliavvlmf ivcqlpwaiy livnqykeiq vgtqvvagnv cnllaslhaa
481 snfflycvls dkyrktvrel itgyryrrrh arnntslyvp htttltqin gdhygsnygg
541 agsrrnrnta rlia
```

Takahashi **XP_017013816.2**

```
  1 mtmsttstat lteaaaaaaa eaatlgnatv gemfsdadma evrhvvqril vpcvfvigll
 61 gnsvsiyvlt rkrmrcttni yltalaitdi ayltcqlils lqhydypkyn fkiywqlygy
121 fvwlcdsfgy isiyiavcft ierfiairyp lkrqtfctes lakkviaava ifcllstlst
181 afehtitvgw kqiddayqpc nqtvanispt pppsaaatpp lvtpplptpa tiwqsqdltt
241 esttagssnl ladwgsgsgd gepdniprqr rhwqssgfvt lptlrktlee qeqdqdvqeg
301 egelgsrvte sllqllrrkr saqnsnsnnt dafafnvtey cqnvtfynhg lselgkdely
361 snlwnmftll vfvvfpllll atfntflill vhrsknlrgd ltnassirrt krksnsgikg
421 svsqenrvti tliavvlmfi vcqlpwaiyl vvnqymdiqv gtqvvagnvc nllaslhaas
481 nfflycvlsd kyrktvreli tgyryrrrha rnntslyvph ttttltqing dhygsnygga
541 gsrrnrntnr lia
```

Biarmipes **XP_016946298.1**

```
1 mtmsstvtat lteaaaaaea eaatlgnata gemfsdadma evrhvvqril vpcvfvigll
      61 gnsvsiyvlt rkrmrcttni yltalaitdi ayltcqlils lqhydypkyh lkiywqlygy
     121 fvwlcdsfgy isiyiavcft ierfiairyp lkrqtfctes lakkviaava vfcllstlst
     181 afehtitvgs rlideayqpc nqsvanispt ppafvaatpp lvtpplstpa tiwksqdstt
     241 esttagssnl lvdwgsgsgd gepeniprhr rhwhssgfvt lptlrktlee qdqegeagep
     301 gsrvtesllq llrrkrstgn nnninntdaf afnvteycqn vtfynhgpse lgmdelysnl
     361 wnlftllvfv vlplllatf ntflillvhr sknlrgdltn assirrtkrk sssgikgsvs
     421 qenrvtitli avvlmfivcq lpwaiylvvn qymdiqfgtq vvagnvcnll aslhaasnff
     481 lycvlsdkyr ktvrelitgy rykrrharnn tslyvpqttt tltqingdhy gsnyggagsr
     541 rnrntnrlia
```

Eugracilis **XP_017074826.1**

```
1 mtmsttstlt atatatatet lgnatvgemf sdadmaevrh vvqrilvpcv fvigllgnsv
      61 siyvltrkrm rcttniylta laitdiaylt cqlilslqhy dytkfhveiy wqlygyfvwl
     121 cdsfgyisiy iavcftierf iairyplkrq tfcteslakk viaavaffcl lstlstafeh
     181 titvgsklid dayqpcnqtl anyspmptss vaatpplatp plptpatiwq stelntestt
     241 tagksnvlid wgsgsgdgea eniprhrrqw qsrgfvtlpt lrktleeqet dqqqdeeqgs
     301 satesllqll lrskrsaenn nnnnnninn tdafafnvte ycqnvtiynh glselgqdel
     361 ysnlwnmftl lvfvvlplll latfnsflil lvhrsknlrg dltnassirr tkrksntgik
     421 gsvsqenrvt itliavvlmf ivcqlpwaiy lvvnqymdiq vgtqvvagnv cnllaslhaa
     481 snfflycvls dkyrktvrel itgyryrrrh arnntslyvp httttltqin gdhygsnygg
     541 agsrrnrnin rlia
```

Rhopaloa **XP_044317201.1**

```
       1 mevaatlgna tvgemfsdad maevrhvvqr ilvpcvfvig llgnsvsiyv ltrkrmrctt
      61 niyltalait diayltcqli lslqhydypk yhfeiyyqlf gyfvwlcdsf gyisiyiavc
     121 ftierfiair yplkrqtfct eslakkviaa valfcllstl stafehtitv gykliddayq
     181 pcnlteanis ptppappfav atpplatppr ptpatisqdf ttesttagss nllvdwgsgs
     241 gdgepdnipr hrrhwqssgf vtlpnlrktl eeqdqdqeve egeqgsrvte sllqllrrkr
     301 nnninntdaf afnvteycqn vtiynhgtsa landelysnl wnlftllvfv vfplllatf
     361 nsflillvhr skslrgdltn assirrtkrk snsglkgsvs qenrvtitli avvlmfivcq
     421 lpwaiylvls qymnfqlgtq vvagnvcnll aslhaasnff lycvlsdkyr ktvrelitgy
     481 ryrrrhirnn tslyvpqttt tltqingdhy gsnyggagsr rsrntsrlit
```

Fichsuphila **XP_017038922.1**

```
           1 msstststtt atltateaat vgnatvgemf sdaemaevrh vvqrilvpcv fvigllgnsv
      61 siyvltrkrm rcttniylta laitdiaylt cqlilsiqhy dyakyhlqif welygyfvwl
     121 cdsfgyisiy iavcftierf iairyplkrq tfcteslakk viaavavfcl lstlstafeh
     181 titvgwklid gayqpcnqte anfspmppsf algtpplatp platpplatp ptvwqdfste
     241 sptaiagsss qlidwgsgsg dgepeniprh rrhwqstgfv tlpalrrtle eqeqqvrqpa
     301 eeeeeqdegs rvtesllqll rskrsadnnn dynndnsnnt dafafnvtey cqnmtvfghg
     361 yselgmdely snlwsmvtll ifvvlpllll atfnsflill vhrsknlrgd ltnassirrt
     421 krksnsgiks svsqehrvti tliavvlmfi vcqlpwaiyl vveqymtiqv stqvvagnvc
     481 nllaslhaas nfflycvlsd kyrktvreli tgyryrhrhn rnntslyvph ttttltqing
     541 dhyggggghy ggagsrrtrn ttgrlia
```

Elegans **XP_017114263.2**

```
1 msststst ea apeaatlgna tvgemfsdad maevrhvvqr ilvpcvfvig llgnsvsiyv
      61 ltrkrmrctt niyltalait diayltcqli lslqhydypk yhleiyyqly gyfvwlcdsf
     121 gyisiyiavc ftierfiair yplkrqtfct eslakkviaa vavfcllstl stafehtitv
     181 gwkliddayq pcnqtvanfs ptppvpsfav atpplatppl ptpatisqhf stesttagss
     241 nllfdwgsgs gdgepenipr hrrhwqssgf vtlpnlrktl eeqdqdeave eqgsrvtesl
     301 lqllrrkrnn ndnnnnnnnn nnnnnnnnnn nnnnnnnnnn nnnnnnntda fafnvteycq
     361 nvtiynhgts elgkdelysn lwnmftllvf vvfplllat fnsflillvh rskslrgdlt
     421 nassirrtkr ksnsgikgsv sqenrvtitl iavvlmfifc qlpwaiylil sqymdfqigt
     481 qvvagnvcnl laslhaasnf flycvlsdky rktvrelitg yryrrrharn ntsvyvphtt
     541 ttltqingdh ygsnyggags rrsrntsrli t
```

Kikkawei **XP_017024118.1**

```
  1 msstssqssd gnetvgemfs eaetaevrhv vqrilvpcvf vigllgnsvs iyvltrkrmr
      61 cttniyltal aitdiayltf qlilslqhyd yvkyhceiyw qlygifvwlc dssgyisiyi
     121 avcftierfi airyplkrqt fcteslakkv iaavslfcll stlstafeht ikvdwklidg
     181 ayqncnqtla nvspmppsts vtpalstppm vtpptiwkek eleqnfftqe sstakssrll
```

```
241 fdfgsgsgdg epvniprlrr hwqssgfvtp ptlrktlqdq erelenggki gqkqelvqvt
301 tesllqllrs krstdhhnit dafafnvtey cqnmtlynhg pselgmnely aniwnvftll
361 vfvvlplllll vtfnsflill vhrskslrgd ltnassirrt krksssgikg svsqenrvti
421 tliavvlmfi vcqlpwaiyl ilstymeiqv gtqlvagnvf nllaalnaas nfflycvlsd
481 kyrktvreli tgyryrrrha rnntslyvph ttttlshing hdhggshygg agsrrsrnnn
541 ksmatgrlia
```

Bipectinate XP_043066026.1

```
1 mtmfslltat ltatatatat atvtatatst eseaeaatln eteaagtemf sdadvvevrr
61 vvqrilvpcv fvigllgnsv siyvltrkrm rcttniylta laitdiaylt fqlilsfqhy
121 dyvkyhceiy wqfygyfvwl cdscayisiy iavcftierf iairyplkrq tfcteslakk
181 viaavalfcl lstlstafeh sfarkfrlid ddyrpcnqtl anvspqplal tpslsmqaps
241 pldsttplpt amlptpvtph mqdltrkpek plgvlvsdvf sgggsgdggv peykaekrwp
301 lhspgfrttp pmskvlqqqd idqdedqdqn pdhvteslqf vsrrlkslse vntteafafn
361 iteycqnmtf flhtsselgm ndlyasvwnl ftllvfvllp llllatfnsf lillvhrskn
421 lrgdltnans irrtkrksns gitgtvsqen rvtitliavv llfivcqlpw aiylilgqym
481 eiapstqvva gnvcnllaaf naasnfflyc vlsdkyrktv relitgyryr rrhmrnntsl
541 yvphttttlt hingdrgggg syyggagnrr srnksaalgr lia
```

Anannassae XP_032309300.1

```
1 mtmfslltat ltatatatat atatvtpaea eteaeaagse mfsdadvaev rrvvqrilvp
61 cvfvigllgn svsiyvltrk rmrcttniyl salaitdiay ltfqlilslq hydyfkyhse
121 iywqlygyvv wlcdssayis iyiavcftie rfiairyplk rqtfctesla kkviaavalf
181 cllstlstaf ehsyargyrl iddayrpcnq tlanvspqpq plipslhmqa pspsasttpl
241 ptvmlptpvt phwqdsrdsq dlagrstekp lgglflaadv fsgggsgdgg vpekrwhlps
301 sgfrttpsms kviqeqdidq dqdqspdvvt eslqfhsrrl rstsdvntte afafnvteyc
361 qnvtyffhas selgmnylya svwnmftllv fvlspllla tfnsflillv hrsknlrgdl
421 tnansirrtk rksntgitgt vsqenrvtit liavvllfiv cqlpwaiyli leqymeihgs
481 tqvvagnicn llaafnaasn fflycvlsdk yrktvrelit gyryrrrhmr nntslyvpht
541 tttlthingd rggggsyygg agnrrsqnks aamgrlia
```

Mojavensis XP_015016788.1

```
1 mstlaaatst stampvdaia aakmfsdadv aevrhvvqri lvpcvfvigl lgnsvsiyvl
61 trkrmrcttn iyltalaitd iayltfvlil slkhyeyiky hcelywllyg fvvwlcdaca
121 yisiyiavcf tierfiairy plkrqtfcte slakkviaav alfcllttls tafehtydin
181 wklidgayrp cnltlanvsp mptptptstp mptpaqtmtt awhieepagn patsrylepf
241 dlssdsgsgs gsgagepdhi psrqrrlpaf tsasvgvtaa atepttasat asllqlsrar
301 rsednynynn dannnnnnnn ndsgndnnnn nnnnnsafaf niteycqnmt fytnglsslg
361 qnalyfniws vytlivfvll plfvlatfnc flillvhrsk slrgdltnas sirrtkrksn
421 sgltgsvsqe nrvtitliav vllfivcqlp waiylilvqy vdiemniqri agnvcnllva
481 inaaanffly cvlsdkyrkt vrelvtgyry rhrharnnis lyvphtsttl ngdgassvgs
541 gggynaygga ssrrcrgkaa varrlia
```

Virilism XP_015026487.1

```
1 mttlaatsts tampvdgigt akmfsdadva evrhvvqril vpcvfvigll gnsvsiyvlt
61 rkrmrcttni ylsalaitdi ayltfvlils lkhyeyikyh celywrlygf imwlcdacay
121 isiyiavcft ierfiairyp lkrqtfctes lakkviaava lfcllttlst afehtydinw
181 klidgayrpc nltlanvspt paqptlataw hvdehannaa tprylepfdl ssgsgsgage
241 sdhipsrqhr lpastsasvg vtaaatepat aaatasllql srarrsedny dseassnnny
301 nynnnsafaf niteycqnmt vytnglsslg qnalyfniws vytlivfvvl plfvlatfnc
361 flillvhrsk slrgdltnas sirrtkrksn sgitgsvsqe nrvtitliav vllfivcqlp
421 waiylilvqy veiemniqri agnvcnllva inaaanffly cvlsdkyrkt vrelitgyry
481 rhrharnnis lyaphttttl ngdgaggga sgygssysga ssrrcraksa varrlia
```

Grimshawi XP_043072003.1

```
1 mttltatsta mpgdgnaaak mfsdadvaev rhvvqrilvp cvfvigllgn svsiyvltrk
61 rmrcttniyl salaitdiay ltfvlilslk hyeyikyhce lywrlygfvm wlcdacayis
121 iyiavcftie rfiairyplk rqtfctesla kkviaavalf cllttlstaf ehtydidwkl
181 idgayrpcnq tlanvsptpa hqtpgtawhv nenasmsntt psnveppldl gsgagesdhi
241 psrqrrlpas tsafvgvtap ateptaaaat vsflhlsrar rneenydnda nfknnysnnn
301 nnnnsnsnnn nnnnnsnknn snssafefnv teycqnmtvy andlsslgqn alyiniwsvy
361 slivfvllpl lvlatfncfl illvhrsksl rgdltnassm rrtkrkstsg ltgsvsqenr
421 vtitliavvl lfivcqlpwa iylilvqymd ielniqriag nvcnllvain aaanfflycv
481 lsdkyrktvr elitgyryhh rharnnfsly mphtttttng dsasasggas gygsyrnans
541 rrcrptgrli a
```
