## Supplementary material for "The incidence of candidate binding sites for β-arrestin in Drosophila neuropeptide GPCRs": S10 Text

**_S10. Text._ Multi-species analysis of Tk R 86C**
**Supporting Figure 12**

CLUSTAL Line-ups; Genbank Reference IDs below
5th, 6th and 7th Predicted TM domain in YELLOW
BBS sequences in RED

```
Grimshawi         ---------------------MPEIVDTELLVNCTILAVLRFELNSIVNITLLNSLNRTE        39
Bipectinate       ---------------------MSETVDTELLVNCTILAVRRFELNSIVNTTLLSSLNRTE        39
Anannassae        ---------------------MSETVDTELLVNCTILAVRRFELNSIVNTTLLSSLNRSE        39
Serrata           ---------------------MSEIVDTELLVNCTILAVRRFELNSIVNTTLLNSLNRTE        39
Kikkawei          ---------------------MSEIVDTELLVNCTILAVRRFELNSIVNTTLLNSLNRTE        39
Rhopaloa          ---------------------MSEIVDTELLVNCTILAVRRFELNSIVNTTLLGSLNRTE        39
Elegans           ---------------------MSEIVDTELLVNCTILAVRRFELNSIVNTTLLGSLNRTE        39
Suzuki            MWHKCGGIITQRSLIPIKGSRMSEIVDTELLVNCTILAVRRFELNSIVNTTLLGSLNRTE        60
Biarmipes         MWRKCGGIITQRSLIPIKGCRMSEIVDTELLVNCTILAVRRFELNSIVNTTLLGSLNRTE        60
Takahashi         ---------------------MSEIVDTELLVNCTILAVRRFELNSIVNTTLLSSLNRTE        39
Eugracilis        ---------------------MSEIVDTELLVNCTILAVRRFELNSIVNTTLLGSLNRTE        39
Fichsuphila       MWHECSGIITQRPRIVIKVWKMSEIVDTELLVNCTILAVRRFELNSIVNTTLLGTLNRTE        60
Erecta            ---------------------MSEIVDTELLVNCTILAVRRFELNSIVNTTLLGSLNRTE        39
Mauritania        ---------------------MSEIVDTELLVNCTILAVRRFELNSIVNTTLLGSLNRTE        39
Melanogaster      ---------------------MSEIVDTELLVNCTILAVRRFELNSIVNTTLLGSLNRTE        39
Simulans          ---------------------MSEIVDTELLVNCTILAVRRFELNSIVNTTLLGSLNRTE        39
Sechellia         ---------------------MSEIVDTELLVNCTILAVRRFELNSIVNTTLLGSLNRTE        39
Mojavensis        ---------------------MSEIVDTELLVNCTILAVRRFELNSIVNTTLLNSLNRTE        39
Virilis          ---------------------MSEIVDTELLVNCTILAVRRFELNTIVNTTLLNTLNRTE         39
                                       * * ************** *****:*** ***.:***:*

Grimshawi         VVSLLTSIIDNRDNLESINEAKDFLTECLFPSPTRPYELPWEQKTIWAIVFGLMMFVAIA        99
Bipectinate       VVGLLSGIIENRDNLENINEAKDFLTECLFPSPTRPYELPWEQKTIWAIIFGLMMFVAIA        99
Anannassae        VVGLLSGIIENRDNLESINEAKDFLTECLFPSPTRPYELPWEQKTIWAIIFGLMMFVAIA        99
Serrata           VVSLLSSIIDNRDNLGSINEAKDFLTECLFPSPTRPYELPWEQKTIWAIIFGLMMFVAIA        99
Kikkawei          VVSLLSSIIDNPDNLGSINEAKDFLTECLFPSPTRPYELPWEQKTIWAIIFGLMMFVAIA        99
Rhopaloa          VVSLLSSIIDNRDNLDSINEAKDFLTECLFPSPTRPYELPWEQKTIWAIIFGLMMFVAIA        99
Elegans           VVSLLSSIIENRDNLQSINEAKDFLTECLFPSPTRPYELPWEQKTIWAIIFGLMMFVAIA        99
Suzuki            VVSLLSSIIDNRDNLESINEAKDFLTECLFPSPTRPYELPWEQKTIWAIIFGLMMFVAIA        120
Biarmipes         VVSLLSSIIDNRDNLESINEAKDFLTECLFPSPTRPYELPWEQKTIWAIIFGLMMFVAIA        120
Takahashi         VVSLLSSIIDNRDNLESINEAKDFLTECLFPSPTRPYELPWEQKTIWAIIFGLMMFVAIA        99
Eugracilis        VVSLLSSIIDNRDNLESINEAKDFLTECLFPSPTRPYELPWEQKTIWAIIFGLMMFVAIA        99
Fichsuphila       VVSLLSSIIENRDNLQSINEAKDFLTECLFPSPTRPYELPWEQKTIWAIIFGLMMFVAIA        120
Erecta            VVNLLSSIIDNRDNLESINEAKDFLTECLFPSPTRPYELPWEQKTIWAIIFGLMMFVAIA        99
Mauritania        VVSLLSSIIDNRDNLESINEAKDFLTECLFPSPTRPYELPWEQKTIWAIIFGLMMFVAIA        99
Melanogaster      VVSLLSSIIDNRDNLESINEAKDFLTECLFPSPTRPYELPWEQKTIWAIIFGLMMFVAIA        99
Simulans          VVSLLSSIIDNRDNLESINEAKDFLTECLFPSPTRPYELPWEQKTIWAIIFGLMMFVAIA        99
Sechellia         VVSLLSSIIDNRDNLESINEAKDFLTECLFPSPTRPYELPWEQKTIWAIIFGLMMFVAIA        99
Mojavensis        VVSLLSGIIENRDNLGSINEAKDFLTECLFPSPTRPYELPWEQKTIWAIVFGLMMFVAIA        99
Virilis          VVGLLSGIIENRDNLDSINEAKDFLTECLFPSPTRPYELPWEQKTIWAIVFGLMMFVAIA         99
                  **.**:.**:* *** .*******************************:**********

Grimshawi         GNGIVLWIVTGHRSMRTVTNYFLLNLSIADLLMSSLNCVFNFIFMVNSDWPFGSIYCTIN        159
Bipectinate       GNGIVLWIVTGHRSMRTVTNYFLLNLSIADLLMSSLNCVFNFIFMVNSDWPFGSIYCTIN        159
Anannassae        GNGIVLWIVTGHRSMRTVTNYFLLNLSIADLLMSSLNCVFNFIFMVNSDWPFGSIYCTIN        159
Serrata           GNGIVLWIVTGHRSMRTVTNYFLLNLSIADLLMSSLNCVFNFIFMLNSDWPFGSIYCTIN        159
Kikkawei          GNGIVLWIVTGHRSMRTVTNYFLLNLSIADLLMSSLNCVFNFIFMLNSDWPFGSIYCTIN        159
Rhopaloa          GNGIVLWIVTGHRSMRTVTNYFLLNLSIADLLMSSLNCVFNFIFMLNSDWPFGSIYCTIN        159
Elegans           GNGIVLWIVTGHRSMRTVTNYFLLNLSIADLLMSSLNCVFNFIFMLNSDWPFGSIYCTIN        159
Suzuki            GNGIVLWIVTGHRSMRTVTNYFLLNLSIADLLMSSLNCVFNFIFMLNSDWPFGSIYCTIN        180
Biarmipes         GNGIVLWIVTGHRSMRTVTNYFLLNLSIADLLMSSLNCVFNFIFMLNSDWPFGSIYCTIN        180
Takahashi         GNGIVLWIVTGHRSMRTVTNYFLLNLSIADLLMSSLNCVFNFIFMLNSDWPFGSIYCTIN        159
Eugracilis        GNGIVLWIVTGHRSMRTVTNYFLLNLSIADLLMSSLNCVFNFIFMLNSDWPFGSIYCTIN        159
Fichsuphila       GNGIVLWIVTGHRSMRTVTNYFLLNLSIADLLMSSLNCVFNFIFMLNSDWPFGSIYCTIN        180
Erecta            GNGIVLWIVTGHRSMRTVTNYFLLNLSIADLLMSSLNCVFNFIFMLNSDWPFGSIYCTIN        159
Mauritania        GNGIVLWIVTGHRSMRTVTNYFLLNLSIADLLMSSLNCVFNFIFMLNSDWPFGSIYCTIN        159
Melanogaster      GNGIVLWIVTGHRSMRTVTNYFLLNLSIADLLMSSLNCVFNFIFMLNSDWPFGSIYCTIN        159
```

```
Simulans          GNGIVLWIVTGHRSMRTVTNYFLLNLSIADLLMSSLNCVFNFIFMLNSDWPFGSIYCTIN      159
Sechellia         GNGIVLWIVTGHRSMRTVTNYFLLNLSIADLLMSSLNCVFNFIFMLNSDWPFGSIYCTIN      159
Mojavensis        GNGIVLWIVTGHRSMRTVTNYFLLNLSIADLLMSSLNCVFNFIFMLNSDWPFGSIYCTIN      159
Virilis          GNGIVLWIVTGHRSMRTVTNYFLLNLSIADLLMSTLNCVFNFIFMVNSDWPFGSIYCTIN       159
                  **********************************:**********:**************

Grimshawi         NFVANVTVSTSVFTLVAISFDRYIAIVHPLKRRTSRRKVRFILVMIWALSCVLSAPCLLY      219
Bipectinate       NFVANVTVSTSVFTLVAISFDRYIAIVHPLKRRTSRRKVRFILVLIWALSCVLSAPCLLY      219
Anannassae        NFVANVTVSTSVFTLVAISFDRYIAIVHPLKRRTSRRKVRFILVLIWALSCVLSAPCLLY      219
Serrata           NFVANVTVSTSVFTLVAISFDRYIAIVHPLKRRTSRRKVRFILVLIWVLSCVLSAPCLLY      219
Kikkawei          NFVANVTVSTSVFTLVAISFDRYIAIVHPLKRRTSRRKVRIILVLIWALSCVLSAPCLLY      219
Rhopaloa          NFVANVTVSTSVFTLVAISFDRYIAIVHPLKRRTSRRKVRIILVLIWALSCVLSAPCLLY      219
Elegans           NFVANVTVSTSVFTLVAISFDRYIAIVHPLKRRTSRRKVRIILVLIWALSCVLSAPCLLY      219
Suzuki            NFVANVTVSTSVFTLVAISFDRYIAIVHPLKRRTSRRKVRIILVLIWALSCVLSAPCLLY      240
Biarmipes         NFVANVTVSTSVFTLVAISFDRYIAIVHPLKRRTSRRKVRIILVLIWALSCVLSAPCLLY      240
Takahashi         NFVANVTVSTSVFTLVAISFDRYIAIVHPLKRRTSRRKVRIILVLIWALSCVLSAPCLLY      219
Eugracilis        NFVANVTVSTSVFTLVAISFDRYIAIVHPLKRRTSRRKVRIILVLIWALSCVLSAPCLLY      219
Fichsuphila       NFVANVTVSTSVFTLVAISFDRYIAIVHPLKRRTSRRKVRIILVLIWVLSCVLSAPCLLY      240
Erecta            NFVANVTVSTSVFTLVAISFDRYIAIVHPLKRRTSRRKVRIILVLIWALSCVLSAPCLLY      219
Mauritania        NFVANVTVSTSVFTLVAISFDRYIAIVHPLKRRTSRRKVRIILVLIWALSCVLSAPCLLY      219
Melanogaster      NFVANVTVSTSVFTLVAISFDRYIAIVHPLKRRTSRRKVRIILVLIWALSCVLSAPCLLY      219
Simulans          NFVANVTVSTSVFTLVAISFDRYIAIVHPLKRRTSRRKVRIILVLIWALSCVLSAPCLLY      219
Sechellia         NFVANVTVSTSVFTLVAISFDRYIAIVHPLKRRTSRRKVRIILVLIWALSCVLSAPCLLY      219
Mojavensis        NFVANVTVSTSVFTLVAISFDRYIAIVHPLKRRTSRRKVRFILVLIWALSCVLSAPCLLY      219
Virilis          NFVANVTVSTSVFTLVAISFDRYIAIVHPLKRRTSRRKVRFILVLIWALSCVLSAPCLLY       219
                  ****************************************:***:**.************

Grimshawi         SSIMTKHYYNGKSRTVCFMMWPDGRYPTSMADYVYNLTILVLTYGIPMIVMLICYTLMGR      279
Bipectinate       SSIMTKHYYNGKSRTVCFMMWPDGRYPTSMADYAYNLIILVLTYGIPMIVMLICYTLMGR      279
Anannassae        SSIMTKHYYNGKSRTVCFMMWPDGRYPTSMADYAYNLIILVLTYGIPMIVMLICYTLMGR      279
Serrata           SSIMTKHYYNGKSRTVCFMMWPDGRYPTSMADYAYNLIILVLTYGIPMIVMLICYSLMGR      279
Kikkawei          SSIMTKQYYNGKSRTVCFMMWPDGRYPTSMADYVYNLIILVLTYGIPMIVMLICYSLMGR      279
Rhopaloa          SSIMTKHYYNGKSRTVCFMMWPDGRYPTSMADYAYNLIILILTYGIPMIVMLICYSLMGR      279
Elegans           SSIMTKHYYNGKSRTVCFMMWPDGRYPTSMADYAYNLIILVLTYGIPMIVMLICYSLMGR      279
Suzuki            SSIMTKHYYNGKSRTVCFMMWPDGRYPTSMADYAYNLIILVLTYGVPMIVMLICYSLMGR      300
Biarmipes         SSIMTKHYYNGKSRTVCFMMWPDGRYPTSMADYAYNLIILVLTYGIPMIVMLICYSLMGR      300
Takahashi         SSIMTKHYYNGKSRTVCFMMWPDGRYPTSMADYAYNLIILVLTYGIPMIVMLICYSLMGR      279
Eugracilis        SSIMTKHYYNGKSRTVCFMMWPDGRYPTSMADYAYNLIILVLTYGIPMIVMLICYSLMGR      279
Fichsuphila       SSIMTKHYYNGKSRTVCFMMWPDGRYPTSMADYAYNLIILVLTYGIPMIVMLICYSLMGR      300
Erecta            SSIMTKHYYNGKSRTVCFMMWPDGRYPTSMADYAYNLIILVLTYGIPMIVMLICYSLMGR      279
Mauritania        SSIMTKHYYNGKSRTVCFMMWPDGRYPTSMADYAYNLIILVLTYGIPMIVMLICYSLMGR      279
Melanogaster      SSIMTKHYYNGKSRTVCFMMWPDGRYPTSMADYAYNLIILVLTYGIPMIVMLICYSLMGR      279
Simulans          SSIMTKHYYNGKSRTVCFMMWPDGRYPTSMADYAYNLIILVLTYGIPMIVMLICYSLMGR      279
Sechellia         SSIMTKHYYNGKSRTVCFMMWPDGRYPTSMADYAYNLIILVLTYGIPMIVMLICYSLMGR      279
Mojavensis        SSIMTKHYYNGKSRTVCFMMWPDGRYPTSMADYVYNLTLLVLTYGIPMIVMLVCYSLMGR      279
Virilis          SSIMTKHYYNGKSRTVCFMMWPDGRYPTSMTDYVYNVTILVLTYGIPMIVMLICYSLMGR       279
                  ******:***********************:**.**: :*:****:******:**:****

Grimshawi         VLWGSRSIGENTDRQMESMKSKRKVVRMFIAVVSIFAICWLPYHLFFIYAYHNNQVASTK      339
Bipectinate       VLWGSRSIGENTDRQMESMKSKRKVVRMFIAIVSIFAICWLPYHLFFIYAYHNNHVASTK      339
Anannassae        VLWGSRSIGENTDRQMESMKSKRKVVRMFIAIVSIFAICWLPYHLFFIYAYHNNHVASTK      339
Serrata           VLWGSRSIGENTDRQMESMKSKRKVVRMFIAIVSIFAICWLPYHLFFIYAYHNNQVASTK      339
Kikkawei          VLWGSRSIGENTDRQMESMKSKRKVVRMFIAIVSIFAICWLPYHLFFIYAYHNNQVASTK      339
Rhopaloa          VLWGSRSIGENTDRQMESMKSKRKVVRMFIAIVSIFAICWLPYHLFFIYAYHNNQVASTK      339
Elegans           VLWGSRSIGENTDRQMESMKSKRKVVRMFIAIVSIFAICWLPYHLFFIYAYHNNQVASTK      339
Suzuki            VLWGSRSIGENTDRQMESMKSKRKVVRMFIAIVSIFAICWLPYHLFFIYAYHNNQVASTK      360
Biarmipes         VLWGSRSIGENTDRQMESMKSKRKVVRMFIAIVSIFAICWLPYHLFFIYAYHNNQVASTK      360
Takahashi         VLWGSRSIGENTDRQMESMKSKRKVVRMFIAIVSIFAICWLPYHLFFIYAYHNNQVASTK      339
Eugracilis        VLWGSRSIGENTDRQMESMKSKRKVVRMFIAIVSIFAICWLPYHLFFIYAYHNNQVASTK      339
Fichsuphila       VLWGSRSIGENTDRQMESMKSKRKVVRMFIAIVSIFAICWLPYHLFFIYAYHNNQVASTK      360
Erecta            VLWGSRSIGENTDRQMESMKSKRKVVRMFIAIVSIFAICWLPYHLFFIYAYHNNQVASTK      339
Mauritania        VLWGSRSIGENTDRQMESMKSKRKVVRMFIAIVSIFAICWLPYHLFFIYAYHNNQVASTK      339
Melanogaster      VLWGSRSIGENTDRQMESMKSKRKVVRMFIAIVSIFAICWLPYHLFFIYAYHNNQVASTK      339
Simulans          VLWGSRSIGENTDRQMESMKSKRKVVRMFIAIVSIFAICWLPYHLFFIYAYHNNQVASTK      339
Sechellia         VLWGSRSIGENTDRQMESMKSKRKVVRMFIAIVSIFAICWLPYHLFFIYAYHNNQVASTK      339
Mojavensis        VLWGSRSIGENTERQIESMKSKRKVVRMFIAIVSIFAICWLPYHMFFIYAYHNNQVASTK      339
Virilis          VLWGSRSIGENTDRQMESMKSKRKVVRMFIAIVSIFAICWLPYHLFFIYAYHNNQVASTK       339
                  ************:**:***************:************:*********:*****

Grimshawi         YVQHMYLGFYWLAMSNAMVNPIIYYWMNKRFRMYFQRIVCCCCLGFIRYRFDSPKSQATN      399
Bipectinate       YVQHMYLGFYWLAMSNAMVNPIIYYWMNKRFRMYFQRIICCCCMGFTRHRFDSPKSRLTN      399
```

```
Anannassae        YVQHMYLGFYWLAMSNAMVNPIIYYWMNKRFRMYFQRIICCCCMGFTRHRFDSPKSRLTN    399
Serrata           YVQHMYLGFYWLAMSNAMVNPIIYYWMNKRFRMYFQRIICCCCVGLTRHRFDSPKSRLTN    399
Kikkawei          YVQHMYLGFYWLAMSNAMVNPIIYYWMNKRFRMYFQRIICCCCVGLTRHRFDSPKSRLTN    399
Rhopaloa          YVQHMYLGFYWLAMSNAMVNPLIYYWMNKRFRMYFQRIICCCCVSLTRHRFDSPKSRLTN    399
Elegans           YVQHMYLGFYWLAMSNAMVNPLIYYWMNKRFRMYFQRIICCCCVGLTRHRFDSPKSRLTN    399
Suzuki            YVQHMYLGFYWLAMSNAMVNPLIYYWMNKRFRMYFQRIICCCCVSLTRHRFDSPKSRLTN    420
Biarmipes         YVQHMYLGFYWLAMSNAMVNPLIYYWMNKRFRMYFQRIICCCCVGLTRHRFDSPKSRLTN    420
Takahashi         YVQHMYLGFYWLAMSNAMVNPLIYYWMNKRFRMYFQRIICCCCVGLTRHRFDSPKSRLTN    399
Eugracilis        YVQHMYLGFYWLAMSNAMVNPLIYYWMNKRFRMYFQRIICCCCVGLTRHRFDSPKSRLTN    399
Fichsuphila       YVQHMYLGFYWLAMSNAMVNPLIYYWMNKRFRMYFQRIICCCCVGLTRHRFDSPKSRLTN    420
Erecta            YVQHMYLGFYWLAMSNAMVNPLIYYWMNKRFRMYFQRIICCCCVGLTRHRFDSPKSRLTN    399
Mauritania        YVQHMYLGFYWLAMSNAMVNPLIYYWMNKRFRMYFQRIICCCCVGLTRHRFDSPKSRLTN    399
Melanogaster      YVQHMYLGFYWLAMSNAMVNPLIYYWMNKRFRMYFQRIICCCCVGLTRHRFDSPKSRLTN    399
Simulans          YVQHMYLGFYWLAMSNAMVNPLIYYWMNKRFRMYFQRIICCCCVGLTRHRFDSPKSRLTN    399
Sechellia         YVQHMYLGFYWLAMSNAMVNPLIYYWMNKRFRMYFQRIICCCCVGLTRHRFDSPKSRLTN    399
Mojavensis        YVQHMYLGFYWLAMSNAMVNPIIYYWMNKRFRMYFQRIIFCCCMGLTRYKFDSPKSRLAN    399
Virilis          YVQHMYLGFYWLAMSNAMVNPIIYYWMNKRFRMYFQRIIFCCCLGLMRYRFESPKSRMAN     399
                  *********************:****************: ***:.: *::*:****: :*

Grimshawi         KNSSHRNTR-------------------------------------------------AE    410
Bipectinate       KNSSNRHTR-------------------------------------------------AE    410
Anannassae        KNSSHRHTR-------------------------------------------------AE    410
Serrata           KNSSHRHTR-------------------------------------------------AE    410
Kikkawei          KNSSHRHTR-------------------------------------------------AE    410
Rhopaloa          KNSSHRHTR-------------------------------------------------AE    410
Elegans           KNSSHRHTR-------------------------------------------------AE    410
Suzuki            KNSSHRHTR-------------------------------------------------AE    431
Biarmipes         KNSSHRHTR-------------------------------------------------AE    431
Takahashi         KNSSHRHTR-------------------------------------------------AE    410
Eugracilis        KNSSHRHTR-------------------------------------------------AE    410
Fichsuphila       KNSSHRHTR-------------------------------------------------AE    431
Erecta            KNSSHRHTR-------------------------------------------------AE    410
Mauritania        KNSSHRHTRGGYTVAHSLPNSSPPTTQTILAVLAQTQPMPQTQLLLSHYSPHPTQPSAAE    459
Melanogaster      KNSSNRHTR-------------------------------------------------AE    410
Simulans          KNSSHRHTR-------------------------------------------------AE    410
Sechellia         KNSSHRHTR-------------------------------------------------AE    410
Mojavensis        KNSSNRNTR-------------------------------------------------VE    410
Virilis          KNSSNRHTR-------------------------------------------------AE     410
                  ****:*:**                                                 .*

Grimshawi         TKSQWKRSTMETQIQQVPTSSSRDR--------------QAGAVQGLNHTAVECIIERP-    455
Bipectinate       TKSQWKRSTMETQIQAPPITHTNSSKDKEREKHLQQV-----QTELHHRKAVECIMEGPG    465
Anannassae        TKSQWKRSTMETQIQAPPITHTNSSKDRERERHLQQ-------TELQHRKAVECIMEGPG    463
Serrata           TKSQWKRSTMETQIQQVPTTNSCRDQRQMTQSQS--L-----GGVATPRPAVECIMERP-    462
Kikkawei          TKSQWKRSTMETQIHQVPTTNSCRDQRQLTQSQS--LGAHSGGGVGTPRPAVECIMERP-    467
Rhopaloa          NKSQWKRSTMETQIQQAPLTNSGRDQRTAQP---------LKAVDGPSRVGVECIMERP-    460
Elegans           NKSQWKRSTMETQIQQAPLTNSGRDPRTPQP---------LKVVDGASRVGVECIMERP-    460
Suzuki            TKSQWKRSTMETQIQQAPVSNSGRDQRMRDGYANQPAQAGVGGAGGPNRAAVECIMERP-    490
Biarmipes         TKSQWKRSTMETQIQQAPVSNSGRDQRMRDGYPS---QAQAGVAGGPNRAAVECIMERP-    487
Takahashi         TKSQWKRSTMETQIQQAPVSNSCRDPRMREATA-QPQTCGVGGP-NANRSAVECIMERP-    467
Eugracilis        TKSQWKRSTMETQIQQAPANSTGRDQRN----A-QQTPTGAGIG-GPNRTAVECIMERP-    463
Fichsuphila       TKSQWKRSTMETQIQQAPLTSSGRDQRN----A-QTQ--SLGGA-GSNRTAVECIMERP-    482
Erecta            TKSQWKRSTMETQIQQAPVTSSGRDQRS----A-QQQQHHPSGA-ATNRAAVECIMERP-    463
Mauritania        TKSQWKRSTMETQIQQAPVTSSCREQRS----A-QQQ--QPPGS-GTNRAAVECIMERP-    510
Melanogaster      TKSQWKRSTMETQIQQAPVTSSCREQRS----A-QQQ--QPPGS-GTNRAAVECIMERP-    461
Simulans          TKSQWKRSTMETQIQQAPVTSSCREQRS----A-QQQ--QPPGS-GTNRAAVECIMERP-    461
Sechellia         TKSQWKRSTMETQIQQAPVTSSCREQRS----A-QQQ--QPPGS-ETNRAAVECIMERP-    461
Mojavensis        TKSQWKRSTMETQIQQVPTTSSRER---------------KAAGAAVNHMAVECVIERP-    454
Virilis          TKSQWKRSTMETQIQQMPKTSSRDK---------------DAGVQGLNHTAVECIIERP-     454
                  .*************:  * . :                         : .***::* *

Grimshawi         --VDESGSPICLSINNSAGERQRVKIKYISCDEDNNPIEHESESGHSHGHTHRHSHRHSL    513
Bipectinate       GNADGNSSPLCLSINNSMGERQRVKIKYISCDEDNNPVEHSPKQL---------------    510
Anannassae        GHADGNSSPLCLSINNSVGERQRVKIKYISCDEDNNPVENSPKQL---------------    508
Serrata           --MDGNSSPLCLSINNSVGERQRVKIKYISCDEDNNPVELEHSSGPKQL-----------    509
Kikkawei          --LDGNSSPLCLSINNSVGERQRVKIKYISCDEDNNPVELEHSSGPKQL-----------    514
Rhopaloa          --ADGNSSPLCLSINNSVGERQRVKIKYISCDEDNNPVEHTPKQ----L-----------    503
Elegans           --ADGNSSPLCLSINNSVGERQRVKIKYISCDEDNNPVEHSPKH----L-----------    503
Suzuki            --LDGNSSPLCLSINNSLGERQRMKIKYISCDEDNNPVELSPNQ----L-----------    533
Biarmipes         --LDGNSSPLCLSINNSLGERQRLKIKYISCDEDNNPVELSPNQ----L-----------    530
Takahashi         --VDGTSSPLCLSINNSLGERQRVKIKYISCDEDNNPVELSPKQ----L-----------    510
```

```
Eugracilis        --TDGTSSPLCLSINNSVGERQRVKIKYISCDEDNNPVELSPKQ----L-----------        506
Fichsuphila       --ADGTSSPLCLSINNSVGERQRVKIKYISCDEDNNPVEHSPKQ----L-----------        525
Erecta            --ADGSSSPLCLSINNSIGERQRVKIKYISCDEDNNPVELSPKQ----L-----------        506
Mauritania        --ADGSSSPLCLSINNSIGERQRVKIKYISCDEDNNPVELSPKQ----L-----------        553
Melanogaster      --ADGSSSPLCLSINNSIGERQRVKIKYISCDEDNNPVELSPKQ----M-----------        504
Simulans          --ADGSSSPLCLSINNSIGERQRVKIKYISCDEDNNPVELSPKQ----L-----------        504
Sechellia         --ADGSSSPLCLSINNSIGERQRVKIKYISCDEDNNPVELSPKQ----L-----------        504
Mojavensis        --MDDNSSPICLSINNSAGERQRLRIKYISCDEDNNPIEEASESSSNHSHGHSHNHSRSH        512
Virilis          --IDDNSSPICLSIKNSAGERQRVKIKYISCDEDNNPIEEGSENNSSHDSNHSHGHGHSC        512
                     *  ..**:****:** *****::************:*    .
```

Melanogaster **NP_524304.2**

```
  1 mseivdtell vnctilavrr felnsivntt llgslnrtev vsllssiidn rdnlesinea
     61 kdflteclfp sptrpyelpw eqktiwaiif glmmfvaiag ngivlwivtg hrsmrtvtny
    121 fllnlsiadl lmsslncvfn fifmlnsdwp fgsiyctinn fvanvtvsts vftlvaisfd
    181 ryiaivhplk rrtsrrkvri ilvliwalsc vlsapcllys simtkhyyng ksrtvcfmmw
    241 pdgryptsma dyaynliilv ltygipmivm licyslmgrv lwgsrsigen tdrqmesmks
    301 krkvvrmfia ivsifaicwl pyhlffiyay hnnqvastky vqhmylgfyw lamsnamvnp
    361 liyywmnkrf rmyfqriicc ccvgltrhrf dspksrltnk nssnrhtrae tksqwkrstm
    421 etqiqqapvt sscreqrsaq qqqppgsgtn raavecimer padgsssplc lsinnsiger
    481 qrvkikyisc dednnpvels pkqm
```

Simulans **XP_002103960.1**

```
  1 mseivdtell vnctilavrr felnsivntt llgslnrtev vsllssiidn rdnlesinea
     61 kdflteclfp sptrpyelpw eqktiwaiif glmmfvaiag ngivlwivtg hrsmrtvtny
    121 fllnlsiadl lmsslncvfn fifmlnsdwp fgsiyctinn fvanvtvsts vftlvaisfd
    181 ryiaivhplk rrtsrrkvri ilvliwalsc vlsapcllys simtkhyyng ksrtvcfmmw
    241 pdgryptsma dyaynliilv ltygipmivm licyslmgrv lwgsrsigen tdrqmesmks
    301 krkvvrmfia ivsifaicwl pyhlffiyay hnnqvastky vqhmylgfyw lamsnamvnp
    361 liyywmnkrf rmyfqriicc ccvgltrhrf dspksrltnk nsshrhtrae tksqwkrstm
    421 etqiqqapvt sscreqrsaq qqqppgsgtn raavecimer padgsssplc lsinnsiger
    481 qrvkikyisc dednnpvels pkql
```

Suzuki **XP_036674593.1**

```
        1 mwhkcggiit qrslipikgs rmseivdtel lvnctilavr rfelnsivnt tllgslnrte
       61 vvsllssiid nrdnlesine akdflteclf psptrpyelp weqktiwaii fglmmfvaia
      121 gngivlwivt ghrsmrtvtn yfllnlsiad llmsslncvf nfifmlnsdw pfgsiyctin
      181 nfvanvtvst svftlvaisf dryiaivhpl krrtsrrkvr iilvliwals cvlsapclly
      241 ssimtkhyyn gksrtvcfmm wpdgryptsm adyaynliil vltygvpmiv mlicyslmgr
      301 vlwgsrsige ntdrqmesmk skrkvvrmfi aivsifaicw lpyhlffiya yhnnqvastk
      361 yvqhmylgfy wlamsnamvn pliyywmnkr frmyfqriic cccvsltrhr fdspksrltn
      421 knsshrhtra etksqwkrst metqiqqapv snsgrdqrmr dgyanqpaqa gvggaggpnr
      481 aavecimerp ldgnssplcl sinnslgerq rmkikyiscd ednnpvelsp nql
```

Mauritania **XP_033163933.1**

```
1 mseivdtell vnctilavrr felnsivntt llgslnrtev vsllssiidn rdnlesinea
      61 kdflteclfp sptrpyelpw eqktiwaiif glmmfvaiag ngivlwivtg hrsmrtvtny
     121 fllnlsiadl lmsslncvfn fifmlnsdwp fgsiyctinn fvanvtvsts vftlvaisfd
     181 ryiaivhplk rrtsrrkvri ilvliwalsc vlsapcllys simtkhyyng ksrtvcfmmw
     241 pdgryptsma dyaynliilv ltygipmivm licyslmgrv lwgsrsigen tdrqmesmks
     301 krkvvrmfia ivsifaicwl pyhlffiyay hnnqvastky vqhmylgfyw lamsnamvnp
     361 liyywmnkrf rmyfqriicc ccvgltrhrf dspksrltnk nsshrhtrgg ytvahslpns
     421 spttqtila vlaqtqpmpq tqlllshysp hptqpsaaet ksqwkrstme tqiqqapvts
     481 screqrsaqq qqppgsgtnr aavecimerp adgsssplcl sinnsigerq rvkikyiscd
     541 ednnpvelsp kql
```

Sechellia **XP_032578092.1**

```
  1 mseivdtell vnctilavrr felnsivntt llgslnrtev vsllssiidn rdnlesinea
     61 kdflteclfp sptrpyelpw eqktiwaiif glmmfvaiag ngivlwivtg hrsmrtvtny
    121 fllnlsiadl lmsslncvfn fifmlnsdwp fgsiyctinn fvanvtvsts vftlvaisfd
    181 ryiaivhplk rrtsrrkvri ilvliwalsc vlsapcllys simtkhyyng ksrtvcfmmw
    241 pdgryptsma dyaynliilv ltygipmivm licyslmgrv lwgsrsigen tdrqmesmks
    301 krkvvrmfia ivsifaicwl pyhlffiyay hnnqvastky vqhmylgfyw lamsnamvnp
    361 liyywmnkrf rmyfqriicc ccvgltrhrf dspksrltnk nsshrhtrae tksqwkrstm
    421 etqiqqapvt sscreqrsaq qqqppgsetn raavecimer padgsssplc lsinnsiger
```

481 qrvkikyisc dednnpvels pkql

Serrata **XP_020798481.1**

```
1 mseivdtell vnctilavrr felnsivntt llnslnrtev vsllssiidn rdnlgsinea
       61 kdflteclfp sptrpyelpw eqktiwaiif glmmfvaiag ngivlwivtg hrsmrtvtny
      121 fllnlsiadl lmsslncvfn fifmlnsdwp fgsiyctinn fvanvtvsts vftlvaisfd
      181 ryiaivhplk rrtsrrkvrf ilvliwvlsc vlsapcllys simtkhyyng ksrtvcfmmw
      241 pdgryptsma dyaynliilv ltygipmivm licyslmgrv lwgsrsigen tdrqmesmks
      301 krkvvrmfia ivsifaicwl pyhlffiyay hnnqvastky vqhmylgfyw lamsnamvnp
      361 iiyywmnkrf rmyfqriicc ccvgltrhrf dspksrltnk nsshrhtrae tksqwkrstm
      421 etqiqqvptt nscrdqrqmt qsqslggvat prpavecime rpmdgnsspl clsinnsvge
      481 rqrvkikyis cdednnpvel ehssgpkql
```

Erecta **XP_001980652.1**

```
  1 mseivdtell vnctilavrr felnsivntt llgslnrtev vnllssiidn rdnlesinea
       61 kdflteclfp sptrpyelpw eqktiwaiif glmmfvaiag ngivlwivtg hrsmrtvtny
      121 fllnlsiadl lmsslncvfn fifmlnsdwp fgsiyctinn fvanvtvsts vftlvaisfd
      181 ryiaivhplk rrtsrrkvri ilvliwalsc vlsapcllys simtkhyyng ksrtvcfmmw
      241 pdgryptsma dyaynliilv ltygipmivm licyslmgrv lwgsrsigen tdrqmesmks
      301 krkvvrmfia ivsifaicwl pyhlffiyay hnnqvastky vqhmylgfyw lamsnamvnp
      361 liyywmnkrf rmyfqriicc ccvgltrhrf dspksrltnk nsshrhtrae tksqwkrstm
      421 etqiqqapvt ssgrdqrsaq qqqhhpsgaa tnraavecim erpadgsssp lclsinnsig
      481 erqrvkikyi scdednnpve lspkql
```

Takahashi **XP_017004985.2**

```
    1 mseivdtell vnctilavrr felnsivntt llsslnrtev vsllssiidn rdnlesinea
       61 kdflteclfp sptrpyelpw eqktiwaiif glmmfvaiag ngivlwivtg hrsmrtvtny
      121 fllnlsiadl lmsslncvfn fifmlnsdwp fgsiyctinn fvanvtvsts vftlvaisfd
      181 ryiaivhplk rrtsrrkvri ilvliwalsc vlsapcllys simtkhyyng ksrtvcfmmw
      241 pdgryptsma dyaynliilv ltygipmivm licyslmgrv lwgsrsigen tdrqmesmks
      301 krkvvrmfia ivsifaicwl pyhlffiyay hnnqvastky vqhmylgfyw lamsnamvnp
      361 liyywmnkrf rmyfqriicc ccvgltrhrf dspksrltnk nsshrhtrae tksqwkrstm
      421 etqiqqapvs nscrdprmre ataqpqtcgv ggpnanrsav ecimerpvdg tssplclsin
      481 nslgerqrvk ikyiscdedn npvelspkql
```

Biarmipes **XP_016959801.1**

```
1 mwrkcggiit qrslipikgc rmseivdtel lvnctilavr rfelnsivnt tllgslnrte
       61 vvsllssiid nrdnlesine akdflteclf psptrpyelp weqktiwaii fglmmfvaia
      121 gngivlwivt ghrsmrtvtn yfllnlsiad llmsslncvf nfifmlnsdw pfgsiyctin
      181 nfvanvtvst svftlvaisf dryiaivhpl krrtsrrkvr iilvliwals cvlsapclly
      241 ssimtkhyyn gksrtvcfmm wpdgryptsm adyaynliil vltygipmiv mlicyslmgr
      301 vlwgsrsige ntdrqmesmk skrkvvrmfi aivsifaicw lpyhlffiya yhnnqvastk
      361 yvqhmylgfy wlamsnamvn pliyywmnkr frmyfqriic cccvgltrhr fdspksrltn
      421 knsshrhtra etksqwkrst metqiqqapv snsgrdqrmr dgypsqaqag vaggpnraav
      481 ecimerpldg nssplclsin nslgerqrlk ikyiscdedn npvelspnql
```

Eugracilis **XP_017083539.1**

```
   1 mseivdtell vnctilavrr felnsivntt llgslnrtev vsllssiidn rdnlesinea
       61 kdflteclfp sptrpyelpw eqktiwaiif glmmfvaiag ngivlwivtg hrsmrtvtny
      121 fllnlsiadl lmsslncvfn fifmlnsdwp fgsiyctinn fvanvtvsts vftlvaisfd
      181 ryiaivhplk rrtsrrkvri ilvliwalsc vlsapcllys simtkhyyng ksrtvcfmmw
      241 pdgryptsma dyaynliilv ltygipmivm licyslmgrv lwgsrsigen tdrqmesmks
      301 krkvvrmfia ivsifaicwl pyhlffiyay hnnqvastky vqhmylgfyw lamsnamvnp
      361 liyywmnkrf rmyfqriicc ccvgltrhrf dspksrltnk nsshrhtrae tksqwkrstm
      421 etqiqqapan stgrdqrnaq qtptgagigg pnrtavecim erptdgtssp lclsinnsvg
      481 erqrvkikyi scdednnpve lspkql
```

Rhopaloa **XP_016989968.2**

```
1 mseivdtell vnctilavrr felnsivntt llgslnrtev vsllssiidn rdnldsinea
       61 kdflteclfp sptrpyelpw eqktiwaiif glmmfvaiag ngivlwivtg hrsmrtvtny
      121 fllnlsiadl lmsslncvfn fifmlnsdwp fgsiyctinn fvanvtvsts vftlvaisfd
      181 ryiaivhplk rrtsrrkvri ilvliwalsc vlsapcllys simtkhyyng ksrtvcfmmw
```

241 pdgryptsma dyaynliili ltygipmivm licyslmgrv lwgsrsigen tdrqmesmks
301 krkvvrmfia ivsifaicwl pyhlffiyay hnnqvastky vqhmylgfyw lamsnamvnp
361 liyywmnkrf rmyfqriicc ccvsltrhrf dspksrltnk nsshrhtrae nksqwkrstm
421 etqiqqaplt nsgrdqrtaq plkavdgpsr vgvecimerp adgnssplcl sinnsvgerq
481 rvkikyiscd ednnpvehtp kql

Fichsuphila **XP_017057393.1**

1 mwhecsgiit qrprivikvw kmseivdtel lvnctilavr rfelnsivnt tllgtlnrte
61 vvsllssiie nrdnlqsine akdflteclf psptrpyelp weqktiwaii fglmmfvaia
121 gngivlwivt ghrsmrtvtn yfllnlsiad llmsslncvf nfifmlnsdw pfgsiyctin
181 nfvanvtvst svftlvaisf dryiaivhpl krrtsrrkvr iilvliwvls cvlsapclly
241 ssimtkhyyn gksrtvcfmm wpdgryptsm adyaynliil vltygipmiv mlicyslmgr
301 vlwgsrsige ntdrqmesmk skrkvvrmfi aivsifaicw lpyhlffiya yhnnqvastk
361 yvqhmylgfy wlamsnamvn pliyywmnkr frmyfqriic cccvgltrhr fdspksrltn
421 knsshrhtra etksqwkrst metqiqqapl tssgrdqrna qtqslggags nrtavecime
481 rpadgtsspl clsinnsvge rqrvkikyis cdednnpveh spkql

Elegans **XP_017112407.1**

1 mseivdtell vnctilavrr felnsivntt llgslnrtev vsllssiien rdnlqsinea
61 kdflteclfp sptrpyelpw eqktiwaiif glmmfvaiag ngivlwivtg hrsmrtvtny
121 fllnlsiadl lmsslncvfn fifmlnsdwp fgsiyctinn fvanvtvsts vftlvaisfd
181 ryiaivhplk rrtsrrkvri ilvliwalsc vlsapcllys simtkhyyng ksrtvcfmmw
241 pdgryptsma dyaynliilv ltygipmivm licyslmgrv lwgsrsigen tdrqmesmks
301 krkvvrmfia ivsifaicwl pyhlffiyay hnnqvastky vqhmylgfyw lamsnamvnp
361 liyywmnkrf rmyfqriicc ccvgltrhrf dspksrltnk nsshrhtrae nksqwkrstm
421 etqiqqaplt nsgrdprtpq plkvvdgasr vgvecimerp adgnssplcl sinnsvgerq
481 rvkikyiscd ednnpvehsp khl

Kikkawei **XP_017020386.1**

1 mseivdtell vnctilavrr felnsivntt llnslnrtev vsllssiidn pdnlgsinea
61 kdflteclfp sptrpyelpw eqktiwaiif glmmfvaiag ngivlwivtg hrsmrtvtny
121 fllnlsiadl lmsslncvfn fifmlnsdwp fgsiyctinn fvanvtvsts vftlvaisfd
181 ryiaivhplk rrtsrrkvri ilvliwalsc vlsapcllys simtkqyyng ksrtvcfmmw
241 pdgryptsma dyvynliilv ltygipmivm licyslmgrv lwgsrsigen tdrqmesmks
301 krkvvrmfia ivsifaicwl pyhlffiyay hnnqvastky vqhmylgfyw lamsnamvnp
361 iiyywmnkrf rmyfqriicc ccvgltrhrf dspksrltnk nsshrhtrae tksqwkrstm
421 etqihqvptt nscrdqrqlt qsqslgahsg ggvgtprpav ecimerpldg nssplclsin
481 nsvgerqrvk ikyiscdedn npvelehssg pkql

Bipectinate **XP_017104778.2**

1 msetvdtell vnctilavrr felnsivntt llsslnrtev vgllsgiien rdnleninea
61 kdflteclfp sptrpyelpw eqktiwaiif glmmfvaiag ngivlwivtg hrsmrtvtny
121 fllnlsiadl lmsslncvfn fifmvnsdwp fgsiyctinn fvanvtvsts vftlvaisfd
181 ryiaivhplk rrtsrrkvrf ilvliwalsc vlsapcllys simtkhyyng ksrtvcfmmw
241 pdgryptsma dyaynliilv ltygipmivm licytlmgrv lwgsrsigen tdrqmesmks
301 krkvvrmfia ivsifaicwl pyhlffiyay hnnhvastky vqhmylgfyw lamsnamvnp
361 iiyywmnkrf rmyfqriicc ccmgftrhrf dspksrltnk nssnrhtrae tksqwkrstm
421 etqiqappit htnsskdker ekhlqqvqte lhhrkaveci megpggnadg nssplclsin
481 nsmgerqrvk ikyiscdedn npvehspkql

Anannassae **XP_001952989.1**

1 msetvdtell vnctilavrr felnsivntt llsslnrsev vgllsgiien rdnlesinea
61 kdflteclfp sptrpyelpw eqktiwaiif glmmfvaiag ngivlwivtg hrsmrtvtny
121 fllnlsiadl lmsslncvfn fifmvnsdwp fgsiyctinn fvanvtvsts vftlvaisfd
181 ryiaivhplk rrtsrrkvrf ilvliwalsc vlsapcllys simtkhyyng ksrtvcfmmw
241 pdgryptsma dyaynliilv ltygipmivm licytlmgrv lwgsrsigen tdrqmesmks
301 krkvvrmfia ivsifaicwl pyhlffiyay hnnhvastky vqhmylgfyw lamsnamvnp
361 iiyywmnkrf rmyfqriicc ccmgftrhrf dspksrltnk nsshrhtrae tksqwkrstm
421 etqiqappit htnsskdrer erhlqqtelq hrkavecime gpgghadgns splclsinns
481 vgerqrvkik yiscdednnp venspkql

Mojavensis **XP_001998315.1**

```
1 mseivdtell vnctilavrr felnsivntt llnslnrtev vsllsgiien rdnlgsinea
       61 kdflteclfp sptrpyelpw eqktiwaivf glmmfvaiag ngivlwivtg hrsmrtvtny
      121 fllnlsiadl lmsslncvfn fifmlnsdwp fgsiyctinn fvanvtvsts vftlvaisfd
      181 ryiaivhplk rrtsrrkvrf ilvliwalsc vlsapcllys simtkhyyng ksrtvcfmmw
      241 pdgryptsma dyvynltllv ltygipmivm lvcyslmgrv lwgsrsigen terqiesmks
      301 krkvvrmfia ivsifaicwl pyhmffiyay hnnqvastky vqhmylgfyw lamsnamvnp
      361 iiyywmnkrf rmyfqriifc ccmgltrykf dspksrlank nssnrntrve tksqwkrstm
      421 etqiqqvptt ssrerkaaga avnhmavecv ierpmddnss piclsinnsa gerqrlriky
      481 iscdednnpi eeasesssnh shghshnhsr shthahrcsh rqkttgtqel
```

Virilis [XP_002053610.1]

```
1 mseivdtell vnctilavrr felntivntt llntlnrtev vgllsgiien rdnldsinea
       61 kdflteclfp sptrpyelpw eqktiwaivf glmmfvaiag ngivlwivtg hrsmrtvtny
      121 fllnlsiadl lmstlncvfn fifmvnsdwp fgsiyctinn fvanvtvsts vftlvaisfd
      181 ryiaivhplk rrtsrrkvrf ilvliwalsc vlsapcllys simtkhyyng ksrtvcfmmw
      241 pdgryptsmt dyvynvtilv ltygipmivm licyslmgrv lwgsrsigen tdrqmesmks
      301 krkvvrmfia ivsifaicwl pyhlffiyay hnnqvastky vqhmylgfyw lamsnamvnp
      361 iiyywmnkrf rmyfqriifc cclglmryrf espksrmank nssnrhtrae tksqwkrstm
      421 etqiqqmpkt ssrdkdagvq glnhtaveci ierpiddnss piclsiknsa gerqrvkiky
      481 iscdednnpi eegsennssh dsnhshghgh scgrshgqna kaiqql
```

Grimshawi [XP_001994653.1]

```
   1 mpeivdtell vnctilavlr felnsivnit llnslnrtev vslltsiidn rdnlesinea
      61 kdflteclfp sptrpyelpw eqktiwaivf glmmfvaiag ngivlwivtg hrsmrtvtny
     121 fllnlsiadl lmsslncvfn fifmvnsdwp fgsiyctinn fvanvtvsts vftlvaisfd
     181 ryiaivhplk rrtsrrkvrf ilvmiwalsc vlsapcllys simtkhyyng ksrtvcfmmw
     241 pdgryptsma dyvynltilv ltygipmivm licytlmgrv lwgsrsigen tdrqmesmks
     301 krkvvrmfia vvsifaicwl pyhlffiyay hnnqvastky vqhmylgfyw lamsnamvnp
     361 iiyywmnkrf rmyfqrivcc cclgfiryrf dspksqatnk nsshrntrae tksqwkrstm
     421 etqiqqvpts ssrdrqagav qglnhtavec iierpvdesg spiclsinns agerqrvkik
     481 yiscdednnp iehesesghs hghthrhshr hslsnsygrs raqnaalvqq l
```
