## Supplementary material for "The incidence of candidate binding sites for β-arrestin in Drosophila neuropeptide GPCRs": S11 Text

**S11. Text  Multi-species analysis of CAPA R PA isoforms**
**Supporting Figure 13**

CLUSTAL Line-ups; Full sequences or restricted to the 7th TM and CT
Genbank Reference IDs below
Predicted 7th TM domain in YELLOW
BBS sequences in RED

```
bipectinata        RMLAAVVITFFVCWFPFHLQRLWFLYAQNNENYYNVNEWLFSIAGFAYYVSCTINPIVYS     351
anannassae         RMLAAVVITFFVCWFPFHMQRLWFLYAKDNENYYNVNEWLFSIAGFAYYVSCTINPIVYS     352
serrata            RMLAAVVITFFVCWFPFHVQRLWFLYAQENDNYLDINEALFSIAGFAYYVSCTINPIVYS     357
kikkawei           RMLAAVVITFFVCWFPFHVQRLWFLYAQENDNYLDINEALFSIAGFAYYVSCTINPIVYS     357
erecta             RMLAAVVITFFVCWFPFHLQRLIFLYATNMDNYLDINEALFSIAGFAYYVSCTVNPIVYS     357
melanogaster       RMLAAVVITFFVCWFPFHLQRLIFLYAKNMDNYLDINEALFSIAGFAYYVSCTVNPIVYS     357
sechelia           RMLAAVVITFFVCWFPFHLQRLIFLYAKNMENYLDINEALFSIAGFAYYVSCTVNPIVYS     343
simulans           RMLAAVVITFFVCWFPFHLQRLIFLYAKNMENYLDINEALFSIAGFAYYVSCTVNPIVYS     358
mauritania         RMLAAVVITFFVCWFPFHLQRLIFLYAKNMENYLDINEALFSIAGFAYYVSCTVNPIVYS     358
ficusphila         RMLAAVVITFFVCWFPFHLQRLIFLYAKDLDNYLDINEALFSIAGFAYYVSCTVNPIVYS     357
takahashi          RMLAAVVITFFVCWFPFHLQRLIFLYAKDMENYYDINEALFSIAGFAYYVSCTINPIVYS     356
eugracilis         RMLAAVVITFFVCWFPFHLQRLIFLYAKNMENYLDINEALFSIAGFAYYVSCTINPIVYS     357
biarmpes           RMLAAVVITFFVCWFPFHLQRLIFLYAKDMENYLDINEALFSIAGFAYYVSCTVNPIVYS     357
suzuki             RMLAAVVITFFVCWFPFHLQRLIFLYAKNMENYLDINEALFSIAGFAYYISCTVNPIVYS     357
rhopaloa           RMLAAVVITFFVCWFPFHLQRLIFLYAKNLENYLDINEALFSIAGFAYYVSCTINPIVYS     357
elegans            RMLAAVVITFFVCWFPFHLQRLIFLYAKDMDNYLDINEALFSIAGFAYYVSCTINPIVYS     356
grimshawi          RMLAAVVVTFFVCWFPFHLQRLWFLYAKNNDNYQDVNEWLFSIAGFAYYVSCTINPIVYN     328
virilis            RMLAAVVITFFVCWFPFHLQRLWFLYAKNIANYQDVNEWLFSIAGFAYYVSCTINPIVYN     351
mojavensis         RMLAAVVITFFVCWFPFHLQRLWFLYAKNFACFQNVNEWLFSIAGFAYYVSCTINPIVYN     347

bipectinata        VMSRRYRVAFRELLCGRPVGAYYNSGFARDHSSFRESTATSMGNHINYDRVHSVHVRASR     411
anannassae         VMSRRYRVAFRELLCGRPVGAYYNSGFARDHSSFRESTATSMGNHINYDRVHSVHVRASR     412
serrata            VMSRRYRVAFRELLCGRPVGAYYNSGFARDHSSFRESTATVLGKNVNYDRVHSVHVRSSR     417
kikkawei           VMSRRYRVAFRELLCGRPAGAYYNSGFARDHSSFRESTATTLGKNVNYDRVHSVHVRSSR     417
erecta             VMSRRYRVAFRELLCGKAVGAYYNSGFARDHSSFRESS--------AYDRVHSVHVRASR     409
melanogaster       VMSRRYRVAFRELLCGKAVGAYYNSGFARDHSSFRESS--------AYDRVHSVHVRASQ     409
sechelia           VMSRRYRVAFRELLCGKAVGAYYNSGFARDHSSFRESS--------AYDRVHSVHVRSSR     395
simulans           VMSRRYRVAFRELLCGKAVGAYYNSGFARDHSSFRESS--------AYDRVHSVHVRASR     410
mauritania         VMSRRYRVAFRELLCGKAVGAYYNSGFARDHSSFRESS--------AYDRVHSVHVRASR     410
ficusphila         VMSRRYRVAFRELLCGKPVGAYYNSGFARDHSSFRDSTATSMGNNIHYDRVHSVHVRSSR     417
takahashi          VMSRRYRVAFRELLCGKALGAYYNSAFARDHSSFRESTATSLGNNINYDRVHSVHVKSSR     416
eugracilis         VMSRRYRVAFRELLCGKAVGAYYNSGFARDHSSFRESTGTSLGNNINYDRVHSVHVRSSR     417
biarmpes           VMSRRYRVAFRELLCGKPVGAYYNSGFARDHSSFRESTATSLGNNINYDRVHSVHVTSSR     417
suzuki             VMSRRYRVAFRELLCGKAVGAYYNSGFARDHSSFRESTATSLGNNINYDRVHSVHVRSSR     417
rhopaloa           VMSRRYRVAFRELLCGKAVGAYYNSAFARDHSSFRESTHTSLGNNINYDRVHSVHVRSSR     417
elegans            VMSRRYRVAFRELLCGKAVGAYYNSAFARDHSSFRESTATTMGNNINYDRVHSVHVRSSR     416
grimshawi          VMSHRYRVAFKEILCGKKAGSYYNSGFARDQSSFMRNDSSFRKETTT-T-----HLGGS-     382
virilis            VMSQRYRVAFKEILCGKKAGAYYNSGFARDQSSFIRDESSFRRGSSATP-----NLRGS-     406
mojavensis         VMSKRYRIAFKEILCGKKAGAFYNSGFARDQSSFMRDETSFRRNGSTNN-----NNLS--     401

bipectinata        QPNIIKEKDSPSQNRVLIKKTYSLPLPKTLDSTVLS-TTDIVIVLENNSIARNPVCEESK     470
anannassae         QPNIVKETDSPSQNRVLIKKTYSLPLPKTLDSAVLS-TTDIVIVLENNSTARHPVAEESK     471
serrata            HQNNKFETDSISSNRVLIKKTYSLPLPKNADSTVLS-TTDIVIVLENNSTARHTDAEEPK     476
kikkawei           HQNNKYETDSISSNRVLIKKTYSLPLPKNADSTVLS-TTDIVIVLENNSTNRQTDAEEPK     476
erecta             HPN-KFETDSLSANRLLIKKTYSLPLPKNADSTVLS-TTDIVIVLEN----SHTVCEEPK     463
melanogaster       HPN-KFETDSSSANRVLIKKTYSLPLPKNADSTVLS-TTDIVIVLEN----SHTVCEEPK     463
sechelia           HPN-KFETDSPSANRVLIKKTYSLPLPKNADSTVLS-TTDIVIVLEN----SHTVCEEPK     449
simulans           HPN-KFETDSPSANRVLIKKTYSLPLPKNADSTVLS-TTDIVIVLEN----SHKVCEEPK     464
mauritania         HPN-KFETDSPSANRVLIKKTYSLPLPKNADSTVLS-TTDIVIVLEN----SHTVCEEPK     464
ficusphila         HQSSKNEADPLSANRVLIKKTYSLPLPKNSDSAVLS-TTDIVIVLEKNSTARHTICEEPK     476
takahashi          HQNNKNEADSLSANRVLIKKTYSLPLPKNADSTVLS-TTDIVIVLEN----SHTVCEEPK     471
eugracilis         HQNIKNEAESLSANRVLIKKTYSLPLPKNADSTVLS-TTDIVIVLEN----SHTVCEEPK     472
biarmpes           HQNNRNDADSLSANRVLIKKTYSLPLPKNADSTVLS-TTDIVIVLEN----SHTVCEEPK     472
suzuki             HQTNKNDADCLSANRVLIKKTYSLPLPKNADSTVLS-TTDIVIVLEN----SHTVCEEPK     472
rhopaloa           QQNNKNEADS--ANRVLIKKTYSLPLPKNVDSTVLS--TDIVIVLENSSTARHTLCEEPK     473
elegans            HQNNKNEADSISASRMLIKKTYSLPLPKNADSTVLS--TDIVIVLENSTSARHTVCEGAQ     474
grimshawi          ----------KRNSRLH--------PNGMTDYPLLST-TNIVIVLGNSSSRGHQKEINRD     422
virilis            ----------TRYSRVS--------ANGMADCSLLNTTTKIVIVLGNNS---PQRDVDRN     444
mojavensis         ----------SSNVRYN--------RNRVSDCSLLST-TKIVIVLGNR------REANHN     435

bipectinata        VSNDIWIENEETCI      484
```

| | | |
|---|---|---|
| anannassae | VTNDIWIENEETCI | 485 |
| serrata | VDNDIWIEHEETSI | 490 |
| kikkawei | VDNEIWIEHEETSI | 490 |
| erecta | VEKDIWSENEETCI | 477 |
| melanogaster | VENDIWIENEETCI | 477 |
| sechelia | VENDIWIDNEETCI | 463 |
| simulans | VENDIWIENEETCI | 478 |
| mauritania | VENDIWIENEETCI | 478 |
| ficusphila | VENDIWIENQETCI | 490 |
| takahashi | VENDIWIENEETCI | 485 |
| eugracilis | VENDIWIENEETCI | 486 |
| biarmpes | VENDIWIENEETCI | 486 |
| suzuki | VENDIWIENEETCI | 486 |
| rhopaloa | VENDIWIENEETCI | 487 |
| elegans | VENDIWIKNEETCI | 488 |
| grimshawi | IPEEIGIKKEEET- | 435 |
| virilis | IPEETEAKKQEN-- | 456 |
| mojavensis | IPEETDIKKEEP-- | 447 |

Melanogaster NP_996140.1

```
  1 mnsstdptfs elnasftntp dtlfatsvss dpshgfgeed yacgtfncsp kefvafvlgp
     61 qtlplykavl itiifggifi tgvvgnllvc iviirhsamh tatnyylfsl avsdllyllf
    121 glptevflyw hqypdlfgmp fckirafise actyvsvfti vafsmerfla ichplhlyam
    181 vgfkrairii talwivsfis aipfgllsdi qylnypldhs rieesafcsm spkivneipv
    241 fevsfciffv ipmiliilly grmgakirsr tnqklgvqqg tnnretrnsq mrkktvirml
    301 aavvitffvc wfpfhlqrli flyaknmdny ldinealfsi agfayyvsct vnpivysvms
    361 rryrvafrel lcgkavgayy nsgfardhss fressaydrv hsvhvrasqh pnkfetdsss
    421 anrvlikkty slplpknads tvlsttdivi vlenshtvce epkvendiwi eneetci
```

Erecta **XP_001973649.1**

```
    1 mnsstdstfs dlntsftntp dlyttsvsld pshgleeevh yacgtlncsp kefvafvlgp
     61 qtlplykail itiifggifi tgvvgnllvc iviirhsamh tatnyylfsl avsdllyllf
    121 glptevflyw hqypylfgmp fckirafise actyvsvfti vafsmerfla ichplhlyam
    181 vgfkrairii talwiasfis aipfgllsdi qyinypldkv rieesafcsm sikivneipv
    241 fevsfciffv ipmiliivly grmgakirsr tnqklgvqhg tnnretrnsq mkkktvirml
    301 aavvitffvc wfpfhlqrli flyatnmdny ldinealfsi agfayyvsct vnpivysvms
    361 rryrvafrel lcgkavgayy nsgfardhss fressaydrv hsvhvrasrh pnkfetdsls
    421 anrllikkty slplpknads tvlsttdivi vlenshtvce epkvekdiws eneetci
```

Biarmpes **XP_016959554.1**

```
1 mnsstdptfl elntsfsstp dlyatsvsle pshgfeaedy yacgtfnctp teyvafvlgp
     61 qtlplykavl itiifggifi tgvignllvc iviirhsamh tatnyylfsl avsdllyllf
    121 glptevflyw hqypdlfgmp fckirafise actyvsvfti vafsmerfla ichplhlyam
    181 vgfkrairii talwiasfis aipfgllsdi qylrypldgs rieesafcsm speivnvfpv
    241 fevsfcvffv ipmiliilly grmgakirsr tnqklgvqqg tnsresrssq mrkktvirml
    301 aavvitffvc wfpfhlqrli flyakdmeny ldinealfsi agfayyvsct vnpivysvms
    361 rryrvafrel lcgkpvgayy nsgfardhss frestatslg nninydrvhs vhvtssrhqn
    421 nrndadslsa nrvlikktys lplpknadst vlsttdiviv lenshtvcee pkvendiwie
    481 neetci
```

Suzuki **XP_036672159.1**

```
1 mnsstdsnfl elntsfsnkp dlyptsvsle pshgfetedy yacgtfnctp tefvafvlgp
     61 qtlplykail itiifggifi tgvignllvc iviirhsamh tatnyylfsl avsdllyllf
    121 glptevflyw hqypdlfgmp fckirafise actyvsvfti vafsmerfla ichplhlyam
    181 vgfkrairii talwiasfis aipfgllsdi qylkypldgs rieesafcsm speivnvipv
    241 fevsfciffv ipmiliilly grmgakirsr tnqklgvqpg tnnretrnsq mrkktvirml
    301 aavvitffvc wfpfhlqrli flyaknmeny ldinealfsi agfayyisct vnpivysvms
    361 rryrvafrel lcgkavgayy nsgfardhss frestatslg nninydrvhs vhvrssrhqt
    421 nkndadclsa nrvlikktys lplpknadst vlsttdiviv lenshtvcee pkvendiwie
    481 neetci
```

Sechelia **XP_002040831.1**

```
  1 mnsstdlfat svsldttsyg feeedhyacg tfncspkefv afvlgpqtlp lykailitii
     61 fggifitgvv gnllvcivii rhsamhtatn yylfslavsd llyllfglpt evflywhqyp
    121 dlfgmpfcki rafiseacty vsvftivafs merflaichp lhlyamvgfk rairiitalw
    181 iasfisaipf gllsdiqyln ypl ddsriee safcsmspki vneypvfevs fciffvipmi
    241 liillygrmg akirsrtnqk lgvqpgtnnr etrnsqmrkk tvirmlaavv itffvcwfpf
    301 hlqrliflya knmenyldin ealfsiagfa yyvsctvnpi vysvmsrryr vafrellcgk
    361 avgayynsgf ardhssfres saydrvhsvh vrssrhpnkf etdspsanrv likktyslpl
    421 pknadstvls ttdivivlen shtvceepkv endiwidnee tci
```

Simulans **XP_016032860.1**

```
1 mnsstdptfs elntsftntt dlfatsvsld ttshgfeeed hyacgtfncs pkefvafvlg
     61 pqtlplykai litiifggif itgvvgnllv civiirhsam htatnyylfs lavsdllyll
    121 fglptevfly whqypdlfgm pfckirafis eactyvsvft ivafsmerfl aichplhlya
    181 mvgfkrairi italwiasfi saipfgllsd iqylnypldd srieesafcs mspkivneyp
    241 vfevsfciff vipmiliill ygrmgakirs rtnqklgvqp gtnnretrns qmrkktvirm
    301 laavvitffv cwfpfhlqrl iflyaknmen yldinealfs iagfayyvsc tvnpivysvm
    361 srryrvafre llcgkavgay ynsgfardhs sfressaydr vhsvhvrasr hpnkfetdsp
    421 sanrvlikkt yslplpknad stvlsttdiv ivlenshkvc eepkvendiw ieneetci
```

Takahashi **XP_017001909.2**

```
1 mnsstdptfl dlntsfsntp elyatsvsse pshgleaedy yacgtfnctp kefvafvlgp
     61 qtlplykail itiifggifi tgvignllvc iviirhsamh tatnyylfsl avsdllyllf
    121 glptevflyw hqypdlfgmp fckirafise actyvsvfti vafsmerfla ichplhlyam
    181 vgfkraikii tglwivsfis avpfgllsdi qylkypidgs rieesafcsm seivnvyplf
    241 evsfsiffvi pmiliiillyg rmgakirsrt nqklgvqhgt nnretrnsqm rkkavirmla
    301 avvitffvcw fpfhlqrlif lyakdmenyy dinealfsia gfayyvscti npivysvmsr
    361 ryrvafrell cgkalgayyn safardhssf restatslgn ninydrvhsv hvkssrhqnn
    421 kneadslsan rvlikktysl plpknadstv lsttdivivl enshtvceep kvendiwien
    481 eetci
```

Bipectinate **XP_017101267.1**

```
  1 mnstteisfl dlnssiisds fgtssfpvye seteyacgtf nctptefvef vlgpqtlqlw
     61 kailitiifg gifitgiign llvcivivrh samhtatnyy lfslavsdll yllglptev
    121 flfwhqypyl fglpfckira fiseactyvs vftiaafsme rflaichplh lyamvgfkra
    181 lriiallwva sfisaipfgv lsdiiyltyp ldnstiaesa fcsmspeivn vvpifelsfc
    241 iffvipmili lllygrmglk irsrtnqklg vqhgninres rnsqmrkktv irmlaavvit
    301 ffvcwfpfhl qrlwflyaqn nenyynvnew lfsiagfayy vsctinpivy svmsrryrva
    361 frellcgrpv gayynsgfar dhssfresta tsmgnhinyd rvhsvhvras rqpniikekd
    421 spsqnrvlik ktyslplpkt ldstvlsttd ivivlennsi arnpvceesk vsndiwiene
    481 etci
```

Rhopaloa **XP_016979869.1**

```
  1 mnsstdpsvl dlntslssst nsyatsvsle pfqgldsedy yacgtfncsp tefvafvlgp
     61 qtlplykavl itiifggifi tgvignllvc iviirhsamh tatnyylfsl avsdllyllf
    121 glptevflyw hqypdlfgmp fckirafise actyvsvfti vafsmerfla ichplhlyam
    181 vgfkrairii talwiasfis aipfgllsdi qylkypldds rieesafcsm speivnvipv
    241 fevsfciffv ipmilimily grmgakirsr tnqklgvqhg tnnretrnsq mrkkavirml
    301 aavvitffvc wfpfhlqrli flyaknleny ldinealfsi agfayyvsct inpivysvms
    361 rryrvafrel lcgkavgayy nsafardhss fresthtslg nninydrvhs vhvrssrqqn
    421 nkneadsanr vlikktyslp lpknvdstvl stdivivlen sstarhtlce epkvendiwi
    481 eneetci
```

Eugracilis **XP_017072282.1**
```
1 mnsstnptfp efntsfsntp dlyatsvsve ptngletdgy yacgtfncsp tefvafvlgp
     61 qtlplykail itiifggifi tgvignllvc iviirhsamh tatnyylfsl avsdllyllf
    121 glptevflyw hqypdlfgmp fckirafise actyvsvfti vafsmerfla ichplhlyam
    181 vgfkrairii talwiasfis aipfgllsdi qylkypldgs iiqesafcsm spekvnvipv
    241 fevsfciffv ipmiliilly grmgakirsr tnqklgvqhg tnnretrnsq mrkktvirml
    301 aavvitffvc wfpfhlqrli flyaknmeny ldinealfsi agfayyvsct inpivysvms
    361 rryrvafrel lcgkavgayy nsgfardhss frestgtslg nninydrvhs vhvrssrhqn
    421 ikneaeslsa nrvlikktys lplpknadst vlsttdiviv lenshtvcee pkvendiwie
    481 neetci
```

Ficusphila **XP_017042441.1**
```
1 mslstdptll dlntsfstip rlyatsvsle ppndleaedn yacgtfncsp tefvafvlgp
     61 qtlplykail itiifggvfi tgvignllvc iviirhsamh tatnyylfsl avsdllyllf
    121 glptevflyw hqypdlfgmp fckirafise actyvsvfti vafsmerfla ichplhlyam
    181 vgfkrairii talwiasfis aipfgllsdi qylkyppdds rieesafcsm speivnvipv
    241 fevsfciffv ipmiliilly grmgakirsr tnqklgvqhg tnnretrnsq mrkktvirml
    301 aavvitffvc wfpfhlqrli flyakdldny ldinealfsi agfayyvsct vnpivysvms
    361 rryrvafrel lcgkpvgayy nsgfardhss frdstatsmg nnihydrvhs vhvrssrhqs
    421 skneadplsa nrvlikktys lplpknsdsa vlsttdiviv leknstarht iceepkvend
    481 iwienqetci
```
Anannassae **XP_001958016.1**
```
1 mnstteisfl dlnssiisds igtssfpaye seteyyacvt fnctptefve fvlgpqtlql
     61 wkailitivf ggifltgtig nllvcivivr hssmhtatny ylfslavsdl lylllglpte
    121 vflfwhqypy lfglpfckir afiseactyv svftivafsm erflaichpl hlyamvgfkr
    181 alriiallwv asfisaipfg vwseiiylty pldnstiees afcsmtpeiv nlvpifelsf
    241 ciffvipmil iillygrmgl kirsrtnkkl gvqhgninre srnsqmkkka virmlaavvi
    301 tffvcwfpfh mqrlwflyak dnenyynvne wlfsiagfay yvsctinpiv ysvmsrryrv
    361 afrellcgrp vgayynsgfa rdhssfrest atsmgnhiny drvhsvhvra srqpnivket
    421 dspsqnrvli kktyslplpk tldsavlstt divivlenns tarhpvaees kvtndiwien
    481 eetci
```
Serrata **XP_020816118.1**
```
1 mnsstdssfl elntnlsntp dlyatsvnea psygsgteey yacvtfncse sefvafvlgp
     61 qtlplykail itiifggvfi tgvignllvc tviirhsamh tatnyylfsl avsdllyllf
    121 glptevflyw hqypylfgmp fckirafise actyvsvfti vafsmerfla ichplhlyam
    181 vgfkrairii talwiasfis aipfgllsdi qylqypldds rieesafcsm stqtvtmipv
    241 fevsfciffv ipmiliiily grmgakirsr tnqklgvqhg tnnretrnsq lrkktvirml
    301 aavvitffvc wfpfhvqrlw flyaqendny ldinealfsi agfayyvsct inpivysvms
    361 rryrvafrel lcgrpvgayy nsgfardhss frestatvlg knvnydrvhs vhvrssrhqn
    421 nkfetdsiss nrvlikktys lplpknadst vlsttdiviv lennstarht daeepkvdnd
    481 iwieheetsi
```
Kikkawei **XP_017034448.1**
```
1 mnsstdssfl elntnlsntp dlyatsvnea psygsgteey yacvtfncse sefvafvlgp
     61 qtlplykail itiifggvfi tgvignllvc iviirhsamh tatnyylfsl avsdllyllf
    121 glptevflyw hqypylfgmp fckirafise actyvsvfti vafsmerfla ichplhlyam
    181 vgfkrairii talwiasfis aipfgllsdi qylqypldds rieesafcsm stqtvtmipv
    241 fevsfciffv ipmilimily grmgakirsr tnqklgvqhg tnnretrnsq lrkktvirml
    301 aavvitffvc wfpfhvqrlw flyaqendny ldinealfsi agfayyvsct inpivysvms
    361 rryrvafrel lcgrpagayy nsgfardhss frestattlg knvnydrvhs vhvrssrhqn
    421 nkyetdsiss nrvlikktys lplpknadst vlsttdiviv lennstnrqt daeepkvdne
    481 iwieheetsi
```

Mauritania **XP_033158765.1**

1 mnsstdptfs elntsftntt dlfatsvsld ttshgfeeed hyacgtfncs pkefvafvlg
61 pqtlplykai litiifggif itgvvgnllv civiirhsam htatnyylfs lavsdllyll
121 fglptevfly whqypdlfgm pfckirafis eactyvsvft ivafsmerfl aichplhlya
181 mvgfkrairi italwiasfi saipfgllsd iqylnfpldd srieesafcs mspkivneyp
241 vfevsfciff vipmiliill ygrmgakirs rtnqklgvqp gtnnretrns qmrkktvirm
301 laavvitffv cwfpfhlqrl iflyaknmen yldinealfs iagfayyvsc tvnpivysvm
361 srryrvafre llcgkavgay ynsgfardhs sfressaydr vhsvhvrasr hpnkfetdsp
421 sanrvlikkt yslplpknad stvlsttdiv ivlenshtvc eepkvendiw ieneetci

Elegans **XP_017128680.1**

1 mnssteatfl dfntslsntp tsyatsvswa psqgleaedy facgtfncsp tefvafvlgp
61 qtlplykail itiifggifi tgvignllvc iviirhsamh tatnyylfsl avsdllyllf
121 glptevflyw hqypdlfgmp fckirafise actyvsvfti vafsmerfla ichplhlyam
181 vgfkrairii talwiasfis aipfgllsdi qylkypldds rieesafcsm speivnvipv
241 fevsfciffv ipmiliilly grmgakirsr tnqklgvqhg nnretrnsqm rkktvirmla
301 avvitffvcw fpfhlqrlif lyakdmdnyl dinealfsia gfayyvscti npivysvmsr
361 ryrvafrell cgkavgayyn safardhssf restattmgn ninydrvhsv hvrssrhqnn
421 kneadsisas rmlikktysl plpknadstv lstdivivle nstsarhtvc egaqvendiw
481 ikneetci

Grimshawi **XP_001984893.2**

1 mnestyfeld dlgcpqinct kmeftqfilg pqtlplhkav misiifggif itgvlgnvlv
61 cmviirhaam htatnyylfs lavsdliyll lglpievfly whqypflfgl pfcklrafis
121 eactyvsvft ivafsmerfl aichplhvca msgfqralri ttilwivsfl iaipfgikte
181 iqylnypidg slitesafca ielefpekfp lfegsfciff iipmvliiil ygrmgaqirs
241 ratdrlgvqq asrnqatrns qkkkravirm laavvvtffv cwfpfhlqrl wflyaknndn
301 yqdvnewlfs iagfayyvsc tinpivynvm shryrvafke ilcgkkagsy ynsgfardqs
361 sfmrndssfr ketttthlgg skrnsrlhpn gmtdypllst tnivivlgns ssrghqkein
421 rdipeeigik keeet

Virilis

1 mnmnmstnms mdtnlstylg tsdatalpyp gmddygcphm nctamefvqf vlgpqtlplh
61 kallisiifs gifitgvlgn vlvcmviirh aamhtatnyy lfslavsdll ylllglpaev
121 flywhqypyl fglpfcklra fvseactyvs vftivafsme rflaichplh vcamsgfqra
181 lriitalwiv sflsaipfgv kteiqylnfp ndgsrilesa fcsielefpe efplfevsfc
241 iffiipmili illygrmgaq irsratdklgvqqgsrnres rssqkkkrav irmlaavvit
301 ffvcwfpfhl qrlwflyakn ianyqdvnew lfsiagfayy vsctinpivy nvmsqryrva
361 fkeilcgkka gayynsgfar dqssfirdes sfrrgssatp nlrgs***trysr vs***angmadcs
421 llntttkivi vlgnnspqrd vdrnipeete akkqen

Mojavensis **XP_032587092.1**

1 mnmnsslnvn lstqlgtmda taaalpdidd ygcpnlnctp meftqfilgp qtlplhkall
61 isiifsgifi tgvlgnvlvc mviirhaamh tatnyylfsl avsdllylll glptevflyw
121 hqypflfglq fcklrafvse actyvsvfti vafsmerfla icyplhvcam sgfqralrii
181 tvlwivsflt aipfgvktei qylnypidgs rilesafcsm esefpdkypl fegsfiiffi
241 ipmilifvly grmgaqirsr aadqlgvqqg srnresrssq kkkravirml aavvitffvc
301 wfpfhlqrlw flyaknfacf qnvnewlfsi agfayyvsct inpivynvms kryriafkei
361 lcgkkagafy nsgfardqss fmrdetsfrr ngstnnnnls ssnvrynrnr vsdcsllstt
421 kivivlgnrr eanhnipeet dikkeep

CLUSTAL

```
bipectinata       RMLAAVVITFFVCWFPFHLQRLWFLYAQNNENYYNVNEWLFSIAGFAYYVSCTINPIVYS     351
anannassae        RMLAAVVITFFVCWFPFHMQRLWFLYAKDNENYYNVNEWLFSIAGFAYYVSCTINPIVYS     352
serrata           RMLAAVVITFFVCWFPFHVQRLWFLYAQENDNYLDINEALFSIAGFAYYVSCTINPIVYS     357
kikkawei          RMLAAVVITFFVCWFPFHVQRLWFLYAQENDNYLDINEALFSIAGFAYYVSCTINPIVYS     357
erecta            RMLAAVVITFFVCWFPFHLQRLIFLYATNMDNYLDINEALFSIAGFAYYVSCTVNPIVYS     357
melanogaster      RMLAAVVITFFVCWFPFHLQRLIFLYAKNMDNYLDINEALFSIAGFAYYVSCTVNPIVYS     357
sechelia          RMLAAVVITFFVCWFPFHLQRLIFLYAKNMENYLDINEALFSIAGFAYYVSCTVNPIVYS     343
simulans          RMLAAVVITFFVCWFPFHLQRLIFLYAKNMENYLDINEALFSIAGFAYYVSCTVNPIVYS     358
mauritania        RMLAAVVITFFVCWFPFHLQRLIFLYAKNMENYLDINEALFSIAGFAYYVSCTVNPIVYS     358
ficusphila        RMLAAVVITFFVCWFPFHLQRLIFLYAKDLDNYLDINEALFSIAGFAYYVSCTVNPIVYS     357
takahashi         RMLAAVVITFFVCWFPFHLQRLIFLYAKDMENYYDINEALFSIAGFAYYVSCTINPIVYS     356
eugracilis        RMLAAVVITFFVCWFPFHLQRLIFLYAKNMENYLDINEALFSIAGFAYYVSCTINPIVYS     357
biarmpes          RMLAAVVITFFVCWFPFHLQRLIFLYAKDMENYLDINEALFSIAGFAYYVSCTVNPIVYS     357
suzuki            RMLAAVVITFFVCWFPFHLQRLIFLYAKNMENYLDINEALFSIAGFAYYISCTVNPIVYS     357
rhopaloa          RMLAAVVITFFVCWFPFHLQRLIFLYAKNLENYLDINEALFSIAGFAYYVSCTINPIVYS     357
elegans           RMLAAVVITFFVCWFPFHLQRLIFLYAKDMDNYLDINEALFSIAGFAYYVSCTINPIVYS     356
grimshawi         RMLAAVVVTFFVCWFPFHLQRLWFLYAKNNDNYQDVNEWLFSIAGFAYYVSCTINPIVYN     328
virilis           RMLAAVVITFFVCWFPFHLQRLWFLYAKNIANYQDVNEWLFSIAGFAYYVSCTINPIVYN     351
mojavensis        RMLAAVVITFFVCWFPFHLQRLWFLYAKNFACFQNVNEWLFSIAGFAYYVSCTINPIVYN     347
```

| | | |
|---|---|---|
| bipectinata | VMSRRYRVAFRELLCGRPVGAYYNSGFARDHSSFRESTATSMGNHINYDRVHSVHVRASR | 411 |
| anannassae | VMSRRYRVAFRELLCGRPVGAYYNSGFARDHSSFRESTATSMGNHINYDRVHSVHVRASR | 412 |
| serrata | VMSRRYRVAFRELLCGRPVGAYYNSGFARDHSSFRESTATVLGKNVNYDRVHSVHVRSSR | 417 |
| kikkawei | VMSRRYRVAFRELLCGRPAGAYYNSGFARDHSSFRESTATTLGKNVNYDRVHSVHVRSSR | 417 |
| erecta | VMSRRYRVAFRELLCGKAVGAYYNSGFARDHSSFRESS--------AYDRVHSVHVRASR | 409 |
| melanogaster | VMSRRYRVAFRELLCGKAVGAYYNSGFARDHSSFRESS--------AYDRVHSVHVRASQ | 409 |
| sechelia | VMSRRYRVAFRELLCGKAVGAYYNSGFARDHSSFRESS--------AYDRVHSVHVRSSR | 395 |
| simulans | VMSRRYRVAFRELLCGKAVGAYYNSGFARDHSSFRESS--------AYDRVHSVHVRASR | 410 |
| mauritania | VMSRRYRVAFRELLCGKAVGAYYNSGFARDHSSFRESS--------AYDRVHSVHVRASR | 410 |
| ficusphila | VMSRRYRVAFRELLCGKPVGAYYNSGFARDHSSFRDSTATSMGNNIHYDRVHSVHVRSSR | 417 |
| takahashi | VMSRRYRVAFRELLCGKALGAYYNSAFARDHSSFRESTATSLGNNINYDRVHSVHVKSSR | 416 |
| eugracilis | VMSRRYRVAFRELLCGKAVGAYYNSGFARDHSSFRESTGTSLGNNINYDRVHSVHVRSSR | 417 |
| biarmpes | VMSRRYRVAFRELLCGKPVGAYYNSGFARDHSSFRESTATSLGNNINYDRVHSVHVTSSR | 417 |
| suzuki | VMSRRYRVAFRELLCGKAVGAYYNSGFARDHSSFRESTATSLGNNINYDRVHSVHVRSSR | 417 |
| rhopaloa | VMSRRYRVAFRELLCGKAVGAYYNSAFARDHSSFRESTHTSLGNNINYDRVHSVHVRSSR | 417 |
| elegans | VMSRRYRVAFRELLCGKAVGAYYNSAFARDHSSFRESTATTMGNNINYDRVHSVHVRSSR | 416 |
| grimshawi | VMSHRYRVAFKEILCGKKAGSYYNSGFARDQSSFMRNDSSFRKETTT-T-----HLGGS- | 382 |
| virilis | VMSQRYRVAFKEILCGKKAGAYYNSGFARDQSSFIRDESSFRRGSSATP-----NLRGS- | 406 |
| mojavensis | VMSKRYRIAFKEILCGKKAGAFYNSGFARDQSSFMRDETSFRRNGSTNN-----NNLS-- | 401 |

| | | |
|---|---|---|
| bipectinata | QPNIIKEKDSPSQNRVLIKKTYSLPLPKTLDS**TVLS-TTD**IVIVLENNSIARNPVCEESK | 470 |
| anannassae | QPNIVKETDSPSQNRVLIKKTYSLPLPKTLDSAVLS-TTDIVIVLENNSTARHPVAEESK | 471 |
| serrata | HQNNKFE**TDSISS**NRVLIKKTYSLPLPKNADS**TVLS-TTD**IVIVLENNSTARHTDAEEPK | 476 |
| kikkawei | HQNNKYE**TDSISS**NRVLIKKTYSLPLPKNADS**TVLS-TTD**IVIVLENNSTNRQTDAEEPK | 476 |
| erecta | HPN-KFETDSLSANRLLIKKTYSLPLPKNADS**TVLS-TTD**IVIVLEN----**SHTVCE**EPK | 463 |
| melanogaster | HPN-KFETDSSSANRVLIKKTYSLPLPKNADS**TVLS-TTD**IVIVLEN----**SHTVCE**EPK | 463 |
| sechelia | HPN-KFETDSPSANRVLIKKTYSLPLPKNADS**TVLS-TTD**IVIVLEN----**SHTVCE**EPK | 449 |
| simulans | HPN-KFETDSPSANRVLIKKTYSLPLPKNADS**TVLS-TTD**IVIVLEN----**SHKVCE**EPK | 464 |
| mauritania | HPN-KFETDSPSANRVLIKKTYSLPLPKNADS**TVLS-TTD**IVIVLEN----**SHTVCE**EPK | 464 |
| ficusphila | HQSSKNEADPLSANRVLIKKTYSLPLPKNSDS**AVLS-TTD**IVIVLEKNSTARHTICEEPK | 476 |
| takahashi | HQNNKNEADSLSANRVLIKKTYSLPLPKNADS**TVLS-TTD**IVIVLEN----**SHTVCE**EPK | 471 |
| eugracilis | HQNIKNEAESLSANRVLIKKTYSLPLPKNADS**TVLS-TTD**IVIVLEN----**SHTVCE**EPK | 472 |
| biarmpes | HQNNRNDADSLSANRVLIKKTYSLPLPKNADS**TVLS-TTD**IVIVLEN----**SHTVCE**EPK | 472 |
| suzuki | HQTNKNDADCLSANRVLIKKTYSLPLPKNADS**TVLS-TTD**IVIVLEN----**SHTVCE**EPK | 472 |
| rhopaloa | QQNNKNEADS--ANRVLIKKTYSLPLPKNVDSTVLS--TDIVIVLENSSTARHTLCEEPK | 473 |
| elegans | HQNNKNEADSISASRMLIKKTYSLPLPKNADSTVLS--TDIVIVLENSTSARHTVCEGAQ | 474 |
| grimshawi | ----------KRNSRLH--------PNGMTDYPLLST-TNIVIVLGNSSSRGHQKEINRD | 422 |
| virilis | ----------**TRYSRVS**--------ANGMADCSLLNTTTKIVIVLGNNS---PQRDVDRN | 444 |
| mojavensis | ----------SSNVRYN--------RNRV**SDCSLLS**T-TKIVIVLGNR------REANHN | 435 |

| | | |
|---|---|---|
| bipectinata | VSNDIWIENEETCI | 484 |
| anannassae | VTNDIWIENEETCI | 485 |
| serrata | VDNDIWIEHEETSI | 490 |
| kikkawei | VDNEIWIEHEETSI | 490 |
| erecta | VEKDIWSENEETCI | 477 |
| melanogaster | VENDIWIENEETCI | 477 |
| sechelia | VENDIWIDNEETCI | 463 |
| simulans | VENDIWIENEETCI | 478 |
| mauritania | VENDIWIENEETCI | 478 |
| ficusphila | VENDIWIENQETCI | 490 |
| takahashi | VENDIWIENEETCI | 485 |
| eugracilis | VENDIWIENEETCI | 486 |
| biarmpes | VENDIWIENEETCI | 486 |
| suzuki | VENDIWIENEETCI | 486 |
| rhopaloa | VENDIWIENEETCI | 487 |
| elegans | VENDIWIKNEETCI | 488 |
| grimshawi | IPEEIGIKKEEET- | 435 |
| virilis | IPEETEAKKQEN-- | 456 |
| mojavensis | IPEETDIKKEEP-- | 447 |
