## Supplementary material for "The incidence of candidate binding sites for β-arrestin in Drosophila neuropeptide GPCRs": S12 Text

***S12. Text*** **Multi-species analysis of PDF R PA isoforms**
**Supporting Figure 15**

CLUSTAL Line-ups; Genbank Reference IDs below
6th & 7th Predicted TM domain in YELLOW
BBS sequences in RED

```
Serrata           -------------LQLQYAFLGDMTLLSSILDSGG--------GISAQRFTRLLRQSSTS        39
Kikkawei          -------------MQLQYAFAGDMTLLSSILDSGG--------SISAQRFTRLLRQSSTS        39
Bipectinate       ----------------HYADTGNMTLLSSILDSGG--------SVSVQRLTRLLRQSSPS        36
Anannassae        -----------------------MTLLSSILDSGG--------SVSVQRLTRLLRQSSTS        29
Fichsuphila       -----------------------MTLLASILDCGG--------GISAQRFTRLLAQSSLL        29
Rhopaloa          -----------------------MTLLSNILDCGG--------GISAQCLTRSLRESSSP        29
Elegans           MKVAFTSQMNNENNHFHNVVDGEMTLLSNILDCGG--------GIPAQRLARLLRQSSSS        52
Eugracilis        -----------------------MTLLSNILDSGG--------GISAQRFTRLLRQSSSS        29
Erecta            -----------------------MTLLSNILDCGG--------CISAQRFTRLLRQSGSS        29
Melanogaster      -----------------------MTLLSNILDCGG--------CISAQRFTRLLRQSGSS        29
Sechellia         -----------------------MTLLSNILDCGG--------CISDQRFTHLLRQSGSS        29
Simulans          -----------------------MTLLSNILDCGG--------CISAQRFTRLLRQSGSS        29
Mauritania        -----------------------MTLLSNILDCGG--------CISAQRFTRLLRQSGSS        29
Takahashi         -----------------------MTLLSNILDCGG--------GISAQRFARLLRQSSSS        29
Suzuki            -----------------------MTLLSNILDCGG--------GISAQRFTRLLRQSSSS        29
Biarmipes         -----------------------MTLLSNILDCGG--------GISAQRFTRLLRQSSSP        29
Mojavensis        ------------------------MQLAGDSEAGSMPTIITTSTSSNSFLQRIALKAVTA        36
Grimshawii        -----------------------MMQLPGINKPGCIPTISTSSIIIPQ---RFARRAARV        34
Virilis          ------------------------------------------------------------         0


Serrata           SSSSSS-----------SSASGSMLEPTSSQLPINDVLGGGRIPFLHDNGTGESL-----        83
Kikkawei          SSS--S-----------VSASASMLEPTSSQLPNNDVLGGGRIPFLPGNATGESSSSSSP        86
Bipectinate       ASASGT-----------ALASESMVEPTASQM-LNEITDGGRVPSL---GQDLAT-----        76
Anannassae        SSASGP-----------SLASDSISSMVE--------ATSLRVPSL---GHDLST-----        62
Fichsuphila       ASSSASASASASASASGTFASESMLEPTSSHI-----LPTGRVPVLHGFDSTT-------        77
Rhopaloa          GS-PG-------SPESGAFESKSMLEPTSSHI-----LPTGRVPVLHDFNASS-------        69
Elegans           GSTPS-------GSASGPFESKSMLEPTTSHI-----LPTGRVPILHDFGSSS-------        93
Eugracilis        GSSSASAS----ASSYGTLESKSMLEPTSSHT-----LPGGRVPILHDFDS---------        71
Erecta            VP---SPS----APAPGTFESISMLEPTSLHS-----LPTGRVPLLHDFDA---------        68
Melanogaster      GP---SPS----APTAGTFESKSMLEPTSSHS-----LATGRVPLLHDFDA---------        68
Sechellia         GP---SPS----APAPGTFESKSMLEPTSPHS-----LATGRVPLLHDFDA---------        68
Simulans          GP---SPS----APAPGTFESKSMLEPTSSHS-----LATGRVPLLHDFDA---------        68
Mauritania        GP---SPS----APAPGTFESKSMLEPTSSHS-----LATGRVPLLHDFDA---------        68
Takahashi         GSPSASASSSASSSSGTSFESKSMLEPTSSQN-----LPTGRVPILHDFDFDFDP-----        79
Suzuki            GSPSS-------SSSGTTFESKSMLEPTSSHI-----LPTGRVPILHDFDFDS-------        70
Biarmipes         GFLSS-------SSLGTTFESKSMLEPTSSHV-----LPTGRVPVLHDFDFDS-------        70
Mojavensis        ATTTKDRL-------------ISIVETD--SPAGSH-M-----DP--STAMTTT--SITA        71
Grimshawii        ASTSTATV---------TTLSMSMTQSG--RSMLNRTM-----TSIATATTTSTSTSIST        78
Virilis          -----------------------MPT-A--APSSNRTV-----STHHSTTMTSTSTTYGT        29
                                         :

Serrata           -----PLPDADALDPNFVLDGVTSVAKMALEATVK-EVLRDPDPEQILANANATAPWNIT        137
Kikkawei          PLPSPSLPDADALDPNFVLDGVTSVAKMALEATVK--VLRDPDPEQILANANASAPWNIT        144
Bipectinate       -------TTETTGQSNVIIDRVTNVAKMALEATVM-DVLPDPDPDQVLAGVNASAHWNMT        128
Anannassae        -------TT----ETNVIIDRVTNVAKMALEATVM-DVLPDPEPDQVLTGVNASSHWNMN        110
Fichsuphila       ---------TESTTSTYVLDGVASVAQMPLLESTVMDALPDSDPDQAFGNLNVTAPWNLT        128
Rhopaloa          --------TDSP--GTFLLDGVVSVAQMALEPTLTD-LLPDPDPDQVLSNLNASAPWNLT        118
Elegans           --------TESPLTGTYLLDGVASVAEMALEPTVRD-ILSDPDPDKVLSNLNASAPWNLT        144
Eugracilis        --------STTESPGAYLLDGVASVAKMALEPAVMD-VLPDPDTDQVLSNLNITAPWNLT        122
Erecta            --------STTESPGTYVLDGVARVAQLALEPTVMD-GLPDPDTEQVLSNLNSSAPWNLT        119
Melanogaster      --------STTESPGTYVLDGVARVAQLALEPTVMD-ALPDSDTEQVLGNLNSSAPWNLT        119
Sechellia         --------STTESPGTYVLDGVARVAQLALEPTVMD-ALPDPDTEQVLGNLNSSAPWNLT        119
Simulans          --------STTESPGTYVLDGVARVAQLALEPTVMD-ALPDPDTEQVLGNLNSSAPWNLT        119
Mauritania        --------TTTESPGTHVLDGVARVAQLALEPTVMD-ALPDPDTEQVLGNLNSNAPWNLT        119
Takahashi         -------TTTTESPGNYVLDGVASVAKMALEPTVMDTVLPDSDPDQVLSNLNVSAPWNLT        132
Suzuki            --------LTTESTGTHVLDGVASVAKMALEPTVMDAVLPDSDTDQVLSNLNISAPWNIT        122
Biarmipes         --------STTELPRILVLDGVAGVATMALEPTAMDAVLPDSDPDQVLSNFNISAPWNIT        122
Mojavensis        TATATPAATATAS--ELSRSDVLNASIEATTAVGMDTVKTATESGLPTVSGTTLPSWNSS        129
Grimshawii        HSTTSSSTSKLPSQVISGLADVINGTELDMTAAATAV---GLDAAVP-DPSTTSSSWNST        134
```

```
Virilis         TSTTQAALTS-----F---------EPMAVTAAGMDV---TMAGTEPLSSTSIPTFWNSS        72
                                                                   .    ** .

Serrata          --LASAAATNYENCSALFANYTLPQTGLYCNWTWDTLLCWPPTPAGVLARMNCPGGYHGV       195
Kikkawei         --LAAAAATNYENCSALFANYTLPQTGLYCNWTWDTLLCWPPTPAGVLARMNCPGGYHGV       202
Bipectinate      --LSSSTATNYENCSALFANYTLPQTGLYCNWTWDTLLCWPPTPAGVLARMHCPGGYHGV       186
Anannassae       MTLSSASATNYENCSALFANYTLPQTGLYCNWTWDTLLCWPPTPAGVLARMNCPGGYHGV       170
Fichsuphila      --LASAAATNFENCSALFANYTLPQSGLYCNWTWDTLLCWPPTPAGVLARMNCPGGFHGV       186
Rhopaloa         --LASAAATNFENCSALFANYTLPQTGLYCNWTWDTLLCWPPTPAGVLARMNCPAGFHGV       176
Elegans          --LASSAATNFENCSALFANYTLPQTGLYCNWTWDTLLCWPPTPAGVLARMNCPGGFHGV       202
Eugracilis       --LASAAATNFENCSALFVNYTLPQTGLYCNWTWDTLLCWPPTPAGVLARMNCPGGFHGV       180
Erecta           --LSSAAATNFENCSALFVNYTLPQTGLYCNWTWDTLLCWPPTPAGVLARMNCPGGFHGV       177
Melanogaster     --LASAAATNFENCSALFVNYTLPQTGLYCNWTWDTLLCWPPTPAGVLARMNCPGGFHGV       177
Sechellia        --LASAAATNFENCSALFVNYTLPQTGLYCNWTWDTLLCWPPTPAGVLARMNCPGGFHGV       177
Simulans         --LASAAATNFENCSALFVNYTLPQTGLYCNWTWDTLLCWPPTPAGVLARMNCPGGFHGV       177
Mauritania       --LASAAATNFENCSALFVNYTLPQTGLYCNWTWDTLLCWPPTPAGVLARMNCPGGFHGV       177
Takahashi        --LASAAATNFENCSALFVNYTLPQTGLYCNWTWDTLLCWPPTPAGVLARMNCPAGFHGV       190
Suzuki           --LASAAATNFENCSSLFVNYTLPQSGLYCNWTWDTLLCWPPTPAGVLARMNCPGGFHGV       180
Biarmipes        --LASAAATNFENCSSLFVNYTLPQTGLYCNWTWDTLLCWPPTPAGVLARMNCPGGFHGV       180
Mojavensis       --FNSALANNYDNCSAMFANYTHPQTGLYCNWTWDSLLCWPPTPAGNLARMNCPAGYHGV       187
Grimshawii       --FSTASANSYDNCAAMFANYTQPVTGLYCNWTWDSLLCWPPTPAGVMARMYCPAGYHGV       192
Virilis         --INTASANSYDNCSALFANYTQPTTVIYCNWTWDSLLCWPPTPAGATAHMHCPAGYHGV        130
                   : :: *..::**:::*.*** * : :*******:**********  *:* **.*:***

Serrata          DTRKFANRKCELDGRWGSRPNATETNATGWTDYGPCYKPEVIRLMQQMGSREDLDLYIEI       255
Kikkawei         DTRKFANRKCELDGRWGSRPNATETNATGWTDYGPCYKPEVIRLMQQMGSREDLDLYIEI       262
Bipectinate      DTRKFANRKCELDGRWGSRPNATEPSPAGWTDYGPCYKPEVIRLMQSMGSK-DIDIYIDI       245
Anannassae       DTRKFANRKCELDGRWGSRPNATEPSPAGWTDYGPCYKPEVIRLMQSMGSK-DIDIYIDI       229
Fichsuphila      DTRKFAIRKCELDGRWGSRPNATEVSPPGWTDYGPCYKPEIIRLMQQMGNK-DFDLYIDI       245
Rhopaloa         DTRKFAIRKCELDGRWGSRPNATEVSPPGWTDYGPCYKPEIIRLMQQMGSR-DFDTFIDI       235
Elegans          DTRKFAIRKCELDGRWGSRPNATEVSPPGWTDYGPCYKPEIIRLMQQMGSK-DFDLYIDI       261
Eugracilis       DTRKFAIRKCELDGRWGSRPNATEASPPGWTDYGPCYKPEIIRLMQQMGNK-DFDLYIDI       239
Erecta           DTRKFAIRKCELDGRWGSRPNATEVSPPGWTDYGPCYKPEIIRLMQQMGSK-DFDAYIDI       236
Melanogaster     DTRKFAIRKCELDGRWGSRPNATEVNPPGWTDYGPCYKPEIIRLMQQMGSK-DFDAYIDI       236
Sechellia        DTRKFANRKCELDGRWGSRPNATEVSPPGWTDYGPCYKPEIIRLMQQMGSK-DFDAYIDI       236
Simulans         DTRKFAIRKCELDGRWGSRPNATEVSPPGWTDYGPCYKPEIIRLMQQMGSK-DFDAYIDI       236
Mauritania       DTRKFAIRKCELDGRWGSRPNATEVSPPGWTDYGPCYKPEIIRLMQQMGSK-DFDAYIDI       236
Takahashi        DTRKFAIRKCELDGRWGSRPNATEVSPPGWTDYGPCYKPEIIRLMEQMGSK-NFDLYIDI       249
Suzuki           DTRKFAIRKCELDGRWGSRPNATEVSPPGWTDYGPCYKPEIIRLMQQMGSK-DFDLYIDI       239
Biarmipes        DTRKFAIRKCELDGRWGSRPNATEVSPPGWTDYGPCYKPEIIRLMQQMGSK-DFDLYIDI       239
Mojavensis       DTRKFANRKCELDGHWGGRPNETEPNHAGWTDYGPCYKPEVIRLMQEIK---DVNLYIDI       244
Grimshawii       DTRKFAIRKCELDGHWGRRPNDTELNTGGWTDYAPCYKPEVIRLMQEIK---DVNVYMDI       249
Virilis         DTRKFANRKCELDGHWAGRPNSTEQKPTGWTDYGPCYKPEVIRLMQEIK---DVNLYMDI        187
                 ****** *******:*. *** ** .  *****.******:****:.:    :.: :::*

Serrata          ARRTRTLEIVGLCLSLFALIVSLLIFCTFRSLRNNRTKIHKNLFVAMVLQVIIRLTLYLD       315
Kikkawei         AKRTRTLEIVGLCLSLFALIVSLLIFCTFRSLRNNRTKIHKNLFIAMVLQVIIRLTLYLD       322
Bipectinate      ARRTRTLEIVGLWVSLFALVISLLIFCTFRSLRNNRTKIHKNLFIAMVLQVIVRLTLYLD       305
Anannassae       ARRTRTLEIVGLWVSLFALVISLLIFCTFRSLRNNRTKIHKNLFVAMVLQVIVRLTLYLD       289
Fichsuphila      ARKTRTLEIVGLCLSLFALIVSLLIFCTFRSLRNNRTKIHKNLFVAMVLQVIIRLTLYLD       305
Rhopaloa         AKKTRTLEIVGLCLSLFALIVSLLIFCTFRSLRNNRTKIHKNLFVAMVLQVIIRLTLYLD       295
Elegans          ARKTRTLEIVGLCLSLFALIVSLLIFCTFRSLRNNRTKIHKNLFVAMVLQVIIRLTLYLD       321
Eugracilis       ARKTRTLEIVGLCLSLFALIVSLLIFCTFRSLRNNRTKIHKNLFVAMVLQVIIRLTLYLD       299
Erecta           ARRTRTLEIVGLCISLFALIVSLLIFCTFRSLRNNRTKIHKNLFVAMVLQVIIRLTLYLD       296
Melanogaster     ARRTRTLEIVGLCLSLFALIVSLLIFCTFRSLRNNRTKIHKNLFVAMVLQVIIRLTLYLD       296
Sechellia        ARRTRTLEIVGLCLSLFALIVSLLIFCTFRSLRNNRTKIHKNLFVAMVLQVIIRLTLYLD       296
Simulans         ARRTRTLEIVGLCLSLFALIVSLLIFCTFRSLRNNRTKIHKNLFVAMVLQVIIRLTLYLD       296
Mauritania       ARRTRTLEIVGLCLSLFALIVSLLIFCTFRSLRNNRTKIHKNLFVAMVLQVIIRLTLYLD       296
Takahashi        AWKTRTLEIVGLCLSLFALIVSLLIFCTFRSLRNNRTKIHKNLFVAMVLQVIIRLTLYLD       309
Suzuki           ARKTRTLEIVGLCLSLFALIVSLLIFCTFRSLRNNRTKIHKNLFVAMVLQVIIRLTLYLD       299
Biarmipes        ARKTRTLEIVGLCLSLFALIVSLLIFCTFRSLRNNRTKIHKNLFVAMVLQVIIRLTLYLD       299
Mojavensis       AQRTRTLEIIGLCLSLLALIISLVIFCAFRSLRNNRTKIHKNLFIAMVLQVIVRLTLYLD       304
Grimshawii       AQRTRTLEIIGLCLSLLALIISLVIFCAFRSLRNNRTKIHKNLFVAMVLQVIVRLTLYLD       309
Virilis         AQRTRTLEIIGLCLSLFALIISLMIFCAFRSLRNNRTKIHKNLFVAMVLQVIVRLTLYLD        247
                 * :******:** :**:**::**:***:****************:*******:*******

Serrata          QFRRGSKEAATNTSLSVIENTPYLCEASYVLLEYARTAMFMWMFIEGLYLHNMVTVAVFQ       375
Kikkawei         QFRRGNKEAATNTSLSVIENTPYLCEASYVLLEYARTAMFMWMFIEGLYLHNMVTVAVFQ       382
Bipectinate      QYRRGNKEAATNTSLSAIENTPYLCEASYVLLEYARTAMFMWMFIEGLYLHNMVTVAVFQ       365
Anannassae       QYRRGNKEAATNTSLSAIENTPYLCEASYVLLEYARTAMFMWMFIEGLYLHNMVTVAVFQ       349
Fichsuphila      QFRRGNKEAATNTSLSVIENTPYLCEASYVLLEYARTAMFMWMFIEGLYLHNMVTVAVFQ       365
```

```
Rhopaloa         QYRRGNKEAATNTSLSVIENTPYLCEASYVLLEYARTAMFMWMFIEGLYLHNMVTVAVFQ        355
Elegans          QYRRGNKEAATNTSLSVIENTPYLCEASYVLLEYARTAMFMWMFIEGLYLHNMVTVAVFQ        381
Eugracilis       QFRRGNKEAATNTSLSVIENTPYLCEASYVLLEYARTAMFMWMFIEGLYLHNMVTVAVFQ        359
Erecta           QFRRGNKEAATNTSLSAIENTPYLCEASYVLLEYARTAMFMWMFIEGLYLHNMVTVAVFQ        356
Melanogaster     QFRRGNKEAATNTSLSVIENTPYLCEASYVLLEYARTAMFMWMFIEGLYLHNMVTVAVFQ        356
Sechellia        QFRRGNKEAATNTSLSVIENTPYLCEASYVLLEYARTAMFMWMFIEGLYLHNMVTVAVFQ        356
Simulans         QFRRGNKEAATNTSLSVIENTPYLCEASYVLLEYARTAMFMWMFIEGLYLHNMVTVAVFQ        356
Mauritania       QFRRGNKEAATNTSLSVIENTPYLCEASYVLLEYARTAMFMWMFIEGLYLHNMVTVAVFQ        356
Takahashi        QYRRGNKEAATNTSLSVIENTPYLCEASYVLLEYARTAMFMWMFIEGLYLHNMVTVAVFQ        369
Suzuki           QFRRGNKEAATNTSLSVIENTPYLCEASYVLLEYARTAMFMWMFIEGLYLHNMVTVAVFQ        359
Biarmipes        QFRRGNKEAATNTSLSVIENTPYLCEASYVLLEYARTAMFMWMFIEGLYLHNMVTVAVFQ        359
Mojavensis       QFRRGIQ---YNSSLSAIENTPYLCEASYVLLEYARTAMFMWMFIEGLYLHNMVTVAVFQ        361
Grimshawii       QFRRGRPETATNASVSIIENTPYLCEASYVLLEYARTAMFMWMFIEGLYLHNMVTVAVFQ        369
Virilis         QFRRGKSDSANNTSLSVIENTPYLCEASYVLLEYARTAMFMWMFIEGLYLHNMITVAVFQ         307
                 *:***      *:*:* ************************************:******

Serrata          GSFPLKFFSRLGWCVPILMTTVWARCTVIYMDTSLGDCLWNYNLTPYYWILEGPRLAVIL        435
Kikkawei         GSFPLKFFSRLGWCVPILMTTVWARCTVIYMDTSLGDCLWNYNLTPYYWILEGPRLAVIL        442
Bipectinate      GSFPLKFFSRLGWCVPILMTFVWARCTVMYMDTSMGECLWNYNLTPYYWILEGPRLAVIL        425
Anannassae       GSFPLKFFSRLGWCVPILMTFVWARCTVMYMDTSMGECLWNYNLTPYYWILEGPRLAVIL        409
Fichsuphila      GSFPLKFFSRLGWCVPILMTTVWARCTVMYMDTSLGECLWNYNLTPYYWILEGPRLAVIL        425
Rhopaloa         GSFPLKFFSRLGWCVPILMTTVWARCTVMYMDTSLGDCLWNYNLTPYYWILEGPRLAVIL        415
Elegans          GSFPLKFFSRLGWCAPILMTTVWARCTVMYMDTSLGECLWNYNLTPYYWILEGPRLAVIL        441
Eugracilis       GSFPLKFFSRLGWCVPILMTTVWARCTVMYMDTTLGECLWNYNLTPYYWILEGPRLAVIL        419
Erecta           GSFPLKFFSRLGWCVPILMTTVWARCTVMYMDTSLGECLWNYNLTPYYWILEGPRLAVIL        416
Melanogaster     GSFPLKFFSRLGWCVPILMTTVWARCTVMYMDTSLGECLWNYNLTPYYWILEGPRLAVIL        416
Sechellia        GSFPLKFFSRLGWCVPILMTTVWARCTVMYMDTSLGECLWNYNLTPYYWILEGPRLAVIL        416
Simulans         GSFPLKFFSRLGWCVPILMTTVWARCTVMYMDTSLGECLWNYNLTPYYWILEGPRLAVIL        416
Mauritania       GSFPLKFFSRLGWCVPILMTTVWARCTVMYMDTSLGECLWNYNLTPYYWILEGPRLAVIL        416
Takahashi        GSFPLKFFSRLGWGFPILMTTVWARCTVMYMDTSLGECLWNYNLTPYYWILEGPRLAVIL        429
Suzuki           GSFPLKFFSRLGWGFPILMTTVWARCTVMYMDTSLGECLWNYNLTPYYWILEGPRLAVIL        419
Biarmipes        GSFPLKFFSRLGWGFPILMTTVWARCTVMYMDTSLGECLWNYNLTPYYWILEGPRLAVIL        419
Mojavensis       GSFPLIFFSLLGWGMPVVMTFVWVQCTAIFMDTALGDCMWNYNLTPYYWILEGPRLAVIL        421
Grimshawii       GNFPLKLFALLGWGLPVLMTFVWVQCTAIFMDTTVGECMWNYNLTPYYWILEGPRLAVIL        429
Virilis         GNFPLVFFSLLGWGMPVLMTFVWVQCTAIFMDTSLGDCLWNYNLTPYYWILEGPRLTVIM         367
                 *.*** :*: ***  *::** **.:**.::***::*:*:*****************:**:

Serrata          LNFCFLVNIIRVLVMKLRQSQASDIEQTRKAVRAAIVLLPLLGITNILHQVAPLKTATNF        495
Kikkawei         LNFCFLVNIIRVLVMKLRQSQASDIEQTRKAVRAAIVLLPLLGITNILHQLAPLKTATNF        502
Bipectinate      LNFCFLVNIIRVLVVKLRQSQASDIEQTRKAVRAAIVLLPLLGITNLLHQLAPLKTATNF        485
Anannassae       LNFCFLVNIIRVLVVKLRQSQASDIEQTRKAVRAAIVLLPLLGITNLLHQLAPLKTATNF        469
Fichsuphila      LNFCFLVNIIRVLVMKLRQSQASDIEQTRKAVRAAIVLLPLLGITNLLHQLAPLKTATNF        485
Rhopaloa         LNFCFLVNIIRVLVMKLRQSQASDIEQTRKAVRAAIVLLPLLGITNILHQMAPLKTATNF        475
Elegans          LNFCFLVNIIRVLVMKLRQSQASDIEQTRKAVRAAIVLLPLLGITNLLHQLAPLKTATNF        501
Eugracilis       LNFCFLVNIIRVLVMKLRQSQASDIEQTRKAVRAAIVLLPLLGITNLLHQLAPLKTATNF        479
Erecta           LNFCFLVNIIRVLVMKLRQSQASDIEQTRKAVRAAIVLLPLLGITNILHQLAPLKTATNF        476
Melanogaster     LNFCFLVNIIRVLVMKLRQSQASDIEQTRKAVRAAIVLLPLLGITNLLHQLAPLKTATNF        476
Sechellia        LNFCFLVNIIRVLVMKLRQSQASDIEQTRKAVRAAIVLLPLLGITNLLHQLAPLKTATNF        476
Simulans         LNFCFLVNIIRVLVMKLRQSQASDIEQTRKAVRAAIVLLPLLGITNLLHQLAPLKTATNF        476
Mauritania       LNFCFLVNIIRVLVMKLRQSQASDIEQTRKAVRAAIVLLPLLGITNLLHQLAPLKTATNF        476
Takahashi        LNFCFLVNIIRVLVMKLRQSQASDIEQTRKAVRAAIVLLPLLGITNILHQMAPLKTATNF        489
Suzuki           LNFCFLVNIIRVLVMKLRQSQASDIEQTRKAVRAAIVLLPLLGITNLLHQLAPLKTATNF        479
Biarmipes        LNFCFLVNIIRVLVMKLRQSQASDIEQTRKAVRAAIVLLPLLGITNLLHQLAPLKTATNF        479
Mojavensis       LNFFFLVNIIRVLVVKLRQSQASDIEQTRKAVRAAIVLLPLLGITNLLHLVPALKTAWKF        481
Grimshawii       LNFFFLVNIIRVLVVKLRQSQASDIEQTRKAVRAAIVLLPLLGITNLLHLVPALKTAWKF        489
Virilis         LNFFFLVNIIRVLVMKLRQSQASEIEQTRKAVRAAIVLLPLLGITNLLHLVPALKTAWKF         427
                 *** **********:********:**********************:** :  **** :*

Serrata          AVWSYSTYFLTSFQGFFIALIYCFLNGEVRAVLLKSLATQLSVRGHPEWAPKRASMYSGA        555
Kikkawei         AVWSYSTYFLTSFQGFFIALIYCFLNGEVRAVLLKSLATQLSVRGHPEWAPKRASMYSGA        562
Bipectinate      AVWSYGTHFLTSFQGFFIALIYCFLNGEVRAVLLKSLATQMSVRGHPEWVPKRASMYSGA        545
Anannassae       AVWSYGTHFLTSFQGFFIALIYCFLNGEVRAVLLKSLATQMSVRGHPEWVPKRASMYSGA        529
Fichsuphila      AVWSYGTHFLTSFQGFFIALIYCFLNGEVRAVLLKSLATQLSVRGHPEWAPKRASMYSGA        545
Rhopaloa         AVWSYGTHFLTSFQGFFIALIYCFLNGEVRAVLLKSLATQLSVRGHPEWAPKRASMYSGA        535
Elegans          AVWSYGTHFLTSFQGFFIALIYCFLNGEVRAVLLKSLATQLSVRGHPEWAPKRASMYSGA        561
Eugracilis       AVWSYGTHFLTSFQGFFIALIYCFLNGEVRAVLLKSLATQLSVRGHPEWAPKRASMYSGA        539
Erecta           AVWSYGTHFLTSFQGFFIALIYCFLNGEVRAVLLKSLATQLSVRGHPEWAPKRASMYSGA        536
Melanogaster     AVWSYGTHFLTSFQGFFIALIYCFLNGEVRAVLLKSLATQLSVRGHPEWAPKRASMYSGA        536
Sechellia        AVWSYGTHFLTSFQGFFIALIYCFLNGEVRAVLLKSLATQLSVRGHPEWAPKRASMYSGA        536
Simulans         AVWSYGTHFLTSFQGFFIALIYCFLNGEVRAVLLKSLATQLSVRGHPEWAPKRASMYSGA        536
Mauritania       AVWSYGTHFLTSFQGFFIALIYCFLNGEVRAVLLKSLATQLSVRGHPEWAPKRASMYSGA        536
```

```
Takahashi         AVWSYVTHFLTSFQGFFIALIYCFLNGEVRAVLLKSLATQLSVRGHPEWAPKRASMYSGA        549
Suzuki            AVWSYGTHFLTSFQGFFIALIYCFLNGEVRAVLLKSLATQLSVRGHPEWAPKRASMYSGA        539
Biarmipes         AVWSYGTHFLTSFQGFFIALIYCFLNGEVRAVLLKSLATQLSVRGHPEWAPKRASMYSGA        539
Mojavensis        AIWSYVTHFLTSFQGFFIALIYCFLNGEVRAVLLKSLAVWMSVRGHPEWVPKRASMYSAA        541
Grimshawii        AVWSYVTHFLTSFQGFFIALIYCFLNGEVRTVLLKSLAVWMSVRGHPEWAPKRASMYSGA        549
Virilis          AIWSYVTHFLTSFQGFFIALIYCFLNGEVRAVMLKSIAVWLSVRGHPEWAPKRPSMYSGA         487
                  *:*** *:**********************:*:***:*. :********.*** ****.*

Serrata           YNTAPDTDAVQ-QHQQTGENPATGKRISPPNKRLNGRKASSASIVLIHEPQQRQRLIPRL        614
Kikkawei          YNTAPDTDAVLQQQQQPGENPATGKRISPPNKRLNGRKASSASIVMIHEPQQRQRLIPRL        622
Bipectinate       YNTAPDTDAVI---QQPGDNPATGKRISPPNKRLNGRKASSASIVMIHEPQQRHRLLPRL        602
Anannassae        YNTAPDTDAVI---QQPGENPATGKRISPPNKRLNGRKASGASIVMIHEPQQRNRLLPRL        586
Fichsuphila       YNTAPDTDAV----HPAGDPSAPGKRISPPNKRLNGRKPSSASIVMIHEPQQRHRLIPRL        601
Rhopaloa          YNTAPDTDAV----QPAGDPLATGKRISPPNKRLNGRKPSSASIVMIHEPQQRQRLIPRL        591
Elegans           YNTAPDTDAV----QPAGDPSATGKRISPPHKRLNGRKPSSASIVMIHEPPQRQRLIPRL        617
Eugracilis        YNTAPDTDAV----QPAGDPSATGKRISPPNKRLNGRKPSSASIVMIHEPQQRQRLIPRL        595
Erecta            YNTAPDTDAV----QPAGDPSATGKRISPPNKRLNGRKPSSASIVMIHEPQQRQRLMPRL        592
Melanogaster      YNTAPDTDAV----QPAGDPSATGKRISPPNKRLNGRKPSSASIVMIHEPQQRQRLMPRL        592
Sechellia         YNTAPDTDAV----QPVGDPSATGKRISPPNKRLNGRKPSSASIVMIHEPQQRQRLMPRM        592
Simulans          YNTAPDTDAV----QPAGDPSATGKRISPPNKRLNGRKPSSASIVMIHEPQQRQRLMPRM        592
Mauritania        YNTAPDTDAV----QPAGDPSATGKRISPPNKRLNGRKPSSASIVMIHEPQQRQRLMPRM        592
Takahashi         YNTAPDTDAV----QPAGDPSATGKRISPPHKRLNGRKPSSASIVMIHEPQQRQRLIPRL        605
Suzuki            YNTAPDTDAV----QPAGDPSATGKRISPPHKRLNGRKPSSASIVMIHEPQQRHRLIPRL        595
Biarmipes         YNTAPDTDAV----QPAGDPLATGKRISPPHKRLNGRKPSSASIVMIHEPQQRHRLIPRL        595
Mojavensis        YNTAPDTEPPA---QQSVEALSSNIRISPQSRRLNCRKASSVIIVIANEPQRQRAAQQ--        596
Grimshawii        YNTAPDTDVVQ---QP-GEAISTSRRVPPPIKRLKSRKANNVTIVISNEPQQQQQQHQML        605
Virilis          YNTAPDTDPQL---KQ-GDPQQSGKRLSQSTKRSNSRKASSVTIVISTEPQIHR----YV         539
                  *******:      :   :    . *:    :* : ** ... **:  **  :.

Serrata           QSKAREKDKSLDKDRGKRIQLQQSQAEAMVADPTTITTANRIRSKDDDGGNGGGGGGGGS        674
Kikkawei          RSRTREKDKSLDKDREKRTQHEAVS---EPINNVTTTKANRIRSKDEDSSNGRGG----G        675
Bipectinate       HSQRKDRDKIKDRDKDNDRDRQR-DD-RQLEETEGDPATTRIHSKETA------RTGRNR        654
Anannassae        HSQRKGRDKAKDRDMENDKDRKRSDD-RQLEETEADPGTIRIHSKDTE------RASRNR        639
Fichsuphila       QNKNQEKGK--DRVEKADID------------TELDPAITRIQSKESTG----STGSRSR        643
Rhopaloa          QNKAREKGR--DRVEKADA----------ETEPEPDPAITRIHSKE--AG-S-TASRNSR        635
Elegans           QSKAREKGR--DRAERADADAD-AET-NTKTEPEPDPAITRIHSKESTAG-S-TGSRSNR        671
Eugracilis        QNKAREKSK--DRVEKAEPE------------TEPDPAISRIHSKETG-V-GGGTGSRTR        639
Erecta            QNKAREKGK--ERVEKTDKE----------AEPEPDPAISRIHSKEAD-----RARSRTR        635
Melanogaster      QNKAREKGK--DRVEKTDAE----------A--EPDPTISHIHSKEAG-----SARSRTR        633
Sechellia         QNKAREKGK--DRVEKTDAE----------A--EPDPAISRIHSKEAG-----SARSRTR        633
Simulans          QNKAREKGK--DRVEKTDAE----------A--EPDPAIARIHSKEAG-----SARSRTR        633
Mauritania        QNKAREKGK--DRVEKTDAE----------A--EPDPAISRIHSKEAG-----SARSRTR        633
Takahashi         QNKAREKSR--DRVDKADA----------ETDPQADPAISRIHSKESSGG-GGSTGSRNR        652
Suzuki            QNKEREKSK--DRVEKADAA---------ETDANPDPAISRIHSKES-GI-AGSTGSRTR        642
Biarmipes         QNKAREKSR--DRVDKADAA---------ETDPNQDPAISRIHSKET-GI-AGGTGSRTR        642
Mojavensis        --RNNNN-TANGNGNGNGNGSR--NQGKDEPASGSARSGQRIRSKETP-E------QGRS        644
Grimshawii        QQRNTNNNSTNGS-----------SPRTVEDDTGSA-MATRIRSKES----------SAG        643
Virilis          PRRNNNNNRASTGSARVRGI-----LKATEEPASGSA-VGQRIRSTDDGAS------TTGR         587
                    :   .                                 :*:*.:

Serrata           GSKWMMG-ICFRGQKVLRVPSASSVPPESVVFELSEQ   710
Kikkawei          GSKWMMG-ICFRGQKVLRVPSASSVPPESVVFELSEQ   711
Bipectinate       GSKWMMDIICFRGQKVLRVPSASSVPPESVVFELSEQ   691
Anannassae        GSKWMMDIICFRGQKVLRVPSASSVPPESVVFELSEQ   676
Fichsuphila       GSKWIMG-ICFRGQKVLRVPSASSVPPETNV------   673
Rhopaloa          GSKWIMG-ICFRGQKVLRVPSASSVPPESVVFELSEQ   671
Elegans           GSKWIMG-ICFRGQKVLRVPSASSVPPESVVFELSEQ   707
Eugracilis        GSKWIMG-ICFRGQKVLRVPSASSVPPESVVFELSEQ   675
Erecta            GSKWIMG-ICFRGQKVLRVPSASSVPPETNV------   665
Melanogaster      GSKWIMG-ICFRGQKVLRVPSASSVPPESVVFELSEQ   669
Sechellia         GSKWIMG-ICFRGQKVLRVPSASSVPPETNV------   663
Simulans          GSKWIMG-ICFRGQKVLRVPSASSVPPETNV------   663
Mauritania        GSKWIMG-ICFRGQKVLRVPSASSVPPESVVFELSEQ   669
Takahashi         GSKWIMG-ICFRGQKVLRVPSASSVPPESVVFELSEQ   687
Suzuki            GSKWIMG-ICFRGQKVLRVPSASSVPPETNV------   672
Biarmipes         GSKWIMG-ICFRGQKVLRVPSASSVPPETNV------   672
Mojavensis        TGNWMFS-LCFHGQKVLRVPPASSVPPETNV------   674
Grimshawii        RSNWMTN-LCFRGKKVLRVPPASSVPPESVVFELSEL   679
Virilis          NSNWMFG-LCFRGQKVLRVPPASSVPPESVVFELSEQ    623
```

Melanogaster NP_570007.2

```
1 mtllsnildc ggcisaqrft rllrqsgssg pspsaptagt fesksmlept sshslatgrv
       61 pllhdfdast tespgtyvld gvarvaqlal eptvmdalpd sdteqvlgnl nssapwnltl
      121 asaaatnfen csalfvnytl pqtglycnwt wdtllcwppt pagvlarmnc pggfhgvdtr
      181 kfairkceld grwgsrpnat evnppgwtdy gpcykpeiir lmqqmgskdf dayidiarrt
      241 rtleivglcl slfalivsll ifctfrslrn nrtkihknlf vamvlqviir ltlyldqfrr
      301 gnkeaatnts lsvientpyl ceasyvlley artamfmwmf ieglylhnmv tvavfqgsfp
      361 lkffsrlgwc vpilmttvwa rctvmymdts lgeclwnynl tpyywilegp rlavillnfc
      421 flvniirvlv mklrqsqasd ieqtrkavra aivllpllgi tnllhqlapl ktatnfavws
      481 ygthfltsfq gffialiycf lngevravll kslatqlsvr ghpewapkra smysgaynta
      541 pdtdavqpag dpsatgkris ppnkrlngrk pssasivmih epqqrqrlmp rlqnkarekg
      601 kdrvektdae aepdptishi hskeagsars rtrgskwimg icfrgqkvlr vpsassvppe
      661 svvfelseq
```

Simulans   XP_039153334

```
  1 mtllsnildc ggcisaqrft rllrqsgssg pspsapapgt fesksmlept sshslatgrv
       61 pllhdfdast tespgtyvld gvarvaqlal eptvmdalpd pdteqvlgnl nssapwnltl
      121 asaaatnfen csalfvnytl pqtglycnwt wdtllcwppt pagvlarmnc pggfhgvdtr
      181 kfairkceld grwgsrpnat evsppgwtdy gpcykpeiir lmqqmgskdf dayidiarrt
      241 rtleivglcl slfalivsll ifctfrslrn nrtkihknlf vamvlqviir ltlyldqfrr
      301 gnkeaatnts lsvientpyl ceasyvlley artamfmwmf ieglylhnmv tvavfqgsfp
      361 lkffsrlgwc vpilmttvwa rctvmymdts lgeclwnynl tpyywilegp rlavillnfc
      421 flvniirvlv mklrqsqasd ieqtrkavra aivllpllgi tnllhqlapl ktatnfavws
      481 ygthfltsfq gffialiycf lngevravll kslatqlsvr ghpewapkra smysgaynta
      541 pdtdavqpag dpsatgkris ppnkrlngrk pssasivmih epqqrqrlmp rmqnkarekg
      601 kdrvektdae aepdpaiari hskeagsars rtrgskwimg icfrgqkvlr vpsassvppe
      661 tnv
```

Suzuki XP_016939142.1

```
     1 mtllsnildc gggisaqrft rllrqssssg spsssssgtt fesksmlept sshilptgrv
       61 pilhdfdfds lttestgthv ldgvasvakm aleptvmdav lpdsdtdqvl snlnisapwn
      121 itlasaaatn fencsslfvn ytlpqsglyc nwtwdtllcw pptpagvlar mncpggfhgv
      181 dtrkfairkc eldgrwgsrp natevsppgw tdygpcykpe iirlmqqmgs kdfdlyidia
      241 rktrtleivg lclslfaliv sllifctfrs lrnnrtkihk nlfvamvlqv iirltlyldq
      301 frrgnkeaat ntslsvient pylceasyvl leyartamfm wmfieglylh nmvtvavfqg
      361 sfplkffsrl gwgfpilmtt vwarctvmym dtslgeclwn ynltpyywil egprlavill
      421 nfcflvniir vlvmklrqsq asdieqtrka vraaivllpl lgitnllhql aplktatnfa
      481 vwsygthflt sfqgffiali ycflngevra vllkslatql svrghpewap krasmysgay
      541 ntapdtdavq pagdpsatgk rispphkrln grkpssasiv mihepqqrhr liprlqnker
      601 ekskdrveka daaetdanpd paisrihske sgiagstgsr trgskwimgi cfrgqkvlrv
      661 psassvppet nv
```

Mauritania NC_046672.1

```
1 mtllsnildc ggcisaqrft rllrqsgssg pspsapapgt fesksmlept sshslatgrv
       61 pllhdfdatt tespgthvld gvarvaqlal eptvmdalpd pdteqvlgnl nsnapwnltl
      121 asaaatnfen csalfvnytl pqtglycnwt wdtllcwppt pagvlarmnc pggfhgvdtr
      181 kfairkceld grwgsrpnat evsppgwtdy gpcykpeiir lmqqmgskdf dayidiarrt
      241 rtleivglcl slfalivsll ifctfrslrn nrtkihknlf vamvlqviir ltlyldqfrr
      301 gnkeaatnts lsvientpyl ceasyvlley artamfmwmf ieglylhnmv tvavfqgsfp
      361 lkffsrlgwc vpilmttvwa rctvmymdts lgeclwnynl tpyywilegp rlavillnfc
      421 flvniirvlv mklrqsqasd ieqtrkavra aivllpllgi tnllhqlapl ktatnfavws
      481 ygthfltsfq gffialiycf lngevravll kslatqlsvr ghpewapkra smysgaynta
      541 pdtdavqpag dpsatgkris ppnkrlngrk pssasivmih epqqrqrlmp rmqnkarekg
      601 kdrvektdae aepdpaisri hskeagsars rtrgskwimg icfrgqkVLRVPSASSVPPESVVFELSEQ
```

Sechellia XP_032581354.1

```
    1 mtllsnildc ggcisdqrft hllrqsgssg pspsapapgt fesksmlept sphslatgrv
       61 pllhdfdast tespgtyvld gvarvaqlal eptvmdalpd pdteqvlgnl nssapwnltl
      121 asaaatnfen csalfvnytl pqtglycnwt wdtllcwppt pagvlarmnc pggfhgvdtr
      181 kfanrkceld grwgsrpnat evsppgwtdy gpcykpeiir lmqqmgskdf dayidiarrt
      241 rtleivglcl slfalivsll ifctfrslrn nrtkihknlf vamvlqviir ltlyldqfrr
      301 gnkeaatnts lsvientpyl ceasyvlley artamfmwmf ieglylhnmv tvavfqgsfp
      361 lkffsrlgwc vpilmttvwa rctvmymdts lgeclwnynl tpyywilegp rlavillnfc
```

```
421 flvniirvlv mklrqsqasd ieqtrkavra aivllpllgi tnllhqlapl ktatnfavws
481 ygthfltsfq gffialiycf lngevravll kslatqlsvr ghpewapkra smysgaynta
541 pdtdavqpvg dpsatgkris ppnkrlngrk pssasivmih epqqrqrlmp rmqnkarekg
601 kdrvektdae aepdpaisri hskeagsars rtrgskwimg icfrgqkvlr vpsassvppe
661 tnv
```

Serrata KAH8373163.1

```
1 lqlqyaflgd mtllssilds gggisaqrft rllrqsstss ssssssasg smleptssql
61 pindvlgggr ipflhdngtg eslplpdada ldpnfvldgv tsvakmalea tvkevlrdpd
121 peqilanana tapwnitlas aaatnyencs alfanytlpq tglycnwtwd tllcwpptpa
181 gvlarmncpg gyhgvdtrkf anrkceldgr wgsrpnatet natgwtdygp cykpevirlm
241 qqmgsredld lyieiarrtr tleivglcls lfalivslli fctfrslrnn rtkihknlfv
301 amvlqviirl tlyldqfrrg skeaatntsl svientpylc easyvlleya rtamfmwmfi
361 eglylhnmvt vavfqgsfpl kffsrlgwcv pilmttvwar ctviymdtsl gdclwnynlt
421 pyywilegpr lavillnfcf lvniirvlvm klrqsqasdi eqtrkavraa ivllpllgit
481 nilhqvaplk tatnfavwsy styfltsfqg ffialiycfl ngevravllk slatqlsvrg
541 hpewapkras mysgayntap dtdavqqhqq tgenpatgkr isppnkrlng rkassasivl
601 ihepqqrqrl iprlqskare kdksldkdrg kriqlqqsqa eamvadptti ttanrirskd
661 ddggnggggg gggsgskwmm gicfrgqkvl rvpsassvpp esvvfelseq
```

Erecta XP_026838097.1

```
1 mtllsnildc ggcisaqrft rllrqsgssv pspsapapgt fesismlept slhslptgrv
61 pllhdfdast tespgtyvld gvarvaqlal eptvmdglpd pdteqvlsnl nssapwnltl
121 ssaaatnfen csalfvnytl pqtglycnwt wdtllcwppt pagvlarmnc pggfhgvdtr
181 kfairkceld grwgsrpnat evsppgwtdy gpcykpeiir lmqqmgskdf dayidiarrt
241 rtleivglci slfalivsll ifctfrslrn nrtkihknlf vamvlqviir ltlyldqfrr
301 gnkeaatnts lsaientpyl ceasyvlley artamfmwmf ieglylhnmv tvavfqgsfp
361 lkffsrlgwc vpilmttvwa rctvmymdts lgeclwnynl tpyywilegp rlavillnfc
421 flvniirvlv mklrqsqasd ieqtrkavra aivllpllgi tnilhqlapl ktatnfavws
481 ygthfltsfq gffialiycf lngevravll kslatqlsvr ghpewapkra smysgaynta
541 pdtdavqpag dpsatgkris ppnkrlngrk pssasivmih epqqrqrlmp rlqnkarekg
601 kervektdke aepepdpais rihskeadra rsrtrgskwi mgicfrgqkv lrvpsassvp
661 petnv
```

Takahashi NW_025323455.1

```
1 mtllsnildc gggisaqrfa rllrqssssg spsasasssa ssssgtsfes ksmleptssq
61 nlptgrvpil hdfdfdfdpt tttespgnyv ldgvasvakm aleptvmdtv lpdsdpdqvl
121 snlnvsapwn ltlasaaatn fencsalfvn ytlpqtglyc nwtwdtllcw pptpagvlar
181 mncpagfhgv dtrkfairkc eldgrwgsrp natevsppgw tdygpcykpe iirlmeqmgs
241 knfdlyidia wktrtleivg lclslfaliv sllifctfrs lrnnrtkihk nlfvamvlqv
301 iirltlyldq yrrgnkeaat ntslsvient pylceasyvl leyartamfm wmfieglylh
361 nmvtvavfqg sfplkffsrl gwgfpilmtt vwarctvmym dtslgeclwn ynltpyywil
421 egprlavill nfcflvniir vlvmklrqsq asdieqtrka vraaivllpl lgitnilhqm
481 aplktatnfa vwsyvthflt sfqgffiali ycflngevra vllkslatql svrghpewap
541 krasmysgay ntapdtdavq pagdpsatgk rispphkrln grkpssasiv mihepqqrqr
601 liprlqnkar eksrdrvdka daetdpqadp aisrihskes sggggstgsr nrgskwimgi
661 cfrgqkdkcvVLRVPSASSVPPESVVFELSEQ
```

Biarmipes XP_016948718.1

```
1 mtllsnildc gggisaqrft rllrqssspg flsssslgtt fesksmlept sshvlptgrv
61 pvlhdfdfds sttelprilv ldgvagvatm aleptamdav lpdsdpdqvl snfnisapwn
121 itlasaaatn fencsslfvn ytlpqtglyc nwtwdtllcw pptpagvlar mncpggfhgv
181 dtrkfairkc eldgrwgsrp natevsppgw tdygpcykpe iirlmqqmgs kdfdlyidia
241 rktrtleivg lclslfaliv sllifctfrs lrnnrtkihk nlfvamvlqv iirltlyldq
301 frrgnkeaat ntslsvient pylceasyvl leyartamfm wmfieglylh nmvtvavfqg
361 sfplkffsrl gwgfpilmtt vwarctvmym dtslgeclwn ynltpyywil egprlavill
421 nfcflvniir vlvmklrqsq asdieqtrka vraaivllpl lgitnllhql aplktatnfa
481 vwsygthflt sfqgffiali ycflngevra vllkslatql svrghpewap krasmysgay
541 ntapdtdavq pagdplatgk rispphkrln grkpssasiv mihepqqrhr liprlqnkar
601 eksrdrvdka daaetdpnqd paisrihske tgiaggtgsr trgskwimgi cfrgqkvlrv
661 psassvppet nv
```

Eugracilis XP_017067611.1

1 mtllsnilds gggisaqrft rllrqssssg sssasasass ygtlesksml eptsshtlpg
61 grvpilhdfd ssttespgay lldgvasvak malepavmdv lpdpdtdqvl snlnitapwn
121 ltlasaaatn fencsalfvn ytlpqtglyc nwtwdtllcw pptpagvlar mncpggfhgv
181 dtrkfairkc eldgrwgsrp nateasppgw tdygpcykpe iirlmqqmgn kdfdlyidia
241 rktrtleivg lclslfaliv sllifctfrs lrnnrtkihk nlfvamvlqv iirltlyldq
301 frrgnkeaat ntslsvient pylceasyvl leyartamfm wmfieglylh nmvtvavfqg
361 sfplkffsrl gwcvpilmtt vwarctvmym dttlgeclwn ynltpyywil egprlavill
421 nfcflvniir vlvmklrqsq asdieqtrka vraaivllpl lgitnllhql aplktatnfa
481 vwsygthflt sfqgffiali ycflngevra vllkslatql svrghpewap krasmysgay
541 ntapdtdavq pagdpsatgk risppnkrln grkpssasiv mihepqqrqr liprlqnkar
601 ekskdrveka epetepdpai srihsketgv gggtgsrtrg skwimgicfr gqkvlrvpsa
661 ssvppesvvf elseq

Rhopaloa XP_044317065.1

1 mtllsnildc gggisaqclt rslressspg spgspesgaf esksmlepts shilptgrvp
61 vlhdfnasst dspgtflldg vvsvaqmale ptltdllpdp dpdqvlsnln asapwnltla
121 saaatnfenc salfanytlp qtglycnwtw dtllcwpptp agvlarmncp agfhgvdtrk
181 fairkceldg rwgsrpnate vsppgwtdyg pcykpeiirl mqqmgsrdfd tfidiakktr
241 tleivglcls lfalivslli fctfrslrnn rtkihknlfv amvlqviirl tlyldqyrrg
301 nkeaatntsl svientpylc easyvlleya rtamfmwmfi eglylhnmvt vavfqgsfpl
361 kffsrlgwcv pilmttvwar ctvmymdtsl gdclwnynlt pyywilegpr lavillnfcf
421 lvniirvlvm klrqsqasdi eqtrkavraa ivllpllgit nilhqmaplk tatnfavwsy
481 gthfltsfqg ffialiycfl ngevravllk slatqlsvrg hpewapkras mysgayntap
541 dtdavqpagd platgkrisp pnkrlngrkp ssasivmihe pqqrqrlipr lqnkarekgr
601 drvekadaet epepdpaitr ihskeagsta srnsrgskwi mgicfrgqkv lrvpsassvp
661 pesvvfelse q

Fichsuphila XP_017048319.1

1 mtllasildc gggisaqrft rllaqsslla sssasasasa sasasgtfas esmleptssh
61 ilptgrvpvl hgfdsttes ttstyvldgv asvaqmplle stvmdalpds dpdqafgnln
121 vtapwnltla saaatnfenc salfanytlp qsglycnwtw dtllcwpptp agvlarmncp
181 ggfhgvdtrk fairkceldg rwgsrpnate vsppgwtdyg pcykpeiirl mqqmgnkdfd
241 lyidiarktr tleivglcls lfalivslli fctfrslrnn rtkihknlfv amvlqviirl
301 tlyldqfrrg nkeaatntsl svientpylc easyvlleya rtamfmwmfi eglylhnmvt
361 vavfqgsfpl kffsrlgwcv pilmttvwar ctvmymdtsl geclwnynlt pyywilegpr
421 lavillnfcf lvniirvlvm klrqsqasdi eqtrkavraa ivllpllgit nllhqlaplk
481 tatnfavwsy gthfltsfqg ffialiycfl ngevravllk slatqlsvrg hpewapkras
541 mysgayntap dtdavhpagd psapgkrisp pnkrlngrkp ssasivmihe pqqrhrlipr
601 lqnknqekgk drvekadidt eldpaitriq skestgstgs rsrgskwimg icfrgqkvlr
661 vpsassvppe tnv

Elegans NW_024545513.1)
1 mkvaftsqmn nennhfhnvv dgemtllsni ldcgggipaq rlarllrqss ssgstpsgsa
61 sgpfesksml epttshilpt grvpilhdfg ssstespltg tylldgvasv aemaleptvr
121 dilsdpdpdk vlsnlnasap wnltlassaa tnfencsalf anytlpqtgl ycnwtwdtll
181 cwpptpagvl armncpggfh gvdtrkfair kceldgrwgs rpnatevspp gwtdygpcyk
241 peiirlmqqm gskdfdlyid iarktrtlei vglclslfal ivsllifctf rslrnnrtki
301 hknlfvamvl qviirltlyl dqyrrgnkea atntslsvie ntpylceasy vlleyartam
361 fmwmfiegly lhnmvtvavf qgsfplkffs rlgwcapilm ttvwarctvm ymdtslgecl
421 wnynltpyyw ilegprlavi llnfcflvni irvlvmklrq sqasdieqtr kavraaivll
481 pllgitnllh qlaplktatn favwsygthf ltsfqgffia liycflngev ravllkslat
541 qlsvrghpew apkrasmysg ayntapdtda vqpagdpsat gkrispphkr lngrkpssas
601 ivmiheppqr qrliprlqsk arekgrdrae radadadaet ntktepepdp aitrihskes
661 tagstgsrsn rgskwimgic frgqkVLRVPSASSVPPESVVFELSEQ

Kikkawei KAH8343197.1

```
  1 mqlqyafagd mtllssilds ggsisaqrft rllrqsstss sssvsasasm leptssqlpn
       61 ndvlgggrip flpgnatges ssssspplps pslpdadald pnfvldgvts vakmaleatv
      121 kvlrdpdpeq ilananasap wnitlaaaaa tnyencsalf anytlpqtgl ycnwtwdtll
      181 cwpptpagvl armncpggyh gvdtrkfanr kceldgrwgs rpnatetnat gwtdygpcyk
      241 pevirlmqqm gsredldlyi eiakrtrtle ivglclslfa livsllifct frslrnnrtk
      301 ihknlfiamv lqviirltly ldqfrrgnke aatntslsvi entpylceas yvlleyarta
      361 mfmwmfiegl ylhnmvtvav fqgsfplkff srlgwcvpil mttvwarctv iymdtslgdc
      421 lwnynltpyy wilegprlav illnfcflvn iirvlvmklr qsqasdieqt rkavraaivl
      481 lpllgitnil hqlaplktat nfavwsysty fltsfqgffi aliycflnge vravllksla
      541 tqlsvrghpe wapkrasmys gayntapdtd avlqqqqqpg enpatgkris ppnkrlngrk
      601 assasivmih epqqrqrlip rlrsrtrekd ksldkdrekr tqheavsepi nnvtttkanr
      661 irskdedssn grggggskwm mgicfrgqkv lrvpsassvp pesvvfelse q
```

Bipectinate KAH8261948.1

```
  1 hyadtgnmtl lssildsggs vsvqrltrll rqsspsasas gtalasesmv eptasqmlne
       61 itdggrvpsl gqdlatttet tgqsnviidr vtnvakmale atvmdvlpdp dpdqvlagvn
      121 asahwnmtls sstatnyenc salfanytlp qtglycnwtw dtllcwpptp agvlarmhcp
      181 ggyhgvdtrk fanrkceldg rwgsrpnate pspagwtdyg pcykpevirl mqsmgskdid
      241 iyidiarrtr tleivglwvs lfalvislli fctfrslrnn rtkihknlfi amvlqvivrl
      301 tlyldqyrrg nkeaatntsl saientpylc easyvlleya rtamfmwmfi eglylhnmvt
      361 vavfqgsfpl kffsrlgwcv pilmtfvwar ctvmymdtsm geclwnynlt pyywilegpr
      421 lavillnfcf lvniirvlvv klrqsqasdi eqtrkavraa ivllpllgit nllhqlaplk
      481 tatnfavwsy gthfltsfqg ffialiycfl ngevravllk slatqmsvrg hpewvpkras
      541 mysgayntap dtdaviqqpg dnpatgkris ppnkrlngrk assasivmih epqqrhrllp
      601 rlhsqrkdrd kikdrdkdnd rdrqrddrql eetegdpatt rihsketart grnrgskwmm
      661 diicfrgqkv lrvpsassvp pesvvfelse q
```

Anannassae XP_044573162.1

```
1 mtllssilds ggsvsvqrlt rllrqsstss sasgpslasd sissmveats lrvpslghdl
       61 stttetnvii drvtnvakma leatvmdvlp dpepdqvltg vnasshwnmn mtlssasatn
      121 yencsalfan ytlpqtglyc nwtwdtllcw pptpagvlar mncpggyhgv dtrkfanrkc
      181 eldgrwgsrp natepspagw tdygpcykpe virlmqsmgs kdidiyidia rrtrtleivg
      241 lwvslfalvi sllifctfrs lrnnrtkihk nlfvamvlqv ivrltlyldq yrrgnkeaat
      301 ntslsaient pylceasyvl leyartamfm wmfieglylh nmvtvavfqg sfplkffsrl
      361 gwcvpilmtf vwarctvmym dtsmgeclwn ynltpyywil egprlavill nfcflvniir
      421 vlvvklrqsq asdieqtrka vraaivllpl lgitnllhql aplktatnfa vwsygthflt
      481 sfqgffiali ycflngevra vllkslatqm svrghpewvp krasmysgay ntapdtdavi
      541 qqpgenpatg krisppnkrl ngrkasgasi vmihepqqrn rllprlhsqr kgrdkakdrd
      601 mendkdrkrs ddrqleetea dpgtirihsk dterasrnrg skwmmdiicf rgqkvlrvps
      661 assvppesvv felseq
```

Grimshawii EDW00227.1

```
1 mmqlpginkp gciptistss iiipqrfarr aarvaststa tvttlsmsmt qsgrsmlnrt
       61 mtsiatattt ststsisths ttssstsklp sqvisgladv ingteldmta aatavgldaa
      121 vpdpsttsss wnstfstasa nsydncaamf anytqpvtgl ycnwtwdsll cwpptpagvm
      181 armycpagyh gvdtrkfair kceldghwgr rpndtelntg gwtdyapcyk pevirlmqei
      241 kdvnvymdia qrtrtleiig lclsllalii slvifcafrs lrnnrtkihk nlfvamvlqv
      301 ivrltlyldq frrgrpetat nasvsiient pylceasyvl leyartamfm wmfieglylh
      361 nmvtvavfqg nfplklfall gwglpvlmtf vwvqctaifm dttvgecmwn ynltpyywil
      421 egprlavill nffflvniir vlvvklrqsq asdieqtrka vraaivllpl lgitnllhlv
      481 palktawkfa vwsyvthflt sfqgffiali ycflngevrt vllkslavwm svrghpewap
      541 krasmysgay ntapdtdvvq qpgeaistsr rvpppikrlk srkannvtiv isnepqqqqq
      601 qhqmlqqrnt nnnstngssp rtveddtgsa matrirskes sagrsnwmtn lcfrgkkvlr
      661 vppassvppe svvfelsel
```

Mojavensis XP_002010381.2

1 mqlagdseag smptiittst ssnsflqria lkavtaattt kdrlisivet dspagshmdp

```
       61 stamtttsit atatatpaat ataselsrsd vlnasieatt avgmdtvkta tesglptvsg
      121 ttlpswnssf nsalannydn csamfanyth pqtglycnwt wdsllcwppt pagnlarmnc
      181 pagyhgvdtr kfanrkceld ghwggrpnet epnhagwtdy gpcykpevir lmqeikdvnl
      241 yidiaqrtrt leiiglclsl laliislvif cafrslrnnr tkihknlfia mvlqvivrlt
      301 lyldqfrrgi qynsslsaie ntpylceasy vlleyartam fmwmfiegly lhnmvtvavf
      361 qgsfpliffs llgwgmpvvm tfvwvqctai fmdtalgdcm wnynltpyyw ilegprlavi
      421 llnffflvni irvlvvklrq sqasdieqtr kavraaivll pllgitnllh lvpalktawk
```

481 faiwsyvthf ltsfqgffia liycflngev ravllkslav wmsvrghpew vpkrasmysa
541 ayntapdtep paqqsveals snirispqsr rlncrkassv iivianepqr qraaqqrnnn
601 ntangngngn gngsrnqgkd epasgsarsg qrirsketpe qgrstgnwmf slcfhgqkvl
661 rvppassvpp etnv

Virilis XP_032288784.1

1 mptaapssnr tvsthhsttm tststtygtt sttqaaltsf epmavtaagm dvtmagtepl
61 sstsiptfwn ssintasans ydncsalfan ytqpttviyc nwtwdsllcw pptpagatah
121 mhcpagyhgv dtrkfanrkc eldghwagrp nsteqkptgw tdygpcykpe virlmqeikd
181 vnlymdiaqr trtleiiglc lslfaliisl mifcafrslr nnrtkihknl fvamvlqviv
241 rltlyldqfr rgksdsannt slsvientpy lceasyvlle yartamfmwm fieglylhnm
301 itvavfqgnf plvffsllgw gmpvlmtfvw vqctaifmdt slgdclwnyn ltpyywileg
361 prltvimlnf fflvniirvl vmklrqsqas eieqtrkavr aaivllpllg itnllhlvpa
421 lktawkfaiw syvthfltsf qgffialiyc flngevravm lksiavwlsv rghpewapkr
481 psmysgaynt apdtdpqlkq gdpqqsgkrl sqstkrsnsr kassvtivis tepqihryvp
541 rrnnnnrast gsarvrgilk ateepasgsa vgqrirstdd gasttgrnsn wmfglcfrgq
601 kvlrvppass vppesvvfel seq
