## Supplementary material for "The incidence of candidate binding sites for β-arrestin in Drosophila neuropeptide GPCRs": S13 Text

### *S13. Text* Multi-species analysis of CCHa R2 Rs

**Supporting Figure 16**

Blastp search retrieved PA and PB isoforms from *D virilis*; however, not from several of the other species in the extended evolutionary analyses.

The lack of a PA in all species is significant because *D suzuki* included a BBS in the additional (PA-specific) sequence. Direct inspection of genomic DNA from each of the missing species (below) revealed them. Genbank references are included for each. First is shown the Clustal alignment of all 19 species

CLUSTAL Line-ups; Genbank Reference IDs below

Predicted TM domains in YELLOW

BBS sequences in RED

Here is the Clustal line up of PB isoforms:

```
Mojavensis        ----MYAALMDMSQMLAASLAYAPAESSSAAA--------AAGVNLSQLSNSSQLDG---        45
Virilis          MPKNMLAALMDMSQTLAASLAYAPLESNAAATAAA-AAAAAVVLNVSQLGNSSQLDG---         56
Grimshawi         ----MYAALMDMSQTLAASLAYAPMESNSAAAAAAAAAAAALGMNVSQLSNSSQLDG---        53
Bipectinate       -----MSALMDMSQTLALSLLYAPPDATGGVSPNIISIGSGN------GSDGGGNDSSLG        49
Anannassae        ----MSAALMDMSQTLALSLVYAPLDATGGASPNIA-IGNGNDSN--GGGDGGANDSGLV        53
Serrata           ----VDTDVMDIAQALVASLAHAPLDGSGG---------------------GGGNGN---        32
Kikkawei          ----VNTNKMDIAQALVASLAHAPLDGSGN---------------------G----S---        28
Fichsuphila       ----MHAALMDVGQTLAGGS------SAND---------------------TGLLATGGE        29
Rhopaloa          ----MFAALMDVGQTLAGSL------A-----------------------------E-GN        20
Elegans           ----MFAALMDVGQTLAGSL------A-----------------------------TEGN        21
Suzuki            ----MYAALVDVGQTLVAGLVDRA--GANE---------------------SGLLASGQA        33
Biarmipes         ----MYVALVDVGQTLAAGLADG----ANE---------------------SVLMATGHA        31
Takahashi         ----MYAALMDVGQTLAASLAEGGGN---E---------------------SGLLATQ--        30
Erecta            ----MYAALMDVGQTLAARLADGEGN---D---------------------SGLLATRQG        32
Melanogaster      ----MYASLMDVGQTLAARLADSDGNGAND---------------------SGLLATGQG        35
Sechellia         ----MYASLMDVGQTLAARLADGEGNGAND---------------------SGLLATGQG        35
Eugracilis        ----MYASLMDVGQTLAARLADGEGNGAND---------------------SGLLATGQG        35
Simulans          ----MYASLMDVGQTLAARLADGEGNGAND---------------------SGLLATGQG        35
Mauritania        ----MYASLMDVGQTLAARLADGEGNGAND---------------------SGLLATGQG        35
                           :*:.* *.


Mojavensis        -SSVVTVATAAT------VGQHN-A-SIEESSYVKVLDRPETYIVTVLYTLIFIVGVLGN        96
Virilis          -SLATAAATTTTTAVTTSTSTHNAS-GEEYPQYVKVLDRPETYIVTVLYTLIFIVGVLGN         114
Grimshawi         -STGATAAT---------SIPHNVS-AEEYPQYVKVLDRPETYIVTVLYTLIFIVGVLGN        102
Bipectinate       LATG-QQGAATAVGGVLGLGQHNASADGGGGPYVPVLERPETYIVTVLYTLIFIVGVLGN        108
Anannassae        LATG-QQGVATVVGGVLGLGQHNGSADGGIGPYVPVLERPETYIVTVLYTLIFIVGVLGN        112
Serrata           -GSGANDSVLLATGTGAAAEQHNASIDGGMVPYVPVLDRPETYIVTVLYTLIFIVGVLGN        91
Kikkawei          -GSGGNDSVLLAT--GASAEQHNASIDGGMVPYVPVLDRPETYIVTVLYTLIFIVGVLGN        85
Fichsuphila       QEQEQGPGLGLGMGMGMEMGQHNASADGGMVPYVPVLDRPETYIVTVLYTLIFIVGVLGN        89
Rhopaloa          GSGVQGEGLELGPGMGVEVGQHNASADGGIVPYVPVLDRPETYIVTVLYTLIFIVGVLGN        80
Elegans           DSGLLATG-QYEGGVEVGMGQHNSSADGGMVPYVPVLDRPETYIVTVLYTLIFIVGVLGN        80
Suzuki            AEQEQG----------HG-VAHNASADDGMVPYVPVLDRPETYIVTVLYTLIFIVGVLGN        82
Biarmipes         AEQEQG----------HW-VGQNASGDGVMVPYVPVLDRPETYIVTVLYTLIFIVGVLGN        80
Takahashi         -------EGQ---GLDLE-TGHNASADGGIVPYVPVLDRPETYIVTVLYTLIFIVGVLGN        79
Erecta            LEQEQEQEQE---GLALG-MAHNASADGGMVPYVPVLDRPETYIVTVLYTLIFVVGVLGN        88
Melanogaster      LEQ----EQE---GLALD-MGHNASADGGIVPYVPVLDRPETYIVTVLYTLIFIVGVLGN        87
Sechellia         LEQ----EQE---GLALD-MGHNASADGGIVPYVPVLDRPETYIVTVLYTLIFIVGVLGN        87
Eugracilis        LEQ----EQE---GLALD-MGHNASADGGIVPYVPVLDRPETYIVTVLYTLIFIVGVLGN        87
Simulans          LEQ----EQE---GLALD-MGHNASADGGIVPYVPVLDRPETYIVTVLYTLIFIVGVLGN        87
Mauritania        LEQ----EQE---GLALD-MGHNASADGGIVPYVPVLDRPETYIVTVLYTLIFIVGVLGN        87
                                       :* :       ** **:***************:******

Mojavensis        GTLVIIFFRHRSMRNIPNTYILSLALADLLVILVCVPVATIVYTQESWPFERNMCRISEF        156
Virilis          GTLVIIFFRHRSMRNIPNTYILSLALADLLVILVCVPVATIVYTQESWPFERNMCRITEF         174
Grimshawi         GTLVIIFFRHRSMRNIPNTYILSLALADLLVILVCVPVATIVYTQESWPFERNMCRISEF        162
```

```
Bipectinate      GTLVIIFFRHRSMRNIPNTYILSLALADLLVILVCVPVATIVYTQESWPFERNMCRISEF        168
Anannassae       GTLVIIFFRHRSMRNIPNTYILSLALADLLVILVCVPVATIVYTQESWPFERNMCRISEF        172
Serrata          GTLVIIFFRHRSMRNIPNTYILSLALADLLVILVCVPVATIVYTQESWPFERNMCRISEF        151
Kikkawei         GTLVIIFFRHRSMRNIPNTYILSLALADLLVILVCVPVATIVYTQESWPFERNMCRISEF        145
Fichsuphila      GTLVIIFFRHRSMRNIPNTYILSLALADLLVILVCVPVATIVYTQESWPFERNMCRISEF        149
Rhopaloa         GTLVIIFFRHRSMRNIPNTYILSLALADLLVILVCVPVATIVYTQESWPFERNMCRISEF        140
Elegans          GTLVIIFFRHRSMRNIPNTYILSLALADLLVILVCVPVATIVYTQESWPFERNMCRISEF        140
Suzuki           GTLVIIFFRHRSMRNIPNTYILSLALADLLVILVCVPVATIVYTQESWPFDRNMCRISEF        142
Biarmipes        GTLVIIFFRHRSMRNIPNTYILSLALADLLVILVCVPVATIVYTQESWPFERNMCRISEF        140
Takahashi        GTLVIIFFRHRSMRNIPNTYILSLALADLLVILVCVPVATIVYTQESWPFERNMCRISEF        139
Erecta           GTLVIIFFRHRSMRNIPNTYILSLALADLLVILVCVPVATIVYTQESWPFERNMCRISEF        148
Melanogaster     GTLVIIFFRHRSMRNIPNTYILSLALADLLVILVCVPVATIVYTQESWPFERNMCRISEF        147
Sechellia        GTLVIIFFRHRSMRNIPNTYILSLALADLLVILVCVPVATIVYTQESWPFERNMCRISEF        147
Eugracilis       GTLVIIFFRHRSMRNIPNTYILSLALADLLVILVCVPVATIVYTQESWPFERNMCRISEF        147
Simulans         GTLVIIFFRHRSMRNIPNTYILSLALADLLVILVCVPVATIVYTQESWPFERNMCRISEF        147
Mauritania       GTLVIIFFRHRSMRNIPNTYILSLALADLLVILVCVPVATIVYTQESWPFERNMCRISEF        147
                 **************************************************:******:**

Mojavensis       FKDISIGVSVFTLTALSGERYCAIVNPLRKLQTKPLTVFTAVIIWVFAIMLGMPSFVVSD        216
Virilis         FKDISIGVSVFTLTALSGERYCAIVNPLRKLQTKPLTVFTAVIIWVFAIMLGMPSFVVSD         234
Grimshawi        FKDISIGVSVFTLTALSGERYCAIVNPLRKLQTKPLTVFTAVIIWVLAIMLGMPSFVVSD        222
Bipectinate      FKDISIGVSVFTLTALSGERYCAIVNPLRKLQTKPLTVFTAAMIWVLAILLGMPSFLVSN        228
Anannassae       FKDISIGVSVFTLTALSGERYCAIVNPLRKLQTKPLTVFTAAMIWVLAILLGMPSFLFSN        232
Serrata          FKDISIGVSVFTLTALSGERYCAIVNPLRKLQTKPLTVFTAVMIWILAILLGMPSFLVSD        211
Kikkawei         FKDISIGVSVFTLTALSGERYCAIVNPLRKLQTKPLTVFTAVMIWILAILLGMPSFLVSD        205
Fichsuphila      FKDISIGVSVFTLTALSGERYCAIVNPLRKLQTKPLTVFTAVIIWILAILLGMPSFLVSD        209
Rhopaloa         FKDISIGVSVFTLTALSGERYCAIVNPLRKLQTKPLTVFTAVIIWILAILLGMPSFLVSD        200
Elegans          FKDISIGVSVFTLTALSGERYCAIVNPLRKLQTKPLTVFTAVIIWILAILLGMPSFLVSD        200
Suzuki           FKDISIGVSVFTLTALSGERYCAIVNPLRKLQTKPLTVFTAVIIWILAILLGMPSVLFSD        202
Biarmipes        FKDISIGVSVFTLTALSGERYCAIVNPLRKLQTKPLTVFTAAIIWILAILLGMPSVLFSD        200
Takahashi        FKDISIGVSVFTLTALSGERYCAIVNPLRKLQTKPLTVFTAVIIWILAILLGMPSVLFSD        199
Erecta           FKDISIGVSVFTLTALSGERYCAIVNPLRKLQTKPLTVFTAVMIWILAILLGMPSVLFSD        208
Melanogaster     FKDISIGVSVFTLTALSGERYCAIVNPLRKLQTKPLTVFTAVMIWILAILLGMPSVLFSD        207
Sechellia        FKDISIGVSVFTLTALSGERYCAIVNPLRKLQTKPLTVFTAVMIWILAILLGMPSVLFSD        207
Eugracilis       FKDISIGVSVFTLTALSGERYCAIVNPLRKLQTKPLTVFTAVMIWILAILLGMPSVLFSD        207
Simulans         FKDISIGVSVFTLTALSGERYCAIVNPLRKLQTKPLTVFTAVMIWILAILLGMPSVLFSD        207
Mauritania       FKDISIGVSVFTLTALSGERYCAIVNPLRKLQTKPLTVFTAVMIWILAILLGMPSVLFSD        207
                 *****************************************.:**::**:*****.:.*:

Mojavensis       IQSYNITTPNGNITIGVCSPFRSKIYAKYMVVAKASIYYLVPLSIIGVLYIMMAKRLHIS        276
Virilis         IQGYTLPTPNGNITIEVCSPFRSKIYAKYMVVAKASIYYLVPLSIIGVLYIMMAKRLHIS         294
Grimshawi        LQGYTLPTNKGNITIEVCSPFRSKIYAKYMVVAKASIYYFVPLSIIGVLYIMMAKRLHIS        282
Bipectinate      IKSYTVLTPNGNMSIEVCDPFRDPEYAKYMVAAKASIYYLLPLSIIGALYIMMAKRLHIS        288
Anannassae       IKSYTVLTPNGNMSIEVCDPYRDPEYAKYMVAAKASIYYLVPLSIIGALYIMMAKRLHIS        292
Serrata          IKSYPVLTATGNMTIEVCSPFRDPEYAQYMVAAKAFIYYLLPLSIIGALYIMMAKRLHIS        271
Kikkawei         IKSYPVLTANGNMTIEVCSPYRDPEYAQCMVAAKAFIYYLLPLSIIGALYIMMAKRLHIS        265
Fichsuphila      IKSYPVLTAKGNMTIEVCSPFRDPEYAKCMVAAKAFIYYLLPLSIIGALYIMMAKRLHMS        269
Rhopaloa         IKSYPVVTVKGNITIEVCSPYSDPEYAKYMVAAKATIYYLLPLSIIGALYIMMAKRLHMS        260
Elegans          IKSYPVYTANGNMTIEVCSPFRDPEYAKCMVAAKAFIYYLLPLSIIGALYIMMAKRLHMS        260
Suzuki           IKSYPVFTAMGNITIEVCSPFRDPEYAKFMVAAKALIYYLLPLSIIGALYIMMANRLHMS        262
Biarmipes        IKSYPVLTAMGNMTIEVCSPFRDAEYAKFMVAAKALIYYLLPLSIIGALYIMMAKRLHMS        260
Takahashi        IKSYPVATPMGNITIEVCSPFRDREYAKFMVAAKALIYYLLPLSIIGALYIMMAKRLHMS        259
Erecta           IKSYPVFTATGNMTIEVCSPFRDPEYAKFMVAGKALVYYLLPLSIIGALYIMMAKRLHMS        268
Melanogaster     IKSYPVFTATGNMTIEVCSPFRDPEYAKFMVAGKALVYYLLPLSIIGALYIMMAKRLHMS        267
Sechellia        IKSYPVFTATGNMTIEVCSPFRDPEYAKFMVAGKALVYYLLPLSIIGALYIMMAKRLHMS        267
Eugracilis       IKSYPVFTATGNMTIEVCSPFRDPEYAKFMVAGKALVYYLLPLSIIGALYIMMAKRLHMS        267
Simulans         IKSYPVFTATGNMTIEVCSPFRDPEYAKFMVAGKALVYYLLPLSIIGALYIMMAKRLHMS        267
Mauritania       IKSYPVFTATGNMTIEVCSPFRDPEYAKFMVAGKALVYYLLPLSIIGALYIMMAKRLHMS        267
                 ::.* : *  **::* **.*: .  **: **..** :**::******.******:***:*

Mojavensis       ARDMPGEQLSIQSRSQARARRHVARMVVAFVVVFFICFFPYHVFELWYHFYPTAEDDFDD        336
Virilis         ARDMPGEQLSIQSRSQARARRHVARMVVAFVVVFFICFFPYHVFELWYHFYPTAEEDFDD         354
Grimshawi        ARDMPGEQLSIQSRSQARARRHVARMVVAFVVVFFICFFPYHVFELWYHFYPTAEDDFDD        342
Bipectinate      ARDMPGEQQSMQSRSQARARRHVARMVVAFVVVFFICFFPYHVFELWYHFYPTAEEDFDE        348
Anannassae       ARDMPGEQQSMQSRSQARARRYVARMVVAFVVVFFICLFPYYVFELWYHFYPTAEEDFDE        352
Serrata          ARDMPGEQQSMQSRTQARARRHVARMVVAFVVVFFICFFPYHVFELWYHFYPTAEEDFDD        331
Kikkawei         ARDMPGEQQSMQSRTQARARRHVARMVVAFVVVFFICFFPYHVFELWYHFYPTAEEDFDD        325
Fichsuphila      ARNMPGEQQSMQSRTQARARRHVARMVVAFVVVFFICFFPYHVFELWYHFYPTAEEDFDD        329
Rhopaloa         ARNMPGEQQSMQSRTQARARRHVARMVVAFVVVFFICFFPYHVFELWYHFYPTAEEDFDD        320
Elegans          ARNMPGEQQSMQSRTQARARRHVARMVVAFVVVFFICFFPYHVFELWYHFYPTAEEDFDD        320
Suzuki           ARNMPGEQQSMQSRTQARARRHVARMVVAFVVVFFICFFPYHVFELWYHFYPTAEEDFDD        322
```

```
Biarmipes          ARNMPGEQQSMQSRTQARARRHVARMVVAFVVVFFICFFPYHVFELWYHFYPTAEEDFDD        320
Takahashi          ARNMPGEQQSMQSRTQARARRHVARMVVAFVVVFFICFFPYHVFELWYHFYPTAEEDFDD        319
Erecta             ARNMPGEQQSMQSRTQARARLHVARMVVAFVVVFFICFFPYHVFELWYHFYPTAEEDFDE        328
Melanogaster       ARNMPGEQQSMQSRTQARARLHVARMVVAFVVVFFICFFPYHVFELWYHFYPTAEEDFDE        327
Sechellia          ARNMPGEQQSMQSRTQARARLHVARMVVAFVVVFFICFFPYHVFELWYHFYPTAEEDFDE        327
Eugracilis         ARNMPGEQQSMQSRTQARARLHVARMVVAFVVVFFICFFPYHVFELWYHFYPTAEEDFDE        327
Simulans           ARNMPGEQQSMQSRTQARARLHVARMVVAFVVVFFICFFPYHVFELWYHFYPTAEEDFDE        327
Mauritania         ARNMPGEQQSMQSRTQARARLHVARMVVAFVVVFFICFFPYHVFELWYHFYPTAEEDFDE        327
                   **:***** *:***:***** :***************:***:*************:***:

Mojavensis         FWNVVRIVGFCTSFLNSCVNPVALYCVSGVFRQHFNRYLCCICVKRQPHLRQHSTATGIM        396
Virilis           FWHVVRIVGFCTSFLNSCVNPVALYCVSGVFRQHFNRYLCCICVKRQPHLRQHSTATGVM         414
Grimshawi          FWHVVRIVGFCTSFLNSCVNPVALYCVSGVFRQHFNRYLCCFCVKRQPHLRQHSTATGVM        402
Bipectinate        FWNVLRIVGFCTSFLNSCVNPVALYCVSGVFRQHFNRYLCCFCVKRQPHLRQHSTATGIM        408
Anannassae         FWNVLRIVGFCTSFLNSCVNPVALYCVSGVFRQHFNRYLCCFCVKRQPHLRQHSTATGIM        412
Serrata            FWNVLRIVGFCTSFLNSCVNPVALYCVSGVFRQHFNRYLCCICVKRQPHLRQHSTATGMM        391
Kikkawei           FWNVLRIVGFCTSFLNSCVNPVALYCVSGVFRQHFNRYLCCICVKRQPHLRQHSTATGMM        385
Fichsuphila        FWNVLRIVGFCTSFLNSCVNPVALYCVSGVFRQHFNRYLCCICVKRQPHLRQHSTATGMM        389
Rhopaloa           FWNVLRIVGFCTSFLNSCVNPVALYCVSGVFRQHFNRYLCCICVKRQPHLRQHSTATGMM        380
Elegans            FWNVLRIVGFCTSFLNSCVNPVALYCVSGVFRQHFNRYLCCICVKRQPHLRQHSTATGMM        380
Suzuki             FWNVLRIVGFCTSFLNSCVNPVALYCVSGVFRLHFNRYLCCVCVKRHPHLRQHSTG---M        379
Biarmipes          FWNVLRIVGFCTSFLNSCVNPVALYCVSGVFRQHFNRYLCCFCVKRQPHVRQHSTATGMM        380
Takahashi          FWNVLRIVGFCTSFLNSCVNPVALYCVSGVFRQHFNRYLCCICVKRQPHLRQHSTATGMM        379
Erecta             FWNVLRIVGFCTSFLNSCVNPVALYCVSGVFRQHFNRYLCCICVKRQPHLRQHSTATGMM        388
Melanogaster       FWNVLRIVGFCTSFLNSCVNPVALYCVSGVFRQHFNRYLCCICVKRQPHLRQHSTATGMM        387
Sechellia          FWNVLRIVGFCTSFLNSCVNPVALYCVSGVFRQHFNRYLCCICVKRQPHLRQHSTATGMM        387
Eugracilis         FWNVLRIVGFCTSFLNSCVNPVALYCVSGVFRQHFNRYLCCICVKRQPHLRQHSTATGMM        387
Simulans           FWNVLRIVGFCTSFLNSCVNPVALYCVSGVFRQHFNRYLCCICVKRQPHLRQHSTATGMM        387
Mauritania         FWNVLRIVGFCTSFLNSCVNPVALYCVSGVFRQHFNRYLCCICVKRQPHLRQHSTATGMM        387
                   **:*:*************************** ********.****:**:*****.   *

Mojavensis         D-TSVTSMRRSTYVGGGIAAG--GAREGGPRASVHMNNNHGVS--------GAAGGRGGS        445
Virilis           D-TSVTSMRRSTYVGGGGG-GAVGGSLAAHRASLHMNNNHGVAVGGG----GGGGGRGGS         468
Grimshawi          D-TSVTSMRRSTYVGGGGGVGGATGSLAAHRASLHMNNNHGSG---A----GGPGGRTGS        454
Bipectinate        DNTSVMSMRRSTYVGGAG---------GNLRASMHRNSNHGGGGG-----SGLSAGRGAS        454
Anannassae         DNTSVMSMRRSTYVGGAG---------GNLRSSMHRNSNHGGGGG-----AGLPAGRGAS        458
Serrata            DNTSVMSMRRSTYVGGCGTG-------GNLRASLHRNSNQG--GGG---GLGGGAGRGGS        439
Kikkawei           DNTSVMSMRRSTYVGGCGAG-------GNLRASLHRNSNQGVGGGG---ALGGGTGRVGS        435
Fichsuphila        DNTSVMSMRRSTYVGV-GG--------GNLRASLHRNSNHGVGA------AGGGPGRXGS        434
Rhopaloa           DNTSVMSMRRSTYVGGATA--------GHLRASLHRNSNHGGGGGGVGGAGGLGSGRVGS        432
Elegans            DNTSVMSMRRSTYVGGAAA--------GQLRASLHRNSSNHGGG------VGFGSGRVGS        426
Suzuki             DNTTVMSMRRSTYIGSAGGG------VANLRASQHRDSCHGVG-------GGGGTGHADS        426
Biarmipes          DNTSVMSMRRSTYVGGAGGV----GA--NLRASQHRN---------------------SS        413
Takahashi          DNTSVMSMRRSTYVGGAVGC----GPAGNLRASQHRNSNHG---------GGGGSGRVGS        426
Erecta             DNTSVMSMRRSTYVGGT---------AGNLRASLHRSSNHGVGGAAGGG-GGGGSGRVGS        438
Melanogaster       DNTSVMSMRRSTYVGGT---------AGNLRASLHRNSNHGVGGAGGGVGGGVGSGRVGS        438
Sechellia          DNTSVMSMRRSTYVGGT---------AGNLRASLHRNSNHGVAGAGGGVGGG--SGRVGS        436
Eugracilis         DNTSVMSMRRSTYVGGT---------AGNLRASLHRNSNHGVAGAGGGVGGG--SGRVGS        436
Simulans           DNTSVMSMRRSTYVGGT---------AGNLRASLHRNSNHGVGGAGGGVGGG--SGRVGS        436
Mauritania         DNTSVMSMRRSTYVGGT---------AGNLRASLHRNSNHGVGGAGGGVGGG--SGRVGS        436
                   * *:* *******:*               *:* * .                      *

Mojavensis         FHRQDSMALQHAGSVNGHSH-----NANVGAGAGIGRASIINEKR              485
Virilis           FHRQDSMPLQHAGSGNGHAH-----NV-GGPGAGIGRASIINEKS               518
Grimshawi          FHRQDSMPLQHAGSGNGHAH-----NV-GG----IGRASIINEKS              500
Bipectinate        FHGGDSMPMQHSNGHGAAGGTLGGAGVGATGGPSSGRAAAVGEKR              499
Anannassae         FHGGDSMPLQHSNGHGATGGNSGGAGASATGGPSLGRTAAVGEKR              503
Serrata            FHRQDSMPLQHGNGHGAA-GGGAA--SGPSCGPAGR-AVAVGEKR              479
Kikkawei           FHRQDSMPLQHGNGHGAAAGGGTA--SGPSCGSAGR-AVAVGEKR              476
Fichsuphila        FHRQDSMPLQHGNGHGAVASAAG-----C--GSGGR-AAAGGEKR              471
Rhopaloa           FHRQDSMPLQHGNGHGSVAGGAA-------SGSGGR-ATAVGEKR              481
Elegans            FHRQDSMALQHGNGHGAATGGAS-----CAPGSGGR-GPAGGEKR              465
Suzuki             LHRQESMPLQHGNGCVSG----------------GR-AASVGEKS              466
Biarmipes          LHRQASTPLQHGNGHGTGGDETGH---GPGWVSGGRAAA-VGEKR              454
Takahashi          FHRQDSMPLQHGNGHGTGIGAGVG---GAACGSGGRASAAVGEKR              468
Erecta             FHRQDSMPLQHGNAHGAGAGGGSC---GL--GSGGRTAP-VCEKS              489
Melanogaster       FHRQDSMPLQHGNAHGGGAGGGSS---GL--GAGGRTAA-VSEKS              489
Sechellia          FYRQDSMPLQHGNAHGTGAGVGSS---GL--GAGGRTAS-VSEKR              475
Eugracilis         FYRQDSMPLQHGNAHGTGAGVGSS---GL--GAGGRTAS-VSEKR              475
Simulans           FHRQDSMPLQHGNAHGAGAGGGSS---GL--GAGGRTAS-VSEKS              487
Mauritania         FHRQDSMPLQHGNAHGAGAGGGSS---GL--GAGGRTAP-VSEKS              487
```

The above are all PB isoforms; which are listed individual below.
Here are the three isoforms predicted from D melanogaster:
PA (ALTERNATIVE SPLICE AT FINAL AA)

```
MYASLMDVGQ TLAARLADSD GNGANDSGLL ATGQGLEQEQ EGLALDMGHN ASADGGIVPY
VPVLDRPETY IVTVLYTLIF IVGVLGNGTL VIIFFRHRSM RNIPNTYILS LALADLLVIL
VCVPVATIVY TQESWPFERN MCRISEFFKD ISIGVSVFTL TALSGERYCA IVNPLRKLQT
KPLTVFTAVM IWILAILLGM PSVLFSDIKS YPVFTATGNM TIEVCSPFRD PEYAKFMVAG
KALVYYLLPL SIIGALYIMM AKRLHMSARN MPGEQQSMQS RTQARARLHV ARMVVAFVVV
FFICFFPYHV FELWYHFYPT AEEDFDEFWN VLRIVGFCTS FLNSCVNPVA LYCVSGVFRQ
HFNRYLCCIC VKRQPHLRQH STATGMMDNT SVMSMRRSTY VGGTAGNLRA SLHRNSNHGV
GGAGGGVGGG VGSGRVGSFH RQDSMPLQHG NAHGGGAGGG SSGLGAGGRT AAVSEKSFIN
RYESGVMRY
```

PB (SHORTEST)

```
MYASLMDVGQ TLAARLADSD GNGANDSGLL ATGQGLEQEQ EGLALDMGHN ASADGGIVPY
VPVLDRPETY IVTVLYTLIF IVGVLGNGTL VIIFFRHRSM RNIPNTYILS LALADLLVIL
VCVPVATIVY TQESWPFERN MCRISEFFKD ISIGVSVFTL TALSGERYCA IVNPLRKLQT
KPLTVFTAVM IWILAILLGM PSVLFSDIKS YPVFTATGNM TIEVCSPFRD PEYAKFMVAG
KALVYYLLPL SIIGALYIMM AKRLHMSARN MPGEQQSMQS RTQARARLHV ARMVVAFVVV
FFICFFPYHV FELWYHFYPT AEEDFDEFWN VLRIVGFCTS FLNSCVNPVA LYCVSGVFRQ
HFNRYLCCIC VKRQPHLRQH STATGMMDNT SVMSMRRSTY VGGTAGNLRA SLHRNSNHGV
GGAGGGVGGG VGSGRVGSFH RQDSMPLQHG NAHGGGAGGG SSGLGAGGRT AAVSEKR
```

PC Stop suppressed PB

```
1   MYASLMDVGQ TLAARLADSD GNGANDSGLL ATGQGLEQEQ EGLALDMGHN ASADGGIVPY
 61 VPVLDRPETY IVTVLYTLIF IVGVLGNGTL VIIFFRHRSM RNIPNTYILS LALADLLVIL
121 VCVPVATIVY TQESWPFERN MCRISEFFKD ISIGVSVFTL TALSGERYCA IVNPLRKLQT
181 KPLTVFTAVM IWILAILLGM PSVLFSDIKS YPVFTATGNM TIEVCSPFRD PEYAKFMVAG
241 KALVYYLLPL SIIGALYIMM AKRLHMSARN MPGEQQSMQS RTQARARLHV ARMVVAFVVV
301 FFICFFPYHV FELWYHFYPT AEEDFDEFWN VLRIVGFCTS FLNSCVNPVA LYCVSGVFRQ
361 HFNRYLCCIC VKRQPHLRQH STATGMMDNT SVMSMRRSTY VGGTAGNLRA SLHRNSNHGV
421 GGAGGGVGGG VGSGRVGSFH RQDSMPLQHG NAHGGGAGGG SSGLGAGGRT AAVSEKRXGT
481 LIVELLGEEE VVVSLQADEQ L
```

Some but not all PA isoforms were recovered in the original blastp search using melanogaster PA, and the Suzuki PA form contained a BBS in the added 12 AA. So I searched for the missing PA isoforms (sechelia, biarmipes, anannassae, mojavensis, serrata, kikkawae, ficusphelia, bipectinate, Takahashi, eugracilis). Found all - the documentation and data analysis are at the bottom of this file in section called:

"CCHa2-R PA isoform – data capture"

Here is the 1 conserved BBS in biarmipes and Suzuki PA proteins:

```
Mojavensis          FHRQDSMALQHAGSVNGHSH-----NANVGAGAGIGRASIINEKR----YDER-TRY  492
Virilis            FHRQDSMPLQHAGSGNGHAH-----NV-GGPGAGIGRASIINEKSLIKRYDER-TRY   529
Grimshawi           FHRQDSMPLQHAGSGNGHAH-----NV-GG----IGRASIINEKSLIKRYEER-ARY  511
Bipectinate         FHGGDSMPMQHSNGHGAAGGTLGGAGVGATGGPSSGRAAAVGEKSINRFESNR-LRY  510
Anannassae          FHGGDSMPLQHSNGHGATGGNSGGAGASATGGPSLGRTAAVGEKRINRFESNR-MRY  514
Serrata             FHRQDSMPLQHGNGHGAA-GGGAA--SGPSCGPAGR-AVAVGEKSLMGRYVSDRLRY 491
Kikkawei            FHRQDSMPLQHGNGHGAAAGGGTA--SGPSCGSAGR-AVAVGEKSLMGRYVSDRLRY 482
Fichsuphila         FHRQDSMPLQHGNGHGAVASAAG-----C--GSGGR-AAAGGEKSIINHYENDRLRY  482
Rhopaloa            FHRQDSMPLQHGNGHGSVAGGAA-------SGSGGR-ATAVGEKSLINRYESDRIRY  481
Elegans             FHRQDSMALQHGNGHGAATGGAS-----CAPGSGGR-GPAGGEKSLINRYESERIRY  477
Suzuki              LHRQESMPLQHGNGCVSG----------------GR-AASVGEKSLLSRYENDRIRY  478
Biarmipes           LHRQASTPLQHGNGHGTGGDETGH---GPGWVSGGRAAA-VGEKSFISRYENDRIRF 466
Takahashi           FHRQDSMPLQHGNGHGTGIGAGVG---GAACGSGGRASAAVGEKSFINRYESGVMRY 470
Erecta              FHRQDSMPLQHGNAHGAGAGGGSC---GL--GSGGRTAP-VCEKSFINRYESGRMRY  501
Melanogaster        FHRQDSMPLQHGNAHGGGAGGGSS---GL--GAGGRTAA-VSEKSFINRYESGVMRY  501
Sechellia           FYRQDSMPLQHGNAHGTGAGVGSS---GL--GAGGRTAS-VSEKSFINRYESGVMRY 489
Eugracilis          FYRQDSMPLQHGNAHGTGAGVGSS---GL--GAGGRTAS-VSEKSLINKR--DRIRY  485
Simulans            FHRQDSMPLQHGNAHGAGAGGGSS---GL--GAGGRTAS-VSEKSFINRYESGVMRY  499
Mauritania          FHRQDSMPLQHGNAHGAGAGGGSS---GL--GAGGRTAP-VSEKSFINRYESGVMRY  499
```

Melanogaster NP_610199.2

```
        1 myaslmdvgq tlaarladsd gngandsgll atgqgleqeq eglaldmghn asadggivpy
       61 vpvldrpety ivtvlytlif ivgvlgngtl viiffrhrsm rnipntyils laladllvil
      121 vcvpvativy tqeswpfern mcriseffkd isigvsvftl talsgeryca ivnplrklqt
      181 kpltvftavm iwilaillgm psvlfsdiks ypvftatgnm tievcspfrd peyakfmvag
      241 kalvyyllpl siigalyimm akrlhmsarn mpgeqqsmqs rtqararlhv armvvafvvv
      301 fficffpyhv felwyhfypt aeedfdefwn vlrivgfcts flnscvnpva lycvsgvfrq
      361 hfnrylccic vkrqphlrqh statgmmdnt svmsmrrsty vggtagnlra slhrnsnhgv
      421 ggagggvggg vgsgrvgsfh rqdsmplqhg nahgggaggg ssglgaggrt aavseksfin
      481 ryesgvmry
```

Simulans XP_016026076.2

```
1 myaslmdvgq tlaarladge gngandsgll atgqgleqeq eglaldmghn asadggivpy
       61 vpvldrpety ivtvlytlif ivgvlgngtl viiffrhrsm rnipntyils laladllvil
      121 vcvpvativy tqeswpfern mcriseffkd isigvsvftl talsgeryca ivnplrklqt
      181 kpltvftavm iwilaillgm psvlfsdiks ypvftatgnm tievcspfrd peyakfmvag
      241 kalvyyllpl siigalyimm akrlhmsarn mpgeqqsmqs rtqararlhv armvvafvvv
      301 fficffpyhv felwyhfypt aeedfdefwn vlrivgfcts flnscvnpva lycvsgvfrq
      361 hfnrylccic vkrqphlrqh statgmmdnt svmsmrrsty vggtagnlra slhrnsnhgv
      421 ggagggvggg sgrvgsfhrq dsmplqhgna hgagagggss glgaggrtas vseksfinry
      481 esgvmry
```

Suzuki XP_036669887.1

```
1 myaalvdvgq tlvaglvdra ganesgllas gqaaeqeqgh gvahnasadd gmvpyvpvld
       61 rpetyivtvl ytlifivgvl gngtlviiff rhrsmrnipn tyilslalad llvilvcvpv
      121 ativytqesw pfdrnmcris effkdisigv svftltalsg erycaivnpl rklqtkpltv
      181 ftaviiwila illgmpsvlf sdiksypvft amgnitievc spfrdpeyak fmvaakaliy
      241 yllplsiiga lyimmanrlh msarnmpgeq qsmqsrtqar arrhvarmvv afvvvfficf
      301 fpyhvfelwy hfyptaeedf ddfwnvlriv gfctsflnsc vnpvalycvs gvfrlhfnry
      361 lccvcvkrhp hlrqhstgmd nttvmsmrrs tyigsagggv anlrasqhrd schgvggggg
      421 tghadslhrq esmplqhgng cvsggraasv geksllsrye ndriry
```

Mauritania XP_033155286.1

```
1 myaslmdvgq tlaarladge gngandsgll atgqgleqeq eglaldmghn asadggivpy
       61 vpvldrpety ivtvlytlif ivgvlgngtl viiffrhrsm rnipntyils laladllvil
      121 vcvpvativy tqeswpfern mcriseffkd isigvsvftl talsgeryca ivnplrklqt
      181 kpltvftavm iwilaillgm psvlfsdiks ypvftatgnm tievcspfrd peyakfmvag
      241 kalvyyllpl siigalyimm akrlhmsarn mpgeqqsmqs rtqararlhv armvvafvvv
      301 fficffpyhv felwyhfypt aeedfdefwn vlrivgfcts flnscvnpva lycvsgvfrq
      361 hfnrylccic vkrqphlrqh statgmmdnt svmsmrrsty vggtagnlra slhrnsnhgv
      421 ggagggvggg sgrvgsfhrq dsmplqhgna hgagagggss glgaggrtap vseksfinry
      481 esgvmry
```

Sechellia XP_002043393.1

```
1 myaslmdvgq tlaarladge gngandsgll atgqgleqeq eglaldmghn asadggivpy
       61 vpvldrpety ivtvlytlif ivgvlgngtl viiffrhrsm rnipntyils laladllvil
      121 vcvpvativy tqeswpfern mcriseffkd isigvsvftl talsgeryca ivnplrklqt
      181 kpltvftavm iwilaillgm psvlfsdiks ypvftatgnm tievcspfrd peyakfmvag
      241 kalvyyllpl siigalyimm akrlhmsarn mpgeqqsmqs rtqararlhv armvvafvvv
      301 fficffpyhv felwyhfypt aeedfdefwn vlrivgfcts flnscvnpva lycvsgvfrq
      361 hfnrylccic vkrqphlrqh statgmmdnt svmsmrrsty vggtagnlra slhrnsnhgv
      421 agagggvggg sgrvgsfyrq dsmplqhgna hgtgagvgss glgaggrtas vsekr
```

Serrata KAH8390260.1

```
1 vdtdvmdiaq alvaslahap ldgsgggggn gngsgandsv llatgtgaaa eqhnasidgg
       61 mvpyvpvldr petyivtvly tlifivgvlg ngtlviiffr hrsmrnipnt yilslaladl
      121 lvilvcvpva tivytqeswp fernmcrise ffkdisigvs vftltalsge rycaivnplr
      181 klqtkpltvf tavmiwilai llgmpsflvs diksypvlta tgnmtievcs pfrdpeyaqy
      241 mvaakafiyy llplsiigal yimmakrlhi sardmpgeqq smqsrtqara rrhvarmvva
      301 fvvvfficff pyhvfelwyh fyptaeedfd dfwnvlrivg fctsflnscv npvalycvsg
      361 vfrqhfnryl ccicvkrqph lrqhstatgm mdntsvmsmr rstyvggcgt ggnlraslhr
      421 nsnqgggggl gggagrggsf hrqdsmplqh gnghgaaggg aasgpscgpa gravavgek
```

Erecta XP_001970784.2

```
1 myaalmdvgq tlaarladge gndsgllatr qgleqeqeqe qeglalgmah nasadggmvp
     61 yvpvldrpet yivtvlytli fvvgvlgngt lviiffrhrs mrnipntyil slaladllvi
    121 lvcvpvativ ytqeswpfer nmcriseffk disigvsvft ltalsgeryc aivnplrklq
    181 tkpltvftav miwilaillg mpsvlfsdik sypvftatgn mtievcspfr dpeyakfmva
    241 gkalvyyllp lsiigalyim makrlhmsar nmpgeqqsmq srtqararlh varmvvafvv
    301 vfficffpyh vfelwyhfyp taeedfdefw nvlrivgfct sflnscvnpv alycvsgvfr
    361 qhfnrylcci cvkrqphlrq hstatgmmdn tsvmsmrrst yvggtagnlr aslhrssnhg
    421 vggaaggggg ggsgrvgsfh rqdsmplqhg nahgagaggg scglgsggrt apvceksfin
    481 ryesgrmry
```

Takahashi [XP_017003388.2]

```
  1 myaalmdvgq tlaaslaegg gnesgllatq egqgldletg hnasadggiv pyvpvldrpe
     61 tyivtvlytl ifivgvlgng tlviiffrhr smrnipntyi lslaladllv ilvcvpvati
    121 vytqeswpfe rnmcriseff kdisigvsvf tltalsgery caivnplrkl qtkpltvfta
    181 viiwilaill gmpsvlfsdi ksypvatpmg nitievcspf rdreyakfmv aakaliyyll
    241 plsiigalyi mmakrlhmsa rnmpgeqqsm qsrtqararr hvarmvvafv vvfficffpy
    301 hvfelwyhfy ptaeedfddf wnvlrivgfc tsflnscvnp valycvsgvf rqhfnrylcc
    361 icvkrqphlr qhstatgmmd ntsvmsmrrs tyvggavgcg pagnlrasqh rnsnhggggg
    421 sgrvgsfhrq dsmplqhgng hgtgigagvg gaacgsggra saavgekr
```

Biarmipes [XP_016966623.1]

```
  1 myvalvdvgq tlaagladga nesvlmatgh aaeqeqghwv gqnasgdgvm vpyvpvldrp
     61 etyivtvlyt lifivgvlgn gtlviiffrh rsmrnipnty ilslaladll vilvcvpvat
    121 ivytqeswpf ernmcrisef fkdisigvsv ftltalsger ycaivnplrk lqtkpltvft
    181 aaiiwilail lgmpsvlfsd iksypvltam gnmtievcsp frdaeyakfm vaakaliyyl
    241 lplsiigaly immakrlhms arnmpgeqqs mqsrtqarar rhvarmvvaf vvvfficffp
    301 yhvfelwyhf yptaeedfdd fwnvlrivgf ctsflnscvn pvalycvsgv frqhfnrylc
    361 cfcvkrqphv rqhstatgmm dntsvmsmrr styvggaggv ganlrasqhr nsslhrqast
    421 plqhgnghgt ggdetghgpg wvsggraaav gekr
```

Eugracilis [XP_017074910.2]

```
      1 myaslmdvgq tlaarladge gngandsgll atgqgleqeq eglaldmghn asadggivpy
     61 vpvldrpety ivtvlytlif ivgvlgngtl viiffrhrsm rnipntyils laladllvil
    121 vcvpvativy tqeswpfern mcriseffkd isigvsvftl talsgeryca ivnplrklqt
    181 kpltvftavm iwilaillgm psvlfsdiks ypvftatgnm tievcspfrd peyakfmvag
    241 kalvyyllpl siigalyimm akrlhmsarn mpgeqqsmqs rtqararlhv armvvafvvv
    301 fficffpyhv felwyhfypt aeedfdefwn vlrivgfcts flnscvnpva lycvsgvfrq
    361 hfnrylccic vkrqphlrqh statgmmdnt svmsmrrsty vggtagnlra slhrnsnhgv
    421 agagggvggg sgrvgsfyrq dsmplqhgna hgtgagvgss glgaggrtas vsekr
```

Rhopaloa [XP_044315908.1]

```
1 mfaalmdvgq tlagslaegn gsgvqgegle lgpgmgvevg qhnasadggi vpyvpvldrp
     61 etyivtvlyt lifivgvlgn gtlviiffrh rsmrnipnty ilslaladll vilvcvpvat
    121 ivytqeswpf ernmcrisef fkdisigvsv ftltalsger ycaivnplrk lqtkpltvft
    181 aviiwilail lgmpsflvsd iksypvvtvk gnitievcsp ysdpeyakym vaakatiyyl
    241 lplsiigaly immakrlhms arnmpgeqqs mqsrtqarar rhvarmvvaf vvvfficffp
    301 yhvfelwyhf yptaeedfdd fwnvlrivgf ctsflnscvn pvalycvsgv frqhfnrylc
    361 cicvkrqphl rqhstatgmm dntsvmsmrr styvggatag hlraslhrns nhggggggvg
    421 gagglgsgrv gsfhrqdsmp lqhgnghgsv aggaasgsgg ratavgeksl inryesdrir
    481 y
```

Fichsuphila [XP_017051484.2]

```
    1 mhaalmdvgq tlaggssand tgllatggeq eqeqgpglgl gmgmgmemgq hnasadggmv
     61 pyvpvldrpe tyivtvlytl ifivgvlgng tlviiffrhr smrnipntyi lslaladllv
    121 ilvcvpvati vytqeswpfe rnmcriseff kdisigvsvf tltalsgery caivnplrkl
    181 qtkpltvfta viiwilaill gmpsflvsdi ksypvltakg nmtievcspf rdpeyakcmv
    241 aakafiyyll plsiigalyi mmakrlhmsa rnmpgeqqsm qsrtqararr hvarmvvafv
    301 vvfficffpy hvfelwyhfy ptaeedfddf wnvlrivgfc tsflnscvnp valycvsgvf
    361 rqhfnrylcc icvkrqphlr qhstatgmmd ntsvmsmrrs tyvgvgggnl raslhrnsnh
    421 gvgaagggpg rxgsfhrqds mplqhgnghg avasaagcgs ggraaaggek r
```

Elegans [XP_041565851.1]

```
   1 mfaalmdvgq tlagslateg ndsgllatgq yeggvevgmg qhnssadggm vpyvpvldrp
```

```
  61 etyivtvlyt lifivgvlgn gtlviiffrh rsmrnipnty ilslaladll vilvcvpvat
 121 ivytqeswpf ernmcrisef fkdisigvsv ftltalsger ycaivnplrk lqtkpltvft
 181 aviiwilail lgmpsflvsd iksypvytan gnmtievcsp frdpeyakcm vaakafiyyl
 241 lplsiigaly immakrlhms arnmpgeqqs mqsrtqarar rhvarmvvaf vvvfficffp
 301 yhvfelwyhf yptaeedfdd fwnvlrivgf ctsflnscvn pvalycvsgv frqhfnrylc
 361 cicvkrqphl rqhstatgmm dntsvmsmrr styvggaaag qlraslhrns snhgggvgfg
 421 sgrvgsfhrq dsmalqhgng hgaatggasc apgsggrgpa ggekr
```

Kikkawei **KAH8308790.1**

```
   1 vntnkmdiaq alvaslahap ldgsgngsgs ggndsvllat gasaeqhnas idggmvpyvp
  61 vldrpetyiv tvlytlifiv gvlgngtlvi iffrhrsmrn ipntyilsla ladllvilvc
 121 vpvativytq eswpfernmc riseffkdis igvsvftlta lsgerycaiv nplrklqtkp
 181 ltvftavmiw ilaillgmps flvsdiksyp vltangnmti evcspyrdpe yaqcmvaaka
 241 fiyyllplsi igalyimmak rlhisardmp geqqsmqsrt qararrhvar mvvafvvvff
 301 icffpyhvfe lwyhfyptae edfddfwnvl rivgfctsfl nscvnpvaly cvsgvfrqhf
 361 nrylccicvk rqphlrqhst atgmmdntsv msmrrstyvg gcgaggnlra slhrnsnqgv
 421 ggggalgggt grvgsfhrqd smplqhgngh gaaagggtas gpscgsagra vavgek
```

Bipectinate **XP_017092331.2**

```
1 msalmdmsqt lalsllyapp datggvspni isigsgngsd gggndsslgl atgqqgaata
  61 vggvlglgqh nasadggggp yvpvlerpet yivtvlytli fivgvlgngt lviiffrhrs
 121 mrnipntyil slaladllvi lvcvpvativ ytqeswpfer nmcriseffk disigvsvft
 181 ltalsgeryc aivnplrklq tkpltvftaa miwvlaillg mpsflvsnik sytvltpngn
 241 msievcdpfr dpeyakymva akasiyyllp lsiigalyim makrlhisar dmpgeqqsmq
 301 srsqararrh varmvvafvv vfficffpyh vfelwyhfyp taeedfdefw nvlrivgfct
 361 sflnscvnpv alycvsgvfr qhfnrylccf cvkrqphlrq hstatgimdn tsvmsmrrst
 421 yvggaggnlr asmhrnsnhg ggggsglsag rgasfhggds mpmqhsnghg aaggtlggag
 481 vgatggpssg raaavgekr
```

Anannassae **XP_001961345.2**

```
1 msaalmdmsq tlalslvyap ldatggaspn iaigngndsn gggdggands glvlatgqqg
  61 vatvvggvlg lgqhngsadg gigpyvpvle rpetyivtvl ytlifivgvl gngtlviiff
 121 rhrsmrnipn tyilslalad llvilvcvpv ativytqesw pfernmcris effkdisigv
 181 svftltalsg erycaivnpl rklqtkpltv ftaamiwvla illgmpsflf sniksytvlt
 241 pngnmsievc dpyrdpeyak ymvaakasiy ylvplsiiga lyimmakrlh isardmpgeq
 301 qsmqsrsqar arryvarmvv afvvvfficl fpyyvfelwy hfyptaeedf defwnvlriv
 361 gfctsflnsc vnpvalycvs gvfrqhfnry lccfcvkrqp hlrqhstatg imdntsvmsm
 421 rrstyvggag gnlrssmhrn snhgggggag lpagrgasfh ggdsmplqhs nghgatggns
 481 ggagasatgg pslgrtaavg ekr
```

Mojavensis **XP_002006847.2**

```
1 myaalmdmsq mlaaslayap aesssaaaaa gvnlsqlsns sqldgssvvt vataatvgqh
  61 nasieessyv kvldrpetyi vtvlytlifi vgvlgngtlv iiffrhrsmr nipntyilsl
 121 aladllvilv cvpvativyt qeswpfernm criseffkdi sigvsvftlt alsgerycai
 181 vnplrklqtk pltvftavii wvfaimlgmp sfvvsdiqsy nittpngnit igvcspfrsk
 241 iyakymvvak asiyylvpls iigvlyimma krlhisardm pgeqlsiqsr sqararrhva
 301 rmvvafvvvf ficffpyhvf elwyhfypta eddfddfwnv vrivgfctsf lnscvnpval
 361 ycvsgvfrqh fnrylccicv krqphlrqhs tatgimdtsv tsmrrstyvg ggiaaggare
 421 ggprasvhmn nnhgvsgaag grggsfhrqd smalqhagsv nghshnanvg agagigrasi
 481 inekr
```

Virilis **XP_002049178.2**

```
1 mpknmlaalm dmsqtlaasl ayaplesnaa ataaaaaaaa vvlnvsqlgn ssqldgslat
  61 aaatttttav ttststhnas geeypqyvkv ldrpetyivt vlytlifivg vlgngtlvii
 121 ffrhrsmrni pntyilslal adllvilvcv pvativytqe swpfernmcr iteffkdisi
 181 gvsvftltal sgerycaivn plrklqtkpl tvftaviiwv faimlgmpsf vvsdiqgytl
 241 ptpngnitie vcspfrskiy akymvvakas iyylvplsii gvlyimmakr lhisardmpg
 301 eqlsiqsrsq ararrhvarm vvafvvvffi cffpyhvfel wyhfyptaee dfddfwhvvr
 361 ivgfctsfln scvnpvalyc vsgvfrqhfn rylccicvkr qphlrqhsta tgvmdtsvts
 421 mrrstyvggg gggavggsla ahraslhmnn nhgvavgggg ggggrggsfh rqdsmplqha
 481 gsgnghahnv ggpgagigra siinekslik rydertry
```

Grimshawi **XP_001985716.1**

```
1 myaalmdmsq tlaaslayap mesnsaaaaa aaaaaaalgm nvsqlsnssq ldgstgataa
```

```
 61 tsiphnvsae eypqyvkvld rpetyivtvl ytlifivgvl gngtlviiff rhrsmrnipn
121 tyilslalad llvilvcvpv ativytqesw pfernmcris effkdisigv svftltalsg
181 erycaivnpl rklqtkpltv ftaviiwvla imlgmpsfvv sdlqgytlpt nkgnitievc
241 spfrskiyak ymvvakasiy yfvplsiigv lyimmakrlh isardmpgeq lsiqsrsqar
301 arrhvarmvv afvvvfficf fpyhvfelwy hfyptaeddf ddfwhvvriv gfctsflnsc
361 vnpvalycvs gvfrqhfnry lccfcvkrqp hlrqhstatg vmdtsvtsmr rstyvggggg
421 vggatgslaa hraslhmnnn hgsgaggpgg rtgsfhrqds mplqhagsgn ghahnvggig
481 rasiineksl ikryeerary
```

"CCHa2-R PA isoform – data capture":
