## Supplementary material for "The incidence of candidate binding sites for β-arrestin in Drosophila neuropeptide GPCRs": S14 Text

***S14. Text*** **Multi-species analysis of LGR1**
**Supporting Figure 17**

CLUSTAL Line-ups; Genbank Reference IDs below
Predicted TM domains in YELLOW
BBS sequences in RED

```
Mojavensis         --------------------------MKCTPIIARFDFSAV--I-----------LLSLV        21
Virilis           --------------------------MKCMVIISQLDFLTF--I-----------LLSLA         21
Bipectinata        --------------------------MKCVLPKFVFRLLL-HHLLLTKLCWLHGVYATS-        32
Anannassae         --------------------------MKCVLPKFVYRLLL-HHLLLTRICWPHGVYATS-        32
Kikkawei           ------MGRRQTNWVRRKGRGQPKTALTCLSIGFLFRFLFVYHLLLAPLCGPHCVYAMLA        54
Serrata            ------MGRRRTHWVRKGDRRPVKTGIKCLSIGYLLRLLFVHHLLLGSFCGSHRVYAMLA        54
Ficusphelia        MKKH-------TNLHRKEVTWRSRRGTKCLTFEFLFRLLY-NHLLLTSMFGPDCVYATSA        52
Erecta             MEKHPSLPPSQTQTQRRDTTCRSKKGLKCLSFEFQCRLLL-HHLLLTSLSGRHCVYAM--        57
Melanogaster       MEKHPSLS------QRMGTTYRPRKGLKCLSFEFQCRLLL-HHLLLTSLSGRHFVYATSA        53
Sechellia          MEKHPSMS------QRLETTCRPRKGLKCLSFEFQCRLLL-HHLLLTSLSGRHCVYATSA        53
Mauritania         MEKHPSLS------QRMDTTCRPRKGLKCLSFEFQCRLLL-HHLLLTSLSGRHCVYATSA        53
Simulans           MEKHPSLS------QRMETTCRPRKGLKCLSFEFQCRLLL-HHLLLTSLSGRHCVYATSA        53
Eugracilis         MEKHTNLC-------RTESTWRPKKGLKCLSFEYLFRLLL-HHLLLTSLSGPHCVYATSD        52
Takahashi          MEKHTSLP-------RKEASESSRKGLKCLSLEFLFRLLF-HQLLFTSLCGPHCVYATSA        52
Suzuki             MEKHTSQY-------PKESPWRTKRGLKCLTFEFQFRLLL-HHLMLSSLCGPHCVYATSA        52
Biarmipes          MEKHTSQC-------PKEAPWRPKRGLKCLTIEFQFRLLL-HHLLLSSLCGPHCVYATSA        52
Elegans            MEKPAILAR----KDVLRRSRKCSKGLKCLSFDFQFRLLL-HYLLLSSLSGPHCVYATMA        55
Rhopaloa           MEKYTILAR----KRAR---RPSKRILKCLSLEFLYRLLL-HHLLLTSLCDHHYVYATSA        52
                                              .*        :     :

Mojavensis         YCSQQASTNCYDNHDGFNAIINNLPRDAGNA-----------IDTLDTLAQAPPMTTNPP        70
Virilis            HGTRQANTNCHDNNNGFNTVFNNLSIDNGNE-----------TDMSVIMTQAPTMSTNPT        70
Bipectinata        -------ASCHDSYNGFNVFPNGSSDLRP---DP------NPTDVSSGQAP------PPT        70
Anannassae         -------AVCHDSHNGFNVSPDGNSDLSP---DR------NPTTMSSGQAP------PPT        70
Kikkawei           AEGPSDGSNCHDIHHGFDVPSSLNAVT-------------------EAQGGDAPMAVTLT        95
Serrata            AEGQSVRSNSHDVHHGFDVPGRQP-VS-------------------DNQVRGAPITVILP        94
Ficusphelia        VGKALSASNCHDIHHGFDVFPNPNPTPALNAVDSNPNTSSNATAMWLGQSTDTP--VTLT        110
Erecta             -----SASSCHDIHHGFDVYPNPNPT-----------------TVSLGQSTDTP-Q-AEA        93
Melanogaster       VGGALSANNCHDIHHGFDVYPNL--T-----------------AVSLAQSTDTP-L-TAT        92
Sechellia          AGGALSASNCHDIHHGFDVYPNP--T-----------------AVSLAQSTDTP-L-TAA        92
Mauritania         AGGALSASNCHDIHHGFDVYPNP--T-----------------AVSLAQSTDTP-L-TAA        92
Simulans           AGGALSASNCHDIHHGFDVYPNP--T-----------------AVSLAQSTDTP-L-TAA        92
Eugracilis         IGEALSASNCHDIHHGFDVYPNPNAL-------------------SLGQSTDTP--LTQT        91
Takahashi          VGEALGASNCHDILHGFDVYPNPTAG-------------------ALGQSTDSP--LTVT        91
Suzuki             VGEALSASNCHDIHHGFDVYPNPNPSP-------------NLTAVTLGQSTDTP-SVTLT        98
Biarmipes          VGEALSASNCHDIHHGFDVYPNPNLNPTS---------SANGTTVTLGQSRDTP-SVPLT        102
Elegans            GSEALSASNCHDIHHGFDVYPNPTAV-------------------SQGQSTDTP--LTLT        94
Rhopaloa           VGEPISASDCHDIHHGFDVFPNHTAL-------------------PLGQSTDTP--LTST        91
                            .:*  .**:.

Mojavensis         VDASVWKCCCWEATNHN-EFECRCEGEALTRVPQTLKLPLQRLTIASAGLPRLRSMGLKV        129
Virilis            SDASVWKCCCWEATNQN-EFECRCEGEALTRVPQTLKLPLLRLTIASAGLPRLRSMGLKV        129
Bipectinata        LTLPPWKCCCWNATNQSEEAECRCEGEALTRVPQTLTMAMQRLTIATAGLPRLRATGLKV        130
Anannassae         LTLPPWKCCCWNATNQSEEVECRCEGEALTRVPQTLTMAMQRLTIASAGLPRLRATGLKV        130
Kikkawei           PAPSGWKCCCWKAPNQGEEVECRCEGDALTRVPQTLQLPIQRLTIASAGLPRLRFTGLKV        155
Serrata            KVTSRWKCFCSKTPNQSEELECRCEGDSLTRVPQTLQLPIQRLTIASAGLPRLRSTGLKV        154
Ficusphelia        LPPSAWKCCCWNASNQNEEVECRCEGDGLNRVPQTLKLPLQRLTIASAGLPRLRYTGLKV        170
Erecta             MPPSAWKCCCWNASNQVEEVECRCEGDGLNRVPQTLKLPIQRLTIASAGLPRLRHTGLKV        153
Melanogaster       MPRSAWKCCCWNASNQAEEVECRCEGDGLNRVPQTLTLPIQRLTIASAGLPRLRHTGLKV        152
Sechellia          MPRSAWKCCCWNASNQAEEVECRCEGDGLNRVPQTLTLPIQRLTIASAGLPRLRHTGLKV        152
Mauritania         MPRSAWKCCCWNASNQAEEVECRCEGDGLNRVPQTLTLPIQRLTIASAGLPRLRHTGLKV        152
Simulans           MPRSAWKCCCWNASNQAEEVECRCEGDGLNRVPQTLTLPIQRLTIASAGLPRLRHTGLKV        152
Eugracilis         LPPSAWKCCCWNASNQMEEVECRCEGDGLNRVPQTLKLPIQRLTIASAGLPRLRYTGLKV        151
Takahashi          LPPSGWKCCCWNASNQMEEVECRCEGDGLNRVPQTLKLPIQRLTIASAGLPRLRYTGLKV        151
Suzuki             LPPSGWKCCCWNASNQAEEVECRCEGDGLNRVPQTLKLPIQRLTIASAGLPRLRYTGLKV        158
Biarmipes          LPPSGWKCCCWNGGNQAEEVECRCEGDGLNRVPQTLKLPLQRLTIASAGLPRLRYTGLKV        162
Elegans            LPPSGWKCCCWNASNQNEEVECRCGGDGLNRVPQTLKRPIQRLTIASAGLPRLRNTGLKV        154
Rhopaloa           LPPSGWKCCCWKSSNQNEEVECRCEGDGLNRVPQSLKLPIQRLTIASAGLPRVRFTGLKV        151
                        *** * :  *:  * **** *:.*.****:*   : *****:*****:*  ****
```

```
Mojavensis        YAPTLLDVAFIDCLQLETIQSGAFSNLTVLRAIYISNAPKLSYLAKNVFEGISDTIEIIR    189
Virilis           YATTLLDVAFIDCLQLEAIQNGAFSNLTFLRTIYISNAPKLTYLPKNVFEGISDTIEIIR    189
Bipectinata       YAQTLMDVAFTDCLQLELIQDGAFANLKLLRTIYIANAPKLTFLSKDVFFGISDTVEIIR    190
Anannassae        YAQTLLDVAFTDCLQLELIQDGAFANLKLLRTIYIANAPKLSFLSKDVFSGISDTVEIIR    190
Kikkawei          YGPTLLDVAFTDCLKLELIQDGAFANLTLLRTIYISNAPKLTFLSKDVFAGISETVEIIR    215
Serrata           YGPTLLDVAFTDCLKLELIQDGAFANLTMLRTIYISNAPKLTFLSKDVFAGISETVEVIR    214
Ficusphelia       YGPTLLDVTFTDCLQLELIQDGAFANLTLLRTIYISNAPKLTFLSKDVFAGISESVEIIR    230
Erecta            YGPTLLDVAFTDCLQLELIQDGAFANLTLLRTIYITNAPKLTFLSKDVFLGISDTVEIIR    213
Melanogaster      YGSTLLDVAFTDCLQLELIQDGAFANLTLLRTIYITNAPKLTFLSKDVFLGISDTVDIIR    212
Sechellia         YGSTLLDVAFTDCLQLELIQDGAFANLTLLRTIYITNAPKLTFLSRDVFLGISDTVDIIR    212
Mauritania        YGSTLLDVAFTDCLQLELIQDGAFANLTLLRTIYITNAPKLTFLSKDVFLGISDTVDIIR    212
Simulans          YGSTLLDVAFTDCLQLELIQDGAFANLTLLRTIYITNAPKLTFLSKDVFLGISDTVDIIR    212
Eugracilis        YGPTLLDVAFTDCQQLELIQDGAFSNLTLLRTIYITNAPKLTFLSKDVFLGISDTVEIIR    211
Takahashi         YGPTLLDVTFTDCLQLELIQDGAFANLTLLRTIYITNAPKLTFLSKDVFFGISDTVEIIR    211
Suzuki            YGPTLLDVAFTDCLQLELIQDGAFANLTLLRTIYITNAPKLTFLSKDVFFGISGTVEIIR    218
Biarmipes         YGATLLDVAFTDCLQLELIQDGAFANLTLLRTIYITNAPKLTFLSKDVFLGISGTVEVIR    222
Elegans           YGSTLLDVAFTDCLQLELIQDGAFANLTLLRTIYISNAPKLTFLSKDVFFGISGSVEIIR    214
Rhopaloa          YGSSLLDVAFIDCLQLELIQDGAFANLALLRTIYISNAPKLTFLSKDVFSGISGSVEIIR    211
                  *. :*:**:* ** :** **.***:** .**:***:*****::* ::** *** ::::**

Mojavensis        IINSGLKTVPDLGDLPPYNILQMIDLDNNQISRIDSKSIQVKTAQLVLANNEITFIDDSA    249
Virilis           IINSGLTSVPDFGYLPPNNILQMIDLDNNQISRIDSKSIQVKTAQFVLANNDIHFIDDSA    249
Bipectinata       IINSGLTRVPDLTHLPPYNILQMIDLDNNQITRIDAKSIKVKTAQLILANNDISYVDDSA    250
Anannassae        IINSGLTSVPDLTHLPPYNILQMIDLDNNQITRIDSKSIKVKTAQLILANNDISYVDDSA    250
Kikkawei          IINSGLTSVPDLGHLPPHNILQMIDLDNNQITRIDTKSINVKTAQFILANNDISYVDDSA    275
Serrata           IINSGLTRVPDLGHLPPHNILQMIDLDNNQITRIDTKSINVKTAQLILANNDISYVDDSA    274
Ficusphelia       IINSGLTRVPDLGHLPPHNILQMIDLDNNQISRIDTKSIKVKTAQLILANNDISYVDDSA    290
Erecta            IINSGLTRVPDLGHLPPHNILQMIDLDNNQITRIDSKSIKVKTAQLILANNEISYVDDSA    273
Melanogaster      IINSGLTRVPDLGHLPPHNILQMIDLDNNQITRIDSKSIKVKTAQLILTNNEISYVDDSA    272
Sechellia         IINSGLTRVPDLGHLPPHNILQMIDLDNNQITRIDSKSIKVKTAQLILTNNEISYVDDSA    272
Mauritania        IINSGLTRVPDLGHLPPHNILQMIDLDNNQITRIDSKSIKVKTAQLILTNNEISYVDDSA    272
Simulans          IINSGLTRVPDLGHLPPHNILQMIDLDNNQITRIDSKSIKVKTAQLILTNNEISYVDDSA    272
Eugracilis        IINSGLTRVPDLSHLPPNNILQMIDLDNNQITRIDTKSIKVKTAQLILANNEISYIDDSA    271
Takahashi         IINSGLTRVPDLSHLPPHNILQMIDLDNNQITRIDTKSIKVKTAQLILANNDISYVDDSA    271
Suzuki            IINSGLTRVPDLSHLPPHNILQMIDLDNNQITRIDTKSIKVKTAQLILANNDISYVDDSA    278
Biarmipes         IINSGLTRVPDLSHLPPHNILQMIDLDNNQITRIDTKSIKVKTAQLILANNDISYVDDSA    282
Elegans           IINSGLTRVPDLGHLPPHNILQMIDLDNNQITRIDTKSIKVKTAQLILANNDISYVDDSA    274
Rhopaloa          IINSGLTRVPDLGHLPPHNILQMIDLDNNQITRIDTKSINVKTAQLILANNDISYVDDSA    271
                  ******. ***:  *** *************:***:***:*****::*:**:* ::****

Mojavensis        FLGSKIAKLSLNDNHKLTEIHPNAFNGIIDMTELDLSSTSLVRLPSAGLQTLEVLYIANT    309
Virilis           FLGSKIAKLSLKDNRRLTDVHPNAFYGIIDITELDLSSTSLVSLPSAGLQTVEVLYIINT    309
Bipectinata       FCGSKIAKLSLKENRKLKELHTNAFHGIIDITELDLSSTSIVEMPSAGLQTIEALYILNT    310
Anannassae        FFGSKIAKLSLKENRKLKELHTNAFHGIIDITELDLSSTSIVEMPSAGLQTIEALYILNT    310
Kikkawei          FFGSKIAKLSLKDNRKLTDMHPNAFNGIIDIAELDLSSTSLVGLPSVGLQTIEALYIMNT    335
Serrata           FFGSKIAKLSLKDNWKLTEMHPEAFNGIIDITELDLSSTSLVGLPSVGLQTIEALYIMNT    334
Ficusphelia       FFGSKIAKLSLQNNRKLKQMHPNAFDGIIDITELDLSSTSLVGLPSVGLQNIEALYITNT    350
Erecta            FFGSKIAKLSLKENKRLKMMHPNAFDGIIDIAELDLSSTSLVGLPSAGLQNIEALYIQNT    333
Melanogaster      FFGSKIAKLSLKENKKLQMMHPNAFDGIIDITELDLSSTSLVGLPSAGLQNIEALYIQNT    332
Sechellia         FYGSKIAKLSLKENKKLEMMHPNAFDGIIDIAELDLSSTSLVGLPSAGLQNIEALYIQNT    332
Mauritania        FFGSKIAKLSLKENKKLQMMHPNAFDGIIDIAELDLSSTSLVGLPSAGLQNIEALYIQNT    332
Simulans          FFGSKIAKLSLKENKKLQEMHPNAFDGIIDIAELDLSSTSLVGLPSAGLQNIEALYIQNT    332
Eugracilis        FFGSKIAKLSLKDNLKLEMMHPKAFDGIIDITELDLSSTSLVGLPSVGLQTIEALYIQNT    331
Takahashi         FFGSKIAKLSLNDNQKLEHMHPNAFDGIIDITELDLSSTSLVGLPSSGLQTIEALYIQNT    331
Suzuki            FFGSKIAKLSLNDNLRLEHMHPNAFDGI-DITELDLSSTSLVAMPSVGLQNIEALYIQNT    337
Biarmipes         FYGSKIAKLSLNDNLRLEHMHPNAFDGIIDITELDLSSTSLVAMPSVGLQNIEALYIQNT    342
Elegans           FFGSKIAKLSLKDNPKLKELHPNAFDGIIDITELDLSSTSLVGMPSVGLQTIEALYIQNT    334
Rhopaloa          FFGSKIAKLSLNDNPKLKELHPNAFNGIIDITALDLSSTSLIGMPSAGLQTIEALYIQNT    331
                  * *********::* :*  :* :** ** *:: *******:: :** ***.:*.*** **

Mojavensis        HTLKTIPSIYNFQNLQRAHLTHSFHCCAFQFPSRHDPQRHAQRMQEVEKWRDQCKNNRDV    369
Virilis           HTLKTIPSIYNFQNLQRAHLTHSFHCCAFQFPSRHDPKRHAERLQELQKWREQCNVERDL    369
Bipectinata       HTLKTIPSIYNFRNLQRAYLTHSFHCCAFQFPSRHDPLRHAQRMDEIEKWRTQCKGERIS    370
Anannassae        HTLKTIPSIYNFRNLQRAYLTHSFHCCAFQFPSRHDPLRHAQRMLEIEKWRTQCNGERNS    370
Kikkawei          HTLKTIPSIYNFRNLQRAYLTHSFHCCAFQFPSRHDPRRHAQRMLEIEKWREQCNK--GS    393
Serrata           HTLKTIPSIYNFRNLQRAYLTHSFHCCAFQLPSRHDPRRHALRMLEIEKWREQCNK--CS    392
Ficusphelia       HTLKTIPSIYNFRNLQRAYLTHSFHCCAFQFPSRHDPQRHAERMREIEKWREQCKSDHGS    410
Erecta            HTLKTIPSIYNFRNLQRAYLTHSFHCCAFQFPSRHDPQRHAQRMLEIEKWRKQCKSESGS    393
Melanogaster      HTLKTIPSIYNFRNLQRAYLTHSFHCCAFQFPSRHDPQRHAQRMLEIEKWRKQCKSDSGT    392
Sechellia         HTLKTIPSIYNFRNLQRAYLTHSFHCCAFQFPSRHDPQRHAQRMLEIEKWRKQCKSDSGS    392
```

```
Mauritania        HTLKTIPSIYNFRNLQRAYLTHSFHCCAFQFPSRHDPQRHAQRMLEIEKWRKQCKSDSGS        392
Simulans          HTLKTIPSIYNFRNLQRAYLTHSFHCCAFQFPSRHDPQRHAQRMLEIEKWRKQCKSDSGS        392
Eugracilis        HTLKTIPSIYNFRNLQRAYLTHSFHCCAFQFPSRHDPQRHAQRMLEIEKWRKQCKSDSGS        391
Takahashi         HTLKTIPSIYNFRNLQRAYLTHSFHCCAFQFPSRHDPQRHAKRMREIEKWREQCKSDSAS        391
Suzuki            HTLKTIPSIYNFRNLQRAYLTHSFHCCAFQFPSRHDPQRHAQRMLEIEKWRKQCKSESVS        397
Biarmipes         HTLKTIPSIYNFRNLQRAHLTHSFHCCAFQFPSRHDPQRHAERMLEIEKWRKQCKSDSVT        402
Elegans           HTLKTIPSIYNFRNLQRAYLTHSFHCCAFQFPSRHDPQRHAQRMLEIQKWREQCKSDHVS        394
Rhopaloa          HTLKTIPSIYNFRNLQRAYLTHSFHCCAFQFPSRHDPERHAQRMLEIEKWRKQCKSDQVS        391
                  ************:*****:***********:****** *** *: *::*** **:

Mojavensis        HTNYERSLSMS-NLTPLANVSEVQPSGSTINNLLAESTPNTYAYMADSTLNNIGIFHEEI        428
Virilis           YRDVDKKLKKNTSVKDTAGVHYTNQSGTATDNLLTDAASNSYDYMADSTMNNIGIFHEQI        429
Bipectinata       RKERSILNNLKSQSDGYGIFG-SEASPT----DVVSTQFASVDYMADAAMN-LGYFHEEI        424
Anannassae        RRERNIAKKIESQQDDLGTLGGSEALSA----DVLSTPFSSVDYMADAAMN-LGYFHEEI        425
Kikkawei          RKERSLLDNFEAQPEDFGSFGSTEPSVT----EITPYPYASVDYMADS-TN-LGYFHEQI        447
Serrata           RKERALLDYLEAQPEDFGSFGSTEPSMT----ENTPFPIASIDYMADS-TN-LGYFHEQI        446
Ficusphelia       RKERSALDGFEGLPEDFGTFGMPEQSGT----DDTSITYASFDYMSDDTLN-KGTFHEKI        465
Erecta            RQERSTLDSSLSMPEDFGTFSGTDDSAT----DITPITFVSPDYMADDTMN-KGTFHEKI        448
Melanogaster      RKERSTLDNPFNMPEDFGSFGGTDDSAT----DITPITFASFDYMADDTMN-KGTFHEKI        447
Sechellia         RKERSTLDNSFSMPEDFGSFGGTDDSST----DITPITFASFDYMADDTMN-KGTFHEKI        447
Mauritania        RKERSTLDNSFDMPEDFGSFGGTDDSAI----DITPITFASFDYMADDTMN-KGTFHEKI        447
Simulans          RKERSTLDNSFNMPEDFGSFGGTDDSAT----DMTPITFASFDYMADDTMN-KGTFHEKI        447
Eugracilis        RKERSTLDGLDNQPEDFGSFGGSYQSAT----DISPITFSSIDYMADDTMN-KGTFHEKI        446
Takahashi         RKERSTSENGFEMPEDFGSFGSPDEATT----DIVPIKFASFDYMADDTLN-KGTFHEKI        446
Suzuki            RQERSTLDNAYTMPEDFGTFESTDESST----DLVPITFASFDYMADDSLN-KGTFHEKI        452
Biarmipes         RKERSTLENGFTLPEDFGTFGGTDDSST----DLVPITFASFDYMADDTLI-KGTFHEKI        457
Elegans           RKERSTVDNLNSMPEEFGSFGSTDQSAT----DMTSMTYASFDYMSDDTMN-KGTFHEKI        449
Rhopaloa          RKERSIADNLNSVPEEFGSFGMTDQ---------------SLDYMSDDTMN-KGTFHEKI        435
                    :              . .                    :  **:*      * ***:*

Mojavensis        TINPDDDQLAEYCGNFTFRKPHVECYPMPNALNPCEDVMGYQWLRIAVWIVVALAIVGNV        488
Virilis           TINPDDNQLAEYCGNFTFRNPDIQCFPMPNALNPCEDVMGYQWLRISVWIVVALAIVGNL        489
Bipectinata       TLNPDDEQFAEFCGNFTFRKPNVECYPMPNALNPCEDVMGYQWLRISVWVVVALAVVGNV        484
Anannassae        TINPDDEQFAEFCGNFTFRKPSVECYPMPDALNPCEDVMGYQWLRISVWVVVALAVVGNV        485
Kikkawei          TINPDDDQSAEFCGNFTFRKPNIECYPMPNDLNPCEDVMGYQWLRISVWIVVALAVVGNV        507
Serrata           TINPDDKQSAEFCGNFTFRKPNIECYPMPNDLNPCEDVMGYQWLRISVWIVVALSAVGNV        506
Ficusphelia       TLNPEDDMTAELCGNFTFRKPNIECYPMPNDLNPCEDVMGYQWLRISVWIVVALAVVGNV        525
Erecta            VLNPEDDSSAELCGNFTFRKPNIECYPMPNDLNPCEDVMGYQWLRISVWIVVALAVVGNV        508
Melanogaster      ILNPGDDSSAELCGNFTFRKPNIECYPMPNDLNPCEDVMGYQWLRISVWIVVALAVVGNV        507
Sechellia         ILNPGDDSSAELCGNFTFRKPNIECYPMPNDLNPCEDVMGYQWLRISVWIVVALAVVGNV        507
Mauritania        ILNPGDDSSAELCGNFTFRKPNIECYPMPNDLNPCEDVMGYQWLRISVWIVVALAVVGNV        507
Simulans          ILNPDDDSSAELCGNFTFRKPNIECYPMPNDLNPCEDVMGYQWLRISVWIVVALAVVGNV        507
Eugracilis        TLNPEDDMSAELCGNFTFRKPNIECYPMPNDLNPCEDVMGYQWLRISVWIVVALAVVGNV        506
Takahashi         TLDPADDMSAELCGNFTFRKPNIECYPMPNDLNPCEDVMGYQWLRISVWIVVALAVVGNV        506
Suzuki            TLNPEDDMSAELCGNFTFRKPNIECYPMPNDLNPCEDVMGYQWLRISVWIVVALAVVGNV        512
Biarmipes         ILNPEDDMSAELCGNFTFRKPNIECYPMPNDLNPCEDVMGYQWLRISVWIVVALAVVGNV        517
Elegans           TLNPEDDI-DELCGNFTFRKPNIECYPMPNDLNPCEDVMGYQWLRISVWIVVALAVVGNV        508
Rhopaloa          TLNPEDDI-DELCGNFTFRKPNIECYPMPNDLNPCEDVMGYQWLRISVWIVVALAVVGNV        494
                   ::* *.   * *******:* ::*:***: ***************:**:****:  ***:

Mojavensis        AVLTVILSIKSESPSVPRFLICHLAFADLCLGLYLLLIASIDAHSMGEYFNYAFDWQYGL        548
Virilis           AVLTVTLSIKSESPSVPRFLICHLAFADLCLGLYLLLIASIDAHSMGEYFNYAFDWQYGL        549
Bipectinata       AVLTVNLSIRPEITPVARFLMCHLAFADLCLGLYLFLVASIDAHSMGEYFNYAYDWQYGL        544
Anannassae        AVLTVNLSIRPEITPVARFLMCHLAFADLCLGLYLFLVASIDAHSMGEYFNYAYDWQYGL        545
Kikkawei          AVLTVILSIRPESTPVPRFLMCHLAFADLCLGVYLLLVASIDAHSMGEYFNYAYDWQYGL        567
Serrata           AVLTVILSIRPESSPVPRFLMCHLAFADLCLGVYLLLVASIDAHSIGEYFNYAYDWQYGL        566
Ficusphelia       AVLTVILSIRPESTPVPRFLMCHLAFADLCLGVYLLLVASIDAHSMGEYFNYAYDWQYGL        585
Erecta            AVLTVILSIRPESTPVPRFLMCHLAFADLCLGLYLLLVACIDAHSMGEYFNFAYDWQYGL        568
Melanogaster      AVLTVILSIRPESTPVPRFLMCHLAFADLCLGLYLLLVACIDAHSMGEYFNFAYDWQYGL        567
Sechellia         AVLTVILSIRPESTPVPRFLMCHLAFADLCLGLYLLLVACIDAHSMGEYFNFAYDWQYGL        567
Mauritania        AVLTVILSIRPESTPVPRFLMCHLAFADLCLGLYLLLVACIDAHSMGEYFNFAYDWQYGL        567
Simulans          AVLTVILSIRAESTPVPRFLMCHLAFADLCLGLYLLLVACIDAHSMGEYFNFAYDWQYGL        567
Eugracilis        AVLTVILSIRPETTPVPRFLMCHLAFADLCLGVYLLLVASIDAHSKGEYFNFAYDWQYGL        566
Takahashi         AVLTVILSIRPESTPVPRFLMCHLAFADLCLGVYLLLVASIDAHSMGEYFNYAYDWQYGL        566
Suzuki            AVLTVILSIRPESTPVPRFLMCHLAFADLCLGVYLLLVASIDAHSMGEYFNYAYDWQYGL        572
Biarmipes         AVLTVILSIRPESTPVPRFLMCHLAFADLCLGVYLLLVASIDAHSMGEYFNYAYDWQYGL        577
Elegans           AVLTVILSIRPESTPVPRFLMCHLAFADLCLGVYLLFVASIDAHSMGEYFNFAYDWQYGL        568
Rhopaloa          AVLTVILSIRPESTPVPRFLMCHLAFADLCLGVYLLLVASIDAHSMGAYFNYAYDWQYGL        554
                  ***** ***: *   * ***:***********:**:::*.***** * ***:*:******
```

```
Mojavensis      GCKIAGFLTVFASHLSVFTLTVITIERWFAITHAMYLNKRIKLRPAAVIMLMGWIYSIVM     608
Virilis         GCKIAGFLTVFASHLSIFTLTIITIERWFAITHAMYLNKRITLRQAAGIMLTGWIYSIIM     609
Bipectinata     GCKVAGFLTVFASHLSVFTLTLITIERWVAITQAMYLNKRIRLRSASIIMLGGWIYSMVM     604
Anannassae      GCKVAGFLTVFASHLSVFTLTLITIERWVAITQAMYLNKRIRLRSASIIMLGGWIYSMVM     605
Kikkawei        GCKAAGFLTVFASHLSVFTLTVITIERWLAITQAMYLNHRIKLRQAALIMLGGWLYSMFM     627
Serrata         GCKVAGFLTVFASHLSVFTLTVITIERWLAITQAMYLNHRIKMRQAALIMLGGWLYSMVM     626
Ficusphelia     GCKVAGFLTVFASHLSVFTLTVITIERCMAITQAMYLNHRIKLRPAAIIMLCGWIYSMLM     645
Erecta          GCKVAGFLTVFASHLSVFTLTVITIERWLAITQAMYLNHRIKLRPAALIMLGGWIYSMLM     628
Melanogaster    GCKVAGFLTVFASHLSVFTLTVITIERWLAITQAMYLNHRIKLRPAALIMLGGWIYSMLM     627
Sechellia       GCKVAGFLTVFASHLSVFTLTVITIERWLAITQAMYLNHRIKLRPAALIMLGGWIYSMLM     627
Mauritania      GCKVAGFLTVFASHLSVFTLTVITIERWLAITQAMYLNHRIKLRPAALIMLGGWIYSMLM     627
Simulans        GCKVAGFLTVFASHLSVFTLTVITIERWLAITQAMYLNHRIKLRPAALIMLGGWIYSMLM     627
Eugracilis      GCKVAGFLTVFASHLSVFTLTVITIERWLAITQAMYLNHRIKLRPAALIMLGGWIYSMFM     626
Takahashi       GCKVAGFLTVFASHLSVFTLTVITIERWLAITQAMYLNHRIKLRPAALIMLGGWIYSMFM     626
Suzuki          GCKAAGFLTVFASHLSVFTLTVITIERWLAITQAMYLNHRIKLRPAALIMLGGWIYSMFM     632
Biarmipes       GCKVAGFLTVFASHLSVFTLTVITIERWLAITQAMYLNHRIKLRPAALIMLGGWIYSMFM     637
Elegans         GCKVAGFLTVFASHLSVFTLTVITIERWLAITQAMYLTHRIKLRQASIIMLGGWIYSVLM     628
Rhopaloa        GCKVAGFLTVFASLLSVFTLTVITIERWLAITQAMYLTHRIKLRPASFIMLGGWIYSILM     614
                *** ********* **:****:***** .***:****.:** :* *: *** **:**:.*

Mojavensis      SSLPLLGISNYSSTSICLPMEKRDIYDSIYLVLILGCNFLAFTIIAICYSQIYLSLGKET     668
Virilis         SSLPLFGISNYSSTSICLPMEIRDIYDSIYLILILGCNFVAFTIIAICYSQIYLSLGQET     669
Bipectinata     SSLPLFGISNYSSTSICLPMEVRDSFDTVYLIAILGCNGVAFFIIAVCYAKIYLSLGRET     664
Anannassae      SSLPLFGISNYSSTSICLPMEVRDTFDTVYLIGILGCNGVAFFIIAVCYAKIYFSLGRET     665
Kikkawei        SSLPLFGISNYSSTSICLPMENRDAFDTMYLIAILGCNGVAFSIIAVCYAQIYLSLGRET     687
Serrata         SSLPLFGISNYSSTSICLPMENRDAFDTMYLIAILGCNGVAFSIIAVCYAQIYLSLGRET     686
Ficusphelia     SSLPLFGISNYSSTSICLPMENRDVFDTMYLIAILGCNGVAFSIIAVCYAQIYLSLGRET     705
Erecta          SSLPLFGISNYSSTSICLPMENRDVYDTVYLIAILGCNGVAFSIIAVCYAQIYLSLGRET     688
Melanogaster    SSLPLFGISNYSSTSICLPMENRDVYDTIYLIAILGSNGVAFSIIAVCYAQIYLSLGRET     687
Sechellia       SSMPLFGISNYSSTSICLPMENRDVYDTIYLIAILGSNGVAFSIIAVCYAQIYLSLGRET     687
Mauritania      SSMPLFGISNYSSTSICLPMENRDVYDTIYLIAILGSNGVAFSIIAVCYAQIYLSLGRET     687
Simulans        SSMPLFGISNYSSTSICLPMENRDVYDTIYLIAILGSNGVAFSIIAVCYAQIYLSLGRET     687
Eugracilis      SSMPLFGISNYSSTSICLPMENRDVYDTVYLVAIMVCNGVAFSIIAVCYAKIYLSLGRET     686
Takahashi       SSLPLFGISNYSSTSICLPMENRDIYDTVYLIAIMVCNGVAFSIIAVCYAQIYLSLGRET     686
Suzuki          SSLPLFGISNYSSTSICLPMENRDVYDTVYLIAIMVCNGVAFSIIAVCYAQIYLSLGRET     692
Biarmipes       SSLPLFGISNYSSTSICLPMENRDGYDTVYLIAIMVCNGVAFSIIAVCYAQIYLSLGRET     697
Elegans         SSLPLFGISNYSSTSICLPMEIRDGFDTAYLIAILACNGVAFTIIAVCYAQIYLSLGRET     688
Rhopaloa        SSLPLFGISNYSSTSICLPMEIRDVFDTVYLIAILACNGVAFSIIAVCYAQIYLSLGRET     674
                **:**:*************** ** :*: **: *: .* :** ***:**::**:***:**

Mojavensis      RRARQNHLGEMSVAKKMALLVFINFSCGAPIAFFGLTALAGYPLINVTKSKILLVFFYPL     728
Virilis         RRARRNNPGEMSVAKKMLLLVFINFTCGAPIAFFGLTALAGCPLINVTKSKILLVFFYPL     729
Bipectinata     RHARQNNPGELSVAKKMALLVFTNFACWSPIAFFGLTALAGFPLINVTNSKILLVFFYPL     724
Anannassae      RHARQNNPGELSVAKKMSLLVFTNFACWSPIAFFGLTALAGYPLINVTNSKILLVFFYPL     725
Kikkawei        RQARQNNPGELSVAKKMALLVFTNFACWSPIAFFGLTALAGFPLINVTKSKILLVFFYPL     747
Serrata         RQAHQNNPGELSVAKKMALLVFTNFACWSPIAFFGLTALAGFPLINVTKSKILLVFFYPL     746
Ficusphelia     RQARHNNPGELSVAKKMALLVFTNFACWSPIAFFGLTALAGYPLINVTKSKILLVFFYPL     765
Erecta          RQAHQNNPGELSVAKKMALLVFTNFACWSPIAFFGLTALAGYPLINVTKSKILLVFFYPL     748
Melanogaster    RQAHQNSPGELSVAKKMALLVFTNFACWSPIAFFGLTALAGYPLINVTKSKILLVFFYPL     747
Sechellia       RQAHQNNPGELSVAKKMALLVFTNFACWSPIAFFGLTALAGYPLINVTKSKILLVFFYPL     747
Mauritania      RQTHQNNPGELSVAKKMALLVFTNFACWSPIAFFGLTALAGYPLINVTKSKILLVFFYPL     747
Simulans        RQAHQNNPGELSVAKKMALLVFTNFACWSPIAFFGLTALAGYPLINVTKSKILLVFFYPL     747
Eugracilis      RQARHNNPGELSVAKKMALLVFTNFACWSPIAFFGLTALAGYPLINVTKSKILLVFFYPL     746
Takahashi       RQAHHNSPGELSVAKKMALLVFTNFACWSPIAFFGLTALAGYPLINVTKSKILLVFFYPL     746
Suzuki          RQAHHNSPGELSVAKKMALLVFTNFACWSPIAFFGLTALAGYPLINVTKSKILLVFFYPL     752
Biarmipes       RQAHQNSPGELSVAKKMALLVFTNFACWSPIAFFGLTALAGYPLINVTKSKILLVFFYPL     757
Elegans         RQARQNSPGELSVAKKMALLVFTNFACWSPIAFFGLTALAGFPLINVTKSKILLVFFYPL     748
Rhopaloa        RQSRQNNPGEMSVAKKMALLVFTNFACWSPIAFFGLTALAGFPLINVTKSKILLVFFYPL     734
                *::::*  **:****** **** **:* :************ ******:***********

Mojavensis      NSCADPYLYAILTSQYRQDLLTLLSKFGLCRQRALKYKHSDSMHGTSHYTIRGSIEREHS     788
Virilis         NSCADPYLYAILTSQYQQDLLTFLSKLGICRQNALKYKHSDSLHGTSHYTIRGSIEQQSS     789
Bipectinata     NSCADPYLYAILTAQYRQDLYTLLSKLGLCRKNAVNSKDNSSGMGTTRFTIH----RHSS     780
Anannassae      NSCADPYLYAILTAQYRQDLYTLLSKLGLCRTNAVNSKDNSSGMGTTRFTIH----RHSS     781
Kikkawei        NSCADPYLYAILTSQYRQDLFTLLSKIGLCRQNALNYKHSSSAQATTRFTIH----RHSS     803
Serrata         NSCADPYLYAILTSQYRQDLLTLLSKLGLCRQNALNYNHSSSAPATTRFTIH----RHSS     802
Ficusphelia     NSCADPYLYAILTSQYRQDLFTLLSKLGLCRQSALKYKDSLSGQATTRFTIHGSIQRHGS     825
Erecta          NSCADPYLYAILTSQYRQDLFTLLSKLGLCRQSALKYKDSLSGQATTRFTIHGSIQRHSS     808
Melanogaster    NSCADPYLYAILTSQYRQDLFTLLSKLGLCQQSALKYKDSLSGQATTRFTIHGSIQRHSS     807
Sechellia       NSCADPYLYAILTSQYRQDLFTLLSKLGLCRQSALKYKASLSGQATTRFTIHGSIQRHSS     807
Mauritania      NSCADPYLYAILTSQYRQDLFTLLSKLGLCRQSALKYKDSLSGQATTRFTIHGSIQRHSS     807
```

```
Simulans          NSCADPYLYAILTSQYRQDLFTLLSKLGLCRQSALKYKDSLSGQATTRFTIHGSIQRHSS        807
Eugracilis        NSCADPYLYAILTSQYRQDLFTLLSKLGLCRQSALKYKDSLSGHPTSRFTIHGSIQRHGS        806
Takahashi         NSCADPYLYAILTSQYRQDLFTLLSKLGLCRQSALKYKDSLSGHATTRFTIHGSIQRHGS        806
Suzuki            NSCADPYLYAILTSQYRQDLFTLLSKLGLCRQSALKYKDSLSGHATTRFTIHGSIQRHSS        812
Biarmipes         NSCADPYLYAILTSQYRQDLFTLLSKLGLCRQSALKYKDSLSGHATTRFTIHGSIQRHGS        817
Elegans           NSCADPYLYAILTSQYRQDLFTLLSKLGLCRQSALKDKDGSSTRGTTRYTINGSIQRHAS        808
Rhopaloa          NSCADPYLYALLTSQYRQDLFTLLSKLGLCRQSALKYKDSSSAHATSRFTIHNSIHRHGS        794
                  **********:**:**:*** *:***:*:*:   *:: : . *   *:::**.    :. *

Mojavensis        VGQKCQKI-VAAEAQNMLRNNEDYV     812
Virilis           LCQKPQQE-GAAETQTMLKNNEDYV     813
Bipectinata       LTCRIPPALEV-ETQKMLTCNEDYV     804
Anannassae        LTCRIPPALEV-ETQKMLTYSEEYV     805
Kikkawei          LTCKMQAAISV-EAQKMLKNGEDYV     827
Serrata           LTCKMQTVLNA-EAQKMLKNGEDYI     826
Ficusphelia       LTCKMQTVMGT-ETQKMLKCGEDYV     849
Erecta            LTCKMQTVMGA-ETQKMLKNSEDYV     832
Melanogaster      LTCKMQTVMGA-ETQKMLKNSEDYV     831
Sechellia         LTCKMQTLMGA-ETQKMLKNSEDYV     831
Mauritania        LTCKMQTVMGA-ETQKMLKNSEDYV     831
Simulans          LTCKMQTVMGA-ETQKMLKNSEDYV     831
Eugracilis        LTCKMQTTMT-TETQKMLKNSEDYV     830
Takahashi         LTCKMQTVMGATETQKMLKSSEDYV     831
Suzuki            LTCKMQTVMGT-ETQKMLKKSEEYV     836
Biarmipes         LTCKMQTVMGA-ESQKMLKNTEDYV     841
Elegans           LTCKMQTVVGA-ETQKMLKNGEDYV     832
Rhopaloa          LTSKMQTVISA-ETQKMLKNSEDYV     818
```

Variants at this putative BBS do not all have 2 S or Ts, some only 1 – meaning the other species do not contain partial BBS codes at this positions

Melanogaster NP_524393.2

```
mekhpslsqr mgttyrprkg lkclsfefqc rlllhhlllt slsgrhfvya tsavggalsa
       61 nnchdihhgf dvypnltavs laqstdtplt atmprsawkc ccwnasnqae evecrcegdg
      121 lnrvpqtltl piqrltiasa glprlrhtgl kvygstlldv aftdclqlel iqdgafanlt
      181 llrtiyitna pkltflskdv flgisdtvdi iriinsgltr vpdlghlpph nilqmidldn
      241 nqitridsks ikvktaqlil tnneisyvdd saffgskiak lslkenkklq mmhpnafdgi
      301 iditeldlss tslvglpsag lqniealyiq nthtlktips iynfrnlqra ylthsfhcca
      361 fqfpsrhdpq rhaqrmleie kwrkqcksds gtrkerstld npfnmpedfg sfggtddsat
      421 ditpitfasf dymaddtmnk gtfhekiiln pgddssaelc gnftfrkpni ecypmpndln
      481 pcedvmgyqw lrisvwivva lavvgnvavl tvilsirpes tpvprflmch lafadlclgl
      541 ylllvacida hsmgeyfnfa ydwqyglgck vagfltvfas hlsvftltvi tierwlaitq
      601 amylnhrikl rpaalimlgg wiysmlmssl plfgisnyss tsiclpmenr dvydtiylia
      661 ilgsngvafs iiavcyaqiy lslgretrqa hqnspgelsv akkmallvft nfacwspiaf
      721 fgltalagyp linvtkskil lvffyplnsc adpylyailt sqyrqdlftl lsklglcqqs
      781 alkykdslsg qattrftihg siqrhssltc kmqtvmgaet qkmlknsedy v
```

Mauritania XP_033166657.1

```
  1 mekhpslsqr mdttcrprkg lkclsfefqc rlllhhlllt slsgrhcvya tsaaggalsa
       61 snchdihhgf dvypnptavs laqstdtplt aamprsawkc ccwnasnqae evecrcegdg
      121 lnrvpqtltl piqrltiasa glprlrhtgl kvygstlldv aftdclqlel iqdgafanlt
      181 llrtiyitna pkltflskdv flgisdtvdi iriinsgltr vpdlghlpph nilqmidldn
      241 nqitridsks ikvktaqlil tnneisyvdd saffgskiak lslkenkklq mmhpnafdgi
      301 idiaeldlss tslvglpsag lqniealyiq nthtlktips iynfrnlqra ylthsfhcca
      361 fqfpsrhdpq rhaqrmleie kwrkqcksds gsrkerstld nsfdmpedfg sfggtddsai
      421 ditpitfasf dymaddtmnk gtfhekiiln pgddssaelc gnftfrkpni ecypmpndln
      481 pcedvmgyqw lrisvwivva lavvgnvavl tvilsirpes tpvprflmch lafadlclgl
      541 ylllvacida hsmgeyfnfa ydwqyglgck vagfltvfas hlsvftltvi tierwlaitq
      601 amylnhrikl rpaalimlgg wiysmlmssm plfgisnyss tsiclpmenr dvydtiylia
      661 ilgsngvafs iiavcyaqiy lslgretrqt hqnnpgelsv akkmallvft nfacwspiaf
      721 fgltalagyp linvtkskil lvffyplnsc adpylyailt sqyrqdlftl lsklglcrqs
      781 alkykdslsg qattrftihg siqrhssltc kmqtvmgaet qkmlknsedy v
```

Simulans XP_016034243.1

```
1 mekhpslsqr mettcrprkg lkclsfefqc rlllhhlllt slsgrhcvya tsaaggalsa
       61 snchdihhgf dvypnptavs laqstdtplt aamprsawkc ccwnasnqae evecrcegdg
      121 lnrvpqtltl piqrltiasa glprlrhtgl kvygstlldv aftdclqlel iqdgafanlt
      181 llrtiyitna pkltflskdv flgisdtvdi iriinsgltr vpdlghlpph nilqmidldn
```

```
241 nqitridsks ikvktaqlil tnneisyvdd saffgskiak lslkenkklq emhpnafdgi
301 idiaeldlss tslvglpsag lqniealyiq nthtlktips iynfrnlqra ylthsfhcca
361 fqfpsrhdpq rhaqrmleie kwrkqcksds gsrkerstld nsfnmpedfg sfggtddsat
421 dmtpitfasf dymaddtmnk gtfhekiiln pdddssaelc gnftfrkpni ecypmpndln
481 pcedvmgyqw lrisvwivva lavvgnvavl tvilsiraes tpvprflmch lafadlclgl
541 ylllvacida hsmgeyfnfa ydwqyglgck vagfltvfas hlsvftltvi tierwlaitq
601 amylnhrikl rpaalimlgg wiysmlmssm plfgisnyss tsiclpmenr dvydtiylia
661 ilgsngvafs iiavcyaqiy lslgretrqa hqnnpgelsv akkmallvft nfacwspiaf
721 fgltalagyp linvtkskil lvffyplnsc adpylyailt sqyrqdlftl lsklglcrqs
781 alkykdslsg qattrftihg siqrhssltc kmqtvmgaet qkmlknsedy v
```

Sechellia **XP_002041048.1**

```
  1 mekhpsmsqr lettcrprkg lkclsfefqc rlllhhlllt slsgrhcvya tsaaggalsa
 61 snchdihhgf dvypnptavs laqstdtplt aamprsawkc ccwnasnqae evecrcegdg
121 lnrvpqtltl piqrltiasa glprlrhtgl kvygstlldv aftdclqlel iqdgafanlt
181 llrtiyitna pkltflsrdv flgisdtvdi iriinsgltr vpdlghlpph nilqmidldn
241 nqitridsks ikvktaqlil tnneisyvdd safygskiak lslkenkkle mmhpnafdgi
301 idiaeldlss tslvglpsag lqniealyiq nthtlktips iynfrnlqra ylthsfhcca
361 fqfpsrhdpq rhaqrmleie kwrkqcksds gsrkerstld nsfsmpedfg sfggtddsst
421 ditpitfasf dymaddtmnk gtfhekiiln pgddssaelc gnftfrkpni ecypmpndln
481 pcedvmgyqw lrisvwivva lavvgnvavl tvilsirpes tpvprflmch lafadlclgl
541 ylllvacida hsmgeyfnfa ydwqyglgck vagfltvfas hlsvftltvi tierwlaitq
601 amylnhrikl rpaalimlgg wiysmlmssm plfgisnyss tsiclpmenr dvydtiylia
661 ilgsngvafs iiavcyaqiy lslgretrqa hqnnpgelsv akkmallvft nfacwspiaf
721 fgltalagyp linvtkskil lvffyplnsc adpylyailt sqyrqdlftl lsklglcrqs
781 alkykaslsg qattrftihg siqrhssltc kmqtlmgaet qkmlknsedy v
```

Erecta **XP_001979725.1**

```
1 mekhpslpps qtqtqrrdtt crskkglkcl sfefqcrlll hhllltslsg rhcvyamsas
 61 schdihhgfd vypnpnpttv slgqstdtpq aeamppsawk cccwnasnqv eevecrcegd
121 glnrvpqtlk lpiqrltias aglprlrhtg lkvygptlld vaftdclqle liqdgafanl
181 tllrtiyitn apkltflskd vflgisdtve iiriinsglt rvpdlghlpp hnilqmidld
241 nnqitridsk sikvktaqli lanneisyvd dsaffgskia klslkenkrl kmmhpnafdg
301 iidiaeldls stslvglpsa glqniealyi qnthtlktip siynfrnlqr aylthsfhcc
361 afqfpsrhdp qrhaqrmlei ekwrkqckse sgsrqerstl dsslsmpedf gtfsgtddsa
421 tditpitfvs pdymaddtmn kgtfhekivl npeddssael cgnftfrkpn iecypmpndl
481 npcedvmgyq wlrisvwivv alavvgnvav ltvilsirpe stpvprflmc hlafadlclg
541 lylllvacid ahsmgeyfnf aydwqyglgc kvagfltvfa shlsvftltv itierwlait
601 qamylnhrik lrpaalimlg gwiysmlmss lplfgisnys stsiclpmen rdvydtvyli
661 ailgcngvaf siiavcyaqi ylslgretrq ahqnnpgels vakkmallvf tnfacwspia
721 ffgltalagy plinvtkski llvffyplns cadpylyail tsqyrqdlft llsklglcrq
781 salkykdsls gqattrftih gsiqrhsslt ckmqtvmgae tqkmlknsed yv
```

Takahashi **XP_017013746.2**

```
1 mekhtslprk easessrkgl kclsleflfr llfhqllfts lcgphcvyat savgealgas
 61 nchdilhgfd vypnptagal gqstdspltv tlppsgwkcc cwnasnqmee vecrcegdgl
121 nrvpqtlklp iqrltiasag lprlrytglk vygptlldvt ftdclqleli qdgafanltl
181 lrtiyitnap kltflskdvf fgisdtveii riinsgltrv pdlshlpphn ilqmidldnn
241 qitridtksi kvktaqlila nndisyvdds affgskiakl slndnqkleh mhpnafdgii
301 diteldlsst slvglpssgl qtiealyiqn thtlktipsi ynfrnlqray lthsfhccaf
361 qfpsrhdpqr hakrmreiek wreqcksdsa srkerstsen gfempedfgs fgspdeattd
421 ivpikfasfd ymaddtlnkg tfhekitldp addmsaelcg nftfrkpnie cypmpndlnp
481 cedvmgyqwl risvwivval avvgnvavlt vilsirpest pvprflmchl afadlclgvy
541 lllvasidah smgeyfnyay dwqyglgckv agfltvfash lsvftltvit ierwlaitqa
601 mylnhriklr paalimlggw iysmfmsslp lfgisnysst siclpmenrd iydtvyliai
661 mvcngvafsi iavcyaqiyl slgretrqah hnspgelsva kkmallvftn facwspiaff
721 gltalagypl invtkskill vffyplnsca dpylyailts qyrqdlftll sklglcrqsa
781 lkykdslsgh attrftihgs iqrhgsltck mqtvmgatet qkmlkssedy v
```

Suzuki **XP_016932820.2**

```
1 mekhtsqypk espwrtkrgl kcltfefqfr lllhhlmlss lcgphcvyat savgealsas
 61 nchdihhgfd vypnpnpspn ltavtlgqst dtpsvtltlp psgwkcccwn asnqaeevec
121 rcegdglnrv pqtlklpiqr ltiasaglpr lrytglkvyg ptlldvaftd clqleliqdg
181 afanltllrt iyitnapklt flskdvffgi sgtveiirii nsgltrvpdl shlpphnilq
241 midldnnqit ridtksikvk taqlilannd isyvddsaff gskiaklsln dnlrlehmhp
301 nafdgidite ldlsstslva mpsvglqnie alyiqnthtl ktipsiynfr nlqraylths
```

```
     361 fhccafqfps rhdpqrhaqr mleiekwrkq cksesvsrqe rstldnaytm pedfgtfest
     421 desstdlvpi tfasfdymad dslnkgtfhe kitlnpeddm saelcgnftf rkpniecypm
     481 pndlnpcedv mgyqwlrisv wivvalavvg nvavltvils irpestpvpr flmchlafad
     541 lclgvylllv asidahsmge yfnyaydwqy glgckaagfl tvfashlsvf tltvitierw
     601 laitqamyln hriklrpaal imlggwiysm fmsslplfgi snysstsicl pmenrdvydt
     661 vyliaimvcn gvafsiiavc yaqiylslgr etrqahhnsp gelsvakkma llvftnfacw
     721 spiaffglta lagyplinvt kskillvffy plnscadpyl yailtsqyrq dlftllsklg
     781 lcrqsalkyk dslsghattr ftihgsiqrh ssltckmqtv mgtetqkmlk kseeyv
```

Biarmipes XP_016951482.1

```
1 mekhtsqcpk eapwrpkrgl kcltiefqfr lllhhlllss lcgphcvyat savgealsas
      61 nchdihhgfd vypnpnlnpt ssangttvtl gqsrdtpsvp ltlppsgwkc ccwnggnqae
     121 evecrcegdg lnrvpqtlkl plqrltiasa glprlrytgl kvygatlldv aftdclqlel
     181 iqdgafanlt llrtiyitna pkltflskdv flgisgtvev iriinsgltr vpdlshlpph
     241 nilqmidldn nqitridtks ikvktaqlil anndisyvdd safygskiak lslndnlrle
     301 hmhpnafdgi iditeldlss tslvampsvg lqniealyiq nthtlktips iynfrnlqra
     361 hlthsfhcca fqfpsrhdpq rhaermleie kwrkqcksds vtrkerstle ngftlpedfg
     421 tfggtddsst dlvpitfasf dymaddtlik gtfhekiiln peddmsaelc gnftfrkpni
     481 ecypmpndln pcedvmgyqw lrisvwivva lavvgnvavl tvilsirpes tpvprflmch
     541 lafadlclgv ylllvasida hsmgeyfnya ydwqyglgck vagfltvfas hlsvftltvi
     601 tierwlaitq amylnhrikl rpaalimlgg wiysmfmssl plfgisnyss tsiclpmenr
     661 dgydtvylia imvcngvafs iiavcyaqiy lslgretrqa hqnspgelsv akkmallvft
     721 nfacwspiaf fgltalagyp linvtkskil lvffyplnsc adpylyailt sqyrqdlftl
     781 lsklglcrqs alkykdslsg hattrftihg siqrhgsltc kmqtvmgaes qkmlkntedy
     841 v
```

Eugracilis XP_041675437.1

```
   1 mekhtnlcrt estwrpkkgl kclsfeylfr lllhhlllts lsgphcvyat sdigealsas
      61 nchdihhgfd vypnpnalsl gqstdtpltq tlppsawkcc cwnasnqmee vecrcegdgl
     121 nrvpqtlklp iqrltiasag lprlrytglk vygptlldva ftdcqqleli qdgafsnltl
     181 lrtiyitnap kltflskdvf lgisdtveii riinsgltrv pdlshlppnn ilqmidldnn
     241 qitridtksi kvktaqlila nneisyidds affgskiakl slkdnlklem mhpkafdgii
     301 diteldlsst slvglpsvgl qtiealyiqn thtlktipsi ynfrnlqray lthsfhccaf
     361 qfpsrhdpqr haqrmleiek wrkqcksdsg srkerstldg ldnqpedfgs fggsyqsatd
     421 ispitfssid ymaddtmnkg tfhekitlnp eddmsaelcg nftfrkpnie cypmpndlnp
     481 cedvmgyqwl risvwivval avvgnvavlt vilsirpett pvprflmchl afadlclgvy
     541 lllvasidah skgeyfnfay dwqyglgckv agfltvfash lsvftltvit ierwlaitqa
     601 mylnhriklr paalimlggw iysmfmssmp lfgisnysst siclpmenrd vydtvylvai
     661 mvcngvafsi iavcyakiyl slgretrqar hnnpgelsva kkmallvftn facwspiaff
     721 gltalagypl invtkskill vffyplnsca dpylyailts qyrqdlftll sklglcrqsa
     781 lkykdslsgh ptsrftihgs iqrhgsltck mqttmttetq kmlknsedyv
```

Ficusphelia XP_017047222.1

```
   1 mkkhtnlhrk evtwrsrrgt kcltfeflfr llynhlllts mfgpdcvyat savgkalsas
      61 nchdihhgfd vfpnpnptpa lnavdsnpnt ssnatamwlg qstdtpvtlt lppsawkccc
     121 wnasnqneev ecrcegdgln rvpqtlklpl qrltiasagl prlrytglkv ygptlldvtf
     181 tdclqleliq dgafanltll rtiyisnapk ltflskdvfa gisesveiir iinsgltrvp
     241 dlghlpphni lqmidldnnq isridtksik vktaqlilan ndisyvddsa ffgskiakls
     301 lqnnrklkqm hpnafdgiid iteldlssts lvglpsvglq niealyitnt htlktipsiy
     361 nfrnlqrayl thsfhccafq fpsrhdpqrh aermreiekw reqcksdhgs rkersaldgf
     421 eglpedfgtf gmpeqsgtdd tsityasfdy msddtlnkgt fhekitlnpe ddmtaelcgn
     481 ftfrkpniec ypmpndlnpc edvmgyqwlr isvwivvala vvgnvavltv ilsirpestp
     541 vprflmchla fadlclgvyl llvasidahs mgeyfnyayd wqyglgckva gfltvfashl
     601 svftltviti ercmaitqam ylnhriklrp aaiimlcgwi ysmlmsslpl fgisnyssts
     661 iclpmenrdv fdtmyliail gcngvafsii avcyaqiyls lgretrqarh nnpgelsvak
     721 kmallvftnf acwspiaffg ltalagypli nvtkskillv ffyplnscad pylyailtsq
     781 yrqdlftlls klglcrqsal kykdslsgqa ttrftihgsi qrhgsltckm qtvmgtetqk
     841 mlkcgedyv
```

Elegans XP_017119258.1

```
1 mekpailark dvlrrsrkcs kglkclsfdf qfrlllhyll lsslsgphcv yatmagseal
      61 sasnchdihh gfdvypnpta vsqgqstdtp ltltlppsgw kcccwnasnq neevecrcgg
     121 dglnrvpqtl krpiqrltia saglprlrnt glkvygstll dvaftdclql eliqdgafan
     181 ltllrtiyis napkltflsk dvffgisgsv eiiriinsgl trvpdlghlp phnilqmidl
     241 dnnqitridt ksikvktaql ilanndisyv ddsaffgski aklslkdnpk lkelhpnafd
     301 giiditeldl sstslvgmps vglqtiealy iqnthtlkti psiynfrnlq raylthsfhc
```

```
361 cafqfpsrhd pqrhaqrmle iqkwreqcks dhvsrkerst vdnlnsmpee fgsfgstdqs
421 atdmtsmtya sfdymsddtm nkgtfhekit lnpeddidel cgnftfrkpn iecypmpndl
481 npcedvmgyq wlrisvwivv alavvgnvav ltvilsirpe stpvprflmc hlafadlclg
541 vyllfvasid ahsmgeyfnf aydwqyglgc kvagfltvfa shlsvftltv itierwlait
601 qamylthrik lrqasiimlg gwiysvlmss lplfgisnys stsiclpmei rdgfdtayli
661 ailacngvaf tiiavcyaqi ylslgretrq arqnspgels vakkmallvf tnfacwspia
721 ffgltalagf plinvtkski llvffyplns cadpylyail tsqyrqdlft llsklglcrq
781 salkdkdgss trgttrytin gsiqrhaslt ckmqtvvgae tqkmlknged yv
```

Rhopaloa XP_016976568.1

```
  1 mekytilark rarrpskril kclsleflyr lllhhlllts lcdhhyvyat savgepisas
 61 dchdihhgfd vfpnhtalpl gqstdtplts tlppsgwkcc cwkssnqnee vecrcegdgl
121 nrvpqslklp iqrltiasag lprvrftglk vygsslldva fidclqleli qdgafanlal
181 lrtiyisnap kltflskdvf sgisgsveii riinsgltrv pdlghlpphn ilqmidldnn
241 qitridtksi nvktaqlila nndisyvdds affgskiakl slndnpklke lhpnafngii
301 ditaldlsst sligmpsagl qtiealyiqn thtlktipsi ynfrnlqray lthsfhccaf
361 qfpsrhdper haqrmleiek wrkqcksdqv srkersiadn lnsvpeefgs fgmtdqsldy
421 msddtmnkgt fhekitlnpe ddidelcgnf tfrkpniecy pmpndlnpce dvmgyqwlri
481 svwivvalav vgnvavltvi lsirpestpv prflmchlaf adlclgvyll lvasidahsm
541 gayfnyaydw qyglgckvag fltvfaslls vftltvitie rwlaitqamy lthriklrpa
601 sfimlggwiy silmsslplf gisnysstsi clpmeirdvf dtvyliaila cngvafsiia
661 vcyaqiylsl gretrqsrqn npgemsvakk mallvftnfa cwspiaffgl talagfplin
721 vtkskillvf fyplnscadp ylyalltsqy rqdlftllsk lglcrqsalk ykdsssahat
781 srftihnsih rhgsltskmq tvisaetqkm lknsedyv
```

Kikkawei KAH8340697.1

```
  1 mgrrqtnwvr rkgrgqpkta ltclsigflf rflfvyhlll aplcgphcvy amlaaegpsd
 61 gsnchdihhg fdvpsslnav teaqggdapm avtltpapsg wkcccwkapn qgeevecrce
121 gdaltrvpqt lqlpiqrlti asaglprlrf tglkvygptl ldvaftdclk leliqdgafa
181 nltllrtiyi snapkltfls kdvfagiset veiiriinsg ltsvpdlghl pphnilqmid
241 ldnnqitrid tksinvktaq filanndisy vddsaffgsk iaklslkdnr kltdmhpnaf
301 ngiidiaeld lsstslvglp svglqtieal yimnthtlkt ipsiynfrnl qraylthsfh
361 ccafqfpsrh dprrhaqrml eiekwreqcn kgsrkersll dnfeaqpedf gsfgstepsv
421 teitpypyas vdymadstnl gyfheqitin pdddqsaefc gnftfrkpni ecypmpndln
481 pcedvmgyqw lrisvwivva lavvgnvavl tvilsirpes tpvprflmch lafadlclgv
541 ylllvasida hsmgeyfnya ydwqyglgck aagfltvfas hlsvftltvi tierwlaitq
601 amylnhrikl rqaalimlgg wlysmfmssl plfgisnyss tsiclpmenr dafdtmylia
661 ilgcngvafs iiavcyaqiy lslgretrqa rqnnpgelsv akkmallvft nfacwspiaf
721 fgltalagfp linvtkskil lvffyplnsc adpylyailt sqyrqdlftl lskiglcrqn
781 alnykhsssa qattrftihr hssltckmqa aisveaqkml kngedyv
```

Bipectinata XP_017105030.2

```
  1 mkcvlpkfvf rlllhhlllt klcwlhgvya tsaschdsyn gfnvfpngss dlrpdpnptd
 61 vssgqapppt ltlppwkccc wnatnqseea ecrcegealt rvpqtltmam qrltiatagl
121 prlratglkv yaqtlmdvaf tdclqleliq dgafanlkll rtiyianapk ltflskdvff
181 gisdtveiir iinsgltrvp dlthlppyni lqmidldnnq itridaksik vktaqlilan
241 ndisyvddsa fcgskiakls lkenrklkel htnafhgiid iteldlssts ivempsaglq
301 tiealyilnt htlktipsiy nfrnlqrayl thsfhccafq fpsrhdplrh aqrmdeiekw
361 rtqckgeris rkersilnnl ksqsdgygif gseasptdvv stqfasvdym adaamnlgyf
421 heeitlnpdd eqfaefcgnf tfrkpnvecy pmpnalnpce dvmgyqwlri svwvvvalav
481 vgnvavltvn lsirpeitpv arflmchlaf adlclglylf lvasidahsm geyfnyaydw
541 qyglgckvag fltvfashls vftltlitie rwvaitqamy lnkrirlrsa siimlggwiy
601 smvmsslplf gisnysstsi clpmevrdsf dtvyliailg cngvaffiia vcyakiylsl
661 gretrharqn npgelsvakk mallvftnfa cwspiaffgl talagfplin vtnskillvf
721 fyplnscadp ylyailtaqy rqdlytllsk lglcrknavn skdnssgmgt trftihrhss
781 ltcrippale vetqkmltcn edyv
```

Anannassae XP_001964371.3

```
1 mkcvlpkfvy rlllhhlllt ricwphgvya tsavchdshn gfnvspdgns dlspdrnptt
 61 mssgqapppt ltlppwkccc wnatnqseev ecrcegealt rvpqtltmam qrltiasagl
121 prlratglkv yaqtlldvaf tdclqleliq dgafanlkll rtiyianapk lsflskdvfs
181 gisdtveiir iinsgltsvp dlthlppyni lqmidldnnq itridsksik vktaqlilan
241 ndisyvddsa ffgskiakls lkenrklkel htnafhgiid iteldlssts ivempsaglq
301 tiealyilnt htlktipsiy nfrnlqrayl thsfhccafq fpsrhdplrh aqrmleiekw
361 rtqcngerns rrerniakki esqqddlgtl ggsealsadv lstpfssvdy madaamnlgy
421 fheeitinpd deqfaefcgn ftfrkpsvec ypmpdalnpc edvmgyqwlr isvwvvvala
481 vvgnvavltv nlsirpeitp varflmchla fadlclglyl flvasidahs mgeyfnyayd
```

```
541 wqyglgckva gfltvfashl svftltliti erwvaitqam ylnkrirlrs asiimlggwi
601 ysmvmsslpl fgisnyssts iclpmevrdt fdtvyligil gcngvaffii avcyakiyfs
661 lgretrharq nnpgelsvak kmsllvftnf acwspiaffg ltalagypli nvtnskillv
721 ffyplnscad pylyailtaq yrqdlytlls klglcrtnav nskdnssgmg ttrftihrhs
781 sltcrippal evetqkmlty seeyv
```

Mojavensis XP_001998425.1

```
1 mkctpiiarf dfsavillsl vycsqqastn cydnhdgfna iinnlprdag naidtldtla
61 qappmttnpp vdasvwkccc weatnhnefe crcegealtr vpqtlklplq rltiasaglp
121 rlrsmglkvy aptlldvafi dclqletiqs gafsnltvlr aiyisnapkl sylaknvfeg
181 isdtieiiri insglktvpd lgdlppynil qmidldnnqi sridsksiqv ktaqlvlann
241 eitfiddsaf lgskiaklsl ndnhklteih pnafngiidm teldlsstsl vrlpsaglqt
301 levlyianth tlktipsiyn fqnlqrahlt hsfhccafqf psrhdpqrha qrmqevekwr
361 dqcknnrdvh tnyerslsms nltplanvse vqpsgstinn llaestpnty aymadstlnn
421 igifheeiti npdddqlaey cgnftfrkph vecypmpnal npcedvmgyq wlriavwivv
481 alaivgnvav ltvilsikse spsvprflic hlafadlclg lylllliasid ahsmgeyfny
541 afdwqyglgc kiagfltvfa shlsvftltv itierwfait hamylnkrik lrpaavimlm
601 gwiysivmss lpllgisnys stsiclpmek rdiydsiylv lilgcnflaf tiiaicysqi
661 ylslgketrr arqnhlgems vakkmallvf infscgapia ffgltalagy plinvtkski
721 llvffyplns cadpylyail tsqyrqdllt llskfglcrq ralkykhsds mhgtshytir
781 gsierehsvg qkcqkivaae aqnmlrnned yv
```

Virilism XP_032289845.1

```
1 mkcmviisql dfltfillsl ahgtrqantn chdnnngfnt vfnnlsidng netdmsvimt
61 qaptmstnpt sdasvwkccc weatnqnefe crcegealtr vpqtlklpll rltiasaglp
121 rlrsmglkvy attlldvafi dclqleaiqn gafsnltflr tiyisnapkl tylpknvfeg
181 isdtieiiri insgltsvpd fgylppnnil qmidldnnqi sridsksiqv ktaqfvlann
241 dihfiddsaf lgskiaklsl kdnrrltdvh pnafygiidi teldlsstsl vslpsaglqt
301 vevlyiinth tlktipsiyn fqnlqrahlt hsfhccafqf psrhdpkrha erlqelqkwr
361 eqcnverdly rdvdkklkkn tsvkdtagvh ytnqsgtatd nlltdaasns ydymadstmn
421 nigifheqit inpddnqlae ycgnftfrnp diqcfpmpna lnpcedvmgy qwlrisvwiv
481 valaivgnla vltvtlsiks espsvprfli chlafadlcl glylllliasi dahsmgeyfn
541 yafdwqyglg ckiagfltvf ashlsiftlt iitierwfai thamylnkri tlrqaagiml
601 tgwiysiims slplfgisny sstsiclpme irdiydsiyl ililgcnfva ftiiaicysq
661 iylslgqetr rarrnnpgem svakkmlllv finftcgapi affgltalag cplinvtksk
721 illvffypln scadpylyai ltsqyqqdll tflsklgicr qnalkykhsd slhgtshyti
781 rgsieqqssl cqkpqqegaa etqtmlknne dyv
```

Grimshawi EDV91130.1

```
1 mkwttfglqf diltlfvlwv lvccahqast ncydsssgfn tnpnalpaed gnetyttdtt
61 pmptadasvw kcccweatnq nlfpedefec rcegealtrv pqtlklplqr ltiasaglpr
121 lrsiglkvya stlldvaftd cpqletieng afsnltilrt iyisnapklk ylskdvfegi
181 snkidiirii nsgltsvpdl ghlppntilq midldnnqis riesksiqvk taqfvlannd
241 ityiddsafl dskiaklslk qnrrlseihp nafngiidmt eldlsstslv rlpaaglqti
301 evlyimntht lktipsiynf qnlqkahlth sfhccafkfp srhdpqrhkk heeelrklqe
361 qcksdrdmhn nvatniiipt ksmedavgmp ngkgnwgien gdwnnnelid spealpnmdd
421 ymadatmnnl gvfhteitin pddnqmtefc gnftfrkpni ecypvpnaln pcedvmgyew
481 lriavwivva ltivgnvavl tvilsikses psvprflich lafadlclgl yllliasida
541 hsmgqyfnya ydwqygfgck vagfltvfas hlsvftltvi tierwfaith amylnkrikl
601 grasvimitg wlyaitmssl plfgisnyss tsiclpmeqr diydsvyllm ilgsnfiaft
661 iiaicysqiy lslgqetrna rqnnpgemsi akkmallvfi nfscgapiaf fgltalagyp
721 linvtkskil lvffyplnsc adpylyailt sqyrqdlfll fsklgicrks vmkykysdsm
781 phtshftirn sieqpnafph kapngagaet qkmlinnedy v
```

Serrata XP_020813515.1

```
1 mgrrrthwvr kgdrrpvktg ikclsigyll rllfvhhlll gsfcgshrvy amlaaegqsv
61 rsnshdvhhg fdvpgrqpvs dnqvrgapit vilpkvtsrw kcfcsktpnq seelecrceg
121 dsltrvpqtl qlpiqrltia saglprlrst glkvygptll dvaftdclkl eliqdgafan
181 ltmlrtiyis napkltflsk dvfagisetv eviriinsgl trvpdlghlp phnilqmidl
241 dnnqitridt ksinvktaql ilanndisyv ddsaffgski aklslkdnwk ltemhpeafn
301 giiditeldl sstslvglps vglqtiealy imnthtlkti psiynfrnlq raylthsfhc
361 cafqlpsrhd prrhalrmle iekwreqcnk csrkeralld yleaqpedfg sfgstepsmt
421 entpfpiasi dymadstnlg yfheqitinp ddkqsaefcg nftfrkpnie cypmpndlnp
481 cedvmgyqwl risvwivval savgnvavlt vilsirpess pvprflmchl afadlclgvy
541 lllvasidah sigeyfnyay dwqyglgckv agfltvfash lsvftltvit ierwlaitqa
601 mylnhrikmr qaalimlggw lysmvmsslp lfgisnysst siclpmenrd afdtmyliai
661 lgcngvafsi iavcyaqiyl slgretrqah qnnpgelsva kkmallvftn facwspiaff
```

721 gltalagfpl invtkskill vffyplnsca dpylyailts qyrqdlltll sklglcrqna
781 lnynhsssap attrftihrh ssltckmqtv lnaeaqkmlk ngedyi
