## Supplementary material for "The incidence of candidate binding sites for β-arrestin in Drosophila neuropeptide GPCRs": S15 Text

***S15. Text*** **Multi-species analysis of NPF R**
**Supporting Figure 18**

CLUSTAL Line-ups; Genbank Reference IDs below
7th Predicted TM domain in YELLOW
BBS sequences in RED

```
mojavensis        -----MNRTELGSQIL---SSAELYKALSSDATFDTIGESRHLVNYPGLDAGGAYTRNHN    52
grimshawi         -MIIAMNRTEFGSQLPNFDSSVEIFKAIARNSNFDLIGERRHLVNYSQLNALSDVNANNT    59
virilis           ------------------------------------------------------------    0
elegans           -MIISMNQT-------AEHLS--GY--VSSS-NSGRYLDDRHPLDYLDLGPARALN----    43
rhopaloa          -MIISMNQTETGPLAAGEHLS--GY--ASSS-NSGRYLDDRHPLDYLDLGIVHALN----    50
ficusphla         -MIISMNQTETGQLSAGEHLS--GY--ASSG-NSVRYLDDRHPLDYLDLGTVHALN----    50
biarmpes          -MIISMNQTEPGQLAAAEHLG--GY--ASSS-NSGRYLDDRHPLDYLDLGTIHALN----    50
suzuki            -MIISMNQTEPGLLATGEHLG--GY--ASSS-NSGRYLDDRHPLDYLDLGTIHALN----    50
simulans          -MIISMNQTEPTQLAAGEHLS--GY--ASSS-NSVRYLDDRHPLDYLDLGTVHALN----    50
erecta            -MIISMNQTEAAQLAAGEHLG--GY--ASSS-NSVRYLDDRHPLDYLDLGTVHALN----    50
melanogaster      -MIISMNQTEPAQLADGEHLS--GY--ASSS-NSVRYLDDRHPLDYLDLGTVHALN----    50
sechelia          -MIISMNQTEPTQLAAGEHLS--GY--ASSS-NSVRYLDDRHPLDYLDLGTVHALN----    50
mauritania        -MIISMNQTETTQLAAGEHLS--GY--ASSS-NSVRYLDDRHPLDYLDLGTVHALN----    50
takahashi         -MIISMNQTEPGQLAAGEQLG--GY--ASSS-NSGRYLDDRHPLDYLDLGTVHALN----    50
eugracilis        -MIISMNQTEPGPLAAGEHLS--GY--ASSS-NSVRYLDDRHPLDYLDLGMVHALN----    50
bipectinata       -MIIGMNQTESGPLATGDRLS--GY--ASSG-NSVRYLDDRHPLDYLDLGSVVSNA----    50
anannassae        MMIIGMNQTESGPLATGDRLS--GY--ASSG-NSGRYLDDRHPLDYLDLGSVVSNA----    51
serrata           -MIISMNQTESVPLAAGDRLS--GF--AAGGDNSVRYLDDRHPLDYLDLGSVVGTA----    51
kikkawei          -MIISLNQTESVPLAAGDRLS--GF--AGGGDNSVRYLDDRHPLDYLDLGSVVGTV----    51


mojavensis        YDKLNSNYEDSSSSSSNDNDSMTMLLLNST---SNGSYVPAGMDPVLMDQYLHNRSIGSP    109
grimshawi         ---NKSNYN-----ISHSSNNMTIQLLNSTHINVSNIIVSSDLDPVLMDQYQHNRAIESP    111
virilis           ---------------------MTILLLNSTN--NESNFMPADMDPVLMDQYLHNRAIESP    37
elegans           ------------------AT--------AVNTSEMNETGSRPLDPVLIDRFLSNRAVDSP    77
rhopaloa          ------------------TT--------AINTSELNETGSRPLDQVLIGRFLSNRAVDSP    84
ficusphla         ------------------TS--------AMNTSDADETGSRPLDPVLIDRFLSNRAVDSP    84
biarmpes          ------------------TT--------AINTSDLNETGSRPLDPVLIDRFLSNRAVDSP    84
suzuki            ------------------TT--------AINTSDLNETASRPLDPVLIDRFLSNRAVDSP    84
simulans          ------------------TT--------AINTSDLNETGSRPLDPVLIDRFLSNRAVDSP    84
erecta            ------------------TT--------AINTSELNETGSRPLDPVLIDRFLSNRAVDSP    84
melanogaster      ------------------TT--------AINTSDLNETGSRPLDPVLIDRFLSNRAVDSP    84
sechelia          ------------------TT--------AINTSDLNETGSRPLDPVLIDRFLSNRAVDSP    84
mauritania        ------------------TT--------AINTSDLNETGSRPLDPVLIDRFLSNRAVDSP    84
takahashi         ------------------TT--------AINTSDLNETGSRPLDPVLIDRFLSNRAVDSP    84
eugracilis        ------------------TT--------AINTSEMNETGSRPLDPVLIDRFLSNRAVDSP    84
bipectinata       ------------------HAA--LNSTATSNLSDANDTGARPLDPVLIDRFLSNRAVDSP    90
anannassae        ------------------HAA--LNSSANNNFSEANDTGARPLDPVLIDRFLSNRAVDSP    91
serrata           ------------------HAV--LNATA-ANMSELNETGSRPLDPVLIDRYLSNRAVDSP    90
kikkawei          ------------------HAV--INATA-TNMSELNETGSRPLDPVLIDRYLSNRAVDSP    90
                                                     .      :* **:.::  **:: **

mojavensis        WYHLLIAIYGVLIVFGAMGNIMVVIAVLRKPIMRTARNLFILNLAISDLLLCLVTMPLTL    169
grimshawi         WYHLLIAMYSILIVFGAMGNIMVVIAVLRKPLMRTARNLFILNLAISDLLLCLVTMPLTL    171
virilis           WYHLLIAMYSVLIVFGAMGNIMVVIAVVRKPIMRTARNLFILNLAISDLLLCLVTMPLTL    97
elegans           WYHMLISMYGVLIVFGALGNTLVVIAVVRKPIMRTARNLFILNLAISDLLLCLVTMPLTL    137
rhopaloa          WYHMLISMYGVLIVFGALGNTLVVIAVVRKPIMRTARNLFILNLAISDLLLCLVTMPLTL    144
ficusphla         WYHMLISMYGVLIVFGALGNTLVVIAVVRKPIMRTARNLFILNLAISDLLLCLVTMPLTL    144
biarmpes          WYHMLISMYGVLIVFGALGNTLVVIAVVRKPIMRTARNLFILNLAISDLLLCLVTMPLTL    144
suzuki            WYHMLISMYGVLILFGALGNTLVVIAVVRKPIMRTARNLFILNLAISDLLLCLVTMPLTL    144
simulans          WYHMLISMYGVLIVFGALGNTLVVIAVIRKPIMRTARNLFILNLAISDLLLCLVTMPLTL    144
erecta            WYHMLISMYGVLIVFGALGNTLVVIAVIRKPIMRTARNLFILNLAISDLLLCLVTMPLTL    144
melanogaster      WYHMLISMYGVLIVFGALGNTLVVIAVIRKPIMRTARNLFILNLAISDLLLCLVTMPLTL    144
sechelia          WYHMLISMYGVLIVFGALGNTLVVIAVIRKPIMRTARNLFILNLAISDLLLCLVTMPLTL    144
mauritania        WYHMLISMYGVLIVFGALGNTLVVIAVIRKPIMRTARNLFILNLAISDLLLCLVTMPLTL    144
takahashi         WYHMLITMYGVLIVFGALGNTLVVIAVVRKPIMRTARNLFILNLAISDLLLCLVTMPLTL    144
eugracilis        WYHMLISMYGVLIVFGALGNTLVVIAVVRKPIMRTARNLFILNLAISDLLLCLVTMPLTL    144
bipectinata       WYHMLISMYGVLIVFGALGNTLVVIAVVRKPIMRTARNLFILNLAISDLLLCLVTMPLTL    150
anannassae        WYHMLISMYGVLIVFGALGNTLVVIAVVRKPIMRTARNLFILNLAISDLLLCLVTMPLTL    151
serrata           WYHMLITMYGVLILFGALGNTLVVIAVVRKPIMRTARNLFILNLAISDLLLCLVTMPLTL    150
```

```
kikkawei          WYHMLITMYGVLILFGALGNTLVVIAVVRKPIMRTARNLFILNLAISDLLLCLVTMPLTL    150
                  ***:**::*.:**:***:** :*****:***:****************************

mojavensis        MELLSKFWPYGSCATLCKTIATLQALSIFVSTISITAIAFDRYQVIVYPTRDSLQFVGAV    229
grimshawi         MEILSKFWPYGSCASLCKMIATLQALSIFVSTISITAIAFDRYQVIVYPTRDSLQFVGAV    231
virilis           MEILSKFWPYGSCAVLCKTIATLQALSIFVSTISITAIAFDRYQVIVYPTRDSLQFVGAV    157
elegans           MEILSKYWPYGSCSILCKTIAMLQALCIFVSTISITAIAFDRYQVIVYPTRDSLQFVGAV    197
rhopaloa          MEILSKYWPYGSCSILCKTIAMLQALCIFVSTISITAIAFDRYQVIVYPTRDSLQFVGAV    204
ficusphla         MEILSKYWPYGSCSILCKTIAMLQALCIFVSTISITAIAFDRYQVIVYPTRNSLQFVGAV    204
biarmpes          MEILSKYWPYGSCSILCKTIAMLQALCIFVSTISITAIAFDRYQVIVYPTRDSLQFVGAV    204
suzuki            MEILSKYWPYGSCSILCKTIAMLQALCIFVSTISITAIAFDRYQVIVYPTRDSLQFVGAV    204
simulans          MEILSKYWPYGSCSILCKTIAMLQALCIFVSTISITAIAFDRYQVIVYPTRDSLQFVGAV    204
erecta            MEILSKYWPYGSCSILCKTIAMLQALCIFVSTISITAIAFDRYQVIVYPTRDSLQFVGAV    204
melanogaster      MEILSKYWPYGSCSILCKTIAMLQALCIFVSTISITAIAFDRYQVIVYPTRDSLQFVGAV    204
sechelia          MEILSKYWPYGSCSILCKTIAMLQALCIFVSTISITAIAFDRYQVIVYPTRDSLQFVGAV    204
mauritania        MEILSKYWPYGSCSILCKTIAMLQALCIFVSTISITAIAFDRYQVIVYPTRDSLQFVGAV    204
takahashi         MEILSKYWPYGSCSILCKTIAMLQALCIFVSTISITAIAFDRYQVIVYPTRDSLQFVGAV    204
eugracilis        MEILSKYWPYGSCSILCKTIAMLQALCIFVSTISITAIAFDRYQVIVYPTRDSLQFVGAV    204
bipectinata       MEILSKYWPFGSCSILCKTIAMLQALCIFVSTISITAIAFDRYQVIVYPTRDSLQFVGAV    210
anannassae        MEILSKYWPFGSCSILCKTIAMLQALCIFVSTISITAIAFDRYQVIVYPTRDSLQFVGAV    211
serrata           MEILSKYWPYGSCSILCKTIAMLQALCIFVSTISITAIAFDRYQVIVYPTRDSLQFVGAV    210
kikkawei          MEILSKYWPYGSCSILCKTIAMLQALCIFVSTISITAIAFDRYQVIVYPTRDSLQFVGAV    210
                  **:***:**:***: *** ** ****.************************:********

mojavensis        TILAGIWTLALILASPLFIYKQLINMDMPLMLQKFGVPHRISYCIEDWPMSDGRFYYSIF    289
grimshawi         TILAFIWILALILASPLFIYKQLINMDMPAVLQDIGVPNRISYCIEDWPLSDGRFYYSIF    291
virilis           AILAGIWILALTLASPLFIYKQLISMDMPPVLQRLGVPHRISYCIEDWPLSDGRFYYSIF    217
elegans           TILAGIWALALLLASPLFVYKELINTDTPALLQQIGLQDTIPYCIEDWPSRNGRFYYSIF    257
rhopaloa          TILAGIWALALLLASPLFVYKELINTDTPALLQQIGLQDTIPYCIEDWPSRNGRFYYSIF    264
ficusphla         GILVGIWALALLLASPLFVYKELINTDTPALLQQIGLQDTIPYCIEDWPSRNGRFYYSIF    264
biarmpes          TILAGIWALSLLLASPLFVYKELINTDTPALLQQIGLQDTIPYCIEDWPTRNGRFYYSIF    264
suzuki            MILAGIWALSLLLASPLFVYKELINTDTPALLQQIGLQDTIPYCIEDWPTRNGRFYYSIF    264
simulans          TILAGIWALALLLASPLFVYKELINTDTPALLQQIGLQDTIPYCIEDWPSRNGRFYYSIF    264
erecta            TILAGIWALALLLASPLFVYKELINTDTPALLQQIGLQDTIPYCIEDWPSRNGRFYYSIF    264
melanogaster      TILAGIWALALLLASPLFVYKELINTDTPALLQQIGLQDTIPYCIEDWPSRNGRFYYSIF    264
sechelia          TILAGIWALALLLASPLFVYKELINTDTPALLQQIGLQDTIPYCIEDWPSRNGRFYYSIF    264
mauritania        TILAGIWALALLLASPLFVYKELINTDTPALLQQIGLQDTIPYCIEDWPSRNGRFYYSIF    264
takahashi         TILAGIWALALLLASPLFVYKELINTDTPALLQQIGLQDTIPYCIEDWPSRNGRFYYSIF    264
eugracilis        TILAGIWALSLLLASPLFVYKELINTDTPALLQQIGLQDTIPYCIEDWPSRNGRFYYSIF    264
bipectinata       TTLACIWALALLLASPLFIYKELINTDTPTLLQQIGFQDTIPFCIEDWPSSNGRFYYSIF    270
anannassae        TTLACIWALALLLASPLFIYKELINTDTPTLLQQMGFQDTIPFCIEDWPSSNGRFYYSIF    271
serrata           TILACIWVLALLLASPLFIYKEMINTETPQLLQQIGLQDRIPYCIEDWPSSNGRFYYSIF    270
kikkawei          TILACIWALALLLASPLFIYKEMINTETPQLLQQIGLQDRIPYCIEDWPSSNGRFYYSIF    270
                    *. ** *:* ******:**::*. : * :** :*. . * :******  :********

mojavensis        SLCVQYLVPIVIVSIAYFGIYNKLKSRITVVAVQC-SSQRKTERGRRMQRTNRLLISIAI    348
grimshawi         SLCVQYLVPILIVSVAYFGIYNKLKSRITVVTGQS-SSQRKVERGRRMKRTNRLLISIAI    350
virilis           SLCVQYLVPILIVSVAYFGIYNKLKSRITVVTVQS-SSQRKVERGRRMKRTNRLLISIAI    276
elegans           SLCVQYLVPILIVSVAYFGIYNKLKSRITVVAVQAASAQRKVERGRRMKRTNCLLISIAI    317
rhopaloa          SLCVQYLVPILIVSVAYFGIYNKLKSRITVVAVQAASAQRKSERGRRMKRTNCLLISIAI    324
ficusphla         SLCVQYLVPILIVSVAYFGIYNKLKSRITVVAVQAASAQRKVERGRRMKRTNCLLISIAI    324
biarmpes          SLCVQYLVPILIVSVAYFGIYNKLKSRITVVAVQAASAQRKVERGRRMKRTNCLLISIAI    324
suzuki            SLCVQYLVPILIVSVAYFGIYNKLKSRITVVAVQAASAQRKVERGRRMKRTNCLLISIAI    324
simulans          SLCVQYLVPILIVSVAYFGIYNKLKSRITVVAVQASSAQRKVERGRRMKRTNCLLISIAI    324
erecta            SLCVQYLVPILIVSVAYFGIYNKLKSRITVVAVQASSAQRKVERGRRMKRTNCLLISIAI    324
melanogaster      SLCVQYLVPILIVSVAYFGIYNKLKSRITVVAVQASSAQRKVERGRRMKRTNCLLISIAI    324
sechelia          SLCVQYLVPILIVSVAYFGIYNKLKSRITVVAVQASSAQRKVERGRRMKRTNCLLISIAI    324
mauritania        SLCVQYLVPILIVSVAYFGIYNKLKSRITVVAVQASSAQRKVERGRRMKRTNCLLISIAI    324
takahashi         SLCVQYLVPILIVSVAYFGIYNKLKSRITVVAVQAASAQRKVERGRRMKRTNCLLISIAI    324
eugracilis        SLCVQYLVPILIVSVAYFGIYNKLKSRITVVAVQAASAQRKVERGRRMKRTNCLLISIAI    324
bipectinata       SLCVQYLVPILIVSVAYFGIYNKLKSRITVVAVQAASAQRKVERGRRMKRTNCLLISIAV    330
anannassae        SLCVQYLVPILIVSVAYFGIYNKLKSRITVVAVQAASAQRKVERGRRMKRTNCLLISIAV    331
serrata           SLCVQYLVPILIVSVAYFGIYNKLRSRITVVAVQASSAQRKVERGRRMKRTNCLLISIAI    330
kikkawei          SLCVQYLVPILIVSVAYFGIYNKLKSRITVVAVQASSAQRKVERGRRMKRTNCLLISIAI    330
                  **********:***:*********:******: *. *:*** ******:*** ******:

mojavensis        IFGVSWLPLNFINLYADMQRPSAVTPRMIVAYAICHMIGMSSACSNPLLYGWLNDNFRSN    408
grimshawi         IFGVSWLPLNFFNLYADLQHPSAVTQRMLVAYAICHMIGMSSACSNPLLYGWLNDNFRSN    410
virilis           IFGVSWLPLNFFNLYADLQHPSAVTQRMLVAYAICHMIGMSSACSNPLLYGWLNDNFRSN    336
elegans           IFGVSWLPLNFFNLYADMERS-PVTQNMLVRYAICHMIGMSSACSNPLLYGWLNDNFRCN    376
rhopaloa          IFGVSWLPLNFFNLYADMERS-PVTQNMLVRYAICHMIGMSSACSNPLLYGWLNDNFRKE    383
```

```
ficusphla         IFGVSWLPLNFFNLYADMERS-PVTQSMLVRYAICHMIGMSSACSNPLLYGWLNDNFRKE    383
biarmpes          IFGVSWLPLNFFNLYADMERS-PVTQSMLVRYAICHMIGMSSACSNPLLYGWLNDNFRCS    383
suzuki            IFGVSWLPLNFFNLYADMERS-PVTQSMLVRYAICHMIGMSSACSNPLLYGWLNDNFRCN    383
simulans          IFGVSWLPLNFFNLYADMERS-PVTQSMLVRYAICHMIGMSSACSNPLLYGWLNDNFRKE    383
erecta            IFGVSWLPLNFFNLYADMERS-PVTQSMLVRYAICHMIGMSSACSNPLLYGWLNDNFR--    381
melanogaster      IFGVSWLPLNFFNLYADMERS-PVTQSMLVRYAICHMIGMSSACSNPLLYGWLNDNFR--    381
sechelia          IFGVSWLPLNFFNLYADMERS-PVTQSMLVRYAICHMIGMSSACSNPLLYGWLNDNFR--    381
mauritania        IFGVSWLPLNFFNLYADMERS-PVTQSMLVRYAICHMIGMSSACSNPLLYGWLNDNFR--    381
takahashi         IFGVSWLPLNFFNLYADMERS-PVTQSMLVRYAICHMIGMSSACSNPLLYGWLNDNFRKE    383
eugracilis        IFGVSWLPLNFFNLYADMERS-PVTQSMLVRYAICHMIGMSSACSNPLLYGWLNDNFRKE    383
bipectinata       IFGVSWLPLNFFNLYADMQQS-PVTQNMLVKYAICHMIGMSSACSNPLLYGWLNDNFR--    387
anannassae        IFGVSWLPLNFFNLYADMQQS-PVTQNMLVKYAICHMIGMSSACSNPLLYGWLNDNFR--    388
serrata           IFGVSWLPLNFFNLYADMQRP-VMTQKMLVAYAVCHMIGMSSACSNPLLYGWLNDNFRKE    389
kikkawei          IFGVSWLPLNFFNLYADMQRP-VMTQKMLVAYAVCHMIGMSSACSNPLLYGWLNDNFR--    387
                  ***********:*****:::   :*  *:* **:************************

mojavensis        ------------------------VQAAATR-RRRKHEADLSKGELQLLGKSATRGLA--    441
grimshawi         ------------------------VQAAASRHRQRKHDAHLSKGELQLLGNP--SC----    440
virilis           ------------------------MQAAAARRRRK--------HELQLLGNSVQRG----    360
elegans           VHS-----------------------AARRRRKLGAGADLSRGELKLLGPGGAQSGTVG    412
rhopaloa          FHELLCLCSEPTNVALNGHTTGCNVQA-ARRRRK--LGANLSRGELKLLGPGGAQSGTAG    440
ficusphla         FHELLCRCSEPTNVALNGHTTGCNVQAAARRRRK--LGADLSKGELKLLGPGGAQSGTVG    441
biarmpes          V-------------------------QAARRRRK--LGGDLSKGELKLLGPGGAQSGTVG    416
suzuki            V-------------------------QAARRRRK--LGGDDSKGELKLLGPGGAQSGTVG    416
simulans          FQELLCRCSD-TNVALNGHTTGCNVQAAARRRRK--LGAELSKGELKLLGPGGAQSGTAG    440
erecta            ----------------------CNVQAAARRRRK--LGAELSKGELKLLGPGGAQSGTAG    417
melanogaster      ----------------------CNVQAAARRRRK--LGAELSKGELKLLGPGGAQSGTAG    417
sechelia          ----------------------CNVQAAARRRRK--LGAELSKGELKLLGPCGAQSGTAG    417
mauritania        ----------------------CNVQAAARRRRK--LGAELSKGELKLLGPGGAQSGTAG    417
takahashi         FQELLCRCSD-TNVALNGHTTGCNVQAAARRRRK--LGADLSKGELKLLGPGGAQSGTVG    440
eugracilis        FQELLCRCSD-TNVALNGHTTGCNVQAAARRRRK--LGADLSKGELKLLGPGGAQSGTVG    440
bipectinata       ----------------------CNVQAAAARRR-RKLGGDLTKGELKLLGKGGASQCGGG    424
anannassae        ----------------------CNVQAAAARRR-RKLGGDLTKGEQKLLGKGGASQCGG-    424
serrata           FQELLCRCSEPTNVALNGHTTGCNVQAAAARRR-RKLGTDLSKGDLKLLGQSGAQSGAP-    447
kikkawei          ----------------------CNVQAAAARRR-RKLGTDLSKGELKLLGQSGAQSGAP-    423
                                              * * *           : :***

mojavensis        -----TSDGDCLGGVSIAATDFNAR----NGTRSAVTESVALTE-NPMPSELIIILAPP--    489
grimshawi         -----ATDGDCIGGVSIAATEFNTR-HTVNGTRSCITESVALTE-NPMPTEMTTLVPRLQ    493
virilis           -----ASDGDCLGGMSIAATEFNTR-HTVNGTRSAVTESVALTE-SPMPSEMTTLVPR--    411
elegans           ----------GEGVTGLAATDFMTGGHHAGGLRSAITESVALTE--NMPSEVTQLMPRLE    460
rhopaloa          ----------GEGGLGMAATDFMTG-HHEGGLRSAITESVALTE--NMPSEVTKLMPRLE    487
ficusphla         -----------GE-GGLAATDFMTG-HQEGGLRSAITESVALTEH-QVPSEVTKLMPRLE    487
biarmpes          ----------GGE-GGLAATDFMTG-HQEGGLRSAITESVALTDHNPVPSEVTKLMPRLE    464
suzuki            ----------GGE-GGLAATDFMTG-HQEGGLRSAITESVALTDHNPVPSEVTKLMPRLE    464
simulans          -----------GE-GGLAATDFMTG-HHEGGLRSAITESVALTDHNPVPSEVTKLMPR--    485
erecta            -----------GE-GGLAATDFMTG-HHEGGLRSAITESVALTDHNPVPSEVTKLMPR--    462
melanogaster      -----------GE-GGLAATDFMTG-HHEGGLRSAITESVALTDHNPVPSEVTKLMPR--    462
sechelia          -----------GE-GGLAATDFMTG-HHEGGLRSAITESVALTDHNPVPSEVTKLMPRLE    464
mauritania        -----------GE-GGLAATDFMTG-HHEGGLRSAITESVALTDHNPVPSEVTKLMPR--    462
takahashi         -----------GE-GGLAATDFMTG-HQEGGLRSAITESVALTDHNPVPSEVTKLMPRLE    487
eugracilis        -----------GD-GGLAATDFMTG-HQEGGLRSAITESVALTDHNPVPSEVTKLMPRLE    487
bipectinata       ASTFGDGGGPDGGCGSMAATDFMTG-NPEYGLRSAITESVALTE-NPMPSEVTKLMPRLE    482
anannassae        -----------GGCGSMAATDFMTG-NPECGLRSAITESVALTE-NPMPSEITKLMPR--    469
serrata           ------GG--GAGGGSMAATDFMTG-HQECGLRSAITESVALTE-NPMPSEVTKLMPRLE    497
kikkawei          ------GG---AGGGSMAATDFMTG-HQECGLRSAITESMALTE-NPMPSEVTQLMPRLE    472
                                 .:***:* :      * **.:***:***:   :*:*:  * *

mojavensis        --   489
grimshawi         QY   495
virilis           --   411
elegans           QY   462
rhopaloa          QY   489
ficusphla         QY   489
biarmpes          QY   466
suzuki            QY   466
simulans          --   485
erecta            --   462
melanogaster      --   462
sechelia          QY   466
mauritania        --   462
```

```
takahashi         QY   489
eugracilis        QY   489
bipectinata       QY   484
anannassae        --   469
serrata           QY   499
kikkawei          QY   474
```

> melanogaster AAF51909.3

```
  1 miismnqtep aqladgehls gyasssnsvr ylddrhpldy ldlgtvhaln ttaintsdln
       61 etgsrpldpv lidrflsnra vdspwyhmli smygvlivfg algntlvvia virkpimrta
      121 rnlfilnlai sdlllclvtm pltlmeilsk ywpygscsil cktiamlqal cifvstisit
      181 aiafdryqvi vyptrdslqf vgavtilagi walalllasp lfvykelint dtpallqqig
      241 lqdtipycie dwpsrngrfy ysifslcvqy lvpilivsva yfgiynklks ritvvavqas
      301 saqrkvergr rmkrtnclli siaiifgvsw lplnffnlya dmerspvtqs mlvryaichm
      361 igmssacsnp llygwlndnf rcnvqaaarr rrklgaelsk gelkllgpgg aqsgtaggeg
      421 glaatdfmtg hhegglrsai tesvaltdhn pvpsevtklm pr
```

> erecta XP_015009571.1

```
  1 miismnqtea aqlaagehlg gyasssnsvr ylddrhpldy ldlgtvhaln ttaintseln
       61 etgsrpldpv lidrflsnra vdspwyhmli smygvlivfg algntlvvia virkpimrta
      121 rnlfilnlai sdlllclvtm pltlmeilsk ywpygscsil cktiamlqal cifvstisit
      181 aiafdryqvi vyptrdslqf vgavtilagi walalllasp lfvykelint dtpallqqig
      241 lqdtipycie dwpsrngrfy ysifslcvqy lvpilivsva yfgiynklks ritvvavqas
      301 saqrkvergr rmkrtnclli siaiifgvsw lplnffnlya dmerspvtqs mlvryaichm
      361 igmssacsnp llygwlndnf rcnvqaaarr rrklgaelsk gelkllgpgg aqsgtaggeg
      421 glaatdfmtg hhegglrsai tesvaltdhn pvpsevtklm pr
```

> biarmpes XP_043951464.1

```
  1 miismnqtep gqlaaaehlg gyasssnsgr ylddrhpldy ldlgtihaln ttaintsdln
       61 etgsrpldpv lidrflsnra vdspwyhmli smygvlivfg algntlvvia vvrkpimrta
      121 rnlfilnlai sdlllclvtm pltlmeilsk ywpygscsil cktiamlqal cifvstisit
      181 aiafdryqvi vyptrdslqf vgavtilagi walslllasp lfvykelint dtpallqqig
      241 lqdtipycie dwptrngrfy ysifslcvqy lvpilivsva yfgiynklks ritvvavqaa
      301 saqrkvergr rmkrtnclli siaiifgvsw lplnffnlya dmerspvtqs mlvryaichm
      361 igmssacsnp llygwlndnf rcsvqaarrr rklggdlskg elkllgpgga qsgtvgggeg
      421 glaatdfmtg hqegglrsai tesvaltdhn pvpsevtklm prleqy
```

> suzuki XP_036675105.1

```
1 miismnqtep gllatgehlg gyasssnsgr ylddrhpldy ldlgtihaln ttaintsdln
       61 etasrpldpv lidrflsnra vdspwyhmli smygvlilfg algntlvvia vvrkpimrta
      121 rnlfilnlai sdlllclvtm pltlmeilsk ywpygscsil cktiamlqal cifvstisit
      181 aiafdryqvi vyptrdslqf vgavmilagi walslllasp lfvykelint dtpallqqig
      241 lqdtipycie dwptrngrfy ysifslcvqy lvpilivsva yfgiynklks ritvvavqaa
      301 saqrkvergr rmkrtnclli siaiifgvsw lplnffnlya dmerspvtqs mlvryaichm
      361 igmssacsnp llygwlndnf rcnvqaarrr rklggddskg elkllgpgga qsgtvgggeg
      421 glaatdfmtg hqegglrsai tesvaltdhn pvpsevtklm prleqy
```

> sechelia XP_032578233.1

1 miismnqtep tqlaagehls gyasssnsvr ylddrhpldy ldlgtvhaln ttaintsdln
61 etgsrpldpv lidrflsnra vdspwyhmli smygvlivfg algntlvvia virkpimrta
121 rnlfilnlai sdlllclvtm pltlmeilsk ywpygscsil cktiamlqal cifvstisit
181 aiafdryqvi vyptrdslqf vgavtilagi walalllasp lfvykelint dtpallqqig
241 lqdtipycie dwpsrngrfy ysifslcvqy lvpilivsva yfgiynklks ritvvavqas
301 saqrkvergr rmkrtnclli siaiifgvsw lplnffnlya dmerspvtqs mlvryaichm
361 igmssacsnp llygwlndnf rcnvqaaarr rrklgaelsk gelkllgpcg aqsgtaggeg
421 glaatdfmtg hhegglrsai tesvaltdhn pvpsevtklm prleqy

> simulans XP_039150862.1

1 miismnqtep tqlaagehls gyasssnsvr ylddrhpldy ldlgtvhaln ttaintsdln
61 etgsrpldpv lidrflsnra vdspwyhmli smygvlivfg algntlvvia virkpimrta
121 rnlfilnlai sdlllclvtm pltlmeilsk ywpygscsil cktiamlqal cifvstisit
181 aiafdryqvi vyptrdslqf vgavtilagi walalllasp lfvykelint dtpallqqig
241 lqdtipycie dwpsrngrfy ysifslcvqy lvpilivsva yfgiynklks ritvvavqas
301 saqrkvergr rmkrtnclli siaiifgvsw lplnffnlya dmerspvtqs mlvryaichm
361 igmssacsnp llygwlndnf rkefqellcr csdtnvalng httgcnvqaa arrrrklgae
421 lskgelkllg pggaqsgtag gegglaatdf mtghhegglr saitesvalt dhnpvpsevt
481 klmpr

> takahashi XP_016993445.2

1 miismnqtep gqlaageqlg gyasssnsgr ylddrhpldy ldlgtvhaln ttaintsdln
61 etgsrpldpv lidrflsnra vdspwyhmli tmygvlivfg algntlvvia vvrkpimrta
121 rnlfilnlai sdlllclvtm pltlmeilsk ywpygscsil cktiamlqal cifvstisit
181 aiafdryqvi vyptrdslqf vgavtilagi walalllasp lfvykelint dtpallqqig
241 lqdtipycie dwpsrngrfy ysifslcvqy lvpilivsva yfgiynklks ritvvavqaa
301 saqrkvergr rmkrtnclli siaiifgvsw lplnffnlya dmerspvtqs mlvryaichm
361 igmssacsnp llygwlndnf rkefqellcr csdtnvalng httgcnvqaa arrrrklgad
421 lskgelkllg pggaqsgtvg gegglaatdf mtghqegglr saitesvalt dhnpvpsevt
481 klmprleqy

> bipectinata XP_043068022.1

1 miigmnqtes gplatgdrls gyassgnsvr ylddrhpldy ldlgsvvsna haalnstats
61 nlsdandtga rpldpvlidr flsnravdsp wyhmlismyg vlivfgalgn tlvviavvrk
121 pimrtarnlf ilnlaisdll lclvtmpltl meilskywpf gscsilckti amlqalcifv
181 stisitaiaf dryqvivypt rdslqfvgav ttlaciwala lllasplfiy kelintdtpt
241 llqqigfqdt ipfciedwps sngrfyysif slcvqylvpi livsvayfgi ynklksritv
301 vavqaasaqr kvergrrmkr tnclliisiav ifgvswlpln ffnlyadmqq spvtqnmlvk
361 yaichmigms sacsnpllyg wlndnfrcnv qaaaarrrrk lggdltkgel kllgkggasq
421 cgggastfgd gggpdggcgs maatdfmtgn peyglrsait esvaltenpm psevtklmpr
481 leqy

> rhopaloa XP_016970622.1

1 miismnqtet gplaagehls gyasssnsgr ylddrhpldy ldlgivhaln ttaintseln
61 etgsrpldqv ligrflsnra vdspwyhmli smygvlivfg algntlvvia vvrkpimrta
121 rnlfilnlai sdlllclvtm pltlmeilsk ywpygscsil cktiamlqal cifvstisit
181 aiafdryqvi vyptrdslqf vgavtilagi walalllasp lfvykelint dtpallqqig
241 lqdtipycie dwpsrngrfy ysifslcvqy lvpilivsva yfgiynklks ritvvavqaa
301 saqrksergr rmkrtnclli siaiifgvsw lplnffnlya dmerspvtqn mlvryaichm

```
361 igmssacsnp llygwlndnf rkefhellcl cseptnvaln ghttgcnvqa arrrrklgan
421 lsrgelkllg pggaqsgtag gegglgmaat dfmtghhegg lrsaitesva ltenmpsevt
481 klmprleqy
```

> eugracilis XP_017082328.2

```
  1 miismnqtep gplaagehls gyasssnsvr ylddrhpldy ldlgmvhaln ttaintsemn
 61 etgsrpldpv lidrflsnra vdspwyhmli smygvlivfg algntlvvia vvrkpimrta
121 rnlfilnlai sdlllclvtm pltlmeilsk ywpygscsil cktiamlqal cifvstisit
181 aiafdryqvi vyptrdslqf vgavtilagi walslllasp lfvykelint dtpallqqig
241 lqdtipycie dwpsrngrfy ysifslcvqy lvpilivsva yfgiynklks ritvvavqaa
301 saqrkvergr rmkrtnclli siaiifgvsw lplnffnlya dmerspvtqs mlvryaichm
361 igmssacsnp llygwlndnf rkefqellcr csdtnvalng httgcnvqaa arrrrklgad
421 lskgelkllg pggaqsgtvg gdgglaatdf mtghqegglr saitesvalt dhnpvpsevt
481 klmprleqy
```

> ficusphla XP_017046702.1

```
  1 miismnqtet gqlsagehls gyassgnsvr ylddrhpldy ldlgtvhaln tsamntsdad
 61 etgsrpldpv lidrflsnra vdspwyhmli smygvlivfg algntlvvia vvrkpimrta
121 rnlfilnlai sdlllclvtm pltlmeilsk ywpygscsil cktiamlqal cifvstisit
181 aiafdryqvi vyptrnslqf vgavgilvgi walalllasp lfvykelint dtpallqqig
241 lqdtipycie dwpsrngrfy ysifslcvqy lvpilivsva yfgiynklks ritvvavqaa
301 saqrkvergr rmkrtnclli siaiifgvsw lplnffnlya dmerspvtqs mlvryaichm
361 igmssacsnp llygwlndnf rkefhellcr cseptnvaln ghttgcnvqa aarrrrklga
421 dlskgelkll gpggaqsgtv ggegglaatd fmtghqeggl rsaitesval tehqvpsevt
481 klmprleqy
```

> anannassae EDV43731.2

```
  1 mmiigmnqte sgplatgdrl sgyassgnsg rylddrhpld yldlgsvvsn ahaalnssan
 61 nnfseandtg arpldpvlid rflsnravds pwyhmlismy gvlivfgalg ntlvviavvr
121 kpimrtarnl filnlaisdl llclvtmplt lmeilskywp fgscsilckt iamlqalcif
181 vstisitaia fdryqvivyp trdslqfvga vttlaciwal alllasplfi ykelintdtp
241 tllqqmgfqd tipfciedwp ssngrfyysi fslcvqylvp ilivsvayfg iynklksrit
301 vvavqaasaq rkvergrrmk rtnclllisia vifgvswlpl nffnlyadmq qspvtqnmlv
361 kyaichmigm ssacsnplly gwlndnfrcn vqaaaarrrr klggdltkge qkllgkggas
421 qcggggcgsm aatdfmtgnp ecglrsaite svaltenpmp seitklmpr
```

> serrata XP_020798669.1

```
  1 miismnqtes vplaagdrls gfaaggdnsv rylddrhpld yldlgsvvgt ahavlnataa
 61 nmselnetgs rpldpvlidr ylsnravdsp wyhmlitmyg vlilfgalgn tlvviavvrk
121 pimrtarnlf ilnlaisdll lclvtmpltl meilskywpy gscsilckti amlqalcifv
181 stisitaiaf dryqvivypt rdslqfvgav tilaciwvla lllasplfiy kemintetpq
241 llqqiglqdr ipyciedwps sngrfyysif slcvqylvpi livsvayfgi ynklrsritv
301 vavqassaqr kvergrrmkr tncllisiai ifgvswlpln ffnlyadmqr pvmtqkmlva
361 yavchmigms sacsnpllyg wlndnfrkef qellcrcsep tnvalnghtt gcnvqaaaar
421 rrrklgtdls kgdlkllgqs gaqsgapggg agggsmaatd fmtghqecgl rsaitesval
481 tenpmpsevt klmprleqy
```

> kikkawei XP_017016715.1

```
  1 miislnqtes vplaagdrls gfagggdnsv rylddrhpld yldlgsvvgt vhavinatat
 61 nmselnetgs rpldpvlidr ylsnravdsp wyhmlitmyg vlilfgalgn tlvviavvrk
121 pimrtarnlf ilnlaisdll lclvtmpltl meilskywpy gscsilckti amlqalcifv
181 stisitaiaf dryqvivypt rdslqfvgav tilaciwala lllasplfiy kemintetpq
241 llqqiglqdr ipyciedwps sngrfyysif slcvqylvpi livsvayfgi ynklksritv
301 vavqassaqr kvergrrmkr tncllisiai ifgvswlpln ffnlyadmqr pvmtqkmlva
361 yavchmigms sacsnpllyg wlndnfrcnv qaaaarrrrk lgtdlskgel kllgqsgaqs
421 gapggagggs maatdfmtgh qecglrsait esmaltenpm psevtqlmpr leqy
```

> mauritania **XP_033166991.1**

```
  1 miismnqtet tqlaagehls gyasssnsvr ylddrhpldy ldlgtvhaln ttaintsdln
 61 etgsrpldpv lidrflsnra vdspwyhmli smygvlivfg algntlvvia virkpimrta
121 rnlfilnlai sdlllclvtm pltlmeilsk ywpygscsil cktiamlqal cifvstisit
181 aiafdryqvi vyptrdslqf vgavtilagi walalllasp lfvykelint dtpallqqig
241 lqdtipycie dwpsrngrfy ysifslcvqy lvpilivsva yfgiynklks ritvvavqas
301 saqrkvergr rmkrtnclli siaiifgvsw lplnffnlya dmerspvtqs mlvryaichm
361 igmssacsnp llygwlndnf rcnvqaaarr rrklgaelsk gelkllgpgg aqsgtaggeg
421 glaatdfmtg hhegglrsai tesvaltdhn pvpsevtklm pr
```

> elegans **XP_017130917.1**

```
  1 miismnqtae hlsgyvsssn sgrylddrhp ldyldlgpar alnatavnts emnetgsrpl
 61 dpvlidrfls nravdspwyh mlismygvli vfgalgntlv viavvrkpim rtarnlfiln
121 laisdlllcl vtmpltlmei lskywpygsc silcktiaml qalcifvsti sitaiafdry
181 qvivyptrds lqfvgavtil agiwalalll asplfvykel intdtpallq qiglqdtipy
241 ciedwpsrng rfyysifslc vqylvpiliv svayfgiynk lksritvvav qaasaqrkve
301 rgrrmkrtnc llisiaiifg vswlplnffn lyadmerspv tqnmlvryai chmigmssac
361 snpllygwln dnfrcnvhsa arrrrklgag adlsrgelkl lgpggaqsgt vggegvtgla
421 atdfmtgghh agglrsaite svaltenmps evtqlmprle qy
```

> grimshawi **XP_032597472.1**

```
  1 miiamnrtef gsqlpnfdss veifkaiarn snfdligerr hlvnysqlna lsdvnanntn
 61 ksnynishss nnmtiqllns thinvsniiv ssdldpvlmd qyqhnraies pwyhlliamy
121 silivfgamg nimvviavlr kplmrtarnl filnlaisdl llclvtmplt lmeilskfwp
181 ygscaslckm iatlqalsif vstisitaia fdryqvivyp trdslqfvga vtilafiwil
241 alilasplfi ykqlinmdmp avlqdigvpn risyciedwp lsdgrfyysi fslcvqylvp
301 ilivsvayfg iynklksrit vvtgqsssqr kvergrrmkr tnrllisiai ifgvswlpln
361 ffnlyadlqh psavtqrmlv ayaichmigm ssacsnplly gwlndnfrsn vqaaasrhrq
421 rkhdahlskg elqllgnpsc atdgdciggv siaatefntr htvngtrsci tesvaltenp
481 mptemttlvp rlqqy
```

> virilis XP_002058565

```
  1 mtillllnstn nesnfmpadm dpvlmdqylh nraiespwyh lliamysvli vfgamgnimv
 61 viavvrkpim rtarnlfiln laisdlllcl vtmpltlmei lskfwpygsc avlcktiatl
121 qalsifvsti sitaiafdry qvivyptrds lqfvgavail agiwilaltl asplfiykql
181 ismdmppvlq rlgvphrisy ciedwplsdg rfyysifslc vqylvpiliv svayfgiynk
241 lksritvvtv qsssqrkver grrmkrtnrl lisiaiifgv swlplnffnl yadlqhpsav
301 tqrmlvayai chmigmssac snpllygwln dnfrsnmqaa aarrrrkhel qllgnsvqrg
361 asdgdclggm siaatefntr htvngtrsav tesvaltesp mpsemttlvp r
```

> mojavensis XP_043863318.1

```
1 mnrtelgsqi lssaelykal ssdatfdtig esrhlvnypg ldaggaytrn hnydklnsny
      61 edssssssnd ndsmtmllln stsngsyvpa gmdpvlmdqy lhnrsigspw yhlliaiygv
     121 livfgamgni mvviavlrkp imrtarnlfi lnlaisdlll clvtmpltlm ellskfwpyg
     181 scatlcktia tlqalsifvs tisitaiafd ryqvivyptr dslqfvgavt ilagiwtlal
     241 ilasplfiyk qlinmdmplm lqkfgvphri syciedwpms dgrfyysifs lcvqylvpiv
     301 ivsiayfgiy nklksritvv avqcssqrkt ergrrmqrtn rllisiaiif gvswlplnfi
     361 nlyadmqrps avtprmivay aichmigmss acsnpllygw lndnfrsnvq aaatrrrrkh
     421 eadlskgelq llgksatrgl atsdgdclgg vsiaatdfna rngtrsavte svaltenpmp
     481 seliiilapp
```
