## Supplementary material for "The incidence of candidate binding sites for β-arrestin in Drosophila neuropeptide GPCRs": S16 Text

***S16. Text*** **Multi-species analysis of Hector**
**Supporting Figure 18\9**

CLUSTAL Line-ups; Genbank Reference IDs below
Predicted TM domains in YELLOW
BBS sequences in RED
missing mauritania

CLUSTAL

```
bipectinata       -----------mgglsasvtep-----------dmessraqdapqpqdnlriflkhlyae    38
anannasae         -----------mgglsasvtep-----------dme----qeapqpqdnlriflkhlyae    34
kikkawei          mtsi--eeapptmgtnasdsdsdsgsenvamatatasqqsytqpqaqdnlriflrhlyae    58
seratta           ----------------------------mamakasgsqqsssqtqaqdnlriflrhlyae    32
eugracilis        mtrvthietptmvgttasese----senv---------emasqaqtqdnllvflthlyae    47
ficusphelia       ---------mgt-gasssdsg----sekl---------etaaqaqtqdnlriflkhlyae    37
erecta            mtavthteqptm-attssdse----plnl---------dvasqaqtqdnlriflkhlyae    46
melanogaster      -----------m-attssdse----sqnv---------dvasqaqtqdnlriflkhlyae    35
simulans          -----------m-attssdsd----venv---------dvasqaqtqdnlriflkhlyae    35
sechellia         -----------m-attssdsd----venv---------dvasqaqtqdnlriflrhlyae    35
elegans           mttvthfdtptm-gtvasdst----menv---------etatqsqtqdnlriflkhlyae    46
rhopaloa          -----------m-gtvasdsv----penv---------etatqsqtqdnlriflkhlyae    35
takahashi         mtrvshtepptm-gaaasdsq----senv---------eaasqsqmqdnlriflkhlyae    46
Suzuki            mtrvspteaptm-gtaasvse----senv---------evasqtqtqdnlriflkhlyae    46
biarmipes         mtraspieaptm-gtaasese----senv---------eaasqtqtqdnlriflkhlyae    46
grimshawi         ---------------------------------mtasasasmspqpqdnlltflwhlyae    27
virilism          -----------mallraaltma--taayfemtvsga-----mppqpqdnlrtflkhlyae    42
mojavensis        -----------mallraalamt--taayietamtasltkpqprpqpqdnlriflrhlyve    47
                                                              * ****  ** ***.*

bipectinata       cvyryqndtyddpste------satdyesdfpenfspvprylenaalnegsidmrnvdek    92
anannasae         cvyryqndtyddpmte------tatdyesdlpenfspvprylenaalnegsidmrnvdek    88
kikkawei          cvyryqndtyddpsvsv-vgeaaatdydldlpenfspvprylenavlnegaidmrnvdve    117
seratta           cvyryqndtyddpsvsv-dga-aatdyeselpenfspvprylenavlnegaidmrnvdve    90
eugracilis        cvhryqnvtndtedsslslgaatdsend--lkgnfstvprylenaalnegtidmrnvdek    105
ficusphelia       cvyryqnvtyeaddpsvplgaateydvdanapdnfspvpryledavlnegaidmrhvdee    97
erecta            cvfryqnvtydtgdpsfslgpatdydsd--lpenfspvprylenaamnegvidmrsvdee    104
melanogaster      cvfryqnvtydtddpsfslgpatdydsd--lpenfspvprylenaamnegvidmrnvdee    93
simulans          cvfryqnvtydtddpsfslgpatdydsd--lpenfspvprylenaamnegvidmrnvdee    93
sechellia         cvfryqnvtydtddpsfslgpatdydsd--lpenfspvprylenaamnegvidmsnvdek    93
elegans           cvyryknvtydaadpsvslvaatdyesd--ltenfspvpryledavlnegaidmrnvdee    104
rhopaloa          cvyryqndtydaddpsvslgaatdydld--lpesfspvpryledavlnegaidmrnvdee    93
takahashi         cvyryqnvtydtedpsvplgaatdydfd--lpenfspvprylenaalnegaidmrnvdav    104
Suzuki            cvyryqnvtydtddpsvslgaatdydfd--lpenfspvprylenaamnegaidmrnvdae    104
biarmipes         cvyryqnvtydtddpsvslgaatdydfd--lpenfspvprylenaalnegaidmrnvdae    104
grimshawi         cvyryqndtspgqqat------dpgdgniqllepytrmpnylemavlnegtidmqdvnee    81
virilism          cvyryqndtataqlat------epddggl-lletytmipryleqavlnegtidmqdvdee    95
mojavensis        cvyryqndtakgda-k------dsgdeel-mlatfttvpryleqavlnegtidmqdvdee    99
                  **.**:* *                :        :: :*.*** *.:*** ***  *:

bipectinata       kaereelfatiltatmatnqnqddpagqrspgsns-sssrrlfcpldfdgylcwprtpag    151
anannasae         qaekeelfatiltatmatnqnqdepagqrssgnssssgskrlfcpldfdgylcwprtpag    148
kikkawei          qaekeellatvlsgtmatnqqkqshqatt------tttgerlfcplnfdgylcwprtpag    171
seratta           laekeellatvlsvtmatnqqesehhevt--ttnnrsagerlfcplnfdgylcwprtpag    148
eugracilis        laeqeelmatvssatmatnqr-----------------drrlfcpldfdgylcwprtpag    148
ficusphelia       aaekeelmatvftatmatnqr-----------------eerlfcplnfdgylcwprtpag    140
erecta            laekeelmatvvsatmatnqk-----------------ehrlfcplnfdgylcwprtpag    147
melanogaster      laekeelmatvvsatmatnqk-----------------enrlfcplnfdgylcwprtpag    136
simulans          laekeelmatvvsatmatnqk-----------------esrlfcplnfdgylcwprtpag    136
sechellia         laekeelmatvvsatmatnqk-----------------esrlfcplnfdgylcwprtpag    136
elegans           laekeelmatvlsatmatnqq-----------------epllfcplnfdgylcwprtpag    147
rhopaloa          laekeelmatvlsatmatnqrp----------------qqrlfcplnfdgylcwprtpag    137
takahashi         laekeelmatvvsatmatnqrqaeeg---------geeerrlfcplnfdgylcwprtpag    155
Suzuki            laekeelvatvvsatmatnqr----------------eeprlfcplnfdgylcwprtpag    148
biarmipes         qaekeelvatvvsatmatnqr----------------aeprlfcplnfdgylcwprtpag    148
grimshawi         asslnelyatvlsatmapnqhsiqyngnnsna-nttremeilycpvnfdgylcwprtpag    140
virilism          aasenelyatvlsatmatnhhs---------h-n-tsemetlycpvnfdgylcwprtpag    144
mojavensis        aasenelyatihsmtmetnqhs---------r-natsemetlycpvnfdgylcwprtpag    149
```

```
                  :. :** **: : **  *:.                    *:**::*************

bipectinata       TVLSQYCPDFVEGFNSKFLAHKTCQENGSWYRHPETNKTWSNYTNCVDYVDFEFRQFINE    211
anannasae         TVLSQYCPDFVEGFNSKFLAHKTCQENGSWYRHPESNKTWSNYTNCVDYVDLEFRQFINE    208
kikkawei          TVLSQYCPDFVEGFNSKFLAHKTCLENGSWFRHPMTNRTWSNYTNCVDYEDLEFRQFINE    231
seratta           TVLSQYCPDFVEGFNSKFLAHKTCLENGSWFRHPMSNQTWSNYTNCVDYEDLEFRQFINE    208
eugracilis        TVLSQYCPDFVEGFNKKFLAHKTCLENGSWFRHPATNQTWSNYTNCVDHEDLEFRKFINE    208
ficusphelia       TVLSQYCPDFVEGFNSKFLAHKTCLENGSWFRHPVSNQTWSNYTSCVDYEDLEFRQFINE    200
erecta            TVLSQYCPDFVEGFNRKFLAHKTCLENGSWYRHPVSNQTWSNYTNCVDYKDLEFRQFINE    207
melanogaster      TVLSQYCPDFVEGFNRKFLAHKTCLENGSWYRHPVSNQTWSNYTNCVDYEDLEFRQFINE    196
simulans          TVLSQYCPDFVEGFNRKFLAHKTCLENGSWYRHPVSNQTWSNYTNCVDYEDLEFRQFINE    196
sechellia         TVLSQYCPDFVEGFNRKFLAHKTCLENGSWYRHPVSNQTWSNYTNCVDYEDLEFRQFINE    196
elegans           TVLSQYCPDFVEGFNSKFLAHKTCLENGSWFRHPVSNQTWSNYTNCVDYEDLEFRQFINE    207
rhopaloa          TVLSQYCPEFVEGFNSKFLAHKTCLENGSWFRHPVSNQTWSNYTNCVDYEDLEFRQFINE    197
takahashi         TVLSQYCPDFVEGFNSKFLAHKTCLENGSWFRHPVSNQTWSNYTNCVDYEDLEFRQFINE    215
Suzuki            TVLSQYCPDFVEGFNRKFLAHKTCLENGSWFRHPESNQTWSNYTNCVDYEDLEFRQFVNE    208
biarmipes         TVLSQYCPDFVEGFNRKFLAHKTCLENGSWFRHPESNQTWSNYTNCVDYEDLEFRQFVNE    208
grimshawi         TVLSQYCPDFVEGFNTKFLAHKTCMETGSWFRHPVSNQTWSNYTNCVDYEDLQFRQIVNE    200
virilism          TVLSQYCPDFVEGFNSKFLAHKTCLETGSWFRHPVSNQTWSNYTNCVDYEDFQFRQFVNE    204
mojavensis        TVLSQYCPDFVEGFSSKFLAHKTCLENGTWYRHPVSNQTWSNYTNCVDYDDFQFRQFVNE    209
                  ********:*****. ******** *.*:*:*** :*:******.***: *::**:::**

bipectinata       LYVKGYALSLLALLISIIIFLGFKSLRCTRIRIHVHLFASLACTCVAWILWYRLVVEQPE    271
anannasae         LYVKGYAISLLALLISIIIFLGFKSLRCTRIRIHVHLFASLACTCVAWILWYRLVVERPE    268
kikkawei          LYVKGYALSLLALLVSIIIFLGFKSLRCTRIRIHVHLFASLAFTCVAWILWYRLVVERPE    291
seratta           LYVKGYALSLLALLVSILIFLGFKSLRCTRIRIHVHLFASLAFTCVAWILWYRLVVERPE    268
eugracilis        LYVKGYALSLLALLISIIIFLGFKSLRCTRIRIHVHLFASLACTCVAWILWYRLVVERHE    268
ficusphelia       LYVKGYALSLLALLISIIIFLGFKSLRCTRIRIHVHLFASLACTCVAWILWYRLVVERPE    260
erecta            LYVKGYALSLLALLISIIIFLGFKSLRCTRIRIHVHLFASLACTCVAWILWYRLVVERSE    267
melanogaster      LYVKGYALSLLALLISIIIFLGFKSLRCTRIRIHVHLFASLACTCVAWILWYRLVVERSE    256
simulans          LYVKGYALSLLALLISIIIFLGFKSLRCTRIRIHVHLFASLACTCVAWILWYRLVVERSE    256
sechellia         LYVKGYALSLLALLISIIIFLGFKSLRCTRIRIHVHLFASLACTCVAWILWYRLVVERSE    256
elegans           LYVKGYALSLLALLISIIIFLGFKSLRCTRIRIHVHLFASLACTCVAWILWYRLVVERNE    267
rhopaloa          LYVKGYALSLLALLISIIIFLGFKSLRCTRIRIHVHLFASLACTCVAWILWYRLVVERNE    257
takahashi         LYVKGYTLSLLALLISIIIFLGFKSLRCTRIRIHVHLFASLACTCVAWILWYRLVVERGE    275
Suzuki            LYVRGYALSLLALLISIIIFLGFKSLRCTRIRIHVHLFASLACTCVAWILWYRLVVERPE    268
biarmipes         LYVRGYALSLLALLISIIIFLGFKSLRCTRIRIHVHLFASLACTCVAWILWYRLVVERPE    268
grimshawi         LYVKGYALSLLALLISIIIFVGFKSLRCNRIRIHVHLFASLACTCLTWILWYRLVVEHSE    260
virilism          LYVKGYALSLLALFISIVIFLGFKSLRCTRIRIHVHLFASLACTCIAWILWYRLVVEHTE    264
mojavensis        LYVKGYALSLLALIISIVIFLGFKSLRCTRIRIHVHLFGSLACTCIAWILWYRLVVEQTD    269
                  ***:**::*****::**:**:*******.*********.*** **::**********: :

bipectinata       ITAENPLWCIVLHLVVHYFMLVNYFWMFCEGLHLHLVLVVVFVKDTIVMRWFIIISWLSP    331
anannasae         ITAENPLWCILLHLVVHYFMLVNYFWMFCEGLHLHLVLVVVFVKDTIVMRWFIIISWLSP    328
kikkawei          IIADNPLWCIGLHLVVHYFMLVNYFWMFCEGLHLHLVLVVVFVKDTIVMRWFIIISWFSP    351
seratta           IIADNPLWCIGLHLVVHYFMLVNYFWMFCEGLHLHLVLVVVFVKDTIVMRWFIIISWFSP    328
eugracilis        TITENPLWCIGLHLVVHYFMLVNYFWMFCEGLHLHLVLVVVFVKDTIVMRWFIVISWFSP    328
ficusphelia       TIAENQLWCIGLHLVVHYFMLVNYFWMFCEGLHLHLVLVVVFVKDTIVMRWFIVISWFSP    320
erecta            TIAENPLWCIGLHLVVHYFMLVNYFWMFCEGLHLHLVLVVVFVKDAIVMRWFILISWFLP    327
melanogaster      TIAENPLWCIGLHLVVHYFMLVNYFWMFCEGLHLHLVLVVVFVKDTIVMRWFIVISWFSP    316
simulans          TIAENPLWCIGLHLVVHYFMLVNYFWMFCEGLHLHLVLVVVFVKDTIVMRWFIVISWFSP    316
sechellia         TIAENPLWCIGLHLVVHYFMLVNYFWMFCEGLHLHLVLVVVFVKDTIVMRWFIVISWFSP    316
elegans           TIAENPLWCIGLHLVVHYFMLVNYFWMFCEGLHLHLVLVVVFVKDTIVMRWFIVISWLSP    327
rhopaloa          TIAENPLWCSGLHLVVHYFMLVNYFWMFCEGLHLHLVLVVVFVKDTIVMRWFIVISWLSP    317
takahashi         AIAENPLWCIGLHLVVHYFMLVNYFWMFCEGLHLHLVLVVVFVKDTIVMRWFIVISWFSP    335
Suzuki            TIADNPLWCIGLHLVVHYFMLVNYFWMFCEGLHLHLVLVVVFVKDTIVMRWFMVISWLSP    328
biarmipes         TIADNPLWCIGLHLVVHYFMLVNYFWMFCEGLHLHLVLVVVFVKDTIVMRWFMVISWLSP    328
grimshawi         RIAENPNWCIALHLVVHYFMLVNYFWMFCEGLHLHLVLVVVFVKDTIVLRWFKFFSWLSP    320
virilism          QIAENPPWCIALHLVVHYFMLVNYFWMFCEGLHLHLVLVVVFVKDTIVMRWFKLLSWLLP    324
mojavensis        QIAENPPWCIALHLVVHYFMLVNYFWMFCEGLHLHLVLVVVFVKDTIVMRWFKLLSWLSP    329
                    ::*  **  **********************************:**:*** .:**: *

bipectinata       VHIAVVYGLARHFSSPDNEHCWINDSLYLWIFSVPITLSLLASFIFLINVLRVIVRKLHP    391
anannasae         VHFAVVYGLARHFSSPDNEHCWINDSLYLWIFSVPITLSLLASFIFLINVLRVIVRKLHP    388
kikkawei          IHFAIVYGLARHFSDSDNEHCWINDSLYLWIFSVPITLSLVASFIFLINVLRVIVRKLHP    411
seratta           IHFAIVYGLSRHFSDSDNEHCWINDSLYLWIFSVPITLSLLASFIFLINVLRVIVRKLHP    388
eugracilis        IPIAAVYGLARHFSNPDNKHCWINDSLYLWMFSVPITLSLLASFIFLINVLRVIVRKLHP    388
ficusphelia       IPIAIVYGLARHFSSPDNKHCWINDSLYLWIFSVPITLSLLASFIFLINVLRVIVRKLHP    380
erecta            IHITILYGLARHFSTPDNKHCWITDSLYLWIFSVPITLSLLASFIFLINVLRVIVRKLHP    387
melanogaster      IPIAIVYGLARHFSSPDNKHCWITDSLYLWIFSVPITLSLLASFIFLINVLRVIVRKLHP    376
simulans          IPIAIVYGLARHFSSPDNKHCWITDSLYLWIFSVPITLSLLASFIFLINVLRVIVRKLHP    376
sechellia         IPIAIVYGLARHFSSPDNKHCWITDSLYLWIFSVPITLSLLASFIFLINVLRVIVRKLHP    376
elegans           IPVAIVYGLARHFSGPDNKHCWINDSLYLWIFSVPITLSLLASFIFLINVLRVIVRKLHP    387
```

```
rhopaloa        IPIAVVYGLARHFTSPDNKHCWINDSLYLWIFSVPITLSLLASFIFLINVLRVIVRKLHP    377
takahashi       IPVAIVYGLARHFSSPDNKHCWINDSLYLWIFSVPITLSLLASFIFLINVLRVIVRKLHP    395
Suzuki          IPIAVVYGLARHFSSPDNKHCWINDSLYLWIFSVPITLSLLASFIFLINVLRVIVRKLHP    388
biarmipes       IPIAIVYGLARHFSSPDNKHCWINDSLYLWIFSVPITLSLLASFIFLINVLRVIVRKLHP    388
grimshawi       LLFIIPYGVVRHFSANDNKHCWISESFYLWILSVPITLSLLASFIFLINVLRVIVRKLHP    380
virilism        LLFVLPYGVARHFSANDNAHCWMNDSFYLWIFSVPITLSLLASFIFLINVLRVIVRKLHP    384
mojavensis      LVFVLPYGVARHLSVNDNEHCWINDSLYLWIFSVPITLSLLASFIFLINVLRVIVRKLHP    389

bipectinata     QSAQPAPLAIRKAVRATIILVPLFGLQHFLLPYRPDAGTQLDRFYQLLSVVLVSLQGFVV    451
anannasae       QSAQPAPLAIRKAVRATIILVPLFGLQHFLLPYRPDAGTQLDRFYQLLSVVLVSLQGFVV    448
kikkawei        QSAQPAPLAIRKAVRATIILVPLFGLQHFLLPYRPDAGTQLDRFYQLLSVVLVSLQGFVV    471
seratta         QSAQPAPLAIRKAVRATIILVPLFGLQHFLLPYRPDAGTQLDRFYQLLSVVLVSLQGFVV    448
eugracilis      QSAQPAPLAIRKAVRATIILVPLFGLQHFLLPYRPDAGSQLDRFYQMLSVALVSPQGFVV    448
ficusphelia     QSAQPAPLAIRKAVRATIILVPLFGLQHFLLPYRPDAGTQLDRFYQLLSVVLVSLQGFVV    440
erecta          QSAQPAPLAIRKAVRATIILVPLFGLQHFLLPYRPDAGTQLDHFYQMLSVVLVSLQGFVV    447
melanogaster    QSAQPAPLAIRKAVRATIILVPLFGLQHFLLPYRPDAGTQLDHFYQMLSVVLVSLQGFVV    436
simulans        QSAQPAPLAIRKAVRATIILVPLFGLQHFLLPYRPDAGTQLDHFYQMLSVVLVSLQGFVV    436
sechellia       QSAQPAPLAIRKAVRATIILVPLFGLQHFLLPYRPDAGTQLDHFYQMLSVVLVSLQGFVV    436
elegans         QSAQPAPLAIRKAVRATIILVPLFGLQHFLLPYRPDAGTQLDRFYQLLSVVLVSLQGFVV    447
rhopaloa        QSAQPAPLAIRKAVRATIILVPLFGLQHFLLPYRPDAGTRLDRFYQLLSVVLVSLQGLVV    437
takahashi       QSAQPAPLAIRKAVRATIILVPLFGLQHFLLPYRPDAGTQLDHFYQMLSVVLVSLQGFVV    455
Suzuki          QSAQPAPLAIRKAVRATIILVPLFGLQHFLLPYRPDAGTQLDHFYQMLSVILVSLQGFVV    448
biarmipes       QSAQPAPLAIRKAVRATIILVPLFGLQHFLLPYRPDAGTQLDHFYQMLSVILVSLQGFVV    448
grimshawi       QSAHPAPMAIRKAVRATIILVPLFGLQHFLLPYRPEAGTKLDRFYQLMSVVLVSLQGFVV    440
virilism        QSAQPAPLAIRKAVRATIILVPLFGLQHFLLPYRPDAGTQLDRFYQLLSVVLVSLQGFVV    444
mojavensis      QSAQPAPLAIRKAVRATIILVPLFGLQHFLLPYRPEAGTQLDRFYQLLSVVLVSLQGFVV    449

bipectinata     SFLFCFANHDVLFAVRTLLNKLLPSLVSPPPAGSNTGQMATTTPSRELGV    501
anannasae       SFLFCFANHDVLFAVRTLLNKLMPSLVSPPPAGSNTGQMATTTPSRELGV    498
kikkawei        SFLFCFANHDVTFAIRTLLNKWLPSLVAAPPAGSNTGQMATTTPSRELGV    521
seratta         SFLFCFANHDVTFAIRTLLNKWLPSVVAPPPAGSNTGQMATTTPSRELGV    498
eugracilis      SFLFCFANHDVTFAIRTLLNKWLPSLVTPPPAGSNTGQMATTTPSRELGV    498
ficusphelia     SFLFCFVNHDVTFAIRTLLNKMMPTLVTAPPAGSNTGQMATTTPSRELGV    490
erecta          SFLFCFANHDVTFAVRTLLNKWLPSLVIAPPAGSNTGQMATTTPSRELGV    497
melanogaster    SFLFCFANHDVTFAIRTLLNKLLPSLVTPPPAGSNTGQMATTTPSRELGV    486
simulans        SFLFCFANHDVTFAIRTLLNKLLPSLVTPPPAGSNTGQMATTTPSRELGV    486
sechellia       SFLFCFANHDVTFAIRTLLNKLLPSLVTPPPAGSNTGQMATTTPSRELGV    486
elegans         SFLFCFANHDVTFAIRTLLNKWLPRLVTAPPAGSNTGQMATTTPSRELGV    497
rhopaloa        SFLFCFANHDVTFAIRTLLNKWLPRLVTAPPAGSNTGQMATTTPSRELGV    487
takahashi       SFLFCFANHDVTFAIRTLLNKWLPSLVTPPPAGSNTGQMATTTPSRELGV    505
Suzuki          SFLFCFANHDVTFAIRTLLNKLLPSLVTPPPAGSNTGQMATTTPSRELGV    498
biarmipes       SFLFCFANHDVTFAIRTLLNKLLPSLVTPPPAGSNTGQMATTTPSRELGV    498
grimshawi       SFVFCFVNQDVIVAIRTLLNKWMPSLVSAPPAGSNTGQMATTTPSRELGV    490
virilism        SFLFCFANHDVTFAMRTLLNKLMPTLVAPPPAGSNTGQLATTTPSRELGV    494
mojavensis      SFLFCFANHDVTFAIRTMLNKWMPNLIAPPPAGSNTGQLATTTPSRELGV    499
```

melanogaster

```
1 mattssdses qnvdvasqaq tqdnlriflk hlyaecvfry qnvtydtddp sfslgpatdy

       61 dsdlpenfsp vprylenaam negvidmrnv deelaekeel matvvsatma tnqkenrlfc

      121 plnfdgylcw prtpagtvls qycpdfvegf nrkflahktc lengswyrhp vsnqtwsnyt

      181 ncvdyedlef rqfinelyvk gyalsllall isiiiflgfk slrctririh vhlfaslact

      241 cvawilwyrl vversetiae nplwciglhl vvhyfmlvny fwmfceglhl hlvlvvvfvk

      301 dtivmrwfiv iswfspipia ivyglarhfs spdnkhcwit dslylwifsv pitlsllasf

      361 iflinvlrvi vrklhpqsaq paplairkav ratiilvplf glqhfllpyr pdagtqldhf

      421 yqmlsvvlvs lqgfvvsflf cfanhdvtfa irtllnkllp slvtpppags ntgqmatttp

      481 srelgv
```

simulans XP_016039137.1

```
  1 mattssdsdv envdvasqaq tqdnlriflk hlyaecvfry qnvtydtddp sfslgpatdy
       61 dsdlpenfsp vprylenaam negvidmrnv deelaekeel matvvsatma tnqkesrlfc
      121 plnfdgylcw prtpagtvls qycpdfvegf nrkflahktc lengswyrhp vsnqtwsnyt
      181 ncvdyedlef rqfinelyvk gyalsllall isiiiflgfk slrctririh vhlfaslact
      241 cvawilwyrl vversetiae nplwciglhl vvhyfmlvny fwmfceglhl hlvlvvvfvk
```

```
301 dtivmrwfiv iswfspipia ivyglarhfs spdnkhcwit dslylwifsv pitlsllasf
361 iflinvlrvi vrklhpqsaq paplairkav ratiilvplf glqhfllpyr pdagtqldhf
421 yqmlsvvlvs lqgfvvsflf cfanhdvtfa irtllnkllp slvtpppags ntgqmatttp
481 srelgv
```

sechellia **XP_002042869.1**

```
1 mattssdsdv envdvasqaq tqdnlriflr hlyaecvfry qnvtydtddp sfslgpatdy
61 dsdlpenfsp vprylenaam negvidmsnv deklaekeel matvvsatma tnqkesrlfc
121 plnfdgylcw prtpagtvls qycpdfvegf nrkflahktc lengswyrhp vsnqtwsnyt
181 ncvdyedlef rqfinelyvk gyalsllall isiiiflgfk slrctririh vhlfaslact
241 cvawilwyrl vversetiae nplwciglhl vvhyfmlvny fwmfceglhl hlvlvvvfvk
301 dtivmrwfiv iswfspipia ivyglarhfs spdnkhcwit dslylwifsv pitlsllasf
361 iflinvlrvi vrklhpqsaq paplairkav ratiilvplf glqhfllpyr pdagtqldhf
421 yqmlsvvlvs lqgfvvsflf cfanhdvtfa irtllnkllp slvtpppags ntgqmatttp
481 srelgv
```

erecta **XP_001978328.2**

```
1 mtavthteqp tmattssdse plnldvasqa qtqdnlrifl khlyaecvfr yqnvtydtgd
61 psfslgpatd ydsdlpenfs pvprylenaa mnegvidmrs vdeelaekee lmatvvsatm
121 atnqkehrlf cplnfdgylc wprtpagtvl sqycpdfveg fnrkflahkt clengswyrh
181 pvsnqtwsny tncvdykdle frqfinelyv kgyalsllal lisiiiflgf kslrctriri
241 hvhlfaslac tcvawilwyr lvversetia enplwciglh lvvhyfmlvn yfwmfceglh
301 lhlvlvvvfv kdaivmrwfi liswflpihi tilyglarhf stpdnkhcwi tdslylwifs
361 vpitlsllas fiflinvlrv ivrklhpqsa qpaplairka vratiilvpl fglqhfllpy
421 rpdagtqldh fyqmlsvvlv slqgfvvsfl fcfanhdvtf avrtllnkwl pslviappag
481 sntgqmattt psrelgv
```

Suzuki **XP_016923892.1**

```
1 mtrvspteap tmgtaasvse senvevasqt qtqdnlrifl khlyaecvyr yqnvtydtdd
61 psvslgaatd ydfdlpenfs pvprylenaa mnegaidmrn vdaelaekee lvatvvsatm
121 atnqreeprl fcplnfdgyl cwprtpagtv lsqycpdfve gfnrkflahk tclengswfr
181 hpesnqtwsn ytncvdyedl efrqfvnely vrgyalslla llisiiiflg fkslrctrir
241 ihvhlfasla ctcvawilwy rlvverpeti adnplwcigl hlvvhyfmlv nyfwmfcegl
301 hlhlvlvvvf vkdtivmrwf mviswlspip iavvyglarh fsspdnkhcw indslylwif
361 svpitlslla sfiflinvlr vivrklhpqs aqpaplairk avratiilvp lfglqhfllp
421 yrpdagtqld hfyqmlsvil vslqgfvvsf lfcfanhdvt fairtllnkl lpslvtpppa
481 gsntgqmatt tpsrelgv
```

biarmipes **XP_016956301.1**

```
1 mtraspieap tmgtaasese senveaasqt qtqdnlrifl khlyaecvyr yqnvtydtdd
61 psvslgaatd ydfdlpenfs pvprylenaa lnegaidmrn vdaeqaekee lvatvvsatm
121 atnqraeprl fcplnfdgyl cwprtpagtv lsqycpdfve gfnrkflahk tclengswfr
181 hpesnqtwsn ytncvdyedl efrqfvnely vrgyalslla llisiiiflg fkslrctrir
241 ihvhlfasla ctcvawilwy rlvverpeti adnplwcigl hlvvhyfmlv nyfwmfcegl
301 hlhlvlvvvf vkdtivmrwf mviswlspip iaivyglarh fsspdnkhcw indslylwif
361 svpitlslla sfiflinvlr vivrklhpqs aqpaplairk avratiilvp lfglqhfllp
421 yrpdagtqld hfyqmlsvil vslqgfvvsf lfcfanhdvt fairtllnkl lpslvtpppa
481 gsntgqmatt tpsrelgv
```

takahashi **XP_016995732.2**

```
1 mtrvshtepp tmgaaasdsq senveaasqs qmqdnlrifl khlyaecvyr yqnvtydted
61 psvplgaatd ydfdlpenfs pvprylenaa lnegaidmrn vdavlaekee lmatvvsatm
121 atnqrqaeeg geeerrlfcp lnfdgylcwp rtpagtvlsq ycpdfvegfn skflahktcl
181 engswfrhpv snqtwsnytn cvdyedlefr qfinelyvkg ytlsllalli siiiflgfks
241 lrctririhv hlfaslactc vawilwyrlv vergeaiaen plwciglhlv vhyfmlvnyf
301 wmfceglhlh lvlvvvfvkd tivmrwfivi swfspipvai vyglarhfss pdnkhcwind
361 slylwifsvp itlsllasfi flinvlrviv rklhpqsaqp aplairkavr atiilvplfg
421 lqhfllpyrp dagtqldhfy qmlsvvlvsl qgfvvsflfc fanhdvtfai rtllnkwlps
481 lvtpppagsn tgqmatttps relgv
```

elegans **XP_017133493.1**

```
1 mttvthfdtp tmgtvasdst menvetatqs qtqdnlrifl khlyaecvyr yknvtydaad
61 psvslvaatd yesdltenfs pvpryledav lnegaidmrn vdeelaekee lmatvlsatm
121 atnqqepllf cplnfdgylc wprtpagtvl sqycpdfveg fnskflahkt clengswfrh
```

181 pvsnqtwsny tncvdyedle frqfinelyv kgyalsllal lisiiiflgf kslrctriri
241 hvhlfaslac tcvawilwyr lvvernetia enplwciglh lvvhyfmlvn yfwmfceglh
301 lhlvlvvvfv kdtivmrwfi viswlspipv aivyglarhf sgpdnkhcwi ndslylwifs
361 vpitlsllas fiflinvlrv ivrklhpqsa qpaplairka vratiilvpl fglqhfllpy
421 rpdagtqldr fyqllsvvlv slqgfvvsfl fcfanhdvtf airtllnkwl prlvtappag
481 sntgqmattt psrelgv

rhopaloa **XP_016985474.1**
1 mgtvasdsvp envetatqsq tqdnlriflk hlyaecvyry qndtydaddp svslgaatdy
61 dldlpesfsp vpryledavl negaidmrnv deelaekeel matvlsatma tnqrpqqrlf
121 cplnfdgylc wprtpagtvl sqycpefveg fnskflahkt clengswfrh pvsnqtwsny
181 tncvdyedle frqfinelyv kgyalsllal lisiiiflgf kslrctriri hvhlfaslac
241 tcvawilwyr lvvernetia enplwcsglh lvvhyfmlvn yfwmfceglh lhlvlvvvfv
301 kdtivmrwfi viswlspipi avvyglarhf tspdnkhcwi ndslylwifs vpitlsllas
361 fiflinvlrv ivrklhpqsa qpaplairka vratiilvpl fglqhfllpy rpdagtrldr
421 fyqllsvvlv slqglvvsfl fcfanhdvtf airtllnkwl prlvtappag sntgqmattt
481 psrelgv

ficusphelia **XP_017052090.1**
1 mgtgasssds gsekletaaq aqtqdnlrif lkhlyaecvy ryqnvtyead dpsvplgaat
61 eydvdanapd nfspvpryle davlnegaid mrhvdeeaae keelmatvft atmatnqree
121 rlfcplnfdg ylcwprtpag tvlsqycpdf vegfnskfla hktclengsw frhpvsnqtw
181 snytscvdye dlefrqfine lyvkgyalsl lallisiiif lgfkslrctr irihvhlfas
241 lactcvawil wyrlvverpe tiaenqlwci glhlvvhyfm lvnyfwmfce glhlhlvlvv
301 vfvkdtivmr wfiviswfsp ipiaivygla rhfsspdnkh cwindslylw ifsvpitlsl
361 lasfiflinv lrvivrklhp qsaqpaplai rkavratiil vplfglqhfl lpyrpdagtq
421 ldrfyqllsv vlvslqgfvv sflfcfvnhd vtfairtlln kmmptlvtap pagsntgqma
481 tttpsrelgv

eugracilis **XP_017065034.1**
1 mtrvthietp tmvgttases esenvemasq aqtqdnllvf lthlyaecvh ryqnvtndte
61 dsslslgaat dsendlkgnf stvprylena alnegtidmr nvdeklaeqe elmatvssat
121 matnqrdrrl fcpldfdgyl cwprtpagtv lsqycpdfve gfnkkflahk tclengswfr
181 hpatnqtwsn ytncvdhedl efrkfinely vkgyalslla llisiiiflg fkslrctrir
241 ihvhlfasla ctcvawilwy rlvverheti tenplwcigl hlvvhyfmlv nyfwmfcegl
301 hlhlvlvvvf vkdtivmrwf iviswfspip iaavyglarh fsnpdnkhcw indslylwmf
361 svpitlslla sfiflinvlr vivrklhpqs aqpaplairk avratiilvp lfglqhfllp
421 yrpdagsqld rfyqmlsval vspqgfvvsf lfcfanhdvt fairtllnkw lpslvtpppa
481 gsntgqmatt tpsrelgv

kikkawei **XP_017029945.1**
1 mtsieeappt mgtnasdsds dsgsenvama tatasqqsyt qpqaqdnlri flrhlyaecv
61 yryqndtydd psvsvvgeaa atdydldlpe nfspvpryle navlnegaid mrnvdveqae
121 keellatvls gtmatnqqkq shqattttg erlfcplnfd gylcwprtpa gtvlsqycpd
181 fvegfnskfl ahktclengs wfrhpmtnrt wsnytncvdy edlefrqfin elyvkgyals
241 llallvsiii flgfkslrct ririhvhlfa slaftcvawi lwyrlvverp eiiadnplwc
301 iglhlvvhyf mlvnyfwmfc eglhlhlvlv vvfvkdtivm rwfiiiswfs pihfaivygl
361 arhfsdsdne hcwindslyl wifsvpitls lvasfiflin vlrvivrklh pqsaqpapla
421 irkavratii lvplfglqhf llpyrpdagt qldrfyqlls vvlvslqgfv vsflfcfanh
481 dvtfairtll nkwlpslvaa ppagsntgqm atttpsrelg v

seratta **KAH8380938.1**
1 mamakasgsq qsssqtqaqd nlriflrhly aecvyryqnd tyddpsvsvd gaaatdyese
61 lpenfspvpr ylenavlneg aidmrnvdve laekeellat vlsvtmatnq qesehhevtt
121 tnnrsagerl fcplnfdgyl cwprtpagtv lsqycpdfve gfnskflahk tclengswfr
181 hpmsnqtwsn ytncvdyedl efrqfinely vkgyalslla llvsiliflg fkslrctrir
241 ihvhlfasla ftcvawilwy rlvverpeii adnplwcigl hlvvhyfmlv nyfwmfcegl
301 hlhlvlvvvf vkdtivmrwf iiiswfspih faivyglsrh fsdsdnehcw indslylwif
361 svpitlslla sfiflinvlr vivrklhpqs aqpaplairk avratiilvp lfglqhfllp
421 yrpdagtqld rfyqllsvvl vslqgfvvsf lfcfanhdvt fairtllnkw lpsvvapppa
481 gsntgqmatt tpsrelgv

bipectinata **XP_017095188.2**
1 mgglsasvte pdmessraqd apqpqdnlri flkhlyaecv yryqndtydd pstesatdye
61 sdfpenfspv prylenaaln egsidmrnvd ekkaereelf atiltatmat nqnqddpagq
121 rspgsnssss rrlfcpldfd gylcwprtpa gtvlsqycpd fvegfnskfl ahktcqengs
181 wyrhpetnkt wsnytncvdy vdfefrqfin elyvkgyals llallisiii flgfkslrct
241 ririhvhlfa slactcvawi lwyrlvveqp eitaenplwc ivlhlvvhyf mlvnyfwmfc
301 eglhlhlvlv vvfvkdtivm rwfiiiswls pvhiavvygl arhfsspdne hcwindslyl

361 wifsvpitls llasfiflin vlrvivrklh pqsaqpapla irkavratii lvplfglqhf
421 llpyrpdagt qldrfyqlls vvlvslqgfv vsflfcfanh dvlfavrtll nkllpslvsp
481 ppagsntgqm atttpsrelg v

ananassae **XP_001964132.2**

1 mgglsasvte pdmeqeapqp qdnlriflkh lyaecvyryq ndtyddpmte tatdyesdlp
61 enfspvpryl enaalnegsi dmrnvdekqa ekeelfatil tatmatnqnq depagqrssg
121 nssssgskrl fcpldfdgyl cwprtpagtv lsqycpdfve gfnskflahk tcqengswyr
181 hpesnktwsn ytncvdyvdl efrqfinely vkgyaislla llisiiiflg fkslrctrir
241 ihvhlfasla ctcvawilwy rlvverpeit aenplwcill hlvvhyfmlv nyfwmfcegl
301 hlhlvlvvvf vkdtivmrwf iiiswlspvh favvyglarh fsspdnehcw indslylwif
361 svpitlslla sfiflinvlr vivrklhpqs aqpaplairk avratiilvp lfglqhfllp
421 yrpdagtqld rfyqllsvvl vslqgfvvsf lfcfanhdvl favrtllnkl mpslvspppa
481 gsntgqmatt tpsrelgv

virilism **XP_032296576.1**

1 mallraaltm ataayfemtv sgamppqpqd nlrtflkhly aecvyryqnd tataqlatep
61 ddggllllety tmipryleqa vlnegtidmq dvdeeaasen elyatvlsat matnhhshnt
121 semetlycpv nfdgylcwpr tpagtvlsqy cpdfvegfns kflahktcle tgswfrhpvs
181 nqtwsnytnc vdyedfqfrq fvnelyvkgy alsllalfis iviflgfksl rctririhvh
241 lfaslactci awilwyrlvv ehteqiaenp pwcialhlvv hyfmlvnyfw mfceglhlhl
301 vlvvvfvkdt ivmrwfklls wllpllfvlp ygvarhfsan dnahcwmnds fylwifsvpi
361 tlsllasfif linvlrvivr klhpqsaqpa plairkavra tiilvplfgl qhfllpyrpd
421 agtqldrfyq llsvvlvslq gfvvsflfcf anhdvtfamr tllnklmptl vapppagsnt
481 gqlatttpsr elgv

mojavensis **XP_015016349.1**

1 mallraalam ttaayietam tasltkpqpr pqpqdnlrif lrhlyvecvy ryqndtakgd
61 akdsgdeelm latfttvpry leqavlnegt idmqdvdeea asenelyati hsmtmetnqh
121 srnatsemet lycpvnfdgy lcwprtpagt vlsqycpdfv egfsskflah ktclengtwy
181 rhpvsnqtws nytncvdydd fqfrqfvnel yvkgyalsll aliisivifl gfkslrctri
241 rihvhlfgsl actciawilw yrlvveqtdq iaenppwcia lhlvvhyfml vnyfwmfceg
301 lhlhlvlvvv fvkdtivmrw fkllswlspl vfvlpygvar hlsvndnehc windslylwi
361 fsvpitlsll asfiflinvl rvivrklhpq saqpaplair kavratiilv plfglqhfll
421 pyrpeagtql drfyqllsvv lvslqgfvvs flfcfanhdv tfairtmlnk wmpnliappp
481 agsntgqlat ttpsrelgv

grimshawi **XP_001992054.1**

1 mtasasasms pqpqdnlltf lwhlyaecvy ryqndtspgq qatdpgdgni qllepytrmp
61 nylemavlne gtidmqdvne eassInelya tvlsatmapn qhsiqyngnn snanttreme
121 ilycpvnfdg ylcwprtpag tvlsqycpdf vegfntkfla hktcmetgsw frhpvsnqtw
181 snytncvdye dlqfrqivne lyvkgyalsl lallisiiif vgfkslrcnr irihvhlfas
241 lactcltwil wyrlvvehse riaenpnwci alhlvvhyfm lvnyfwmfce glhlhlvlvv
301 vfvkdtivlr wfkffswlsp llfiipygvv rhfsandnkh cwisesfylw ilsvpitlsl
361 lasfiflinv lrvivrklhp qsahpapmai rkavratiil vplfglqhfl lpyrpeagtk
421 ldrfyqlmsv vlvslqgfvv sfvfcfvnqd vivairtlln kwmpslvsap pagsntgqma
481 tttpsrelgv
