## Supplementary material for "The incidence of candidate binding sites for β-arrestin in Drosophila neuropeptide GPCRs": S17 Text

***S17 Text.*** **Multi-species analysis of CG33639**
**Supporting Figure 20**

CLUSTAL Line-ups; Genbank Reference IDs below
Predicted TM domains in YELLOW
BBS sequences in RED

```
Grimshawi         ------------MITRLYNTEEDPVYCSFVWGANVSNGRS-----------HAANSSALY        37
Mojavensis        MERIVRNHQSKTMITRLYNTEEDPAYCSVIWGANLTSSSD-------LVLGTATNATHIY        53
Virilism          ------------MITRLYNTEEDPAYCSFIWGANLTSSND-------LVLGTTANSTHIY        41
Bipectinate       ------------MITRLYHTEEDPAYCSFIGGSNLSS--------SSDVLATSANATPVF        40
Anannassae        ------------MITRLYHTEEDPAYCSFIGGSNISISSSSSSISSSDVLATSANATPVF        48
Serrata           ------------MITRLYNTEEDPAYCSFIWGSNLSSTT--------DVLA-AAGNSSPV        39
Kikkawei          ------------MITRLYNTEEDPAYCSFIWGSNLSSTT--------DVLA-AAGNSSPV        39
Fichsuphila       ------------MITRLYNTEEDPAYCSFIWGGSNLTAT--------AAEVLAANATSIF        40
Eugracilis        ------------MITSLYTTQEDPAYCSFIWGSNL--TT--------SADVLTANATSVF        38
Rhopaloa          ------------MITRLYNTEEDPAYCSFIWGANL--TS--------SADVLA-NATSVF        37
Elegans           ------------MITRLYNTEEDPAYCSFIWGSNL--TS--------SADVLA-NATSAF        37
Erecta            ------------MITRLYNTEEDPAYCSFIWGSNL--TS--------SVDVLAANATSVF        38
Suzuki            ------------MITRLYNTEEDPAYCSFIWGSNL--TS--------SAEVLAANATSVF        38
Biarmipes         ------------MITRLYNTEEDPAYCSFIWGSNL--TS--------SAEVLAANATSIF        38
Takahashi         ------------MITRLYNTEEDPAYCSFIWGSNL--TS--------SADVLAANATSVF        38
Sechellia         ------------MITRLYNTEEDPAYCSFIWGSNL--TS--------SVDVLAANATSVF        38
Melanogaster      ------------MITRLYNTEEDPAYCSFIWGSNL--TS--------SVDVLAANATSVF        38
Simulans          ------------MITRLYNTEEDPAYCSFIWGSNL--TS--------SVDVLAANATSVF        38
Mauritania        ------------MITRLYNTEEDPAYCSFIWGSNL--TS--------SVDVLAANATSVF        38
                              *** ** *:***.***.: *..                  : . :

Grimshawi         PDDMRDDWFADAEDPRTELLRQYCYGYFLPFICASGIIGNVLNLIVLTRRNMRGPSYIYM        97
Mojavensis        ANDLRDDLYADVEDPRTEALREYCYGLMLPIICALGIIGNVLNLIVLTRRNMRGTAYIYM        113
Virilism          ANDLRDDLYADVEDPRTESLREYCYGLMLPVICALGIIGNVLNLIVLTRRNMRGTAYIYM        101
Bipectinate       GSDLRDDFYRDVEDPRTESLREYCYGLLLPIICAMGIIGNVLNLIVLTRRNMRGIAYIYM        100
Anannassae        GSDLRDDFYRDVEDPRTESLREYCYGLLLPIICAMGIIGNVLNLIVLTRRNMRGIAYIYM        108
Serrata           FTDLRDDFYRDVEDPRTESLREYCYGLLLPIICAMGIIGNVLNLIVLTRRNMRGTAYIYM        99
Kikkawei          FTDLRDDFYRDVEDPRTESLREYCYGLLLPITCAMGIIGNVLNLIVLTRRNMRGTAYIYM        99
Fichsuphila       GTDLRDDFYRDVEDPRTESLREYCYGLLLPIICAMGIIGNLLNLVVLTRRNMRGTAYIYM        100
Eugracilis        SSDLRDDFYRDVEDPRTESLREYCYGLVLPIICAMGIIGNVLNLIVLTRRNMRGTAYIYM        98
Rhopaloa          NTDLRDDFYRDVEDPRTESLREYCYGLVLPIICAMGIIGNVFNLIVLTRRNMRGTAYIYM        97
Elegans           SSDMRDDFYRDVEDPRTESLREYCYGLVLPIICAMGIIGNVLNLIVLTRRNMRGTAYIYM        97
Erecta            SSDLRDDFYRDVEDPRTESLREYCYGLVLPIICAMGIIGNVLNLIVLTRRNMRGTAYIYM        98
Suzuki            SSDLRDDFYRDVEDPRTESLREYCYGLVLPIICAMGIIGNVLNLIVLTRRNMRGTAYIYM        98
Biarmipes         SSDLRDDFYRDVEDPRTESLREYCYGLVLPIICAMGIIGNVLNLIVLTRRNMRGTAYIYM        98
Takahashi         SSDLRDDFYRDVEDPRTESLREYCYGLVLPIICAMGIIGNVLNLIVLTRRNMRGTAYIYM        98
Sechellia         SSDLRDDFYRDVEDPRTESLREYCYGLVLPIICAMGIIGNVLNLVVLTRRNMRGTAYIYM        98
Melanogaster      SSDLRDDFYRDVEDPRTESLREYCYGLVLPIICAMGIIGNVLNLVVLTRRNMRGTAYIYM        98
Simulans          GSDLRDDFYRDVEDPRTESLREYCYGLVLPIICAMGIIGNVLNLVVLTRRNMRGTAYIYM        98
Mauritania        SSDLRDDFYRDVEDPRTESLREYCYGLVLPIICAMGIIGNVLNLVVLTRRNMRGTAYIYM        98
                    *:*** : *.****** **:**** .**. ** *****::**:********* :****

Grimshawi         RAYSTAALLAIVFAIPFGIRMLVHKDRGQWEEIGPAFYTAHLELFLGNGCLGVGVMMLLV        157
Mojavensis        RAYSTAALLAIVFAIPFGIRMLVHKDRGQWEEFGPAFYTAHLELFLGNGCLGVGVMMLLV        173
Virilism          RAYSTAALLAIVFAIPFGIRMLVHKDRGQWEEFGPAFYTAHLELFLGNGCLGVGVMMLLV        161
Bipectinate       RAYSTAALLAIVFAIPFGIRMLVHKDRGQWEEFGPAFYTAHLELYLGNGCLGVGVMMLLV        160
Anannassae        RAYSTAALLAIVFAIPFGIRMLVHKDRGQWEEFGPAFYTAHLELYLGNGCLGVGVMMLLV        168
Serrata           RAYSTAALLAIVFAIPFGIRMLVHKDRGQWEEFGPAFYTAHLELYLGNGCLGVGVMMLLV        159
Kikkawei          RAYSTAALLAIVFAIPFGIRMLVHKDRGQWEEFGPAFYTAHLELYLGNGCLGVGVMMLLV        159
Fichsuphila       RAYSTAALLAIVFAIPFGIRMLVHKDRGQWEEFGPAFYTAHLELYLGNGCLGVGVMMLLV        160
Eugracilis        RAYSTAALLAIVFAIPFGIRMLVHKDRGQWEKFGPAFYTAHLELYLGNGCLGIGVMMLLV        158
Rhopaloa          RAYSTAALLAIVFAIPFGIRMLVHKDRGQWEEFGPAFYTAHLELYLGNGCLGVGVMMLLV        157
Elegans           RAYSTAALLAIVFAIPFGIRMLVHKDRGQWEEFGPAFYTAHLELYLGNGCLGVGVMMLLV        157
Erecta            RAYSTAALLAIVFAIPFGIRMLVHKDRGQWEEFGPAFYTAHLELYLGNGCLGVGVMMLLV        158
Suzuki            RAYSTAALLAIVFAIPFGIRMLVHKDRGQWEEFGPAFYTAHLELYLGNGCLGVGVMMLLV        158
Biarmipes         RAYSTAALLAIVFAIPFGIRMLVHKDRGQWEEFGPAFYTAHLELYLGNGCLGVGVMMLLV        158
Takahashi         RAYSTAALLAIVFAIPFGIRMLVHKDRGQWEEFGPAFYTAHLELYLGNGCLGVGVMMLLV        158
Sechellia         RAYSTAALLAIVFAIPFGIRMLVHKDRGQWEEFGPAFYTAHLELYLGNGCLGVGVMMLLV        158
Melanogaster      RAYSTAALLAIVFAIPFGIRMLVHKDRGQWEEFGPAFYTAHLELYLGNGCLGVGVMMLLV        158
Simulans          RAYSTAALLAIVFAIPFGIRMLVHKDRGQWEEFGPAFYTAHLELYLGNGCLGVGVMMLLV        158
Mauritania        RAYSTAALLAIVFAIPFGIRMLVHKDRGQWEEFGPAFYTAHLELYLGNGCLGVGVMMLLV        158
```

```
                  ******************************::***********:*******:*******

Grimshawi         LTIERYVSVCRPGATRPAIGQPGVVIFIICVLTFIFYLPSIFRGELIKCMLTSKNVYVYL        217
Mojavensis        LTIERYVSVCHPGFTRPVMGPPGVVVFVTCLVTFIIYLPSIFRGELIKCMLTSNNVYVYL        233
Virilism          LTIERYVSVCHPGFTRPVMGPPGVVVFLTCFATFIIYLPSIFRGELIKCMLTSNNVYVYL        221
Bipectinate       LTIERYVSVCHPGFSRPVMGPPGVVVFFTCLATVIVYLPSIFRGELIKCMLGSSDVYVYL        220
Anannassae        LTIERYVSVCHPGFSRPVMGPPGVVVFFTCLATVIVYLPSIFRGELIKCMLGSSDVYVYL        228
Serrata           LTIERYVSVCHPGFARPVMGPPGVVVFLTCLATVIVYLPSIFRGELIKCILGSSDVYVYL        219
Kikkawei          LTIERYVSVCHPGFARPVMGPPGVVVFLTCLATVIVYLPSIFRGELIKCILGSSDVYVYL        219
Fichsuphila       LTIERYVSVCHPGFVRPVMGPPGVVVFLTCLATVIVYLPSIFRGELIKCVLHASDVYVYL        220
Eugracilis        LTIERYVSVCHPGFARPVMGPPGVVVFLTCLATVIVYLPSIFRGELIKCMYGSRDVYVYL        218
Rhopaloa          LTIERYVSVCHPGFARPVMGPPGVVVFFTCLATVIVYLPSIFRGELIKCIFGSSDAFVYL        217
Elegans           LTIERYVSVCHPGFARPVMGPPGVVVFLTCLATVIVYLPSIFRGELIKCIFGSSDVFVYL        217
Erecta            LTIERYVSVCHPGFARPVMGPPGVVVFLTCLATVIVYLPSIFRGELIKCILGSSDVYVYL        218
Suzuki            LTIERYVSVCHPGFARPVMGPPGVVVFLTCLATVIVYLPSIFRGELIKCIFGSSDVYVYL        218
Biarmipes         LTIERYVSVCHPGFARPVMGPPGVVVFLTCLATVIVYLPSIFRGELIKCIFGSSDVYVYL        218
Takahashi         LTIERYVSVCHPGFARPVMGPPGVVVFLTCLATVIVYLPSIFRGELIKCIFGSSDVYVYL        218
Sechellia         LTIERYVSVCHPGFARPVMGPPGVVVFLTCLATVIVYLPSIFRGELIKCILGSSDVYVYL        218
Melanogaster      LTIERYVSVCHPGFARPVMGPPGVVVFLTCLATVIVYLPSIFRGELIKCILGSSDVYVYL        218
Simulans          LTIERYVSVCHPGFARPVMGPPGVVVFLTCLATVIVYLPSIFRGELIKCILGSSDVYVYL        218
Mauritania        LTIERYVSVCHPGFARPVMGPPGVVVFLTCLATVIVYLPSIFRGELIKCILGSSDVYVYL        218
                  **********:**  **.:* ****:*. *. *.*.*************:  : :.:***

Grimshawi         RRDNIIYQQTLFYSVYKVLVEIIFKIIPTFIIAGLNLRIMWVYRRTCARRRQMVLTR---        274
Mojavensis        RRDNNIYQRTIFYSVYKIMLEVIFKLVPTVVIAGLNLRIMLVYRRTCERRRQMVLSR---        290
Virilism          RRDNNIYQRTIFYSVYKIMLEVIFKLIPTVLIAGLNLRIMLVYRRTCERRRQMVLTR---        278
Bipectinate       RRDNTIYQQTLFYRVYKIMLEVIFKLIPTVLIGGLNMRIMMVYRRTCERRRQMVLSRPQA        280
Anannassae        RRDNTIYQQTLFYRVYKIMLEVIFKLIPTVLIGGLNMRIMMVYRRTCERRRQMVLSRPQ-        287
Serrata           RRDNTIYQQTIFYRIYKIMLEVIFKLVPTVLIGGLNMRIMMVYRRTCERRRQMVLSRNHG        279
Kikkawei          RRDNTIYQQTIFYRIYKIMLEVIFKLVPTVLIGGLNMRIMMVYRRTCERRRQMVLSRNHG        279
Fichsuphila       RRDNTIYQQTIFYRVYKIMLEVIFKLVPTLVIGFLNMRIMMVYRRTCERRRQMVLSRPH-        279
Eugracilis        RRDNTIYQQTIFYRVYKIMLEVIFKLIPTLVIGGLNMRIMMVYRRTCERRRQMVLSRPH-        277
Rhopaloa          RRDNTIYQQTIFYRVYKIMLEVIFKLVPTLVIGGLNMRIMMVYRRTCERRRQMVLSRPHA        277
Elegans           RRDNTIYQQTIFYRVYKIMLEVIFKLVPTLVIGGLNMRIMMVYRRTCERRRQMVLSRPHG        277
Erecta            RRDNTIYQQTIFYRVYKIMLEVIFKLVPTLVIGGLNMRIMMVYRRTCERRRQMVLTRSHA        278
Suzuki            RRDNTIYQQTIFYRVYKIMLEVIFKLVPTLVIGGLNMRIMMVYRRTCERRRQMVLSRPHA        278
Biarmipes         RRDNTIYQQTIFYRVYKIMLEVIFKLVPTLVIGGLNMRIMMVYRRTCERRRQMVLSRPHA        278
Takahashi         RRDNTIYQQTIFYRVYKIMLEVIFKLVPTLVIGGLNLRIMMVYRRTCERRRQMVLSRPHA        278
Sechellia         RRDNTIYQQTIFYRVYKIMLEVIFKLVPTLVIGGLNMRIMMVYRRTCERRRKMVLSRSHA        278
Melanogaster      RRDNTIYQQTIFYRVYKIMLEVIFKLVPTLVIGGLNMRIMMVYRRTCERRRKMVLSRPHA        278
Simulans          RRDNTIYQQTIFYRVYKIMLEVIFKLVPTLVIGGLNMRIMMVYRRTCERRRKMVLSRPHA        278
Mauritania        RRDNTIYQQTIFYRVYKIMLEVIFKLVPTLVIGGLNMRIMMVYRRTCERRRKMVLSRPHA        278
                  **** ***:*:** :**:::*:***::**.:*. **:*** ****** ***:***:*

Grimshawi         ------------------------AIYAKNEDPRKFAEERRLFLLLGSTSILFLLCISPM        310
Mojavensis        ------------------------ATYVKDDDPRKFAEERRLFLLLGSTSILFLLCVSPM        326
Virilism          ------------------------ANYVKDDDPRKFAEERRLFLLLGSTSILFLLCVSPM        314
Bipectinate       HHHHHHN--------------ANGPGYVKDDDPRKFAEERRLFLLLGSTSILFLVCVSPM        326
Anannassae        ---AHHH--------------QNGPGYVKDDDPRKFAEERRLFLLLGSTSILFLVCVSPM        330
Serrata           HGHGHSHGQGHAQGHA------HTNGYVKDDDPRKFAEERRLFLLLGSTSILFLVCVSPM        333
Kikkawei          HGHGHSHAQGQAQGHA------HTNGYVKDDDPRKFAEERRLFLLLGSTSILFLVCVSPM        333
Fichsuphila       -----------------------AHGYLKDDDPRKFAEERRLFLLLGSTSILFLVCVSPM        316
Eugracilis        -----------------------AHGYMKDDDPRKFAEERRLFLLLGSTSILFLVCVSPM        314
Rhopaloa          HAHGHGHAHGQGHG--------HSHGYLKDDDPRKFAEERRLFLLLGSTSILFLVCVSPM        329
Elegans           HGHGHGHGHGHGHGAQGHGHAHGGHGYLKDDDPRKFAEERRLFLLLGSTSILFLVCVSPM        337
Erecta            YGHGH---------------G-HGHGYLKDDDPRKFAEERRLFLLLGSTSILFLVCVSPM        322
Suzuki            ------------------------HGYMKDDDPRKFAEERRLFLLLGSTSILFLVCVSPM        314
Biarmipes         ------------------------HGYMKDDDPRKFAEERRLFLLLGSTSILFLVCVSPM        314
Takahashi         HGHGHGHGHGHGHGH---GHG-HGHGYMKDDDPRKFAEERRLFLLLGSTSILFLVCVSPM        334
Sechellia         QGHGHGH----GHGH---GHA-HGHGYLKDDDPRKFAEERRLFLLLGSTSILFLVCVSPM        330
Melanogaster      QGHGHGHGHGHGHGH---GHA-HGHGYLKDDDPRKFAEERRLFLLLGSTSILFLVCVSPM        334
Simulans          QGHGHGHGHGHGHGH---GHA-HGHGYLKDDDPRKFAEERRLFLLLGSTSILFLVCVSPM        334
Mauritania        QGHGHGHGHGHGHGH---GHA-HGHGYLKDDDPRKFAEERRLFLLLGSTSILFLVCVSPM        334
                                            * *::***********************:*:***

Grimshawi         AILHMTIASEVLPSFPFQVFRALANLLELINYSITFYIYCLFSEDFRNTLLRTFNWPWVK        370
Mojavensis        AILHMTIASEVLPSFPFQVFRAMANLLELINYSITFYIYCLFSEDFRNTLMRTIKWPWLK        386
Virilism          AILHMTIASEVLPSFPFQVFRALANLLELINYSITFYIYCLFSEDFRNTLMRTIKWPWLK        374
```

```
Bipectinate       AILHMTIASEVYPSFPFQVFRASANLLELINYSLTFYIYCLFSEDFRNTLVRTIKWPWLK        386
Anannassae        AILHMTIASEVYPSFPFQVFRASANLLELINYSLTFYIYCLFSEDFRNTLVRTIKWPWLK        390
Serrata           AILHMTIASEVYPSFPFQVFRASANLLELINYSLTFYIYCLFSEDFRNTLVRTIKWPWLK        393
Kikkawei          AILHMTIASEVYPSFPFQVFRASANLLELINYSLTFYIYCLFSEDFRNTLVRTIKWPWLK        393
Fichsuphila       AILHMTIASEVYPSFPFQVFRASANLLELINYSLTFYIYCLFSEDFRNTLVRTIKWPWLK        376
Eugracilis        AILHMTIASEVYPSFPFQVFRASANLLELINYSLTFYIYCLFSEDFRNTLVRTIKWPWLK        374
Rhopaloa          AILHMTIASEVYPSFPFQVFRASANLLELINYSLTFYIYCLFSEDFRNTLVRTIKWPWLK        389
Elegans           AILHMTIASEVYPSFPFQVFRASANLLELINYSLTFYIYCLFSEDFRNTLVRTIKWPWLK        397
Erecta            AILHMTIASEVYPSFPFQVFRASANLLELINYSLTFYIYCLFSEDFRNTLVRTIKWPWLK        382
Suzuki            AILHMTIASEVYPSFPFQVFRASANLLELINYSLTFYIYCLFSEDFRNTLVRTIKWPWLK        374
Biarmipes         AILHMTIASEVYPSFPFQVFRASANLLELINYSLTFYIYCLFSEDFRNTLVRTIKWPWLK        374
Takahashi         AILHMTIASEVYPSFPFQVFRASANLLELINYSLTFYIYCLFSEDFRNTLVRTIKWPWLK        394
Sechellia         AILHMTIASEVYPSFPFQVFRASANLLELINYSLTFYIYCLFSEDFRNTLVRTIKWPWLK        390
Melanogaster      AILHMTIASEVYPSFPFQVFRASANLLELINYSLTFYIYCLFSEDFRNTLVRTIKWPWLK        394
Simulans          AILHMTIASEVYPSFPFQVFRASANLLELINYSLTFYIYCLFSEDFRNTLVRTIKWPWLK        394
Mauritania        AILHMTIASEVYPSFPFQVFRASANLLELINYSLTFYIYCLFSEDFRNTLVRTIKWPWLK        394
                  *********** ********** **********:****************:**::***:*

Grimshawi         SKLFRRQV-DEVSASPPATATAAVCNPVCPTIHIEHS-----------------------        406
Mojavensis        GKLCHQVDETTQAIMGVPMVRFEKVNMRNHHITTTSGIGR------------SSSI----        430
Virilism          SKLCHQVDET-QTIKGVPMVRFDKVTLRNHHITTTSGIGR------------SSSI----        417
Bipectinate       GKFCHQAE-HEVSASP--PATAGTVAA---AGTATTATANP-----------SPAIPANP        429
Anannassae        GKLCHQADQHEVSASP--PATAGTVAA---GIGTAPATANP-----------SPAIPANP        434
Serrata           GKFCHQAE-HEVSASP--PATAGTVAV---AVAVGHGTGP---------VSNPQPIPAIP        438
Kikkawei          GKFCHQAE-HEVSASP--PATAGTVAV---AVAVGHGTGP---------VSNPQPIPAIP        438
Fichsuphila       GKFCHQVE-HEVSASP--PATAGTVAV---AGTGT-GAG----------HVSYYH-PAIP        418
Eugracilis        GKLCHQAE-NEVSASP--PATAGTVAV---AA----GTG--------TGHISKYHAASIP        416
Rhopaloa          GKCCHQAE-HEVSASP--PATAGTVAV---AVAVA-GTGT------APGHVTNFN-PAIP        435
Elegans           GKFCHQGE-HEVSASP--PATAGTVAV---AVAAT-GTA--------SGHVSNLH-PAIP        441
Erecta            GKFCHQGE-HEVSASP--PATAGTVAV---AGIGT-GTGTGTG------NVSHFH-PVIP        428
Suzuki            GKFCHQAE-HEVSGSP--PATAGTVAV---AGIGI-GPGTGNGPGTSSRTKSNFH-PAIP        426
Biarmipes         GKFCHQAE-HEVSASP--PATAGTVAV---AGIVA-GTGPGT------GTKSNFH-PAIP        420
Takahashi         GKFCHQAE-HEVSASP--PATAGTVAV---AGLGT-GTGIG--------QVSNFH-PAIP        438
Sechellia         GKFCHQAE-HEVSASP--PATAGTVAV---AGT-----GTG--------HVSNFH-PAIP        430
Melanogaster      GKFCHQAE-HEVSASP--PATAGTVAV---AGT-----GNG--------HVSIFH-PAIP        434
Simulans          GKFCHQAE-HEVSASP--PATAGTVAV---AGT-----GTG--------HVSNFH-PAIP        434
Mauritania        GKFCHQAE-HEVSASP--PATAGTVAV---AGT-----GTG--------HVSNFH-PAIP        434
                  .*  ::      :      .

Grimshawi         ------NHPEHANGPFRKGQY-------A---        422
Mojavensis        --------------------------------        430
Virilism          --------------------------------        417
Bipectinate       APILPTDP-HGE-EPERLANGTVR--------        451
Anannassae        APILPTDHPHGEKSPQRIPNGTVR--------        458
Serrata           AVTFTPADPEDHTRPANGVLR-----------        459
Kikkawei          AVTFTPADPEDHARPPNGVLR-----------        459
Fichsuphila       SLTFTPAEPEYRPPVCQWPTSLSWQRDNTNIT        450
Eugracilis        ALTLTPAEPAEDRRP-RIANGVHH--------        439
Rhopaloa          ALILTPAEHEE--RPIRLANGVLH--------        457
Elegans           ALTLTPAEPEDRDRPNRLANGVLH--------        465
Erecta            ALTLTPAEPKDRH---RCANGVLH--------        449
Suzuki            ALTLTPAEPEDRP---RLANGVLH--------        447
Biarmipes         ALTLTPAEPEDRP---RLANGVLH--------        441
Takahashi         ALTLTPAEPEERP---RLANGVLH--------        459
Sechellia         ALTLTPAEPDERP---RCANGVLH--------        451
Melanogaster      ALTLTPAEPDERP---RCANGVLH--------        455
Simulans          ALTLTPAEPDERP---RCANGVLH--------        455
Mauritania        ALTLTPAEPDERP---RCANGVLH--------        455
```

Melanogaster NP_001027070.1

```
1 mitrlyntee dpaycsfiwg snltssvdvl aanatsvfss dlrddfyrdv edprteslre
       61 ycyglvlpii camgiignvl nlvvltrrnm rgtayiymra ystaallaiv faipfgirml
      121 vhkdrgqwee fgpafytahl elylgngclg vgvmmllvlt ieryvsvchp gfarpvmgpp
      181 gvvvfltcla tvivylpsif rgelikcilg ssdvyvylrr dntiyqqtif yrvykimlev
      241 ifklvptlvi gglnmrimmv yrrtcerrk mvlsrphaqg hghghghghg hghghahghg
      301 ylkdddprkf aeerrlflll gstsilflvc vspmailhmt iasevypsfp fqvfrasanl
      361 lelinysltf yiyclfsedf rntlvrtikw pwlkgkfchq aehevsaspp atagtvavag
      421 tgnghvsifh paipaltltp aepderprca ngvlh
```

Simulans XP_016039903.1

```
1 mitrlyntee dpaycsfiwg snltssvdvl aanatsvfgs dlrddfyrdv edprteslre
       61 ycyglvlpii camgiignvl nlvvltrrnm rgtayiymra ystaallaiv faipfgirml
      121 vhkdrgqwee fgpafytahl elylgngclg vgvmmllvlt ieryvsvchp gfarpvmgpp
      181 gvvvfltcla tvivylpsif rgelikcilg ssdvyvylrr dntiyqqtif yrvykimlev
      241 ifklvptlvi gglnmrimmv yrrtcerrrk mvlsrphaqg hghghghghg hghghahghg
      301 ylkdddprkf aeerrlflll gstsilflvc vspmailhmt iasevypsfp fqvfrasanl
      361 lelinysltf yiyclfsedf rntlvrtikw pwlkgkfchq aehevsaspp atagtvavag
      421 tgtghvsnfh paipaltltp aepderprca ngvlh
```

Suzuki XP_036676037.1

```
1 mitrlyntee dpaycsfiwg snltssaevl aanatsvfss dlrddfyrdv edprteslre
       61 ycyglvlpii camgiignvl nlivltrrnm rgtayiymra ystaallaiv faipfgirml
      121 vhkdrgqwee fgpafytahl elylgngclg vgvmmllvlt ieryvsvchp gfarpvmgpp
      181 gvvvfltcla tvivylpsif rgelikcifg ssdvyvylrr dntiyqqtif yrvykimlev
      241 ifklvptlvi gglnmrimmv yrrtcerrrq mvlsrphahg ymkdddprkf aeerrlflll
      301 gstsilflvc vspmailhmt iasevypsfp fqvfrasanl lelinysltf yiyclfsedf
      361 rntlvrtikw pwlkgkfchq aehevsgspp atagtvavag igigpgtgng pgtssrtksn
      421 fhpaipaltl tpaepedrpr langvlh
```

Mauritania XP_033171498.1

```
1 mitrlyntee dpaycsfiwg snltssvdvl aanatsvfss dlrddfyrdv edprteslre
       61 ycyglvlpii camgiignvl nlvvltrrnm rgtayiymra ystaallaiv faipfgirml
      121 vhkdrgqwee fgpafytahl elylgngclg vgvmmllvlt ieryvsvchp gfarpvmgpp
      181 gvvvfltcla tvivylpsif rgelikcilg ssdvyvylrr dntiyqqtif yrvykimlev
      241 ifklvptlvi gglnmrimmv yrrtcerrrk mvlsrphaqg hghghghghg hghghahghg
      301 ylkdddprkf aeerrlflll gstsilflvc vspmailhmt iasevypsfp fqvfrasanl
      361 lelinysltf yiyclfsedf rntlvrtikw pwlkgkfchq aehevsaspp atagtvavag
      421 tgtghvsnfh paipaltltp aepderprca ngvlh
```

Sechellia XP_002039256.1

```
    1 mitrlyntee dpaycsfiwg snltssvdvl aanatsvfss dlrddfyrdv edprteslre
       61 ycyglvlpii camgiignvl nlvvltrrnm rgtayiymra ystaallaiv faipfgirml
      121 vhkdrgqwee fgpafytahl elylgngclg vgvmmllvlt ieryvsvchp gfarpvmgpp
      181 gvvvfltcla tvivylpsif rgelikcilg ssdvyvylrr dntiyqqtif yrvykimlev
      241 ifklvptlvi gglnmrimmv yrrtcerrrk mvlsrshaqg hghghghghg hahghgylkd
      301 ddprkfaeer rlflllgsts ilflvcvspm ailhmtiase vypsfpfqvf rasanllelі
      361 nysltfyiyc lfsedfrntl vrtikwpwlk gkfchqaehe vsasppatag tvavagtgtg
      421 hvsnfhpaip altltpaepd erprcangvl h
```

Serrata KAH8362383.1

```
  1 mitrlyntee dpaycsfiwg snlssttdvl aaagnsspvf tdlrddfyrd vedprteslr
       61 eycyglllpi icamgiignv lnlivltrrn mrgtayiymr aystaallai vfaipfgirm
      121 lvhkdrgqwe efgpafytah lelylgngcl gvgvmmllvl tieryvsvch pgfarpvmgp
      181 pgvvvfltcl atvivylpsi frgelikcil gssdvyvylr rdntiyqqti fyriykimle
      241 vifklvptvl igglnmrimm vyrrtcerrr qmvlsrnhgh ghghshgqgh aqghahtngy
      301 vkdddprkfa eerrlflllg stsilflvcv spmailhmti asevypsfpf qvfrasanll
      361 elinysltfy iyclfsedfr ntlvrtikwp wlkgkfchqa ehevsasppa tagtvavava
      421 vghgtgpvsn pqpipaipav tftpadpedh trpangvlr
```

Erecta XP_026837687.1

```
   1 mitrlyntee dpaycsfiwg snltssvdvl aanatsvfss dlrddfyrdv edprteslre
       61 ycyglvlpii camgiignvl nlivltrrnm rgtayiymra ystaallaiv faipfgirml
      121 vhkdrgqwee fgpafytahl elylgngclg vgvmmllvlt ieryvsvchp gfarpvmgpp
      181 gvvvfltcla tvivylpsif rgelikcilg ssdvyvylrr dntiyqqtif yrvykimlev
      241 ifklvptlvi gglnmrimmv yrrtcerrrq mvltrshayg hghghghgyl kdddprkfae
      301 errlflllgs tsilflvcvs pmailhmtia sevypsfpfq vfrasanlle linysltfyi
      361 yclfsedfrn tlvrtikwpw lkgkfchqge hevsasppat agtvavagig tgtgtgtgnv
      421 shfhpvipal tltpaepkdr hrcangvlh
```

Takahashi XP_017002450.2

```
  1 mitrlyntee dpaycsfiwg snltssadvl aanatsvfss dlrddfyrdv edprteslre
       61 ycyglvlpii camgiignvl nlivltrrnm rgtayiymra ystaallaiv faipfgirml
      121 vhkdrgqwee fgpafytahl elylgngclg vgvmmllvlt ieryvsvchp gfarpvmgpp
      181 gvvvfltcla tvivylpsif rgelikcifg ssdvyvylrr dntiyqqtif yrvykimlev
      241 ifklvptlvi gglnlrimmv yrrtcerrrq mvlsrphahg hghghghghg hghghghghg
      301 ymkdddprkf aeerrlflll gstsilflvc vspmailhmt iasevypsfp fqvfrasanl
      361 lelinysltf yiyclfsedf rntlvrtikw pwlkgkfchq aehevsaspp atagtvavag
      421 lgtgtgigqv snfhpaipal tltpaepeer prlangvlh
```

Biarmipes XP_016962943.1

```
  1 mitrlyntee dpaycsfiwg snltssaevl aanatsifss dlrddfyrdv edprteslre
       61 ycyglvlpii camgiignvl nlivltrrnm rgtayiymra ystaallaiv faipfgirml
      121 vhkdrgqwee fgpafytahl elylgngclg vgvmmllvlt ieryvsvchp gfarpvmgpp
      181 gvvvfltcla tvivylpsif rgelikcifg ssdvyvylrr dntiyqqtif yrvykimlev
      241 ifklvptlvi gglnmrimmv yrrtcerrrq mvlsrphahg ymkdddprkf aeerrlflll
      301 gstsilflvc vspmailhmt iasevypsfp fqvfrasanl lelinysltf yiyclfsedf
      361 rntlvrtikw pwlkgkfchq aehevsaspp atagtvavag ivagtgpgtg tksnfhpaip
      421 altltpaepe drprlangvl h
```

Eugracilis XP_017074309.1

```
1 mitslyttqe dpaycsfiwg snlttsadvl tanatsvfss dlrddfyrdv edprteslre
       61 ycyglvlpii camgiignvl nlivltrrnm rgtayiymra ystaallaiv faipfgirml
      121 vhkdrgqwek fgpafytahl elylgngclg igvmmllvlt ieryvsvchp gfarpvmgpp
      181 gvvvfltcla tvivylpsif rgelikcmyg srdvyvylrr dntiyqqtif yrvykimlev
      241 ifkliptlvi gglnmrimmv yrrtcerrrq mvlsrphahg ymkdddprkf aeerrlflll
      301 gstsilflvc vspmailhmt iasevypsfp fqvfrasanl lelinysltf yiyclfsedf
      361 rntlvrtikw pwlkgklchq aenevsaspp atagtvavaa gtgtghisky haasipaltl
      421 tpaepaedrr priangvhh
```

Rhopaloa XP_016979814.1

```
1 mitrlyntee dpaycsfiwg anltssadvl anatsvfntd lrddfyrdve dprteslrey
       61 cyglvlpiic amgiignvfn livltrrnmr gtayiymray staallaivf aipfgirmlv
      121 hkdrgqweef gpafytahle lylgngclgv gvmmllvlti eryvsvchpg farpvmgppg
      181 vvvfftclat vivylpsifr gelikcifgs sdafvylrrd ntiyqqtify rvykimlevi
      241 fklvptlvig glnmrimmvy rrtcerrrqm vlsrphahah ghghahgqgh ghshgylkdd
      301 dprkfaeerr lflllgstsi lflvcvspma ilhmtiasev ypsfpfqvfr asanllelin
      361 ysltfyiycl fsedfrntlv rtikwpwlkg kcchqaehev sasppatagt vavavavagt
      421 gtapghvtnf npaipalilt paeheerpir langvlh
```

Fichsuphila XP_017042138.2

```
1 mitrlyntee dpaycsfiwg gsnltataae vlaanatsif gtdlrddfyr dvedprtesl
       61 reycyglllp iicamgiign llnlvvltrr nmrgtayiym raystaalla ivfaipfgir
      121 mlvhkdrgqw eefgpafyta hlelylgngc lgvgvmmllv ltieryvsvc hpgfvrpvmg
      181 ppgvvvfltc latvivylps ifrgelikcv lhasdvyvyl rrdntiyqqt ifyrvykiml
      241 evifklvptl vigflnmrim mvyrrtcerr rqmvlsrpha hgylkdddpr kfaeerrlfl
      301 llgstsilfl vcvspmailh mtiasevyps fpfqvfrasa nllelinysl tfyiyclfse
      361 dfrntlvrti kwpwlkgkfc hqvehevsas ppatagtvav agtgtgaghv syyhpaipsl
      421 tftpaepeyr ppvcqwptsl swqrdntnit
```

Elegans XP_017113339.1

```
1 mitrlyntee dpaycsfiwg snltssadvl anatsafssd mrddfyrdve dprteslrey
       61 cyglvlpiic amgiignvln livltrrnmr gtayiymray staallaivf aipfgirmlv
      121 hkdrgqweef gpafytahle lylgngclgv gvmmllvlti eryvsvchpg farpvmgppg
      181 vvvfltclat vivylpsifr gelikcifgs sdvfvylrrd ntiyqqtify rvykimlevi
      241 fklvptlvig glnmrimmvy rrtcerrrqm vlsrphghgh ghghghghgh gaqghghahg
      301 ghgylkdddp rkfaeerrlf lllgstsilf lvcvspmail hmtiasevyp sfpfqvfras
      361 anllelinys ltfyiyclfs edfrntlvrt ikwpwlkgkf chqgehevsa sppatagtva
      421 vavaatgtas ghvsnlhpai paltltpaep edrdrpnrla ngvlh
```

Kikkawei KAH8343513.1

```
1 mitrlyntee dpaycsfiwg snlssttdvl aaagnsspvf tdlrddfyrd vedprteslr
       61 eycyglllpi tcamgiignv lnlivltrrn mrgtayiymr aystaallai vfaipfgirm
```

```
121 lvhkdrgqwe efgpafytah lelylgngcl gvgvmmllvl tieryvsvch pgfarpvmgp
181 pgvvvfltcl atvivylpsi frgelikcil gssdvyvylr rdntiyqqti fyriykimle
241 vifklvptvl igglnmrimm vyrrtcerrr qmvlsrnhgh ghghshaqgq aqghahtngy
301 vkdddprkfa eerrlflllg stsilflvcv spmailhmti asevypsfpf qvfrasanll
361 elinysltfy iyclfsedfr ntlvrtikwp wlkgkfchqa ehevsasppa tagtvavava
421 vghgtgpvsn pqpipaipav tftpadpedh arppngvlr
```

Bipectinate **XP_017097721.2**

```
1 mitrlyhtee dpaycsfigg snlssssdvl atsanatpvf gsdlrddfyr dvedprtesl
61 reycyglllp iicamgiign vlnlivltrr nmrgiayiym raystaalla ivfaipfgir
121 mlvhkdrgqw eefgpafyta hlelylgngc lgvgvmmllv ltieryvsvc hpgfsrpvmg
181 ppgvvvfftc latvivylps ifrgelikcm lgssdvyvyl rrdntiyqqt lfyrvykiml
241 evifkliptv ligglnmrim mvyrrtcerr rqmvlsrpqa hhhhhhnang pgyvkdddpr
301 kfaeerrlfl llgstsilfl vcvspmailh mtiasevyps fpfqvfrasa nllelinysl
361 tfyiyclfse dfrntlvrti kwpwlkgkfc hqaehevsas ppatagtvaa agtattatan
421 pspaipanpa pilptdphge eperlangtv r
```

Anannassae **XP_044573352.1**

```
1 mitrlyhtee dpaycsfigg snisisssss sisssdvlat sanatpvfgs dlrddfyrdv
61 edprteslre ycyglllpii camgiignvl nlivltrrnm rgiayiymra ystaallaiv
121 faipfgirml vhkdrgqwee fgpafytahl elylgngclg vgvmmllvlt ieryvsvchp
181 gfsrpvmgpp gvvvfftcla tvivylpsif rgelikcmlg ssdvyvylrr dntiyqqtlf
241 yrvykimlev ifkliptvli gglnmrimmv yrrtcerrrq mvlsrpqahh hqngpgyvkd
301 ddprkfaeer rlflllgsts ilflvcvspm ailhmtiase vypsfpfqvf rasanlleli
361 nysltfyiyc lfsedfrntl vrtikwpwlk gklchqadqh evsasppata gtvaagigta
421 patanpspai panpapilpt dhphgekspq ripngtvr
```

Mojavensis **XP_015016833.1**

```
1 merivrnhqs ktmitrlynt eedpaycsvi wganltsssd lvlgtatnat hiyandlrdd
61 lyadvedprt ealreycygl mlpiicalgi ignvlnlivl trrnmrgtay iymraystaa
121 llaivfaipf girmlvhkdr gqweefgpaf ytahlelflg ngclgvgvmm llvltieryv
181 svchpgftrp vmgppgvvvf vtclvtfiiy lpsifrgeli kcmltsnnvy vylrrdnniy
241 qrtifysvyk imlevifklv ptvviaglnl rimlvyrrtc errrqmvlsr atyvkdddpr
301 kfaeerrlfl llgstsilfl lcvspmailh mtiasevlps fpfqvframa nllelinysi
361 tfyiyclfse dfrntlmrti kwpwlkgklc hqvdettqai mgvpmvrfek vnmrnhhitt
421 tsgigrsssi
```

Virilism **XP_032295448.1**

```
1 mitrlyntee dpaycsfiwg anltssndlv lgttansthi yandlrddly advedprtes
61 lreycyglml pvicalgiig nvlnlivltr rnmrgtayiy mraystaall aivfaipfgi
121 rmlvhkdrgq weefgpafyt ahlelflgng clgvgvmmll vltieryvsv chpgftrpvm
181 gppgvvvflt cfatfiiylp sifrgelikc mltsnnvyvy lrrdnniyqr tifysvykim
241 levifklipt vliaglnlri mlvyrrtcer rrqmvltran yvkdddprkf aeerrlflll
301 gstsilfllc vspmailhmt iasevlpsfp fqvfralanl lelinysitf yiyclfsedf
361 rntlmrtikw pwlksklchq vdetqtikgv pmvrfdkvtl rnhhitttsg igrsssi
```

Grimshawi **XP_032597329.1**

```
1 mitrlyntee dpvycsfvwg anvsngrsha anssalypdd mrddwfadae dprtellrqy
61 cygyflpfic asgiignvln livltrrnmr gpsyiymray staallaivf aipfgirmlv
121 hkdrgqweei gpafytahle lflgngclgv gvmmllvlti eryvsvcrpg atrpaigqpg
181 vvifiicvlt fifylpsifr gelikcmlts knvyvylrrd niiyqqtlfy svykvlveii
241 fkiiptfiia glnlrimwvy rrtcarrrqm vltraiyakn edprkfaeer rlflllgsts
301 ilfllcispm ailhmtiase vlpsfpfqvf ralanlleli nysitfyiyc lfsedfrntl
361 lrtfnwpwvk sklfrrqvde vsasppatat aavcnpvcpt ihiehsnhpe hangpfrkgq
421 ya
```
