## Supplementary material for "The incidence of candidate binding sites for β-arrestin in Drosophila neuropeptide GPCRs": S18 Text

***S18. Text* Multi-species analysis of CG30340**
**Supporting Figure 21**

CLUSTAL Line-ups; Genbank Reference IDs below
Predicted TM domains in YELLOW
BBS sequences in RED

```
Bipectinate       --------------------------------------MTTFSNGEEFDFSKWDFPAERI    22
Anannassae        --------------------------------------MTTFSTGEEFDFSKWDFPAERI    22
Serrata           ---------MWDQFSVMSICSMQKAVPLRKPTKQEATNMATFSSSDEFDFSKWNFPXERI    51
Kikkawei          --------------------------------------MATFSSSDEFDFSKWDFPEERI    22
Erecta            --------------------------------------MALLSSNDEFDFGKWDFPAERI    22
Melanogaster      --------------------------------------MASVSSSDDFDFGKWDFPAERI    22
Sechellia         --------------------------------------MTSVSSSGDFDFGKWDFPAERI    22
Simulans          --------------------------------------MASVSSSGDFDFGKWDFPAERI    22
Mauritania        --------------------------------------MASVSSSGDFDFGKWDFPAERI    22
Fichsuphila       --------------------------------------MSSFSSSDEFDFSKWDFPAERI    22
Rhopaloa          --------------------------------------MAAYSSSDEFDFGKWDFPAERI    22
Elegans           --------------------------------------MGTNSSNDEFDFSKWDFPAERI    22
Biarmipes         --------------------------------------MTSLLSSNEFDFSKWDFPAERI    22
Eugracilis        --------------------------------------MTSFSSSNEFDFSKWDFPAERI    22
Suzuki            --------------------------------------MTTLSSSNEFDFSKWDFPAERI    22
Takahashi         MALSISQGTYVGSVLGHVNLLLARAVPLRKASKQEANGMAANSSSDEFDFSKWDFPAERI    60
Grimshawi         ---------------------------------------------MDFDFSQWDFPEDRI    15
Mojavensis        -------------------------------------MDAHNYSSIEFDFSQWDFPAERI    23
Virilism          -------------------------------------MTAYNYSIQQFDFSQWDFPAERI    23
                                                               :***.:*:**  :**

Bipectinate       WLHKPDVEITWKICTFVPLIAFGLYGNIVMVYLIVANRSLRTPTNMIIANMAVADLLTLA    82
Anannassae        WLHKPDAEITWKICTFVPLIAFGLYGNIIMVYLIVANRSLRTPTNMIIANMAVADLLTLA    82
Serrata           WLHKPNGEITWKIITFLPLIVFGLYGNFTMVYLIAANRSLRSPTNLIIANMAVADLLTLA    111
Kikkawei          WLHKPNGEITWKIITFLPLIAFGLYGNFTMVYLIAANRSLRSPTNLIIANMAVADLLTLA    82
Erecta            WLHKSSGEIIWKICTFLPLIAFGLYGNFTMLYLIATNRSLRSPTNLIIANMAVADLLTLA    82
Melanogaster      WLHKPNGEITWKICTFLPLIAFGLYGNFSMVYVIATNRSLRSPTNLIIANMAVADLLTLA    82
Sechellia         WLHKPNGEITWKICTFLPLIAFGLYGNFTMLYLIATNRSLRSPTNLIIANMAVADLLTLA    82
Simulans          WLHKPNGEITWKICTFLPLIAFGLYGNFTMLYLIATNRSLRSPTNLIIANMAVADLLTLA    82
Mauritania        WLHKPNGEITWKICTFLPLIAFGLYGNFTMLYLIATNRSLRSPTNLIIANMAVADLLTLA    82
Fichsuphila       WLHKSDGEITWKICTFLPLIAFGLYGNFTMVYLIAANRSLRSPTNLIIANMAVADLLTLA    82
Rhopaloa          WLHKSNGEITLKIGTFLPLIAFGLYGNITMVYLIATNRSLRSPTNLIIANMAVADLLTLA    82
Elegans           WLHKSNGEITLKIGTFLPLIAFGLYGNITMVYLIAVNRSLRSPTNLIIANMAVADLLTLA    82
Biarmipes         WLHKSNGEITWKICTFLPLIAFGLYGNFTMVYLIAANRSLRSPTNLIIANMAVADLLTLA    82
Eugracilis        WLHKPSGEITWKICTFVPLIAFGLYGNFTMVYLIATNRSLRSPTNLIIANMAVADLLTLA    82
Suzuki            WLHKSNGEITWKICTFVPLIAFGLYGNFTMVYLIAANRSLRSPTNLIIANMAVADLLTLA    82
Takahashi         WLHKSNGEITWKICTFVPLIAFGLYGNFTMVYLIAANRSLRSPTNLIIANMAVADLLTLA    120
Grimshawi         WLRIPSGEIAWKVCSFLPLIIFGLYGNSVMIYLIAANRTLRTPTNLIVANMAVADCLTLL    75
Mojavensis        WRHKAIEEIAWKVCSFLPLIIFGLYANYILIYLIATNRALRSPTNLIIANMAMADLLTLL    83
Virilism          WLHKANEEIAWKIISFLPLIIFGLYGNYILIYLIATNRALRSPTNLIIANMAMADFLTLL    83
                  * :    **  *: :*:*** ****.*  ::*:*..**:**:***:*:****:** ***

Bipectinate       ICPAMFMLNDFYQNYQLGCVGCKLEGFLVVVFLITAVLNLSAVSYDRLTAIVLPRETRLT    142
Anannassae        ICPAMFMLNDFYQNYQLGCVGCKLEGFLVVVFLITAVLNLSAVSYDRLTAIVLPRETRLT    142
Serrata           ICPAMFMLNDFYQNYQLGCVGCKLEGFLVVVFLITAVLNLSVVSYDRLTAIVLPMETRLT    171
Kikkawei          ICPAMFMVNDFYQNYQLGCVGCKLEGFLVVVFLITAVLNLSVVSYDRLTAIVLPMETRLT    142
Erecta            ICPAMFMVNDFYQNYQLGCVGCKLEGFLVVLFLITAVLNLSVVSYDRLTAIVLPMETRLT    142
Melanogaster      ICPAMFMVNDFYQNYQLGCVGCKLEGFLVVVFLITAVLNLSVVSYDRLTAIVLPMETRLT    142
Sechellia         ICPAMFMVNDFYQNYQLGCVGCKLEGFLVVVFLITAVLNLSVVSYDRLTAIVLPMETRLT    142
Simulans          ICPAMFMVNDFYQNYQLGCVGCKLEGFLVVVFLITAVLNLSVVSYDRLTAIVLPMETRLT    142
Mauritania        ICPAMFMVNDFYQNYQLGCVGCKLEGFLVVVFLITAVLNLSVVSYDRLTAIVLPMETRLT    142
Fichsuphila       ICPAMFMVNDFYQNYQLGCVGCKLEGFLVVVFLIAAVLNLSVVSYDRLTAIVLPRETRLT    142
Rhopaloa          ICPAMFMVNDFYQNYQLGCVGCKLEGFLVVVFLITAVLNLSVVSYDRLTAIVLPMETRLT    142
Elegans           ICPAMFMVNDFYQNYQLGGVGCKLEGFLVVVFLITAVLNLSVVSYDRLTAIVLPMETRLT    142
Biarmipes         ICPAMFMLNDFYQNYQLGCVGCKLEGFLVVVFLITAVLNLSVVSYDRLTAIVLPMETRLT    142
Eugracilis        ICPAMFMLNDFYQNYQLGCVGCKLEGFLVVVFLITAVLNLSVVSYDRLTAIVLPMETRLT    142
Suzuki            ICPAMFMLNDFYQNYQLGYVGCKMEGFLVVVFLITAVLNLSVVSYDRLTAIVLPMETRLT    142
Takahashi         ICPAMFMVNDFYQNYQLGCVGCKLEGFLVVVFLIAAVLNLSVVSYDRLTAIVLPMETRLT    180
Grimshawi         ICPTMFMINDFYQNYQLGYVGCKMEGFVVVVFLITAVLNLSVVSYDRLTAIVLPLEKRLT    135
Mojavensis        ICPVMFLINDFYQNFQLGWVGCKLEGFLVVVFLITAVLNLSVVSYDRLTAIVLPQETRLT    143
Virilism          ICPAMFLINDFYQNYQLGCVGCKLEGFLVVVFLITAVLNLSVVSYDRLTAIVLPQETRLT    143
```

```
                         ***.**::******:*** ****:***:**:***:******.************ *.***

Bipectinate              TRGAQIVLVSTWISGILLASPLAFYRSYRVRIWKNFTERYCKENTVVLPKYWYVLITILV        202
Anannassae               MRGAQIVVVSTWISGILLASPLAFYRSYKVRIWKNFTERYCKENTAVLPKYWYVLITILV        202
Serrata                  VRGAQIVVVCTWFLGILLASPLALYRSYRVRLWKNFTERYCKENTSILPKYWYVLITILV        231
Kikkawei                 VRGAQIVVVCTWILGILLASPLALYRGYRVRVWKNFTERYCKENTSILPKYWYVLITILV        202
Erecta                   TRGVQIVVVCTWLSGILLASPLAFYRSYKVRVWKNFTERYCKENTSILPKYWYVLITILV        202
Melanogaster             IRGVQIVVVCTWVSGILLASPLAFYRSYRVRVWKNFTERYCKENTSVLPKYWYVLITILV        202
Sechellia                IRGVQIVVVCTWVSGILLASPLAFYRSFRVRVWKNFTERYCKENTSVLPKYWYVLITILV        202
Simulans                 IRGVQIVVVCTWVSGILLASPLAFYRSYRVRVWKNFTERYCKENTSVLPKYWYVLITILV        202
Mauritania               IRGVQIVVVCTWVSGILLASPLAFYRSYRVRVWKNFTERYCKENTSVLPKYWYVLITILV        202
Fichsuphila              VRGAQVVVVCTWVLGILLASPLALYRVYRVRVWKNFTERYCKENTVVLPKYWYVLITILV        202
Rhopaloa                 VRGAQIVVVCTWVLGILLASPLALYRVYRVRVWKNFTERYCKENTVVLPKYWYVLITILV        202
Elegans                  VRGAQIVVVCTWVLGILLASPLALYRAYRVRVWKNFTERYCKENTVVLPKYWYVLITILV        202
Biarmipes                VRGVQIVVVCTWLLGILLASPLAIYRAYRVRIWKNFTERYCKENTSILPKYWYVLITILV        202
Eugracilis               IRGVQVVVVCTWILGILLASPLALYRSYRVRIWKNFTERYCKENTSILPKYWYVLITILV        202
Suzuki                   VRGVQIVVVCTWILGILFASPLALYRAYRVRIWKNFTERYCKENTSILPKYWYVLITILV        202
Takahashi                VRGVQIVVVCTWVLGILLASPLALYRVYRVRIWKNFTERYCKENTSVLPKYWYVLITILV        240
Grimshawi                LRAAKIVIFCTWLAGVLLALPLAIYRDYRVRVWRNFTERYCKENINVLPKYWYVLITVLV        195
Mojavensis               LHGAKIVIACTWLTGLLLALPLAIYREYRVRIWRNFTERYCKENTNVLPKYWYVLITVLV        203
Virilism                 LCGARIVIAGTWLAGLLLALPLAIYRQYRVRIWRNFTERYCKENMTVLPKYWYVLITVLV        203
                           ..::*:  **. *:*:* ***:** ::**:*:**********  :**********:**

Bipectinate              WLPLGIMLICYIAIFYKLDRYEKRLRSRENPLTVSYKRSVAKTLFIVVVVFAALRLPFTI        262
Anannassae               WLPLGIMLICYIAIFYKLDRYEKRLRSRENPLTVSYKRSVAKTLFIVVVVFAALRLPFTI        262
Serrata                  WLPLGIMLICYIAIFYKLDRYEKRILSRENPIQVSYKRSVAKTLFIVVVVXAVLRLPFTI        291
Kikkawei                 WLPLGIMLICYIAIFYKLDRYEKRVLSRENPLQVSYKRSVAKTLFIVVVVFAVLRLPFTI        262
Erecta                   WLPLGIMLICYIAIFYKLDRYEKRVLSRENPLTVSYKRSVAKTLFIVVVVFAALRLPFTI        262
Melanogaster             WLPLGIMLICYIAIFYKLDRYEKRVLSRENPLTVSYKRSVAKTLFIVVVVFAALRLPFTI        262
Sechellia                WLPLGIMLICYIAIFYKLDRYEKRVLSRENPLTVSYKRSVAKTLFIVVAVFAALRLPFTI        262
Simulans                 WLPLGIMLICYIAIFYKLDRYEKRVLSRENPLTVSYKRSVAKTLFIVVAVFAALRLPFTI        262
Mauritania               WLPLGIMLICYIAIFYKLDRYEKRVLSRENPLTVSYKRSVAKTLFIVVAVFAALRLPFTI        262
Fichsuphila              WLPLGIMLICYAGIFYKLDRYEKRVLSRENPLSVSYKRCVAKTLFIVVVVFAVLRLPFTI        262
Rhopaloa                 WLPLGIMLICYIAIFYKLDRYEKRVLSRENPLTVSYKRSVAKTLFIVVVVFAVLRLPFTI        262
Elegans                  WLPLGIMLICYIAIFYKLDRYEKRVLRRENPLTVSYKRSVAKTLFIVVVVFAVLRLPFTI        262
Biarmipes                WLPLGIMLICYIAIFYKLDRYEKRVLSRENPLTVSYKRSVAKTLFIVVVVFAALRLPFTI        262
Eugracilis               WLPLGIMLICYIAIFYKLDRYEKRVLSRENPLTVSYKRSVAKTLFIVVVVFAVLRLPFTI        262
Suzuki                   WLPLGIMLICYIAIFYKLDRYEKRVLSRENPLTVSYKRSVAKTLFIVVVVFAALRLPFTI        262
Takahashi                WLPLGIMLICYIAIFYKLDRYEKRVLSRENPLSVSYKRSVAKTLFIVVVVFAVLRLPFTI        300
Grimshawi                WLPLGIMLICYTAIFIKLDRYEKRVLSRENPLSVNYKRSVAKTLFIVVIVFGVLRLPFTI        255
Mojavensis               WLPLSIMLICYTAIFIKLDRYEKRVLSRENPLTVSYKRSVAKTLFIVVVVFVVLRLPFTI        263
Virilism                 WLPLGIMLICYTAIFVKLDRYEKRVLSRENPLSVRYKRSVAKTLFIVVIVFVLLRLPFTI        263
                         ****.****** .** ********:  ****: * ***.********* *   *******

Bipectinate              LVVLREKYYDEDISVSSGMQLFWYISQYLMFLNAAVNPLIYGFNNENFRRAYNQISWVRR        322
Anannassae               LVVLREKYYDEDISVGSGMQLFWYISQYLMFLNAAVNPLIYGFNNENFRRAYNQISWVRR        322
Serrata                  LVVLREKYFAEDISVSNGXQLFWYISQYLMFLNAAVNPLIYGFNNENFRRAYYQISWVRR        351
Kikkawei                 LVVLREKYYAEDISVSSGMQLFWYISQYLMFLNAAVNPLIYGFNNENFRRAYYQISWVRR        322
Erecta                   LVVLREKYFGEDVSVSSGMQLFWYISQYLMFLNAAVNPLIYGFNNENFRRAYYQISWVRR        322
Melanogaster             LVVLREKYFGEDVSVSSGMQLFWYISQYLMFLNAAVNPLIYGFNNENFRRAYYQISWVRR        322
Sechellia                LVVLREKYFGEEVSVSSGMQLFWYISQYLMFLNAAVNPLIYGFNNENFRRAYYQISWVRR        322
Simulans                 LVVLREKYFGEDVSVSSGMQLFWYISQYLMFLNAAVNPLIYGFNNENFRRAYYQISWVRR        322
Mauritania               LVVLREKYFGEDVSVSSGMQLFWYISQYLMFLNAAVNPLIYGFNNENFRRAYYQISWVRR        322
Fichsuphila              LVVLREKYFDEDVSVSSGMQLFWYISQYLMFLNAAVNPLIYGFNNENFRRAYYQISWVQR        322
Rhopaloa                 LVVLREKYFDEDVSVSSGMQTFWYISQYLMFLNAAVNPLIYGFNNENFRRAYNQISWVRR        322
Elegans                  LVVLREKYFAADVSVSSGMQIFWYISQYLMFLNAAVNPLIYGFNNENFRRAYYQISWVRR        322
Biarmipes                LVVLREKYFDEDVSVSSGMQFFWYFSQYLMFLNAAVNPLIYGFNNENFRRAYHQISWVRR        322
Eugracilis               LVVLREKYFDEDVSVSSGMQLFWYISQYLMFLNAAVNPLIYGFNNENFRKAYYQISWVRR        322
Suzuki                   LVVLREKYFGEDVSVSSGMQLFWYISQYLMFLNAAVNPLIYGFNNENFRRAYYQISWVRR        322
Takahashi                LVVLREKYFDEDVSVSSGMQLFWYISQYLMFLNAAVNPLIYGFNNENFRRAYYQISWVRR        360
Grimshawi                FVVLREKYYNTEVSVDSAMQYFSYFSQYLMFVNAAVNPLIYGFNNENFRRAYAEMSCVKR        315
Mojavensis               FVVQREKYYKTAESVGCGTQYFSYFSQYLMFVNAAVNPIVYGFNNENFRRAYAQIGWVKR        323
Virilism                 FVVLREKYYSTESSVDCGMKYFSYFSQYLIFVNAAVNPIIYGFNNENFRRAYAQIACMQK        323
                         :** ****:    **. . : * *:****:*:******::*********:** ::. :::

Bipectinate              CRETTKLRRESNPEDHCCYCAFMKKGKASIKKAVEPQQPKTVEVDLSRELSTESYPTTKA        382
Anannassae               CRETTKLRKESDPSDHCCYCAFMKKGKAAVKNAEEPKQPKTAEVE-STELSTESYPTTKT        381
Serrata                  CRDAAKMSKISDSSQHCCYCAFMKNGK-LKKPEGATQPKDSVNEDVSQEMITEGPTAACS        410
Kikkawei                 CREAAKMKKSSDTSQHCCYCAFMKKGKSVKKPQGTTQSPENVDKGESLEFTTEGPTTACS        382
Erecta                   WREAAKMKKVSKTSNHCCYCAFMRKGKR--SPE-AAQQAGTVKGDVSKDISSEKES-AKS        378
Melanogaster             WRDATQMKKFSRSPDHCCYCAFMKNGKR--TSE-AAQKAGNLEKDISKDMSSAQQS-AKS        378
```

```
Sechellia         WRDAAKMKKVSGSRNHCCYCAFMKKGKR--T-----QQAGNLERDISKDMSSEQQS-AKS        374
Simulans          WRDAAKMKKVSGSTNHCCYCAFMKKGKR--T-----QQAGNLERDISKDMSSEQQS-AKS        374
Mauritania        WRDAAKMKKVSGSTNHCCYCAFMKKGKR--T-----QQAGNLERDISKDMSSEQPS-AKS        374
Fichsuphila       CRNAAKMTKESKTPSHCCYCAFMKKGQP--KAE-EPERPGNVGEDVGKTLSTGEPT-AKS        378
Rhopaloa          CREAAKMKKVSKTSNHCCYCNFMKKGKR--KAE-APQEPEKVEEDMSKDLSSEKPT-AES        378
Elegans           WRAAGKMKKVSKTSKHCCYCAFMKKGKR--KME-EPQQPKNVEEDMSKDISSEEAI-ART        378
Biarmipes         CREAAKMKKISGTSKHCCYCDFIKRGKA--KADGQAQQTGDVERDLSRDLSTEVPT-AKS        379
Eugracilis        WKDASKMKKASG-SKHCCYCAFMKKGER--KAE-EPQQPANVDKDLSKDISIEEPT-AKT        377
Suzuki            WREAAKMKKLSGKSKHCCYCDFMKRGKP--KTD-EPHQSGNVERDMSKELSTEEPT-AKS        378
Takahashi         WKDAAKMKKRSVTSKHCCYCAFMKKGKR--KTE-EPQQPGNVERDMSNDMSTEEPT-AKS        416
Grimshawi         RRA-K-----GNRVHHCCYCDFIKKNKNNKQTEANANAEKSCAKEISQSATAETKR-LQE        368
Mojavensis        RRAAS-----ANRAHNCCYCDFVKNRNGA-----VVTADQNLNKEISQSAVEETKN-LEE        372
Virilism          RRAAN-----ANRIHHCLYCDFIQNNKSG-----QANAEQRSDKEISQSAARETKK-LGA        372
                   :        .    :* ** *::. :                   .

Bipectinate       TERIRDEPGDNLVPEIEADGFI        404
Anannassae        TERIRDEPGDSLVPDIEADGFI        403
Serrata           TEIVREEPGDIQVPEVDADGFI        432
Kikkawei          TERIREDQGDMQVPEVDADGFI        404
Erecta            TKIMENDPTGLLVSEIEADGFI        400
Melanogaster      TKIVENE----FVSEIEADGFI        396
Sechellia         TKIVQNEPT-ILVSEIGADGFI        395
Simulans          TKIVQNEPTDLLVSEMGADGFI        396
Mauritania        SKIVKNEPTDLLVSEIGADGFI        396
Fichsuphila       TERARDDRAEQLVSEIEAEGFI        400
Rhopaloa          TETIRDEPADLLVSEIEANGFI        400
Elegans           TEGSRDDAADVLSSEIEADGYI        400
Biarmipes         TERIQDDRADILPSEVEVDGYI        401
Eugracilis        TERIRDDPADSLAPEIEADGFI        399
Suzuki            TKRIRDDPTDLLATEIEADGFI        400
Takahashi         TERIQDDLDYSVASGIEADGFI        438
Grimshawi         TSNLDGSVEDTLVAQIDGEGFI        390
Mojavensis        TTDNLSIAKESLVTRLNSDGFI        394
Virilism          TSN----IDETLMPQLKGEGFI        390
```

Melanogaster NP_724812.2

```
   1 masvsssddf dfgkwdfpae riwlhkpnge itwkictflp liafglygnf smvyviatnr
      61 slrsptnlii anmavadllt laicpamfmv ndfyqnyqlg cvgcklegfl vvvflitavl
     121 nlsvvsydrl taivlpmetr ltirgvqivv vctwvsgill asplafyrsy rvrvwknfte
     181 ryckentsvl pkywyvliti lvwlplgiml icyiaifykl dryekrvlsr enpltvsykr
     241 svaktlfivv vvfaalrlpf tilvvlreky fgedvsvssg mqlfwyisqy lmflnaavnp
     301 liygfnnenf rrayyqiswv rrwrdatqmk kfsrspdhcc ycafmkngkr tseaaqkagn
     361 lekdiskdms saqqsakstk ivenefvsei eadgfi
```

Simulans XP_002080785.2

```
1 masvsssgdf dfgkwdfpae riwlhkpnge itwkictflp liafglygnf tmlyliatnr
      61 slrsptnlii anmavadllt laicpamfmv ndfyqnyqlg cvgcklegfl vvvflitavl
     121 nlsvvsydrl taivlpmetr ltirgvqivv vctwvsgill asplafyrsy rvrvwknfte
     181 ryckentsvl pkywyvliti lvwlplgiml icyiaifykl dryekrvlsr enpltvsykr
     241 svaktlfivv avfaalrlpf tilvvlreky fgedvsvssg mqlfwyisqy lmflnaavnp
     301 liygfnnenf rrayyqiswv rrwrdaakmk kvsgstnhcc ycafmkkgkr tqqagnlerd
     361 iskdmsseqq sakstkivqn eptdllvsem gadgfi
```

Suzuki XP_016928738.1

```
  1 mttlsssnef dfskwdfpae riwlhksnge itwkictfvp liafglygnf tmvyliaanr
      61 slrsptnlii anmavadllt laicpamfml ndfyqnyqlg yvgckmegfl vvvflitavl
     121 nlsvvsydrl taivlpmetr ltvrgvqivv vctwilgilf asplalyray rvriwknfte
     181 ryckentsil pkywyvliti lvwlplgiml icyiaifykl dryekrvlsr enpltvsykr
     241 svaktlfivv vvfaalrlpf tilvvlreky fgedvsvssg mqlfwyisqy lmflnaavnp
     301 liygfnnenf rrayyqiswv rrwreaakmk klsgkskhcc ycdfmkrgkp ktdephqsgn
     361 verdmskels teeptakstk rirddptdll ateieadgfi
```

Mauritania XP_033156480.1

```
1 masvsssgdf dfgkwdfpae riwlhkpnge itwkictflp liafglygnf tmlyliatnr
      61 slrsptnlii anmavadllt laicpamfmv ndfyqnyqlg cvgcklegfl vvvflitavl
     121 nlsvvsydrl taivlpmetr ltirgvqivv vctwvsgill asplafyrsy rvrvwknfte
```

181 ryckentsvl pkywyvliti lvwlplgiml icyiaifykl dryekrvlsr enpltvsykr
241 svaktlfivv avfaalrlpf tilvvlreky fgedvsvssg mqlfwyisqy lmflnaavnp
301 liygfnnenf rrayyqiswv rrwrdaakmk kvsgstnhcc ycafmkkgkr tqqagnlerd
361 iskdmsseqp saksskivkn eptdllvsei gadgfi

Sechellia XP_002033117.2
1 mtsvsssgdf dfgkwdfpae riwlhkpnge itwkictflp liafglygnf tmlyliatnr
61 slrsptnlii anmavadllt laicpamfmv ndfyqnyqlg cvgcklegfl vvvflitavl
121 nlsvvsydrl taivlpmetr ltirgvqivv vctwvsgill asplafyrsf rvrvwknfte
181 ryckentsvl pkywyvliti lvwlplgiml icyiaifykl dryekrvlsr enpltvsykr
241 svaktlfivv avfaalrlpf tilvvlreky fgeevsvssg mqlfwyisqy lmflnaavnp
301 liygfnnenf rrayyqiswv rrwrdaakmk kvsgsrnhcc ycafmkkgkr tqqagnlerd
361 iskdmsseqq sakstkivqn eptilvseig adgfi

Serrata XP_020818415.1
1 mwdqfsvmsi csmqkavplr kptkqeatnm atfsssdefd fskwnfpxer iwlhkpngei
61 twkiitflpl ivfglygnft mvyliaanrs lrsptnliia nmavadlltl aicpamfmln
121 dfyqnyqlgc vgcklegflv vvflitavln lsvvsydrlt aivlpmetrl tvrgaqivvv
181 ctwflgilla splalyrsyr vrlwknfter yckentsilp kywyvlitil vwlplgimli
241 cyiaifykld ryekrilsre npiqvsykrs vaktlfivvv vxavlrlpft ilvvlrekyf
301 aedisvsngx qlfwyisqyl mflnaavnpl iygfnnenfr rayyqiswvr rcrdaakmsk
361 isdssqhccy cafmkngklk kpegatqpkd svnedvsqem itegptaacs teivreepgd
421 iqvpevdadg fi

Erecta XP_001969173.1
1 mallssndef dfgkwdfpae riwlhkssge iiwkictflp liafglygnf tmlyliatnr
61 slrsptnlii anmavadllt laicpamfmv ndfyqnyqlg cvgcklegfl vvlflitavl
121 nlsvvsydrl taivlpmetr lttrgvqivv vctwlsgill asplafyrsy kvrvwknfte
181 ryckentsil pkywyvliti lvwlplgiml icyiaifykl dryekrvlsr enpltvsykr
241 svaktlfivv vvfaalrlpf tilvvlreky fgedvsvssg mqlfwyisqy lmflnaavnp
301 liygfnnenf rrayyqiswv rrwreaakmk kvsktsnhcc ycafmrkgkr speaaqqagt
361 vkgdvskdis sekesakstk imendptgll vseieadgfi

Takahashi XP_017012153.2
1 malsisqgty vgsvlghvnl llaravplrk askqeangma ansssdefdf skwdfpaeri
61 wlhksngeit wkictfvpli afglygnftm vyliaanrsl rsptnliian mavadlltla
121 icpamfmvnd fyqnyqlgcv gcklegflvv vfliaavlnl svvsydrlta ivlpmetrlt
181 vrgvqivvvc twvlgillas plalyrvyrv riwknftery ckentsvlpk ywyvlitilv
241 wlplgimlic yiaifykldr yekrvlsren plsvsykrsv aktlfivvvv favlrlpfti
301 lvvlrekyfd edvsvssgmq lfwyisqylm flnaavnpli ygfnnenfrr ayyqiswvrr
361 wkdaakmkkr svtskhccyc afmkkgkrkt eepqqpgnve rdmsndmste eptaksteri
421 qddldysvas gieadgfi

Biarmipes XP_016967821.1
1 mtsllssnef dfskwdfpae riwlhksnge itwkictflp liafglygnf tmvyliaanr
61 slrsptnlii anmavadllt laicpamfml ndfyqnyqlg cvgcklegfl vvvflitavl
121 nlsvvsydrl taivlpmetr ltvrgvqivv vctwllgill asplaiyray rvriwknfte
181 ryckentsil pkywyvliti lvwlplgiml icyiaifykl dryekrvlsr enpltvsykr
241 svaktlfivv vvfaalrlpf tilvvlreky fdedvsvssg mqffwyfsqy lmflnaavnp
301 liygfnnenf rrayhqiswv rrcreaakmk kisgtskhcc ycdfikrgka kadgqaqqtg
361 dverdlsrdl stevptakst eriqddradi lpsevevdgy i

Eugracilis XP_017066482.1
1 mtsfsssnef dfskwdfpae riwlhkpsge itwkictfvp liafglygnf tmvyliatnr
61 slrsptnlii anmavadllt laicpamfml ndfyqnyqlg cvgcklegfl vvvflitavl
121 nlsvvsydrl taivlpmetr ltirgvqvvv vctwilgill asplalyrsy rvriwknfte
181 ryckentsil pkywyvliti lvwlplgiml icyiaifykl dryekrvlsr enpltvsykr
241 svaktlfivv vvfavlrlpf tilvvlreky fdedvsvssg mqlfwyisqy lmflnaavnp
301 liygfnnenf rkayyqiswv rrwkdaskmk kasgskhccy cafmkkgerk aeepqqpanv
361 dkdlskdisi eeptaktter irddpadsla peieadgfi

Rhopaloa **XP_016973955.1**

```
1 maaysssdef dfgkwdfpae riwlhksnge itlkigtflp liafglygni tmvyliatnr
       61 slrsptnlii anmavadllt laicpamfmv ndfyqnyqlg cvgcklegfl vvvflitavl
      121 nlsvvsydrl taivlpmetr ltvrgaqivv vctwvlgill asplalyrvy rvrvwknfte
      181 ryckentvvl pkywyvliti lvwlplgiml icyiaifykl dryekrvlsr enpltvsykr
      241 svaktlfivv vvfavlrlpf tilvvlreky fdedvsvssg mqtfwyisqy lmflnaavnp
      301 liygfnnenf rraynqiswv rrcreaakmk kvsktsnhcc ycnfmkkgkr kaeapqepek
      361 veedmskdls sekptaeste tirdepadll vseieangfi
```

Fichsuphila **XP_017044914.1**

```
    1 mssfsssdef dfskwdfpae riwlhksdge itwkictflp liafglygnf tmvyliaanr
       61 slrsptnlii anmavadllt laicpamfmv ndfyqnyqlg cvgcklegfl vvvfliaavl
      121 nlsvvsydrl taivlpretr ltvrgaqvvv vctwvlgill asplalyrvy rvrvwknfte
      181 ryckentvvl pkywyvliti lvwlplgiml icyagifykl dryekrvlsr enplsvsykr
      241 cvaktlfivv vvfavlrlpf tilvvlreky fdedvsvssg mqlfwyisqy lmflnaavnp
      301 liygfnnenf rrayyqiswv qrcrnaakmt kesktpshcc ycafmkkgqp kaeeperpgn
      361 vgedvgktls tgeptakste rarddraeql vseieaegfi
```

Elegans **XP_017120155.2**

```
     1 mgtnssndef dfskwdfpae riwlhksnge itlkigtflp liafglygni tmvyliavnr
       61 slrsptnlii anmavadllt laicpamfmv ndfyqnyqlg gvgcklegfl vvvflitavl
      121 nlsvvsydrl taivlpmetr ltvrgaqivv vctwvlgill asplalyray rvrvwknfte
      181 ryckentvvl pkywyvliti lvwlplgiml icyiaifykl dryekrvlrr enpltvsykr
      241 svaktlfivv vvfavlrlpf tilvvlreky faadvsvssg mqifwyisqy lmflnaavnp
      301 liygfnnenf rrayyqiswv rrwraagkmk kvsktskhcc ycafmkkgkr kmeepqqpkn
      361 veedmskdis seeaiartte gsrddaadvl sseieadgyi
```

Kikkawei **XP_017026956.1**

```
1 matfsssdef dfskwdfpee riwlhkpnge itwkiitflp liafglygnf tmvyliaanr
       61 slrsptnlii anmavadllt laicpamfmv ndfyqnyqlg cvgcklegfl vvvflitavl
      121 nlsvvsydrl taivlpmetr ltvrgaqivv vctwilgill asplalyrgy rvrvwknfte
      181 ryckentsil pkywyvliti lvwlplgiml icyiaifykl dryekrvlsr enplqvsykr
      241 svaktlfivv vvfavlrlpf tilvvlreky yaedisvssg mqlfwyisqy lmflnaavnp
      301 liygfnnenf rrayyqiswv rrcreaakmk kssdtsqhcc ycafmkkgks vkkpqgttqs
      361 penvdkgesl efttegptta csteriredq gdmqvpevda dgfi
```

Bipectinate **XP_017105995.2**

```
1 mttfsngeef dfskwdfpae riwlhkpdve itwkictfvp liafglygni vmvylivanr
       61 slrtptnmii anmavadllt laicpamfml ndfyqnyqlg cvgcklegfl vvvflitavl
      121 nlsavsydrl taivlpretr lttrgaqivl vstwisgill asplafyrsy rvriwknfte
      181 ryckentvvl pkywyvliti lvwlplgiml icyiaifykl dryekrlrsr enpltvsykr
      241 svaktlfivv vvfaalrlpf tilvvlreky ydedisvssg mqlfwyisqy lmflnaavnp
      301 liygfnnenf rraynqiswv rrcrettklr resnpedhcc ycafmkkgka sikkavepqq
      361 pktvevdlsr elstesyptt katerirdep gdnlvpeiea dgfi
```

Anannassae **XP_032306759.1**

```
1 mttfstgeef dfskwdfpae riwlhkpdae itwkictfvp liafglygni imvylivanr
       61 slrtptnmii anmavadllt laicpamfml ndfyqnyqlg cvgcklegfl vvvflitavl
      121 nlsavsydrl taivlpretr ltmrgaqivv vstwisgill asplafyrsy kvriwknfte
      181 ryckentavl pkywyvliti lvwlplgiml icyiaifykl dryekrlrsr enpltvsykr
      241 svaktlfivv vvfaalrlpf tilvvlreky ydedisvgsg mqlfwyisqy lmflnaavnp
      301 liygfnnenf rraynqiswv rrcrettklr kesdpsdhcc ycafmkkgka avknaeepkq
      361 pktaeveste lstesypttk tterirdepg dslvpdiead gfi
```

Mojavensis **XP_032585686.1**

```
1 mdahnyssie fdfsqwdfpa eriwrhkaie eiawkvcsfl pliifglyan yiliyliatn
       61 ralrsptnli ianmamadll tllicpvmfl indfyqnfql gwvgcklegf lvvvflitav
      121 lnlsvvsydr ltaivlpqet rltlhgakiv iactwltgll lalplaiyre yrvriwrnft
      181 eryckentnv lpkywyvlit vlvwlplsim licytaifik ldryekrvls renpltvsyk
      241 rsvaktlfiv vvvfvvlrlp ftifvvqrek yyktaesvgc gtqyfsyfsq ylmfvnaavn
      301 pivygfnnen frrayaqigw vkrrraasan rahnccycdf vknrngavvt adqnlnkeis
      361 qsaveetknl eettdnlsia keslvtrlns dgfi
```

Virilism **XP_032293011.1**

```
      1 mtaynysiqq fdfsqwdfpa eriwlhkane eiawkiisfl pliifglygn yiliyliatn
       61 ralrsptnli ianmamadfl tllicpamfl indfyqnyql gcvgcklegf lvvvflitav
      121 lnlsvvsydr ltaivlpqet rltlcgariv iagtwlagll lalplaiyrq yrvriwrnft
      181 eryckenmtv lpkywyvlit vlvwlplgim licytaifvk ldryekrvls renplsvryk
      241 rsvaktlfiv vivfvllrlp ftifvvlrek yystessvdc gmkyfsyfsq ylifvnaavn
      301 piiygfnnen frrayaqiac mqkrraanan rihhclycdf iqnnksgqan aeqrsdkeis
      361 qsaaretkkl gatsnidetl mpqlkgegfi
```

Grimshawi **XP_001995037.1**

```
1 mdfdfsqwdf pedriwlrip sgeiawkvcs flpliifgly gnsvmiylia anrtlrtptn
       61 livanmavad cltllicptm fmindfyqny qlgyvgckme gfvvvvflit avlnlsvvsy
      121 drltaivlpl ekrltlraak ivifctwlag vllalplaiy rdyrvrvwrn fteryckeni
      181 nvlpkywyvl itvlvwlplg imlicytaif ikldryekrv lsrenplsvn ykrsvaktlf
      241 ivvivfgvlr lpftifvvlr ekyyntevsv dsamqyfsyf sqylmfvnaa vnpliygfnn
      301 enfrrayaem scvkrrrakg nrvhhccycd fikknknnkq teananaeks cakeisqsat
      361 aetkrlqets nldgsvedtl vaqidgegfi
```
