## Supplementary material for "The incidence of candidate binding sites for β-arrestin in Drosophila neuropeptide GPCRs": S19 Text

### ***S19. Text*** **Multi-species analysis of PDF R PD isoforms**
### **Supporting Figure 22**

CLUSTAL Line-ups; Genbank Reference IDs below
5th, 6th and 7th Predicted TM domain in YELLOW
BBS sequences in RED

```
Serrata          MTLLSSILDSGG--------GISAQRFTRLLRQSSTSSSSSSSS---------SASASGS    43
Kikkawei         MTLLSSILDSGG--------SISAQRFTRLLRQSSTSSSS-------------SVSASAS    39
Rhopaloa         MTLLSNILDCGG--------GISAQCLTRSLRESSSPGS-P-------GSPESGAFESKS    44
Elegans          MTLLSNILDCGG--------GIPAQRLARLLRQSSSSGSTP-------SGSASGPFESKS    45
Fichsuphila      MTLLASILDCGG--------GISAQRFTRLLAQSSLLASSSASASASASASASGTFASES    52
Takahashi        MTLLSNILDCGG--------GISAQRFARLLRQSSSSGSPSASASSSASSSSGTSFESKS    52
Suzuki           MTLLSNILDCGG--------GISAQRFTRLLRQSSSSGSPSS-------SSSGTTFESKS    45
Biarmipes        MTLLSNILDCGG--------GISAQRFTRLLRQSSSPGFLSS-------SSLGTTFESKS    45
Eugracilis       MTLLSNILDSGG--------GISAQRFTRLLRQSSSSGSSSASAS----ASSYGTLESKS    48
Erecta           MTLLSNILDCGG--------CISAQRFTRLLRQSGSSVP---SPS----APAPGTFESIS    45
Melanogaster     MTLLSNILDCGG--------CISAQRFTRLLRQSGSSGP---SPS----APTAGTFESKS    45
Sechellia        MTLLSNILDCGG--------CISDQRFTHLLRQSGSSGP---SPS----APAPGTFESKS    45
Simulans         ------------------------------------------------------------    0
Mauritania       MTLLSNILDCGG--------CISAQRFTRLLRQSGSSGP---SPS----APAPGTFESKS    45
Bipectinate      MTLLSSILDSGG--------SVSVQRLTRLLRQSSPSASA-----------SGTALASES    41
Anannassae       MTLLSSILDSGG--------SVSVQRLTRLLRQSSTSSSA-----------SGPSLASDS    41
Mojavensis       -MQLAGDSEAGSMPTIITTSTSSNSFLQRIALKAVTAATTTKDR---------L----IS    46
Virilism         ------------------------------------------------------------    0
Grimshawi        MMQLPGINKPGCIPTISTSSIIIPQ---RFARRAARVASTSTAT---------VTTLSMS    48


Serrata          MLEPTSSQLPINDVLGGGRIPFLHDNGTGES----------LPLPDADALDPNFVLDGV-    92
Kikkawei         MLEPTSSQLPNNDVLGGGRIPFLPGNATGESSSSSSPPLPSPSLPDADALDPNFVLDGV-    98
Rhopaloa         MLEPTSSHI-LPT----GRVPVLHDFNASS---------------TDSP--GTFLLDGV-    81
Elegans          MLEPTTSHI-LPT----GRVPILHDFGSSS---------------TESPLTGTYLLDGV-    84
Fichsuphila      MLEPTSSHI-LPT----GRVPVLHGFDST----------------TTESTTSTYVLDGV-    90
Takahashi        MLEPTSSQN-LPT----GRVPILHDFDFDFDPT------------TTTESPGNYVLDGV-    94
Suzuki           MLEPTSSHI-LPT----GRVPILHDFDFDS---------------LTTESTGTHVLDGV-    84
Biarmipes        MLEPTSSHV-LPT----GRVPVLHDFDFDS---------------STTELPRILVLDGV-    84
Eugracilis       MLEPTSSHT-LPG----GRVPILHDFDS-----------------STTESPGAYLLDGV-    85
Erecta           MLEPTSLHS-LPT----GRVPLLHDFDA-----------------STTESPGTYVLDGV-    82
Melanogaster     MLEPTSSHS-LAT----GRVPLLHDFDA-----------------STTESPGTYVLDGV-    82
Sechellia        MLEPTSPHS-LAT----GRVPLLHDFDA-----------------STTESPGTYVLDGV-    82
Simulans         --------------------PLLHDFDA-----------------STTESPGTYVLDGV-    22
Mauritania       MLEPTSSHS-LAT----GRVPLLHDFDA-----------------TTTESPGTHVLDGV-    82
Bipectinate      MVEPTASQM-LNEITDGRRVPSL---GQDLATT------------TETTGQSNVIIDRV-    84
Anannassae       ISSMVE--------ATSLRVPSL---GHDLSTT------------T----ETNVIIDRV-    73
Mojavensis       IVETDSPA--GS-----H-MDP--STAMTTT--S--ITATATATPAATATA-S--ELSRS    89
Virilism         MPT-AAPS--SN-----RTVSTHHSTTMTSTSTT--YGTTSTTQAALTS------FE---    41
Grimshawi        MTQSGRSM--LN-----RTMTSIATATTTSTSTS--ISTHSTTSSSTSKLP-SQVISGLA    98


Serrata          --TSVAKMALEATVK-EVLRDPDPEQILANANATAPWNITL--ASAAATNYENCSALFAN    147
Kikkawei         --TSVAKMALEATVK--VLRDPDPEQILANANASAPWNITL--AAAAATNYENCSALFAN    152
Rhopaloa         --VSVAQMALEPTLT-DLLPDPDPDQVLSNLNASAPWNLTL--ASAAATNFENCSALFAN    136
Elegans          --ASVAEMALEPTVR-DILSDPDPDKVLSNLNASAPWNLTL--ASSAATNFENCSALFAN    139
Fichsuphila      --ASVAQMPLLESTVMDALPDSDPDQAFGNLNVTAPWNLTL--ASAAATNFENCSALFAN    146
Takahashi        --ASVAKMALEPTVMDTVLPDSDPDQVLSNLNVSAPWNLTL--ASAAATNFENCSALFVN    150
Suzuki           --ASVAKMALEPTVMDAVLPDSDTDQVLSNLNISAPWNITL--ASAAATNFENCSSLFVN    140
Biarmipes        --AGVATMALEPTAMDAVLPDSDPDQVLSNFNISAPWNITL--ASAAATNFENCSSLFVN    140
Eugracilis       --ASVAKMALEPAVMD-VLPDPDTDQVLSNLNITAPWNLTL--ASAAATNFENCSALFVN    140
Erecta           --ARVAQLALEPTVMD-GLPDPDTEQVLSNLNSSAPWNLTL--SSAAATNFENCSALFVN    137
Melanogaster     --ARVAQLALEPTVMD-ALPDSDTEQVLGNLNSSAPWNLTL--ASAAATNFENCSALFVN    137
Sechellia        --ARVAQLALEPTVMD-ALPDPDTEQVLGNLNSSAPWNLTL--ASAAATNFENCSALFVN    137
Simulans         --ARVAQLALEPTVMD-ALPDPDTEQVLGNLNSSAPWNLTL--ASAAATNFENCSALFVN    77
Mauritania       --ARVAQLALEPTVMD-ALPDPDTEQVLGNLNSNAPWNLTL--ASAAATNFENCSALFVN    137
Bipectinate      --TNVAKMALEATVMD--------DQVLAGVNASAHWNMT--LSSSTATNYENCSALFAN    132
Anannassae       --TNVAKMALEATVMDV-LPDPEPDQVLTGVNASSHWNMNMTLSSASATNYENCSALFAN    130
Mojavensis       DVLNASIEATTAVGMDTVKTATESGLPTVSGTTLPSWNSSF--NSALANNYDNCSAMFAN    147
```

```
Virilism        ------PMAVTAAGMDV---TMAGTEPLSSTSIPTFWNSSI--NTASANSYDNCSALFAN    90
Grimshawi       DVINGTELDMTAAATAV---GLDAAVP-DPSTTSSSWNSTF--STASANSYDNCAAMFAN    152
                                               .    ** .    :: *..::**:::*.*

Serrata         YTLPQTGLYCNWTWDTLLCWPPTPAGVLARMNCPGGYHGVDTRKFANRKCELDGRWGSRP    207
Kikkawei        YTLPQTGLYCNWTWDTLLCWPPTPAGVLARMNCPGGYHGVDTRKFANRKCELDGRWGSRP    212
Rhopaloa        YTLPQTGLYCNWTWDTLLCWPPTPAGVLARMNCPAGFHGVDTRKFAIRKCELDGRWGSRP    196
Elegans         YTLPQTGLYCNWTWDTLLCWPPTPAGVLARMNCPGGFHGVDTRKFAIRKCELDGRWGSRP    199
Fichsuphila     YTLPQSGLYCNWTWDTLLCWPPTPAGVLARMNCPGGFHGVDTRKFAIRKCELDGRWGSRP    206
Takahashi       YTLPQTGLYCNWTWDTLLCWPPTPAGVLARMNCPAGFHGVDTRKFAIRKCELDGRWGSRP    210
Suzuki          YTLPQSGLYCNWTWDTLLCWPPTPAGVLARMNCPGGFHGVDTRKFAIRKCELDGRWGSRP    200
Biarmipes       YTLPQTGLYCNWTWDTLLCWPPTPAGVLARMNCPGGFHGVDTRKFAIRKCELDGRWGSRP    200
Eugracilis      YTLPQTGLYCNWTWDTLLCWPPTPAGVLARMNCPGGFHGVDTRKFAIRKCELDGRWGSRP    200
Erecta          YTLPQTGLYCNWTWDTLLCWPPTPAGVLARMNCPGGFHGVDTRKFAIRKCELDGRWGSRP    197
Melanogaster    YTLPQTGLYCNWTWDTLLCWPPTPAGVLARMNCPGGFHGVDTRKFAIRKCELDGRWGSRP    197
Sechellia       YTLPQTGLYCNWTWDTLLCWPPTPAGVLARMNCPGGFHGVDTRKFANRKCELDGRWGSRP    197
Simulans        YTLPQTGLYCNWTWDTLLCWPPTPAGVLARMNCPGGFHGVDTRKFAIRKCELDGRWGSRP    137
Mauritania      YTLPQTGLYCNWTWDTLLCWPPTPAGVLARMNCPGGFHGVDTRKFAIRKCELDGRWGSRP    197
Bipectinate     YTLPQTGLYCNWTWDTLLCWPPTPAGVLARMHCPGGYHGVDTRKFANRKCELDGRWGSRP    192
Anannassae      YTLPQTGLYCNWTWDTLLCWPPTPAGVLARMNCPGGYHGVDTRKFANRKCELDGRWGSRP    190
Mojavensis      YTHPQTGLYCNWTWDSLLCWPPTPAGNLARMNCPAGYHGVDTRKFANRKCELDGHWGGRP    207
Virilism        YTQPTTVIYCNWTWDSLLCWPPTPAGATAHMHCPAGYHGVDTRKFANRKCELDGHWAGRP    150
Grimshawi       YTQPVTGLYCNWTWDSLLCWPPTPAGVMARMYCPAGYHGVDTRKFAIRKCELDGHWGRRP    212
                ** * : :*******:**********  *:* **.*:********* *******:*. **

Serrata         NATETNATGWTDYGPCYKPEVIRLMQQMGSREDLDLYIEIARRTRTLEIVGLCLSLFALI    267
Kikkawei        NATETNATGWTDYGPCYKPEVIRLMQQMGSREDLDLYIEIAKRTRTLEIVGLCLSLFALI    272
Rhopaloa        NATEVSPPGWTDYGPCYKPEIIRLMQQMGSR-DFDTFIDIAKKTRTLEIVGLCLSLFALI    255
Elegans         NATEVSPPGWTDYGPCYKPEIIRLMQQMGSK-DFDLYIDIARKTRTLEIVGLCLSLFALI    258
Fichsuphila     NATEVSPPGWTDYGPCYKPEIIRLMQQMGNK-DFDLYIDIARKTRTLEIVGLCLSLFALI    265
Takahashi       NATEVSPPGWTDYGPCYKPEIIRLMEQMGSK-NFDLYIDIAWKTRTLEIVGLCLSLFALI    269
Suzuki          NATEVSPPGWTDYGPCYKPEIIRLMQQMGSK-DFDLYIDIARKTRTLEIVGLCLSLFALI    259
Biarmipes       NATEVSPPGWTDYGPCYKPEIIRLMQQMGSK-DFDLYIDIARKTRTLEIVGLCLSLFALI    259
Eugracilis      NATEASPPGWTDYGPCYKPEIIRLMQQMGNK-DFDLYIDIARKTRTLEIVGLCLSLFALI    259
Erecta          NATEVSPPGWTDYGPCYKPEIIRLMQQMGSK-DFDAYIDIARRTRTLEIVGLCISLFALI    256
Melanogaster    NATEVNPPGWTDYGPCYKPEIIRLMQQMGSK-DFDAYIDIARRTRTLEIVGLCLSLFALI    256
Sechellia       NATEVSPPGWTDYGPCYKPEIIRLMQQMGSK-DFDAYIDIARRTRTLEIVGLCLSLFALI    256
Simulans        NATEVSPPGWTDYGPCYKPEIIRLMQQMGSK-DFDAYIDIARRTRTLEIVGLCLSLFALI    196
Mauritania      NATEVSPPGWTDYGPCYKPEIIRLMQQMGSK-DFDAYIDIARRTRTLEIVGLCLSLFALI    256
Bipectinate     NATEPSPAGWTDYGPCYKPEVIRLMQSMGSK-DIDIYIDIARRTRTLEIVGLWVSLFALV    251
Anannassae      NATEPSPAGWTDYGPCYKPEVIRLMQSMGSK-DIDIYIDIARRTRTLEIVGLWVSLFALV    249
Mojavensis      NETEPNHAGWTDYGPCYKPEVIRLMQEIK---DVNLYIDIAQRTRTLEIIGLCLSLLALI    264
Virilism        NSTEQKPTGWTDYGPCYKPEVIRLMQEIK---DVNLYMDIAQRTRTLEIIGLCLSLFALI    207
Grimshawi       NDTELNTGGWTDYAPCYKPEVIRLMQEIK---DVNVYMDIAQRTRTLEIIGLCLSLLALI    269
                * ** .  *****.******:****:.:    :.: :::** :******:** :**:**:

Serrata         VSLLIFCTFRSLRNNRTKIHKNLFVAMVLQVIIRLTLYLDQFRRGSKEAATNTSLSVIEN    327
Kikkawei        VSLLIFCTFRSLRNNRTKIHKNLFIAMVLQVIIRLTLYLDQFRRGNKEAATNTSLSVIEN    332
Rhopaloa        VSLLIFCTFRSLRNNRTKIHKNLFVAMVLQVIIRLTLYLDQYRRGNKEAATNTSLSVIEN    315
Elegans         VSLLIFCTFRSLRNNRTKIHKNLFVAMVLQVIIRLTLYLDQYRRGNKEAATNTSLSVIEN    318
Fichsuphila     VSLLIFCTFRSLRNNRTKIHKNLFVAMVLQVIIRLTLYLDQFRRGNKEAATNTSLSVIEN    325
Takahashi       VSLLIFCTFRSLRNNRTKIHKNLFVAMVLQVIIRLTLYLDQYRRGNKEAATNTSLSVIEN    329
Suzuki          VSLLIFCTFRSLRNNRTKIHKNLFVAMVLQVIIRLTLYLDQFRRGNKEAATNTSLSVIEN    319
Biarmipes       VSLLIFCTFRSLRNNRTKIHKNLFVAMVLQVIIRLTLYLDQFRRGNKEAATNTSLSVIEN    319
Eugracilis      VSLLIFCTFRSLRNNRTKIHKNLFVAMVLQVIIRLTLYLDQFRRGNKEAATNTSLSVIEN    319
Erecta          VSLLIFCTFRSLRNNRTKIHKNLFVAMVLQVIIRLTLYLDQFRRGNKEAATNTSLSAIEN    316
Melanogaster    VSLLIFCTFRSLRNNRTKIHKNLFVAMVLQVIIRLTLYLDQFRRGNKEAATNTSLSVIEN    316
Sechellia       VSLLIFCTFRSLRNNRTKIHKNLFVAMVLQVIIRLTLYLDQFRRGNKEAATNTSLSVIEN    316
Simulans        VSLLIFCTFRSLRNNRTKIHKNLFVAMVLQVIIRLTLYLDQFRRGNKEAATNTSLSVIEN    256
Mauritania      VSLLIFCTFRSLRNNRTKIHKNLFVAMVLQVIIRLTLYLDQFRRGNKEAATNTSLSVIEN    316
Bipectinate     ISLLIFCTFRSLRNNRTKIHKNLFIAMVLQVIVRLTLYLDQYRRGNKEAATNTSLSAIEN    311
Anannassae      ISLLIFCTFRSLRNNRTKIHKNLFVAMVLQVIVRLTLYLDQYRRGNKEAATNTSLSAIEN    309
Mojavensis      ISLVIFCAFRSLRNNRTKIHKNLFIAMVLQVIVRLTLYLDQFRRGIQ---YNSSLSAIEN    321
Virilism        ISLMIFCAFRSLRNNRTKIHKNLFVAMVLQVIVRLTLYLDQFRRGKSDSANNTSLSVIEN    267
Grimshawi       ISLVIFCAFRSLRNNRTKIHKNLFVAMVLQVIVRLTLYLDQFRRGRPETATNASVSIIEN    329
                :**:***:****************:*******:********:***      *:*:* ***

Serrata         TPYLCEASYVLLEYARTAMFMWMFIEGLYLHNMVTVAVFQGSFPLKFFSRLGWCVPILMT    387
Kikkawei        TPYLCEASYVLLEYARTAMFMWMFIEGLYLHNMVTVAVFQGSFPLKFFSRLGWCVPILMT    392
Rhopaloa        TPYLCEASYVLLEYARTAMFMWMFIEGLYLHNMVTVAVFQGSFPLKFFSRLGWCVPILMT    375
Elegans         TPYLCEASYVLLEYARTAMFMWMFIEGLYLHNMVTVAVFQGSFPLKFFSRLGWCAPILMT    378
```

```
Fichsuphila       TPYLCEASYVLLEYARTAMFMWMFIEGLYLHNMVTVAVFQGSFPLKFFSRLGWCVPILMT    385
Takahashi         TPYLCEASYVLLEYARTAMFMWMFIEGLYLHNMVTVAVFQGSFPLKFFSRLGWGFPILMT    389
Suzuki            TPYLCEASYVLLEYARTAMFMWMFIEGLYLHNMVTVAVFQGSFPLKFFSRLGWGFPILMT    379
Biarmipes         TPYLCEASYVLLEYARTAMFMWMFIEGLYLHNMVTVAVFQGSFPLKFFSRLGWGFPILMT    379
Eugracilis        TPYLCEASYVLLEYARTAMFMWMFIEGLYLHNMVTVAVFQGSFPLKFFSRLGWCVPILMT    379
Erecta            TPYLCEASYVLLEYARTAMFMWMFIEGLYLHNMVTVAVFQGSFPLKFFSRLGWCVPILMT    376
Melanogaster      TPYLCEASYVLLEYARTAMFMWMFIEGLYLHNMVTVAVFQGSFPLKFFSRLGWCVPILMT    376
Sechellia         TPYLCEASYVLLEYARTAMFMWMFIEGLYLHNMVTVAVFQGSFPLKFFSRLGWCVPILMT    376
Simulans          TPYLCEASYVLLEYARTAMFMWMFIEGLYLHNMVTVAVFQGSFPLKFFSRLGWCVPILMT    316
Mauritania        TPYLCEASYVLLEYARTAMFMWMFIEGLYLHNMVTVAVFQGSFPLKFFSRLGWCVPILMT    376
Bipectinate       TPYLCEASYVLLEYARTAMFMWMFIEGLYLHNMVTVAVFQGSFPLKFFSRLGWCVPILMT    371
Anannassae        TPYLCEASYVLLEYARTAMFMWMFIEGLYLHNMVTVAVFQGSFPLKFFSRLGWCVPILMT    369
Mojavensis        TPYLCEASYVLLEYARTAMFMWMFIEGLYLHNMVTVAVFQGSFPLIFFSLLGWGMPVVMT    381
Virilism          TPYLCEASYVLLEYARTAMFMWMFIEGLYLHNMITVAVFQGNFPLVFFSLLGWGMPVLMT    327
Grimshawi         TPYLCEASYVLLEYARTAMFMWMFIEGLYLHNMVTVAVFQGNFPLKLFALLGWGLPVLMT    389
                  *********************************:*******.*** :*: ***  *::**

Serrata           TVWARCTVIYMDTSLGDCLWNYNLTPYYWILEGPRLAVILLNFCFLVNIIRVLVMKLRQS    447
Kikkawei          TVWARCTVIYMDTSLGDCLWNYNLTPYYWILEGPRLAVILLNFCFLVNIIRVLVMKLRQS    452
Rhopaloa          TVWARCTVMYMDTSLGDCLWNYNLTPYYWILEGPRLAVILLNFCFLVNIIRVLVMKLRQS    435
Elegans           TVWARCTVMYMDTSLGECLWNYNLTPYYWILEGPRLAVILLNFCFLVNIIRVLVMKLRQS    438
Fichsuphila       TVWARCTVMYMDTSLGECLWNYNLTPYYWILEGPRLAVILLNFCFLVNIIRVLVMKLRQS    445
Takahashi         TVWARCTVMYMDTSLGECLWNYNLTPYYWILEGPRLAVILLNFCFLVNIIRVLVMKLRQS    449
Suzuki            TVWARCTVMYMDTSLGECLWNYNLTPYYWILEGPRLAVILLNFCFLVNIIRVLVMKLRQS    439
Biarmipes         TVWARCTVMYMDTSLGECLWNYNLTPYYWILEGPRLAVILLNFCFLVNIIRVLVMKLRQS    439
Eugracilis        TVWARCTVMYMDTTLGECLWNYNLTPYYWILEGPRLAVILLNFCFLVNIIRVLVMKLRQS    439
Erecta            TVWARCTVMYMDTSLGECLWNYNLTPYYWILEGPRLAVILLNFCFLVNIIRVLVMKLRQS    436
Melanogaster      TVWARCTVMYMDTSLGECLWNYNLTPYYWILEGPRLAVILLNFCFLVNIIRVLVMKLRQS    436
Sechellia         TVWARCTVMYMDTSLGECLWNYNLTPYYWILEGPRLAVILLNFCFLVNIIRVLVMKLRQS    436
Simulans          TVWARCTVMYMDTSLGECLWNYNLTPYYWILEGPRLAVILLNFCFLVNIIRVLVMKLRQS    376
Mauritania        TVWARCTVMYMDTSLGECLWNYNLTPYYWILEGPRLAVILLNFCFLVNIIRVLVMKLRQS    436
Bipectinate       FVWARCTVMYMDTSMGECLWNYNLTPYYWILEGPRLAVILLNFCFLVNIIRVLVVKLRQS    431
Anannassae        FVWARCTVMYMDTSMGECLWNYNLTPYYWILEGPRLAVILLNFCFLVNIIRVLVVKLRQS    429
Mojavensis        FVWVQCTAIFMDTALGDCMWNYNLTPYYWILEGPRLAVILLNFFFLVNIIRVLVVKLRQS    441
Virilism          FVWVQCTAIFMDTSLGDCLWNYNLTPYYWILEGPRLTVIMLNFFFLVNIIRVLVMKLRQS    387
Grimshawi         FVWVQCTAIFMDTTVGECMWNYNLTPYYWILEGPRLAVILLNFFFLVNIIRVLVVKLRQS    449
                   **.:**.::***::*:*:*****************:**:*** **********:*****

Serrata           QASDIEQTRKAVRAAIVLLPLLGITNILHQVAPLKTATNFAVWSYSTYFLTSFQGFFIAL    507
Kikkawei          QASDIEQTRKAVRAAIVLLPLLGITNILHQLAPLKTATNFAVWSYSTYFLTSFQGFFIAL    512
Rhopaloa          QASDIEQTRKAVRAAIVLLPLLGITNILHQMAPLKTATNFAVWSYGTHFLTSFQGFFIAL    495
Elegans           QASDIEQTRKAVRAAIVLLPLLGITNLLHQLAPLKTATNFAVWSYGTHFLTSFQGFFIAL    498
Fichsuphila       QASDIEQTRKAVRAAIVLLPLLGITNLLHQLAPLKTATNFAVWSYGTHFLTSFQGFFIAL    505
Takahashi         QASDIEQTRKAVRAAIVLLPLLGITNILHQMAPLKTATNFAVWSYVTHFLTSFQGFFIAL    509
Suzuki            QASDIEQTRKAVRAAIVLLPLLGITNLLHQLAPLKTATNFAVWSYGTHFLTSFQGFFIAL    499
Biarmipes         QASDIEQTRKAVRAAIVLLPLLGITNLLHQLAPLKTATNFAVWSYGTHFLTSFQGFFIAL    499
Eugracilis        QASDIEQTRKAVRAAIVLLPLLGITNLLHQLAPLKTATNFAVWSYGTHFLTSFQGFFIAL    499
Erecta            QASDIEQTRKAVRAAIVLLPLLGITNILHQLAPLKTATNFAVWSYGTHFLTSFQGFFIAL    496
Melanogaster      QASDIEQTRKAVRAAIVLLPLLGITNLLHQLAPLKTATNFAVWSYGTHFLTSFQGFFIAL    496
Sechellia         QASDIEQTRKAVRAAIVLLPLLGITNLLHQLAPLKTATNFAVWSYGTHFLTSFQGFFIAL    496
Simulans          QASDIEQTRKAVRAAIVLLPLLGITNLLHQLAPLKTATNFAVWSYGTHFLTSFQGFFIAL    436
Mauritania        QASDIEQTRKAVRAAIVLLPLLGITNLLHQLAPLKTATNFAVWSYGTHFLTSFQGFFIAL    496
Bipectinate       QASDIEQTRKAVRAAIVLLPLLGITNLLHQLAPLKTATNFAVWSYGTHFLTSFQGFFIAL    491
Anannassae        QASDIEQTRKAVRAAIVLLPLLGITNLLHQLAPLKTATNFAVWSYGTHFLTSFQGFFIAL    489
Mojavensis        QASDIEQTRKAVRAAIVLLPLLGITNLLHLVPALKTAWKFAIWSYVTHFLTSFQGFFIAL    501
Virilism          QASEIEQTRKAVRAAIVLLPLLGITNLLHLVPALKTAWKFAIWSYVTHFLTSFQGFFIAL    447
Grimshawi         QASDIEQTRKAVRAAIVLLPLLGITNLLHLVPALKTAWKFAVWSYVTHFLTSFQGFFIAL    509
                  ***:**********************:** :  **** :**:*** *:************

Serrata           IYCFLNGEVRAVLLKSLATQLSVRGHPEWAPKRASMYSGAYNTAPDTDAVQ-QHQQTGEN    566
Kikkawei          IYCFLNGEVRAVLLKSLATQLSVRGHPEWAPKRASMYSGAYNTAPDTDAVLQQQQQPGEN    572
Rhopaloa          IYCFLNGEVRAVLLKSLATQLSVRGHPEWAPKRASMYSGAYNTAPDTDAV----QPAGDP    551
Elegans           IYCFLNGEVRAVLLKSLATQLSVRGHPEWAPKRASMYSGAYNTAPDTDAV----QPAGDP    554
Fichsuphila       IYCFLNGEVRAVLLKSLATQLSVRGHPEWAPKRASMYSGAYNTAPDTDAV----HPAGDP    561
Takahashi         IYCFLNGEVRAVLLKSLATQLSVRGHPEWAPKRASMYSGAYNTAPDTDAV----QPAGDP    565
Suzuki            IYCFLNGEVRAVLLKSLATQLSVRGHPEWAPKRASMYSGAYNTAPDTDAV----QPAGDP    555
Biarmipes         IYCFLNGEVRAVLLKSLATQLSVRGHPEWAPKRASMYSGAYNTAPDTDAV----QPAGDP    555
Eugracilis        IYCFLNGEVRAVLLKSLATQLSVRGHPEWAPKRASMYSGAYNTAPDTDAV----QPAGDP    555
Erecta            IYCFLNGEVRAVLLKSLATQLSVRGHPEWAPKRASMYSGAYNTAPDTDAV----QPAGDP    552
Melanogaster      IYCFLNGEVRAVLLKSLATQLSVRGHPEWAPKRASMYSGAYNTAPDTDAV----QPAGDP    552
Sechellia         IYCFLNGEVRAVLLKSLATQLSVRGHPEWAPKRASMYSGAYNTAPDTDAV----QPVGDP    552
```

```
Simulans          IYCFLNGEVRAVLLKSLATQLSVRGHPEWAPKRASMYSGAYNTAPDTDAV----QPAGDP      492
Mauritania        IYCFLNGEVRAVLLKSLATQLSVRGHPEWAPKRASMYSGAYNTAPDTDAV----QPAGDP      552
Bipectinate       IYCFLNGEVRAVLLKSLATQMSVRGHPEWVPKRASMYSGAYNTAPDTDAVI---QQPGDN      548
Anannassae        IYCFLNGEVRAVLLKSLATQMSVRGHPEWVPKRASMYSGAYNTAPDTDAVI---QQPGEN      546
Mojavensis        IYCFLNGEVRAVLLKSLAVWMSVRGHPEWVPKRASMYSAAYNTAPDTEPPA---QQSVEA      558
Virilism          IYCFLNGEVRAVMLKSIAVWLSVRGHPEWAPKRPSMYSGAYNTAPDTDPQL---KQ-GDP      503
Grimshawi         IYCFLNGEVRTVLLKSLAVWMSVRGHPEWAPKRASMYSGAYNTAPDTDVVQ---QP-GEA      565
                  **********:*:***:*. :********.*** ****.********:      :   :

Serrata           PATGKRISPPNKRLNGRKASSASIVLIHEPQQRQRLIPRLQSKAREKDKSL--DKDRGKR      624
Kikkawei          PATGKRISPPNKRLNGRKASSASIVMIHEPQQRQRLIPRLRSRTREKDKSL--DKDREKR      630
Rhopaloa          LATGKRISPPNKRLNGRKPSSASIVMIHEPQQRQRLIPRLQNKAREKG--R--DRVEKAD      607
Elegans           SATGKRISPPHKRLNGRKPSSASIVMIHEPQQRQRLIPRLQSKAREKG--R--DRAERAD      610
Fichsuphila       SAPGKRISPPNKRLNGRKPSSASIVMIHEPQQRHRLIPRLQNKNQEKG--K--DRVEKAD      617
Takahashi         SATGKRISPPHKRLNGRKPSSASIVMIHEPQQRQRLIPRLQNKAREKS--R--DRVDKAD      621
Suzuki            SATGKRISPPHKRLNGRKPSSASIVMIHEPQQRHRLIPRLQNKEREKS--K--DRVEKAD      611
Biarmipes         LATGKRISPPHKRLNGRKPSSASIVMIHEPQQRHRLIPRLQNKAREKS--R--DRVDKAD      611
Eugracilis        SATGKRISPPNKRLNGRKPSSASIVMIHEPQQRQRLIPRLQNKAREKS--K--DRVEKAE      611
Erecta            SATGKRISPPNKRLNGRKPSSASIVMIHEPQQRQRLMPRLQNKAREKG--K--ERVEKTD      608
Melanogaster      SATGKRISPPNKRLNGRKPSSASIVMIHEPQQRQRLMPRLQNKAREKG--K--DRVEKTD      608
Sechellia         SATGKRISPPNKRLNGRKPSSASIVMIHEPQQRQRLMPRMQNKAREKG--K--DRVEKTD      608
Simulans          SATGKRISPPNKRLNGRKPSSASIVMIHEPQQRQRLMPRMQNKAREKG--K--DRVEKTD      548
Mauritania        SATGKRISPPNKRLNGRKPSSASIVMIHEPQQRQRLMPRMQNKAREKG--K--DRVEKTD      608
Bipectinate       PATGKRISPPNKRLNGRKASSASIVMIHEPQQRHRLLPRLHSQRKDRDKIK--DRDKEND      606
Anannassae        PATGKRISPPNKRLNGRKASGASIVMIHEPQQRNRLLPRLHSQRKGRDKAK--DRDMEND      604
Mojavensis        LSSNIRISPQSRRLNCRKASSVIIVIANEPQRQRAAQQ----RNNNN-TANGNGNGNGNG      613
Virilism          QQSGKRLSQSTKRSNSRKASSVTIVISTEPQIHR----YVPRRNNNNRASTGSARVRGI-      558
Grimshawi         ISTSRRVPPPIKRLKSRKANNVTIVISNEPQQQQQQHQMLQQRNTNNNSTNGS-------      618
                     . *:    :* : ** ... **:  **  :.        :   .

Serrata           IQLQQSQAEAMVADPTTITTANRIRSKDDDGGNGGGGGGGGSGSKWMMG-ICFRGQKDKC      683
Kikkawei          TQHEAVS---EPINNVTTTKANRIRSKDEDSSNGRGG----GGSKWMMG-ICFRGQKDKC      682
Rhopaloa          A----------ETEPEPDPAITRIHSKE--AGS--TASRNSRGSKWIMG-ICFRGQKDKC      652
Elegans           ADA-DAE-TNTKTEPEPDPAITRIHSKESTAGS--TGSRSNRGSKWIMG-ICFRGQKDKC      665
Fichsuphila       I------------DTELDPAITRIQSKESTG----STGSRSRGSKWIMG-ICFRGQKDKC      660
Takahashi         A----------ETDPQADPAISRIHSKESSGGGG-STGSRNRGSKWIMG-ICFRGQKDKC      669
Suzuki            AA---------ETDANPDPAISRIHSKES-GIAG-STGSRTRGSKWIMG-ICFRGQKDKC      659
Biarmipes         AA---------ETDPNQDPAISRIHSKET-GIAG-GTGSRTRGSKWIMG-ICFRGQKDKC      659
Eugracilis        PE------------TEPDPAISRIHSKET-GVGG-GTGSRTRGSKWIMG-ICFRGQKDKC      656
Erecta            KE----------AEPEPDPAISRIHSKEA-D----RARSRTRGSKWIMG-ICFRGQKDKC      652
Melanogaster      AE----------A--EPDPTISHIHSKEA-G----SARSRTRGSKWIMG-ICFRGQKDKC      650
Sechellia         AE----------A--EPDPAISRIHSKEA-G----SARSRTRGSKWIMG-ICFRGQKDKC      650
Simulans          AE----------A--EPDPAIARIHSKEA-G----SARSRTRGSKWIMG-ICFRGQKDKC      590
Mauritania        AE----------A--EPDPAISRIHSKEA-G----SARSRTRGSKWIMG-ICFRGQKDKC      650
Bipectinate       RDRQR-D-DRQLEETEADPATTRIHSKETAR------TGRNRGSKWMMDIICFRGQKDKC      658
Anannassae        KDRKRSD-DRQLEETEADPGTIRIHSKDTER------ASRNRGSKWMMDIICFRGQKDKC      657
Mojavensis        SRNQG----KDEPASGSARSGQRIRSKETP-------EQGRSTGNWMFS-LCFHGQKNKC      661
Virilism          --LKA----TEEPASGSA-VGQRIRSTDDGA------STTGRNSNWMFG-LCFRGQKNKC      604
Grimshawi         --SPR----TVEDDTGSA-MATRIRSKES----------SAGRSNWMTN-LCFRGKKNKC      660
                                        :*:*.:               .:*: . :**:*:*:**

Serrata           VMAGGQKPQQIFMTSQLPPTSTLAAIATTTITTTTGSATSATAT-------AAAKATTTI      736
Kikkawei          VMSGGQKPQQIFMTSQLPPTSTLAAVATTTITTT-GSATSATAA-------AAAK-ATAI      733
Rhopaloa          VMPGSQKTQQIFMTSQLAPTSTLAAVATTTITTTTT-----------AAAKTTIATIATI      701
Elegans           VMPGSQKTQQIFMTSQLAPTSTLAAVATTTITTTTT-----------AAAKTTIATIATI      714
Fichsuphila       VMPGSQKTQQIFMTSQLPPTSTLTPVATTTITTTTT----------AAAAKTTIATSATI      710
Takahashi         VMPGSQKTQQIFMTSQLPPTSTLAAVATTTITTTTT----------AAAAKTTIATIATI      719
Suzuki            VMPGSQKTQQIFMTSQLPPTSTLAAVATTTITTSTT----------AAAAKTTIATIATI      709
Biarmipes         VMPGSQKTQQIFMTSQLPPTSTLAAVATTTITTTTT----------AVAAKTTIATIATI      709
Eugracilis        VMPGSQKTQQIFMTSQMAPTSTLAAVATTTITTTTTTT--------AAAAKTTIATIATI      708
Erecta            VMPGSQKTQQIFMTSQMPPTSTLAVVATTMATTATTTS-------AAAAKTTIASIATI      705
Melanogaster      VMPGSQKTQQIFMTSQMPPTSTLAAVATTTITTTSTTT----------AAKTTIASIATI      700
Sechellia         VMPGSQKTQQIFMTSQMPPTSTLAAVATTTITTTT-----------AAPAKTTIASIATI      699
Simulans          VMPGSQKTQQIFMTSQMPPTSTLAAVATTTITTTSTTT--------AAAAKTTIASIATI      642
Mauritania        VMPGSQKTQQIFMTSQMPPTSTLAAVATTTITTTSTTT--------AAAAKTTIASIATI      702
Bipectinate       VMPGSQKSQQIFMTSQLPPTSKLAAITTTSLSTTTATTTTTIATVAAAAASVKTTTATTI      718
Anannassae        VMPGPQKPQQIFMTSQLPPTSKLAAITTTSLSTTTIAATTTTIATIAAAASV--KTTTTI      715
Mojavensis        VMPGA---QQIFMTSQLSATTSMATPTTTTTTTTT----------------TAIATITTT      702
Virilism          VIPNAQVSQQIFMTSQLP---------TATAATTTTTTT-----VAAAAAATTTTAATAV      656
Grimshawi         VMPGAQVSQQIFMTSQLPTSTTALATITTTTATATAATTAAAAATTTT--RAAVAAVTTT      718
                  *: .    ********:  ::      **  ::                  .     ::
```

| | | |
|---|---|---|
| Serrata | **AT**ITNSK—AKAKAKAIAKNNAK-------PKA | 760 |
| Kikkawei | ATKTKSK—AKAKAKAITKNHAK-------PKA | 757 |
| Rhopaloa | **AT**ITKS----KAKAKAIAKSHQM------PKS | 723 |
| Elegans | **AT**ITKS----KAKAKAIAKGHQM------PKS | 736 |
| Fichsuphila | **AT**ITKS----KTKAKAISKNYQI------PKA | 732 |
| Takahashi | **AT**ITKS----KAKAKAIAKSHQM------PKA | 741 |
| Suzuki | **AT**ITKS----KAKAKAIAKSHQM------PKA | 731 |
| Biarmipes | **AT**ITKS----KARAKAIASSHQI------PKA | 731 |
| Eugracilis | **AT**ITKS----KAKAKAITK---------MPKA | 727 |
| Erecta | **AT**ITKS----KAKAKAISKSH----------- | 722 |
| Melanogaster | **AT**MTKS----KAKAKAISKSHQI----QMPKA | 724 |
| Sechellia | **AT**ITKS----KAKAKAISKSHQI----QMPKA | 723 |
| Simulans | **AT**ITKS----KAKAKAISKSHQI----QMPKA | 666 |
| Mauritania | **AT**ITKS----KAKAKAISKSHQI----QMPKA | 726 |
| Bipectinate | **ST**ITKS----KAKAKAITKNHRT------PKA | 740 |
| Anannassae | **ST**ITKS----KAKAKAITKNPRT------PKA | 737 |
| Mojavensis | **S**------------------------------ | 703 |
| Virilism | AAA**TTTSRIS**SAAAAAAAAAAAILQKQQTPKA | 682 |
| Grimshawi | TALAAAAAAAAVAAAANTTKAAIA-------- | 742 |

Length of PDFR-PD CT

| | |
|---|---|
| Serrata | =760-512 |
| Kikkawei | =757-517 |
| Rhopaloa | =723-500 |
| Elegans | =736-503 |
| Fichsuphila | =732-510 |
| Takahashi | =741-514 |
| Suzuki | =731-504 |
| Biarmipes | =731-504 |
| Eugracilis | =727-504 |
| Erecta | =722-501 |
| Melanogaster | =724-501 |
| Sechellia | =723-501 |
| Simulans | =666-441 |
| Mauritania | =726-501 |
| Bipectinate | =740-496 |
| Anannassae | =737-494 |
| Mojavensis | =703-506 |
| Virilism | =682-552 |
| Grimshawi | =742-514 |

| | |
|---|---|
| Serrata | 248 |
| Kikkawei | 240 |
| Rhopaloa | 223 |
| Elegans | 233 |
| Fichsuphila | 222 |
| Takahashi | 227 |
| Suzuki | 227 |
| Biarmipes | 227 |
| Eugracilis | 223 |
| Erecta | 221 |
| Melanogaster | 223 |
| Sechellia | 222 |
| Simulans | 225 |
| Mauritania | 225 |
| Bipectinate | 244 |
| Anannassae | 243 |
| Mojavensis | 200 |
| Virilism | 220 |
| Grimshawi | 228 |

Melanogaster NP_001284826.1

```
1 mtllsnildc ggcisaqrft rllrqsgssg pspsaptagt fesksmlept sshslatgrv
       61 pllhdfdast tespgtyvld gvarvaqlal eptvmdalpd sdteqvlgnl nssapwnltl
      121 asaaatnfen csalfvnytl pqtglycnwt wdtllcwppt pagvlarmnc pggfhgvdtr
      181 kfairkceld grwgsrpnat evnppgwtdy gpcykpeiir lmqqmgskdf dayidiarrt
```

```
241 rtleivglcl slfalivsll ifctfrslrn nrtkihknlf vamvlqviir ltlyldqfrr
301 gnkeaatnts lsvientpyl ceasyvlley artamfmwmf ieglylhnmv tvavfqgsfp
361 lkffsrlgwc vpilmttvwa rctvmymdts lgeclwnynl tpyywilegp rlavillnfc
421 flvniirvlv mklrqsqasd ieqtrkavra aivllpllgi tnllhqlapl ktatnfavws
481 ygthfltsfq gffialiycf lngevravll kslatqlsvr ghpewapkra smysgaynta
541 pdtdavqpag dpsatgkris ppnkrlngrk pssasivmih epqqrqrlmp rlqnkarekg
601 kdrvektdae aepdptishi hskeagsars rtrgskwimg icfrgqkdkc vmpgsqktqq
661 ifmtsqmppt stlaavatti tttsttttaa kttiasiati atmtkskaka kaiskshqiq
721 mpka
```

Simulans XP_016037874.1

```
61 pllhdfdast tespgtyvld gvarvaqlal eptvmdalpd pdteqvlgnl nssapwnltl
121 asaaatnfen csalfvnytl pqtglycnwt wdtllcwppt pagvlarmnc pggfhgvdtr
181 kfairkceld grwgsrpnat evsppgwtdy gpcykpeiir lmqqmgskdf dayidiarrt
241 rtleivglcl slfalivsll ifctfrslrn nrtkihknlf vamvlqviir ltlyldqfrr
301 gnkeaatnts lsvientpyl ceasyvlley artamfmwmf ieglylhnmv tvavfqgsfp
361 lkffsrlgwc vpilmttvwa rctvmymdts lgeclwnynl tpyywilegp rlavillnfc
421 flvniirvlv mklrqsqasd ieqtrkavra aivllpllgi tnllhqlapl ktatnfavws
481 ygthfltsfq gffialiycf lngevravll kslatqlsvr ghpewapkra smysgaynta
541 pdtdavqpag dpsatgkris ppnkrlngrk pssasivmih epqqrqrlmp rmqnkarekg
601 kdrvektdae aepdpaiari hskeagsars rtrgskwimg icfrgqkdkc vmpgsqktqq
661 ifmtsqmppt stlaavatti tttsttttaa aakttiasia tiatitkska kakaiskshq
721 iqmpka
```

Suzuki XP_016939141.1

```
1 mtllsnildc gggisaqrft rllrqsssg spssssgtt fesksmlept sshilptgrv
61 pilhdfdfds lttestgthv ldgvasvakm aleptvmdav lpdsdtdqvl snlnisapwn
121 itlasaaatn fencsslfvn ytlpqsglyc nwtwdtllcw pptpagvlar mncpggfhgv
181 dtrkfairkc eldgrwgsrp natevsppgw tdygpcykpe iirlmqqmgs kdfdlyidia
241 rktrtleivg lclslfaliv sllifctfrs lrnnrtkihk nlfvamvlqv iirltlyldq
301 frrgnkeaat ntslsvient pylceasyvl leyartamfm wmfieglylh nmvtvavfqg
361 sfplkffsrl gwgfpilmtt vwarctvmym dtslgeclwn ynltpyywil egprlavill
421 nfcflvniir vlvmklrqsq asdieqtrka vraaivllpl lgitnllhql aplktatnfa
481 vwsygthflt sfqgffiali ycflngevra vllkslatql svrghpewap krasmysgay
541 ntapdtdavq pagdpsatgk rispphkrln grkpssasiv mihepqqrhr liprlqnker
601 ekskdrveka daaetdanpd paisrihske sgiagstgsr trgskwimgi cfrgqkdkcv
661 mpgsqktqqi fmtsqlppts tlaavattit tstttaaaak ttiatiatia titkskakak
721 aiakshqmpk a
```

Mauritania XP_033169658.1

```
1 mtllsnildc ggcisaqrft rllrqsgssg pspsapapgt fesksmlept sshslatgrv
61 pllhdfdatt tespgthvld gvarvaqlal eptvmdalpd pdteqvlgnl nsnapwnltl
121 asaaatnfen csalfvnytl pqtglycnwt wdtllcwppt pagvlarmnc pggfhgvdtr
181 kfairkceld grwgsrpnat evsppgwtdy gpcykpeiir lmqqmgskdf dayidiarrt
241 rtleivglcl slfalivsll ifctfrslrn nrtkihknlf vamvlqviir ltlyldqfrr
301 gnkeaatnts lsvientpyl ceasyvlley artamfmwmf ieglylhnmv tvavfqgsfp
361 lkffsrlgwc vpilmttvwa rctvmymdts lgeclwnynl tpyywilegp rlavillnfc
421 flvniirvlv mklrqsqasd ieqtrkavra aivllpllgi tnllhqlapl ktatnfavws
481 ygthfltsfq gffialiycf lngevravll kslatqlsvr ghpewapkra smysgaynta
541 pdtdavqpag dpsatgkris ppnkrlngrk pssasivmih epqqrqrlmp rmqnkarekg
601 kdrvektdae aepdpaisri hskeagsars rtrgskwimg icfrgqkdkc vmpgsqktqq
661 ifmtsqmppt stlaavatti tttsttttaa aakttiasia tiatitkska kakaiskshq
721 iqmpka
```

Sechellia XP_032581353.1

```
1 mtllsnildc ggcisdqrft hllrqsgssg pspsapapgt fesksmlept sphslatgrv
61 pllhdfdast tespgtyvld gvarvaqlal eptvmdalpd pdteqvlgnl nssapwnltl
121 asaaatnfen csalfvnytl pqtglycnwt wdtllcwppt pagvlarmnc pggfhgvdtr
181 kfanrkceld grwgsrpnat evsppgwtdy gpcykpeiir lmqqmgskdf dayidiarrt
241 rtleivglcl slfalivsll ifctfrslrn nrtkihknlf vamvlqviir ltlyldqfrr
301 gnkeaatnts lsvientpyl ceasyvlley artamfmwmf ieglylhnmv tvavfqgsfp
361 lkffsrlgwc vpilmttvwa rctvmymdts lgeclwnynl tpyywilegp rlavillnfc
421 flvniirvlv mklrqsqasd ieqtrkavra aivllpllgi tnllhqlapl ktatnfavws
481 ygthfltsfq gffialiycf lngevravll kslatqlsvr ghpewapkra smysgaynta
541 pdtdavqpvg dpsatgkris ppnkrlngrk pssasivmih epqqrqrlmp rmqnkarekg
```

601 kdrvektdae aepdpaisri hskeagsars rtrgskwimg icfrgqkdkc vmpgsqktqq
661 ifmtsqmppt stlaavatti tttttaapak ttiasiatia titkskakak aiskshqiqm
721 pka

Serrata XP_020802784.1

1 mtllssilds gggisaqrft rllrqsstss sssssssasa sgsmleptss qlpindvlgg
61 gripflhdng tgeslplpda daldpnfvld gvtsvakmal eatvkevlrd pdpeqilana
121 natapwnitl asaaatnyen csalfanytl pqtglycnwt wdtllcwppt pagvlarmnc
181 pggyhgvdtr kfanrkceld grwgsrpnat etnatgwtdy gpcykpevir lmqqmgsred
241 ldlyieiarr trtleivglc lslfalivsl lifctfrslr nnrtkihknl fvamvlqvii
301 rltlyldqfr rgskeaatnt slsvientpy lceasyvlle yartamfmwm fieglylhnm
361 vtvavfqgsf plkffsrlgw cvpilmttvw arctviymdt slgdclwnyn ltpyywileg
421 prlavillnf cflvniirvl vmklrqsqas dieqtrkavr aaivllpllg itnilhqvap
481 lktatnfavw systyfltsf qgffialiyc flngevravl lkslatqlsv rghpewapkr
541 asmysgaynt apdtdavqqh qqtgenpatg krisppnkrl ngrkassasi vlihepqqrq
601 rliprlqska rekdksldkd rgkriqlqqs qaeamvadpt tittanrirs kdddggnggg
661 gggggsgskw mmgicfrgqk dkcvmaggqk pqqifmtsql pptstlaaia tttittttgs
721 atsatataaa katttiatit nskakakaka iaknnakpka

Eugracilis XP_017067614.1

```
1 mtllsnilds gggisaqrft rllrqssssg sssasasass ygtlesksml eptsshtlpg
61 grvpilhdfd ssttespgay lldgvasvak malepavmdv lpdpdtdqvl snlnitapwn
121 ltlasaaatn fencsalfvn ytlpqtglyc nwtwdtllcw pptpagvlar mncpggfhgv
181 dtrkfairkc eldgrwgsrp nateasppgw tdygpcykpe iirlmqqmgn kdfdlyidia
241 rktrtleivg lclslfaliv sllifctfrs lrnnrtkihk nlfvamvlqv iirltlyldq
301 frrgnkeaat ntslsvient pylceasyvl leyartamfm wmfieglylh nmvtvavfqg
361 sfplkffsrl gwcvpilmtt vwarctvmym dttlgeclwn ynltpyywil egprlavill
421 nfcflvniir vlvmklrqsq asdieqtrka vraaivllpl lgitnllhql aplktatnfa
481 vwsygthflt sfqgffiali ycflngevra vllkslatql svrghpewap krasmysgay
541 ntapdtdavq pagdpsatgk risppnkrln grkpssasiv mihepqqrqr liprlqnkar
601 ekskdrveka epetepdpai srihsketgv gggtgsrtrg skwimgicfr gqkdkcvmpg
661 sqktqqifmt sqmaptstla avattitttt ttttaaaakt tiatiatiat itkskakaka
721 itkmpka
```

Rhopaloa XP_016982481.1

```
1 mtllsnildc gggisaqclt rslressspg spgspesgaf esksmlepts shilptgrvp
61 vlhdfnasst dspgtflldg vvsvaqmale ptltdllpdp dpdqvlsnln asapwnltla
121 saaatnfenc salfanytlp qtglycnwtw dtllcwpptp agvlarmncp agfhgvdtrk
181 fairkceldg rwgsrpnate vsppgwtdyg pcykpeiirl mqqmgsrdfd tfidiakktr
241 tleivglcls lfalivslli fctfrslrnn rtkihknlfv amvlqviirl tlyldqyrrg
301 nkeaatntsl svientpylc easyvlleya rtamfmwmfi eglylhnmvt vavfqgsfpl
361 kffsrlgwcv pilmttvwar ctvmymdtsl gdclwnynlt pyywilegpr lavillnfcf
421 lvniirvlvm klrqsqasdi eqtrkavraa ivllpllgit nilhqmaplk tatnfavwsy
481 gthfltsfqg ffialiycfl ngevravllk slatqlsvrg hpewapkras mysgayntap
541 dtdavqpagd platgkrisp pnkrlngrkp ssasivmihe pqqrqrlipr lqnkarekgr
601 drvekadaet epepdpaitr ihskeagsta srnsrgskwi mgicfrgqkd kcvmpgsqkt
661 qqifmtsqla ptstlaavat titttttttaa akttiatiat iatitkskak akaiakshqm
721 pks
```

Fichsuphila XP_017048315.1

```
1 mtllasildc gggisaqrft rllaqsslla sssasasasa sasasgtfas esmleptssh
61 ilptgrvpvl hgfdsttes ttstyvldgv asvaqmplle stvmdalpds dpdqafgnln
121 vtapwnltla saaatnfenc salfanytlp qsglycnwtw dtllcwpptp agvlarmncp
181 ggfhgvdtrk fairkceldg rwgsrpnate vsppgwtdyg pcykpeiirl mqqmgnkdfd
241 lyidiarktr tleivglcls lfalivslli fctfrslrnn rtkihknlfv amvlqviirl
301 tlyldqfrrg nkeaatntsl svientpylc easyvlleya rtamfmwmfi eglylhnmvt
361 vavfqgsfpl kffsrlgwcv pilmttvwar ctvmymdtsl geclwnynlt pyywilegpr
421 lavillnfcf lvniirvlvm klrqsqasdi eqtrkavraa ivllpllgit nllhqlaplk
481 tatnfavwsy gthfltsfqg ffialiycfl ngevravllk slatqlsvrg hpewapkras
541 mysgayntap dtdavhpagd psapgkrisp pnkrlngrkp ssasivmihe pqqrhrlipr
601 lqnknqekgk drvekadidt eldpaitriq skestgstgs rsrgskwimg icfrgqkdkc
661 vmpgsqktqq ifmtsqlppt stltpvatti ttttttaaaa kttiatsati atitksktka
721 kaisknyqip ka
```

Elegans XP_017114250.1

```
1 mtllsnildc gggipaqrla rllrqssssg stpsgsasgp fesksmlept tshilptgrv
61 pilhdfgsss tespltgtyl ldgvasvaem aleptvrdil sdpdpdkvls nlnasapwnl
121 tlassaatnf encsalfany tlpqtglycn wtwdtllcwp ptpagvlarm ncpggfhgvd
181 trkfairkce ldgrwgsrpn atevsppgwt dygpcykpei irlmqqmgsk dfdlyidiar
241 ktrtleivgl clslfalivs llifctfrsl rnnrtkihkn lfvamvlqvi irltlyldqy
301 rrgnkeaatn tslsvientp ylceasyvll eyartamfmw mfieglylhn mvtvavfqgs
361 fplkffsrlg wcapilmttv warctvmymd tslgeclwny nltpyywile gprlavilln
421 fcflvniirv lvmklrqsqa sdieqtrkav raaivllpll gitnllhqla plktatnfav
481 wsygthflts fqgffialiy cflngevrav llkslatqls vrghpewapk rasmysgayn
541 tapdtdavqp agdpsatgkr ispphkrlng rkpssasivm iheppqrqrl iprlqskare
601 kgrdraerad adadaetntk tepepdpait rihskestag stgsrsnrgs kwimgicfrg
661 qkdkcvmpgs qktqqifmts qlaptstlaa vattittttt taaakttiat iatiatitks
721 kakakaiakg hqmpks
```

Kikkawei XP_017037981.1

```
1 mtllssilds ggsisaqrft rllrqsstss sssvsasasm leptssqlpn ndvlgggrip
61 flpgnatges sssspplps pslpdadald pnfvldgvts vakmaleatv kvlrdpdpeq
121 ilananasap wnitlaaaaa tnyencsalf anytlpqtgl ycnwtwdtll cwpptpagvl
181 armncpggyh gvdtrkfanr kceldgrwgs rpnatetnat gwtdygpcyk pevirlmqqm
241 gsredldlyi eiakrtrtle ivglclslfa livsllifct frslrnnrtk ihknlfiamv
```

```
301 lqviirltly ldqfrrgnke aatntslsvi entpylceas yvlleyarta mfmwmfiegl
361 ylhnmvtvav fqgsfplkff srlgwcvpil mttvwarctv iymdtslgdc lwnynltpyy
421 wilegprlav illnfcflvn iirvlvmklr qsqasdieqt rkavraaivl lpllgitnil
481 hqlaplktat nfavwsysty fltsfqgffi aliycflnge vravllksla tqlsvrghpe
541 wapkrasmys gayntapdtd avlqqqqqpg enpatgkris ppnkrlngrk assasivmih
601 epqqrqrlip rlrsrtrekd ksldkdrekr tqheavsepi nnvtttkanr irskdedssn
661 grggggskwm mgicfrgqkd kcvmsggqkp qqifmtsqlp ptstlaavat ttitttgsat
721 sataaaaaka taiatktksk akakakaitk nhakpka
```

Bipectinate XP_017095129.2

```
  1 mtllssilds ggsvsvqrlt rllrqsspsa sasgtalase smveptasqm lneitdgrrv
     61 pslgqdlatt tettgqsnvi idrvtnvakm aleatvmddq vlagvnasah wnmtlsssta
    121 tnyencsalf anytlpqtgl ycnwtwdtll cwpptpagvl armhcpggyh gvdtrkfanr
    181 kceldgrwgs rpnatepspa gwtdygpcyk pevirlmqsm gskdidiyid iarrtrtlei
    241 vglwvslfal visllifctf rslrnnrtki hknlfiamvl qvivrltlyl dqyrrgnkea
    301 atntslsaie ntpylceasy vlleyartam fmwmfiegly lhnmvtvavf qgsfplkffs
    361 rlgwcvpilm tfvwarctvm ymdtsmgecl wnynltpyyw ilegprlavi llnfcflvni
    421 irvlvvklrq sqasdieqtr kavraaivll pllgitnllh qlaplktatn favwsygthf
    481 ltsfqgffia liycflngev ravllkslat qmsvrghpew vpkrasmysg ayntapdtda
    541 viqqpgdnpa tgkrisppnk rlngrkassa sivmihepqq rhrllprlhs qrkdrdkikd
    601 rdkendrdrq rddrqleete adpattrihs ketartgrnr gskwmmdiic frgqkdkcvm
    661 pgsqksqqif mtsqlpptsk laaitttsls tttattttti atvaaaaasv ktttattist
    721 itkskakaka itknhrtpka
```

Anannassae XP_014759839.1

```
  1 mtllssilds ggsvsvqrlt rllrqsstss sasgpslasd sissmveats lrvpslghdl
     61 sttteтnvii drvtnvakma leatvmdvlp dpepdqvltg vnasshwnmn mtlssasatn
    121 yencsalfan ytlpqtglyc nwtwdtllcw pptpagvlar mncpggyhgv dtrkfanrkc
    181 eldgrwgsrp natepspagw tdygpcykpe virlmqsmgs kdidiyidia rrtrtleivg
    241 lwvslfalvi sllifctfrs lrnnrtkihk nlfvamvlqv ivrltlyldq yrrgnkeaat
    301 ntslsaient pylceasyvl leyartamfm wmfieglylh nmvtvavfqg sfplkffsrl
    361 gwcvpilmtf vwarctvmym dtsmgeclwn ynltpyywil egprlavill nfcflvniir
    421 vlvvklrqsq asdieqtrka vraaivllpl lgitnllhql aplktatnfa vwsygthflt
    481 sfqgffiali ycflngevra vllkslatqm svrghpewvp krasmysgay ntapdtdavi
    541 qqpgenpatg krisppnkrl ngrkasgasi vmihepqqrn rllprlhsqr kgrdkakdrd
    601 mendkdrkrs ddrqleetea dpgtirihsk dterasrnrg skwmmdiicf rgqkdkcvmp
    661 gpqkpqqifm tsqlpptskl aaitttslst ttiaatttti atiaaaasvk ttttistitk
    721 skakakaitk nprtpka
```

Mojavensis XP_015016735.1

```
1 mqlagdseag smptiittst ssnsflqria lkavtaattt kdrlisivet dspagshmdp
     61 stamtttsit atatatpaat ataselsrsd vlnasieatt avgmdtvkta tesglptvsg
    121 ttlpswnssf nsalannydn csamfanyth pqtglycnwt wdsllcwppt pagnlarmnc
    181 pagyhgvdtr kfanrkceld ghwggrpnet epnhagwtdy gpcykpevir lmqeikdvnl
    241 yidiaqrtrt leiiglclsl laliislvif cafrslrnnr tkihknlfia mvlqvivrlt
    301 lyldqfrrgi qynsslsaie ntpylceasy vlleyartam fmwmfiegly lhnmvtvavf
    361 qgsfpliffs llgwgmpvvm tfvwvqctai fmdtalgdcm wnynltpyyw ilegprlavi
    421 llnffflvni irvlvvklrq sqasdieqtr kavraaivll pllgitnllh lvpalktawk
    481 faiwsyvthf ltsfqgffia liycflngev ravllkslav wmsvrghpew vpkrasmysa
    541 ayntapdtep paqqsveals snirispqsr rlncrkassv iivianepqr qraaqqrnnn
    601 ntangngngn gngsrnqgkd epasgsarsg qrirsketpe qgrstgnwmf slcfhgqknk
    661 cvmpgaqqif mtsqlsatts matptttttt ttttaiatit tts
```

```
       R   N   H   N   N   N   K   L   N   S   R   S   S   S   S   R   E   N   I   A     F1
      A   T   I   T   T   T   S   *   T   A   A   A   A   A   V   V   K   I   L   Q     F2
        Q   P   *   Q   Q   Q   A   E   Q   P   Q   Q   Q   Q   S   *   K   Y   C   K   F3
     1 CGCAACCATAACAACAACAAGCTGAACAGCCGCAGCAGCAGCAGTCGTGAAAATATTGCA 60
       ----:----|----:----|----:----|----:----|----:----|----:----|

       K   A   K   I   L   T   V   N   L   K   N   C   *   T   P   K   A   *   R   E     F1
        K   P   K   Y   *   L   *   T   L   R   T   V   K   R   L   K   L   N   A   S    F2
         S   Q   N   T   N   C   K   P   *   E   L   L   N   A   *   S   L   T   R   A   F3
    61 AAAGCCAAAATACTAACTGTAAACCTTAAGAACTGTTAAACGCCTAAAGCTTAACGCGAG 120
```

Virilism XP_032288761.1

```
1 mptaapssnr tvsthhsttm tststtygtt sttqaaltsf epmavtaagm dvtmagtepl
      61 sstsiptfwn ssintasans ydncsalfan ytqpttviyc nwtwdsllcw pptpagatah
     121 mhcpagyhgv dtrkfanrkc eldghwagrp nsteqkptgw tdygpcykpe virlmqeikd
     181 vnlymdiaqr trtleiiglc lslfaliisl mifcafrslr nnrtkihknl fvamvlqviv
     241 rltlyldqfr rgksdsannt slsvientpy lceasyvlle yartamfmwm fieglylhnm
     301 itvavfqgnf plvffsllgw gmpvlmtfvw vqctaifmdt slgdclwnyn ltpyywileg
     361 prltvimlnf fflvniirvl vmklrqsqas eieqtrkavr aaivllpllg itnllhlvpa
     421 lktawkfaiw syvthfltsf qgffialiyc flngevravm lksiavwlsv rghpewapkr
     481 psmysgaynt apdtdpqlkq gdpqqsgkrl sqstkrsnsr kassvtivis tepqihryvp
     541 rrnnnnrast gsarvrgilk ateepasgsa vgqrirstdd gasttgrnsn wmfglcfrgq
     601 knkcvipnaq vsqqifmtsq lptataattt ttttvaaaaa attttaatav aaatttsris
     661 saaaaaaaaa aailqkqqtp ka
```

Grimshawi XP_001991602.2

```
  1 mmqlpginkp gciptistss iiipqrfarr aarvaststa tvttlsmsmt qsgrsmlnrt
      61 mtsiatattt ststsisths ttssstsklp sqvisgladv ingteldmta aatavgldaa
     121 vpdpsttsss wnstfstasa nsydncaamf anytqpvtgl ycnwtwdsll cwpptpagvm
     181 armycpagyh gvdtrkfair kceldghwgr rpndtelntg gwtdyapcyk pevirlmqei
     241 kdvnvymdia qrtrtleiig lclsllalii slvifcafrs lrnnrtkihk nlfvamvlqv
     301 ivrltlyldq frrgrpetat nasvsiient pylceasyvl leyartamfm wmfieglylh
     361 nmvtvavfqg nfplklfall gwglpvlmtf vwvqctaifm dttvgecmwn ynltpyywil
     421 egprlavill nffflvniir vlvvklrqsq asdieqtrka vraaivllpl lgitnllhlv
     481 palktawkfa vwsyvthflt sfqgffiali ycflngevrt vllkslavwm svrghpewap
     541 krasmysgay ntapdtdvvq qpgeaistsr rvpppikrlk srkannvtiv isnepqqqqq
     601 qhqmlqqrnt nnnstngssp rtveddtgsa matrirskes sagrsnwmtn lcfrgkknkc
     661 vmpgaqvsqq ifmtsqlpts ttalatittt tatataatta aaaattttra avaavtttta
     721 laaaaaaaav aaaanttkaa ia
```
