## Supplementary material for "The incidence of candidate binding sites for β-arrestin in Drosophila neuropeptide GPCRs": S20 Text

### *S20. Text* Multi-species analysis of AstA R2 PA isoforms
### Supporting Figure 23

CLUSTAL Line-ups; Genbank Reference IDs below

Predicted TM domains in YELLOW

BBS sequences in RED

```
Grimshawi         MNQSYIMENNTPSMSNSSWLELKLTT---NNSTGNSSLSLDA--EEEEAIRQTVRWLVPL     55
Virilis           MNLS---NITLTLPSNSSWLESQLELTTASSNLDNSTLSSFY--AEAEAIRATVRWVVPF     55
Mojavensis        MELS---NLTTTLATQLHQ----LATSSLNHSIGPDNSTALQ--SEEEAIRATVRWLVPI     51
Bipectinate       ------MDNTTLATPDP----------LATMATYDDNLTSFFSEEERQAIEGTLIWLVPF     44
Anannassae        ------MDNTTLATSEL----------LATMATYDDNFTSFFSEEERQAIEGTLIWLVPF     44
Serrata           ------MENTTLIAGET-----LANISLANITQSDENITSFFTEEEWLALTGILPWLVGF     49
Kikkawei          ------MENTTLITGET-----LANISLANTTRSDENITSFFTEEEWLALTGILPWLVGF     49
Eugracilis        ------MENM--DNSTM-----VVDLV-VNSTRNDENMTSFFTDEEWVAIRGTLPWIVIF     46
Ficusphila        ------MENATFFPAKM-----LGSLA-SNATRNDENLTTFFTDEEWLAINSTLPWVVGF     48
Takahashi         ------MENA------T-----IMLNK-INSSRMEENSTSYFTDEEWLSISSTLPWIVCF     42
Rhopaloa          ------MDNITVSPAEM-----LGNIS-INASRNDENLTSFFTDEEWLAINGTLPWIVGF     48
Elegans           ------MDNITESAAEM-----LANIS-LNASRNDENLTSFFTDEEWLAISGTLPWIVGF     48
Erecta            -----------MENTAL-----LANIS-LNATRNEENITSFFTDEEWLAISGTLPWIVGF     43
Mauritania        -----------MENTTM-----LANIS-LNATRNEENITSFFTDEEWLAINGTLPWIVGF     43
Sechelia          -----------MENTTM-----LANIS-LNATRNEENITSFFTDEEWLAINGTLPWIVGF     43
Melanogaster      -----------MENTTM-----LANIS-LNATRNEENITSFFTDEEWLAINGTLPWIVGF     43
simulans          -----------MENTTM-----LANIS-LNATRNEENITSFFTDEEWLAINGTLPWIVGF     43
Suzuki            --------MDTMENSTL-----LGKIS-LNASRSDENITSFFTDEEWLAISGTLPWIVVF     46
biarmipes         --------MDTMDNATL-----LGKIS-LNASRNDENITSFFTDEEWLAINGTLPWIVAF     46
                                               .     .. :      *  ::   : *:* :

Grimshawi         FFGIIAISGFFFGNLLVILVVLLNKNMHSTTNLLIVNLAAADLLFVIFCVPFTAVDYVAD     115
Virilis           FFGIIAISGF-FGNLLVILVVLLNKNMHSTTNLLIVNLAAADLLFVIFCVPFTAIDYVTQ     114
Mojavensis        FFGIIAVSGF-FGNLLVILVVLLNKNMHSTTNLLIVNLAAADLLFVIFCVPFTAVDYVTQ     110
Bipectinate       FFGIIALTGF-FGNLLVILVVVFNKNMRSTTNLMIVNLAVADLLFVIFCIPFTATDYVTK     103
Anannassae        FFGIIALTGF-FGNLLVILVVVFNKNMRSTTNLMIVNLAVADLLFVIFCIPFTATDYVTK     103
Serrata           TFGAIAITGF-LGNLLVILVVVFNNNMRSTTNLMIVNLAAADLLFVILCIPFTAADYVTD     108
Kikkawei          TFGAIAITGF-LGNLLVILVVVFNNNMRSTTNLMIVNLAVADLLFVILCIPFTAADYVTD     108
Eugracilis        LFGVIAITGL-LGNLLVILVVVFNKNMRSTTNLMIVNLAAADLLFVILCIPFTATDYMVY     105
Ficusphila        FFGAIAITGF-FGNLLVILVVVFNKSMRSTTNLMIVNLAVADLLFVILCIPFTATDYMVP     107
Takahashi         FFGVIAITGF-FGNLLVILVVVFNKNMRSTTNLMIVNLAAADLMFVILCIPFTATDYMVY     101
Rhopaloa          FFGAIAITGF-FGNLLVILVVVFNKNMRSTTNLMIVNLAAADLMFVILCIPFTATDYMVY     107
Elegans           FFGAIAITGF-FGNLLVILVVVFNKNMRSTTNLMIVNLAAADLMFVILCIPFTATDYMVY     107
Erecta            FFGVIAITGF-FGNLLVILVVVFNNNMRSTTNLMIVNLAAADLMFVILCIPFTATDYMVY     102
Mauritania        FFGVIAITGF-FGNLLVILVVVFNNNMRSTTNLMIVNLAAADLMFVILCIPFTATDYMVY     102
Sechelia          FFGVIAITGF-FGNLLVILVVVFNNNMRSTTNLMIVNLAAADLMFVILCIPFTATDYMVY     102
Melanogaster      FFGVIAITGF-FGNLLVILVVVFNNNMRSTTNLMIVNLAAADLMFVILCIPFTATDYMVY     102
simulans          FFGVIAITGF-FGNLLVILVVVFNNNMRSTTNLMIVNLAAADLMFVILCIPFTATDYMVY     102
Suzuki            FFGAIAITGF-FGNLLVILVVVFNKNMRSTTNLMIVNLAAADLMFVILCIPFTATDYMVY     105
biarmipes         FFGAIAITGF-FGNLLVILVVVFNKNMRSTTNLMIVNLAAADLMFVVLCIPFTATDYMVY     105
                   ** **::*: :*********::*:.*:*****:*****.***:**::*:**** **:.

Grimshawi         HWPFGTIWCRIVQYLIVVTAYASIYTLVLMSIDRFLAVVHPIRSRMLRTEHITKIAIFTL     175
Virilis           HWPFGKMWCRSVQYLIVVTAYASIYTLVLMSIDRFLAVVHPIRSRMLRTEHITKIAIFTL     174
Mojavensis        HWPFGKIWCRSVQYLIVVTAYASIYTLVLMSIDRFLAVVHPIRSRMLRTEHITKIAIFTL     170
Bipectinate       VWPFGLFWCRSVQYLIVVTAFASIYTLVLMSIDRFLAVVHPIRSRMMRTENITMIAIVTL     163
Anannassae        VWPFGLFWCRSVQYLIVVTAFASIYTLVLMSIDRFLAVVHPIRSRMMRTENITMIAIVTL     163
Serrata           YWPFGGIWCRCVQYLIVVTAFASIYTLVLMSIDRFLAVVHPIRSRMMRTENITMVAIVTL     168
Kikkawei          YWPFGGFWCRSVQYLIVVTAFASIYTLVLMSIDRFLAVVHPIRSRMMRTENITMVAIVTL     168
Eugracilis        YWPYGRFWCRSVQYLIVVTAFASIYTLVLMSIDRFLAVVHPIRSRMMRTENITLIAIVTL     165
Ficusphila        YWPYGRFWCHSVQYLIVVTAFASIYTLVLMSIDRFLAVVHPIRSRMMRTENITLIAIVTL     167
Takahashi         YWPYGVFWCRSVQYLIVVTAFASIYTLVLMSIDRFLAVVHPIRSRMMRTENITLIAIVTL     161
Rhopaloa          YWPFGRFWCRSVQYLIVVTAFASIYTLVLMSIDRFLAVVHPIRSRMLRTENITLIAIVTL     167
Elegans           YWPFGRFWCRSVQYLIVVTAFASIYTLVLMSIDRFLAVVHPIRSRMMRTENITLIAIVTL     167
Erecta            YWPYGRFWCRSVQYLIVVTAFASIYTLVLMSIDRFLAVVHPIRSRMMRTENITLIAIVTL     162
Mauritania        YWPYGRFWCRSVQYLIVVTAFASIYTLVLMSIDRFLAVVHPIRSRMMRTENITLIAIVTL     162
Sechelia          YWPYGRFWCRSVQYLIVVTAFASIYTLVLMSIDRFLAVVHPIRSRMMRTENITLIAIVTL     162
Melanogaster      YWPYGRFWCRSVQYLIVVTAFASIYTLVLMSIDRFLAVVHPIRSRMMRTENITLIAIVTL     162
simulans          YWPYGRFWCRSVQYLIVVTAFASIYTLVLMSIDRFLAVVHPIRSRMMRTENITLIAIVTL     162
Suzuki            YWPYGRFWCRSVQYLIVVTAFASIYTLVLMSIDRFLAVVHPIRSRMMRTENITLIAIVTL     165
biarmipes         YWPYGRFWCRSVQYLIVVTAFASIYTLVLMSIDRFLAVVHPIRSRMMRTENITLIAIVTL     165
                   **:* :**: *********:*************************:***:** :**.**
```

```
Grimshawi         WTVVLTVSMPVTFTHDLVVHHNYKTNSTYAKCHFIE-NDLLDWLTFQVSFFISSYLLPLM        234
Virilis           WTVVLTVSMPVTFAHDVVVDYDNQTNVTYAMCRYID-NDVLDLSTFQVSFFISSYLLPLM        233
Mojavensis        WTVVLTVSMPVTFAHDVVVDYDNQTNVTYAMCRFID-NEVLEQSTFQVSFFISSYLLPLM        229
Bipectinate       WAVILVVSTPVGFVNDMVVLYDNKTNITYVACTYDAGNELLSPRSFQISFFISSYLLPLM        223
Anannassae        WAVILVVSTPVGFVNDMVVSYDNKTNITYVLCTFDAGNELLSPRSFQISFFVSSYLLPLM        223
Serrata           WVVILVVSVPVAFINDFAVMSDGKR--SLGMCTFKP-NDYISPRTFQVSFFISSYLLPLM        225
Kikkawei          WVVILVVSVPVAFINDFAVMSDGKR--SIGMCTFKA-NDYISPRTFQVSFFISSYLLPLM        225
Eugracilis        WIVVLIISVPVALTHDVVVDHDVKKNITYGMCTFKP-NDYVTEKSYHVIFFITSYLLPLM        224
Ficusphila        WIVVLVVSVPVALNHDVVEAIDEKKNITYGLCTYSP-NEFLSSRTYHVTFFVSSYLMPLM        226
Takahashi         WIVVLVVSLPVFFAHDVMVQYDAKKNITYGMCQFTE-NDFMGPKTYQVTFFTSSYLLPLM        220
Rhopaloa          WIVVLVVSMPVAFTHDVVVDYDAKKNITYGMCMFTN-NDFLGSRTYQVTFFISSYMLPLM        226
Elegans           WIVVLVVSVPVAFTHDVVVESDAKKNITYGMCMFIT-NDIMDSRTYHVTFFISSYMLPLM        226
Erecta            WIVVLVVSVPVAFTHDVVVDYDAKKNITYGMCTFTS-NEFLSPRTYQVTFFISSYLLPLM        221
Mauritania        WIVVLVVSVPVAFTHDVVVDYDAKKNITYGMCTFTT-NDFLSPRTYQVTFFISSYLLPLM        221
Sechelia          WIVVLVVSVPVAFTHDVVVDYDAKKNITYGMCTFTT-NDFLSPRTYQVTFFISSYLLPLM        221
Melanogaster      WIVVLVVSVPVAFTHDVVVDYDAKKNITYGMCTFTT-NDFLGPRTYQVTFFISSYLLPLM        221
simulans          WIVVLVVSVPVAFTHDVVVDYDAKKNITYGMCTFTT-NDFLSPRTYQVTFFISSYLLPLM        221
Suzuki            WIVVLVISMPVFFTHDVMVDTDAKKNITYGMCTFAT-NDFLGPKTYQVTFFISSYLLPLM        224
biarmipes         WIVVLVVSMPVFFTHDVMEDTDAKKNITYGMCTFAT-NDFLGPKTYQVTFFVSSYLLPLM        224
                  * *:* :* ** : :*.    : :   :   * :   *: :   :::: ** :**::***

Grimshawi         VISGLYVRMIMRLWRQGSGVRMSKESQRGRKRVTRLVVVVVIAFASLWLPVQL-------        287
Virilis           VISGLYVRMIMRLWHQGTGVRMSKESQRGRKRVTRLVVVVVIAFASLWLPVQL-------        286
Mojavensis        VISGLYVRMIMRLWHQGTGVRMSKESQRGRKRVTRLVVVVVIAFASLWLPVQL-------        282
Bipectinate       IISGLYVRMIMRLWHQGSGVRMSKESQRGRKRVTRLVVVVVIAFASLWLPVQL-------        276
Anannassae        IISGLYVRMIMRLWHQGSGVRMSKESQRGRKRVTRLVVVVVIAFASLWLPVQL-------        276
Serrata           IISGLYVRMIMRLWHQGSGVRMSKESQRGRKRVTRLVVVVVIAFASLWLPIQL-------        278
Kikkawei          IISGLYVRMIMRLWHQGSGVRMSKESQRGRKRVTRLVVVVVIAFASLWLPIQVSILFVHS        285
Eugracilis        IISGLYMRMIMRLWRQGTGVRMSKESQRGRKRVTRLVVVVVIAFASLWLPIQL-------        277
Ficusphila        VISGLYMRMIMRLWRQGTGVRMSKESQRGRKRVTRLVVVVVIAFASLWLPVQL-------        279
Takahashi         IISGLYMRMIMRLWRQGTGVRMSKESQRGRKRVTRLVVVVVIAFASLWLPVQL-------        273
Rhopaloa          IISGLYMRMIMRLWRQGTGVRMSKESQRGRKRVTRLVIVVVIAFASLWLPVQL-------        279
Elegans           IISGLYMRMIMRLWRQGTGVRMSKESQRGRKRVTRLVIVVVIAFASLWLPVQL-------        279
Erecta            IISGLYMRMIMRLWRQGTGVRMSKESQRGRKRVTRLVVVVVIAFASLWLPVQL-------        274
Mauritania        IISGLYMRMIMRLWHQGTGVRMSKESQRGRKRVTRLVVVVVIAFASLWLPVQL-------        274
Sechelia          IISGLYMRMIMRLWRQGTGVRMSKESQRGRKRVTRLVVVVVIAFASLWLPVQL-------        274
Melanogaster      IISGLYMRMIMRLWRQGTGVRMSKESQRGRKRVTRLVVVVVIAFASLWLPVQL-------        274
simulans          IISGLYMRMIMRLWRQGTGVRMSKESQRGRKRVTRLVVVVVIAFASLWLPVQL-------        274
Suzuki            IISGLYMRMIMRLWHQGTGVRMSKESQRGRKRVTRLVVVVVIAFASLWLPVQL-------        277
biarmipes         IISGLYMRMIMRLWHQGTGVRMSKESQRGRKRVTRLVVVVVIAFASLWLPVQL-------        277
                  :*****:*******:**:*******************:************:*:

Grimshawi         --------------ILLLKALNIYVANSMLSVILQIVAQTMAYTSSSINPLLYAFLSDNF        333
Virilis           --------------ILLLKALDMYEINSMFNVILQIVAHTMAYTSSCINPLLYAFLSDNF        332
Mojavensis        --------------ILLLKALDLYETNNMFSVILQIVAHTMAYTSSCINPLLYAFLSDNF        328
Bipectinate       --------------ILLLKALNLYKADTLFKIILQISAQTLAYSSSCINPLLYAFLSDNF        322
Anannassae        --------------ILLLKALNLYQADTLFKVILQISAQTLAYSSSCINPLLYAFLSDNF        322
Serrata           --------------ILLLKSLGVIQINSLPKVVMQVSAQTLAYSSSCINPLLYAFLSDNF        324
Kikkawei          LSNTIKFHLNCPQLILLLKSLDIIQINSLPKVVMQVSAQTLAYSSSCINPLLYAFLSDNF        345
Eugracilis        --------------ILLLKSLEILTTDTLIKLIIQVAAQTLAYTSSCINPLLYAFLSENF        323
Ficusphila        --------------ILLLKALDVIEMNSLNKLIIQVSAQTLAYSSSCINPLLYAFLSENF        325
Takahashi         --------------ILLLKSLNVIETNTLSKLIIQVTAQTLAYSSSCINPLLYAFLSENF        319
Rhopaloa          --------------ILLFKALDVIEMNSLTKLVIQVTAQTLAYSSSCINPLLYAFLSENF        325
Elegans           --------------ILLFKALGVVEMNSLTKLVIQVTAQTLAYSSSCINPLLYAFLSENF        325
Erecta            --------------ILLLKSLDVIETNTLTKLVIQVTAQTLAYSSSCINPLLYAFLSENF        320
Mauritania        --------------ILLLKSLDIIETNTLTKLVIQVTAQTLAYSSSCINPLLYAFLSENF        320
Sechelia          --------------ILLLKSLDVIETNTLTKLVIQVTAQTLAYSSSCINPLLYAFLSENF        320
Melanogaster      --------------ILLLKSLDVIETNTLTKLVIQVTAQTLAYSSSCINPLLYAFLSENF        320
simulans          --------------ILLLKSLDVIETNTLTKLVIQVTAQTLAYSSSCINPLLYAFLSENF        320
Suzuki            --------------LLLFKSLGVIETNTLTKLIIQVTAQTLAYSSSCINPLLYAFLSENF        323
biarmipes         --------------ILLFKTLGVIETNTLTKLIIQVTAQTLAYSSSCINPLLYAFLSENF        323
                                :**:*:* :   :.: .:::*: *:*:**:**.**********:**

Grimshawi         RKAFHKAINCSNRYQDYTSDLPPPRKTSCGRTSTTGL  370
Virilis           RKAFYKAINCSNRYHNYTSDLPPPRKTSCGRTSTTGL  369
Mojavensis        RKAFYKAINCSNRYQNYTSDLPPPRKTSCGRTSTTGL  365
Bipectinate       RKAFYKAINCSSRYQNYTSDLPPPRKTSCARTSTTGL  359
Anannassae        RKAFYKAINCSSRYQNYTSDLPPPRKTSCARTSTTGL  359
Serrata           RKAFYKAINCSPRYQNYTSDLPPPRKTSCARTSTTGL  361
Kikkawei          RKAFYKAINCSPRYQNYTSDLPPPRKTSCARTSTTGL  382
```

```
Eugracilis        RKAFYKAINCSSRYQNYTSDLPPPRKTSCARTSTTGL  360
Ficusphila        RKAFYKAINCSSRYQNYTSDLPPPRKTSCARTSTTGL  362
Takahashi         RKAFYKAINCSSRYQNYTSDLPPPRKTSCARTSTTGL  356
Rhopaloa          RKAFYKAINCSSRYQNYTSDLPPPRKTSCARTSTTGL  362
Elegans           RKAFYKAINCSSRYQNYTSDLPPPRKTSCARTSTTGL  362
Erecta            RKAFYKAVNCSSRYQNYTSDLPPPRKTSCARTSTTGL  357
Mauritania        RKAFYKAVNCSSRYQNYTSDLPPPRKTSCARTSTTGL  357
Sechelia          RKAFFKAVNCSSRYQNYTSDLPPPRKTSCARTSTTGL  357
Melanogaster      RKAFYKAVNCSSRYQNYTSDLPPPRKTSCARTSTTGL  357
simulans          RKAFYKAVNCSSRYQNYTSDLPPPRKTSCARTSTTGL  357
Suzuki            RKAFYKAINCSSRYQNYTSDLPPPRKTSCGRTSTTGL  360
biarmipes         RKAFYKAINCSSRYQNYTSDLPPPRKTSCARTSTTGL  360
```

Melanogaster NP_001263042.1

```
  1 menttmlani slnatrneen itsfftdeew laingtlpwi vgfffgviai tgffgnllvi
 61 lvvvfnnnmr sttnlmivnl aaadlmfvil cipftatdym vyywpygrfw crsvqylivv
121 tafasiytlv lmsidrflav vhpirsrmmr tenitliaiv tlwivvlvvs vpvafthdvv
181 vdydakknit ygmctfttnd flgprtyqvt ffissyllpl miisglymrm imrlwrqgtg
241 vrmskesqrg rkrvtrlvvv vviafaslwl pvqlillllks ldvietntlt klviqvtaqt
301 layssscinp llyaflsenf rkafykavnc ssryqnytsd lppprktsca rtsttgl
```

Erecta XP_001981439.1

```
1 mentallani slnatrneen itsfftdeew laisgtlpwi vgfffgviai tgffgnllvi
 61 lvvvfnnnmr sttnlmivnl aaadlmfvil cipftatdym vyywpygrfw crsvqylivv
121 tafasiytlv lmsidrflav vhpirsrmmr tenitliaiv tlwivvlvvs vpvafthdvv
181 vdydakknit ygmctftsne flsprtyqvt ffissyllpl miisglymrm imrlwrqgtg
241 vrmskesqrg rkrvtrlvvv vviafaslwl pvqlillllks ldvietntlt klviqvtaqt
301 layssscinp llyaflsenf rkafykavnc ssryqnytsd lppprktsca rtsttgl
```

Suzuki XP_016926809.2

```
  1 mdtmenstll gkislnasrs denitsfftd eewlaisgtl pwivvfffga iaitgffgnl
 61 lvilvvvfnk nmrsttnlmi vnlaaadlmf vilcipftat dymvyywpyg rfwcrsvqyl
121 ivvtafasiy tlvlmsidrf lavvhpirsr mmrtenitli aivtlwivvl vismpvffth
181 dvmvdtdakk nitygmctfa tndflgpkty qvtffissyl lplmiisgly mrmimrlwhq
241 gtgvrmskes qrgrkrvtrl vvvvviafas lwlpvqllll fkslgvietn tltkliiqvt
301 aqtlaysssc inpllyafls enfrkafyka incssryqny tsdlppprkt scgrtsttgl
```

Sechelia XP_002037108.1

```
1 menttmlani slnatrneen itsfftdeew laingtlpwi vgfffgviai tgffgnllvi
 61 lvvvfnnnmr sttnlmivnl aaadlmfvil cipftatdym vyywpygrfw crsvqylivv
121 tafasiytlv lmsidrflav vhpirsrmmr tenitliaiv tlwivvlvvs vpvafthdvv
181 vdydakknit ygmctfttnd flsprtyqvt ffissyllpl miisglymrm imrlwrqgtg
241 vrmskesqrg rkrvtrlvvv vviafaslwl pvqlillllks ldvietntlt klviqvtaqt
301 layssscinp llyaflsenf rkaffkavnc ssryqnytsd lppprktsca rtsttgl
```

simulans XP_002105245.1

```
  1 menttmlani slnatrneen itsfftdeew laingtlpwi vgfffgviai tgffgnllvi
 61 lvvvfnnnmr sttnlmivnl aaadlmfvil cipftatdym vyywpygrfw crsvqylivv
121 tafasiytlv lmsidrflav vhpirsrmmr tenitliaiv tlwivvlvvs vpvafthdvv
181 vdydakknit ygmctfttnd flsprtyqvt ffissyllpl miisglymrm imrlwrqgtg
241 vrmskesqrg rkrvtrlvvv vviafaslwl pvqlillllks ldvietntlt klviqvtaqt
301 layssscinp llyaflsenf rkafykavnc ssryqnytsd lppprktsca rtsttgl
```

biarmipes XP_016950481.1

```
1 mdtmdnatll gkislnasrn denitsfftd eewlaingtl pwivafffga iaitgffgnl
 61 lvilvvvfnk nmrsttnlmi vnlaaadlmf vvlcipftat dymvyywpyg rfwcrsvqyl
121 ivvtafasiy tlvlmsidrf lavvhpirsr mmrtenitli aivtlwivvl vvsmpvffth
181 dvmedtdakk nitygmctfa tndflgpkty qvtffvssyl lplmiisgly mrmimrlwhq
241 gtgvrmskes qrgrkrvtrl vvvvviafas lwlpvqlill fktlgvietn tltkliiqvt
301 aqtlaysssc inpllyafls enfrkafyka incssryqny tsdlppprkt scartsttgl
```

Takahashi XP_017006243.1

```
  1 menatimlnk inssrmeens tsyftdeewl sisstlpwiv cfffgviait gffgnllvil
 61 vvvfnknmrs ttnlmivnla aadlmfvilc ipftatdymv yywpygvfwc rsvqylivvt
121 afasiytlvl msidrflavv hpirsrmmrt enitliaivt lwivvlvvsl pvffahdvmv
181 qydakknity gmcqftendf mgpktyqvtf ftssyllplm iisglymrmi mrlwrqgtgv
241 rmskesqrgr krvtrlvvvv viafaslwlp vqlillllksl nvietntlsk liiqvtaqtl
```

301 ayssscinpl lyaflsenfr kafykaincs sryqnytsdl ppprktscar tsttgl

Bipectinate XP_017092876.2
1 mdnttlatpd platmatydd nltsffseee rqaiegtliw lvpfffgiia ltgffgnllv
61 ilvvvfnknm rsttnlmivn lavadllfvi fcipftatdy vtkvwpfglf wcrsvqyliv
121 vtafasiytl vlmsidrfla vvhpirsrmm rtenitmiai vtlwavilvv stpvgfvndm
181 vvlydnktni tyvactydag nellsprsfq isffissyll plmiisglyv rmimrlwhqg
241 sgvrmskesq rgrkrvtrlv vvvviafasl wlpvqlilll kalnlykadt lfkiilqisa
301 qtlaysssci npllyaflsd nfrkafykai ncssryqnyt sdlppprkts cartsttgl

Rhopaloa XP_016982221.1
1 mdnitvspae mlgnisinas rndenltsff tdeewlaing tlpwivgfff gaiaitgffg
61 nllvilvvvf nknmrsttnl mivnlaaadl mfvilcipft atdymvyywp fgrfwcrsvq
121 ylivvtafas iytlvlmsid rflavvhpir srmlrtenit liaivtlwiv vlvvsmpvaf
181 thdvvvdyda kknitygmcm ftnndflgsr tyqvtffiss ymlplmiisg lymrmimrlw
241 rqgtgvrmsk esqrgrkrvt rlvivvviaf aslwlpvqli llfkaldvie mnsltklviq
301 vtaqtlayss scinpllyaf lsenfrkafy kaincssryq nytsdlpppr ktscartstt
361 gl

Eugracilis XP_017083059.1
1 menmdnstmv vdlvvnstrn denmtsfftd eewvairgtl pwiviflfgv iaitgllgnl
61 lvilvvvfnk nmrsttnlmi vnlaaadllf vilcipftat dymvyywpyg rfwcrsvqyl
121 ivvtafasiy tlvlmsidrf lavvhpirsr mmrtenitli aivtlwivvl iisvpvalth
181 dvvvdhdvkk nitygmctfk pndyvteksy hviffitsyl lplmiisgly mrmimrlwrq
241 gtgvrmskes qrgrkrvtrl vvvvviafas lwlpiqlill lksleilttd tlikliiqva
301 aqtlaytssc inpllyafls enfrkafyka incssryqny tsdlppprkt scartsttgl

Ficusphila XP_017038669.1
1 menatffpak mlgslasnat rndenltttff tdeewlains tlpwvvgfff gaiaitgffg
61 nllvilvvvf nksmrsttnl mivnlavadl lfvilcipft atdymvpywp ygrfwchsvq
121 ylivvtafas iytlvlmsid rflavvhpir srmmrtenit liaivtlwiv vlvvsvpval
181 nhdvveaide kknityglct yspneflssr tyhvtffvss ylmplmvisg lymrmimrlw
241 rqgtgvrmsk esqrgrkrvt rlvvvvviaf aslwlpvqli lllkaldvie mnslnkliiq
301 vsaqtlayss scinpllyaf lsenfrkafy kaincssryq nytsdlpppr ktscartstt
361 gl

Anannassae XP_001955066.1
1 mdnttlatse llatmatydd nftsffseee rqaiegtliw lvpfffgiia ltgffgnllv
61 ilvvvfnknm rsttnlmivn lavadllfvi fcipftatdy vtkvwpfglf wcrsvqyliv
121 vtafasiytl vlmsidrfla vvhpirsrmm rtenitmiai vtlwavilvv stpvgfvndm
181 vvsydnktni tyvlctfdag nellsprsfq isffvssyll plmiisglyv rmimrlwhqg
241 sgvrmskesq rgrkrvtrlv vvvviafasl wlpvqlilll kalnlyqadt lfkvilqisa
301 qtlaysssci npllyaflsd nfrkafykai ncssryqnyt sdlppprkts cartsttgl

Serrata XP_020809023.1
1 menttliage tlanislani tqsdenitsf fteeewlalt gilpwlvgft fgaiaitgfl
61 gnllvilvvv fnnnmrsttn lmivnlaaad llfvilcipf taadyvtdyw pfggiwcrcv
121 qylivvtafa siytlvlmsi drflavvhpi rsrmmrteni tmvaivtlwv vilvvsvpva
181 findfavmsd gkrslgmctf kpndyisprt fqvsffissy llplmiisgl yvrmimrlwh
241 qgsgvrmske sqrgrkrvtr lvvvvviafa slwlpiqlil llkslgviqi nslpkvvmqv
301 saqtlaysss cinpllyafl sdnfrkafyk aincspryqn ytsdlppprk tscartsttg
361 l

Kikkawei KAH8347379.1
menttlitge tlanislant trsdenitsf fteeewlalt gilpwlvgft fgaiaitgfl
61 gnllvilvvv fnnnmrsttn lmivnlavad llfvilcipf taadyvtdyw pfggfwcrsv
121 qylivvtafa siytlvlmsi drflavvhpi rsrmmrteni tmvaivtlwv vilvvsvpva
181 findfavmsd gkrsigmctf kandyisprt fqvsffissy llplmiisgl yvrmimrlwh
241 qgsgvrmske sqrgrkrvtr lvvvvviafa slwlpiqvsi lfvhslsnti kfhlncpqli
301 lllksldiiq inslpkvvmq vsaqtlayss scinpllyaf lsdnfrkafy kaincspryq
361 nytsdlpppr ktscartstt gl

Mauritania XP_033166501.1
1 menttmlani slnatrneen itsfftdeew laingtlpwi vgfffgviai tgffgnllvi
61 lvvvfnnnmr sttnlmivnl aaadlmfvil cipftatdym vyywpygrfw crsvqylivv

```
      121 tafasiytlv lmsidrflav vhpirsrmmr tenitliaiv tlwivvlvvs vpvafthdvv
      181 vdydakknit ygmctfttnd flsprtyqvt ffissyllpl miisglymrm imrlwhqgtg
      241 vrmskesqrg rkrvtrlvvv vviafaslwl pvqlillllks ldiietntlt klviqvtaqt
      301 layssscinp llyaflsenf rkafykavnc ssryqnytsd lppprktsca rtsttgl
```

Elegans XP_017131735.1

```
  1 mdnitesaae mlanislnas rndenltsff tdeewlaisg tlpwivgfff gaiaitgffg
       61 nllvilvvvf nknmrsttnl mivnlaaadl mfvilcipft atdymvyywp fgrfwcrsvq
      121 ylivvtafas iytlvlmsid rflavvhpir srmmrtenit liaivtlwiv vlvvsvpvaf
      181 thdvvvesda kknitygmcm fitndimdsr tyhvtffiss ymlplmiisg lymrmimrlw
      241 rqgtgvrmsk esqrgrkrvt rlvivvviaf aslwlpvqli llfkalgvve mnsltklviq
      301 vtaqtlayss scinpllyaf lsenfrkafy kaincssryq nytsdlpppr ktscartstt
      361 gl
```

Grimshawi XP_001995830.3

```
1 mnqsyimenn tpsmsnsswl elklttnnst gnsslsldae eeeairqtvr wlvplffgii
       61 aisgfffgnl lvilvvllnk nmhsttnlli vnlaaadllf vifcvpftav dyvadhwpfg
      121 tiwcrivqyl ivvtayasiy tlvlmsidrf lavvhpirsr mlrtehitki aiftlwtvvl
      181 tvsmpvtfth dlvvhhnykt nstyakchfi endlldwltf qvsffissyl lplmvisgly
      241 vrmimrlwrq gsgvrmskes qrgrkrvtrl vvvvviafas lwlpvqlill lkalniyvan
      301 smlsvilqiv aqtmaytsss inpllyafls dnfrkafhka incsnryqdy tsdlppprkt
      361 scgrtsttgl
```

Virilis XP_002053983.1

```
     1 mnlsnitltl psnsswlesq lelttassnl dnstlssfya eaeairatvr wvvpfffgii
       61 aisgffgnll vilvvllnkn mhsttnlliv nlaaadllfv ifcvpftaid yvtqhwpfgk
      121 mwcrsvqyli vvtayasiyt lvlmsidrfl avvhpirsrm lrtehitkia iftlwtvvlt
      181 vsmpvtfahd vvvdydnqtn vtyamcryid ndvldlstfq vsffissyll plmvisglyv
      241 rmimrlwhqg tgvrmskesq rgrkrvtrlv vvvviafasl wlpvqlilll kaldmyeins
      301 mfnvilqiva htmaytssci npllyaflsd nfrkafykai ncsnryhnyt sdlppprkts
      361 cgrtsttgl
```

Mojavensis XP_002000968.1

```
  1 melsnltttl atqlhqlats slnhsigpdn stalqseeea iratvrwlvp iffgiiavsg
       61 ffgnllvilv vllnknmhst tnllivnlaa adllfvifcv pftavdyvtq hwpfgkiwcr
      121 svqylivvta yasiytlvlm sidrflavvh pirsrmlrte hitkiaiftl wtvvltvsmp
      181 vtfahdvvvd ydnqtnvtya mcrfidnevl eqstfqvsff issyllplmv isglyvrmim
      241 rlwhqgtgvr mskesqrgrk rvtrlvvvvv iafaslwlpv qlilllkald lyetnnmfsv
      301 ilqivahtma ytsscinpll yaflsdnfrk afykaincsn ryqnytsdlp pprktscgrt
      361 sttgl
```
